# Design of SARS-CoV-2 papain-like protease inhibitor with antiviral efficacy in a mouse model

**DOI:** 10.1101/2023.12.01.569653

**Authors:** Bin Tan, Xiaoming Zhang, Ahmadullah Ansari, Prakash Jadhav, Haozhou Tan, Kan Li, Ashima Chopra, Alexandra Ford, Xiang Chi, Francesc Xavier Ruiz, Eddy Arnold, Xufang Deng, Jun Wang

## Abstract

The emergence of SARS-CoV-2 variants and drug-resistant mutants calls for additional oral antivirals. The SARS-CoV-2 papain-like protease (PL^pro^) is a promising but challenging drug target. In this study, we designed and synthesized 85 noncovalent PL^pro^ inhibitors that bind to the newly discovered Val70^Ub^ site and the known BL2 groove pocket. Potent compounds inhibited PL^pro^ with inhibitory constant K_i_ values from 13.2 to 88.2 nM. The co-crystal structures of PL^pro^ with eight leads revealed their interaction modes. The *in vivo* lead **Jun12682** inhibited SARS-CoV-2 and its variants, including nirmatrelvir-resistant strains with EC_50_ from 0.44 to 2.02 µM. Oral treatment with **Jun12682** significantly improved survival and reduced lung viral loads and lesions in a SARS-CoV-2 infection mouse model, suggesting PL^pro^ inhibitors are promising oral SARS-CoV-2 antiviral candidates.

**One-Sentence Summary:** Structure-guided design of SARS-CoV-2 PL^pro^ inhibitors with *in vivo* antiviral efficacy in a mouse model.

---

The COVID-19 pandemic is a timely call for the urgent need for orally bioavailable antivirals. The FDA has approved three direct-acting antivirals, including two viral RNA-dependent RNA polymerase inhibitors, remdesivir and molnupiravir, and one viral main protease (M^pro^) inhibitor, nirmatrelvir (*1*). Remdesivir is administered by intravenous injection, and its use is limited to hospitalized patients. The clinical efficacy of remdesivir is also controversial (*2*). Molnupiravir is a mutagen and should not be used in pregnant women (*3*). Nirmatrelvir is co-administered with ritonavir, a CYP3A4 inhibitor, to improve the *in vivo* half-life (*4*). For this reason, Paxlovid, a combination of nirmatrelvir and ritonavir, has drug-drug interaction concerns. Ensiltrelvir is a second-generation M^pro^ inhibitor approved in Japan (*5*). Although it is highly efficacious in clinical trials (*6*), ensitrelvir is a potent CYP3A4 inhibitor that may lead to severe adverse drug-drug interactions with other medications (*7, 8*). Mutant SARS-CoV-2 viruses with resistance against remdesivir or nirmatrelvir have been identified from viral passage experiments in cell culture (*9-11*) and drug-treated COVID-19 patients (*12-15*). Therefore, additional antivirals with alternative mechanisms of action are urgently needed to combat drug-resistant and emerging SARS-CoV-2 variants.

The papain-like protease (PL^pro^) is one of the two viral cysteine proteases encoded by SARS-CoV-2. PL^pro^ cleaves the viral non-structural (nsp) polyproteins at the nsp1/2, nsp2/3, and nsp3/4 junctions and is pivotal for viral replication (*16*). In addition, PL^pro^ suppresses the host immune response through cleaving ubiquitin and interferon-stimulated gene 15 (ISG-15) modifications from host proteins (*17-19*). The SARS-CoV-2 PL^pro^ sequence is 82.9% identical to SARS-CoV-1 PL^pro^. PL^pro^ is highly conserved among SARS-CoV-2 variants (*20*), rendering it a high-profile antiviral drug target (*19, 21*). However, despite decades of medicinal chemistry optimization and high-throughput screening, no drug-like PL^pro^ inhibitor has shown *in vivo* antiviral efficacy in SARS-CoV-2-infected animal models (*20*). PL^pro^ substrates contain a consensus motif LXGG↓(N/K/X) (*16*). The S1 and S2 subsites of PL^pro^ form a narrow tunnel for binding two glycines (*22*). The absence of binding pockets near the catalytic cysteine Cys111 presents a challenge in designing highly potent PL^pro^ inhibitors. In this study, we describe the design of potent PL^pro^ inhibitors by exploiting a novel drug-binding site that accommodates the ubiquitin Val70 side chain (Val70^Ub^). We validate PL^pro^ as a viable drug target by demonstrating the *in vivo* antiviral efficacy of a designed PL^pro^ inhibitor **Jun12682** with oral administration in SARS-CoV-2 infected mice.

## Discovery of the binding site for ubiquitin Val70 as a new drug binding site on PL^pro^

Before our study, noncovalent and covalent PL^pro^ inhibitors were reported (*20, 23*). One potent noncovalent PL^pro^ inhibitor is **XR8-24**, which has an IC_50_ of 0.56 µM in the fluorescence resonance energy transfer (FRET) enzymatic assay and an EC_50_ of 1.2 µM in the antiviral assay (*24*). Compound 7 (**Cp7**) is a rationally designed covalent PL^pro^ inhibitor with a fumarate reactive warhead that inhibits PL^pro^ with an IC^_50_^ of 0.094 µM and SARS-CoV-2 replication with an EC_50_ of 1.1 µM (*25*). Inspired by these results, we designed a hybrid covalent PL^pro^ inhibitor **Jun11313** by converting the naphthalene in **Cp7** to 3-phenylthiophene (Fig. 1A); **Jun11313** potently inhibited PL^pro^ with an IC_50_ of 0.12 µM.

**Fig. 1.**
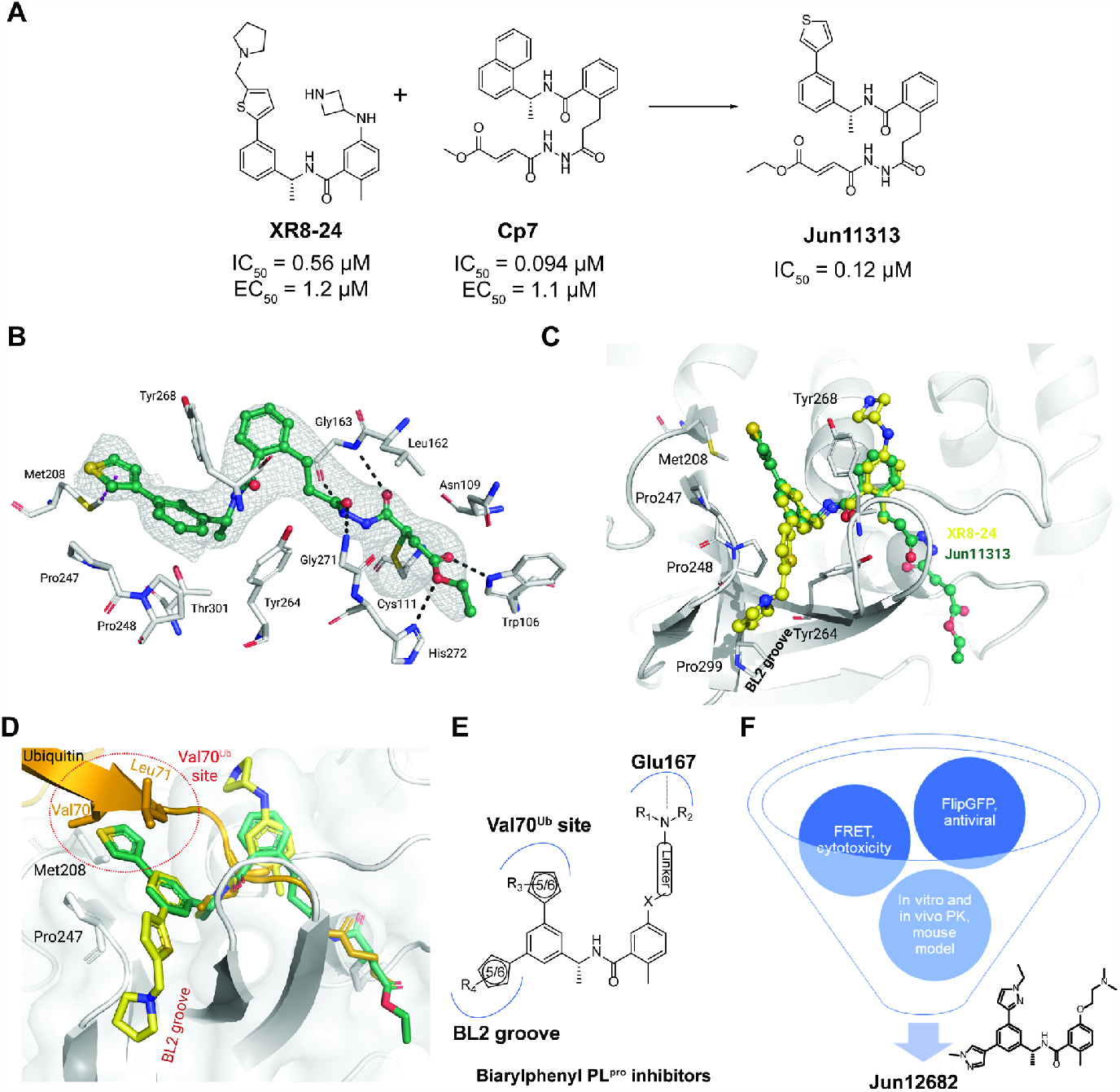
X-ray crystal structure of the covalent inhibitor Jun11313 with SARS-CoV-2 PL^pro^ and structure-based design of biarylphenyl SARS-CoV-2 PL^pro^ inhibitors. (**A**) Design of the hybrid covalent inhibitor **Jun11313** based on **XR8-24** and **Cp7**. (**B**) Atomic model of the **Jun11313** (in green sticks and spheres) binding site in PL^pro^ (light grey sticks, residues within a 5 Å distance of the inhibitor), with hydrogen bonds displayed as black dashed lines. **Jun11313** polder map (an unbiased difference map (*36*)) is displayed as a grey mesh with 4σ contour. (**C**) Superposition of the PL^pro^-**Jun11313** structure to the structure of the PL^pro^-**XR8-24** complex (PDB 7LBS), with **XR8-24** in yellow sticks and spheres, with the relevant residues for binding of both compounds indicated. (**D**) Superimposed X-ray crystal structures of SARS-CoV-2 PL^pro^ with **Jun11313** (green) (PDB: 8UVM), **XR8-24** (yellow) (PDB: 7LBS), and ubiquitin (orange) (PDB: 6XAA). (**E**) Generic chemical structure of the designed biarylphenyl PL^pro^ inhibitor. Critical interactions are highlighted. (**F**) Flow chart for the lead optimziation of PL^pro^ inhibitors. **Jun12682** was selected as the *in vivo* lead candidate.

The lead covalent compound **Jun11313** was co-crystallized with SARS-CoV-2 PL^pro^ to visualize the key interactions for binding. We determined the co-crystal structure at 2.85 Å resolution (PDB: 8UVM). As expected, the fumarate ester electrophile forms a covalent bond with the catalytic Cys111 (Fig. 1B). **Jun11313** forms hydrogen bonds with main-chain atoms of Leu162, Gly163, Tyr268, and Gly271, as well as with side-chain atoms of Trp106 and His272. **Jun11313** also makes extensive van der Waals contacts with Leu162, Tyr264, Pro247, Pro248, and Thr301 (Fig. 1B).

Surprisingly, the thienyl group in **Jun11313** is oriented towards the opposite side of the BL2 groove compared to the pyrrolidine-substituted thienyl group in **XR8-24** (Fig. 1C). The phenylthiophene in **XR8-24** makes extensive contacts with the BL2 groove, namely van der Waals interactions with residues surrounding the cavity (Pro248, Tyr264, and Tyr268; Fig. 1C). In comparison, the phenylthienyl group of **Jun11313** is oriented towards Pro247, making van der Waals contacts with both Pro248 and Pro247, and CH–π and S–π interactions with Met208 (Fig. 1B). While the strength of these interactions is difficult to estimate and has been understudied in drug design (*26, 27*), here they may be the culprit for **Jun11313**’s unexpected conformation (Fig. 1B). If the methyl pyrrolidine group in **XR8-24** were arranged as the phenylthienyl group of **Jun11313**, it should orient in a very solvent-exposed region to avoid a clash with Met208 (fig. S1A).

Superimposition of PL^pro^ structures complexed with ubiquitin and **Jun11313** revealed that the thienyl group occupies the same hydrophobic site as Val70 from ubiquitin (Fig. 1D). Therefore, we designate this pocket as the Val70^Ub^ site. The comparison of the co-crystal structures of PL^pro^ with **Jun11313** and with ubiquitin suggests that this site is essential for binding both the ubiquitin substrate and this inhibitor. Similarly, the superimposition of PL^pro^ structures complexed with ISG15 showed that the thienyl group interacts with the analogous Asn151-Leu152 site from ISG15 (fig. S1B).

## Rational design of biarylphenyl PL^pro^ inhibitors

The unexpected binding pose of **Jun11313** in PL^pro^ led us to hypothesize that potent PL^pro^ inhibitors can be designed by simultaneously engaging the BL2 groove pocket and the Val70^Ub^ hydrophobic site (Fig. 1E). It is noteworthy that the Val70^Ub^ site has not been explored for PL^pro^ inhibitor design (*20*). We designed and synthesized a library of 85 biarylphenyl benzamide compounds (Fig. S2). Aryl substitutions were installed on the 3 and 5 positions of the phenylethylamine ring to engage hydrophobic interactions with residues in the BL2 groove and the ubiquitin Val70^Ub^ binding site (Fig. 1E). In addition, diverse amines were installed on the meta-position of the benzoic acid to engage electrostatic interactions with Glu167 (*24*). The central 1-phenylethyl benzamide core structure was kept intact to maintain the critical hydrogen bonds and π-π interactions with the BL2 loop. All synthesized compounds were initially tested in the FRET-based enzymatic assay and the cytotoxicity assay in Vero E6 cells (Fig. 1F). Promising lead compounds were then tested in a secondary FlipGFP PL^pro^ assay and SARS-CoV-2 antiviral assay (*28*). FlipGFP PL^pro^ is a cell-based assay that validates intracellular PL^pro^ target engagement (*22*). Next, leads were profiled for *in vitro* microsomal stability and *in vivo* oral pharmacokinetic (PK) properties in mice. **Jun12682** was finalized as the *in vivo* lead for the SARS-CoV-2 infection mouse model study.

A complete list of the designed biarylphenyl PL^pro^ inhibitors is shown in fig. S2, with representative examples in Table 1. Among the 85 compounds tested in the FRET assay, 26 had IC_50_ < 100 nM, 42 had IC_50_ between 100 – 200 nM, and 14 had IC_50_ between 200 – 400 nM. The control compound **GRL0617** had IC_50_ values of 1.92 µM. The inhibitory constant K_i_ was determined for potent compounds (Table 1). The first designed compound, **Jun11875**, inhibited PL^pro^ with a K_i_ of 13.2 nM, a 104-fold improvement over **GRL0617** (K_i_ = 1,374 nM). However, thiophene-containing compounds were generally cytotoxic in Vero E6 cells (CC_50_ < 20 µM). Next, we examined several symmetric and asymmetric five and six-membered aromatic substitutions at the 3 and 5-positions to mitigate cellular cytotoxicity while maintaining enzymatic inhibition. Among the list of heterocycle substitutions examined, pyrazole was shown to confer potent PL^pro^ inhibition and reduced cellular cytotoxicity. It is noted that despite the potent PL^pro^ enzymatic inhibition for **Jun1247** (IC_50_ = 165.3 nM), **Jun121210** (IC_50_ = 81.9 nM), **Jun121911** (IC_50_ = 73.2 nM), **Jun12208** (IC_50_ = 91.6 nM), and **Jun12242** (IC_50_ = 90.5 nM), their cellular activities were moderate to weak as shown by the FlipGFP assay (EC_50_ > 10 µM) (fig. S2), which might due to poor membrane permeability. The most potent compounds **Jun12199** and **Jun12197** had EC_50_ of 0.8 and 0.6 µM, respectively, more than 20-fold improved from **GRL0617** (EC_50_ = 22.4 µM). The *in vivo* lead **Jun12682** inhibited PL^pro^ with a K_i_ of 37.7 nM and displayed an EC_50_ of 1.1 µM in the FlipGFP PL^pro^ assay.

**Table 1.**
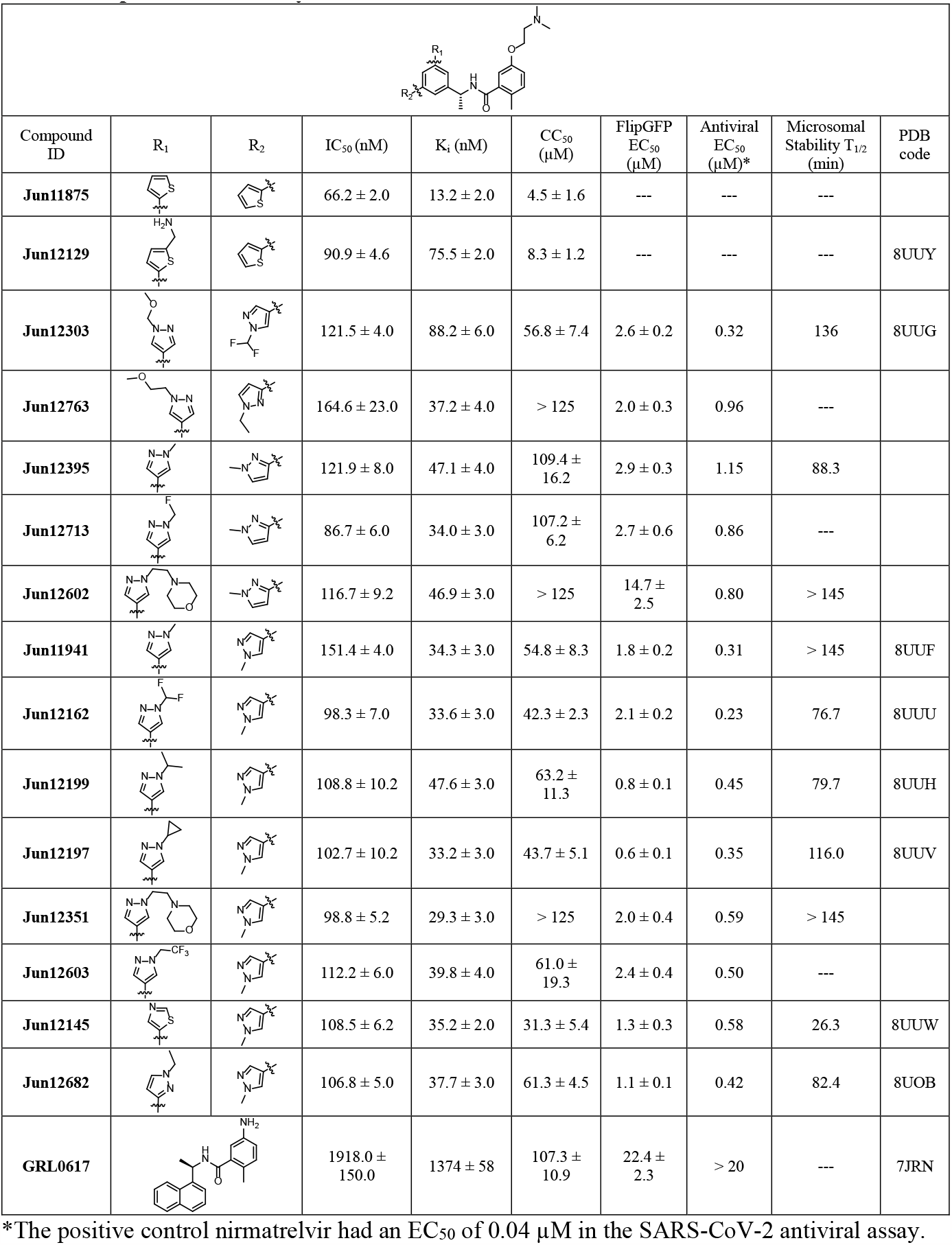
Representative biarylbenzamide series of SARS-CoV-2 PL^pro^ inhibitors.

|  |  |  |  |  |  |  |  |  |  |
| --- | --- | --- | --- | --- | --- | --- | --- | --- | --- |
| Compound ID | R<sub>1</sub> | R<sub>2</sub> | IC<sub>50</sub> (nM) | K<sub>i</sub> (nM) | CC<sub>50</sub> (μM) | FlipGFP EC<sub>50</sub> (μM) | Antiviral EC<sub>50</sub> (μM)* | Microsomal Stability T<sub>1/2</sub> (min) | PDB code |
| Jun11875 |  |  | 66.2 ± 2.0 | 13.2 ± 2.0 | 4.5 ± 1.6 | --- | --- | --- |  |
| Jun12129 |  |  | 90.9 ± 4.6 | 75.5 ± 2.0 | 8.3 ± 1.2 | --- | --- | --- | 8UUY |
| Jun12303 |  |  | 121.5 ± 4.0 | 88.2 ± 6.0 | 56.8 ± 7.4 | 2.6 ± 0.2 | 0.32 | 136 | 8UUG |
| Jun12763 |  |  | 164.6 ± 23.0 | 37.2 ± 4.0 | > 125 | 2.0 ± 0.3 | 0.96 | --- |  |
| Jun12395 |  |  | 121.9 ± 8.0 | 47.1 ± 4.0 | 109.4 ± 16.2 | 2.9 ± 0.3 | 1.15 | 88.3 |  |
| Jun12713 |  |  | 86.7 ± 6.0 | 34.0 ± 3.0 | 107.2 ± 6.2 | 2.7 ± 0.6 | 0.86 | --- |  |
| Jun12602 |  |  | 116.7 ± 9.2 | 46.9 ± 3.0 | > 125 | 14.7 ± 2.5 | 0.80 | > 145 |  |
| Jun11941 |  |  | 151.4 ± 4.0 | 34.3 ± 3.0 | 54.8 ± 8.3 | 1.8 ± 0.2 | 0.31 | > 145 | 8UUF |
| Jun12162 |  |  | 98.3 ± 7.0 | 33.6 ± 3.0 | 42.3 ± 2.3 | 2.1 ± 0.2 | 0.23 | 76.7 | 8UUU |
| Jun12199 |  |  | 108.8 ± 10.2 | 47.6 ± 3.0 | 63.2 ± 11.3 | 0.8 ± 0.1 | 0.45 | 79.7 | 8UUH |
| Jun12197 |  |  | 102.7 ± 10.2 | 33.2 ± 3.0 | 43.7 ± 5.1 | 0.6 ± 0.1 | 0.35 | 116.0 | 8UUV |
| Jun12351 |  |  | 98.8 ± 5.2 | 29.3 ± 3.0 | > 125 | 2.0 ± 0.4 | 0.59 | > 145 |  |
| Jun12603 |  |  | 112.2 ± 6.0 | 39.8 ± 4.0 | 61.0 ± 19.3 | 2.4 ± 0.4 | 0.50 | --- |  |
| Jun12145 |  |  | 108.5 ± 6.2 | 35.2 ± 2.0 | 31.3 ± 5.4 | 1.3 ± 0.3 | 0.58 | 26.3 | 8UUW |
| Jun12682 |  |  | 106.8 ± 5.0 | 37.7 ± 3.0 | 61.3 ± 4.5 | 1.1 ± 0.1 | 0.42 | 82.4 | 8UOB |
| GRL0617 |  |  | 1918.0 ± 150.0 | 1374 ± 58 | 107.3 ± 10.9 | 22.4 ± 2.3 | > 20 | --- | 7JRN |

\*The positive control nirmatrelvir had an EC<sub>50</sub> of 0.04 μM in the SARS-CoV-2 antiviral assay.

Prioritized lead compounds were tested in the SARS-CoV-2 antiviral assay and had EC_50_ from 0.23 to 1.15 µM. **Jun12682** had EC_50_ values of 0.42 and 0.51 µM in the icSARS-CoV-2-nLuc reporter virus and plaque assays, respectively (fig. S3). To examine the potential of PL^pro^ inhibitors in inhibiting SARS-CoV-2 variants and drug-resistant mutants, we selected two PL^pro^ inhibitors **Jun11941** and **Jun12682** to test against SARS-CoV-2 delta and omicron variants and three recombinant SARS-CoV-2 viruses that are resistant to nirmatrelvir, rNsp5-S144M, rNsp5-L50F/E166V, and rNsp5-L50F/E166V/L167F (Fig. 2, A to C). S144M and E166V are nirmatrelvir-resistant mutations identified from enzymatic assays and viral passage experiments (*9, 10*). **Jun11941** and **Jun12682** showed consistent antiviral activity against these viruses with EC_50_ fold-increases less than 1.5 and 2.0, respectively, compared to wild-type (WT). In comparison, the rNsp5-S144M, rNsp5-L50F/E166V, and rNsp5-L50F/E166V/L167F showed significant resistance to nirmatrelvir with EC_50_ fold-increases of 12.5, 24.2, and 21.7, respectively, compared to WT.

**Fig. 2.**
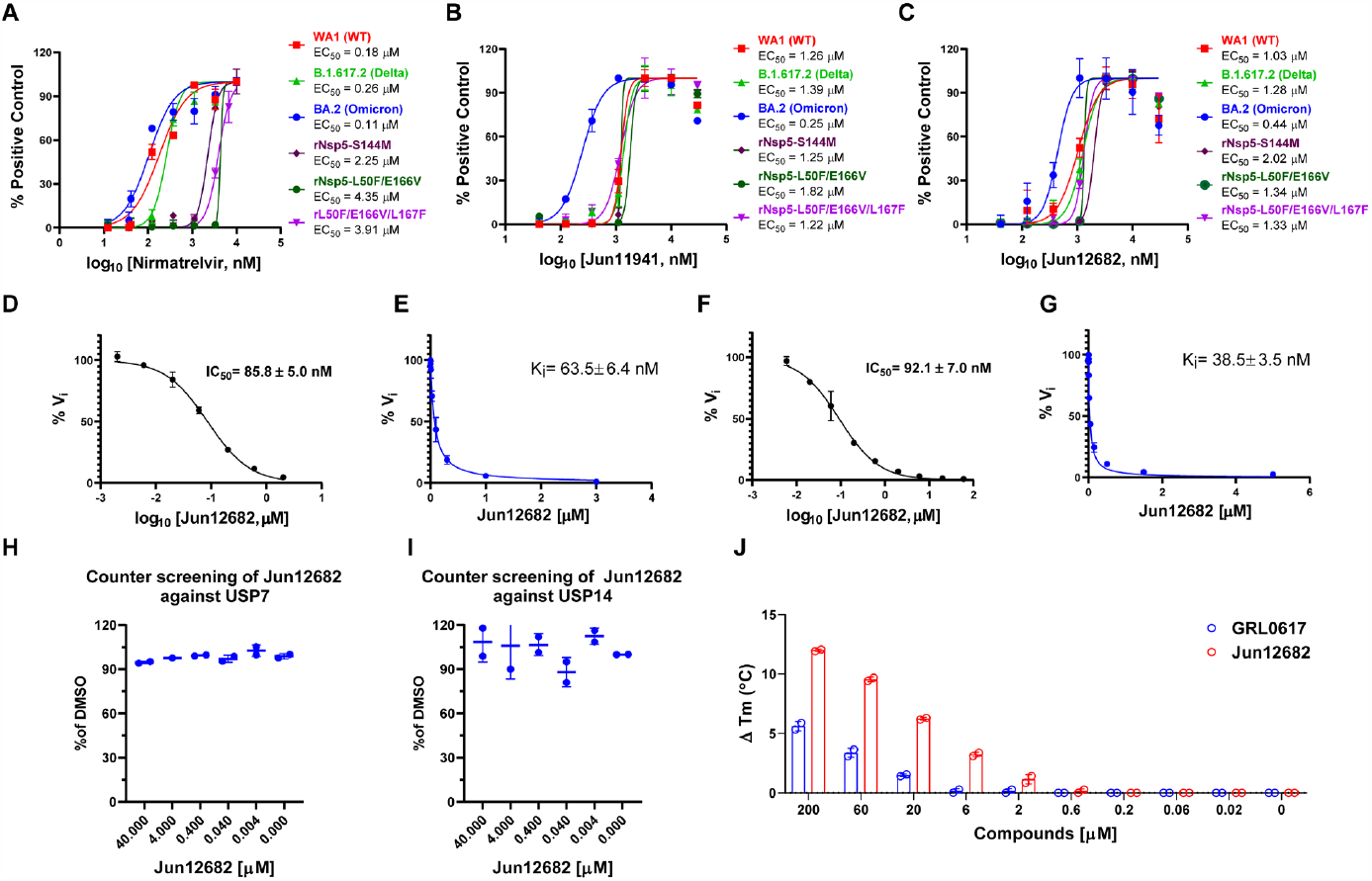
Antiviral activity of PL^pro^ inhibitors against SARS-CoV-2 variants and mechanistic studies of Jun12682. Antiviral activity of nirmatrelvir (**A**), **Jun11941** (**B**), and **Jun12682** (**C**) against SARS-CoV-2 WA1 strain (WT) (USA-WA1/2020), Omicron strain (BA.2 strain, lineage B.1.1.529, BA.2; HCoV-19/USA/CO-CDPHE-2102544747/2021), Delta strain (Lineage B.1.617.2; hCoV-19/USA/MD-HP05647/2021), and nirmatrelvir-resistant strains rNsp5-S144M, rNsp5-L50F/E166V, and rNsp5-L50F/E166V/L167F. (**D**) IC_50_ curve of **Jun12682** in inhibiting SARS-CoV-2 PL^pro^ hydrolysis of Ub-AMC. (**E**) K_i_ curve of **Jun12682** in inhibiting SARS-CoV-2 PL^pro^ hydrolysis of Ub-AMC. (**F**) IC_50_ curve of **Jun12682** in inhibiting SARS-CoV-2 PL^pro^ hydrolysis of ISG15-AMC. (**G**) K_i_ curve of **Jun12682** in inhibiting SARS-CoV-2 PL^pro^ hydrolysis of ISG15-AMC. (**H**) Counter screening of **Jun12682** in inhibiting USP7 hydrolysis of Ub-AMC. (**I**) Counter screening of **Jun12682** in inhibiting USP14 hydrolysis of Ub-AMC. (**J**) Differential scanning fluorimetry assay of **Jun12682** and GRL0617 in stabilizing the SARS-CoV-2 PL^pro^.

## Mechanism of action of Jun12682 in inhibiting SARS-CoV-2 PL^pro^

PL^pro^ is known to antagonize the host innate immune response upon viral infection by hydrolyzing the isopeptide bond between ubiquitin and ISG-15 to lysine side chains of host proteins (*17, 18*). To characterize whether **Jun12682** inhibits the deubiquitinating and deISGylating activities of SARS-CoV-2 PL^pro^, we performed the PL^pro^ enzymatic assay using Ub-AMC and ISG15-AMC substrates (*22*). **Jun12682** showed potent enzymatic inhibition with K_i_ values of 63.5 and 38.5 nM in the ubiquitin-AMC (7-amino-4-methylcoumarin) and ISG15-AMC FRET assays (Fig. 2, D to G). The results are consistent with **GRL0617** and **XR8-24** *(22, 24)*. To profile the off-target effect, **Jun12682** was tested against two closest human structural homologs of PL^pro^, USP7, and USP14 (*24, 29*). **Jun12682** displayed no inhibition of USP7 and USP14-catalyzed Ub-AMC hydrolysis at up to 40 µM (Fig. 2, H and I). **Jun12682** also showed dose-dependent stabilization of SARS-CoV-2 PL^pro^ and was more potent than GRL0617 in the differential scanning fluorimetry assay (Fig. 2J).

## X-ray crystal structures of SARS-CoV-2 PL^pro^ with inhibitors

The X-ray co-crystal structures of SARS-CoV-2 PL^pro^ were solved for eight biarylphenyl PL^pro^ inhibitors, **Jun11941, Jun12129, Jun12303, Jun12162, Jun12199, Jun12197, Jun12145**, and **Jun12682** (resolution range of 2.5-3.1 Å, Fig. 3, table S1). The phenylthienyl group of **Jun11313** binds to the Val70^Ub^ site of SARS-CoV-2 PL^pro^ in the region where residues Val70^Ub^ and Leu71^Ub^ at the end of a β sheet in ubiquitin interact with PL^pro^ (an analogous situation happens with residues Asn151 and Leu152 of ISG-15, fig. S1B) (Fig. 1B). This is a unique property of **Jun11313** due to the unusual binding conformation of the phenylthienyl group, which we have subsequently exploited in a new compound series (Fig. 3), exemplified by **Jun12682**. The 2.52 Å resolution co-crystal structure of PL^pro^ with **Jun12682** reveals the “best of both worlds”, with the N-ethyl and N-methyl pyrazole substituents in the phenyl moiety binding toward the Met208/Pro247 direction (Val70^Ub^ site) and the Pro248/Tyr264/Tyr268 direction (BL2 groove), respectively (Fig. 3A). While Met208 remains involved in CH–π and S–π interactions with the pyrazole ring, it also displays a van der Waals contact with the methyl substituent. Moreover, the phenyl group of **Jun12682** makes a π–π interaction with the side chain of Tyr264 (Fig. 3A). Fig. 3B-I shows the unbiased electron density of PL^pro^ inhibitors **Jun12145, Jun12303, Jun12199, Jun12162, Jun12129, Jun12197**, and **Jun11941**, in a similar conformation to **Jun12682**. The disubstituted phenyl group has two alternative conformations in the **Jun12145** and **Jun12129** complex structures, as indicated by the refined electron density (Fig. 3C, G; fig. S4). Despite the differences between compounds, the interactions depicted in Fig. 3A between PL^pro^ and **Jun12682** are conserved in all complexes (fig. S4). The crystallographic data and refinement statistics are listed in table S1.

**Fig. 3.**
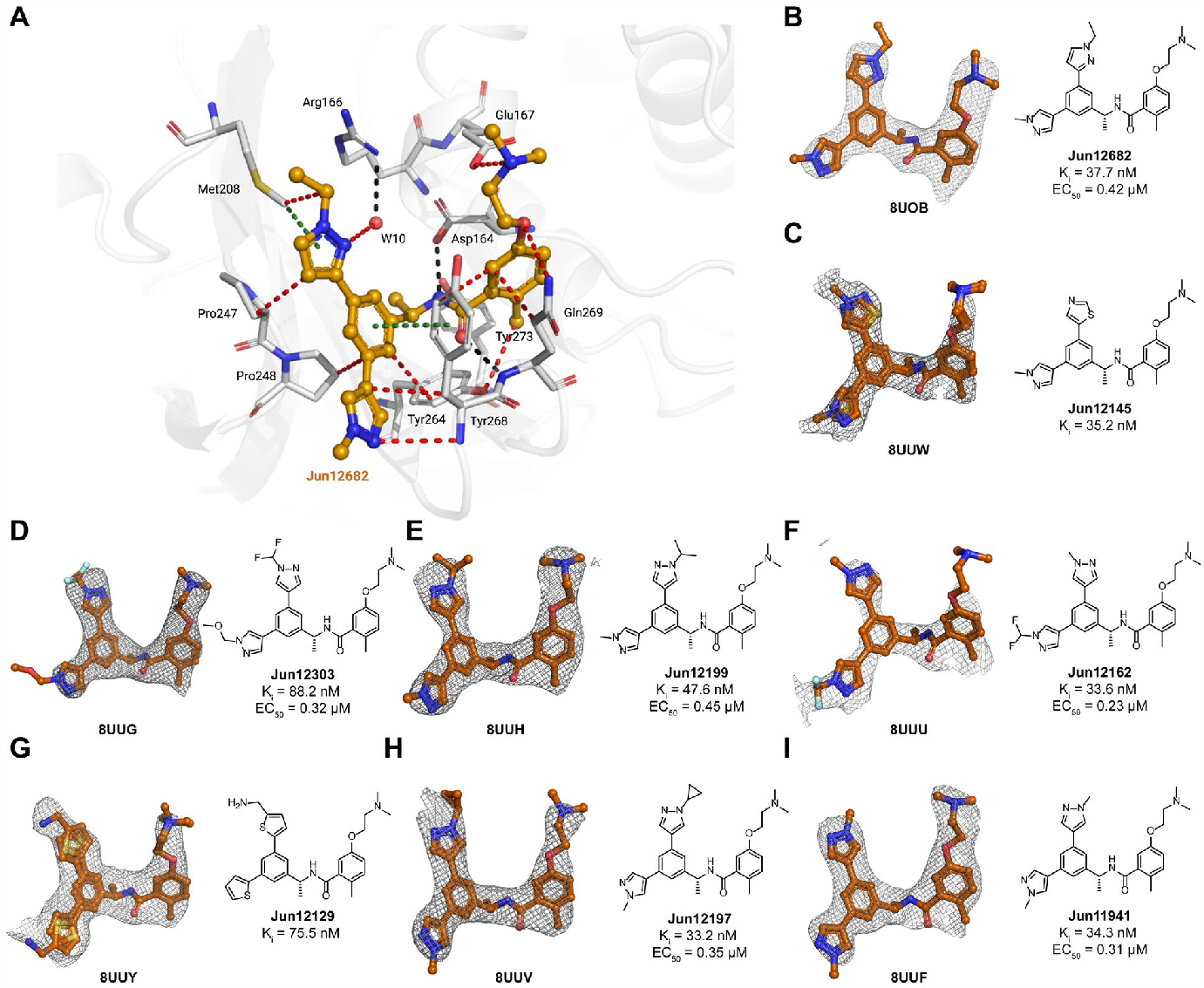
X-ray crystal structures of SARS-CoV-2 PL^pro^ with biarylphenyl PL^pro^ inhibitors. (**A**) Atomic model of the **Jun12682** (in orange sticks and spheres) binding site in SARS-CoV-2 PL^pro^ (residues within 5 Å of the inhibitor are shown in light gray sticks), with hydrogen bonds displayed in black dashed lines, van der Waals contacts as red dashed lines, and π–π interactions as light green dashed lines. (**B**-**I**) Gallery of the polder maps of the indicated inhibitors, displayed as a grey mesh with contour levels between 2 and 4σ. The PL^pro^ inhibitory constant K_i_ in the FRET enzymatic assay and the SARS-CoV-2 antiviral activity EC_50_ in Caco-2 cells were shown for each compound.

## *In vitro* and *in vivo* pharmacokinetic (PK) profiling identified Jun12682 as an *in vivo* lead candidate

PL^pro^ inhibitors with potent SARS-CoV-2 antiviral activity (EC_50_ ≤ 1 µM) and a high selectivity index (SI > 50) were selected for *in vitro* microsomal stability assay and *in vivo* oral PK studies in mice. Most pyrazole-containing PL^pro^ inhibitors showed high stability in the mouse microsomal stability assay (T_1/2_ > 60 min) (Table 1). Next, ten compounds were advanced to the *in vivo* oral snap PK in C57BL/6J mice (Fig. 4A, B; table S2). Compounds were dosed to 3 male C57BL/6J mice per group at 50mg/kg through oral gavage, and blood samples were collected at 0.5, 1, 3, and 5 h, and the drug concentration was quantified by LC-MS/MS. **Jun12199** and **Jun12682** showed the highest *in vivo* plasma concentrations. Given the faster absorption of **Jun12682** compared to **Jun12199, Jun12682** was selected as an *in vivo* lead candidate. A 24-hour *in vivo* PK study was conducted to determine the oral bioavailability of **Jun12682** (Fig. 4C; table S3). **Jun12682** had a rapid absorption after p.o. administration (50 mg/kg) and reached the maximum plasma concentration at 1.67 h (T_max_), with a peak plasma concentration (C_max_) of 4537 ng/mL. The clearance of **Jun12682** was moderate, with a half-life (t_1/2_) of 2.01 h (table S3). The plasma drug concentration was above the antiviral EC_90_ value (1.59 µM) for 7 h. The oral bioavailability of **Jun12682** was 72.8%.

**Fig. 4.**
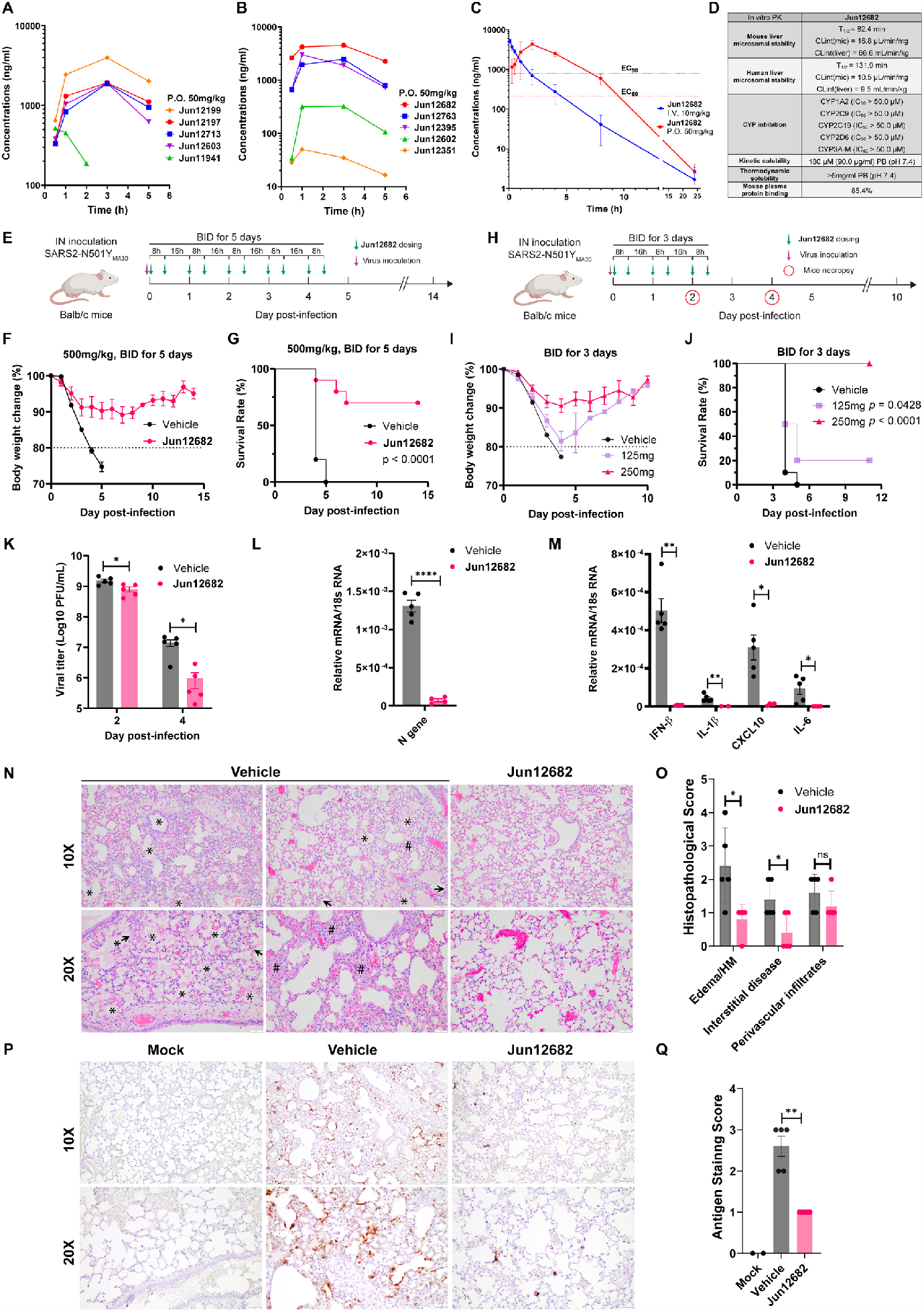
*In vitro* and *in vivo* PK profiling of PL^pro^ inhibitors, and *in vivo* antiviral efficacy of Jun12682. (**A**) Plasma drug concentration of **Jun12199, Jun12197, Jun12713, Jun12603**, and **Jun11941** in C57BL/6J mice (6 – 8 weeks old) following p.o. administration of 50 mg/kg of compound in 0.5% methylcellulose and 2% Tween 80 in water. (**B**) Plasma drug concentration of **Jun12682, Jun12763, Jun12395, Jun12602**, and **Jun12351** in C57BL/6J mice (6 – 8 weeks old) following p.o. administration of 50 mg/kg of compound in 0.5% methylcellulose and 2% Tween 80 in water. (**C**) Plasma drug concentration of **Jun12682** in C57BL/6J mice (6 – 8 weeks old) following p.o. administration of 50 mg/kg and i.v. injection of 10 mg/kg. (**D**) *In vitro* PK parameters of **Jun12682**. (**E**) Experimental design for the 5-day treatment experiment. Ten mice per group were intranasally (IN) inoculated with 5,600 PFU of SARS2-N501Y_MA30_ and subsequently orally administered 500 mg/kg **Jun12682** or vehicle twice a day (BID) for five days (BID_5). (**F**) Body weight loss and (**G**) survival rate of the BID_5 mice experiment. (**H**) Experimental design for the 3-day treatment experiment. Ten mice per group were intranasally (IN) inoculated with 5,600 PFU of SARS2-N501Y_MA30_ and subsequently orally administered 125, 250 mg/kg **Jun12682** or vehicle BID for three days (BID_3). (**I**) Body weight loss and (**J**) survival rate of the BID_3 mice experiment. Data in **F, G, I**, and **J** are pool of two independent experiments (n=10) and are shown as mean ± s.e.m. The p values in **G** and **J** were determined using a log-rank (Mantel–Cox) test. (**K**) Viral titers in lungs collected at 2 and 4 DPI from vehicle- or 250 mg/kg **Jun12682**-treated mice (n=5 each group). Data are mean ± s.e.m and analyzed with unpaired t test with Welch’s correction. *, p< 0.05; ***, p<0.001. (**L-M**) Quantitative PCR analysis of viral nucleocapsid gene (**L**) and cellular cytokines (**M**) in lungs collected at 2 DPI from vehicle- or 250mg/kg **Jun12682**-treated mice (n=5 each group). (**N**-**Q**) Lungs collected at 4 DPI from vehicle- or 250mg/kg **Jun12682**-treated mice (n=5 each group) were stained with haematoxylin and eosin (H&E) (**N**) or immunostained for SARS-CoV-2 nucleocapsid (**P**), and the pathological lesions and staining were quantified (**O** and **Q**, respectively). **N**, H&E stained lungs from vehicle-treated infected mice exhibited airway edema (asterisks), hyaline membranes (HM, arrowheads), and interstitial thickness (number sign). Scale bars, 100 μm (top) and 50 μm (bottom). **O**, Summary scores of lung lesions (n = 5 for each group). **P**, Lungs from vehicle- or **Jun12682**-treated mice (n = 5 for each treatment group) were immunostained to detect SARS-CoV-2 nucleocapsid protein. Scale bars, 100 μm (top) and 50 μm (bottom). **Q**, Summary scores of nucleocapsid immunostaining of lungs. Data in **L, M, O** and **Q** are mean ± sem and analyzed with unpaired t test with Welch’s correction. ns, not significant; *, p<0.05; **, p<0.01; ***, p<0.001; ****, p<0.0001.

Further profiling of the *in vitro* PK properties revealed that **Jun12682** was highly stable in the human microsomes (T_1/2_ = 131.9 min, CLint(mic) = 10.5 µL/min/mg) and had high selectivity for CYP1A2, 2C9, 2C19, 2D6, and 3A-M (IC_50_ > 50.0 µM) (Fig. 4D). **Jun12682** had a kinetic solubility of 180 µM and a thermodynamic solubility greater than 5mg/ml. The mouse plasma protein binding was 85.4%. Overall, **Jun12682** showed favorable *in vitro* and *in vivo* PK properties amenable for the *in vivo* antiviral efficacy study.

## *In vivo* antiviral efficacy of Jun12682 in a SARS-CoV-2 infection mouse model

To assess the *in vivo* antiviral efficacy of **Jun12682**, we utilized a lethal SARS-CoV-2 mouse model described in (*30*). This model involves infecting young Balb/c mice (9 to 12 weeks old) with a mouse-adapted SARS2-N501Y_MA30_, resulting in severe lung disease resembling human patients’ lung injuries. This model has been widely used for evaluating SARS-CoV-2 therapeutics and vaccine candidates (*31-34*). To this end, young Balb/c mice were intranasally infected with 5,600 PFUs of SARS2-N501Y_MA30_ and orally administered **Jun12682** at 500 mg/kg twice daily for 5 days (Fig. 4E). The weight loss plot (Fig. 4F) illustrates that mice administered with the vehicle experienced rapid body weight loss exceeding 20%, leading to a 100% fatality rate by 5 DPI (day post inoculation) (Fig. 4G). In contrast, most mice treated with **Jun12682** exhibited reduced weight loss, resulting in a significantly improved survival rate (0% vs. 70%, p < 0.0001) (n = 10, two independent studies). To further evaluate **Jun12682**’s *in vivo* efficacy, two lower dosages (125, 250 mg/kg) were tested with reduced dosing times from 5 days to 3 days twice daily (Fig. 4H). Mice treated with a dose of 250 mg/kg showed an average of 10% maximum weight loss, providing evident protection from infection compared to the vehicle- and 125 mg/kg-treated groups, which exhibited over 20% weight loss (Fig. 4I). Survival analyses demonstrated that **Jun12682**-treated mice had statistically higher survival rates compared to the vehicle group (0% survival): 125 mg/kg (20%, p = 0.0428), and 250 mg/kg (100%, p < 0.0001) (Fig. 4J). The three-day treatment of 250 mg/kg dose conferred even better protection than the five-day treatment of 500 mg/kg dose, possibly due to reduced drug toxicity. Lung viral load analyses revealed that at 2 DPI, the vehicle-treated mice had robust infections in the lungs (mean lung titer of log_10_ 9.17 ± 0.124 PFUs/ml), while the 250 mg/kg **Jun12682**-treated mice had statistically lower lung viral titers (mean lung titers of log_10_ 8.87 ± 0.194 PFUs/ml, p = 0.0199) (Fig. 4K). The antiviral effect of the 250 mg/kg treatment was more evident at 4 DPI with over a log viral titer reduction comparing to the vehicle treatment (mean lung titers of log_10_ 7.05 ± 0.401 and log_10_ 5.73 ± 0.528 PFUs/ml for the vehicle and 250 mg/kg groups, respectively, p = 0.00247) (Fig. 4K), corroborating the weight loss and survival data (Fig. 4, I and J).

Quantitative PCR analysis of the RNA samples extracted from 2 DPI mice lungs showed that the 250mg/kg treatment significantly reduced the viral N gene level (Fig. 4L) and the expression of multiple inflammatory cytokines, including IFN-β, IL-1β, IL-6, and CXCL10 (*35*) (Fig. 4M). Histopathological analysis revealed that lungs from the vehicle-treated, SARS2-N501Y_MA30_–infected mice at 4 DPI exhibited multifocal pulmonary lesions, including lymphocytic perivascular cuffing, pulmonary edema, hyaline membrane formation, and interstitial thickening and inflammation compared with **Jun12682**-treated mice (Fig. 4N, O; fig. S5). Immunohistochemical analysis using a monoclonal antibody to detect SARS-CoV-2 nucleocapsid (N) in the lungs demonstrated strong and expansive antigen staining in lungs from vehicle-treated, infected mice, whereas **Jun12682** treatment considerably decreased viral antigen staining levels with a few sporadic positive cells (Fig. 4P, Q; fig. S6), consistent with the lung viral titer results (Fig. 4K). Mock-infected lungs were negative for nucleocapsid staining. Overall, the reduced viral replication in the lung and the expression of inflammatory cytokines (Fig. 4K to M) corroborates with the reduced lung inflammation and N protein staining at 4 DPI (Fig. 4N to Q). In summary, these findings demonstrate that oral administration of our rationally designed PL^pro^ inhibitor **Jun12682** efficiently inhibited SARS-CoV-2 replication and mitigated SARS-CoV-2 induced lung lesions *in vivo*, and it represents a promising candidate for further development as orally bioavailable SARS-CoV-2 antivirals. PL^pro^ inhibitors can be used alone or in combination with existing RdRp and M^pro^ inhibitors to combat SARS-CoV-2 variants and drug-resistant mutants.

## Acknowledgments

We thank Shannon Cowan and Caden Miller at the Immunopathology Core of the Oklahoma Center for Respiratory and Infectious Diseases for their technical assistance.

## Funding

National Institutes of Health grant U19AI171110 (J.W., E.A.)

National Institutes of Health grant R01AI158775 (J.W., X.D.)

National Institute of Food and Agriculture grants 2023-67015-39096 and 2023-70432-39482 (X.D.)

USDA-ARS Non-Assistance Cooperative Agreement 58-5030-3-047 (X.D.)

Center for Advancement of Science & Technology (OCAST) grant HR23-096 (X.D.)

## Author contributions

J.W., X.D., and E.A. conceived and supervised the research and designed the experiments; J.W., B.T., and P.J. designed the inhibitors; B.T., and P.J. performed chemical syntheses, separation, purification, and structural characterizations; A.A., A.C., and F.X.R. performed gene expression, protein purification, crystallization, and diffraction data collection; A.A., A.C., and F.X.R., and E.A. determined and analyzed the crystal structures; B.T., H.T., and K.L. performed enzymatic inhibition assays, DSF assays, and cellular cytotoxicity assays; X.Z. performed *in vitro* cellular antiviral assays and *in vivo* antiviral studies; A.F. performed the histopathology and immunohistochemistry (IHC) assessment; X.C. performed the mouse tissue analysis and generated the recombinant SARS-CoV-2 viruses; J.W., B.T., X.Z., A.A., A.F., X.C., F.X.R., X.D., and E.A. analyzed and discussed the data with the assistance of P.J., H.T., K.L., and A.C.; and J.W., A.A., F.X.R., E.A., and X.D. wrote the manuscript with the assistance of B.T., X.Z., P.J., H.T., K.L., A.C., and A.F.

## Data and materials availability

All data are available in the main text or the supplementary materials. The PDB accession numbers for the coordinates of SARS-CoV-2 PL^pro^ in complex with PL^pro^ inhibitors are 8UUW (**Jun12145**), 8UUY (**Jun12129**), 8UUV (**Jun12197**), 8UUU (**Jun12162**), 8UUH (**Jun12199**), 8UUG (**Jun12303**), 8UUF (**Jun11941**), 8UOB (**Jun12682**), and 8UVM (**Jun11313**).

## Supplementary Materials

Materials and Methods

Supplementary Text

Figs. S1 to S6

Tables S1 to S3

References (*37–50*)

