## Supplemental File for "Design of SARS-CoV-2 papain-like protease inhibitor with antiviral efficacy in a mouse model"

**The PDF file includes:**

Materials and Methods
Supplementary Text
Figs. S1 to S6
Tables S1 to S3
References (*37-50*)

#### Materials and Methods

*Cell lines and virus*

A Vero-E6 cell line, a gift from Dr. Susan Baker (Loyola University Chicago), was grown in Dulbecco's modified Eagle medium (DMEM) (Corning, 10013CM) containing 10% heat-inactivated fetal bovine serum (FBS) (Gibco, 10-438-026), 1% Pen/Strep (30-001-CI), 1× nonessential amino acid (NEAA) (Cytiva HyClone, SH30238.01). A Vero-E6 line expressing human angiotensin-converting enzyme 2 (hACE2) and human transmembrane protease, serine 2 (hTMPRSS2) (Vero-AT), obtained through NIH-BEI Resources (NR-54970) were grown in DMEM containing 10% FBS, 1% Pen/Strep, 1× NEAA, and 10 μg/mL puromycin (InVivogen, ant-pr-1) to maintain the expression of hTMPRSS2 and hACE2. A Caco-2 line expressing hACE2 and hTMPRSS2 (Caco2-AT) (*37*), a gift from Dr. Mohsan Saeed (Boston University), was propagated in DMEM containing 10% FBS, 1% Pen/Strep, 1× NEAA, 1 μg/mL puromycin, and 1 μg/mL blasticidin (InVivogen, ant-bl-05).

The mouse-adapted SARS2-N501Y$_{MA30}$ virus, a gift from Dr. Stanley Perlman (University of Iowa), was propagated once with Vero-AT cells and titrated with Vero-E6 cells. The following SARS-CoV-2 strains/isolates were obtained through BEI Resources, NIAID, NIH: Washington strain 1 (WA1) (NR-52281), recombinant SARS-CoV-2 expressing nano-luciferase reporter (icSARS-CoV-2-nLuc) (NR-54003), Delta strain (NR-55672), Omicron BA.2 strain (NR-56520). These viruses were propagated once with Vero-AT cells to obtain large viral stocks and were titrated with Vero-AT cells. The following recombinant SARS-CoV-2 viruses were generated previously (PMID: 37637734) or in this study using a reverse genetics system developed by Dr. Ralph Baric's group at the University of North Carolina at Chapel Hill (BEI, NR-52281): recombinant wild-type SARS-CoV-2 WA1 strain (rSARS-CoV-2), and recombinant Nsp5/Mpro mutant viruses (rNsp5$^{S144M}$, rNsp5$^{L50F/E166V}$, and rNsp5$^{L50F/E166V/L167F}$). These viruses were titrated with Vero-E6 cells and full-genome sequenced using the ARTIC method (*38, 39*).

*Biocontainment*

All procedures with live SARS-CoV-2 were performed in certified biosafety level 3 (BSL3) facilities at Oklahoma State University using biosafety protocols approved by the Institutional Biosafety Committee (IBC), which comprises scientists, biosafety and compliance experts, and members of the local community. All research personnel received rigorous biosafety, biosecurity, and BSL3 training before participating in experiments. Personal protective equipment, including scrubs, disposable overalls, shoe covers, double-layered gloves, and powered air-purifying respirators, were used. Biosecurity measures are built in the environment through building and security systems and are reinforced through required training programs, standing meetings, and emergency exercises. The researchers involved in working with live viruses received the SARS-CoV-2 mRNA vaccine before the study was started. Finally, all researchers were medically cleared by the Oklahoma State University Occupational Health Program.

*Enzymatic Assays*

The SARS-CoV-2 PL$^{pro}$ enzymatic assays were carried out as follows: The assay was carried out in 96-well plates with 100 μL of 200 nM PL$^{pro}$ protein in PL$^{pro}$ reaction buffer [50 mM HEPES (pH 7.5), 0.01% Triton X-100, and 5 mM DTT]. Then, 1 μl of testing compound at various concentrations was added to each well and incubated at 30°C for 30 min. The enzymatic reaction was initiated by adding 1 µL of 1 mM FRET substrate Dabcyl-FTLRGG/APTKV-Edans (the final substrate concentration is 10 µM). The reaction was monitored in a Cytation 5 image reader with filters for excitation at 360/40 nm and emission at 460/40 nm at 30°C for 1 hour. The initial velocity of the enzymatic reaction with and without testing compounds was calculated by linear regression for the first 15 min of the kinetic progress curve. The $IC_{50}$ values were calculated by plotting the initial velocity against various concentrations of testing compounds with a dose-response function in Prism 8 software.

For determination of $K_m$ and $V_{max}$, $PL^{pro}$ was diluted in the reaction buffer (50 mM Hepes (pH 7.5), 0.01% Triton X-100, and 5 mM DTT) to the final concentration of 200 nM, and various concentrations (1.5625, 3.125, 6.25, 12.5, 25, 50, 100, 200 µM) of $PL^{pro}$ FRET substrate were added to initiate the reaction. The reaction was monitored every 71 seconds for 1 hour at 30 °C in Cytation 5 image reader with excitation wavelength at 360/40 nm and emission wavelength at 460/40 nm. The initial velocity of the enzymatic reaction at each concentration of FRET substrate was determined in the first 15 min by linear regression. The $K_m$ and $V_{max}$ were determined by fitting the curves with nonlinear regression of initial velocity vs. concentration of FRET substrate using the Michaelis-Menten equation in Prism 8.

For the $K_i$ measurement, 10 µL 200 nM SARS-CoV-2 $PL^{pro}$ protein was added to 190 µl of $PL^{pro}$ reaction buffer containing testing compound and the FRET substrate, and the reaction was monitored for 2 h. The final FRET substrate concentration in this assay is 20 µM. Detailed curve fitting and $K_i$ determination was described previously (*40, 41*). The equation used for Morrison plot curve fitting is: $Y=V_o*(1-((((E_t + X+(K_i*(1+(S/K_m))))-(((E_t + X+(K_i*(1+(S/K_m))))\^2)-4*E_t*X)\^0.5))/(2*E_t)))$. Y: enzyme activity; $V_0$: velocity in the absence of inhibitor; $E_t$: enzyme concentration (0.2 µM); X: concentration of inhibitor (µM); $K_i$: dissociation of inhibitor (µM); S: substrate concentration (20 µM); $K_m$: the concentration of substrate which permits the enzyme to achieve half $V_{max}$ (34.08 µM).

The enzymatic assay of SARS-CoV-2 $PL^{pro}$ with Ub-AMC (UBPBIO, M3010) substrate was carried out in 384-well plate format. compound with various concentrations was mixed with 50 nM SARS-CoV-2 $PL^{pro}$ dissolved in $PL^{pro}$ reaction buffer. After incubation at 30°C for 15 min, the reaction was initiated by adding 2.5 µM Ub-AMC substrate. The fluorescence was monitored using the excitation of 360/40nm and emission of 460/40nm at 30°C for 1 hour. The first 1000 seconds of initial velocity was calculated for plotting $IC_{50}$. For the $K_i$ measurement, compound with various concentrations was mixed with 2.5 µM Ub-AMC substrate in $PL^{pro}$ reaction buffer, and the reaction was initiated by the addition of 10 nM SARS-CoV-2 $PL^{pro}$. The fluorescence was monitored immediately in the plate reader for 4 hours. The first 4000 seconds of initial velocity was calculated for plotting the $K_i$.

The enzymatic assay of SARS-CoV-2 $PL^{pro}$ with ISG15-AMC (R&D, UL553050) substrate was carried out in 384-well plate format. Compound with various concentrations was mixed with 2 nM SARS-CoV-2 $PL^{pro}$ in $PL^{pro}$ reaction buffer. After incubation at 30°C for 15 min, the reaction was initiated by adding 0.5 µM ISG15-AMC substrate. The fluorescence was monitored using the excitation of 360/40nm and emission of 460/40nm at 30°C for 1 hour. The first 500 seconds of initial velocity was calculated for plotting the $IC_{50}$. For the $K_i$ measurement, compound with various concentrations was mixed with 0.1 µM ISG15-AMC substrate in $PL^{pro}$ reaction buffer, and the reaction was initiated by adding 0.2 nM SARS-CoV-2 $PL^{pro}$. The fluorescence was monitored immediately in the plate reader with the same condition for 4 hours. The first 2500 seconds of initial velocity was calculated for plotting the $K_i$.

The counter screening of the compound against USP7(R&D, E519025) and USP14 (Enzo, BMLUW98400100) were performed in 384-well plate format. For the USP7 assay, compound with various concentrations was mixed with 40 nM USP7 dissolved in the USP7 reaction buffer (HEPES 50 mM, pH 7.5, BSA 0.1 mg/mL, Triton X-100 0.01%, DTT 5mM, Glycerol 5%). After incubation at 30°C for 15 min, the reaction was initiated by adding 5 µM Ub-AMC substrate. For the USP14 assay, the procedure was similar, with 1.7 µM of USP14 assayed and 4 µM Ub-AMC. The fluorescence was monitored in Cytation 5 plate reader with excitation of 360/40nm and emission of 460/40nm at 30°C for 1 hour. For both assays, the 3000 seconds of initial velocity was calculated and normalized with that of the DMSO-treated group to determine the percentage inhibition.

*Differential Scanning Fluorimetry (DSF)*

The thermal shift assay was performed using Thermo Fisher QuantStudio 5 Real-time PCR system. Compounds with various concentrations were mixed with 4 µM SARS-CoV-2 $PL^{pro}$ in the reaction buffer (HEPES 50mM, pH 7.5, DTT 5mM, and Triton X-100 0.01%). After incubation at 30°C for 1 hour, 1x SYPRO orange dye was added. The fluorescence was monitored under a temperature gradient from 25 °C to 95 °C at an increment of 0.05 °C/s. The melting temperature ($T_m$) was determined by the mid-log of the transition phase from the native to denatured protein by the Boltzmann model in Protein Thermal Shift Software v1.3.

*Cell-Based FlipGFP-$PL^{pro}$ Assay*

Plasmid pcDNA3-TEV-flipGFP-T2A-mCherry was ordered from Addgene (catalog no.124429). SARS-CoV-2 $PL^{pro}$ cleavage site LRGGAPTK was introduced into pcDNA3-FlipGFP-T2A-mCherry via overlapping PCRs to generate a fragment with SacI and *Hin*dIII sites at the ends. SARS-CoV-2 $PL^{pro}$ expression plasmid pcDNA3.1 SARS2 $PL^{pro}$ was ordered fromGenscript (Piscataway NJ) with codon optimization. For transfection, 96-well Greiner plate (catalog no. 655090) was seeded with HEK293T cells to overnight 70-80% confluency. A total of 9 µL of Opti-MEM, 0.1 µL of 500 ng/µL pcDNA3-flipGFP-T2A-mCherry plasmid, 0.1 µL of 500 ng/µL protease expression plasmid pcDNA3.1, and 0.3 µL of transIT-293 (Mirus) were used each well of a 96-well plate. Three hours after transfection in a cell culture incubator (humidified, 5% $CO_2$/95% air, 37°C), 1 µL of testing compound was added to each well at 100-fold dilution. Images were acquired 48h after transfection with a Celigo Image Cytometer (Nexcelom) and were analyzed with Gen5 3.10 software (Biotek). SARS-CoV-2 $PL^{pro}$ protease activity was calculated by the ratio of GFP signal sum intensity over the mCherry signal sum intensity. The FlipGFP- $PL^{pro}$ assay $IC_{50}$ value was calculated by plotting the GFP/mCherry signal over the applied compound concentration with a four-parameter dose-response function in Prism 8. The mCherry signal alone was utilized to determine the compound cytotoxicity (*28*).

*Antiviral assays with live SARS-CoV-2*

Three assays were employed for antiviral effect evaluation: cell viability assay, reporter virus assay, and antiviral plaque assay. For the cell viability assay, Vero-AT cells at a density of $1.5x10^4$ cells/well were batched inoculated with SARS-CoV-2 at a multiplicity of infection (MOI) of 0.2 and then added to 96-well plates (Corning, #3598) in 50 µL per well of DMEM containing 4% FBS and 2µM CP-100356, a P-glycoprotein inhibitor. For the reporter virus assay, Caco2-AT cells at a density of $1.5x10^4$ cells/well were batched inoculated with icSARS-CoV-2-nLuc at a MOI of 0.2 and then added to 96-well plates in 50 µL per well of DMEM containing 4% FBS and 2 µM

CP-100356. To prepare compound solutions, the test compounds and positive control (nirmatrelvir) were 3-fold serially diluted in DMEM, starting at 30 µM final concentration for the test compounds or 3.3 µM for the positive control. After dilution, 50 µL diluted compound was transferred and mixed (1:1 volume ratio) with the cell-virus mixture that was pre-seeded in 96-well plates. Cells were incubated for 24 hours (Caco2-AT) or 48 hours (Vero-AT) at 37 °C with 5% $CO_2$ and then subjected to a Celltiter-Glo cell viability assay (Promega, G9243) or Nano-Glo Luciferase assay (Promega, N1130) on a Promega GloMax Discover microplate reader (Promega, GM3000) following the instruction of the manufacturer.

For the antiviral plaque assay, Vero-AT cells were seeded in 12-well plates a day prior to infection and the drug dissolved in DMSO was serially diluted in DMEM with 3-fold dilutions between test concentrations, starting at 10 µM final concentration. Cells in 12-well plates were incubated with approximately 20 plaque-forming units (PFUs) per well of each virus for 1 hour. After incubation, the inoculum was removed and 1 mL 1X DMEM-1.2% Avicel (FMC polymers) mixture containing serially-diluted compound and 2uM P-glycoprotein inhibitor, CP-100356 was added to each well. After 48 hours of incubation at 37°C, the DMEM-Avicel mixture was removed and the cells were stained using 0.1% crystal violet solution. Plates were photographed and measured for the area of cells affected by infection using ImageJ.

*Mouse experiments*

All mouse experiments with SARS-CoV-2 were performed in a certified biosafety level 3 (BSL3) facility at Oklahoma State University. All animal studies were reviewed and approved by the Oklahoma State University Animal Care and Use Committee and met stipulations of the Guide for the Care and Use of Laboratory Animals. Nine to twelve-week-old female Balb/c mice (Strain #: 000651) were procured from the Jackson laboratory. Mice were briefly anesthetized with isoflurane and inoculated intranasally (i.n.) with 5600 PFUs of SARS2-N501$Y_{MA30}$ in a total volume of 50 µl DMEM. PLpro inhibitor (Jun-12-68-2) was dissolved with 0.5% methylcellulose solution containing 2% Tween-80 and was administered via oral gavage using 20G/30mm plastic feeding tubes (Instech, FTP203050) for 3 or 5 days after virus inoculation. Animal weight and health were monitored daily. A group of 5 mice for each treatment were euthanized at 2- and 4 days post-infection (DPI) for necropsy. The left lungs were collected in pre-filled bead tubes (Fisher Scientific, 15-340-153) filled with 1 mL DMEM for viral load determination, and the rest of the lungs were transfused with 1 mL zinc-buffered formalin solution (Fisher Scientific, STLBFZ1) and then removed and fixed with 30 ml zinc-buffered formalin solution for at least two days before removal from BSL3 in accordance with an approved IBC protocol.

*Lung viral titer by plaque assay*

The lung tissues were homogenized in pre-filled bead tubes using an automated homogenizer (Fisherbrand™, 15-340-164), and followed by centrifugation (400 xg for 5 minutes). The clarified tissue homogenate supernatants were aliquoted and stored at -80 °C or subjected to standard plaque assay with Vero-AT cells. The supernatants were serially diluted in DMEM and inoculated onto Vero-AT cells in twelve-well plates and maintained at 37 °C in 5% $CO_2$ for 1 hour with gentle rocking every 15 min. After removing the inoculum, plates were overlaid with 1.2 % agarose (Fisherbrand, BP160-500)-1X DMEM mix containing 2% FBS. After 2 days, overlays were removed, and plaques were visualized by staining with 0.1% crystal violet. Viral titers were quantified as PFUs per ml tissue.

*Histology and immunohistochemistry (IHC)*

Fixed tissues were trimmed, and processed using a Leica ASP300s processor (Leica Biosystems, Leico Biosystems Inc., 1700 Leider Lane, Buffalo Grove, IL 60089 United States) on a delayed short cycle program and embedded in paraffin (Leica Surgipath Paraplast Infiltration and Embedding Medium; Leica Biosystems). Paraffin blocks were cut into 4 µm-thick sections and mounted on VistaVision HistoBond adhesive glass slides from VWR (Radnor, PA). Hematoxylin and eosin (H&E) staining was performed following standard operating procedures with the Sakura Finetek DRS601 (Sakura Finetek USA, Inc., 1750 West 214th Street, Torrance, CA 90501). For IHC, slides were rehydrated with water, following HIER (Heat Induced Epitope Retrieval) performed at 95C$^o$ for 20 minutes in Citrate Unmasking solution (H-3300, Vector Laboratories, Newark, CA). SARS-CoV-2 Nucleocapsid antibody [HL448] (Genetex, GTX635686) was diluted 1:5000 in TBS-Tween 20 buffer with 10% normal goat serum and slides were incubated for 1 hour at room temperature (RT). Slides were washed in TBS-Tween 20 buffer and then quenched of endogenous peroxidase using 0.3% $H_2O_2$ for 10 minutes. Slides were washed and detection was carried out using VECTASTAIN® Elite® ABC-HRP Kit, Peroxidase (Rabbit IgG) (Vector Laboratories, PK-6101) per the manufacturer's instructions. Hematoxylin diluted 1:10 was used as a counterstain. HE stained tissues were evaluated by a board-certified veterinary pathologist for three parameters: presence of edema or hyaline membranes, perivascular lymphoid inflammation, and interstitial pneumonia. Edema or hyaline membranes were evaluated using a distribution-based ordinal scoring with 0, none; 1, <25%; 2, 26-50%, 3, 51-75%, 4, >75% of tissue affected. Perivascular lymphoid inflammation and interstitial pneumonia were evaluated using a severity-based ordinal scoring system on a scale of zero to three: 0 (absent), 1 (mild), 2 (moderate), 3 (severe). IHC was ordinally scored on the percentage distribution of staining in the tissues: 0, absent; 1, 0–25%; 2, 26–50%; 3, 51–75%; 4, >75% of tissue.

*RNA extraction and Real-time PCR quantification*

Total RNA was extracted from the lung homogenate supernatants using TRIzol (Invitrogen, 15596018) and followed by RNA purification using the ReliaPrep™ RNA Cleanup/Concentration Kit (Promega, Z1073). A total of 1000 ng RNA was used for cDNA synthesis using the RT² HT First Strand Kit (QIAGEN, 330411) which contains a component to eliminate genomic DNA contamination. Quantitative PCR was performed with specific primers (Table S1) using PowerUp SYBR Green Master mix (Fisher, A25918) on QuanStudio 6 Pro (ThermoFisher, A43160). Cycle threshold values were normalized to 18S RNA levels by using the $2^{-\Delta Ct}$ method.

| Gene | Forward (5'->3') | Reverse (5'->3') |
|---|---|---|
| SARS-CoV-2 N gene | AAGCTGGACTTCCCTATGGTG | CGATTGCAGCATTGTTAGCAGG |
| Mouse IL-6 | GCTACCAAACTGGATATAATCAGGA | CCAGGTAGCTATGGTACTCCAGAA |
| Mouse IL-1b | TGGACCTTCCAGGATGAGGACA | GTTCATCTCGGAGCCTGTAGTG |
| Mouse CXCL10 | GCTTCCCTATGGCCCTCATT | GCCGTCATTTTCTGCCTCAT |
| Mouse IFN-b | TCAGAATGAGTGGTGGTTGC | GACCTTTCAAATGCAGTAGATTCA |

*In vitro PK studies*

The *in vitro* PK studies were performed by Wuxi Apptec.

**1. Plasma Protein Binding Assay-HTD Method**

1. The test compounds or positive control warfarin were spiked into frozen plasma at the final concentration of 2 μM.
2. An aliquot of 150 μL of the compound-spiked plasma sample was added to one side of the chamber in a 96-well equilibrium dialysis plate (HTD dialysis), and an equal volume of dialysis buffer was added to the other side of the chamber. An aliquot of plasma sample was harvested before the incubation and used as T0 samples for recovery calculation. Triplicate incubations were performed.
3. The plate was then incubated in a humidified incubator with 5% $CO_2$ at 37°C for 4 hours.
4. After incubation, 50 μL samples were taken from the plasma side as well as the buffer side.
5. The plasma sample was mixed with an equal volume of blank buffer; buffer samples were mixed with an equal volume of blank plasma.
6. The matrix-matched samples were quenched with stop solution containing internal standard (IS).
7. Samples were analyzed by LC-MS/MS. Test compound concentrations in plasma and buffer samples were determined based on peak area ratio of analyte to IS without a standard curve.

**2. Microsomal CYP Inhibition (IC50 determination) Using 5 in 1 Cocktail Substrates**

1) CYP450 enzymatic activities were determined using 5in1 probe substrate cocktail. For each reaction, enzymatic activities in the presence of the test compound at 8 concentrations (e.g. 0.00, 0.050, 0.150, 0.500, 1.50, 5.00, 15.0, or 50.0 μM) were measured in singlet (n=1). A known inhibitor for each isoform, tested at a single concentration (3.00 μM) in duplicate (n=2), was included as positive control.
2) Incubation mixture containing pooled human liver microsomes (Corning, Xenotech, or other qualified vendors; pooled from multiple donors) at 0.200 mg/ml, probe substrates, and standard inhibitors (listed in the following table) or the test compound were warm up at 37.0°C for 10 minutes. The reactions were initiated by the addition of the NADPH (1.00 mM).

| CYP isoform | Probe Substrate | Substrate Final Conc. (μM) | Standard Inhibitor | Inhibitor Final Conc. (μM) |
|---|---|---|---|---|
| 1A2 | Phenacetin | 10.0 | α-Naphthoflavone | 3.00 |
| 2C9 | Diclofenac | 5.00 | sulfaphenazole | 3.00 |
| 2C19 | S-Mephenytoin | 30.0 | (+)-N-3-benzylnirvanol | 3.00 |
| 2D6 | Dextromethorphan | 5.00 | quinidine | 3.00 |
| 3A4 | Midazolam | 2.00 | ketoconazole | 3.00 |

3) After the mixture was incubated at 37.0°C for 10 minutes, ice cold acetonitrile containing the internal standard (IS) was added to terminate the reactions.
4) The metabolites generated from the probe substrates were measured by LC-MS/MS and were assessed based on peak area ratios of analyte/IS.

5) The remaining activity (expressed as % of control activity) was calculated; $IC_{50}$ values of the test compound were determined using SigmaPlot or XLfit with 3- or 4- parameter logistic sigmoidal equation.

**3. Kinetic Solubility Assay**

1. The Kinetic solubility assay employs the shake flask method followed by HPLC-UV analysis. The following were the step-wise procedure:

– Weigh and dissolve the samples in 100% DMSO as the stock solution of 10 mM. About 10 µL (compound/Media) of stock solution was needed in this assay.

– Add the test compounds and controls (10 mM in DMSO, 10 µL/vial) into the buffer (490 µL/well) placed in a Mini-Uniprep filter. Vortex the samples of kinetic solubility for 2 minutes.

– Incubate and shake the solubility solutions on an orbital shaker with 800 rpm at room temperature for 24 hours.

– Centrifuge at 4000rpm, 20 °C for 10 min

– Transfer 400 µL (with or without dilution) of each solubility supernatant into 96- deep well for analysis after the samples were directly filtered by the syringeless filter device

– Determine the test compound concentration of the filtrate using HPLC-UV.

– Inject at least 5 UV standard solutions into HPLC from low to high concentration subsequently and then test the Kinetic solubility supernatant in duplicate.

– Use the QC samples to monitor Kinetic solubility determination process.

**4. Liver Microsome Metabolic Stability Assay (NADPH)**

1) Test compounds were incubated at 37.0°C with liver microsomes (pooled from multiple donors) at 1.00 µM in the presence of NADPH (~1.00 mM) at 0.500 mg/ml microsomal protein.

2) Positive controls include testosterone (3A4 substrate), propafenone (2D6) and diclofenac (2C9). They were incubated with microsomes in the presence of NADPH.

3) Time samples (0, 5, 15, 30, 45, and 60 minutes) were removed, immediately mixed with cold acetonitrile containing internal standard (IS). Test compounds incubated with microsomes without NADPH for 60 min are also included.

4) Single point for each test condition (n=1).

5) Samples were analyzed by LC/MS/MS; disappearance of test compound was assessed base on peak area ratios of analyte/IS (no standard curve).

6) Using the following equation to calculate the microsome clearance:

$$C_t = C_0 \bullet e^{-k_e \bullet t}$$

$$T_{1/2} = \frac{Ln2}{-k_e} = \frac{0.693}{-k_e}$$

$$CL_{\text{int}(mic)} = \frac{0.693}{In\ vitro\ T_{1/2}} \bullet \frac{1}{mg\,/\,mL\ microsomal\ \ protein\ in\ reaction\ system}$$

The mg microsomal protein / g liver weight is 45 for 5 species

The liver weight values will use 40 g/kg, 30 g/kg, 32 g/kg, 20 g/kg and 88 g/kg for rat, monkey, dog, human and mouse, respectively.

The liver clearance will be calculated using CLint(mic) with

$$CL_{\text{int}(liver)} = CL_{\text{int}(mic)} \bullet \frac{mg\ microsomes}{g\ liver} \bullet \frac{g\ liver}{kg\ body\ weight}$$

*<u>In vivo PK studies</u>*

The *in vivo* PK studies were performed by Pharmaron US.

Male C57BL/6J mice (approximately 6-8 weeks old, 20-30 g, supplied by SPF Biotechology Co. Ltd, Beijing, China) were used for the pharmacokinetic studies . Mice (n=3 oral or intraperitoneal per group) each received either oral gavage administration or intraperitoneal administration of test compound. The dose formulation used for oral dosing was 0.5% methylcellulose (400 cP)/2% Tween 80 in water, administered at a dose level of 50 mg/kg. Serial blood samples (ca. 25 µL blood in to tubes containing heparin-Na as anticoagulant) were collected up to h after the start of dosing via direct vein puncture of the dorsal metatarsal vein. The samples were collected via dorsal metatarsal vein at all time points. At the end of the study, the mice were euthanized by inhalation of rising concentrations of carbon dioxide. Blood was centrifuged to yield plasma. Plasma samples were stored at −75°C prior to analysis.

Plasma sample analysis

Plasma samples (10 uL for mouse) were extracted using protein precipitation with 200 µL of acetonitrile containing an analytical internal standard. An aliquot of the supernatant was analysed by reverse phase LC-MS/MS. Samples were assayed against calibration standards prepared in control plasma and the assay was qualified through inclusion of quality control (QC) samples.

PK data analysis from PK studies

PK parameters were obtained from the plasma concentration–time profiles using non-compartmental analysis with Phoenix (WinNonlin) pharmacokinetic software version 8.3.1.5014 (Pharsight, Mountain View, CA).

All the procedures related to animal handling, care, and the treatment in this study were performed according to Animal care and Use Application (AUP) approved by the Institutional Animal Care and Use Committee (IACUC) of Pharmaron following the guidance of the Association for Assessment and Accreditation of Laboratory Animal Care (AAALAC). Pharmaron's IACUC policies are compliance with laws and regulations of local government for animal research, and also compliance with the guidelines of the Guide for the Care and Use of Laboratory Animals.

*<u>Cytotoxicity assay in Vero E6 cells</u>*

The cytotoxicity of compounds was determined using the neutral red uptake assay (*22*). Briefly, 252,000 cells/mL of Vero cells grown in DMEM with 10% fetal bovine serum (FBS) were dispensed into 96-well cell culture plates at 100 µL/well. Twelve hours later, the growth DMEM medium was aspirated and washed with 100 µL PBS buffer. After aspiration, 100 µL fresh DMEM medium with 2% FBS was first added, followed by the addition of 1 µL serially diluted compounds. Another 100 µL fresh DMEM medium with 2% FBS was added as an overlay as soon as the compounds were added. After 48 hours of incubation at 37 °C, the medium was aspirated and replaced with 100 µL serum-free DMEM medium containing 40 µg/mL neutral red and incubated at 37 °C for 2 hours. After aspiration and washing with 100 µL PBS buffer, 100 µL neutral red destain solution was added to extract the neutral red from the cells. Shake the plate till a homogeneous solution formed in each well, the amount of neutral red taken up was determined by measuring the absorbance at 540 nm using a BioTek Microplate Epoch 2 Reader

(Agilent). The $CC_{50}$ values were calculated from best-fit dose-response curves with the variable slope in Prism 9.

Recombinant protein production

The $PL^{pro}$ bacterial expression plasmid was obtained through BEI Resources, NIAID, NIH: Vector pMCSG53 Containing the SARS-Related Coronavirus 2, Wuhan-Hu-1 Papain-Like Protease Gene, NR-52897. Protein expression and purification were done similarly as reported (*42*). Briefly, the pMCSG53-$PL^{pro}$ plasmid was transformed into the *E. coli* BL21(DE3) strain (Invitrogen) and cultured in LB medium supplemented with ampicillin (100 µg/ml). The transformed cells were induced with 1 mM IPTG at an OD600 of 0.8, followed by expression at 16 °C for 18 hours.
Bacterial cells were harvested by centrifugation at 7000 × g and cell pellets were resuspended in a 12.5 ml lysis buffer (500 mM NaCl, 5% (v/v) glycerol, 50 mM HEPES(2-[4-(2-Hydroxyethyl)piperazin-1-yl]ethane-1-sulfonic acid) pH 8.0, 20 mM imidazole pH 8.0, 1 mM TCEP, 1 µM $ZnCl_2$) per liter culture and sonicated at 120 W for 10 min (4 s ON, 20 s OFF). The lysate was clarified by centrifugation at 37000 × g for 60 min at 4 °C. Ni-NTA purification was performed according to the manufacturer's recommendations (Qiagen, Valencia, CA USA) with the lysis buffer. Bound $PL^{pro}$ was eluted with 20 ml of lysis buffer supplemented to 500 mM imidazole pH 7.5, followed by Tobacco Etch Virus (TEV) protease treatment at 1:25 protease:protein ratio at 4 °C overnight, and a reverse Ni-NTA purification. Size exclusion chromatography was performed on a Superdex 200 increase 10/300 GL column equilibrated in lysis buffer. Peak fractions were pooled, buffer-exchanged to a final buffer 150 mM NaCl, 50 mM tris(hydroxymethyl)aminomethane (Tris) pH 7.5, 1 µM $ZnCl_2$, 4 mM TCEP and concentrated at 20 mg/ml. In the case of the PLpro-**Jun11313** structure, the buffer used was identical, except that 20 mM HEPES pH 7.5 was utilized and $PL^{pro}$ was concentrated at 8 mg/ml.

Crystallization of SARS-CoV-2 $PL^{pro}$ with inhibitors

For co-crystallization of the $PL^{pro}$-**Jun11313** complex, 10-fold molar excess of inhibitor was added to $PL^{pro}$ and incubated for 1-2 hr on ice. The complex was then clarified by spinning at 15,000 × g for 20 minutes before the crystallization screening. The sitting drop vapor diffusion method was dispensed with the help of an Oryx8 robot (Douglas Instruments Ltd) in 96-well Intelli-Plate (Art Robbins Instruments). Cocrystallizations were attempted with the protein-to-matrix ratio of 1:1, 2:1 and 1:2 at 4 °C and 20 °C in a focused screen based on the SARS-CoV-2 $PL^{pro}$ PDB deposited conditions. After several crystallization optimization steps, crystals started growing in a hanging drop vapor-diffusion setup consisting of 4µl protein against 2µl well solution. After two weeks, hexagonal shaped cocrystals grew with protein at 8 mg/ml, inhibitor at 2.2 mM and a well solution containing sodium citrate dibasic trihydrate pH 6.7, ammonium sulfate 2.5 M at 4 °C, in the space group $P3_221$ with four copies of the complex in the asymmetric unit.
For cocrystallization with the rest of inhibitors, a similar initial approach was used (sitting drop), except the protein (20 mg/ml) was mixed with inhibitors at 5.7 mM (molar ratio of 2:1), and a well solution containing 0.1 M Bis-Tris pH 5.5-6.5, 200 mM zinc acetate, 8-12% PEG 8000 at 4 °C. Pyramidal shaped crystals (~150µm) grew in 1 hour, belonging to the $I4_122$ space group, with one copy of the complex in the asymmetric unit. Crystals selected for data collection were washed in the crystallization buffer supplemented with 20% glycerol and flash-cooled in liquid nitrogen.

Data collection, structure determination, and refinement

Single-wavelength X-ray diffraction data for **Jun11313**, **Jun12129**, **Jun12145**, **Jun12197** and **Jun 12162** were collected at 100K temperature at the beamline 23-ID-B at Advance Photon Source at Argonne National Laboratory (Lemont, IL), and **Jun12199**, **Jun12303**, **Jun11941** and **Jun12682** were collected at Beamline 17-ID-1(AMX) at NSLS-II at Brookhaven National Laboratory (Upton, NY), respectively. All data sets were collected remotely, followed by crystal rastering before data collection to find the best diffraction spot on the crystals.

The dataset for the $PL^{pro}$-**Jun11313** complex was indexed, integrated, and scaled using HKL2000 (*43*), and the rest of the data sets were processed by using Dials (*44*), Aimless (*45*) or autoPROC from Global Phasing (*46*). The structure coordinates were obtained by molecular replacement method using PDB 7NFV as a template in Phaser (*47*). The obtained initial maps were used to generate the unbiased polder difference map for the inhibitor (*36*). The inhibitor electron density was visualized and examined, and the inhibitor molecules were fitted by using Coot (*48*). To obtain a better model and converging Rwork/Rfree values, the datasets were subjected to refinement iteratively by manual model building in Coot followed by refinement in Phenix (*49*). Solvent molecules were added in the last few refinement cycles. Statistics of diffraction data processing and the model refinement are given in table S1. The inhibitor restraints were generated with the Grade online server, and the interactions were assessed with the help of the PLIP online server (*50*) and manually in Coot. PLpro-inhibitor complex structures were analyzed, and the figures showing the protein-inhibitor cocrystal structures were made with PyMOL.

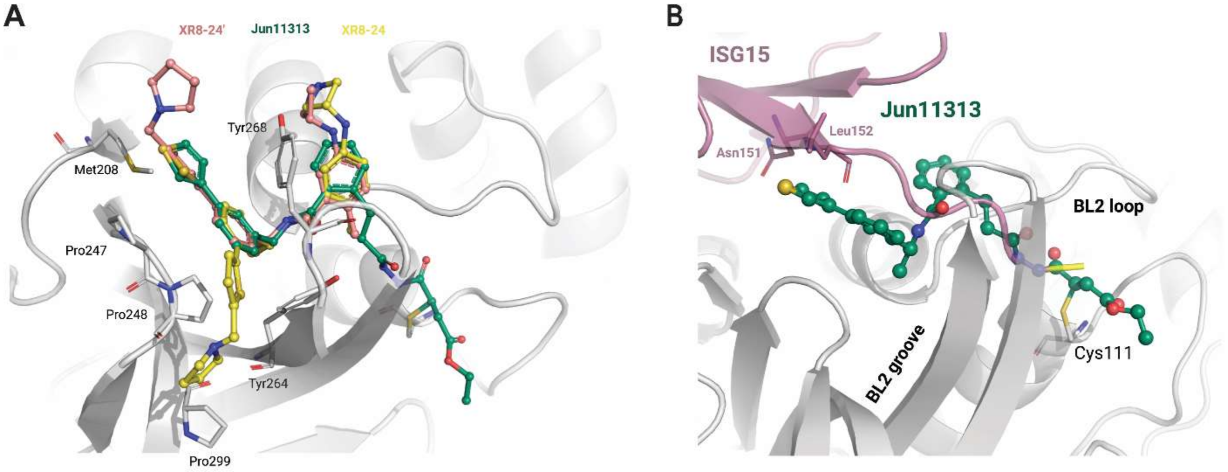


**Fig. S1. Comparison of Jun11313**, **XR8-24, and ISG15 binding to SARS-CoV-2 PL$^{pro}$.** (**A**) Superposition of the PL$^{pro}$-**Jun11313** structure to the structure of the PL$^{pro}$-**XR8-24** complex (PDB 7LBS), with **XR8-24** in yellow sticks and spheres, with the relevant residues for binding of both compounds indicated. Additionally, **XR8-24'** is displayed as light pink sticks and spheres, representing the docking of **XR8-24** with a phenyl-substituted moiety in the opposite direction to the experimental data, pointing towards the solvent. (**B**) Superposition of the PL$^{pro-}$ **Jun11313** structure to the structure of SARS-CoV-2 PL$^{pro}$ in complex with ISG15 (magenta ribbons, PDB 7RBS). The phenyl thienyl group of **Jun11313** is binding in the ISG15 binding site (analogous to the Ubiquitin binding site) of SARS-CoV-2 PL$^{pro}$ in the region where residues Asn151$^{ISG15}$ and Leu152$^{ISG15}$ at the end of a β sheet in ISG15 interact with PL$^{pro}$. The BL2 loop and BL2 groove elements of PL$^{pro}$ are highlighted alongside the catalytic residue Cys111.

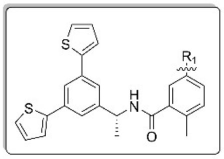

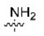

**Jun11924**
$IC_{50}$ = 177.7 ± 10.0 nM
$CC_{50}$ = 36.4 ± 1.3 μM

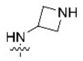

**Jun11922**
$IC_{50}$ = 171.2 ± 14.0 nM
$CC_{50}$ = 11.0 ± 0.4 μM

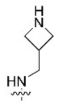

**Jun1238**
$IC_{50}$ = 339.8 ± 29.0 nM
$CC_{50}$ = 5.9 ± 0.7 μM

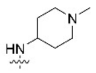

**Jun12112**
$IC_{50}$ = 184.1 ± 22.0 nM
$CC_{50}$ = 7.9 ± 1.4 μM

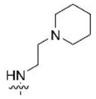

**Jun1271**
$IC_{50}$ = 116.6 ± 16.0 nM
$CC_{50}$ = 4.6 ± 0.4 μM

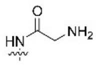

**Jun11923**
$IC_{50}$ = 147.2 ± 7.0 nM
$CC_{50}$ = 11.6 ± 0.7 μM

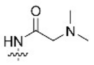

**Jun12111**
$IC_{50}$ = 118.2 ± 12.0 nM
$CC_{50}$ = 8.23 ± 0.26 μM

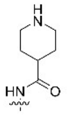

**Jun11945**
$IC_{50}$ = 162.8 ± 0.5 nM
$CC_{50}$ = 13.5 ± 0.7 μM

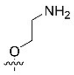

**Jun11874**
$IC_{50}$ = 65.1 ± 1.4 nM
$CC_{50}$ = 7.0 ± 1.1 μM

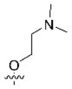

**Jun11875**
$IC_{50}$ = 66.2 ± 2.0 nM
$CC_{50}$ = 4.5 ± 1.6 μM

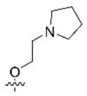

**Jun11899**
$IC_{50}$ = 75.9 ± 2.0 nM
$CC_{50}$ = 6.5 ± 0.3 μM

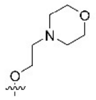

**Jun118910**
$IC_{50}$ = 367.1 ± 3.0 nM
$CC_{50}$ = 96.8 ± 3.6 μM

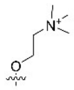

**Jun11898**
$IC_{50}$ = 132.6 ± 4.0 nM
$CC_{50}$ = 19.1 ± 4.2 μM

---

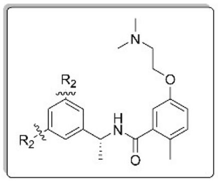

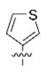

**Jun11931**
$IC_{50}$ = 99.3 ± 4.0 nM
$CC_{50}$ = 9.4 ± 1.0 μM

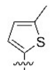

**Jun119415**
$IC_{50}$ = 143.2 ± 9.0 nM
$CC_{50}$ = 7.6 ± 0.3 μM

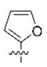

**Jun11903**
$IC_{50}$ = 202.2 ± 8.0 nM
$CC_{50}$ = 7.7 ± 0.4 μM

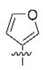

**Jun11932**
$IC_{50}$ = 83.7 ± 6.0 nM
$CC_{50}$ = 16.0 ± 1.2 μM

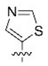

**Jun11934**
$IC_{50}$ = 109.0 ± 7.0 nM
$CC_{50}$ = 37.1 ± 4.4 μM
$T_{1/2}$ = 17.9 min

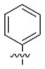

**Jun11933**
$IC_{50}$ = 103.0 ± 7.0 nM
$CC_{50}$ = 13.8 ± 1.9 μM

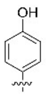

**Jun11943**
$IC_{50}$ = 167.3 ± 7.0 nM
$CC_{50}$ = 9.1 ± 0.3 μM

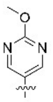

**Jun1218**
$IC_{50}$ = 1.95 ± 0.08 μM
$CC_{50}$ = 89.8 ± 3.0 μM

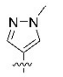

**Jun11941**
$IC_{50}$ = 151.1 ± 4.0 nM
$CC_{50}$ = 54.8 ± 8.3 μM
FlipGFP $EC_{50}$ = 1.8 ± 0.2 μM
$T_{1/2}$ > 145 min

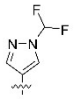

**Jun11942**
$IC_{50}$ = 178.5 ± 8.0 nM
$CC_{50}$ = 160.4 ± 59.2 μM
FlipGFP $EC_{50}$ = 2.2 ± 0.2 μM
$T_{1/2}$ = 45.5 min

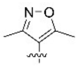

**Jun1212**
$IC_{50}$ = 390.3 ± 25.0 nM
$CC_{50}$ = 17.8 ± 0.3 μM

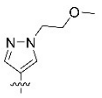

**Jun1215**
$IC_{50}$ = 181.7 ± 6.0 nM
$CC_{50}$ = 77.6 ± 5.5 μM
FlipGFP $EC_{50}$ = 4.7 ± 0.4 μM

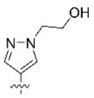

**Jun1247**
$IC_{50}$ = 165.3 ± 9.0 nM
$CC_{50}$ > 125 μM
FlipGFP $EC_{50}$ > 60 μM

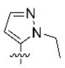

**Jun12382**
$IC_{50}$ = 443.4 ± 44.0 nM
$CC_{50}$ = 31.0 ± 2.1 μM

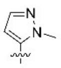

**Jun1246**
$IC_{50}$ = 463.6 ± 39.1 nM
$CC_{50}$ = 39.6 ± 4.7 μM

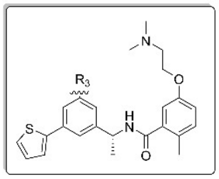

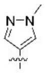

**Jun1211**
$IC_{50}$ = 152.7 ± 6.0 nM
$CC_{50}$ = 9.8 ± 1.0 μM

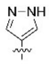

**Jun1244**
$IC_{50}$ = 81.9 ± 8.0 nM
$CC_{50}$ = 9.0 ± 0.7 μM
$T_{1/2}$ = 6.6 min

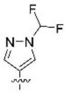

**Jun1251**
$IC_{50}$ = 74.7 ± 4.0 nM
$CC_{50}$ = 7.6 ± 0.4 μM
$T_{1/2}$ = 25 min

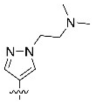

**Jun1252**
$IC_{50}$ = 116.4 ± 10.0 nM
$CC_{50}$ = 8.2 ± 0.2 μM
$T_{1/2}$ = 66.8 min

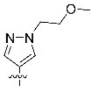

**Jun1254**
$IC_{50}$ = 111.8 ± 11.0 nM
$CC_{50}$ = 9.7 ± 2.2 μM

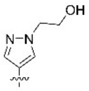

**Jun1257**
$IC_{50}$ = 113.5 ± 8.0 nM
$CC_{50}$ = 18.0 ± 1.6 μM
$T_{1/2}$ = 55.6 min

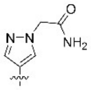

**Jun121210**
$IC_{50}$ = 81.9 ± 5.7 nM
$CC_{50}$ > 125 μM
FlipGFP $EC_{50}$ = 13.6 ± 1.9 μM

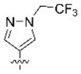

**Jun12149**
$IC_{50}$ = 73.8 ± 3.0 nM
$CC_{50}$ = 6.3 ± 1.1 μM

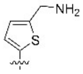

**Jun12129**
$IC_{50}$ = 90.9 ± 4.6 nM
$CC_{50}$ = 8.3 ± 1.2 μM

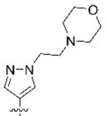

**Jun121910**
$IC_{50}$ = 66.4 ± 3.0 nM
$CC_{50}$ = 17.7 ± 2.1 μM
FlipGFP $EC_{50}$ = 1.1 ± 0.2 μM
$T_{1/2}$ = 28.5 min

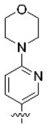

**Jun12511**
$IC_{50}$ = 91.5 ± 17.0 nM
$CC_{50}$ = 6.2 ± 0.2 μM
$T_{1/2}$ = 91.2 min

---

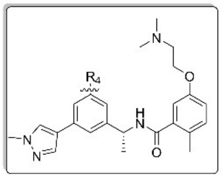

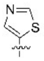

**Jun12145**
$IC_{50}$ = 108.5 ± 6.2 nM
$CC_{50}$ = 31.3 ± 5.4 μM
FlipGFP $EC_{50}$ = 1.3 ± 0.3 μM
$T_{1/2}$ = 26.3 min

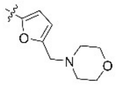

**Jun12681**
$IC_{50}$ = 120.2 ± 5.1 nM
$CC_{50}$ = 114.5 ± 3.4 μM
FlipGFP $EC_{50}$ = 1.8 ± 0.3 μM

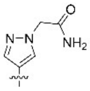

**Jun121911**
$IC_{50}$ = 73.2 ± 3.2 nM
$CC_{50}$ > 125 μM
FlipGFP $EC_{50}$ > 60 μM

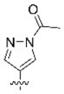

**Jun12336**
$IC_{50}$ = 118.2 ± 6.4 nM
$CC_{50}$ > 125 μM
$T_{1/2}$ < 2.5 min

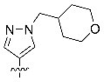

**Jun12467**
$IC_{50}$ = 73.26 ± 6 nM
$CC_{50}$ = 165.2 ± 26.5 μM
FlipGFP $EC_{50}$ = 1.7 ± 0.4 μM
$T_{1/2}$ = 139.1 min

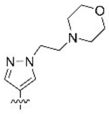

**Jun12351**
$IC_{50}$ = 98.8 ± 5.2 nM
$CC_{50}$ > 125 μM
FlipGFP $EC_{50}$ = 2.0 ± 0.4 μM
$T_{1/2}$ > 145 min

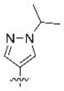

**Jun12199**
$IC_{50}$ = 108.8 ± 10.2 nM
$CC_{50}$ = 63.2 ± 11.3 μM
FlipGFP $EC_{50}$ = 0.8 ± 0.1 μM
$T_{1/2}$ = 79.7 min

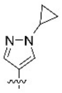

**Jun12197**
$IC_{50}$ = 102.7 ± 10.2 nM
$CC_{50}$ = 43.7 ± 5.1 μM
FlipGFP $EC_{50}$ = 0.6 ± 0.1 μM
$T_{1/2}$ = 116.0 min

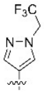

**Jun12603**
$IC_{50}$ = 112.2 ± 6.0 nM
$CC_{50}$ = 61.0 ± 19.3 μM
FlipGFP $EC_{50}$ = 2.4 ± 0.4 μM

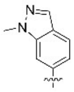

**Jun12446**
$IC_{50}$ = 132.5 ± 4.2 nM
$CC_{50}$ = 8.6 ± 0.5 μM

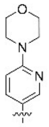

**Jun12278**
$IC_{50}$ = 88.1 ± 11.5 nM
$CC_{50}$ = 37.3 ± 3.8 μM
FlipGFP $EC_{50}$ = 1.0 ± 0.2 μM
$T_{1/2}$ = 71.3 min

---

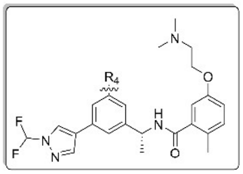

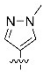

**Jun12162**
$IC_{50}$ = 98.3 ± 7.0 nM
$CC_{50}$ = 42.3 ± 2.3 μM
FlipGFP $EC_{50}$ = 2.1 ± 0.2 μM
$T_{1/2}$ = 76.7 min
SARS-CoV-2
$EC_{50}$ = 0.33 μM (A549)

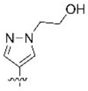

**Jun12208**
$IC_{50}$ = 91.6 ± 6.0 nM
$CC_{50}$ = 167.8 ± 15.8 μM
FlipGFP $EC_{50}$ = 16.9 ± 3.3 μM

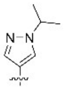

**Jun12235**
$IC_{50}$ = 162.2 ± 10.0 nM
$CC_{50}$ = 16.5 ± 6.8 μM

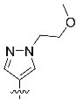

**Jun12281**
$IC_{50}$ = 111.9 ± 9.0 nM
$CC_{50}$ = 59.2 ± 10.0 μM
FlipGFP $EC_{50}$ = 1.8 ± 0.5 μM
$T_{1/2}$ = 75.6 min

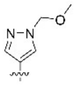

**Jun12303**
$IC_{50}$ = 121.5 ± 4.0 nM
$CC_{50}$ = 56.8 ± 7.4 μM
FlipGFP $EC_{50}$ = 2.6 ± 0.2 μM
$T_{1/2}$ = 136 min

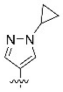

**Jun12284**
$IC_{50}$ = 96.6 ± 6.0 nM
$CC_{50}$ = 30.4 ± 9.5 μM
FlipGFP $EC_{50}$ = 2.1 ± 0.4 μM

---

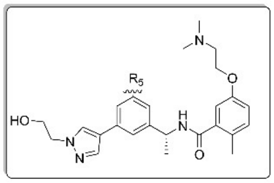

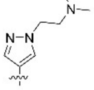

**Jun12165**
$IC_{50}$ = 145.0 ± 10.0 nM
$CC_{50}$ > 125 μM

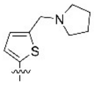

**Jun12168**
$IC_{50}$ = 72.4 ± 3.0 nM
$CC_{50}$ = 34.5 ± 10.2 μM
FlipGFP $EC_{50}$ = 9.7 ± 1.5 μM

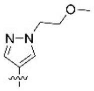

**Jun12163**
$IC_{50}$ = 102.0 ± 7.0 nM
$CC_{50}$ > 125 μM

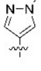

**Jun12164**
$IC_{50}$ = 92.4 ± 4.0 nM
$CC_{50}$ > 125 μM

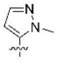

**Jun12198**
$IC_{50}$ = 142.0 ± 19.0 nM
$CC_{50}$ > 125 μM

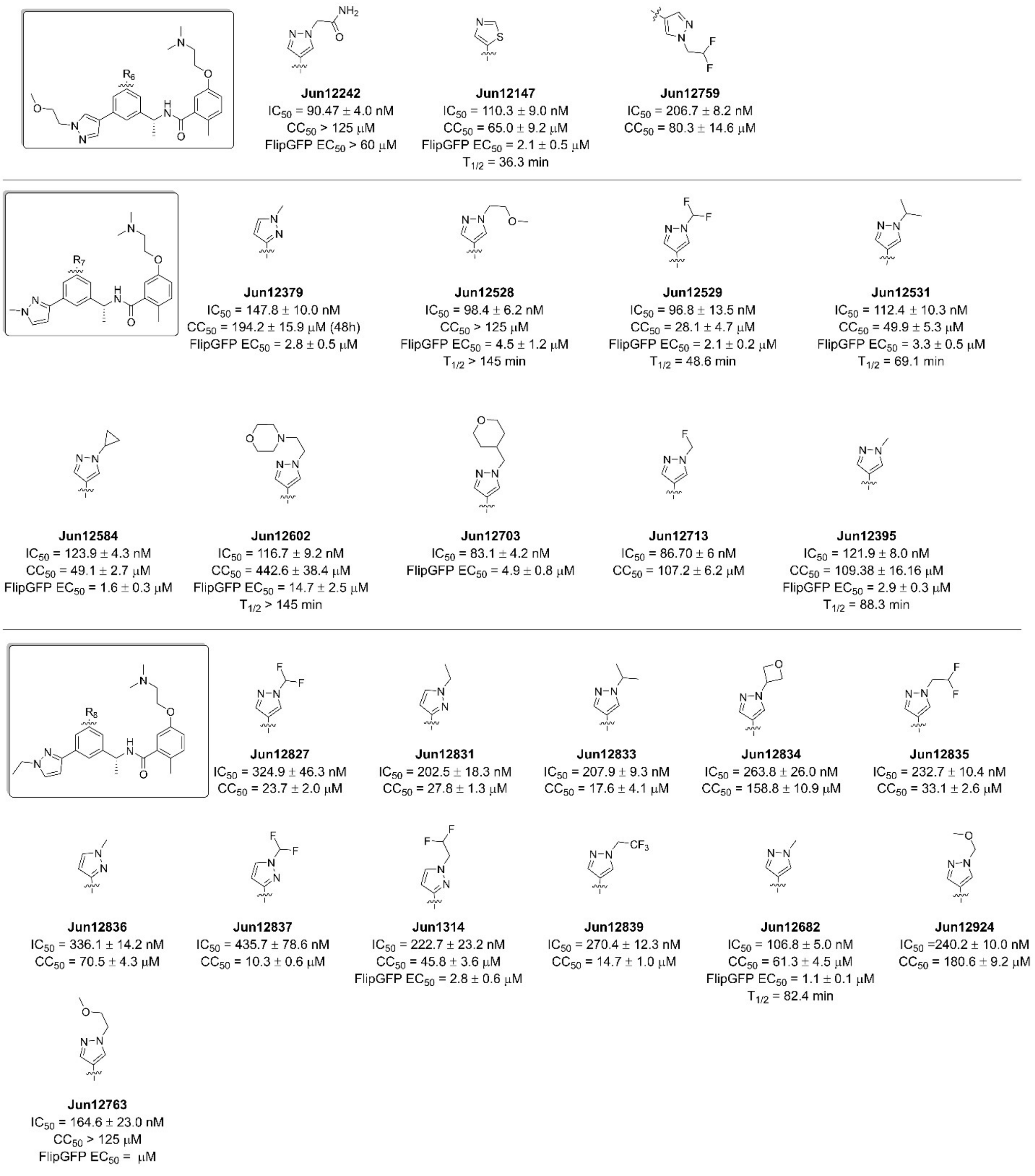


**Fig. S2. Lead optimization of biarylphenyl series of SARS-CoV-2 PL$^{pro}$ inhibitors.** $IC_{50}$ values were obtained in the FRET-based enzymatic assay as described in the materials and methods section. $CC_{50}$ values were determined using the neutral red method in Vero E6 cells with a 48 h incubation. FlipGFP $EC_{50}$ values were measured in 293T cells transfected with the pcDNA3-flipGFP-T2A-mCherry plasmid and PL$^{pro}$ expression plasmid pcDNA3.1. The results are mean ± standard deviation of three replicates.

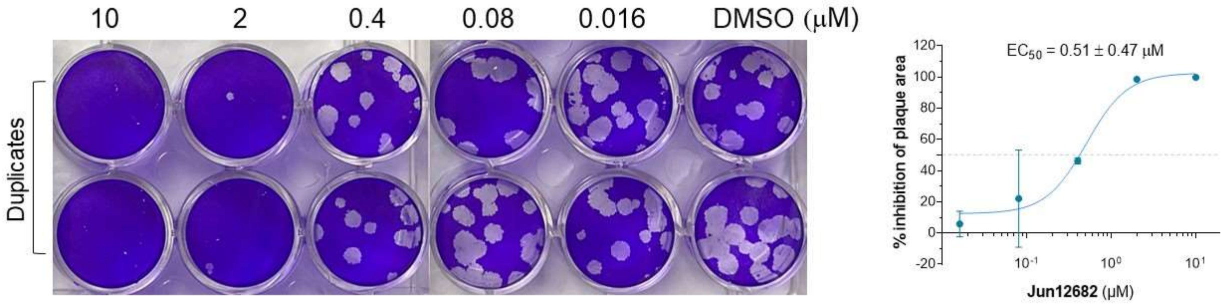


**Fig. S3. Plaque assay of Jun12682 against SARS-CoV-2.** Vero-AT cells were seeded in 12-well plates a day prior to infection and the drug dissolved in DMSO was serially diluted in DMEM with 3-fold dilutions between test concentrations.

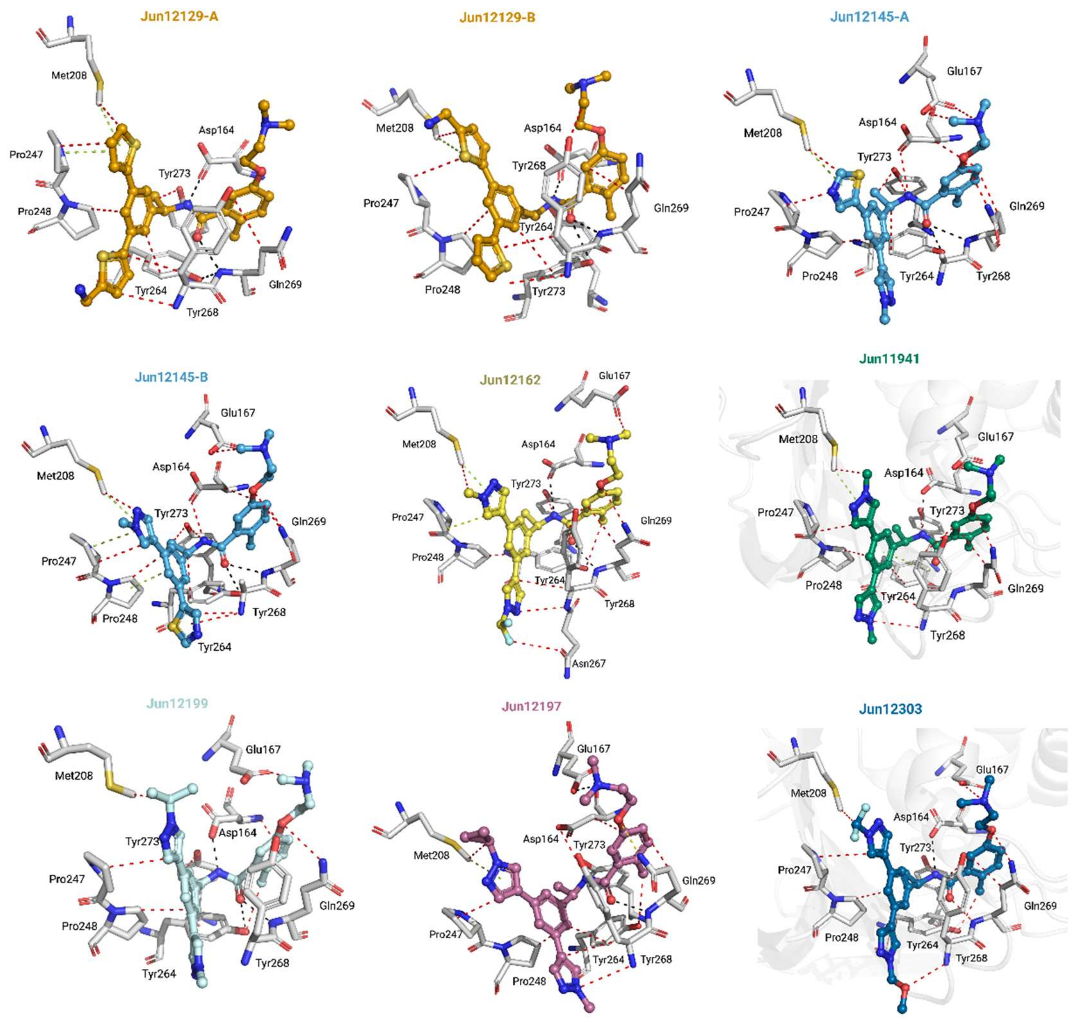


**Fig. S4. Binding of biarylphenyl benzamides to SARS-CoV-2 PL$^{pro}$.** Interactions of **Jun 12682** derivatives with the PL$^{pro}$ protein are almost conserved. Hydrogen bonds are represented in black dashed lines, van der Waals contacts as red dashed lines, and π–π interactions as light green dashed. PL$^{pro}$ complex structures of **Jun12129** and **Jun12145** possess two conformations of the inhibitor, designated by A and B conformers. The occupancies for **Jun 12129** conformations A and B are 0.64 and 0.36, and **Jun12145** are 0.67 and 0.33.

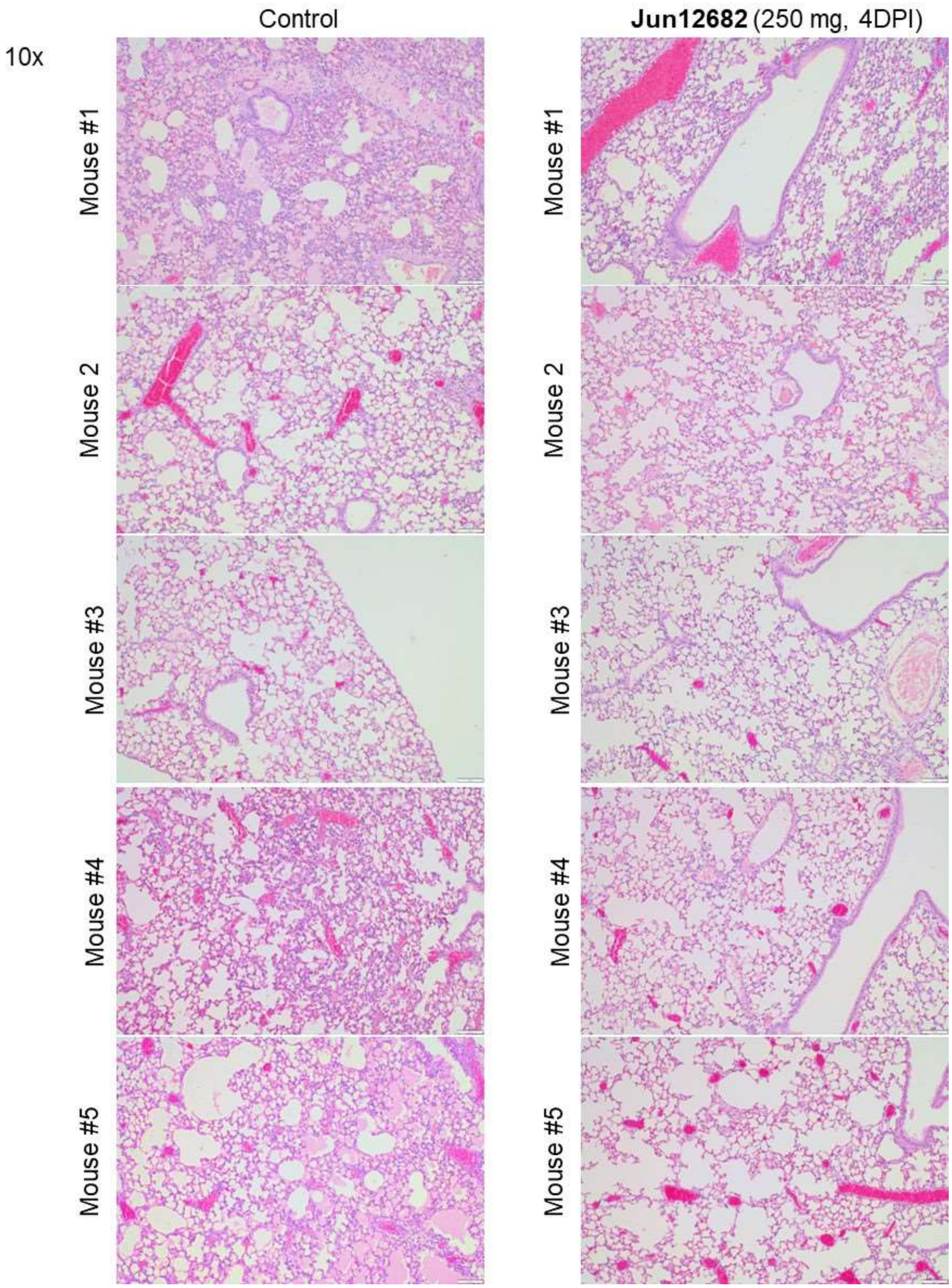


**Fig. S5. H&E staining of individual mouse lung sections.** Lungs collected at 4 DPI from vehicle- or 250mg/kg **Jun12682**-treated mice (n=5 each group) were stained with haematoxylin and eosin (H&E). Scale bars, 50 µm (10X).

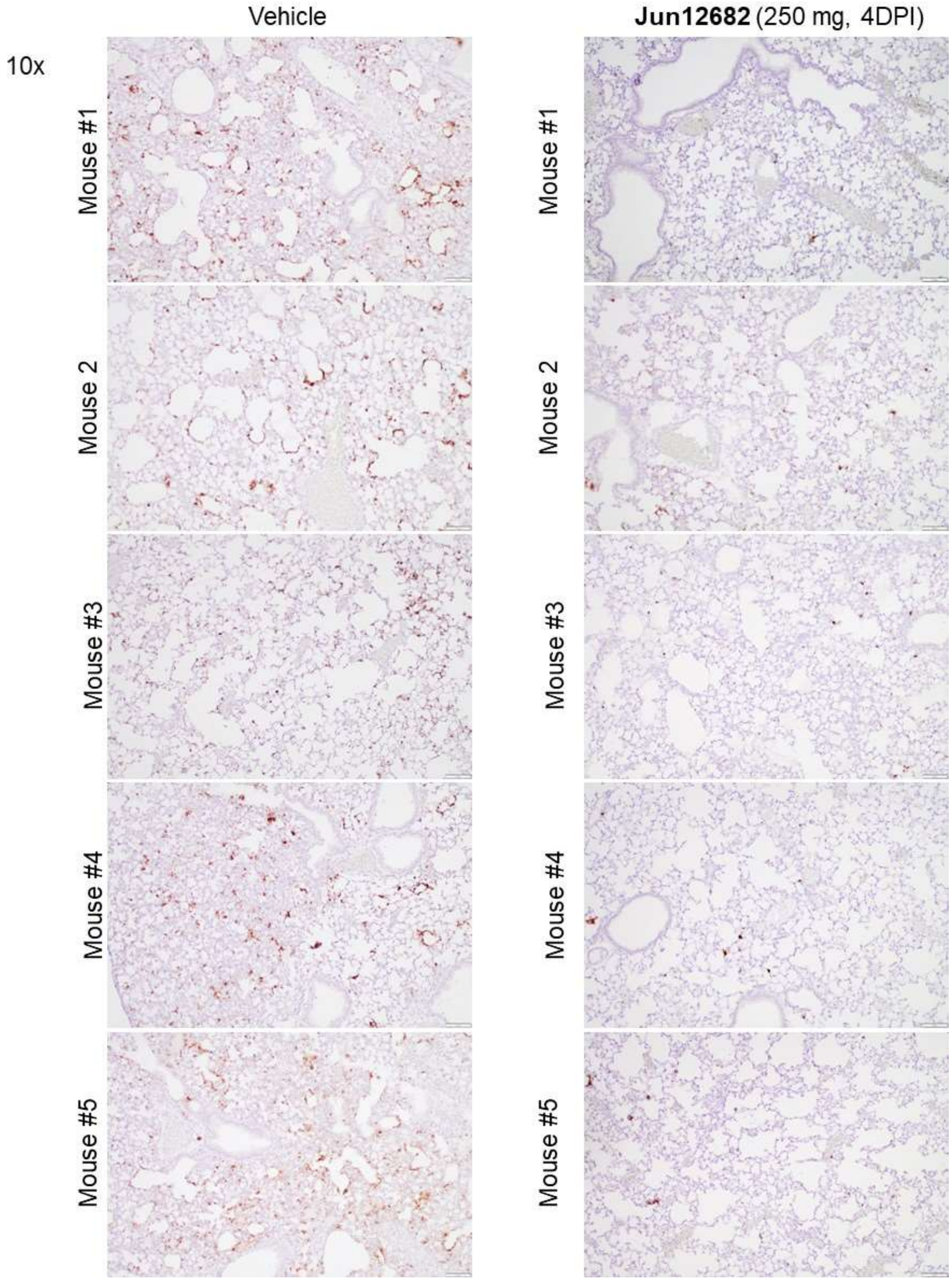


**Fig. S6. Viral antigen staining of individual mouse lung sections.** Lungs collected at 4 DPI from vehicle- or 250mg/kg **Jun12682**-treated mice (n=5 each group) were stained with a monoclonal antibody against SARS-CoV-2 nucleocapsid protein. Scale bars, 50 µm (10X).

#### Table S1. X-ray data collection and refinement statistics

| PDB Accession code | 8UUW | 8UUY | 8UUV | 8UUU |
|---|---|---|---|---|
| **Inhibitor** | **Jun12145** | **Jun12129** | **Jun12197** | **Jun12162** |
| **Data Statistics** | | | | |
| **Wavelength** | 1.033 | 1.033 | 1.033 | 1.033 |
| **Resolution range** | 109.18 - 3.20 (3.42-3.20) | 100.92 - 3.05 (3.26 - 3.051) | 29.05 - 3.01 (3.19 - 3.01) | 29.35 - 3.01 (3.19 - 3.01) |
| **Space group** | I $4_1$ 2 2 | I $4_1$ 2 2 | I $4_1$ 2 2 | I $4_1$ 2 2 |
| **Unit cell dimensions a b c [Å]** | 115.028 115.028 218.357 | 113.68 113.68 219.955 | 114.212 114.212 218.533 | 115.393 115.393 220.691 |
| **Unit cell dimensions α β γ (°)** | 90 90 90 | 90 90 90 | 90 90 90 | 90 90 90 |
| **Unique reflections** | 12514 (2233) | 14155 (2508) | 14671 (2235) | 15235 (2404) |
| **Multiplicity** | 9.6 (8.3) | 26.4 (27.2) | 25.5 (24.6) | 25.6 (25.4) |
| **Completeness (%)** | 99.9 (99.80) | 100.0 (100) | 99.3 (96.3) | 99.7 (99.1) |
| **Mean I/sigma(I)** | 4.5 (1.5) | 6.8 (1.0) | 19.1 (4.60) | 17.03 (5.1) |
| **Wilson B-factor** | 68.26 | 86.1 | 74.45 | 65.43 |
| **R-merge** | 0.620 (3.791) | 0.508(5.381) | 0.148 (0.838) | 0.189 (0.813) |
| **R-meas** | 0.656 (4.055) | 0.518(5.482) | 0.151 (0.856) | 0.192 (0.830) |
| **R-pim** | 0.209 (1.403) | 0.100(1.037) | 0.030 (0.170) | 0.038 (0.162) |
| **CC1/2** | 0.931 (0.434) | 0.997(0.677) | 0.999 (0.975) | 0.998 (0.954) |
| **Refinement Statistics** | | | | |
| **Reflections used in refinement** | 12339 (1183) | 14106 (1379) | 14644 (1357) | 15190 (1506) |
| **Reflections used for R-free** | 613 (58) | 704 (65) | 715 (49) | 755 (57) |
| **R-work** | 0.2745 (0.3451) | 0.2400 (0.3671) | 0.2144 (0.3257) | 0.2141 (0.2900) |
| **R-free** | 0.3149 (0.4997) | 0.2575 (0.3731) | 0.2426 (0.3734) | 0.2386 (0.3405) |
| **Number of non-hydrogen atoms** | 2503 | 2663 | 2543 | 2522 |
| **macromolecules** | 2330 | 2469 | 2431 | 2390 |
| **ligands** | 85 | 101 | 45 | 62 |
| **solvent** | 88 | 93 | 67 | 70 |
| **Protein residues** | 296 | 312 | 307 | 305 |
| **RMS(bonds)** | 0.005 | 0.002 | 0.002 | 0.004 |
| **RMS(angles)** | 0.89 | 0.42 | 0.44 | 0.64 |
| **Ramachandran favored (%)** | 96.85 | 95.81 | 97.36 | 95.02 |
| **Ramachandran allowed (%)** | 3.15 | 4.19 | 2.64 | 4.98 |
| **Ramachandran outliers (%)** | 0 | 0 | 0 | 0 |
| **Rotamer outliers (%)** | 0.4 | 0.37 | 0 | 0 |
| **Clashscore** | 9.9 | 3.8 | 6.01 | 4.42 |
| **Average B-factor** | 76.49 | 90.92 | 78.36 | 66.85 |
| **macromolecules** | 77.35 | 91.36 | 78.79 | 66.98 |
| **ligands** | 68.34 | 90.09 | 69.85 | 69.85 |
| **solvent** | 61.63 | 80.18 | 59.79 | 64.06 |

| PDB Accession code | 8UUH | 8UUG | 8UUF | 8UOB | 8UVM |
|---|---|---|---|---|---|
| **Inhibitor** | **Jun12199** | **Jun12303** | **Jun11941** | **Jun12682** | **Jun11313** |
| **Data Statistics** | | | | | |
| **Wavelength** | 0.9201 | 0.9201 | 0.9201 | 0.9201 | 1.033 |
| **Resolution range** | 101.59 - 2.8 (2.95 - 2.8) | 29.05 - 2.743 (2.89 - 2.743) | 29.08 - 2.84 (2.99 - 2.84) | 34.74 - 2.52 (2.62 - 2.52) | 42.00 - 2.85 (2.952 - 2.85) |
| **Space group** | $I\,4_1\,2\,2$ | $I\,4_1\,2\,2$ | $I\,4_1\,2\,2$ | $I\,4_1\,2\,2$ | P 3 2 1 |
| **Unit cell dimensions a b c [Å]** | 114.713 114.713 218.719 | 114.2 114.2 218.557 | 114.285 114.285 219.48 | 115.825 115.825 219.317 | 190.997 190.997 112.926 |
| **Unit cell dimensions α β γ (°)** | 90 90 90 | 90 90 90 | 90 90 90 | 90 90 90 | 90 90 120 |
| **Unique reflections** | 18424 (2643) | 19362 (2774) | 17605 (2480) | 25626 (2834) | 54651 (2264) |
| **Multiplicity** | 26.7 (27.3) | 13.5 (13.8) | 13.3 (13.6) | 15.2 (15.4) | 9.8 (4.8) |
| **Completeness (%)** | 100 (100) | 99.9 (99.6) | 99.76 (98.2) | 100 (100) | 99.8 (96.7) |
| **Mean I/sigma(I)** | 9.51 (1.2) | 17.5 (3.2) | 17.65 (3.8) | 7.61 (0.72) | 3.2 (0.4) |
| **Wilson B-factor** | 64.65 | 60.06 | 63.01 | 57.81 | 59.43 |
| **R-merge** | 0.346 (3.758) | 0.118 (0.883) | 0.115 (0.746) | 0.304 (4.946) | 0.187(0.944) |
| **R-meas** | 0.352 (3.829) | 0.123 (0.947) | 0.119 (0.775) | 0.315 (5.11) | 0.193(0.998) |
| **R-pim** | 0.068 (0.730) | 0.03315 (0.252) | 0.033 (0.207) | 0.081 (1.300) | 0.043 (0.309) |
| **CC1/2** | 0.997 (0.581) | 0.999 (0.876) | 0.999 (0.905) | 0.995 (0.400) | 0.996 (0.472) |
| **Refinement Statistics** | | | | | |
| **Reflections used in refinement** | 18341 (1796) | 19338 (1879) | 17548 (1708) | 25575 (2490) | 54651 (2264) |
| **Reflections used for R-free** | 951 (90) | 939 (102) | 848 (86) | 1308 (110) | 2793 (129) |
| **R-work** | 0.2282 (0.3309) | 0.2122 (0.2733) | 0.2118 (0.2894) | 0.2063 (0.3694) | 0.2381 (0.3246) |
| **R-free** | 0.2640 (0.3806) | 0.2362 (0.2704) | 0.2387 (0.3824) | 0.2442 (0.4062) | 0.2576 (0.3724) |
| **Number of non-hydrogen atoms** | 2553 | 2580 | 2598 | 2699 | 10153 |
| **macromolecules** | 2427 | 2435 | 2452 | 2460 | 9791 |
| **ligands** | 64 | 56 | 47 | 52 | 177 |
| **solvent** | 62 | 89 | 99 | 187 | 185 |
| **Protein residues** | 307 | 311 | 312 | 311 | 1258 |
| **RMS(bonds)** | 0.002 | 0.002 | 0.002 | 0.003 | 0.002 |
| **RMS(angles)** | 0.46 | 0.52 | 0.47 | 0.58 | 0.43 |
| **Ramachandran favored (%)** | 95.68 | 95.47 | 97.1 | 97.09 | 95.03 |
| **Ramachandran allowed (%)** | 4.32 | 4.21 | 2.9 | 2.91 | 4.97 |
| **Ramachandran outliers (%)** | 0 | 0.32 | 0 | 0 | 0 |
| **Rotamer outliers (%)** | 0.77 | 0.38 | 0.75 | 0 | 0.19 |
| **Clashscore** | 3.1 | 4.54 | 3.71 | 4.08 | 4.5 |
| **Average B-factor** | 69.62 | 67.1 | 69.94 | 77.09 | 65.23 |
| **macromolecules** | 69.69 | 67.17 | 70.38 | 77.43 | 65.35 |
| **ligands** | 72.24 | 71.43 | 63.77 | 70.68 | 75.85 |
| **solvent** | 62.45 | 62.13 | 66.62 | 74.39 | 48.55 |

Statistics for the highest-resolution shell are shown in parentheses.

Rmerge=ΣΣ|$ni$=1$hkl$ Ii – $I$|/ΣΣ$Ini$=1$hkl$i

c Rpim= Σ√1$n$−1$hkl$ Σ|$Ini$=1i - $I(hkl)$ΣΣ$Ini$=1$hkl$i

#### Table S2. *In vivo* oral snap PK of PL$^{pro}$ inhibitors in C57BL/6J mice

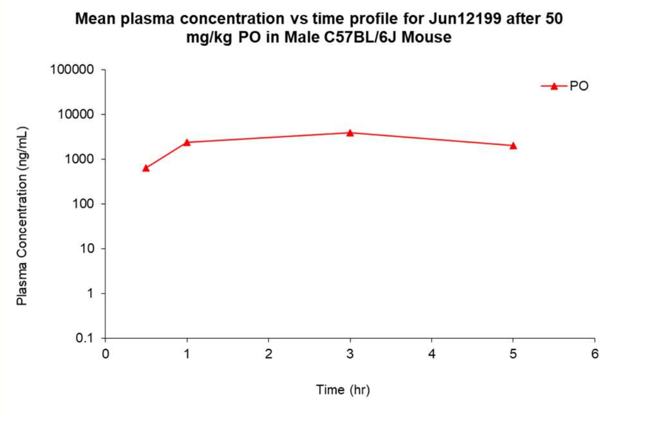


| PK parameters | Unit | Mean |
|---|---|---|
| $T_{1/2}$ | h | NA |
| $T_{max}$ | h | 3.00 |
| $C_{max}$ | ng/mL | 3980 |
| $AUC_{last}$ | h*ng/mL | 13355 |
| $AUC_{Inf}$ | h*ng/mL | NA |
| $AUC\_{_{\%Extrap}}\_obs$ | % | NA |
| $MRT_{Inf}\_obs$ | h | NA |
| $AUC_{last}/D$ | h*mg/mL | 267 |
| $AUC_{inf}/D$ | h*mg/mL | NA |
| F | % | NA |

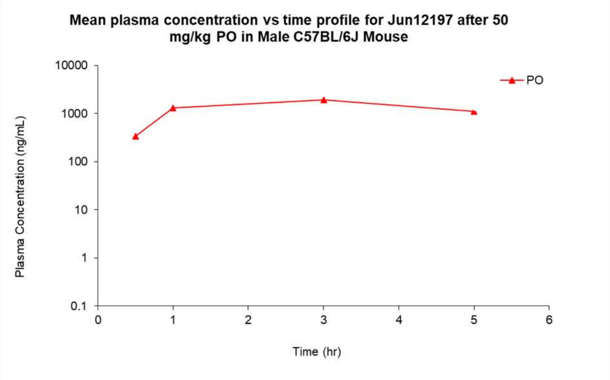


| PK parameters | Unit | Mean |
|---|---|---|
| $T_{1/2}$ | h | NA |
| $T_{max}$ | h | 3.00 |
| $C_{max}$ | ng/mL | 1930 |
| $AUC_{last}$ | h*ng/mL | 6787 |
| $AUC_{Inf}$ | h*ng/mL | NA |
| $AUC\_{_{\%Extrap}}\_obs$ | % | NA |
| $MRT_{Inf}\_obs$ | h | NA |
| $AUC_{last}/D$ | h*mg/mL | 136 |
| $AUC_{inf}/D$ | h*mg/mL | NA |
| F | % | NA |

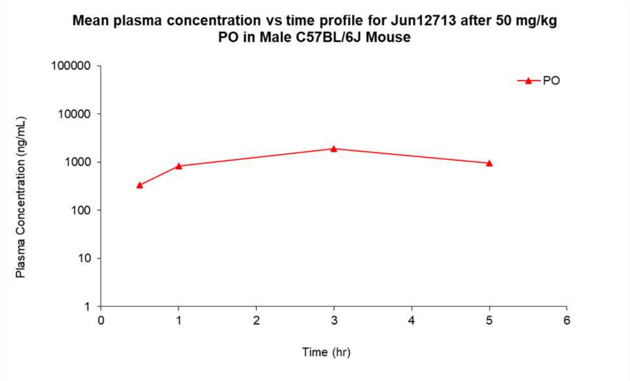


| PK parameters | Unit | Mean |
|---|---|---|
| $T_{1/2}$ | h | NA |
| $T_{max}$ | h | 3.00 |
| $C_{max}$ | ng/mL | 1890 |
| $AUC_{last}$ | h*ng/mL | 5940 |
| $AUC_{Inf}$ | h*ng/mL | NA |
| $AUC\_{_{\%Extrap}}\_obs$ | % | NA |
| $MRT_{Inf}\_obs$ | h | NA |
| $AUC_{last}/D$ | h*mg/mL | 119 |
| $AUC_{inf}/D$ | h*mg/mL | NA |
| F | % | NA |

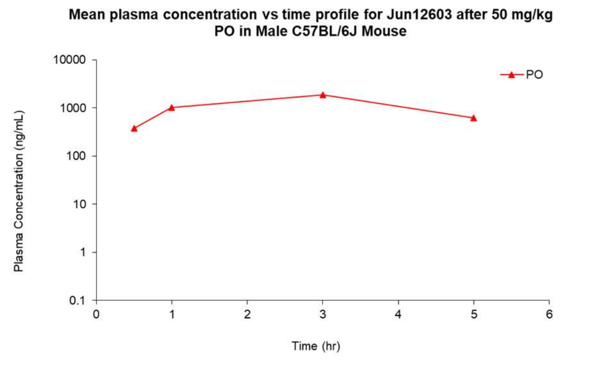


| PK parameters | Unit | Mean |
|---|---|---|
| $T_{1/2}$ | h | NA |
| $T_{max}$ | h | 3.00 |
| $C_{max}$ | ng/mL | 1910 |
| $AUC_{last}$ | h*ng/mL | 5935 |
| $AUC_{Inf}$ | h*ng/mL | NA |
| $AUC_{}_{\%Extrap}$_obs | % | NA |
| $MRT_{Inf}$_obs | h | NA |
| $AUC_{last}$/D | h*mg/mL | 119 |
| $AUC_{inf}$/D | h*mg/mL | NA |
| F | % | NA |

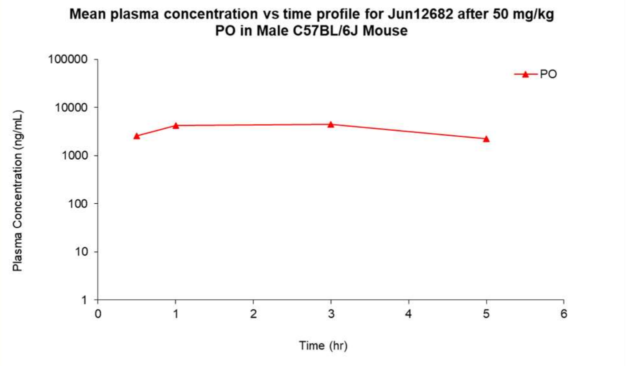


| PK parameters | Unit | Mean |
|---|---|---|
| $T_{1/2}$ | h | NA |
| $T_{max}$ | h | 3.00 |
| $C_{max}$ | ng/mL | 4520 |
| $AUC_{last}$ | h*ng/mL | 17895 |
| $AUC_{Inf}$ | h*ng/mL | NA |
| $AUC_{}_{\%Extrap}$_obs | % | NA |
| $MRT_{Inf}$_obs | h | NA |
| $AUC_{last}$/D | h*mg/mL | 358 |
| $AUC_{inf}$/D | h*mg/mL | NA |
| F | % | NA |

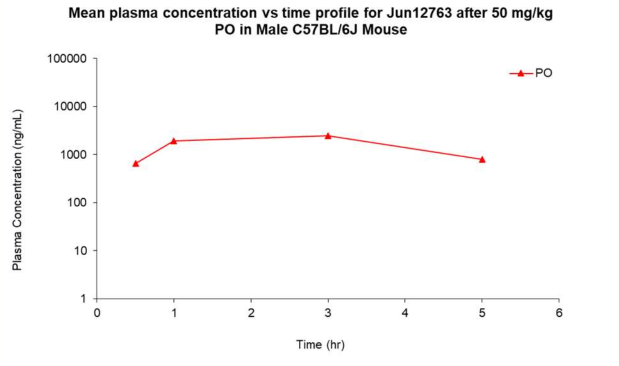


| PK parameters | Unit | Mean |
|---|---|---|
| $T_{1/2}$ | h | NA |
| $T_{max}$ | h | 3.00 |
| $C_{max}$ | ng/mL | 2480 |
| $AUC_{last}$ | h*ng/mL | 8541 |
| $AUC_{Inf}$ | h*ng/mL | NA |
| $AUC_{}_{\%Extrap}$_obs | % | NA |
| $MRT_{Inf}$_obs | h | NA |
| $AUC_{last}$/D | h*mg/mL | 171 |
| $AUC_{inf}$/D | h*mg/mL | NA |
| F | % | NA |

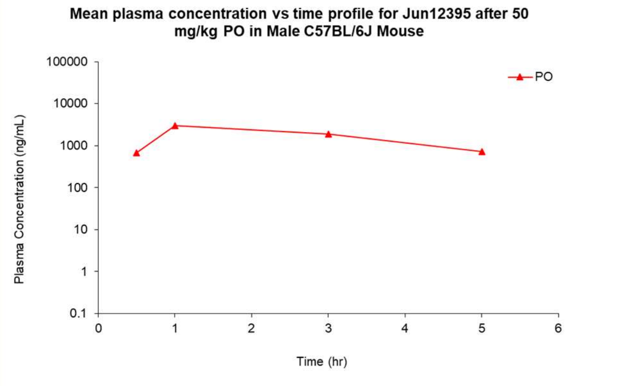


| PK parameters | Unit | Mean |
|---|---|---|
| $T_{1/2}$ | h | 1.97 |
| $T_{max}$ | h | 1.00 |
| $C_{max}$ | ng/mL | 3000 |
| $AUC_{last}$ | h*ng/mL | 8637 |
| $AUC_{Inf}$ | h*ng/mL | 10722 |
| $AUC_{}_{\%Extrap}$_obs | % | 19.4 |
| $MRT_{Inf}$_obs | h | 3.30 |
| $AUC_{last}$/D | h*mg/mL | 173 |
| $AUC_{inf}$/D | h*mg/mL | 214 |
| F | % | NA |

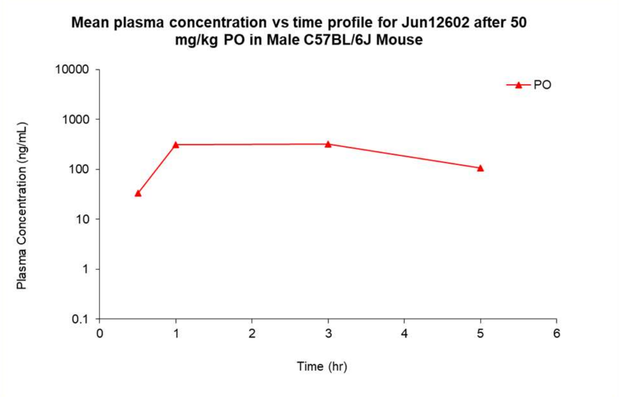


| PK parameters | Unit | Mean |
|---|---|---|
| $T_{1/2}$ | h | NA |
| $T_{max}$ | h | 3.00 |
| $C_{max}$ | ng/mL | 323 |
| $AUC_{last}$ | h*ng/mL | 1164 |
| $AUC_{Inf}$ | h*ng/mL | NA |
| AUC_$_{\%Extrap}$_obs | % | NA |
| $MRT_{Inf}$_obs | h | NA |
| $AUC_{last}$/D | h*mg/mL | 23.3 |
| $AUC_{inf}$/D | h*mg/mL | NA |
| F | % | NA |

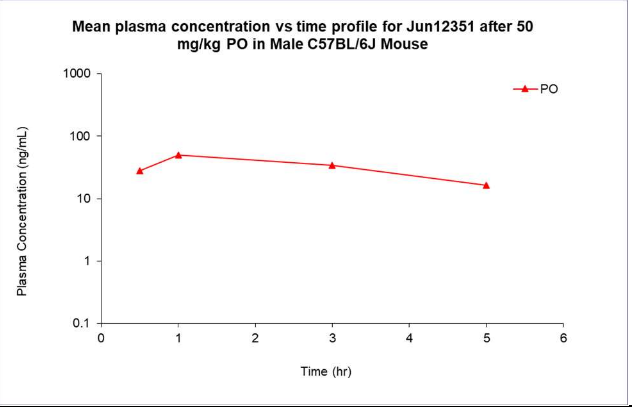


| PK parameters | Unit | Mean |
|---|---|---|
| $T_{1/2}$ | h | 2.50 |
| $T_{max}$ | h | 1.00 |
| $C_{max}$ | ng/mL | 49.7 |
| $AUC_{last}$ | h*ng/mL | 162 |
| $AUC_{Inf}$ | h*ng/mL | 221 |
| AUC_$_{\%Extrap}$_obs | % | 26.8 |
| $MRT_{Inf}$_obs | h | 3.93 |
| $AUC_{last}$/D | h*mg/mL | 3.24 |
| $AUC_{inf}$/D | h*mg/mL | 4.42 |
| F | % | NA |

**Table S3. *In vivo* oral PK of $PL^{pro}$ inhibitor Jun12682 in C57BL/6J mice.**

**Summary of Jun12682 IV plasma PK parameters**

| PK parameters | Unit | Mouse 1 | Mouse 2 | Mouse 3 | Mean | SD | CV(%) |
|---|---|---|---|---|---|---|---|
| Cl_obs | mL/min/kg | 28.5 | 28.3 | 23.7 | 26.8 | 2.7 | 10.1 |
| $T_{1/2}$ | h | 2.36 | 2.71 | 3.22 | 2.76 | 0.43 | 15.7 |
| $C_0$ | ng/mL | 5890 | 6957 | 5898 | 6249 | 614 | 9.82 |
| $AUC_{last}$ | h*ng/mL | 5840 | 5889 | 7011 | 6247 | 662 | 10.6 |
| $AUC_{Inf}$ | h*ng/mL | 5843 | 5891 | 7028 | 6254 | 671 | 10.7 |
| $AUC_{_\%Extrap_}$obs | % | 0.0416 | 0.0395 | 0.245 | 0.109 | 0.118 | 109 |
| $MRT_{Inf}$_obs | h | 1.64 | 1.32 | 2.31 | 1.76 | 0.51 | 28.7 |
| $AUC_{last}$/D | h*mg/mL | 584 | 589 | 701 | 625 | 66 | 10.6 |
| $AUC_{inf}$/D | h*mg/mL | 584 | 589 | 703 | 625 | 67 | 10.7 |
| $V_{ss}$_obs | L/kg | 2.81 | 2.24 | 3.29 | 2.78 | 0.52 | 18.9 |

**Summary of Jun12682 PO plasma PK parameters**

| PK parameters | Unit | Mouse 4 | Mouse 5 | Mouse 6 | Mean | SD | CV(%) |
|---|---|---|---|---|---|---|---|
| $T_{1/2}$ | h | 2.17 | 1.80 | 2.04 | 2.01 | 0.19 | 9.38 |
| $T_{max}$ | h | 2.00 | 2.00 | 1.00 | 1.67 | 0.58 | 34.6 |
| $C_{max}$ | ng/mL | 5220 | 3100 | 5290 | 4537 | 1245 | 27.4 |
| $AUC_{last}$ | h*ng/mL | 21521 | 18152 | 28590 | 22755 | 5327 | 23.4 |
| $AUC_{Inf}$ | h*ng/mL | 21532 | 18155 | 28600 | 22762 | 5330 | 23.4 |
| $AUC_{_\%Extrap_}$obs | % | 0.0517 | 0.0152 | 0.0339 | 0.0336 | 0.0183 | 54.4 |
| $MRT_{Inf}$_obs | h | 4.15 | 4.11 | 4.05 | 4.10 | 0.05 | 1.17 |
| $AUC_{last}$/D | h*mg/mL | 430 | 363 | 572 | 455 | 107 | 23.4 |
| $AUC_{inf}$/D | h*mg/mL | 431 | 363 | 572 | 455 | 107 | 23.4 |
| F | % | 68.9 | 58.1 | 91.5 | 72.8 | 17.0 | 23.4 |

**General Information of Chemical Synthesis**

All chemicals were purchased from commercial vendors and used without further purification unless otherwise noted. $^1H$ and $^{13}C$ NMR spectra were recorded on a Bruker-400 or -500 NMR spectrometer. Chemical shifts are reported in parts per million referenced with respect to residual solvent ($CD_3OD$) 3.31 ppm, (DMSO-$d_6$) 2.50 ppm, and ($CDCl_3$) 7.26 ppm or from internal standard tetramethylsilane (TMS) 0.00 ppm. The following abbreviations were used in reporting spectra: s, singlet; d, doublet; t, triplet; q, quartet; m, multiplet; dd, doublet of doublets; ddd, doublet of doublet of doublets. All reactions were carried out under Ar atmosphere, unless otherwise stated. HPLC-grade solvents were used for all reactions. Flash column chromatography was performed using silica gel (230-400 mesh, Merck). Low-resolution mass spectra were obtained using an ESI technique on a 3200 Q Trap LC/MS/MS system (Applied Biosystems). The purity was assessed by using Shimadzu LC-MS with Waters XTerra MS C-18 column (part #186000538), 50 × 2.1 mm, at a flow rate of 0.3 mL/min; $\lambda$ = 250 and 220 nm; mobile phase A, 0.1% formic acid in $H_2O$, and mobile phase B', 0.1% formic in 60% isopropanol, 30% $CH_3CN$ and 9.9% H2O. All compounds submitted for testing were confirmed to be > 95.0% purity by LC-MS traces. All compounds were characterized by proton and carbon NMR and HRMS.

#### Synthesis of PL$^{pro}$ inhibitors

***Scheme 1. Synthesis of SARS-CoV-2 PL$^{pro}$ inhibitors.***

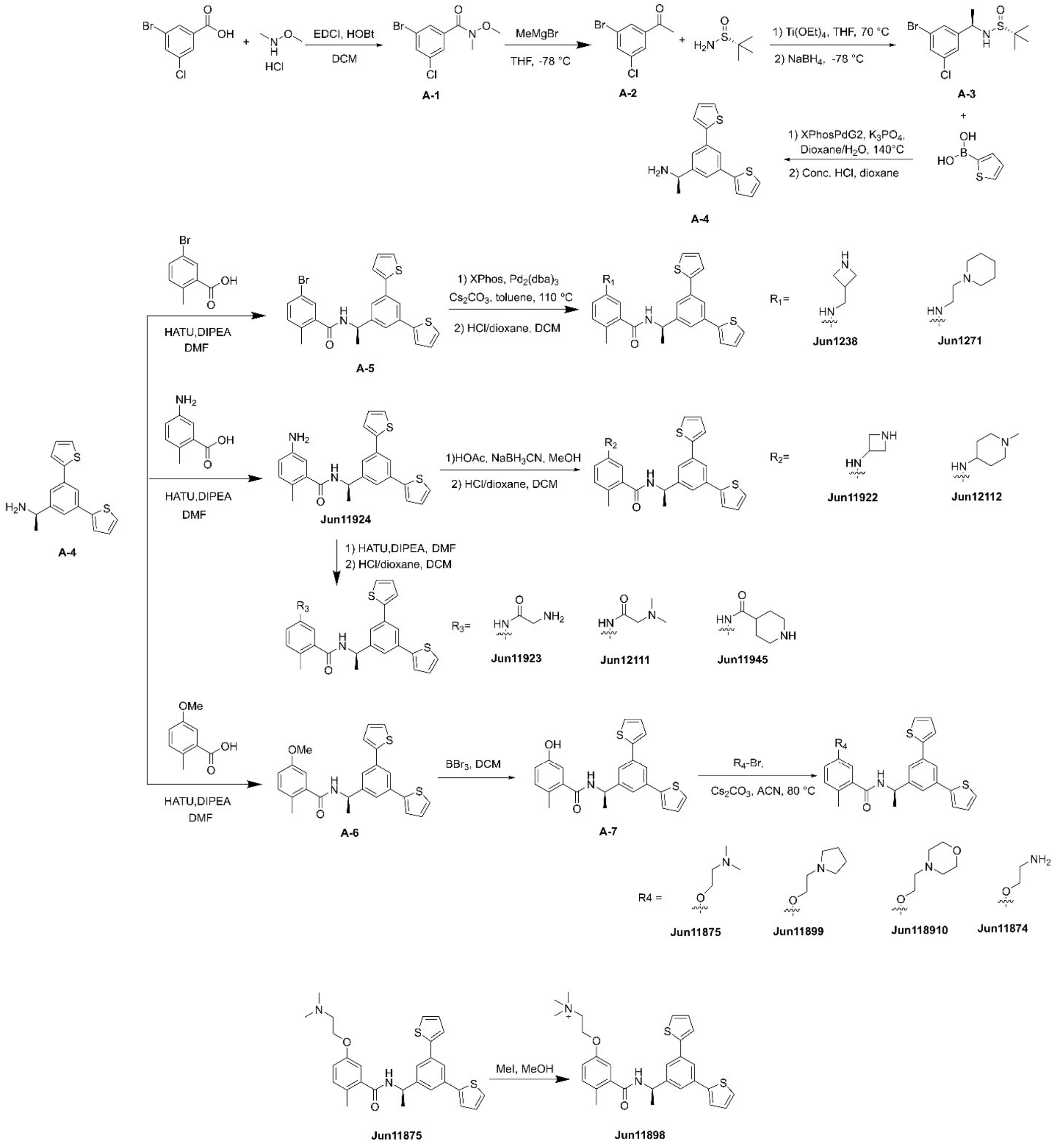

***Synthetic Procedures for Scheme 1***

***3-bromo-5-chloro-N-methoxy-N-methylbenzamide*** (**A-1**). To a solution of 3-bromo-5-chlorobenzoic acid (1.0 equiv), *N*,*O*-dimethylhydroxylamine hydrochloride (1.2 equiv), HOBt (1.3 equiv) and EDCI (1.3 equiv) in DCM (0.1 M) was added dropwise of triethylamine (3 equiv) at 0 °C. The mixture was stirred at rt for 24 hours. After the reaction was completed, the organic layer was washed with water and brine, dried over $Na_2SO_4$, filtered, and concentrated under vacuum to give the product **A-1** for the next step without purification.

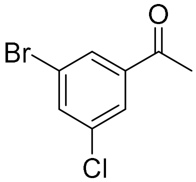


***1-(3-bromo-5-chlorophenyl)ethan-1-one*** (**A-2**). To a solution of 3-bromo-5-chloro-*N*-methoxy-*N*-methylbenzamide (**A-1**, 1.0 equiv) in dry ether at -78 °C under $N_2$, the methylmagnesium bromide (3 M solution in diethyl ether, 3.0 equiv) was added slowly. After warming to room temperature and stirred for 2 hours, the reaction mixture was quenched with ice-cold saturated $NH_4Cl$ and washed with water and brine, dried over $Na_2SO_4$, filtered, and concentrated under vacuum to give the crude residue, which was purified by silica gel column chromatography (Hexanes/EtOAc = 5:1) to provide the desired product **A-2** as a white solid. ***1-(3-bromo-5-chlorophenyl)ethan-1-one*** (**A-2**). White solid, 75% yield for two steps. $^1$H NMR (400 MHz, $CDCl_3$) δ 7.95 (d, J = 1.7 Hz, 1H), 7.85 (s, 1H), 7.70 (s, 1H), 2.59 (s, 3H). $^{13}$C NMR (101 MHz, $CDCl_3$) δ 195.22, 139.59, 135.79, 135.54, 129.68, 127.21, 123.26, 26.62. $C_8H_6BrClO$, MS calculated for m/z $[M+H]^+$: 232.9 (calculated), 233.1 (found).

***(S)-N-((R)-1-(3-bromo-5-chlorophenyl)ethyl)-2-methylpropane-2-sulfinamide*** (**A-3**). To a solution of $Ti(OEt)_4$ (1.44 equiv) and 1-(3-bromo-5-chlorophenyl)ethan-1-one (1.00 equiv) in THF (0.1 M) under an Ar atmosphere was added (*R*)-2-methylpropane-2-sulfinamide (1.50 equiv) and the mixture was heated to 70 °C for overnight. Upon completion, as determined by TLC, the mixture was cooled to -78 °C, and $NaBH_4$ (3.00 equiv) was added carefully. The mixture was stirred at -78 °C for 3 h and then slowly warmed to room temperature. After another 3 h, MeOH was added dropwise until the gas was no longer evolved. The resulting suspension was filtered through a plug of Celite, and the filter cake was washed with EtOAc. The filtrate was washed with brine, and the brine layer was extracted with EtOAc. The combined organic phase was dried over $Na_2SO_4$, filtered, and concentrated under vacuum to give a residue that was purified by flash silica gel column chromatography (Hexanes/EtOAc = 3:1) to get the desired product **A-3** as a light yellow solid.

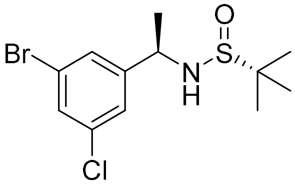


***1-(3-bromo-5-chlorophenyl)ethan-1-one*** (**A-3**). White solid, 80% yield. $^1$H NMR (400 MHz, $CDCl_3$) δ 7.44 (t, J = 1.9 Hz, 1H), 7.38 (d, J = 1.6 Hz, 1H), 7.27 (d, J = 1.8 Hz, 1H), 4.47 (m,

1H), 3.46 (s, 1H), 1.50 (d, J = 6.6 Hz, 3H), 1.24 (s, 9H). $^{13}$C NMR (101 MHz, $CDCl_3$) δ 147.66, 135.42, 130.77, 128.16, 125.74, 123.02, 60.37, 55.74, 53.40, 22.79, 22.58, 21.03, 14.20. $C_{12}H_{17}BrClNOS$, MS calculated for m/z $[M+H]^+$: 338.0 (calculated), 338.0 (found).

***(R)-1-(3,5-di(thiophen-2-yl)phenyl)ethan-1-amine*** **(A-4)**. To a solution of (*S*)-*N*-((*R*)-1-(3-bromo-5-chlorophenyl)ethyl)-2-methylpropane-2-sulfinamide (**A-3**) (1.0 equiv) in dioxane/$H_2O$ (4:1) in a microwave reaction vial was added 2-thienylboronic acid (2.5 equiv) and tripotassium phosphate (1.8 equiv). The mixture was purged with nitrogen for 5 min, then XPhosPdG2 (0.1 equiv) was added. The resulting mixture was heated in the biotage microwave reactor at 140 °C for 90 min. LCMS indicated that the starting material was completely consumed. The reaction mixture was diluted with EtOAc and extracted with water and brine. The organic layers were dried over anhydrous $Na_2SO_4$, filtered, and concentrated under reduced pressure to give a residue for the next step without purification. The crude was dissolved in 1,4-dioxane (0.1 M) and then concentrated HCl was added. The reactions were stirred at room temperature. When the starting material was consumed entirely as monitored by LCMS, solvent was removed under vacuum to give a white solid **A-4**, which was used in the next step without purification.

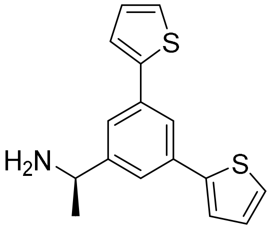


***(R)-1-(3,5-di(thiophen-2-yl)phenyl)ethan-1-amine*** (**A-4**). White solid, 98% yield. $^1$H NMR (400 MHz, $CDCl_3$) δ 7.85 (d, *J* = 1.8 Hz, 1H), 7.71 (d, *J* = 1.7 Hz, 2H), 7.53 (d, *J* = 3.6 Hz, 2H), 7.43 (d, *J* = 5.0 Hz, 2H), 7.19-6.90 (m, 2H), 4.55 (q, *J* = 6.8 Hz, 1H), 1.71 (d, *J* = 6.7 Hz, 3H). $^{13}$C NMR (101 MHz, $CDCl_3$) δ 143.94, 141.37, 137.45, 129.37, 126.82, 125.45, 124.22, 123.89, 52.18, 49.64, 49.43, 49.21, 49.00, 48.79, 48.57, 48.36, 20.84. $C_{16}H_{16}NS_2$, MS calculated for m/z $[M+H]^+$: 286.1 (calculated), 286.0 (found).

***(R)-5-bromo-N-(1-(3,5-di(thiophen-2-yl)phenyl)ethyl)-2-methylbenzamide*** (**A-5**). 5-bromo-2-methylbenzoic acid (1.1 equiv), HATU (1.1 equiv) and DIPEA (3.0 equiv) were dissolved in DMF (0.2 M) and stirred for 15 min. After that (*R*)-1-(3-bromo-5-chlorophenyl)ethan-1-amine (**A-4**) (1.0 equiv) was added and stirred at room temperature overnight. When the starting material was consumed entirely as monitored by LCMS, the mixture was diluted with EtOAc and was then washed with saturated aq. $NaHCO_3$, water, and brine. The organic layer was dried over $Na_2SO_4$, filtered, and concentrated under vacuum to give a residue that was purified by silica gel column chromatography (Hexanes/EtOAc = 4:1) to provide the desired product **A-5** as a slightly yellow solid.

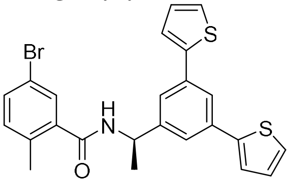


***(R)-5-bromo-N-(1-(3,5-di(thiophen-2-yl)phenyl)ethyl)-2-methylbenzamide*** (**A-5**). White solid, 88% yield. $^1$H NMR (400 MHz, $CDCl_3$) δ 7.75 (s, 1H), 7.51 (s, 3H), 7.46-7.41 (m, 1H), 7.37 (d, J = 3.6 Hz, 2H), 7.32 (d, J = 4.9 Hz, 2H), 7.22-6.85 (m, 3H), 6.09 (d, J = 8.0 Hz, 1H), 5.34 (td, J = 14.2, 13.8, 6.9 Hz, 1H), 2.37 (s, 3H), 1.65 (d, J = 7.0 Hz, 3H). $^{13}$C NMR (101 MHz, DMSO-$d_6$) δ 167.30, 147.20, 143.34, 139.64, 135.14, 135.12, 133.09, 132.39, 129.95, 128.99, 126.52,

124.81, 122.90, 121.11, 118.70, 48.86, 22.86, 19.20. $C_{24}H_{21}BrNOS_2$, MS calculated for m/z $[M+H]^+$: 482.0 (calculated), 482.0 (found).

***tert-butyl (R)-3-(((3-((1-(3,5-di(thiophen-2-yl)phenyl)ethyl)carbamoyl)-4-methylphenyl)amino)methyl)azetidine-1-carboxylate*** **(Jun1238).** To a solution of (*R*)-5-bromo-N-(1-(3,5-di(thiophen-2-yl)phenyl)ethyl)-2-methylbenzamide (**A-5**, 1.0 equiv) in dry toluene (8 mL) in a 15 mL sealed tube, tert-butyl 3-(aminomethyl)azetidine-1-carboxylate (1.5 equiv), XPhos (0.04 equiv), tris(dibenzylideneacetone)dipalladium (0.02 equiv) and $Cs_2CO_3$ (2.0 equiv) were added. The reaction mixture was degassed and purged with argon, and then the tube was sealed and heated to 110° C overnight. The mixture was diluted with EtOAc and was then washed with water and brine. The organic layer was dried over $Na_2SO_4$, filtered, and concentrated. The residue was purified by silica gel column chromatography (Hexanes/EtOAc = 1:1) to provide the desired intermediate. The intermediate was disolved in DCM and HCl (4M in dioxane, 10 equiv) was added. When the starting material was consumed entirely as monitored by LCMS, the mixture was diluted with DCM, adjusted to pH = 10, and then washed with water and brine. The organic layer was dried over $Na_2SO_4$, filtered, and concentrated under vacuum to give a residue which was purified by Prep-HPLC to give the final product **Jun1238** as a white solid.

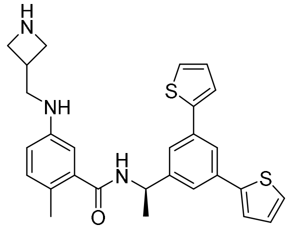

***(R)-5-((azetidin-3-ylmethyl)amino)-N-(1-(3,5-di(thiophen-2-yl)phenyl)ethyl)-2-methylbenzamide*** (**Jun1238**). White solid, 58% yield for two steps. $^1$H NMR (400 MHz, MeOD-$d_4$) δ 7.76 (d, J = 1.7 Hz, 1H), 7.64 (d, J = 1.7 Hz, 2H), 7.55 (d, J = 8.0 Hz, 2H), 7.49 (d, J = 3.6 Hz, 2H), 7.42 (dd, J = 12.0, 6.3 Hz, 4H), 7.11 (dd, J = 5.1, 3.5 Hz, 2H), 5.26 (q, J = 6.9 Hz, 1H), 4.14 (d, J = 10.1 Hz, 1H), 4.02 (t, J = 5.5 Hz, 1H), 3.80 (d, J = 6.9 Hz, 1H), 3.67-3.64 (m, 1H), 3.44 (q, J = 7.7, 6.9 Hz, 1H), 2.39 (s, 3H), 1.62 (d, J = 7.0 Hz, 3H). $^{13}$C NMR (101 MHz, MeOD-$d_4$) δ 168.62, 145.60, 143.34, 138.56, 137.78, 135.48, 132.51, 132.47, 127.95, 125.02, 123.64, 123.54, 122.48, 121.44, 121.09, 60.80, 52.78, 49.33, 28.89, 21.03, 18.11. $C_{28}H_{29}N_3OS_2$, MS calculated for m/z $[M+H]^+$: 488.2 (calculated), 488.1 (found).

***(R)-N-(1-(3,5-di(thiophen-2-yl)phenyl)ethyl)-2-methyl-5-((2-(piperidin-1-yl)ethyl)amino)benzamide*** (**Jun1271**). **Jun1271** was synthesized using the same coupling procedure as described for **Jun1238** with the starting material 2-(piperidin-1-yl)ethan-1-amine.

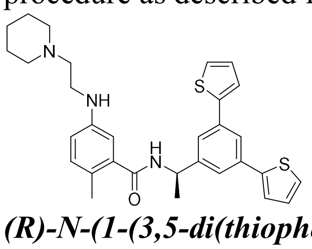

***(R)-N-(1-(3,5-di(thiophen-2-yl)phenyl)ethyl)-2-methyl-5-((2-(piperidin-1-yl)ethyl)amino)benzamide*** (**Jun1271**). White solid, 62% yield. $^1$H NMR (400 MHz, MeOD-$d_4$) δ 7.75 (d, J = 1.9 Hz, 1H), 7.61 (d, J = 1.7 Hz, 2H), 7.45 (d, J = 3.6 Hz, 2H), 7.38 (d, J = 5.1 Hz, 2H), 7.10 (dd, J = 5.1, 3.6 Hz, 2H), 7.01-6.95 (m, 1H), 6.65 (dd, J = 5.4, 2.9 Hz, 2H), 5.22 (q, J = 7.1 Hz, 1H), 3.40-3.33 (m, 2H), 2.96 (t, J = 6.5 Hz, 2H), 2.90 (t, J = 6.1 Hz, 4H), 2.23 (s, 3H), 1.70 (p, J = 5.8 Hz, 4H), 1.57 (d, J = 7.1 Hz, 3H), 1.52 (dt, J = 9.9, 5.3 Hz, 2H). $^{13}$C NMR (101

MHz, MeOD-$d_4$) δ 171.41, 146.09, 145.81, 143.45, 137.22, 135.37, 131.19, 127.94, 124.98, 123.52, 123.49, 122.39, 121.23, 114.44, 110.93, 56.08, 53.50, 49.14, 38.78, 23.43, 22.05, 21.13, 17.43. $C_{31}H_{35}N_3OS_2$, MS calculated for m/z $[M+H]^+$: 529.2 (calculated), 529.2 (found).

***(R)-5-amino-N-(1-(3,5-di(thiophen-2-yl)phenyl)ethyl)-2-methylbenzamide*** (**Jun11924**). 5-amino-2-methylbenzoic acid (1.1 equiv), HATU (1.1 equiv) and DIPEA (3 equiv) were dissolved in DMF (0.2 M) and stirred for 15 min. Then (*R*)-1-(3-bromo-5-chlorophenyl)ethan-1-amine (**A-4**) (1.0 equiv) was added and stirred at room temperature overnight. When the starting material was consumed entirely as monitored by LCMS, the mixture was diluted with EtOAc and was then washed with saturated aq. $NaHCO_3$, water, and brine. The organic layer was dried over $Na_2SO_4$, filtered, and concentrated under vacuum to give a residue that was purified by silica gel column chromatography (Hexanes/EtOAc = 2:1) to provide the desired product **Jun11924** as a slightly yellow solid.

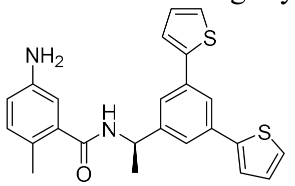


***(R)-5-amino-N-(1-(3,5-di(thiophen-2-yl)phenyl)ethyl)-2-methylbenzamide*** (**Jun11924**). White solid, 70% yield. $^1H$ NMR (400 MHz, $CDCl_3$) δ 7.76 (d, J = 1.8 Hz, 1H), 7.51 (t, J = 1.9 Hz, 3H), 7.42 (dd, J = 8.2, 2.2 Hz, 1H), 7.40-7.35 (m, 2H), 7.32 (d, J = 5.0 Hz, 2H), 7.13-7.04 (m, 3H), 6.08 (d, J = 8.0 Hz, 1H), 5.35 (p, J = 7.1 Hz, 1H), 2.38 (s, 3H), 1.66 (d, J = 6.9 Hz, 3H). $^{13}C$ NMR (101 MHz, $CDCl_3$) δ 169.50, 144.90, 144.28, 143.73, 137.05, 135.48, 131.81, 128.15, 125.29, 125.14, 123.76, 122.99, 122.62, 116.72, 113.54, 48.91, 21.97, 18.83. $C_{24}H_{22}N_2OS_2$, MS calculated for m/z $[M+H]^+$: 419.1 (calculated), 419.1 (found).

***(R)-5-(azetidin-3-ylamino)-N-(1-(3,5-di(thiophen-2-yl)phenyl)ethyl)-2-methylbenzamide*** **(Jun11922).** To a solution of (*R*)-5-amino-*N*-(1-(3,5-di(thiophen-2-yl)phenyl)ethyl)-2-methylbenzamide (**Jun11924**) (1.0 equiv) and tert-butyl 3-oxoazetidine-1-carboxylate (2.0 equiv) in dry MeOH (2 mL), HOAc (500 μL) was added and the solution was stirred for 3 hours at 50 ºC before $NaBH_3CN$ (3.0 equiv) was added. The mixture was poured into $H_2O$ (10 mL) and extracted with EtOAc (20 mL). The organic phase was concentrated, and the residue was purified by flash silica gel column chromatography to provide the Boc-protected intermediate. The intermediate was dissolved in DCM (2 mL), and HCl (4M in dioxane, 10 equiv) was added. When the starting material was consumed entirely as monitored by LCMS, the mixture was diluted with DCM, adjusted to pH = 10, and then washed with water and brine. The organic layer was dried over $Na_2SO_4$, filtered, and concentrated under vacuum to give a residue which was purified by Prep-HPLC to give the final product **Jun11922** as a white solid.

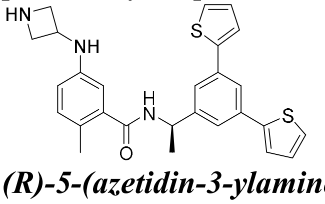


***(R)-5-(azetidin-3-ylamino)-N-(1-(3,5-di(thiophen-2-yl)phenyl)ethyl)-2-methylbenzamide*** (**Jun11922**). White solid, 63% yield for two steps. $^1H$ NMR (400 MHz, MeOD-$d_4$) δ 7.66 (d, J = 1.7 Hz, 1H), 7.52 (d, J = 1.6 Hz, 2H), 7.37 (d, J = 3.6 Hz, 2H), 7.30 (d, J = 5.2 Hz, 3H), 7.13 (d, J = 8.0 Hz, 1H), 7.06 (s, 1H), 7.00 (dd, J = 5.1, 3.6 Hz, 2H), 4.58 (p, J = 7.3 Hz, 1H), 4.25 (q, J = 11.3, 9.1 Hz, 3H), 4.14-3.69 (m, 1H), 2.19 (s, 3H), 1.50 (d, J = 7.0 Hz, 3H). $^{13}C$ NMR (101 MHz, MeOD-$d_4$) δ 170.02, 145.58, 143.35, 137.51, 135.46, 132.09, 128.00, 125.10, 123.67,

122.42, 121.43, 119.48, 116.56, 58.58, 51.02, 49.62, 21.12, 17.83. $C_{27}H_{27}N_3OS_2$, MS calculated for m/z $[M+H]^+$: 474.2 (calculated), 474.1 (found).

***(R)-N-(1-(3,5-di(thiophen-2-yl)phenyl)ethyl)-2-methyl-5-((1-methylpiperidin-4-yl)amino)benzamide*** (**Jun12112**). **Jun12112** was synthesized using the same reductive amination procedure as described for **Jun11922** with the starting material 1-methylpiperidin-4-one.

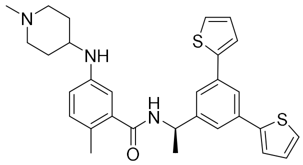


***(R)-N-(1-(3,5-di(thiophen-2-yl)phenyl)ethyl)-2-methyl-5-((1-methylpiperidin-4-yl)amino)benzamide*** (**Jun12112**). White solid, 65% yield. $^1$H NMR (400 MHz, $CDCl_3$) δ 7.73 (d, J = 1.7 Hz, 1H), 7.51 (d, J = 1.8 Hz, 2H), 7.36 (d, J = 3.6 Hz, 2H), 7.31 (d, J = 5.1 Hz, 2H), 7.09 (dd, J = 5.1, 3.5 Hz, 2H), 6.99 (dd, J = 8.2, 4.4 Hz, 1H), 6.66 (d, J = 2.5 Hz, 1H), 6.58 (dd, J = 8.3, 2.3 Hz, 1H), 6.21 (d, J = 8.1 Hz, 1H), 5.32 (q, J = 7.0 Hz, 1H), 3.32 (tt, J = 9.7, 4.4 Hz, 1H), 3.04 (t, J = 5.6 Hz, 1H), 2.93 (tt, J = 13.7, 6.3 Hz, 1H), 2.58 (d, J = 11.8 Hz, 3H), 2.51 (td, J = 12.1, 9.7, 3.7 Hz, 2H), 2.31 (s, 3H), 2.19-1.99 (m, 2H), 1.87 (d, J = 13.7 Hz, 2H), 1.62 (d, J = 6.9 Hz, 3H). $^{13}$C NMR (101 MHz, $CDCl_3$) δ 169.48, 144.86, 144.75, 143.68, 143.62, 137.31, 135.52, 131.97, 128.16, 125.35, 125.31, 123.79, 123.76, 122.94, 122.64, 115.84, 112.30, 59.35, 57.54, 54.64, 49.04, 27.71, 22.04, 18.81. $C_{30}H_{33}N_3OS_2$, MS calculated for m/z $[M+H]^+$: 516.2 (calculated), 516.2 (found).

***(R)-5-(2-aminoacetamido)-N-(1-(3,5-di(thiophen-2-yl)phenyl)ethyl)-2-methylbenzamide*** **(Jun11923).** (tert-butoxycarbonyl)glycine (1.1 equiv), HATU (1.1 equiv) and DIPEA (3 equiv) were dissolved in DMF (0.2 M) and stirred for 15 min. Then (*R*)-5-amino-*N*-(1-(3,5-di(thiophen-2-yl)phenyl)ethyl)-2-methylbenzamide (**Jun11924**) (1.0 equiv) was added and stirred at room temperature overnight. When the starting material was consumed entirely as monitored by LCMS, the mixture was diluted with EtOAc and was then washed with saturated aq. $NaHCO_3$, water, and brine. The organic layer was dried over $Na_2SO_4$, filtered, and concentrated under vacuum to give the Boc-protected intermediate. The intermediate was dissolved in DCM (2 mL), and HCl (4M in dioxane, 10 equiv) was added. When the starting material was consumed entirely as monitored by LCMS, the mixture was diluted with DCM, adjusted to pH = 10, and then washed with water and brine. The organic layer was dried over $Na_2SO_4$, filtered, and concentrated under vacuum to give a residue which was purified by Prep-HPLC to give the final product **Jun11923** as a white solid.

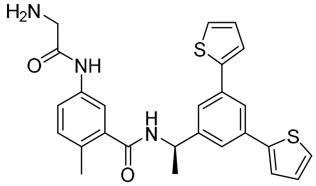


***(R)-5-(2-aminoacetamido)-N-(1-(3,5-di(thiophen-2-yl)phenyl)ethyl)-2-methylbenzamide*** (**Jun11923**). White solid, 75% yield. $^1$H NMR (400 MHz, MeOD-$d_4$) δ 7.79 (s, 1H), 7.65 (d, J = 9.7 Hz, 2H), 7.50 (s, 3H), 7.39 (d, J = 31.1 Hz, 3H), 7.23 (d, J = 8.3 Hz, 1H), 7.14 (s, 2H), 5.34-5.19 (m, 1H), 3.86 (s, 2H), 3.67 (s, 2H), 2.38 (d, J = 29.3 Hz, 3H), 1.62 (t, J = 7.4 Hz, 3H). $^{13}$C NMR (101 MHz, MeOD-$d_4$) δ 145.71, 143.35, 135.36, 131.48, 127.82, 124.88, 123.44, 122.32, 121.29, 120.77, 120.28, 117.47, 52.02, 20.94, 17.57, 7.78. $C_{26}H_{25}N_3O_2S_2$, MS calculated for m/z $[M+H]^+$: 476.1 (calculated), 476.2 (found).

***(R)-N-(1-(3,5-di(thiophen-2-yl)phenyl)ethyl)-5-(2-(dimethylamino)acetamido)-2-methylbenzamide*** (**Jun12111**). **Jun12111** was synthesized using the same amide coupling condition as described for **Jun11923** using the starting material dimethylglycine.

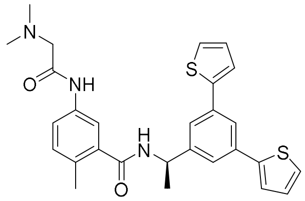

***(R)-N-(1-(3,5-di(thiophen-2-yl)phenyl)ethyl)-5-(2-(dimethylamino)acetamido)-2-methylbenzamide*** (**Jun12111**). White solid, 77% yield. $^1$H NMR (400 MHz, $CDCl_3$) δ 9.07 (s, 1H), 7.72 (d, J = 1.8 Hz, 1H), 7.58 (d, J = 2.3 Hz, 1H), 7.53 (d, J = 1.8 Hz, 2H), 7.36 (d, J = 3.6 Hz, 2H), 7.28 (d, J = 5.0 Hz, 2H), 7.12 (d, J = 8.3 Hz, 1H), 7.07 (dd, J = 5.1, 3.5 Hz, 2H), 6.62 (d, J = 8.0 Hz, 1H), 5.34 (m, 1H), 3.01-2.92 (m, 2H), 2.37 (s, 3H), 2.31 (s, 6H), 1.62 (d, J = 7.0 Hz, 3H). $^{13}$C NMR (101 MHz, $CDCl_3$) δ 168.90, 168.81, 144.88, 143.73, 136.80, 135.48, 135.44, 131.64, 131.53, 128.09, 125.21, 123.74, 122.99, 122.62, 120.91, 117.97, 63.43, 49.06, 45.96, 38.59, 22.02, 19.30. $C_{28}H_{29}N_3O_2S_2$, MS calculated for m/z $[M+H]^+$: 504.2 (calculated), 504.1 (found).

***(R)-N-(3-((1-(3,5-di(thiophen-2-yl)phenyl)ethyl)carbamoyl)-4-methylphenyl)piperidine-4-carboxamide*** (**Jun11945**). **Jun11945** was synthesized using the same procedures as described for **Jun11923** using the starting material 1-(tert-butoxycarbonyl)piperidine-4-carboxylic acid.

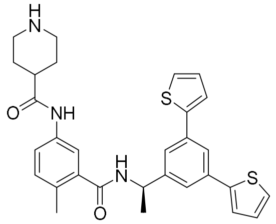

***(R)-N-(3-((1-(3,5-di(thiophen-2-yl)phenyl)ethyl)carbamoyl)-4-methylphenyl)piperidine-4-carboxamide*** (**Jun11945**). White solid, 80% yield for two steps. $^1$H NMR (400 MHz, MeOD-$d_4$) δ 7.78 (d, J = 1.8 Hz, 1H), 7.68 (d, J = 2.3 Hz, 1H), 7.64 (d, J = 1.6 Hz, 2H), 7.53-7.45 (m, 3H), 7.43 (d, J = 5.0 Hz, 2H), 7.22 (d, J = 8.3 Hz, 1H), 7.14 (dd, J = 5.2, 3.6 Hz, 2H), 5.27 (q, J = 7.2 Hz, 1H), 3.47 (dt, J = 13.3, 3.8 Hz, 2H), 3.09 (td, J = 12.5, 3.6 Hz, 2H), 2.75 (dq, J = 8.0, 5.4, 3.9 Hz, 1H), 2.34 (s, 3H), 2.18-1.83 (m, 4H), 1.61 (d, J = 7.2 Hz, 3H). $^{13}$C NMR (101 MHz, MeOD-$d_4$) δ 172.85, 170.53, 145.85, 143.41, 136.83, 136.02, 135.42, 131.09, 130.82, 127.95, 124.99, 123.57, 122.32, 121.32, 118.71, 49.25, 42.93, 40.13, 37.72, 21.15, 17.87. $C_{30}H_{31}N_3O_2S_2$, MS calculated for m/z $[M+H]^+$: 530.2 (calculated), 530.1 (found).

***(R)-5-bromo-N-(1-(3,5-di(thiophen-2-yl)phenyl)ethyl)-2-methylbenzamide*** (**A-6**). 5-methoxy-2-methylbenzoic acid (1.1 equiv), HATU (1.1 equiv) and DIPEA (3 equiv) were dissolved in DMF (0.2 M) and stirred for 15 min. After that (*R*)-1-(3-bromo-5-chlorophenyl)ethan-1-amine (**A-4**) was added and the solution was stirred at room temperature overnight. When the starting material was consumed entirely as monitored by LCMS, the mixture was diluted with EtOAc and was then washed with saturated aq. $NaHCO_3$, water, and brine. The organic layer was dried over $Na_2SO_4$, filtered, and concentrated under vacuum to give a residue which was purified by silica gel column chromatography (Hexanes/EtOAc = 4:1) to provide the desired product **A-6** as a slightly yellow solid.

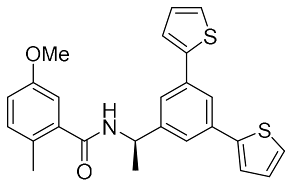

***(R)-N-(1-(3,5-di(thiophen-2-yl)phenyl)ethyl)-5-methoxy-2-methylbenzamide*** (**A-6**). White solid, 83% yield. $^{1}$H NMR (400 MHz, $CDCl_3$) δ 7.74 (t, J = 1.8 Hz, 1H), 7.51 (d, J = 1.7 Hz, 2H), 7.44-7.29 (m, 4H), 7.15-7.05 (m, 3H), 6.93 (d, J = 2.8 Hz, 1H), 6.84 (dd, J = 8.4, 2.8 Hz, 1H), 5.35 (m, 1H), 3.77 (s, 3H), 2.36 (s, 3H), 1.63 (d, J = 6.9 Hz, 3H). $^{13}$C NMR (101 MHz, $CDCl_3$) δ 169.06, 157.54, 144.68, 143.70, 137.18, 135.57, 132.04, 128.10, 127.56, 125.29, 123.74, 122.95, 122.74, 115.50, 112.32, 55.47, 49.03, 21.98, 18.87. $C_{25}H_{24}NO_2S_2$, MS calculated for m/z $[M+H]^+$: 434.1 (calculated), 434.1 (found).

***(R)-N-(1-(3,5-di(thiophen-2-yl)phenyl)ethyl)-5-hydroxy-2-methylbenzamide*** (**A-7**). (*R*)-5-bromo-*N*-(1-(3,5-di(thiophen-2-yl)phenyl)ethyl)-2-methylbenzamide (**A-6**) (1.0 equiv) was dissolved in DCM at 0 °C and $BBr_3$ (1 M solution in DCM, 3 equiv) was added slowly and the solution was stirred for 3 h. After the reaction was complete, the reaction was quenched with water and was then washed with DCM, water, and brine. The organic layer was dried over $Na_2SO_4$, filtered, and concentrated to give a residue that was purified by silica gel column chromatography (Hexanes/EtOAc =1:1) to provide the desired product **A-7** as a slightly yellow liquid.

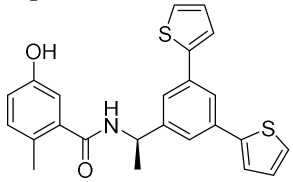

***(R)-N-(1-(3,5-di(thiophen-2-yl)phenyl)ethyl)-5-hydroxy-2-methylbenzamide*** (**A-7**). White solid, 90% yield. $^{1}$H NMR (400 MHz, $CDCl_3$) δ 7.74 (s, 1H), 7.66 (d, J = 1.8 Hz, 1H), 7.43 (d, J = 1.7 Hz, 2H), 7.28-7.26 (m, 2H), 7.24 (d, J = 1.3 Hz, 1H), 7.02 (dd, J = 5.1, 3.6 Hz, 2H), 6.84 (d, J = 8.3 Hz, 1H), 6.77 (d, J = 2.7 Hz, 1H), 6.64 (dd, J = 8.3, 2.7 Hz, 1H), 6.32 (d, J = 7.9 Hz, 1H), 5.23 (m, 1H), 2.22 (s, 3H), 1.53 (s, 3H). $^{13}$C NMR (101 MHz, $CDCl_3$) δ 172.35, 156.25, 147.17, 144.81, 138.75, 136.69, 132.68, 129.17, 126.89, 126.21, 124.77, 123.69, 122.68, 117.69, 114.77, 50.41, 22.42, 18.76. $C_{24}H_{22}NO_2S_2$, MS calculated for m/z $[M+H]^+$: 420.1 (calculated), 420.1 (found).

***(R)-N-(1-(3,5-di(thiophen-2-yl)phenyl)ethyl)-5-(2-(dimethylamino)ethoxy)-2-methylbenzamide*** (**Jun11875**). To a solution of *(R)-N-(1-(3,5-di(thiophen-2-yl)phenyl)ethyl)-5-hydroxy-2-methylbenzamide* (**A-7**) (1.0 equiv) and (2-bromoethyl)dimethylamine hydrobromide (1.1 equiv) in dry ACN, $Cs_2CO_3$ (2.0 equiv) was added. The reaction was stirred at 80 °C for overnight. After cooling down, the reaction was quenched with MeOH (1 mL). The mixture was diluted with EtOAc and was then washed with saturated aq. $NaHCO_3$, water, and brine. The organic layer was dried over $Na_2SO_4$, filtered, and concentrated to give a residue, which was purified by Prep-HPLC to afford the final product **Jun11875** as a white solid.

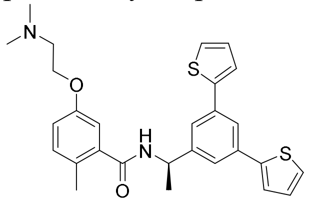

**(R)-N-(1-(3,5-di(thiophen-2-yl)phenyl)ethyl)-5-(2-(dimethylamino)ethoxy)-2-methylbenzamide** (**Jun11875**). White solid, 85% yield. $^{1}$H NMR (400 MHz, MeOD-d$_4$) δ 7.78 (d, J = 1.8 Hz, 1H), 7.62 (d, J = 1.7 Hz, 2H), 7.44 (dd, J = 24.7, 4.4 Hz, 4H), 7.20-7.09 (m, 3H), 6.94 (d, J = 7.9 Hz, 2H), 5.25 (q, J = 7.0 Hz, 1H), 4.12 (t, J = 5.3 Hz, 2H), 2.89 (t, J = 5.3 Hz, 2H), 2.43 (s, 6H), 2.29 (s, 3H), 1.59 (d, J = 7.1 Hz, 3H). $^{13}$C NMR (101 MHz, MeOD-d$_4$) δ 170.59, 156.16, 145.88, 143.42, 137.48, 135.38, 131.47, 127.85, 127.63, 124.90, 123.44, 122.33, 121.24, 115.71, 112.77, 64.15, 57.12, 49.10, 47.89, 47.84, 47.68, 47.63, 47.46, 47.42, 47.38, 47.17, 46.96, 43.71, 20.99, 17.39. $C_{28}H_{30}N_2O_2S_2$, MS calculated for m/z $[M+H]^+$: 491.2 (calculated), 491.1 (found).

***(R)-N-(1-(3,5-di(thiophen-2-yl)phenyl)ethyl)-2-methyl-5-(2-(pyrrolidin-1-yl)ethoxy)benzamide*** (**Jun11899**). **Jun11899** was synthesized using the same alkylation procedures described for **Jun11875** with the starting material 1-(2-bromoethyl)pyrrolidine.

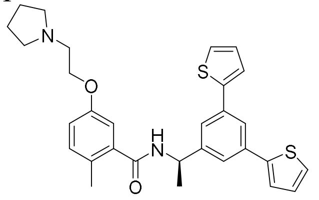

***(R)-N-(1-(3,5-di(thiophen-2-yl)phenyl)ethyl)-2-methyl-5-(2-(pyrrolidin-1-yl)ethoxy)benzamide*** (**Jun11899**). White solid, 75% yield. $^{1}$H NMR (400 MHz, MeOD-d$_4$) δ 7.66 (t, J = 1.7 Hz, 1H), 7.51 (d, J = 1.7 Hz, 2H), 7.36 (dd, J = 3.7, 1.1 Hz, 2H), 7.29 (dd, J = 5.1, 1.1 Hz, 2H), 7.10-6.92 (m, 3H), 6.90-6.74 (m, 2H), 5.13 (q, J = 7.0 Hz, 1H), 4.04 (t, J = 5.3 Hz, 2H), 3.00 (t, J = 5.3 Hz, 2H), 2.85-2.67 (m, 4H), 2.18 (s, 3H), 1.83-1.70 (m, 4H), 1.47 (d, J = 7.1 Hz, 3H). $^{13}$C NMR (101 MHz, MeOD-d$_4$) δ 170.57, 156.25, 145.94, 143.45, 137.53, 135.40, 131.49, 127.88, 127.60, 124.93, 123.47, 122.35, 121.26, 115.69, 112.79, 65.29, 54.29, 54.19, 49.11, 47.41, 47.19, 46.98, 22.71, 21.03, 17.43. $C_{30}H_{32}N_2O_2S_2$, MS calculated for m/z $[M+H]^+$: 517.2 (calculated), 517.1 (found).

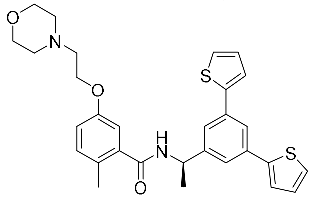

***(R)-N-(1-(3,5-di(thiophen-2-yl)phenyl)ethyl)-2-methyl-5-(2-morpholino-ethoxy)benzamide*** (**Jun118910**). **Jun118910** was synthesized using the same alkylation procedures described for **Jun11875** with the starting material 4-(2-bromoethyl)morpholine.

***(R)-N-(1-(3,5-di(thiophen-2-yl)phenyl)ethyl)-2-methyl-5-(2-morpholino-ethoxy)benzamide*** (**Jun118910**). White solid, 79% yield. $^{1}$H NMR (400 MHz, MeOD-d$_4$) δ 7.68 (d, J = 1.8 Hz, 1H), 7.52 (d, J = 1.7 Hz, 2H), 7.39-7.35 (m, 2H), 7.31 (d, J = 5.0 Hz, 2H), 7.09-6.94 (m, 3H), 6.88-6.70 (m, 2H), 5.15 (m, 1H), 4.02 (t, J = 5.4 Hz, 2H), 3.59 (t, J = 4.7 Hz, 4H), 2.68 (t, J = 5.4 Hz, 2H), 2.47 (t, J = 4.8 Hz, 4H), 2.19 (s, 3H), 1.49 (d, J = 7.1 Hz, 3H). $^{13}$C NMR (101 MHz, MeOD-d$_4$) δ 170.72, 156.58, 145.92, 143.48, 137.44, 135.40, 131.40, 130.95, 127.84, 127.25, 124.88, 123.42, 122.33, 121.26, 115.74, 112.68, 66.19, 65.24, 57.28, 53.71, 20.98, 17.35. $C_{30}H_{32}N_2O_3S_2$, MS calculated for m/z $[M+H]^+$: 533.2 (calculated), 533.1 (found).

***(R)-5-(2-aminoethoxy)-N-(1-(3,5-di(thiophen-2-yl)phenyl)ethyl)-2-methylbenzamide*** (**Jun11874**). **Jun11874** was synthesized using the same alkylation procedures described for **Jun11875** with the starting material of tert-butyl (2-bromoethyl)carbamate. The Boc-protected intermediate was dissolved in DCM (2 mL), and HCl (4M in dioxane, 10 equiv) was added.

When the starting material was consumed entirely as monitored by LCMS, the mixture was diluted with DCM, adjusted to pH = 10, and then washed with water and brine. The organic layer was dried over $Na_2SO_4$, filtered, and concentrated under vacuum to give a residue which was purified by Prep-HPLC to give the final product **Jun11874** as a white solid.

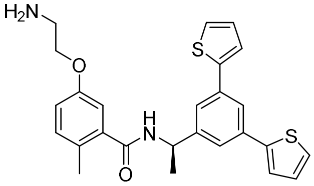

***(R)-5-(2-aminoethoxy)-N-(1-(3,5-di(thiophen-2-yl)phenyl)ethyl)-2-methylbenzamide*** (**Jun11874**). White solid, 88% yield. $^1$H NMR (400 MHz, MeOD-$d_4$) δ 7.78 (d, J = 1.8 Hz, 1H), 7.62 (d, J = 1.7 Hz, 2H), 7.48 (d, J = 3.6 Hz, 2H), 7.42 (d, J = 5.1 Hz, 2H), 7.20 (d, J = 8.3 Hz, 1H), 7.13 (dd, J = 5.1, 3.6 Hz, 2H), 7.04-6.85 (m, 2H), 5.28 (m, 1H), 4.21 (t, J = 5.0 Hz, 2H), 3.34 (d, J = 4.7 Hz, 2H), 2.31 (s, 3H), 1.60 (d, J = 7.1 Hz, 3H). $^{13}$C NMR (101 MHz, MeOD-$d_4$) δ 170.51, 155.88, 145.84, 143.42, 137.65, 135.41, 131.57, 128.02, 127.91, 124.97, 123.50, 122.40, 121.30, 115.77, 112.87, 66.74, 64.09, 49.22, 47.44, 47.23, 47.01, 38.93, 21.02, 17.47. $C_{26}H_{26}N_2O_2S_2$, MS calculated for m/z $[M+H]^+$: 463.1 (calculated), 463.1 (found).

***(R)-2-(3-((1-(3,5-di(thiophen-2-yl)phenyl)ethyl)carbamoyl)-4-methylphenoxy)-N,N,N-trimethylethan-1-aminium*** (**Jun11898**). To a solution of (*R*)-*N*-(1-(3,5-di(thiophen-2-yl)phenyl)ethyl)-5-(2-(dimethylamino)ethoxy)-2-methylbenzamide (**Jun11875**) (1.0 equiv) in dry MeOH, MeI (1.5 equiv) was added and the solution was stirred at room temperature for overnight. When the starting material was consumed entirely as monitored by LCMS, the mixture was concentrated under vacuum to give a residue, which was purified by Prep-HPLC to afford the final product **Jun11898** as a white solid.

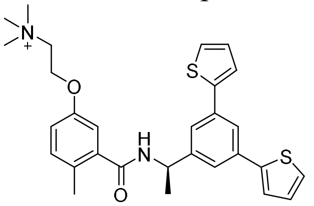

***(R)-2-(3-((1-(3,5-di(thiophen-2-yl)phenyl)ethyl)carbamoyl)-4-methylphenoxy)-N,N,N-trimethylethan-1-aminium*** (**Jun11898**). White solid, 50% yield. $^1$H NMR (400 MHz, MeOD-$d_4$) δ 7.67 (d, J = 1.8 Hz, 1H), 7.53 (d, J = 1.7 Hz, 2H), 7.38 (d, J = 3.6 Hz, 2H), 7.31 (d, J = 5.1 Hz, 2H), 7.08 (d, J = 8.2 Hz, 1H), 7.01 (dd, J = 5.1, 3.6 Hz, 2H), 6.88 (d, J = 8.1 Hz, 2H), 5.15 (q, J = 7.0 Hz, 1H), 4.36 (h, J = 2.4 Hz, 2H), 3.84-3.61 (m, 2H), 3.15 (s, 9H), 2.20 (s, 3H), 1.50 (d, J = 7.1 Hz, 3H). $^{13}$C NMR (101 MHz, MeOD-$d_4$) δ 170.32, 155.28, 145.91, 143.42, 137.62, 135.41, 131.67, 128.42, 127.92, 124.99, 123.54, 122.41, 121.26, 115.78, 113.07, 65.15, 62.02, 53.49, 53.45, 49.23, 47.42, 47.20, 46.99, 21.07, 17.49. $C_{29}H_{33}N_2O_2S_2$, MS calculated for m/z $[M]^+$: 505.2(calculated), 505.1 (found).

***Scheme 2. Synthesis of SARS-CoV-2 PL$^{pro}$ inhibitors.***

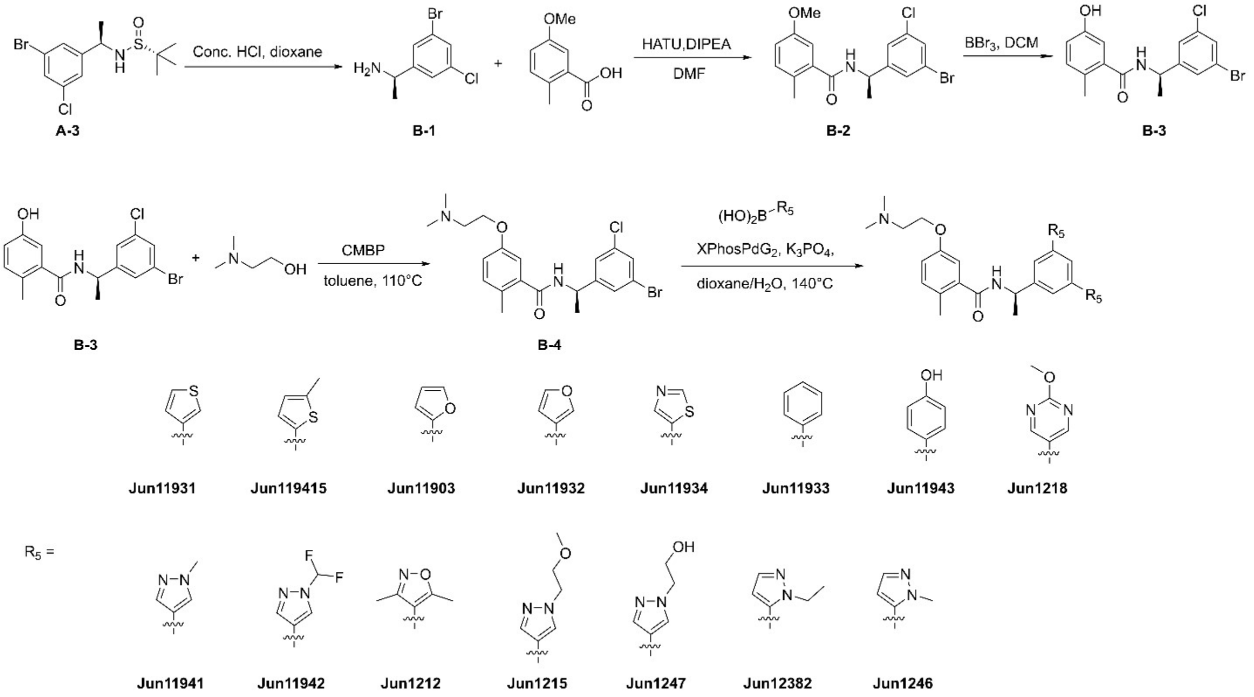


***Synthetic Procedures for Scheme 2***

***(R)-1-(3-bromo-5-chlorophenyl)ethan-1-amine*** **(B-1)**. To a solution of compound **A-3** (1 equiv) in 1,4-dioxane (0.1 M), concentrated HCl was added. The reactions were stirred at room temperature for 10 min. When the starting material was consumed entirely as monitored by LCMS, the mixture was concentrated under vacuum to give the desired compound **B-1** as a white solid, which was used for the next step without further purification.

***(R)-N-(1-(3-bromo-5-chlorophenyl)ethyl)-5-methoxy-2-methylbenzamide*** (**B-2**). 5-methoxy-2-methylbenzoic acid (1.1 equiv), HATU (1.1 equiv) and DIPEA (3 equiv) were dissolved in DMF (0.2 M) and stirred for 15 min. After that compound **B-1** was added and the solution was stirred at room temperature overnight. When the starting material was consumed entirely as monitored by LCMS, the mixture was diluted with EtOAc and was then washed with saturated aq. $NaHCO_3$, water, and brine. The organic layer was dried over $Na_2SO_4$, filtered, and concentrated under vacuum to give a residue that was purified by silica gel column chromatography (Hexanes/EtOAc = 3:1) to provide the desired product **B-2** as a slightly yellow solid.

***1-(3-bromo-5-chlorophenyl)ethan-1-one*** (**B-2**). White solid, 86% yield. $^1$H NMR (400 MHz, DMSO-d$_6$) δ 8.75 (d, J = 7.8 Hz, 1H), 7.59 (d, J = 1.8 Hz, 2H), 7.49 (d, J = 1.7 Hz, 1H), 7.15 (d, J = 8.3 Hz, 1H), 6.92 (dd, J = 8.3, 2.7 Hz, 1H), 6.89 (d, J = 2.8 Hz, 1H), 5.08 (m, 1H), 3.76 (s, 3H), 2.19 (s, 3H), 1.42 (d, J = 7.0 Hz, 3H). $^{13}$C NMR (101 MHz, DMSO-d$_6$) δ 168.63, 157.40, 150.05, 138.01, 134.56, 131.96, 129.44, 128.33, 127.26, 125.86, 122.64, 115.20, 113.16, 55.69, 48.30, 38.70, 22.54, 18.73. $C_{17}H_{17}BrClNO_2$, MS calculated for m/z $[M+H]^+$: 382.0209 (calculated), 382.0 (found).

***(R)-N-(1-(3-bromo-5-chlorophenyl)ethyl)-5-hydroxy-2-methylbenzamide*** (**B-3**). Compound **B-2** (1 equiv) was dissolved in DCM at 0 °C, and $BBr_3$ (1 M solution in DCM, 3 equiv) was added slowly and the solution was stirred for 3 h. When the starting material was consumed entirely as monitored by LCMS, the reaction was quenched with water and washed with DCM, water, and brine, respectively. The organic layer was dried over $Na_2SO_4$, filtered, and concentrated. The residue was purified by silica gel column chromatography (Hexanes/EtOAc = 1:1) to provide the desired product **B-3** as a colorless liquid.

***1-(3-bromo-5-chlorophenyl)ethan-1-one*** (**B-3**). White solid, 89% yield. $^1$H NMR (400 MHz, DMSO-$d_6$) δ 9.43 (s, 1H), 8.74 (d, J = 7.9 Hz, 1H), 7.58 (d, J = 5.0 Hz, 2H), 7.49 (d, J = 1.9 Hz, 1H), 7.02 (d, J = 8.1 Hz, 1H), 6.75 (dd, J = 10.7, 2.6 Hz, 2H), 5.09 (m, 1H), 2.16 (s, 3H), 1.41 (d, J = 7.2 Hz, 3H). $^{13}$C NMR (101 MHz, DMSO-$d_6$) δ 168.93, 155.42, 150.10, 137.99, 134.57, 131.85, 129.41, 128.31, 125.83, 125.37, 122.63, 116.68, 114.34, 60.20, 48.22, 22.49, 21.17, 18.71, 14.51. $C_{16}H_{15}BrClNO_2$, MS calculated for m/z $[M+H]^+$: 368.0 (calculated), 368.1 (found).

***(R)-N-(1-(3-bromo-5-chlorophenyl)ethyl)-5-(2-(dimethylamino)ethoxy)-2-methylbenzamide*** (**B-4**). To a solution of compound **B-3** (1 equiv) in dry toluene in a 15 mL sealed tube, 2-(dimethylamino)ethanol (1.5 equiv) and cyanomethylene trimethylphosphorane (CMMP) (2.0 equiv) were added. The reaction mixture was degassed and purged with argon, and then the tube was sealed and heated to 110° C overnight. When the starting material was consumed entirely as monitored by LCMS, the mixture was diluted with EtOAc and washed with water and brine. The organic layer was dried over $Na_2SO_4$, filtered, and concentrated. The residue was purified by silica gel column chromatography (DCM/MeOH = 20:1) to provide the desired product **B-4** as a white solid.

***(R)-N-(1-(3-bromo-5-chlorophenyl)ethyl)-5-(2-(dimethylamino)ethoxy)-2-methylbenzamide*** (**B-4**). White solid, 94% yield. $^1$H NMR (400 MHz, DMSO-$d_6$) δ 8.76 (d, J = 7.9 Hz, 1H), 7.70 (s, 1H), 7.62 (s, 2H), 7.14 (d, J = 8.2 Hz, 1H), 6.95-6.88 (m, 2H), 5.07 (m, 1H), 4.05 (t, J = 5.7 Hz, 2H), 2.64 (t, J = 5.7 Hz, 2H), 2.23 (s, 6H), 2.19 (s, 3H), 1.41 (d, J = 7.0 Hz, 3H). $^{13}$C NMR (101 MHz, DMSO-$d_6$) δ 168.57, 156.59, 150.33, 137.90, 132.05, 131.99, 128.70, 127.41, 122.86, 115.78, 113.83, 66.34, 58.05, 48.26, 45.90, 22.56, 18.80. $C_{20}H_{24}BrClN_2O_2$, MS calculated for m/z $[M+H]^+$: 439.0788 (calculated), 439.0795 (found).

**General procedures for the Suzuki coupling I**. To a solution of compound **B-4** (1 equiv) in dioxane/$H_2O$ (4:1) in a microwave reaction vial was added corresponding boronic acid or ester (2.5 equiv) and tripotassium phosphate (1.8 equiv). The mixture was purged with nitrogen for 5 min. Then XPhosPdG2 (0.1 equiv) was added, and the solution was heated in the biotage microwave reactor at 140 °C for 90 min. After the reaction was completed, the reaction mixture was diluted with EtOAc and washed with water and brine. The organic layer was separated, dried over anhydrous $Na_2SO_4$, filtered, and concentrated under vacuum to give a residue which was purified by Prep-HPLC to give the final product.

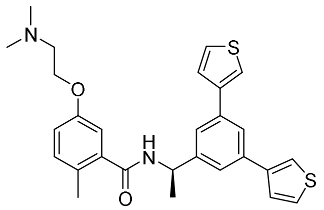

***(R)-N-(1-(3,5-di(thiophen-3-yl)phenyl)ethyl)-5-(2-(dimethylamino)ethoxy)-2-methylbenzamide*** (**Jun11931**). White solid, 90% yield. $^1$H NMR (400 MHz, DMSO-$d_6$) δ 8.76 (d, J = 8.3 Hz, 1H), 7.95 (d, J = 11.6 Hz, 3H), 7.68 (d, J = 13.8 Hz, 6H), 7.14 (d, J = 8.1 Hz, 1H), 6.91 (d, J = 7.8 Hz, 2H), 5.27-5.20 (m, 1H), 4.07 (t, J = 5.8 Hz, 2H), 2.69 (t, J = 5.7 Hz, 2H), 2.26 (s, 6H), 2.23 (s, 3H), 1.51 (d, J = 6.9 Hz, 3H). $^{13}$C NMR (101 MHz, DMSO-$d_6$) 168.46, 156.53, 146.61, 141.98, 138.49, 136.21, 131.91, 127.48, 127.39, 126.83, 123.20, 122.74, 121.67, 115.82, 113.68, 66.15, 57.92, 48.89, 45.75, 23.32, 18.84. $C_{28}H_{30}N_2O_2S_2$, HRMS calculated for m/z $[M+H]^+$: 491.1827(calculated), 491.1832(found)

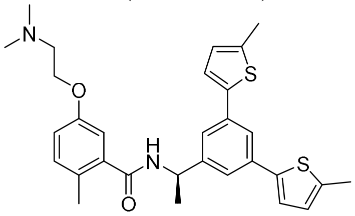


***(R)-N-(1-(3,5-bis(5-methylthiophen-2-yl)phenyl)ethyl)-5-(2-(dimethylamino)ethoxy)-2-methylbenzamide*** (**Jun119415**). White solid, 86% yield. $^1$H NMR (400 MHz, $CDCl_3$) δ 7.61 (d, J = 1.9 Hz, 1H), 7.43 (s, 2H), 7.15 (d, J = 3.6 Hz, 2H), 7.12 (s, 1H), 6.92 (d, J = 2.8 Hz, 1H), 6.82 (d, J = 5.6 Hz, 1H), 6.74 (d, J = 3.6 Hz, 2H), 6.28 (d, J = 8.0 Hz, 1H), 5.36-5.30 (m, 1H), 4.35-4.33 (m, 2H), 3.46-3.43 (m, 2H), 2.91 (s, 6H), 2.51 (s, 6H), 2.36 (s, 3H), 1.64 (d, J = 6.9 Hz, 3H). $^{13}$C NMR (101 MHz, $CDCl_3$) δ 141.30, 139.99, 135.75, 132.33, 126.28, 123.50, 122.10, 115.84, 62.88, 56.42, 49.30, 43.68, 21.76, 18.92, 15.47. $C_{30}H_{34}N_2O_2S_2$, HRMS calculated for m/z $[M+H]^+$: 519.2140(calculated), 519.2144(found)

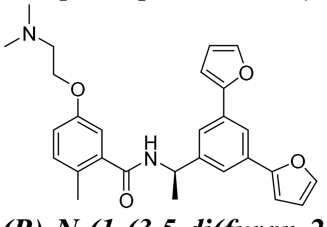


***(R)-N-(1-(3,5-di(furan-2-yl)phenyl)ethyl)-5-(2-(dimethylamino)ethoxy)-2-methylbenzamide*** (**Jun11903**). White solid, 92% yield. $^1$H NMR (400 MHz, DMSO-$d_6$) δ 8.82 (d, J = 8.0 Hz, 1H), 7.90 (d, J = 2.1 Hz, 1H), 7.79 (s, 2H), 7.68 (s, 2H), 7.15 (d, J = 7.6 Hz, 1H), 7.04 (s, 2H), 6.93 (d, J = 3.4 Hz, 2H), 6.64 (s, 2H), 5.21-5.14 (m, 1H), 4.09 (t, J = 5.8 Hz, 2H), 2.75 (t, J = 5.7 Hz, 2H), 2.30 (s, 6H), 2.21 (s, 3H), 1.49 (d, J = 6.9 Hz, 3H). $^{13}$C NMR (101 MHz, DMSO-$d_6$) δ 168.50, 156.45, 153.23, 146.93, 143.55, 138.39, 131.92, 131.40, 128.70, 127.50, 122.86, 120.66, 117.24, 115.80, 113.74, 112.62, 106.81, 65.89, 57.79, 48.78, 45.57, 22.78, 18.79. $C_{28}H_{30}N_2O_4$, HRMS calculated for m/z $[M+H]^+$: 459.2284(calculated), 459.2291(found)

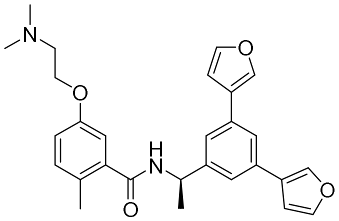


***(R)-N-(1-(3,5-di(thiophen-3-yl)phenyl)ethyl)-5-(2-(dimethylamino)ethoxy)-2-methylbenzamide*** (**Jun11932**). White solid, 90% yield. $^1$H NMR (400 MHz, DMSO-$d_6$) δ 8.71 (d, J = 8.5 Hz, 1H), 8.23 (s, 2H), 7.77 (d, J = 11.2 Hz, 3H), 7.55 (s, 2H), 7.14 (d, J = 7.4 Hz, 1H), 7.02 (s, 2H), 6.91 (d, J = 5.1 Hz, 2H), 5.21-5.14 (m, 1H), 4.06 (t, J = 5.7 Hz, 2H), 2.68 (t, J = 5.7 Hz, 2H), 2.25 (s, 6H), 2.21 (s, 3H), 1.49 (d, J = 6.9 Hz, 3H). $^{13}$C NMR (101 MHz, DMSO-$d_6$) δ 168.45, 156.53,

146.51, 144.74, 139.93, 138.49, 132.90, 131.89, 127.35, 126.31, 122.41, 121.61, 115.78, 113.68, 109.24, 66.17, 57.94, 48.83, 45.76, 23.23, 18.80. $C_{28}H_{30}N_2O_4$, HRMS calculated for m/z $[M+H]^+$: 459.2284(calculated), 459.2297(found)

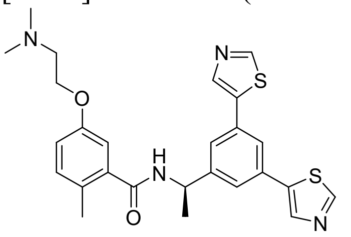


***(R)-N-(1-(3,5-di(thiazol-5-yl)phenyl)ethyl)-5-(2-(dimethylamino)ethoxy)-2-methylbenzamide*** (**Jun11934**). White solid, 78% yield. $^1$H NMR (400 MHz, DMSO-$d_6$) δ 8.90 (s, 2H), 8.59 (d, J = 8.1 Hz, 1H), 8.21 (s, 2H), 7.70 (s, 1H), 7.43 (s, 2H), 6.91 (d, J = 8.5 Hz, 1H), 6.69 (d, J = 6.7 Hz, 2H), 4.99-4.91 (m, 1H), 3.85 (d, J = 5.9 Hz, 2H), 2.25 (s, 2H), 2.07 (s, 6H), 1.97 (s, 3H), 1.25 (d, J = 7.0 Hz, 3H). $^{13}$C NMR (101 MHz, DMSO-$d_6$) δ 168.57, 156.46, 154.55, 147.91, 140.66, 138.42, 138.34, 132.29, 131.95, 127.45, 124.98, 123.00, 115.91, 113.68, 65.84, 57.71, 48.64, 45.52, 23.00, 18.83. $C_{26}H_{28}N_4O_2S_2$, HRMS calculated for m/z $[M+H]^+$: 493.1732(calculated), 493.1740(found)

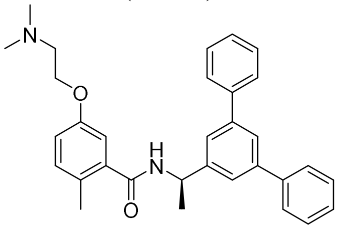


***(R)-N-(1-([1,1':3',1''-terphenyl]-5'-yl)ethyl)-5-(2-(dimethylamino)ethoxy)-2-methylbenzamide*** (**Jun11933**). White solid, 88% yield. $^1$H NMR (400 MHz, DMSO-$d_6$) δ 8.82 (d, J = 8.3 Hz, 1H), 7.77 (d, J = 7.6 Hz, 5H), 7.71 (s, 2H), 7.50 (t, J = 7.5 Hz, 4H), 7.41 (d, J = 7.3 Hz, 2H), 7.14 (d, J = 8.2 Hz, 1H), 6.92 (d, J = 7.7 Hz, 2H), 5.32-5.25 (m, 1H), 4.06 (t, J = 5.7 Hz, 2H), 2.69 (t, J = 5.6 Hz, 2H), 2.26 (s, 6H), 2.23 (s, 3H), 1.54 (d, J = 7.0 Hz, 3H). $^{13}$C NMR (101 MHz, DMSO-$d_6$) δ 168.48, 156.52, 146.77, 141.44, 140.75, 138.47, 131.92, 129.39, 128.06, 127.39, 124.22, 124.02, 115.80, 113.68, 66.08, 57.91, 48.92, 45.72, 23.23, 18.84. $C_{32}H_{34}N_2O_2$, HRMS calculated for m/z $[M+H]^+$: 479.2698(calculated), 479.2709(found)

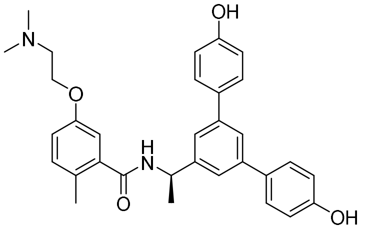


***(R)-N-(1-(4,4''-dihydroxy-[1,1':3',1''-terphenyl]-5'-yl)ethyl)-5-(2-(dimethylamino)ethoxy)-2-methylbenzamide*** (**Jun11943**). White solid, 88% yield. $^1$H NMR (400 MHz, DMSO-$d_6$) δ 9.59 (s, 2H), 8.77 (d, J = 8.3 Hz, 1H), 7.59-7.52 (m, 7H), 7.14 (d, J = 8.3 Hz, 1H), 6.92-6.86 (m, 6H), 5.25-5.18 (m, 1H), 4.06 (d, J = 5.9 Hz, 2H), 2.68 (t, J = 5.6 Hz, 2H), 2.26 (s, 6H), 2.22 (s, 3H), 1.50 (d, J = 7.0 Hz, 3H). $^{13}$C NMR (101 MHz, DMSO-$d_6$) δ 168.41, 157.66, 156.51, 146.37, 141.27, 138.51, 131.90, 131.65, 128.35, 127.38, 122.61, 122.51, 116.15, 115.79, 113.65, 66.13, 57.92, 48.93, 46.03, 45.73, 23.22, 18.83. $C_{32}H_{34}N_2O_4$, HRMS calculated for m/z $[M+H]^+$: 511.2597(calculated), 511.2605(found)

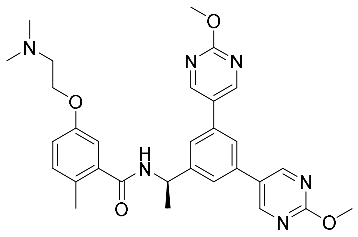

***(R)-N-(1-(3,5-bis(2-methoxypyrimidin-5-yl)phenyl)ethyl)-5-(2-(dimethylamino)ethoxy)-2-methylbenzamide*** (**Jun1218)**. White solid, 84% yield. $^1$H NMR (400 MHz, DMSO-d$_6$) δ 8.99 (s, 4H), 8.96 (s, 1H), 8.71 (d, J = 8.2 Hz, 1H), 7.93 (s, 1H), 7.76 (s, 2H), 7.14 (d, J = 8.4 Hz, 1H), 6.93 (s, 1H), 5.27-5.20 (m, 1H), 4.25 (t, J = 5.0 Hz, 2H), 3.92 (s, 6H), 3.44 (q, J = 4.7 Hz, 2H), 2.80 (s, 6H), 2.17 (s, 3H), 1.46 (d, J = 7.1 Hz, 3H). $^{13}$C NMR (101 MHz, DMSO-d$_6$) δ 168.40, 165.20, 158.85, 158.51, 158.00, 157.95, 155.70, 147.24, 138.67, 135.32, 131.97, 128.15, 127.58, 127.50, 124.16, 123.18, 118.26, 115.98, 114.09, 62.87, 55.90, 55.25, 55.22, 48.98, 43.27, 23.50, 18.82. $C_{30}H_{34}N_6O_4$, HRMS calculated for m/z [M+H]$^+$: 543.2720(calculated), 543.2727(found)

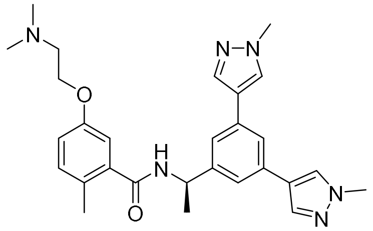

***(R)-N-(1-(3,5-bis(1-methyl-1H-pyrazol-4-yl)phenyl)ethyl)-5-(2 (dimethylamino)ethoxy)-2-methylbenzamide*** (**Jun11941**). White solid, 79% yield. $^1$H NMR (400 MHz, DMSO-d$_6$) δ 8.68 (d, J = 8.3 Hz, 1H), 8.15 (s, 2H), 7.89 (s, 2H), 7.66 (s, 1H), 7.43 (s, 2H), 7.14 (d, J = 8.1 Hz, 1H), 6.92 (d, J = 7.9 Hz, 2H), 5.18-5.11 (m, 1H), 4.08 (t, J = 5.8 Hz, 2H), 3.88 (s, 6H), 2.72 (t, J = 5.7 Hz, 2H), 2.28 (s, 6H), 2.22 (s, 3H), 1.48 (d, J = 7.1 Hz, 3H). $^{13}$C NMR (101 MHz, DMSO-d$_6$) δ 168.40, 156.49, 146.33, 138.55, 136.51, 133.50, 131.88, 128.30, 127.40, 122.51, 120.92, 120.49, 115.74, 113.73, 66.01, 57.84, 48.83, 45.64, 39.14, 23.14, 23.00, 18.82. $C_{28}H_{34}N_6O_2$, HRMS calculated for m/z [M+H]$^+$: 487.2821(calculated), 487.2830(found)

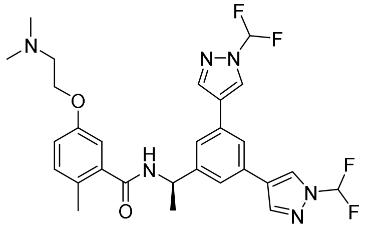

***(R)-N-(1-(3,5-bis(1-(difluoromethyl)-1H-pyrazol-4-yl)phenyl)ethyl)-5-(2-(dimethylamino)ethoxy)-2-methylbenzamide*** (**Jun11942**). White solid, 86% yield. $^1$H NMR (400 MHz, CDCl$_3$) δ 8.11 (s, 2H), 7.96 (s, 2H), 7.54 (s, 1H), 7.46 (s, 2H), 7.38 (s, 1H), 7.27 (s, 1H), 7.12-7.08 (m, 1H), 6.98 (d, J = 2.7 Hz, 1H), 6.87 (dd, J = 8.4, 2.8 Hz, 1H), 6.15 (d, J = 7.8 Hz, 1H), 5.39-5.31 (m, 1H), 4.03 (t, J = 5.6 Hz, 2H), 2.71 (t, J = 5.6 Hz, 2H), 2.36 (s, 3H), 2.32 (s, 6H), 1.64 (d, J = 6.9 Hz, 3H). $^{13}$C NMR (101 MHz, CDCl$_3$) δ 168.94, 156.78, 145.16, 139.66, 139.64, 139.62, 136.85, 132.42, 132.12, 127.99, 125.37, 123.11, 123.09, 122.85, 115.71, 113.63, 113.58, 111.09, 108.60, 66.08, 58.27, 49.11, 45.80, 21.97, 18.89. $C_{28}H_{30}F_4N_6O_2$, HRMS calculated for m/z [M+H]$^+$: 559.2446(calculated), 559.2451(found)

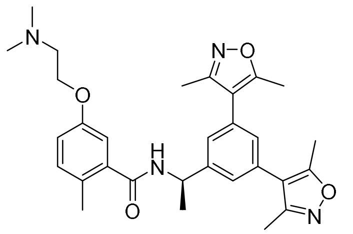

***(R)-N-(1-(3,5-bis(3,5-dimethylisoxazol-4-yl)phenyl)ethyl)-5-(2-(dimethylamino)ethoxy)-2-methylbenzamide*** (**Jun1212**). White solid, 77% yield. $^{1}H$ NMR (400 MHz, DMSO-$d_6$) δ 8.74 (d, J = 8.2 Hz, 1H), 7.35 (s, 2H), 7.21 (s, 1H), 7.12 (d, J = 8.3 Hz, 1H), 6.93-6.88 (m, 2H), 5.22-5.14 (m, 1H), 4.23 (t, J = 5.1 Hz, 2H), 3.44 (s, 2H), 2.79 (s, 6H), 2.40 (s, 6H), 2.23 (s, 6H), 2.14 (s, 3H), 1.44 (d, J = 7.0 Hz, 3H). $^{13}C$ NMR (101 MHz, DMSO-$d_6$) δ 168.32, 165.86, 158.63, 155.67, 146.49, 138.53, 132.00, 130.93, 128.21, 127.85, 126.11, 116.12, 115.92, 114.03, 62.88, 55.89, 48.65, 43.27, 22.98, 18.81, 11.95, 11.04. $C_{30}H_{36}N_4O_4$, HRMS calculated for m/z $[M+H]^+$: 517.2815(calculated), 517.2821(found)

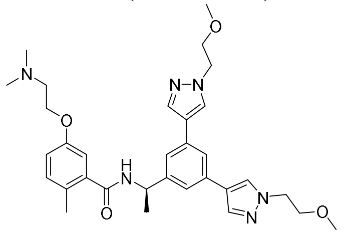

***(R)-N-(1-(3,5-bis(1-(2-methoxyethyl)-1H-pyrazol-4-yl)phenyl)ethyl)-5-(2-(dimethylamino)ethoxy)-2-methylbenzamide*** (**Jun1215**). White solid, 74% yield. $^{1}H$ NMR (400 MHz, DMSO-$d_6$) δ 8.69 (d, J = 8.3 Hz, 1H), 8.18 (s, 2H), 7.93 (s, 2H), 7.68 (s, 1H), 7.44 (s, 2H), 7.14 (d, J = 8.1 Hz, 1H), 6.91 (d, J = 8.0 Hz, 2H), 5.18-5.11 (m, 1H), 4.29 (t, J = 5.3 Hz, 4H), 4.07 (t, J = 5.7 Hz, 2H), 3.72 (t, J = 5.3 Hz, 4H), 3.25 (s, 6H), 2.70 (t, J = 5.7 Hz, 2H), 2.27 (s, 6H), 2.22 (s, 3H), 1.48 (d, J = 7.0 Hz, 3H). $^{13}C$ NMR (101 MHz, DMSO-$d_6$) δ 168.40, 156.50, 146.33, 138.56, 136.68, 133.48, 131.88, 128.12, 127.37, 122.26, 120.92, 120.41, 115.71, 113.71, 73.99, 70.97, 66.04, 58.47, 57.86, 51.85, 48.84, 46.03, 45.67, 25.42, 23.19, 18.82, 9.50. $C_{32}H_{42}N_6O_4$, HRMS calculated for m/z $[M+H]^+$: 575.3346(calculated), 575.3352(found)

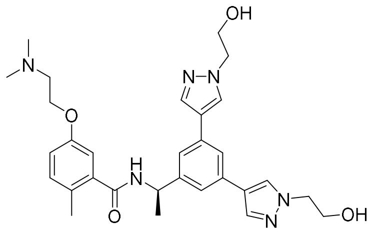

***(R)-N-(1-(3,5-bis(1-(2-hydroxyethyl)-1H-pyrazol-4-yl)phenyl)ethyl)-5-(2-(dimethylamino)ethoxy)-2-methylbenzamide*** (**Jun1247**). White solid, 72% yield. $^{1}H$ NMR (400 MHz, DMSO-$d_6$) δ 8.72 (d, J = 8.3 Hz, 1H), 8.19 (s, 2H), 7.93 (s, 2H), 7.69 (s, 1H), 7.45 (s, 2H), 7.19 (d, J = 8.4 Hz, 1H), 6.97 (d, J = 10.8 Hz, 2H), 5.20-5.12 (m, 1H), 4.30 (q, J = 5.9 Hz, 4H), 4.18 (t, J = 5.6 Hz, 2H), 3.78 (t, J = 5.6 Hz, 2H), 3.73 (t, J = 5.3 Hz, 2H), 3.50 (q, J = 4.6 Hz, 2H), 2.86 (s, 6H), 2.24 (s, 3H), 1.49 (d, J = 7.1 Hz, 3H). $^{13}C$ NMR (101 MHz, DMSO-$d_6$) δ 168.40, 158.76, 158.42, 155.67, 146.54, 142.67, 138.55, 138.45, 132.00, 131.22, 128.18, 126.94, 126.57, 115.98, 114.04, 106.63, 62.85, 55.93, 48.70, 43.29, 38.09, 23.03, 18.82. $C_{30}H_{38}N_6O_4$, HRMS calculated for m/z $[M+H]^+$: 547.3033(calculated), 547.3040(found)

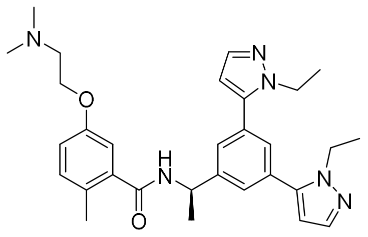

***(R)-N-(1-(3,5-bis(1-ethyl-1H-pyrazol-5-yl)phenyl)ethyl)-5-(2-(dimethylamino)ethoxy)-2-methylbenzamide*** (**Jun12382**). White solid, 74% yield. $^{1}$H NMR (400 MHz, DMSO-$d_6$) δ 8.84 (d, J = 8.1 Hz, 1H), 7.55 (d, J = 9.1 Hz, 4H), 7.43 (d, J = 1.8 Hz, 1H), 7.19 (d, J = 8.2 Hz, 1H), 6.98 (d, J = 8.7 Hz, 2H), 6.44 (s, 2H), 5.30-5.23 (m, J = 7.2 Hz, 1H), 4.31 (t, J = 4.9 Hz, 2H), 4.19 (q, J = 7.1 Hz, 4H), 3.51 (q, J = 4.6 Hz, 2H), 2.87 (s, 6H), 2.20 (s, 3H), 1.53 (d, J = 7.0 Hz, 3H), 1.33 (t, J = 7.2 Hz, 6H). $^{13}$C NMR (101 MHz, DMSO-$d_6$) δ 168.42, 155.69, 146.62, 142.26, 138.69, 138.49, 131.98, 131.46, 128.18, 127.16, 126.80, 115.88, 114.10, 106.66, 62.90, 55.89, 48.64, 44.62, 43.26, 22.70, 18.77, 15.93. $C_{30}H_{38}N_6O_2$, HRMS calculated for m/z $[M+H]^+$: 515.3134 (calculated), 515.3140 (found)

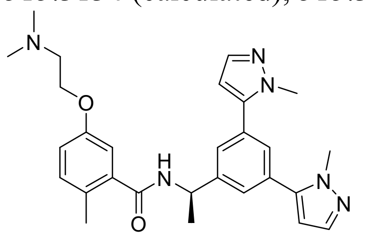

***(R)-N-(1-(3,5-bis(1-methyl-1H-pyrazol-5-yl)phenyl)ethyl)-5-(2-(dimethylamino)ethoxy)-2-methylbenzamide*** (**Jun1246**). White solid, 78% yield. $^{1}$H NMR (400 MHz, DMSO-$d_6$) δ 8.78 (s, 4H), 8.76 (s, 2H), 7.77 (s, 1H), 7.63 (s, 2H), 7.44 (s, 1H), 7.20 (d, J = 8.2 Hz, 1H), 6.98 (s, 2H), 5.28-5.21 (m, 1H), 4.32 (d, J = 4.9 Hz, 2H), 4.05 (s, 12H), 3.51 (t, J = 6.4 Hz, 2H), 2.86 (d, J = 3.2 Hz, 6H), 2.69 (s, 2H), 2.24 (s, 4H), 1.50 (d, J = 6.9 Hz, 3H). $^{13}$C NMR (101 MHz, DMSO-$d_6$) δ 168.40, 158.76, 158.42, 155.67, 146.54, 142.67, 138.55, 138.45, 132.00, 131.22, 128.18, 126.94, 126.57, 115.98, 114.04, 106.63, 62.85, 55.93, 48.70, 43.29, 38.09, 23.03, 18.82. $C_{28}H_{34}N_6O_2$, HRMS calculated for m/z $[M+H]^+$: 487.2821 (calculated), 487.2828 (found)

***Scheme 3. Synthesis of SARS-CoV-2 PL$^{pro}$ inhibitors***

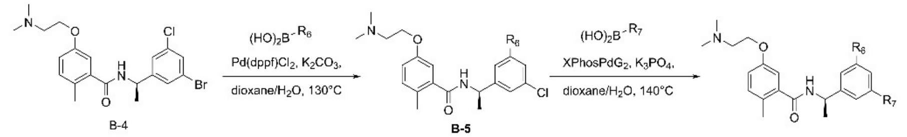


B-4 B-5

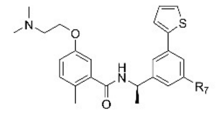

$R_7$ =

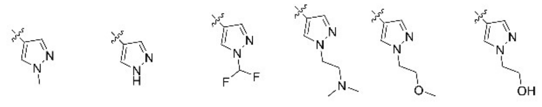

Jun1211 Jun1244 Jun1251 Jun1252 Jun1254 Jun1257

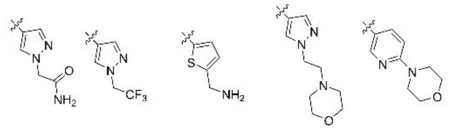

Jun121210 Jun12149 Jun12129 Jun121910 Jun12511

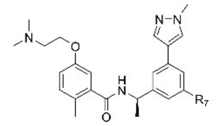

$R_7$ =

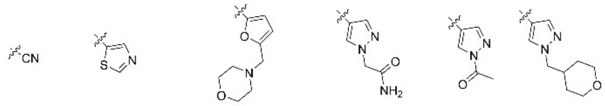

Jun12338 Jun12145 Jun12681 Jun121911 Jun12336 Jun12467

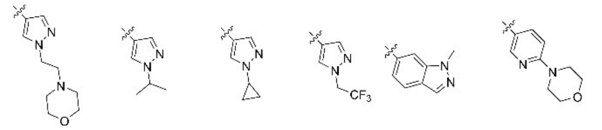

Jun12351 Jun12199 Jun12197 Jun12603 Jun12446 Jun12278

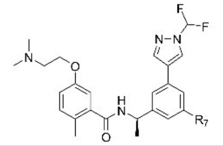

$R_7$ =

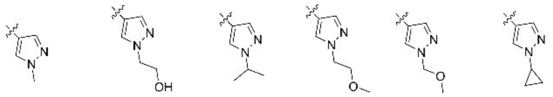

Jun12162 Jun12208 Jun12235 Jun12281 Jun12303 Jun12284

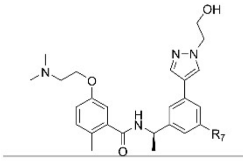

$R_7$ =

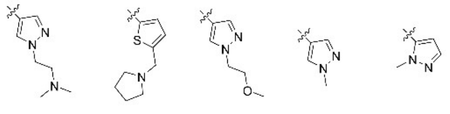

Jun12165 Jun12168 Jun12163 Jun12164 Jun12198

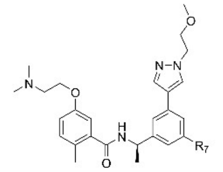

$R_7$ =

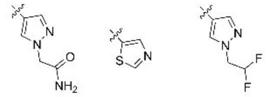

Jun12242 Jun12147 Jun12759

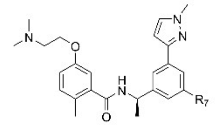

$R_7$ =

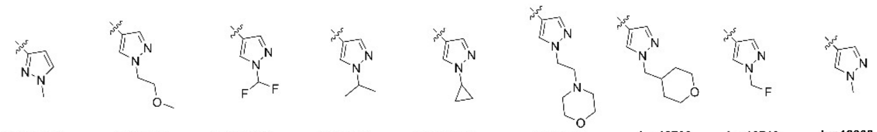

Jun12379 Jun12528 Jun12529 Jun12531 Jun12584 Jun12602 Jun12703 Jun12713 Jun12308

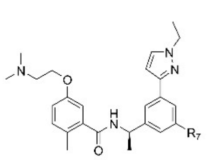

$R_7$ =

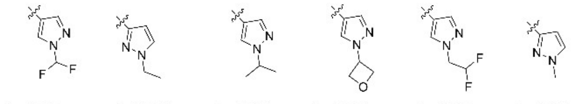

Jun12827 Jun12831 Jun12833 Jun12834 Jun12835 Jun12836

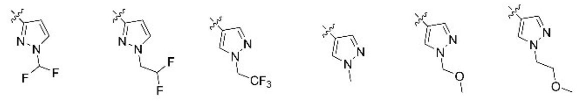

Jun12837 Jun1314 Jun12839 Jun12682 Jun12924 Jun12763

*Synthetic Procedure for Scheme 3*

**General procedures for the Suzuki coupling II.** To a solution of compound **B-4** (1 equiv) in dioxane/$H_2O$ (4:1) in a microwave reaction vial was added corresponding boronic acid or ester (1.2 equiv) and potassium carbonate (1.8 equiv). The mixture was purged with nitrogen for 5 min. Then Pd(dppf)$Cl_2$ (0.1 equiv) was added, and the solution was heated in the biotage microwave reactor at 130 °C for 90 min. The reaction mixture was diluted with EtOAc and extracted with aqueous $NaHCO_3$ solution and brine. The organic layer was separated, dried over anhydrous $Na_2SO_4$, filtered, and concentrated under vacuum to get intermediate **B-5**. To a solution of intermediate **B-5** (1 equiv) in dioxane/$H_2O$ (4:1) in a microwave reaction vial was added corresponding boronic acid or ester (1.2 equiv) and tripotassium phosphate (1.8 equiv). The mixture was purged with nitrogen for 5 min. Then XPhosPdG2 (0.1 equiv) was added, and the solution was heated in the biotage microwave reactor at 140 °C for 90 min. After the reaction was completed, the reaction mixture was diluted with EtOAc and washed with water and brine. The organic layer was separated, dried over anhydrous $Na_2SO_4$, filtered, and concentrated under vacuum to give a residue which was purified by Prep-HPLC to give the final product.

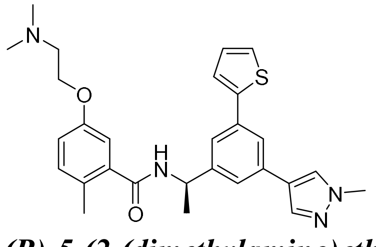

***(R)-5-(2-(dimethylamino)ethoxy)-2-methyl-N-(1-(3-(1-methyl-1H-pyrazol-4-yl)-5-(thiophen-2-yl)phenyl)ethyl)benzamide*** (**Jun1211**). White solid, 68% yield for two steps. $^1$H NMR (400 MHz, DMSO-$d_6$) δ 8.73 (d, J = 8.1 Hz, 1H), 8.17 (s, 1H), 7.87 (s, 1H), 7.67 (s, 1H), 7.51 (t, J = 4.7 Hz, 2H), 7.47 (s, 1H), 7.43 (s, 1H), 7.4-7.09 (m, 2H), 6.90 (s, 1H), 5.13-5.06 (m, 1H), 4.24 (t, J = 5.0 Hz, 2H), 3.82 (s, 3H), 3.45-3.42 (m, 2H), 2.86 (s, 1H), 2.79 (s, 6H), 2.17 (s, 2H), 1.42 (d, J = 7.0 Hz, 3H). $^{13}$C NMR (101 MHz, DMSO-$d_6$) δ 168.36, 158.79, 158.46, 155.68, 146.75, 143.95, 138.77, 136.65, 134.82, 133.87, 131.95, 128.87, 128.63, 128.17, 126.16, 124.37, 122.68, 122.02, 121.28, 120.73, 115.92, 114.02, 62.87, 55.91, 48.79, 43.27, 39.16, 22.99, 18.84. $C_{28}H_{32}N_4O_2S$, HRMS calculated for m/z $[M+H]^+$: 489.2324 (calculated), 489.2330 (found)

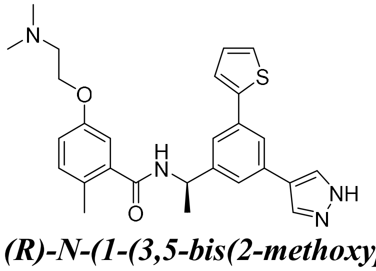

***(R)-N-(1-(3,5-bis(2-methoxypyrimidin-5-yl)phenyl)ethyl)-5-(2-(dimethylamino)ethoxy)-2-methylbenzamide*** (**Jun1244**). White solid, 66% yield for two steps. $^1$H NMR (400 MHz, $CDCl_3$) δ 8.65 (d, J = 2.3 Hz, 4H), 7.49 (s, 2H), 7.43 (s, 1H), 7.32 (s, 1H), 7.12 (d, J = 8.3 Hz, 1H), 6.99 (d, J = 2.8 Hz, 1H), 6.83 (dd, J = 8.4, 2.8 Hz, 1H), 6.63 (d, J = 7.7 Hz, 1H), 5.41-5.34 (m, 1H), 4.38 (dt, J = 6.8, 3.7 Hz, 2H), 3.47 (q, J = 4.8 Hz, 2H), 2.93 (s, 6H), 2.36 (s, 3H), 1.65 (d, J = 6.9 Hz, 3H). $^{13}$C NMR (101 MHz, $CDCl_3$) δ 168.70, 160.11, 155.61, 155.54, 155.07, 145.86, 137.02, 136.60, 132.39, 129.68, 122.78, 122.67, 122.29, 121.71, 116.05, 113.46, 62.99, 56.57, 49.42, 43.77, 43.39, 37.75, 37.55, 22.17, 19.03. $C_{27}H_{30}N_4O_2S$, HRMS calculated for m/z $[M+H]^+$: 475.2168 (calculated), 475.2173 (found)

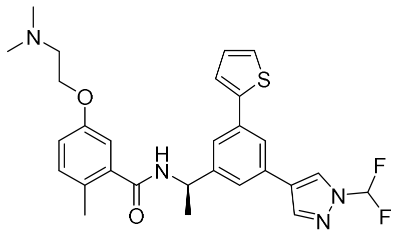

***(R)-N-(1-(3-(1-(difluoromethyl)-1H-pyrazol-4-yl)-5-(thiophen-2-yl)phenyl)ethyl)-5-(2-(dimethylamino)ethoxy)-2-methylbenzamide*** (**Jun1251**). White solid, 68% yield for two steps. $^{1}$H NMR (400 MHz, $CDCl_3$) δ 8.14 (s, 1H), 7.98 (s, 1H), 7.63 (s, 1H), 7.57 (s, 1H), 7.50 (s, 1H), 7.37 (d, J = 3.6 Hz, 1H), 7.32 (d, J = 5.0 Hz, 1H), 7.12-7.08 (m, 2H), 6.97 (d, J = 2.8 Hz, 1H), 6.82 (d, J = 8.4 Hz, 1H), 6.51 (d, J = 7.9 Hz, 1H), 5.38-5.31 (m, 1H), 4.37-4.34 (m, 2H), 3.44-3.42 (m, 2H), 2.91 (s, 6H), 2.36 (s, 3H), 1.66 (d, J = 6.9 Hz, 3H). $^{13}$C NMR (101 MHz, $CDCl_3$) δ 168.67, 155.03, 145.16, 143.66, 139.73, 137.25, 135.66, 132.35, 132.19, 129.66, 128.13, 125.48, 125.31, 123.75, 123.20, 123.08, 122.66, 115.95, 113.53, 111.08, 62.94, 56.63, 49.32, 43.82, 21.98, 18.94. $C_{28}H_{30}F_2N_4O_2S$, HRMS calculated for m/z $[M+H]^+$: 525.2136 (calculated), 525.2141 (found)

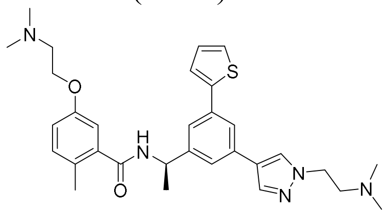

***(R)-5-(2-(dimethylamino)ethoxy)-N-(1-(3-(1-(2-(dimethylamino)ethyl)-1H-pyrazol-4-yl)-5-(thiophen-2-yl)phenyl)ethyl)-2-methylbenzamide*** (**Jun1252**). White solid, 65% yield for two steps. $^{1}$H NMR (400 MHz, DMSO-$d_6$) δ 8.82 (d, J = 8.1 Hz, 1H), 8.40 (s, 1H), 8.09 (s, 1H), 7.76 (s, 1H), 7.57 (dd, J = 13.5, 6.1 Hz, 4H), 7.19 (q, J = 6.0, 4.7 Hz, 2H), 6.98 (d, J = 8.3 Hz, 2H), 5.21-5.13 (m, 1H), 4.58 (t, J = 6.3 Hz, 2H), 4.32 (t, J = 5.0 Hz, 2H), 3.63 (t, J = 6.3 Hz, 2H), 3.51 (s, 2H), 2.86 (s, 6H), 2.83 (s, 6H), 2.23 (s, 3H), 1.49 (d, J = 7.0 Hz, 3H). $^{13}$C NMR (101 MHz, DMSO-$d_6$) δ 168.39, 158.74, 158.43, 155.70, 146.91, 143.86, 138.74, 137.72, 134.91, 133.40, 131.96, 128.89, 128.77, 128.14, 126.26, 124.39, 122.95, 122.52, 121.41, 120.81, 115.91, 114.03, 62.90, 56.11, 55.88, 48.84, 46.66, 43.26, 43.15, 23.00, 18.84. $C_{31}H_{39}N_5O_2S$, HRMS calculated for m/z $[M+H]^+$: 546.2903 (calculated), 546.2910(found)

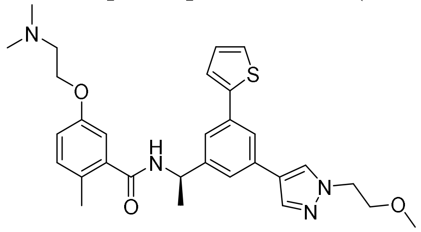

***(R)-5-(2-(dimethylamino)ethoxy)-N-(1-(3-(1-(2-methoxyethyl)-1H-pyrazol-4-yl)-5-(thiophen-2-yl)phenyl)ethyl)-2-methylbenzamide*** (**Jun1254**). White solid, 75% yield for two steps. $^{1}$H NMR (400 MHz, DMSO-$d_6$) δ 8.88 (d, J = 8.2 Hz, 1H), 8.33 (s, 1H), 8.05 (s, 1H), 7.84 (s, 1H), 7.69 – 7.62 (m, 3H), 7.58 (s, 1H), 7.30 – 7.24 (m, 2H), 7.06 (d, J = 8.1 Hz, 2H), 5.29-5.21 (m, 1H), 4.39 (dt, J = 10.1, 5.0 Hz, 4H), 3.88 (s, 2H), 3.81 (t, J = 5.2 Hz, 3H), 3.59 (s, 2H), 2.94 (s, 6H), 2.32 (s, 3H), 1.57 (d, J = 7.0 Hz, 3H). $^{13}$C NMR (101 MHz, DMSO-$d_6$) δ 168.35, 155.68, 146.74, 143.95, 138.79, 136.80, 134.82, 133.86, 131.95, 128.86, 128.42, 128.16, 126.15, 124.41, 122.70,

121.79, 121.30, 120.71, 115.92, 114.02, 70.91, 62.88, 58.46, 55.89, 51.83, 48.80, 43.27, 23.02, 18.84. $C_{30}H_{36}N_4O_3S$, HRMS calculated for m/z $[M+H]^+$: 533.2586 (calculated), 533.2594(found)

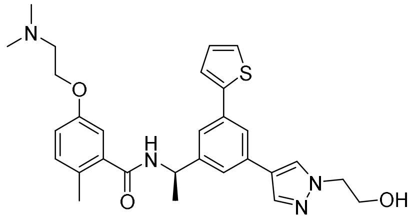

***(R)-5-(2-(dimethylamino)ethoxy)-N-(1-(3-(1-(2-hydroxyethyl)-1H-pyrazol-4-yl)-5-(thiophen-2-yl)phenyl)ethyl)-2-methylbenzamide*** (**Jun1257**). White solid, 72% yield for two steps. $^1$H NMR (400 MHz, DMSO-$d_6$) δ 9.73 (s, 1H), 8.79 (d, J = 8.2 Hz, 1H), 8.25 (s, 1H), 7.97 (s, 1H), 7.76 (s, 1H), 7.60-7.55 (m, 3H), 7.49 (s, 1H), 7.21-7.17 (m, 2H), 6.98 (d, J = 10.2 Hz, 2H), 5.21-5.13 (m, 1H), 4.31 (t, J = 5.0 Hz, 2H), 4.19 (t, J = 5.7 Hz, 2H), 3.79 (t, J = 5.6 Hz, 2H), 3.51 (s, 2H), 2.86 (s, 6H), 2.25 (s, 3H), 1.49 (d, J = 7.1 Hz, 3H). $^{13}$C NMR (101 MHz, DMSO-$d_6$) δ 168.98, 168.35, 155.68, 146.76, 143.93, 138.78, 137.07, 134.81, 133.82, 131.95, 129.47, 128.88, 128.18, 126.16, 124.41, 122.66, 122.03, 121.39, 120.72, 115.96, 113.99, 62.86, 55.91, 54.48, 48.78, 43.28, 23.00, 18.84. $C_{29}H_{34}N_4O_3S$, HRMS calculated for m/z $[M+H]^+$: 519.2430 (calculated), 519.2436 (found)

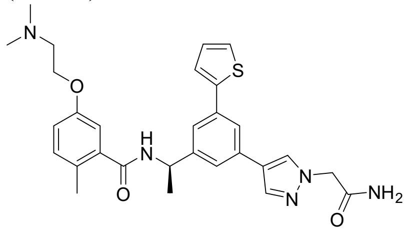

***(R)-N-(1-(3-(1-(2-amino-2-oxoethyl)-1H-pyrazol-4-yl)-5-(thiophen-2-yl)phenyl)ethyl)-5-(2-(dimethylamino)ethoxy)-2-methylbenzamide*** (**Jun121210**). White solid, 52% yield for two steps. $^1$H NMR (400 MHz, DMSO-$d_6$) δ 9.82 (s, 1H), 8.80 (d, J = 8.1 Hz, 1H), 8.24 (s, 1H), 7.98 (s, 1H), 7.77 (s, 1H), 7.59 (dd, J = 12.2, 3.7 Hz, 4H), 7.51 (s, 1H), 7.31 (s, 1H), 7.21-7.16 (m, 2H), 6.97 (s, 1H), 5.22-5.15 (m, 1H), 4.81 (s, 2H), 4.31 (t, J = 5.0 Hz, 2H), 3.51 (d, J = 5.0 Hz, 2H), 2.86 (s, 6H), 2.25 (s, 3H), 1.50 (d, J = 7.0 Hz, 3H). $^{13}$C NMR (101 MHz, DMSO-$d_6$) δ 168.98, 168.35, 155.68, 146.76, 143.93, 138.78, 137.07, 134.81, 133.82, 131.95, 129.47, 128.88, 128.18, 126.16, 124.41, 122.66, 122.03, 121.39, 120.72, 115.96, 113.99, 62.86, 55.91, 54.48, 48.78, 43.28, 23.00, 18.84. $C_{29}H_{33}N_5O_3S$, HRMS calculated for m/z $[M+H]^+$: 532.2382 (calculated), 532.2388 (found)

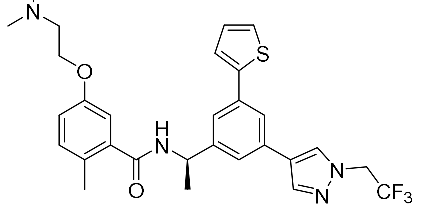

***(R)-5-(2-(dimethylamino)ethoxy)-2-methyl-N-(1-(3-(thiophen-2-yl)-5-(1-(2,2,2-trifluoroethyl)-1H-pyrazol-4-yl)phenyl)ethyl)benzamide*** (**Jun12149**). White solid, 62% yield for two steps. $^1$H NMR (400 MHz, DMSO-$d_6$) δ 8.74 (d, J = 8.1 Hz, 1H), 8.33 (s, 1H), 8.07 (s, 1H), 7.73 (d, J = 2.0 Hz, 1H), 7.55 (d, J = 3.6 Hz, 1H), 7.51 (d, J = 5.4 Hz, 2H), 7.49-7.46 (m, 1H), 7.12 (q, J = 6.2, 4.5 Hz, 2H), 6.91 (d, J = 7.4 Hz, 2H), 5.16-5.09 (m, 3H), 4.25 (t, J = 4.9 Hz, 2H), 3.44 (q, J = 4.4 Hz, 2H), 2.79 (s, 6H), 2.17 (s, 3H), 1.43 (d, J = 6.9 Hz, 3H). $^{13}$C NMR (101 MHz, DMSO-

$d_6$) δ 168.37, 158.81, 158.48, 155.69, 146.87, 143.79, 138.77, 138.63, 134.93, 133.03, 131.95, 129.73, 128.88, 128.16, 126.26, 124.54, 123.18, 122.98, 121.83, 121.01, 115.93, 114.03, 62.88, 55.91, 52.33, 51.99, 48.79, 43.27, 22.97, 18.82. $C_{29}H_{31}F_3N_4O_2S$, HRMS calculated for m/z $[M+H]^+$: 557.2198(calculated), 557.2205(found)

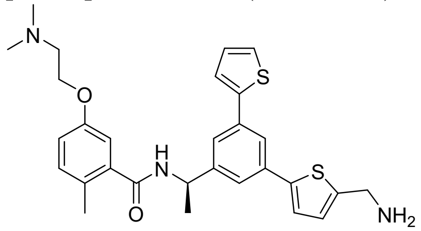


***(R)-N-(1-(3-(5-(aminomethyl)thiophen-2-yl)-5-(thiophen-2-yl)phenyl)ethyl)-5-(2-(dimethylamino)ethoxy)-2-methylbenzamide*** (**Jun12129**). White solid, 66% yield for two steps. $^1H$ NMR (400 MHz, DMSO-$d_6$) δ 8.89 (d, J = 7.9 Hz, 1H), 8.47 (s, 3H), 7.77 (s, 1H), 7.65 (s, 1H), 7.62-7.57 (m, 4H), 7.28 (d, J = 3.7 Hz, 1H), 7.20 (d, J = 4.4 Hz, 2H), 6.98 (s, 1H), 5.23-5.16 (m, 1H), 4.34-4.30 (m, 4H), 3.52 (s, 2H), 2.87 (s, 6H), 2.24 (s, 3H), 1.51 (d, J = 6.9 Hz, 3H). $^{13}C$ NMR (101 MHz, DMSO-$d_6$) δ 168.47, 159.06, 158.74, 155.71, 147.46, 144.40, 143.22, 138.63, 135.64, 135.20, 134.66, 131.96, 130.79, 129.04, 128.16, 126.66, 124.83, 124.73, 123.20, 122.95, 120.99, 115.96, 114.01, 62.91, 55.85, 48.76, 43.23, 37.52, 22.83, 18.86. $C_{29}H_{33}N_3O_2S_2$, HRMS calculated for m/z $[M+H]^+$: 520.2092 (calculated), 520.2099 (found)

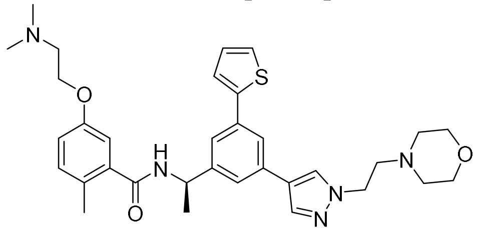


***(R)-5-(2-(dimethylamino)ethoxy)-2-methyl-N-(1-(3-(1-(2-morpholinoethyl)-1H-pyrazol-4-yl)-5-(thiophen-2-yl)phenyl)ethyl)benzamide*** (**Jun121910**). White solid, 78% yield for two steps. $^1H$ NMR (400 MHz, DMSO-$d_6$) δ 8.81 (d, J = 8.1 Hz, 1H), 8.38 (s, 1H), 8.09 (s, 1H), 7.76 (s, 1H), 7.61-7.53 (m, 4H), 7.23-7.16 (m, 2H), 6.98 (d, J = 7.7 Hz, 2H), 5.21-5.14 (m, 1H), 4.62 (t, J = 6.4 Hz, 2H), 4.32 (t, J = 5.0 Hz, 2H), 3.69 (t, J = 6.5 Hz, 2H), 3.52 (d, J = 5.3 Hz, 2H), 2.87 (s, 6H), 2.24 (s, 3H), 1.50 (d, J = 7.0 Hz, 3H). $^{13}C$ NMR (101 MHz, DMSO-$d_6$) δ 168.39, 159.00, 158.66, 155.70, 146.90, 143.87, 138.75, 137.71, 134.92, 133.41, 131.95, 128.88, 128.68, 128.17, 126.24, 124.38, 122.96, 122.53, 121.44, 120.84, 118.43, 115.92, 114.05, 63.79, 62.92, 55.91, 55.46, 51.94, 48.85, 46.21, 43.28, 22.96, 18.83. $C_{33}H_{41}N_5O_3S$, HRMS calculated for m/z $[M+H]^+$: 588.3008 (calculated), 588.3015 (found)

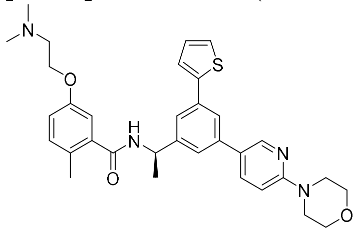


***(R)-5-(2-(dimethylamino)ethoxy)-2-methyl-N-(1-(3-(6-morpholinopyridin-3-yl)-5-(thiophen-2-yl)phenyl)ethyl)benzamide*** (**Jun12511**). White solid, 74% yield for two steps. $^1H$ NMR (400 MHz, $CDCl_3$) δ 8.45 (s, 1H), 8.11 (dd, J = 9.3, 2.2 Hz, 1H), 7.67 (s, 1H), 7.58 (s, 1H), 7.51 (s, 1H), 7.38 (d, J = 3.6 Hz, 1H), 7.33 (d, J = 5.0 Hz, 1H), 7.13-7.10 (m, 2H), 7.03 (d, J = 9.3 Hz, 1H), 6.97 (d, J = 2.7 Hz, 1H), 6.82 (dd, J = 8.3, 2.7 Hz, 1H), 6.75 (d, J = 7.6 Hz, 1H), 5.37-5.33

(m, 1H), 4.35 (q, J = 6.8, 5.9 Hz, 2H), 3.90 (t, J = 4.8 Hz, 4H), 3.73 (t, J = 5.0 Hz, 4H), 3.47 (q, J = 4.2 Hz, 2H), 2.92 (s, 6H), 2.36 (s, 3H), 1.65 (d, J = 7.0 Hz, 3H). $^{13}$C NMR (101 MHz, DMSO-$d_6$) δ 168.43, 155.69, 146.93, 143.20, 142.81, 138.61, 138.43, 134.82, 131.98, 131.57, 129.04, 128.18, 126.66, 125.83, 124.94, 124.10, 123.64, 115.96, 114.03, 106.50, 62.89, 55.89, 48.73, 43.26, 38.00, 22.94, 18.84. $C_{33}H_{38}N_4O_3S$, HRMS calculated for m/z $[M+H]^+$: 571.2743 (calculated), 571.2750 (found)

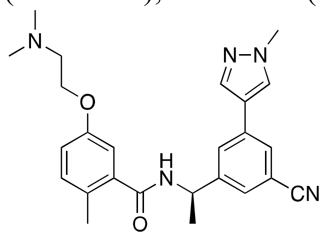

***(R)-N-(1-(3-cyano-5-(1-methyl-1H-pyrazol-4-yl)phenyl)ethyl)-5-(2-(dimethylamino)ethoxy)-2-methylbenzamide*** (**Jun12338**). White solid, 60% yield for two steps. $^1$H NMR (400 MHz, $CDCl_3$) δ 8.79 (d, J = 7.9 Hz, 1H), 8.27 (s, 1H), 7.98 (s, 1H), 7.96 (s, 2H), 7.64 (s, 1H), 7.20 (d, J = 8.4 Hz, 1H), 6.99 (d, J = 6.1 Hz, 2H), 5.21-5.13 (m, 1H), 4.32 (t, J = 4.9 Hz, 2H), 3.88 (s, 3H), 3.52 (q, J = 4.8 Hz, 2H), 2.87 (d, J = 3.7 Hz, 6H), 2.22 (s, 3H), 1.47 (d, J = 7.0 Hz, 3H). $^{13}$C NMR (101 MHz, DMSO-$d_6$) δ 168.47, 159.01, 158.66, 155.70, 147.47, 138.39, 136.86, 134.59, 131.99, 129.09, 128.21, 127.96, 127.24, 127.07, 120.58, 119.40, 115.98, 114.11, 112.38, 62.87, 55.90, 48.59, 43.26, 39.24, 22.70, 18.77. $C_{25}H_{29}N_5O_2$, HRMS calculated for m/z $[M+H]^+$: 432.2399 (calculated), 432.2406(found)

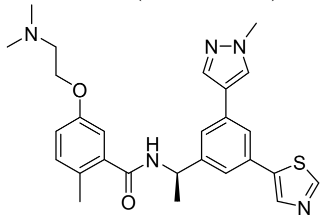

***(R)-5-(2-(dimethylamino)ethoxy)-2-methyl-N-(1-(3-(1-methyl-1H-pyrazol-4-yl)-5-(thiazol-5-yl)phenyl)ethyl)benzamide*** (**Jun12145**). White solid, 55% yield for two steps. $^1$H NMR (400 MHz, DMSO-$d_6$) δ 9.11 (s, 1H), 8.78 (d, J = 8.2 Hz, 1H), 8.38 (s, 1H), 8.25 (s, 1H), 7.96 (s, 1H), 7.81 (d, J = 1.8 Hz, 1H), 7.61 (s, 1H), 7.51 (d, J = 1.8 Hz, 1H), 7.20 (d, J = 8.1 Hz, 1H), 6.98 (d, J = 8.3 Hz, 2H), 5.22-5.14 (m, 1H), 4.32 (t, J = 4.9 Hz, 2H), 3.89 (s, 3H), 3.52 (s, 2H), 2.86 (s, 6H), 2.24 (s, 3H), 1.49 (d, J = 7.0 Hz, 3H). $^{13}$C NMR (101 MHz, DMSO-$d_6$) δ 168.38, 158.86, 158.52, 155.70, 154.07, 147.00, 140.03, 139.20, 138.74, 136.72, 134.10, 131.96, 131.80, 128.72, 128.16, 123.40, 122.53, 121.81, 121.73, 115.95, 114.03, 62.89, 55.93, 48.78, 43.29, 39.18, 23.03, 18.81. $C_{27}H_{31}N_5O_2S$, HRMS calculated for m/z $[M+H]^+$: 490.2277(calculated), 490.2283(found)

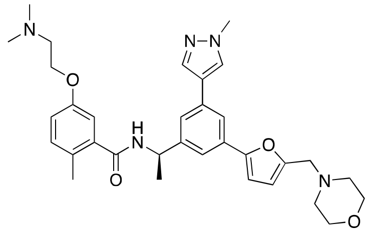

***(R)-5-(2-(dimethylamino)ethoxy)-2-methyl-N-(1-(3-(1-methyl-1H-pyrazol-4-yl)-5-(5-(morpholinomethyl)furan-2-yl)phenyl)ethyl)benzamide*** (**Jun12681**). White solid, 65% yield for two steps. $^1$H NMR (400 MHz, DMSO-$d_6$) δ 8.80 (d, J = 8.1 Hz, 1H), 8.22 (s, 1H), 7.92 (s, 1H), 7.74 (s, 1H), 7.60 (s, 2H), 7.49 (s, 1H), 7.34 (d, J = 3.7 Hz, 1H), 7.19 (d, J = 8.2 Hz, 1H), 6.98 (d,

J = 6.3 Hz, 2H), 5.21-5.14 (p, 1H), 4.62 (s, 2H), 4.32 (t, J = 5.0 Hz, 2H), 3.89 (s, 3H), 2.87 (s, 6H), 2.23 (s, 3H), 1.50 (d, J = 7.0 Hz, 3H). $^{13}$C NMR (101 MHz, DMSO-d$_6$) δ 168.39, 159.15, 158.81, 155.70, 146.99, 146.90, 138.69, 136.60, 134.14, 134.03, 131.93, 128.62, 128.15, 124.64, 123.21, 121.91, 121.54, 120.80, 115.90, 114.04, 63.84, 62.88, 55.85, 53.62, 50.92, 48.82, 43.22, 39.17, 22.85, 18.83. $C_{33}H_{41}N_5O_4$, HRMS calculated for m/z $[M+H]^+$: 572.3637(calculated), 572.3644 (found)

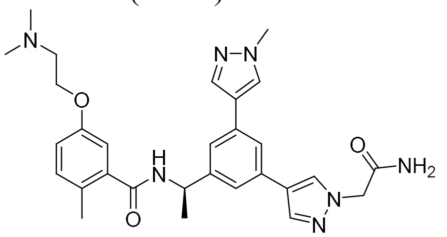

***(R)-N-(1-(3-(1-(2-amino-2-oxoethyl)-1H-pyrazol-4-yl)-5-(1-methyl-1H-pyrazol-4-yl)phenyl)ethyl)-5-(2-(dimethylamino)ethoxy)-2-methylbenzamide*** (**Jun121911**). White solid, 45% yield for two steps. $^1$H NMR (400 MHz, DMSO-d$_6$) δ 8.71 (d, J = 8.3 Hz, 1H), 8.17 (d, J = 5.9 Hz, 2H), 7.92 (d, J = 10.1 Hz, 2H), 7.69 (s, 1H), 7.58 (s, 1H), 7.45 (d, J = 9.5 Hz, 2H), 7.30 (s, 1H), 7.19 (d, J = 8.3 Hz, 1H), 6.97 (d, J = 10.6 Hz, 2H), 5.20-5.13 (m, 1H), 4.80 (s, 2H), 4.30 (t, J = 4.8 Hz, 2H), 3.88 (s, 3H), 3.50 (q, J = 5.0 Hz, 2H), 2.86 (s, 6H), 2.24 (s, 3H), 1.49 (d, J = 6.9 Hz, 3H). $^{13}$C NMR (101 MHz, DMSO-d$_6$) δ 169.00, 168.27, 155.67, 146.20, 138.85, 136.92, 136.53, 133.52, 133.47, 131.92, 129.16, 128.38, 128.17, 122.48, 114.01, 62.83, 55.92, 54.48, 48.86, 43.27, 39.14, 23.12, 18.81. $C_{29}H_{35}N_7O_3$, HRMS calculated for m/z $[M+H]^+$: 530.2880 (calculated), 530.2886 (found)

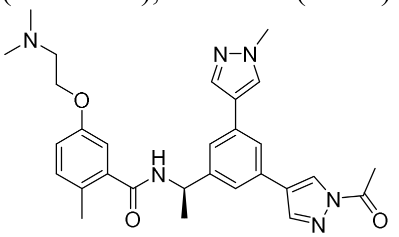

***(R)-N-(1-(3-(1-acetyl-1H-pyrazol-4-yl)-5-(1-methyl-1H-pyrazol-4-yl)phenyl)ethyl)-5-(2-(dimethylamino)ethoxy)-2-methylbenzamide*** (**Jun12336**). White solid, 40% yield for two steps. $^1$H NMR (400 MHz, DMSO-d$_6$) δ 8.94 (s, 1H), 8.42 (s, 1H), 8.21 (s, 1H), 8.08 (s, 1H), 7.95 (s, 1H), 7.90 (d, J = 8.2 Hz, 2H), 7.65 (s, 1H), 7.55 (s, 1H), 7.20 (d, J = 8.2 Hz, 1H), 6.98 (d, J = 8.2 Hz, 2H), 5.22-5.14 (m, 1H), 4.32-4.29 (m, 3H), 3.89 (s, 6H), 3.50 (q, J = 5.0 Hz, 3H), 2.86 (d, J = 4.4 Hz, 9H), 2.69 (s, 3H), 2.24 (d, J = 3.3 Hz, 4H), 1.49 (d, J = 4.2 Hz, 3H). $^{13}$C NMR (101 MHz, DMSO-d$_6$) δ 168.31, 158.88, 158.52, 155.69, 146.44, 142.61, 138.87, 138.82, 136.65, 133.79, 131.95, 131.32, 128.54, 128.15, 126.37, 125.01, 122.52, 122.22, 121.70, 117.62, 114.02, 62.83, 55.94, 48.88, 43.29, 23.30, 23.18, 21.93, 18.78. $C_{29}H_{34}N_6O_3$, HRMS calculated for m/z $[M+H]^+$: 515.2770 (calculated), 515.2778 (found)

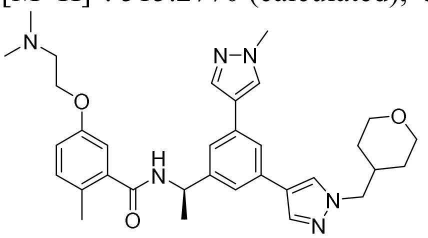

***(R)-5-(2-(dimethylamino)ethoxy)-2-methyl-N-(1-(3-(1-methyl-1H-pyrazol-4-yl)-5-(1-((tetrahydro-2H-pyran-4-yl)methyl)-1H-pyrazol-4-yl)phenyl)ethyl)benzamide*** (**Jun12467**).

White solid, 60% yield for two steps. $^{1}$H NMR (400 MHz, DMSO-$d_6$) δ 8.71 (d, J = 8.2 Hz, 1H), 8.18 (d, J = 13.1 Hz, 2H), 7.91 (d, J = 11.3 Hz, 2H), 7.68 (d, J = 1.9 Hz, 1H), 7.44 (s, 2H), 7.19 (d, J = 8.2 Hz, 1H), 6.97 (d, J = 8.1 Hz, 2H), 5.19-5.12 (m, 1H), 4.31 (t, J = 4.9 Hz, 2H), 4.04 (d, J = 7.0 Hz, 2H), 3.88 (s, 3H), 3.84 (dd, J = 11.2, 4.0 Hz, 2H), 3.51 (q, J = 4.8 Hz, 2H), 3.27 (t, J = 11.5 Hz, 2H), 2.87 (s, 6H), 2.24 (s, 3H), 1.49 (d, J = 7.0 Hz, 3H), 1.43 (dd, J = 12.9, 3.7 Hz, 2H), 1.26 (qd, J = 12.2, 4.5 Hz, 2H). $^{13}$C NMR (101 MHz, DMSO-$d_6$) δ 1 168.28, 159.01, 158.65, 155.68, 146.19, 138.84, 136.63, 136.51, 133.51, 131.92, 128.33, 128.15, 128.12, 122.49, 122.12, 121.02, 120.94, 120.53, 115.85, 114.05, 67.00, 62.85, 57.41, 55.90, 48.92, 43.25, 39.13, 36.19, 30.47, 23.09, 18.80. $C_{33}H_{42}N_6O_3$, HRMS calculated for m/z $[M+H]^+$: 571.3397(calculated), 571.3405(found)

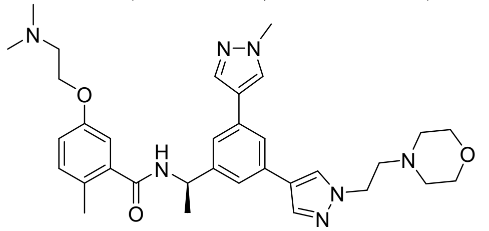

***(R)-5-(2-(dimethylamino)ethoxy)-2-methyl-N-(1-(3-(1-methyl-1H-pyrazol-4-yl)-5-(1-(2-morpholinoethyl)-1H-pyrazol-4-yl)phenyl)ethyl)benzamide*** (**Jun12351**). White solid, 82% yield for two steps. $^{1}$H NMR (400 MHz, DMSO-$d_6$) δ 8.70 (d, J = 8.2 Hz, 1H), 8.30 (s, 1H), 8.13 (s, 1H), 8.04 (s, 1H), 7.87 (s, 1H), 7.68 (d, J = 1.7 Hz, 1H), 7.45 (d, J = 11.8 Hz, 2H), 7.19 (d, J = 8.3 Hz, 1H), 6.98 (dd, J = 11.7, 3.5 Hz, 2H), 5.19-5.11 (m, J = 7.1 Hz, 1H), 4.58 (t, J = 6.5 Hz, 2H), 4.30 (t, J = 5.0 Hz, 2H), 3.89 (s, 4H), 3.65 (s, 2H), 3.50 (d, J = 5.0 Hz, 2H), 2.85 (s, 6H), 2.22 (s, 3H), 1.48 (d, J = 7.0 Hz, 3H).. $^{13}$C NMR (101 MHz, DMSO-$d_6$) δ 168.31, 155.69, 146.33, 138.79, 137.60, 136.49, 133.61, 133.04, 131.93, 128.41, 128.32, 128.14, 122.97, 122.44, 121.25, 121.12, 120.60, 115.85, 114.06, 63.79, 62.87, 55.88, 55.44, 51.92, 48.93, 46.19, 43.25, 39.16, 23.07, 18.81. $C_{33}H_{43}N_7O_3$, MS calculated for m/z $[M+H]^+$: 586.3506 (calculated), 586.3512 (found).

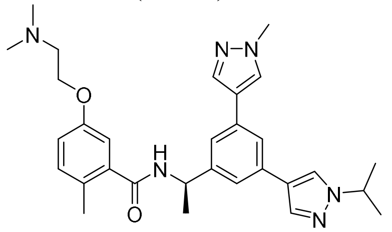

***(R)-5-(2-(dimethylamino)ethoxy)-N-(1-(3-(1-isopropyl-1H-pyrazol-4-yl)-5-(1-methyl-1H-pyrazol-4-yl)phenyl)ethyl)-2-methylbenzamide*** (**Jun12199**). White solid, 77% yield for two steps. $^{1}$H NMR (400 MHz, DMSO-$d_6$) δ 8.69 (d, J = 8.3 Hz, 1H), 8.25 (s, 1H), 8.16 (s, 1H), 7.89 (s, 2H), 7.68 (s, 1H), 7.44 (d, J = 9.3 Hz, 2H), 7.20 (d, J = 8.2 Hz, 1H), 6.97 (d, J = 8.6 Hz, 2H), 5.20-5.12 (m, 1H), 4.55-4.49 (m 1H), 4.31 (t, J = 5.0 Hz, 2H), 3.89 (s, 3H), 3.51 (q, J = 4.6 Hz, 2H), 2.86 (s, 6H), 2.24 (s, 3H), 1.48 (t, J = 7.2 Hz, 9H). $^{13}$C NMR (101 MHz, DMSO-$d_6$) δ 168.27, 155.69, 146.11, 138.88, 136.52, 136.00, 133.72, 133.48, 131.92, 128.31, 128.17, 125.13, 122.54, 121.99, 121.00, 120.87, 120.53, 115.87, 114.09, 62.87, 55.94, 53.61, 48.92, 43.29, 39.13, 23.16, 23.12, 18.80. $C_{30}H_{38}N_6O_2$, HRMS calculated for m/z $[M+H]^+$: 515.3134(calculated), 515.3144(found)

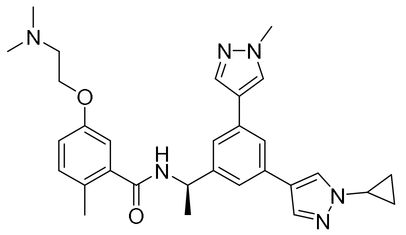

***(R)-N-(1-(3-(1-cyclopropyl-1H-pyrazol-4-yl)-5-(1-methyl-1H-pyrazol-4-yl)phenyl)ethyl)-5-(2-(dimethylamino)ethoxy)-2-methylbenzamide*** (**Jun12197**). White solid, 73% yield for two steps. $^{1}$H NMR (400 MHz, DMSO-d$_6$) δ 8.68 (d, J = 8.3 Hz, 1H), 8.26 (s, 1H), 8.15 (s, 1H), 7.88 (d, J = 3.6 Hz, 2H), 7.68 (s, 1H), 7.43 (d, J = 6.2 Hz, 2H), 7.20 (d, J = 8.3 Hz, 1H), 6.97 (d, J = 11.6 Hz, 2H), 5.19-5.11 (m, 1H), 4.31 (t, J = 4.9 Hz, 2H), 3.88 (s, 3H), 3.75 (tt, J = 7.4, 3.9 Hz, 1H), 3.50 (q, J = 5.0 Hz, 2H), 2.86 (s, 6H), 2.24 (s, 3H), 1.48 (d, J = 7.1 Hz, 3H), 1.08 (q, J = 4.3 Hz, 2H), 1.00 (q, J = 7.3, 6.0 Hz, 2H). $^{13}$C NMR (101 MHz, DMSO-d$_6$) δ 168.26, 155.68, 146.12, 138.88, 136.53, 133.50, 133.40, 131.92, 128.32, 128.17, 127.44, 122.50, 122.22, 121.00, 120.58, 115.88, 114.09, 62.86, 55.96, 48.89, 43.31, 39.14, 33.24, 23.13, 18.80, 6.73, 6.71. $C_{30}H_{36}N_6O_2$, HRMS calculated for m/z [M+H]$^+$: 513.2978(calculated), 513.2983 (found)

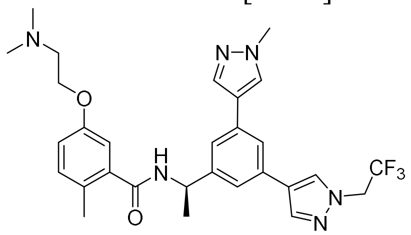

***(R)-5-(2-(dimethylamino)ethoxy)-2-methyl-N-(1-(3-(1-methyl-1H-pyrazol-4-yl)-5-(1-(2,2,2-trifluoroethyl)-1H-pyrazol-4-yl)phenyl)ethyl)benzamide*** (**Jun12603**). White solid, 69% yield for two steps. $^{1}$H NMR (400 MHz, DMSO-d$_6$) δ 8.72 (d, J = 8.3 Hz, 1H), 8.32 (s, 1H), 8.19 (s, 1H), 8.09 (s, 1H), 7.92 (s, 1H), 7.72 (s, 1H), 7.48 (s, 2H), 7.19 (d, J = 8.1 Hz, 1H), 6.98 (d, J = 7.4 Hz, 2H), 5.18 (m, 3H), 4.31 (t, J = 4.9 Hz, 2H), 3.89 (s, 3H), 3.51 (q, J = 4.8 Hz, 2H), 2.87 (s, 6H), 2.24 (s, 3H), 1.50 (d, J = 6.9 Hz, 3H). $^{13}$C NMR (101 MHz, DMSO-d$_6$) δ 168.31, 155.69, 146.33, 138.82, 138.51, 136.57, 133.65, 132.69, 131.92, 129.39, 128.43, 128.15, 123.64, 122.39, 121.51, 121.23, 120.78, 115.87, 114.04, 62.85, 55.90, 48.89, 43.25, 39.13, 23.07, 18.78. $C_{29}H_{33}F_3N_6O_2$, MS calculated for m/z [M+H]$^+$: 555.2695 (calculated), 555.2700 (found).

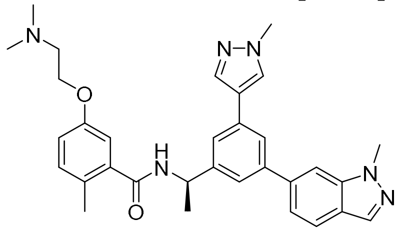

***(R)-5-(2-(dimethylamino)ethoxy)-2-methyl-N-(1-(3-(1-methyl-1H-indazol-6-yl)-5-(1-methyl-1H-pyrazol-4-yl)phenyl)ethyl)benzamide*** (**Jun12446**). White solid, 72% yield for two steps. $^{1}$H NMR (400 MHz, DMSO-d$_6$) δ 8.79 (d, J = 8.2 Hz, 1H), 8.25 (s, 1H), 8.08 (s, 1H), 7.96 (d, J = 3.5 Hz, 2H), 7.86-7.84 (m, 2H), 7.64 (d, J = 7.6 Hz, 2H), 7.52 (d, J = 8.5 Hz, 1H), 7.20 (d, J = 8.3 Hz, 1H), 6.98 (d, J = 7.0 Hz, 2H), 5.28-5.21 (m, 1H), 4.30 (t, J = 5.0 Hz, 2H), 4.13 (s, 3H), 3.90 (s, 3H), 3.50 (q, J = 4.5 Hz, 2H), 2.85 (s, 6H), 2.25 (s, 3H), 1.54 (d, J = 7.0 Hz, 3H), 1.16 (t, J = 7.3 Hz, 3H). $^{13}$C NMR (101 MHz, DMSO-d$_6$) δ 168.33, 155.69, 146.39, 141.61, 140.73, 138.84, 136.69, 133.71, 132.71, 131.94, 128.57, 128.15, 123.30, 123.22, 122.78, 122.55, 122.42, 121.54, 120.60, 115.88, 114.07, 107.88, 62.84, 55.91, 49.00, 43.27, 41.81, 39.17, 35.89, 23.23,

18.85, 11.46. $C_{32}H_{36}N_6O_2$, MS calculated for m/z $[M+H]^+$: 537.2978 (calculated), 537.2985 (found).

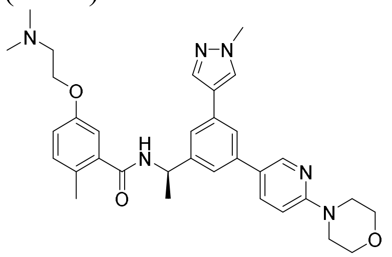

***(R)-5-(2-(dimethylamino)ethoxy)-2-methyl-N-(1-(3-(1-methyl-1H-pyrazol-4-yl)-5-(6-morpholinopyridin-3-yl)phenyl)ethyl)benzamide*** (**Jun12278**). White solid, 81% yield for two steps. $^1H$ NMR (400 MHz, DMSO-$d_6$) δ 8.74 (d, J = 8.3 Hz, 1H), 8.51 (s, 1H), 8.22 (s, 1H), 8.08 (d, J = 9.0 Hz, 1H), 7.94 (s, 1H), 7.73 (d, J = 1.7 Hz, 1H), 7.57 (s, 1H), 7.52 (s, 1H), 7.20 (d, J = 8.1 Hz, 1H), 7.11 (d, J = 9.1 Hz, 1H), 6.98 (d, J = 8.4 Hz, 2H), 5.25-5.17 (m, 1H), 4.31 (t, J = 5.0 Hz, 2H), 3.89 (s, 3H), 3.75 (t, J = 4.8 Hz, 4H), 3.57 (t, J = 4.9 Hz, 4H), 3.51 (s, 2H), 2.86 (s, 6H), 2.24 (s, 3H), 1.50 (d, J = 6.9 Hz, 3H). $^{13}C$ NMR (101 MHz, DMSO-$d_6$) δ 168.32, 158.98, 158.62, 155.69, 146.52, 138.82, 137.75, 136.67, 133.82, 131.93, 128.55, 128.15, 125.83, 122.37, 122.12, 121.66, 121.31, 117.75, 115.89, 114.07, 66.24, 62.87, 55.93, 48.96, 45.84, 43.28, 39.15, 23.23, 18.81. $C_{33}H_{40}N_6O_3$, HRMS calculated for m/z $[M+H]^+$: 569.3240 (calculated), 569.3248(found)

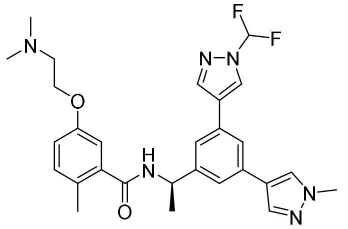

***(R)-N-(1-(3-(1-(difluoromethyl)-1H-pyrazol-4-yl)-5-(1-methyl-1H-pyrazol-4-yl)phenyl)ethyl)-5-(2-(dimethylamino)ethoxy)-2-methylbenzamide*** (**Jun12162**). White solid, 77% yield for two steps. $^1H$ NMR (400 MHz, DMSO-$d_6$) δ 8.75 (s, 1H), 8.69 (d, J = 8.0 Hz, 1H), 8.31 (s, 1H), 8.18 (s, 1H), 7.92 (s, 1H), 7.85-7.81 (m, 1H), 7.55 (d, J = 13.8 Hz, 2H), 7.19 (d, J = 8.3 Hz, 1H), 6.97 (d, J = 7.0 Hz, 2H), 5.21-5.14 (m, 1H), 4.31 (t, J = 4.9 Hz, 2H), 3.89 (s, 3H), 3.50 (s, 2H), 2.86 (s, 6H), 2.23 (s, 3H), 1.49 (d, J = 7.1 Hz, 3H). $^{13}C$ NMR (101 MHz, DMSO-$d_6$) δ 168.31, 158.82, 158.49, 155.69, 146.37, 140.44, 138.80, 136.59, 133.74, 131.93, 131.75, 128.44, 128.16, 125.66, 125.10, 122.29, 122.03, 121.64, 121.20, 115.90, 114.06, 111.13, 62.86, 55.92, 48.90, 43.27, 39.17, 23.13, 18.79. $C_{28}H_{32}F_2N_6O_2$, HRMS calculated for m/z $[M+H]^+$: 523.2633(calculated), 523.2639(found)

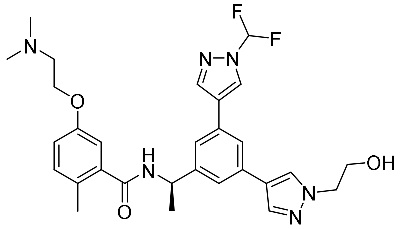

***(R)-N-(1-(3-(1-(difluoromethyl)-1H-pyrazol-4-yl)-5-(1-(2-hydroxyethyl)-1H-pyrazol-4-yl)phenyl)ethyl)-5-(2-(dimethylamino)ethoxy)-2-methylbenzamide*** (**Jun12208**). White solid, 73% yield for two steps. $^1H$ NMR (400 MHz, DMSO-$d_6$) δ 8.77 (s, 1H), 8.71 (d, J = 8.6 Hz, 1H), 8.32 (s, 1H), 8.20 (s, 1H), 7.95 (s, 1H), 7.86 (d, J = 11.2 Hz, 2H), 7.56 (d, J = 5.6 Hz, 2H), 7.20 (d, J = 8.1 Hz, 1H), 6.98 (d, J = 7.0 Hz, 2H), 5.22-5.15 (m, 2H), 4.31 (t, J = 4.9 Hz, 2H), 4.18 (t,

J = 5.5 Hz, 2H), 3.78 (t, J = 5.5 Hz, 2H), 3.50 (q, J = 4.8 Hz, 2H), 2.86 (s, 6H), 2.24 (s, 3H), 1.50 (d, J = 7.0 Hz, 3H). $^{13}$C NMR (101 MHz, DMSO-d$_6$) δ 168.30, 155.69, 146.36, 140.45, 138.82, 136.60, 133.85, 131.93, 131.73, 128.24, 128.15, 125.67, 125.11, 121.96, 121.90, 121.60, 121.13, 115.91, 114.04, 62.85, 60.49, 55.91, 54.85, 48.90, 43.26, 23.17, 18.80. $C_{29}H_{34}F_2N_6O_3$, HRMS calculated for m/z [M+H]$^+$: 553.2739(calculated), 553.2745(found)

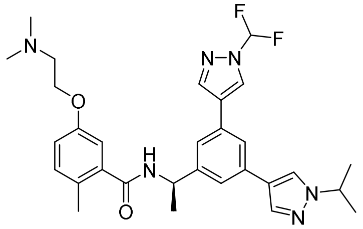

***(R)-N-(1-(3-(1-(difluoromethyl)-1H-pyrazol-4-yl)-5-(1-isopropyl-1H-pyrazol-4-yl)phenyl)ethyl)-5-(2-(dimethylamino)ethoxy)-2-methylbenzamide*** (**Jun12235**). White solid, 78% yield for two steps. $^1$H NMR (400 MHz, DMSO-d$_6$) δ 8.75 (s, 1H), 8.70 (d, J = 8.5 Hz, 1H), 8.30 (d, J = 13.1 Hz, 2H), 7.93 (s, 1H), 7.84 (s, 1H), 7.56 (s, 2H), 7.20 (d, J = 8.2 Hz, 1H), 6.98 (d, J = 6.9 Hz, 2H), 5.22-5.15 (m, 1H), 4.56-4.51 (m, 1H), 4.31 (t, J = 4.9 Hz, 2H), 3.50 (q, J = 4.9 Hz, 2H), 2.86 (s, 6H), 2.24 (s, 3H), 1.50 (d, J = 7.1 Hz, 3H), 1.47 (d, J = 6.8 Hz, 6H). $^{13}$C NMR (101 MHz, DMSO-d$_6$) δ 168.30, 155.69, 146.31, 140.44, 138.82, 136.08, 133.93, 131.93, 131.70, 128.15, 125.62, 125.24, 125.14, 122.06, 121.77, 121.54, 121.16, 117.80, 115.88, 114.89, 114.06, 111.14, 62.84, 55.92, 53.65, 48.92, 43.26, 23.16, 18.79. $C_{30}H_{36}F_2N_6O_2$, HRMS calculated for m/z [M+H]$^+$: 551.2946(calculated), 551.2950(found)

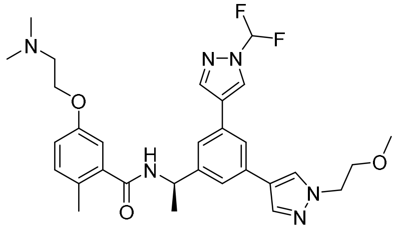

***(R)-N-(1-(3-(1-(difluoromethyl)-1H-pyrazol-4-yl)-5-(1-(2-methoxyethyl)-1H-pyrazol-4-yl)phenyl)ethyl)-5-(2-(dimethylamino)ethoxy)-2-methylbenzamide*** (**Jun12281**). White solid, 81% yield for two steps. $^1$H NMR (400 MHz, DMSO-d$_6$) δ 8.85 (s, 1H), 8.79 (d, *J* = 7.8 Hz, 1H), 8.40 (s, 1H), 8.28 (s, 1H), 8.04 (s, 1H), 7.92 (s, 1H), 7.64 (d, *J* = 10.2 Hz, 2H), 7.27 (d, *J* = 8.2 Hz, 1H), 7.05 (d, *J* = 7.2 Hz, 2H), 5.29-5.23 (m, 1H), 4.72 (s, 4H), 4.38 (q, *J* = 5.0 Hz, 3H), 3.80 (t, *J* = 5.2 Hz, 2H), 3.59 (t, *J* = 4.8 Hz, 2H), 2.94 (s, 6H), 2.32 (s, 3H), 1.58 (d, *J* = 7.1 Hz, 3H). $^{13}$C NMR (101 MHz, DMSO-d$_6$) δ 168.32, 155.70, 146.40, 140.46, 138.80, 136.76, 133.74, 131.92, 131.75, 128.25, 128.15, 125.69, 125.11, 122.06, 121.66, 121.16, 115.89, 114.05, 70.96, 62.86, 58.47, 55.87, 51.90, 48.92, 43.23, 23.16, 18.79. $C_{30}H_{36}F_2N_6O_3$, HRMS calculated for m/z [M+H]$^+$: 567.2895(calculated), 567.2901(found)

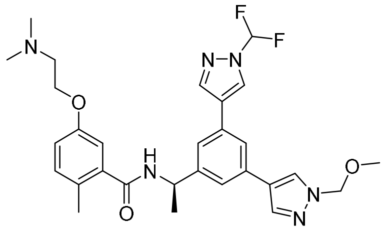

***(R)-N-(1-(3-(1-(difluoromethyl)-1H-pyrazol-4-yl)-5-(1-(methoxymethyl)-1H-pyrazol-4-yl)phenyl)ethyl)-5-(2-(dimethylamino)ethoxy)-2-methylbenzamide*** (**Jun12303**). White solid, 65% yield for two steps. $^1$H NMR (400 MHz, DMSO-d$_6$) δ 8.70 (s, 1H), 8.63 (d, J = 8.0 Hz, 1H),

8.34 (s, 1H), 8.25 (s, 1H), 8.00 (s, 1H), 7.82 (d, J = 8.8 Hz, 2H), 7.53 (s, 2H), 7.12 (d, J = 8.8 Hz, 1H), 6.91 (d, J = 4.9 Hz, 2H), 5.35 (s, 2H), 5.12 (m, 1H), 4.24 (t, J = 4.9 Hz, 2H), 3.43 (d, J = 4.8 Hz, 2H), 3.21 (s, 3H), 2.79 (s, 6H), 2.16 (s, 3H), 1.43 (d, J = 6.9 Hz, 3H). $^{13}$C NMR (101 MHz, DMSO-$d_6$) δ 168.32, 155.69, 146.43, 140.44, 138.79, 137.87, 133.31, 131.93, 131.82, 128.49, 128.15, 125.69, 125.05, 123.19, 122.27, 121.98, 121.42, 118.35, 115.91, 114.05, 111.14, 81.88, 62.85, 56.61, 55.91, 48.92, 43.26, 23.16, 18.79. $C_{29}H_{34}F_2N_6O_3$, MS calculated for m/z $[M+H]^+$: 553.2739 (calculated), 553.2745 (found).

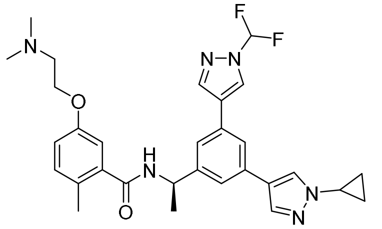

***(R)-N-(1-(3-(1-cyclopropyl-1H-pyrazol-4-yl)-5-(1-(difluoromethyl)-1H-pyrazol-4-yl)phenyl)ethyl)-5-(2-(dimethylamino)ethoxy)-2-methylbenzamide*** (**Jun12284**). White solid, 75% yield for two steps. $^1$H NMR (400 MHz, $CDCl_3$) δ 8.17 (s, 1H), 8.01 (d, J = 2.9 Hz, 2H), 7.84 (s, 1H), 7.53 (s, 2H), 7.50 (s, 1H), 7.14 (d, J = 8.4 Hz, 1H), 7.06 (s, 1H), 6.89-6.81 (m, 2H), 5.38-5.31 (m, 1H), 4.69-4.63 (m, 1H), 4.40 (s, 2H), 3.47 (s, 2H), 2.96 (s, 6H), 2.36 (s, 3H), 1.68 (d, J = 7.0 Hz, 3H), 1.60 (d, J = 6.7 Hz, 4H). $^{13}$C NMR (101 MHz, DMSO-$d_6$) δ 168.33, 155.69, 146.37, 140.43, 138.77, 136.61, 133.61, 131.93, 131.72, 128.15, 127.59, 125.63, 125.11, 122.07, 122.00, 121.64, 121.20, 115.89, 114.05, 62.84, 55.89, 48.93, 43.24, 33.24, 23.17, 18.78, 6.74, 6.72. $C_{30}H_{34}F_2N_6O_2$, HRMS calculated for m/z $[M+H]^+$: 549.2789(calculated), 549.2794(found)

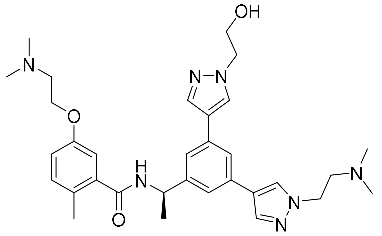

***(R)-5-(2-(dimethylamino)ethoxy)-N-(1-(3-(1-(2-(dimethylamino)ethyl)-1H-pyrazol-4-yl)-5-(1-(2-hydroxyethyl)-1H-pyrazol-4-yl)phenyl)ethyl)-2-methylbenzamide*** (**Jun12165**). White solid, 66% yield for two steps. $^1$H NMR (400 MHz, DMSO-$d_6$) δ 8.73 (d, J = 8.2 Hz, 1H), 8.33 (s, 1H), 8.16 (s, 1H), 8.06 (s, 1H), 7.91 (s, 1H), 7.70 (s, 1H), 7.46 (d, J = 17.9 Hz, 2H), 7.19 (d, J = 8.1 Hz, 1H), 6.97 (d, J = 8.4 Hz, 2H), 5.19-5.12 (m, 1H), 4.58 (t, J = 6.2 Hz, 2H), 4.31 (t, J = 4.9 Hz, 2H), 4.18 (t, J = 5.5 Hz, 2H), 3.78 (t, J = 5.5 Hz, 2H), 3.63 (t, J = 6.3 Hz, 2H), 3.51 (t, J = 5.1 Hz, 2H), 2.85 (d, J = 11.1 Hz, 12H), 2.23 (s, 3H), 1.49 (d, J = 6.9 Hz, 3H). $^{13}$C NMR (101 MHz, DMSO-$d_6$) δ 168.30, 158.94, 158.61, 155.69, 146.34, 138.81, 137.62, 136.49, 133.73, 133.02, 131.92, 128.49, 128.14, 128.10, 122.99, 122.05, 121.23, 121.08, 120.53, 118.54, 115.87, 115.59, 114.04, 62.87, 60.49, 56.11, 55.89, 54.82, 48.93, 46.63, 43.25, 43.14, 23.10, 18.81. $C_{32}H_{43}N_7O_3$, HRMS calculated for m/z $[M+H]^+$: 574.3506(calculated), 574.3511(found)

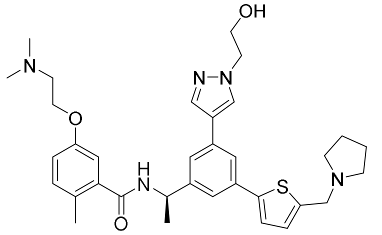

***(R)-5-(2-(dimethylamino)ethoxy)-N-(1-(3-(1-(2-hydroxyethyl)-1H-pyrazol-4-yl)-5-(5-(pyrrolidin-1-ylmethyl)thiophen-2-yl)phenyl)ethyl)-2-methylbenzamide*** (**Jun12168**). White solid, 60% yield for two steps. $^{1}$H NMR (400 MHz, DMSO-$d_6$) δ 10.03 (s, 1H), 8.80 (d, J = 8.1 Hz, 1H), 8.23 (s, 1H), 7.95 (s, 1H), 7.75 (s, 1H), 7.61 (s, 1H), 7.59 (d, J = 3.6 Hz, 1H), 7.48 (s, 1H), 7.35 (d, J = 3.7 Hz, 1H), 7.19 (d, J = 8.1 Hz, 1H), 6.98 (d, J = 7.6 Hz, 2H), 5.21-5.14 (m, 1H), 4.63 (s, 2H), 4.32 (t, J = 4.9 Hz, 2H), 4.19 (t, J = 5.5 Hz, 2H), 3.78 (t, J = 5.5 Hz, 2H), 3.51 (t, J = 5.5 Hz, 3H), 3.46 (s, 1H), 2.86 (s, 6H), 2.23 (s, 3H), 2.03 (d, J = 9.9 Hz, 2H), 1.91-1.88 (m, 2H), 1.50 (d, J = 7.1 Hz, 3H). $^{13}$C NMR (101 MHz, DMSO-$d_6$) δ 168.38, 155.70, 146.88, 146.43, 138.73, 136.61, 134.14, 132.98, 131.94, 128.40, 128.14, 124.57, 123.10, 121.54, 121.50, 115.92, 114.04, 62.88, 60.46, 55.88, 54.83, 53.14, 51.34, 48.82, 43.25, 23.07, 18.84. $C_{34}H_{43}N_5O_3S$, HRMS calculated for m/z $[M+H]^+$: 602.3165(calculated), 602.3171 (found)

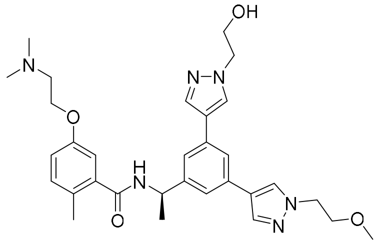

***(R)-5-(2-(dimethylamino)ethoxy)-N-(1-(3-(1-(2-hydroxyethyl)-1H-pyrazol-4-yl)-5-(1-(2-methoxyethyl)-1H-pyrazol-4-yl)phenyl)ethyl)-2-methylbenzamide*** (**Jun12163**). White solid, 76% yield for two steps. $^{1}$H NMR (400 MHz, DMSO-$d_6$) δ 9.83 (s, 1H), 8.72 (d, J = 8.3 Hz, 1H), 8.19 (s, 2H), 7.93 (d, J = 2.4 Hz, 2H), 7.69 (s, 1H), 7.44 (d, J = 3.4 Hz, 2H), 7.19 (d, J = 8.3 Hz, 1H), 6.98 (d, J = 11.1 Hz, 2H), 5.20-5.12 (m, 1H), 4.30 (q, J = 5.8 Hz, 4H), 4.18 (t, J = 5.4 Hz, 2H), 3.78 (t, J = 5.5 Hz, 2H), 3.73 (t, J = 5.2 Hz, 2H), 3.50 (d, J = 5.0 Hz, 2H), 3.25 (s, 3H), 2.86 (s, 6H), 2.24 (s, 3H), 1.49 (d, J = 7.0 Hz, 3H). $^{13}$C NMR (101 MHz, DMSO-$d_6$) δ 168.27, 158.86, 158.52, 155.68, 146.17, 138.87, 136.68, 136.54, 133.62, 133.49, 131.92, 128.14, 122.27, 122.11, 120.93, 120.44, 115.88, 114.04, 70.96, 62.85, 60.50, 58.47, 55.92, 54.80, 51.84, 48.90, 43.27, 23.15, 18.81. $C_{31}H_{40}N_6O_4$, HRMS calculated for m/z $[M+H]^+$: 561.3189(calculated), 561.3195 (found)

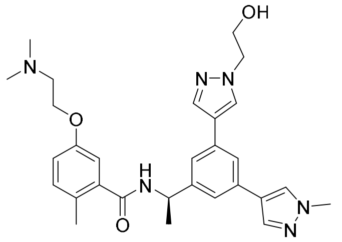

***(R)-5-(2-(dimethylamino)ethoxy)-N-(1-(3-(1-(2-hydroxyethyl)-1H-pyrazol-4-yl)-5-(1-methyl-1H-pyrazol-4-yl)phenyl)ethyl)-2-methylbenzamide*** (**Jun12164**). White solid, 74% yield for two steps. $^{1}$H NMR (400 MHz, DMSO-$d_6$) δ 9.82 (s, 1H), 8.71 (d, J = 8.3 Hz, 1H), 8.17 (s, 2H), 7.91 (d, J = 7.1 Hz, 2H), 7.68 (s, 1H), 7.44 (d, J = 6.7 Hz, 2H), 7.19 (d, J = 8.2 Hz, 1H), 6.99-6.96 (m, 2H), 5.19-5.12 (m, 1H), 4.31 (t, J = 4.9 Hz, 2H), 4.18 (t, J = 5.5 Hz, 2H), 3.88 (s, 3H), 3.78 (t, J = 5.5 Hz, 2H), 3.50 (d, J = 4.9 Hz, 2H), 2.86 (s, 6H), 2.24 (s, 3H), 1.48 (d, J = 7.0 Hz, 3H). $^{13}$C NMR (101 MHz, DMSO-$d_6$) δ 168.27, 158.83, 158.49, 155.68, 146.17, 138.86, 136.52, 133.62, 133.50, 131.92, 128.35, 128.16, 128.11, 122.50, 122.11, 120.92, 120.47, 115.88, 114.04, 62.85, 60.50, 55.92, 54.80, 48.89, 43.27, 39.14, 23.13, 18.81. $C_{29}H_{36}N_6O_3$, HRMS calculated for m/z $[M+H]^+$: 517.2927(calculated), 517.2933(found)

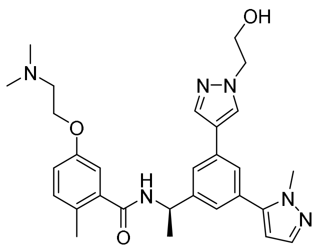

***(R)-5-(2-(dimethylamino)ethoxy)-N-(1-(3-(1-(2-hydroxyethyl)-1H-pyrazol-4-yl)-5-(1-methyl-1H-pyrazol-5-yl)phenyl)ethyl)-2-methylbenzamide*** (**Jun12198**). White solid, 77% yield for two steps. $^{1}$H NMR (400 MHz, DMSO-$d_6$) δ 8.77 (d, J = 8.1 Hz, 1H), 8.24 (s, 1H), 7.96 (s, 1H), 7.67 (s, 1H), 7.61 (s, 1H), 7.50 (s, 1H), 7.36 (s, 1H), 7.19 (d, J = 8.3 Hz, 1H), 6.97 (d, J = 10.8 Hz, 2H), 6.43 (d, J = 2.1 Hz, 1H), 5.25-5.18 (m, 1H), 4.30 (t, J = 4.8 Hz, 2H), 4.17 (t, J = 5.5 Hz, 2H), 3.89 (s, 3H), 3.78 (t, J = 5.5 Hz, 2H), 3.51 (q, J = 4.8 Hz, 2H), 2.86 (s, 6H), 2.23 (s, 3H), 1.50 (d, J = 7.1 Hz, 3H). $^{13}$C NMR (101 MHz, DMSO-$d_6$) δ 168.34, 155.68, 146.36, 143.28, 138.72, 138.34, 136.71, 133.75, 131.95, 131.30, 128.42, 128.19, 124.09, 123.76, 123.31, 121.58, 115.94, 114.05, 106.27, 62.88, 60.46, 55.94, 54.80, 48.80, 43.30, 37.98, 23.05, 18.80. $C_{29}H_{36}N_6O_3$, HRMS calculated for m/z $[M+H]^+$: 517.2927(calculated), 517.2934 (found)

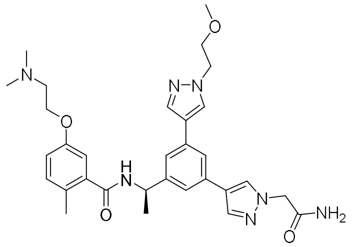

***(R)-N-(1-(3-(1-(2-amino-2-oxoethyl)-1H-pyrazol-4-yl)-5-(1-(2-methoxyethyl)-1H-pyrazol-4-yl)phenyl)ethyl)-5-(2-(dimethylamino)ethoxy)-2-methylbenzamide*** (**Jun12242**). White solid, 58% yield for two steps. $^{1}$H NMR (400 MHz, DMSO-$d_6$) δ 8.72 (d, J = 8.2 Hz, 1H), 8.18 (d, J = 8.1 Hz, 2H), 7.94 (s, 2H), 7.70 (s, 1H), 7.58 (s, 1H), 7.45 (d, J = 6.3 Hz, 2H), 7.31 (s, 1H), 7.19 (d, J = 8.3 Hz, 1H), 6.97 (d, J = 8.4 Hz, 2H), 5.19-5.13 (m, 1H), 4.80 (s, 2H), 4.30 (q, J = 5.4 Hz, 4H), 3.72 (t, J = 5.2 Hz, 2H), 3.50 (q, J = 4.9 Hz, 2H), 3.25 (s, 3H), 2.85 (s, 6H), 2.24 (s, 3H), 1.49 (d, J = 7.0 Hz, 3H). $^{13}$C NMR (101 MHz, DMSO-$d_6$) 169.01, 168.27, 155.67, 146.20, 138.85, 136.94, 136.69, 133.50, 133.47, 131.92, 129.19, 128.16, 122.48, 122.24, 121.07, 120.92, 120.47, 117.94, 115.91, 114.00, 70.95, 62.82, 58.46, 55.91, 54.48, 51.85, 48.88, 43.26, 23.13, 18.81. $C_{31}H_{39}N_7O_4$, HRMS calculated for m/z $[M+H]^+$: 574.3142(calculated), 571.3146(found)

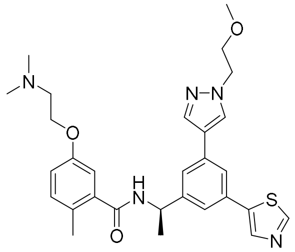

***(R)-5-(2-(dimethylamino)ethoxy)-N-(1-(3-(1-(2-methoxyethyl)-1H-pyrazol-4-yl)-5-(thiazol-5-yl)phenyl)ethyl)-2-methylbenzamide*** (**Jun12147**). White solid, 54% yield for two steps. $^{1}$H NMR (400 MHz, DMSO-$d_6$) δ 9.19 (s, 1H), 8.86 (d, J = 8.1 Hz, 1H), 8.47 (s, 1H), 8.35 (s, 1H), 8.07 (s, 1H), 7.89 (s, 1H), 7.70 (s, 1H), 7.59 (s, 1H), 7.28 (d, J = 8.1 Hz, 1H), 7.05 (s, 1H), 5.30-5.22 (m, 1H), 4.38 (q, J = 5.2 Hz, 4H), 3.90 (s, 3H), 3.81 (t, J = 5.2 Hz, 3H), 3.58 (q, J = 4.6 Hz, 2H), 2.93 (s, 6H), 2.32 (s, 3H), 1.57 (d, J = 7.0 Hz, 3H). $^{13}$C NMR (101 MHz, DMSO-$d_6$) δ

168.36, 155.69, 154.06, 146.97, 140.06, 139.18, 138.77, 136.87, 134.09, 131.95, 131.80, 128.51, 128.15, 123.41, 122.54, 121.71, 121.58, 115.95, 114.02, 70.90, 62.86, 58.46, 55.94, 51.86, 48.77, 43.29, 23.04, 18.81. $C_{29}H_{35}N_5O_3S$, HRMS calculated for m/z $[M+H]^+$: 534.2539(calculated), 534.2545(found)

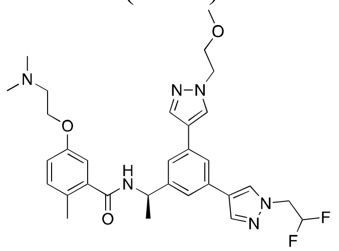

***(R)-N-(1-(3-(1-(2,2-difluoroethyl)-1H-pyrazol-4-yl)-5-(1-(2-methoxyethyl)-1H-pyrazol-4-yl)phenyl)ethyl)-5-(2-(dimethylamino)ethoxy)-2-methylbenzamide*** (**Jun12759**). White solid, 77% yield for two steps. $^1$H NMR (400 MHz, DMSO-$d_6$) δ 8.79 (d, J = 8.3 Hz, 1H), 8.34 (s, 1H), 8.27 (s, 1H), 8.11 (s, 1H), 8.01 (s, 1H), 7.79 (s, 1H), 7.54 (s, 2H), 7.27 (d, J = 8.2 Hz, 1H), 7.05 (d, J = 9.4 Hz, 2H), 5.28-5.21 (m, 1H), 4.75 (t, J = 13.4 Hz, 2H), 4.38 (q, J = 5.4 Hz, 4H), 3.94 (s, 3H), 3.81 (t, J = 5.2 Hz, 2H), 3.58 (q, J = 4.6 Hz, 2H), 2.93 (s, 6H), 2.32 (s, 3H), 1.57 (d, J = 7.0 Hz, 3H). $^{13}$C NMR (101 MHz, DMSO-$d_6$) δ 168.30, 158.87, 158.53, 155.68, 146.26, 138.85, 137.81, 136.70, 133.59, 133.02, 131.92, 129.07, 128.18, 128.15, 123.13, 122.18, 121.31, 121.14, 120.64, 115.88, 114.47, 114.04, 70.95, 62.84, 58.46, 55.93, 51.85, 48.90, 43.27, 23.09, 18.79. $C_{31}H_{38}F_2N_6O_3$, HRMS calculated for m/z $[M+H]^+$: 581.3052(calculated), 581.3060(found)

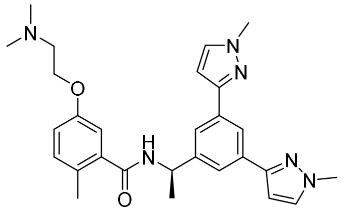

***(R)-N-(1-(3,5-bis(1-methyl-1H-pyrazol-3-yl)phenyl)ethyl)-5-(2-(dimethylamino)ethoxy)-2-methylbenzamide*** (**Jun12379**). White solid, 78% yield. $^1$H NMR (400 MHz, DMSO-$d_6$) δ 8.84 (d, J = 8.2 Hz, 1H), 8.06 (s, 1H), 7.76 (s, 4H), 7.19 (d, J = 8.1 Hz, 1H), 6.97 (d, J = 8.1 Hz, 2H), 6.73 (d, J = 1.9 Hz, 2H), 5.22-5.15 (m, 1H), 4.31 (t, J = 5.1 Hz, 2H), 3.90 (s, 6H), 3.88 (s, 1H), 3.51 (s, 2H), 2.86 (s, 6H), 2.24 (s, 3H), 1.50 (d, J = 7.0 Hz, 3H). $^{13}$C NMR (101 MHz, DMSO-$d_6$) δ 168.29, 159.01, 158.66, 155.65, 150.50, 150.46, 146.01, 138.80, 138.76, 134.24, 132.78, 131.91, 128.28, 122.32, 120.54, 115.84, 114.94, 114.09, 103.00, 62.84, 55.89, 48.91, 43.25, 39.11, 39.08, 22.87, 18.83. $C_{28}H_{34}N_6O_2$, HRMS calculated for m/z $[M+H]^+$: 487.2821(calculated), 487.2830 (found)

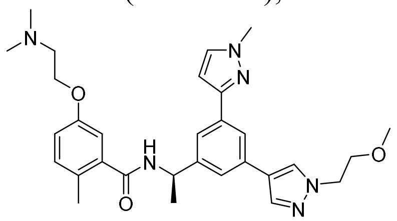

***(R)-5-(2-(dimethylamino)ethoxy)-N-(1-(3-(1-(2-methoxyethyl)-1H-pyrazol-4-yl)-5-(1-methyl-1H-pyrazol-3-yl)phenyl)ethyl)-2-methylbenzamide*** (**Jun12528**). White solid, 68% yield for two steps. $^1$H NMR (400 MHz, DMSO-$d_6$) δ 8.80 (d, J = 8.3 Hz, 1H), 8.21 (s, 1H), 7.93 (s, 1H), 7.83 (s, 1H), 7.76 (s, 1H), 7.70 (s, 1H), 7.53 (s, 1H), 7.19 (d, J = 8.1 Hz, 1H), 6.98 (d, J = 7.6 Hz, 2H), 6.76 (s, 1H), 5.21-5.14 (m, 1H), 4.31 (q, J = 5.8 Hz, 4H), 3.90 (s, 3H), 3.73 (t, J = 5.3 Hz, 2H),

3.54-3.47 (m, 2H), 3.25 (s, 3H), 2.86 (s, 6H), 2.24 (s, 3H), 1.50 (d, J = 7.0 Hz, 3H). $^{13}$C NMR (101 MHz, DMSO-d$_6$) δ 168.28, 155.67, 150.52, 146.15, 138.83, 136.61, 134.49, 133.29, 132.72, 131.91, 128.22, 128.17, 122.43, 122.20, 120.90, 120.50, 115.84, 114.07, 103.11, 70.94, 62.85, 58.44, 55.88, 51.78, 48.90, 43.24, 39.10, 22.99, 18.82. $C_{30}H_{38}N_6O_3$, HRMS calculated for m/z [M+H]$^+$: 531.3083(calculated), 531.3087 (found)

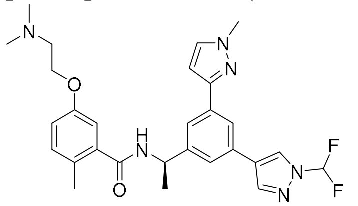

***(R)-N-(1-(3-(1-(difluoromethyl)-1H-pyrazol-4-yl)-5-(1-methyl-1H-pyrazol-3-yl)phenyl)ethyl)-5-(2-(dimethylamino)ethoxy)-2-methylbenzamide*** (**Jun12529**). White solid, 77% yield for two steps. $^1$H NMR (400 MHz, DMSO-d$_6$) δ 8.73 (d, J = 5.6 Hz, 2H), 8.25 (s, 1H), 7.88 (s, 1H), 7.79 (s, 1H), 7.74 (s, 1H), 7.70 (s, 1H), 7.59 (s, 1H), 7.12 (d, J = 9.0 Hz, 1H), 6.91 (d, J = 6.2 Hz, 2H), 6.72 (d, J = 2.2 Hz, 1H), 5.16-5.09 (m, 1H), 4.24 (t, J = 5.0 Hz, 2H), 3.83 (s, 3H), 3.44 (q, J = 4.0 Hz, 2H), 2.79 (s, 6H), 2.17 (s, 3H), 1.44 (d, J = 7.0 Hz, 3H).$^{13}$C NMR (101 MHz, DMSO-d$_6$) δ 168.33, 155.67, 150.31, 146.36, 140.49, 138.77, 134.70, 132.78, 131.92, 131.57, 128.25, 125.91, 124.97, 123.17, 121.89, 121.26, 115.87, 114.09, 103.26, 62.86, 55.88, 48.90, 43.24, 39.12, 22.98, 18.82. $C_{28}H_{32}F_2N_6O_2$, HRMS calculated for m/z [M+H]$^+$: 523.2633(calculated), 523.2641 (found)

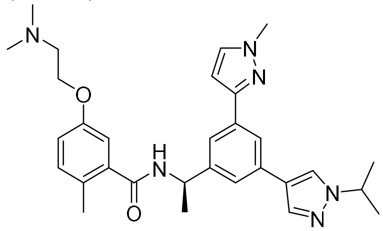

***(R)-5-(2-(dimethylamino)ethoxy)-N-(1-(3-(1-isopropyl-1H-pyrazol-4-yl)-5-(1-methyl-1H-pyrazol-3-yl)phenyl)ethyl)-2-methylbenzamide*** (**Jun12531**). White solid, 76% yield for two steps. $^1$H NMR (400 MHz, DMSO-d$_6$) δ 8.72 (d, J = 8.2 Hz, 1H), 8.21 (s, 1H), 7.82 (s, 1H), 7.77 (s, 1H), 7.68 (s, 1H), 7.62 (s, 1H), 7.47 (s, 1H), 7.12 (d, J = 8.2 Hz, 1H), 6.90 (d, J = 7.3 Hz, 2H), 6.68 (d, J = 2.2 Hz, 1H), 5.14-5.06 (m, 1H), 4.49-4.42 (m, 1H), 4.24 (t, J = 5.0 Hz, 2H), 3.82 (s, 3H), 2.79 (s, 6H), 2.17 (s, 3H), 1.42 (d, J = 6.9 Hz, 3H), 1.40 (d, J = 6.7 Hz, 6H). $^{13}$C NMR (101 MHz, DMSO-d$_6$) δ 168.28, 155.67, 150.57, 146.09, 138.82, 135.95, 134.46, 133.50, 132.70, 131.91, 128.23, 125.23, 122.46, 121.94, 120.79, 120.52, 115.84, 114.09, 103.11, 62.86, 55.88, 53.62, 48.93, 43.24, 39.10, 23.15, 23.00, 18.83. $C_{30}H_{38}N_6O_2$, HRMS calculated for m/z [M+H]$^+$: 515.3134(calculated), 515.3141 (found)

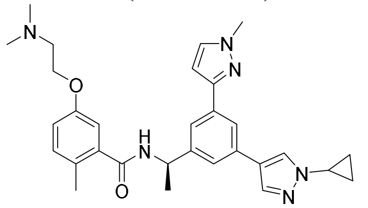

***(R)-N-(1-(3-(1-cyclopropyl-1H-pyrazol-4-yl)-5-(1-methyl-1H-pyrazol-3-yl)phenyl)ethyl)-5-(2-(dimethylamino)ethoxy)-2-methylbenzamide*** (**Jun12584**). White solid, 74% yield for two steps. $^1$H NMR (400 MHz, DMSO-d$_6$) δ 8.78 (d, J = 8.2 Hz, 1H), 8.30 (s, 1H), 7.89 (s, 1H), 7.83 (s, 1H), 7.75 (s, 1H), 7.70 (s, 1H), 7.54 (s, 1H), 7.19 (d, J = 8.1 Hz, 1H), 6.98 (d, J = 7.8 Hz, 2H),

6.75 (s, 1H), 5.21-5.13 (m, 1H), 4.31 (t, J = 5.0 Hz, 2H), 3.89 (s, 3H), 3.76 (tt, J = 7.6, 3.9 Hz, 1H), 3.51 (q, J = 4.7 Hz, 2H), 2.87 (s, 6H), 2.24 (s, 3H), 1.49 (d, J = 7.0 Hz, 3H), 1.12-1.08 (m, 2H), 1.00 (q, J = 7.2 Hz, 2H). $^{13}$C NMR (101 MHz, DMSO-$d_6$) δ 168.29, 155.66, 150.53, 146.10, 138.82, 136.50, 134.48, 133.19, 132.70, 131.91, 128.24, 127.47, 122.50, 122.16, 120.88, 120.60, 115.84, 114.09, 103.13, 62.85, 55.90, 48.91, 43.25, 39.09, 33.27, 23.00, 18.82, 6.73, 6.71, 6.63. $C_{30}H_{36}N_6O_2$, HRMS calculated for m/z $[M+H]^+$: 513.2978(calculated), 513.2984(found)

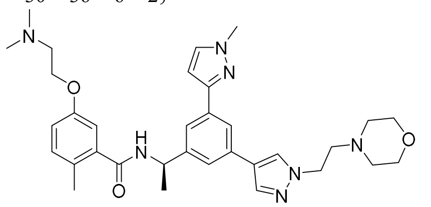

***(R)-5-(2-(dimethylamino)ethoxy)-2-methyl-N-(1-(3-(1-methyl-1H-pyrazol-3-yl)-5-(1-(2-morpholinoethyl)-1H-pyrazol-4-yl)phenyl)ethyl)benzamide*** (**Jun12602**). White solid, 81% yield for two steps. $^1$H NMR (400 MHz, $CDCl_3$) δ 8.01 (s, 1H), 7.89 (s, 2H), 7.76 (s, 1H), 7.49 (s, 1H), 7.44 (d, J = 6.7 Hz, 2H), 7.13 (d, J = 8.3 Hz, 1H), 7.03 (d, J = 2.7 Hz, 1H), 6.85 – 6.80 (m, 2H), 5.34-5.27 (m , 1H), 4.75 (t, J = 5.9 Hz, 2H), 4.39 (d, J = 9.2 Hz, 2H), 4.00 (s, 3H), 3.95 (t, J = 4.9 Hz, 5H), 3.69 (t, J = 5.6 Hz, 3H), 3.48 (q, J = 8.7, 7.3 Hz, 2H), 3.12 (s, 3H), 2.95 (s, 6H), 2.37 (s, 3H), 1.65 (d, J = 7.0 Hz, 3H). $^{13}$C NMR (101 MHz, DMSO-$d_6$) δ 168.32, 159.14, 158.81, 155.67, 150.44, 146.30, 138.77, 137.52, 134.57, 132.83, 132.77, 131.92, 128.47, 128.22, 122.94, 122.65, 121.06, 120.56, 118.31, 115.83, 115.38, 114.08, 103.04, 63.74, 62.88, 55.85, 55.40, 51.87, 48.94, 46.15, 43.22, 39.10, 22.96, 18.82. $C_{33}H_{43}N_7O_3$, MS calculated for m/z $[M+H]^+$: 586.3506 (calculated), 586.3510 (found).

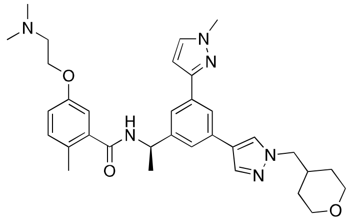

***(R)-5-(2-(dimethylamino)ethoxy)-2-methyl-N-(1-(3-(1-methyl-1H-pyrazol-3-yl)-5-(1-((tetrahydro-2H-pyran-4-yl)methyl)-1H-pyrazol-4-yl)phenyl)ethyl)benzamide*** (**Jun12703**). White solid, 70% yield for two steps. $^1$H NMR (400 MHz, DMSO-$d_6$) δ 8.74 (d, J = 8.2 Hz, 1H), 8.31 (s, 1H), 8.25 (s, 1H), 7.96 (d, J = 12.5 Hz, 1H), 7.75 (s, 1H), 7.50 (d, J = 9.0 Hz, 1H), 7.26 (d, J = 8.3 Hz, 1H), 7.07-6.99 (m, 2H), 5.25-5.18 (m, 1H), 4.58 (p, J = 6.7 Hz, 1H), 4.37 (t, J = 5.0 Hz, 1H), 4.10 (d, J = 7.1 Hz, 1H), 3.90 (dd, J = 11.6, 3.1 Hz, 1H), 3.56 (d, J = 5.0 Hz, 1H), 2.92 (d, J = 4.0 Hz, 3H), 2.30 (s, 2H), 2.15 (ddd, J = 11.2, 7.5, 3.9 Hz, 1H), 1.53 (q, J = 6.8, 6.3 Hz, 7H), 1.48 (d, J = 3.6 Hz, 1H), 1.33 (qd, J = 12.0, 4.4 Hz, 1H). $^{13}$C NMR (101 MHz, DMSO-$d_6$) δ 168.32, 155.65, 150.51, 146.14, 138.79, 136.58, 134.48, 133.28, 132.73, 131.92, 128.23, 122.47, 122.05, 120.88, 120.52, 117.93, 115.85, 114.08, 103.10, 67.00, 62.83, 57.35, 55.91, 48.94, 43.26, 39.09, 36.15, 30.44, 22.94, 18.80. $C_{33}H_{42}N_6O_3$, MS calculated for m/z $[M+H]^+$: 571.3397 (calculated), 571.3404(found).

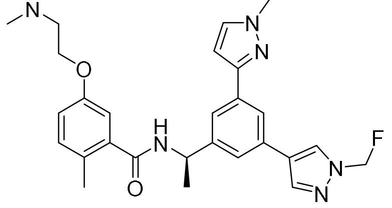

***(R)-5-(2-(dimethylamino)ethoxy)-N-(1-(3-(1-(fluoromethyl)-1H-pyrazol-4-yl)-5-(1-methyl-1H-pyrazol-3-yl)phenyl)ethyl)-2-methylbenzamide*** (**Jun12713**). White solid, 79% yield for two steps. $^1H$ NMR (400 MHz, DMSO-$d_6$) δ 8.82 (d, J = 8.2 Hz, 1H), 8.33 (t, J = 1.9 Hz, 1H), 8.24 (s, 1H), 7.93 (d, J = 3.6 Hz, 2H), 7.79 (s, 1H), 7.64 (s, 1H), 7.19 (d, J = 8.2 Hz, 1H), 7.12-7.08 (m, 1H), 6.98 (d, J = 7.4 Hz, 2H), 5.23-5.16 (m, 1H), 4.31 (t, J = 5.0 Hz, 2H), 3.90 (s, 3H), 3.51 (q, J = 4.7 Hz, 2H), 2.87 (s, 6H), 2.23 (s, 3H), 1.51 (d, J = 7.0 Hz, 3H). $^{13}C$ NMR (101 MHz, DMSO-$d_6$) δ 168.34, 155.65, 150.35, 146.31, 139.98, 138.75, 134.62, 132.78, 132.25, 131.93, 129.66, 128.24, 124.43, 122.88, 121.60, 120.96, 115.87, 114.06, 103.17, 62.83, 55.90, 48.89, 43.25, 39.10, 22.92, 18.80. $C_{28}H_{33}FN_6O_2$, MS calculated for m/z $[M+H]^+$: 505.2727 (calculated), 505.2733 (found).

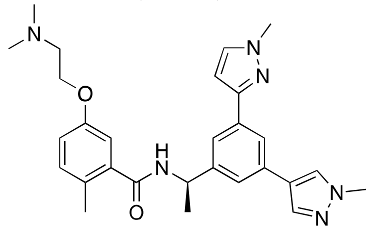

***(R)-5-(2-(dimethylamino)ethoxy)-2-methyl-N-(1-(3-(1-methyl-1H-pyrazol-3-yl)-5-(1-methyl-1H-pyrazol-4-yl)phenyl)ethyl)benzamide*** (**Jun12308**). White solid, 82% yield for two steps. $^1H$ NMR (400 MHz, DMSO-$d_6$) δ 8.77 (d, J = 8.3 Hz, 1H), 8.19 (s, 1H), 7.88 (s, 1H), 7.81 (d, J = 1.7 Hz, 1H), 7.75 (d, J = 2.2 Hz, 1H), 7.68 (d, J = 1.9 Hz, 1H), 7.51 (d, J = 1.9 Hz, 1H), 7.19 (d, J = 8.2 Hz, 1H), 7.00-6.94 (m, 2H), 6.73 (d, J = 2.3 Hz, 1H), 5.20-5.13 (m, 1H), 4.30 (t, J = 5.0 Hz, 2H), 3.89 (d, J = 3.1 Hz, 6H), 3.50 (q, J = 4.9 Hz, 2H), 2.85 (d, J = 4.3 Hz, 6H), 2.23 (s, 3H), 1.49 (d, J = 7.0 Hz, 3H). $^{13}C$ NMR (101 MHz, DMSO-$d_6$) δ 168.28, 155.66, 150.51, 146.09, 138.85, 136.46, 134.49, 133.31, 132.72, 131.92, 128.40, 128.25, 122.42, 120.90, 120.55, 115.87, 114.10, 103.09, 62.85, 55.97, 48.88, 43.31, 39.11, 22.95, 18.82. $C_{28}H_{34}N_6O_2$, HRMS calculated for m/z $[M+H]^+$: 487.2821(calculated), 487.2826 (found)

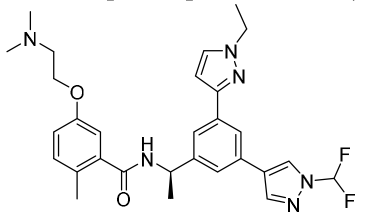

***(R)-N-(1-(3-(1-(difluoromethyl)-1H-pyrazol-4-yl)-5-(1-ethyl-1H-pyrazol-3-yl)phenyl)ethyl)-5-(2-(dimethylamino)ethoxy)-2-methylbenzamide*** (**Jun12827**). White solid, 83% yield for two steps. $^1H$ NMR (400 MHz, DMSO-$d_6$) δ 8.78 (d, J = 8.1 Hz, 2H), 8.32 (s, 1H), 7.95 (s, 1H), 7.86 (s, 1H), 7.80 (s, 2H), 7.65 (s, 1H), 7.19 (d, J = 8.2 Hz, 1H), 6.98 (d, J = 7.6 Hz, 2H), 6.79 (d, J = 2.2 Hz, 1H), 5.23-5.15 (m, 1H), 4.31 (t, J = 5.1 Hz, 2H), 4.19 (q, J = 7.2 Hz, 2H), 3.50 (q, J = 4.8 Hz, 2H), 2.86 (s, 6H), 2.24 (s, 3H), 1.51 (d, J = 7.0 Hz, 3H), 1.42 (t, J = 7.3 Hz, 3H). $^{13}C$ NMR (101 MHz, DMSO-$d_6$) δ 168.33, 155.67, 150.13, 146.29, 140.50, 138.81, 134.81, 131.92, 131.56, 131.21, 128.24, 125.90, 124.99, 123.13, 121.89, 121.31, 115.87, 114.10, 103.12, 62.85, 55.91, 48.90, 46.83, 43.26, 22.96, 18.82, 16.01. $C_{29}H_{34}F_2N_6O_2$, MS calculated for m/z $[M+H]^+$: 537.2789 (calculated), 537.2795 (found).

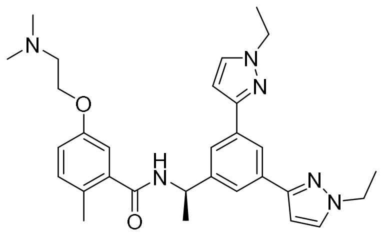

***(R)-N-(1-(3,5-bis(1-ethyl-1H-pyrazol-3-yl)phenyl)ethyl)-5-(2-(dimethylamino)ethoxy)-2-methylbenzamide*** (**Jun12831**). White solid, 82% yield. $^1$H NMR (400 MHz, DMSO-d$_6$) δ 8.82 (d, J = 8.2 Hz, 1H), 8.04 (s, 1H), 7.80 (d, J = 2.3 Hz, 2H), 7.75 (s, 2H), 7.19 (d, J = 8.2 Hz, 1H), 7.00 – 6.94 (m, 2H), 6.73 (d, J = 2.3 Hz, 2H), 5.22-5.14 (m, J = 7.1 Hz, 1H), 4.30 (t, J = 5.1 Hz, 2H), 4.19 (q, J = 7.2 Hz, 4H), 3.50 (q, J = 4.8 Hz, 2H), 2.85 (d, J = 3.9 Hz, 6H), 2.24 (s, 3H), 1.50 (d, J = 7.0 Hz, 3H), 1.42 (t, J = 7.2 Hz, 6H). $^{13}$C NMR (101 MHz, DMSO-d$_6$) δ 168.32, 155.65, 150.31, 145.95, 138.82, 134.34, 131.90, 131.21, 128.26, 122.32, 120.58, 115.82, 114.10, 102.87, 62.84, 55.89, 48.94, 46.82, 43.24, 22.87, 18.83, 16.04. $C_{30}H_{38}N_6O_2$, MS calculated for m/z [M+H]$^+$: 515.3134 (calculated), 515.3141 (found).

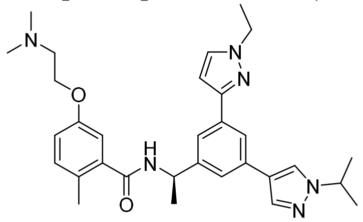

***(R)-5-(2-(dimethylamino)ethoxy)-N-(1-(3-(1-ethyl-1H-pyrazol-3-yl)-5-(1-isopropyl-1H-pyrazol-4-yl)phenyl)ethyl)-2-methylbenzamide*** (**Jun12833**). White solid, 71% yield for two steps. $^1$H NMR (400 MHz, DMSO-d$_6$) δ 8.78 (d, J = 8.2 Hz, 1H), 8.28 (s, 1H), 7.90 (s, 1H), 7.84 (s, 1H), 7.80 (s, 1H), 7.69 (s, 1H), 7.54 (s, 1H), 7.19 (d, J = 8.2 Hz, 1H), 6.98 (d, J = 7.8 Hz, 2H), 6.75 (s, 1H), 5.21-5.13 (m, 1H), 4.56-4.49 (m, 1H), 4.31 (t, J = 5.1 Hz, 2H), 4.18 (q, J = 7.3 Hz, 2H), 3.51 (s, 2H), 2.86 (s, 6H), 2.24 (s, 3H), 1.50 (d, J = 6.9 Hz, 3H), 1.47 (d, J = 6.7 Hz, 6H), 1.42 (t, J = 7.2 Hz, 3H). $^{13}$C NMR (101 MHz, DMSO-d$_6$) δ 168.30, 155.66, 150.39, 146.04, 138.84, 135.96, 134.57, 133.49, 131.91, 131.11, 128.22, 125.23, 122.42, 121.95, 120.80, 120.57, 115.82, 114.10, 102.99, 62.85, 55.89, 53.62, 48.94, 46.80, 43.25, 23.15, 22.98, 18.82, 16.03. $C_{31}H_{40}N_6O_2$, MS calculated for m/z [M+H]$^+$: 529.3291(calculated), 529.3297 (found).

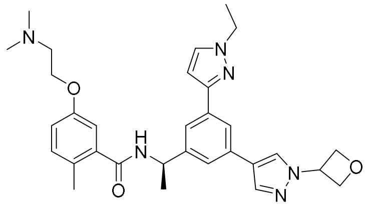

***(R)-5-(2-(dimethylamino)ethoxy)-N-(1-(3-(1-ethyl-1H-pyrazol-3-yl)-5-(1-(oxetan-3-yl)-1H-pyrazol-4-yl)phenyl)ethyl)-2-methylbenzamide*** (**Jun12834**). White solid, 78% yield for two steps. $^1$H NMR (400 MHz, DMSO-d$_6$) δ 8.61 (d, J = 8.2 Hz, 1H), 8.28 (s, 1H), 7.89 (s, 1H), 7.69 (s, 1H), 7.62 (s, 1H), 7.54 (s, 1H), 7.39 (s, 1H), 7.01 (d, J = 8.3 Hz, 1H), 6.80 (d, J = 7.9 Hz, 2H), 6.58 (s, 1H), 5.47-5.40 (m, 1H), 5.04-4.96 (m, 1H), 4.77 (d, J = 3.6 Hz, 2H), 4.13 (d, J = 5.3 Hz, 2H), 4.01 (q, J = 7.2 Hz, 2H), 2.68 (s, 6H), 2.07 (s, 3H), 1.32 (d, J = 7.0 Hz, 3H), 1.24 (t, J = 7.2 Hz, 3H). $^{13}$C NMR (101 MHz, DMSO-d$_6$) δ 168.32, 155.66, 150.31, 146.14, 138.82, 137.46, 134.65, 132.98, 131.91, 131.15, 128.23, 127.21, 122.81, 122.55, 121.08, 120.73, 115.84, 114.09, 102.99, 77.10, 62.85, 55.89, 55.15, 48.91, 46.81, 43.25, 22.96, 18.82, 16.02. $C_{31}H_{38}N_6O_3$, MS calculated for m/z [M+H]$^+$: 543.3084 (calculated), 543.3091 (found).

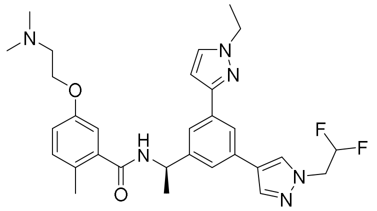

***(R)-N-(1-(3-(1-(2,2-difluoroethyl)-1H-pyrazol-4-yl)-5-(1-ethyl-1H-pyrazol-3-yl)phenyl)ethyl)-5-(2-(dimethylamino)ethoxy)-2-methylbenzamide*** (**Jun12835**). White solid, 75% yield for two steps. $^{1}$H NMR (400 MHz, DMSO-d$_6$) δ 8.79 (d, J = 8.2 Hz, 1H), 8.29 (s, 1H), 8.03 (s, 1H), 7.85 (s, 1H), 7.80 (s, 1H), 7.72 (s, 1H), 7.55 (s, 1H), 7.19 (d, J = 8.2 Hz, 1H), 6.98 (d, J = 8.2 Hz, 2H), 6.76 (d, J = 2.1 Hz, 1H), 5.47 (s, 1H), 5.22-5.14 (m, 1H), 4.67 (td, J = 15.2, 3.8 Hz, 2H), 4.31 (d, J = 5.2 Hz, 2H), 4.19 (q, J = 7.2 Hz, 2H), 3.50 (d, J = 5.0 Hz, 2H), 2.86 (s, 6H), 2.24 (s, 3H), 1.50 (d, J = 7.0 Hz, 3H), 1.42 (t, J = 7.2 Hz, 3H). $^{13}$C NMR (101 MHz, DMSO-d$_6$) δ 168.32, 155.67, 150.26, 146.20, 138.82, 137.75, 134.67, 132.79, 131.91, 131.18, 129.12, 128.22, 123.07, 122.54, 121.24, 120.69, 115.84, 114.49, 114.08, 102.99, 62.85, 55.89, 48.91, 46.82, 43.24, 22.94, 18.81, 16.02. $C_{30}H_{36}F_2N_6O_2$, MS calculated for m/z $[M+H]^+$: 551.2946(calculated), 551.2952 (found).

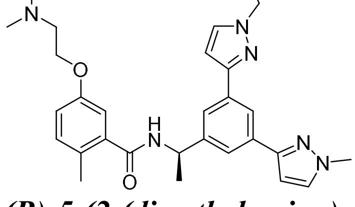

***(R)-5-(2-(dimethylamino)ethoxy)-N-(1-(3-(1-ethyl-1H-pyrazol-3-yl)-5-(1-methyl-1H-pyrazol-3-yl)phenyl)ethyl)-2-methylbenzamide*** (**Jun13836**). White solid, 70% yield for two steps. $^{1}$H NMR (400 MHz, DMSO-d$_6$) δ 8.83 (d, J = 8.2 Hz, 1H), 8.05 (s, 1H), 7.80 (d, J = 2.3 Hz, 1H), 7.76 (s, 3H), 7.19 (d, J = 8.1 Hz, 1H), 6.97 (d, J = 8.7 Hz, 2H), 6.73 (s, 2H), 5.22-5.15 (m, 1H), 4.31 (t, J = 5.0 Hz, 2H), 4.19 (q, J = 7.3 Hz, 2H), 3.90 (s, 3H), 3.50 (q, J = 4.9 Hz, 2H), 2.86 (s, 6H), 2.24 (s, 3H), 1.50 (d, J = 7.0 Hz, 3H), 1.42 (t, J = 7.2 Hz, 3H). $^{13}$C NMR (101 MHz, DMSO-d$_6$) δ 168.29, 155.65, 150.48, 150.29, 145.98, 138.82, 134.34, 134.22, 132.77, 131.90, 131.22, 128.27, 122.34, 122.30, 120.55, 115.83, 114.10, 103.01, 102.85, 62.84, 55.90, 48.92, 46.82, 43.25, 39.10, 22.87, 18.82, 16.03. $C_{29}H_{36}N_6O_2$, MS calculated for m/z $[M+H]^+$: 501.2978 (calculated), 501.2989 (found).

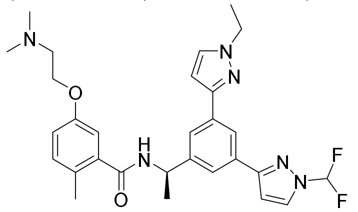

***(R)-N-(1-(3-(1-(difluoromethyl)-1H-pyrazol-3-yl)-5-(1-ethyl-1H-pyrazol-3-yl)phenyl)ethyl)-5-(2-(dimethylamino)ethoxy)-2-methylbenzamide*** (**Jun12837**). White solid, 68% yield for two steps. $^{1}$H NMR (400 MHz, DMSO-d$_6$) δ 8.86 (d, J = 8.1 Hz, 1H), 8.33 (s, 1H), 8.14 (s, 1H), 7.88 (s, 2H), 7.86 (s, 1H), 7.82 (s, 1H), 7.19 (d, J = 8.2 Hz, 1H), 7.10 (d, J = 2.7 Hz, 1H), 6.98 (d, J = 9.5 Hz, 2H), 6.77 (s, 1H), 5.25-5.17 (m, 1H), 4.30 (d, J = 5.3 Hz, 2H), 4.20 (q, J = 7.3 Hz, 2H), 3.50 (s, 2H), 2.85 (s, 6H), 2.23 (s, 3H), 1.51 (d, J = 7.0 Hz, 3H), 1.43 (t, J = 7.2 Hz, 3H). $^{13}$C NMR (101 MHz, DMSO-d$_6$) δ 168.34, 155.66, 149.98, 146.34, 138.77, 134.64, 132.58, 131.93, 131.34, 128.26, 123.66, 122.95, 121.27, 115.88, 114.07, 106.09, 103.01, 62.83, 55.90, 48.91,

46.86, 43.26, 22.85, 18.81, 16.05. $C_{29}H_{34}F_2N_6O_2$, MS calculated for m/z $[M+H]^+$: 537.2789 (calculated), 537.2796 (found).

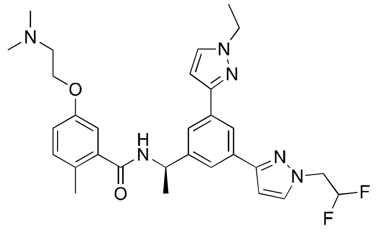

***(R)-N-(1-(3-(1-(2,2-difluoroethyl)-1H-pyrazol-3-yl)-5-(1-ethyl-1H-pyrazol-3-yl)phenyl)ethyl)-5-(2-(dimethylamino)ethoxy)-2-methylbenzamide*** (**Jun1314**). White solid, 66% yield for two steps. $^1H$ NMR (400 MHz, DMSO-$d_6$) δ 8.77 (d, J = 8.2 Hz, 1H), 8.28 (s, 1H), 8.02 (s, 1H), 7.84 (s, 1H), 7.80 (d, J = 2.3 Hz, 1H), 7.71 (s, 1H), 7.54 (s, 1H), 7.19 (d, J = 8.2 Hz, 1H), 7.00 – 6.93 (m, 2H), 6.75 (d, J = 2.3 Hz, 1H), 5.21-5.14 (m, 1H), 4.67 (td, J = 15.2, 3.8 Hz, 2H), 4.30 (t, J = 5.0 Hz, 2H), 4.18 (q, J = 7.3 Hz, 3H), 3.50 (q, J = 5.0 Hz, 2H), 2.85 (d, J = 4.1 Hz, 6H), 2.24 (s, 3H), 1.49 (d, J = 7.0 Hz, 3H), 1.42 (t, J = 7.2 Hz, 3H). $^{13}C$ NMR (101 MHz, DMSO-$d_6$) δ 168.40, 168.33, 155.66, 151.58, 150.22, 146.07, 138.80, 134.44, 133.72, 131.91, 131.25, 128.26, 122.71, 122.57, 120.76, 115.84, 114.09, 103.92, 102.89, 62.86, 55.88, 48.94, 46.84, 43.24, 22.86, 18.82, 16.04. $C_{30}H_{36}F_2N_6O_2$, MS calculated for m/z $[M+H]^+$: 551.2946 (calculated), 551.2951 (found).

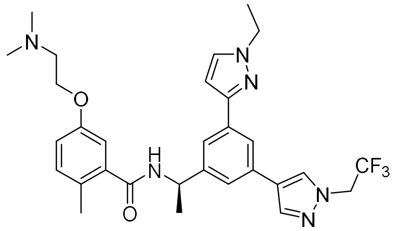

***(R)-5-(2-(dimethylamino)ethoxy)-N-(1-(3-(1-ethyl-1H-pyrazol-3-yl)-5-(1-(2,2,2-trifluoroethyl)-1H-pyrazol-4-yl)phenyl)ethyl)-2-methylbenzamide*** (**Jun12839**). White solid, 72% yield for two steps. $^1H$ NMR (400 MHz, $CDCl_3$) δ 7.97 (s, 1H), 7.91 (s, 1H), 7.85 (s, 1H), 7.75 (s, 1H), 7.56 (s, 1H), 7.51 (d, J = 2.4 Hz, 1H), 7.13 (d, J = 8.4 Hz, 1H), 6.98 (d, J = 2.7 Hz, 1H), 6.83 (dd, J = 8.4, 2.6 Hz, 1H), 6.66 (d, J = 6.6 Hz, 2H), 5.41-5.34 (m, J = 7.1 Hz, 1H), 4.78 (q, J = 8.3 Hz, 2H), 4.41-4.36 (m, 2H), 4.32 (q, J = 7.3 Hz, 2H), 3.45 (d, J = 4.7 Hz, 2H), 2.92 (d, J = 2.5 Hz, 6H), 2.36 (s, 3H), 1.68 (d, J = 6.8 Hz, 3H), 1.57 (t, J = 7.3 Hz, 3H). $^{13}C$ NMR (101 MHz, DMSO-$d_6$) δ 168.34, 155.67, 150.21, 146.25, 138.80, 138.49, 134.71, 132.46, 131.91, 131.19, 129.51, 128.21, 123.55, 122.66, 121.43, 120.80, 115.84, 114.07, 103.03, 62.84, 55.88, 48.91, 46.82, 43.24, 22.93, 18.80, 16.01. $C_{30}H_{35}F_3N_6O_2$, MS calculated for m/z $[M+H]^+$: 569.2852 (calculated), 569.2860 (found).

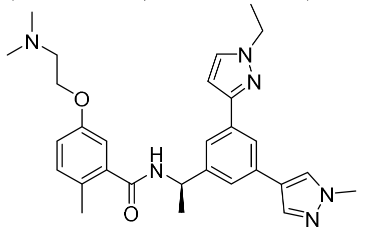

***(R)-5-(2-(dimethylamino)ethoxy)-N-(1-(3-(1-ethyl-1H-pyrazol-3-yl)-5-(1-methyl-1H-pyrazol-4-yl)phenyl)ethyl)-2-methylbenzamide*** (**Jun12682**). White solid, 74% yield for two steps. $^1H$ NMR (400 MHz, DMSO-$d_6$) δ 8.71 (d, J = 8.2 Hz, 1H), 8.12 (s, 1H), 7.82 (s, 1H), 7.77-7.71 (m, 2H), 7.62 (s, 1H), 7.45 (s, 1H), 7.12 (d, J = 8.2 Hz, 1H), 6.90 (d, J = 7.3 Hz, 2H), 6.67 (d, J = 2.2 Hz, 1H), 5.13-5.06 (m, 1H), 4.24 (t, J = 5.0 Hz, 2H), 4.11 (q, J = 7.2 Hz, 2H), 3.81 (s, 3H), 3.43

(s, 2H), 2.79 (s, 6H), 2.17 (s, 3H), 1.42 (d, J = 7.0 Hz, 3H), 1.35 (t, J = 7.3 Hz, 3H). $^{13}$C NMR (101 MHz, DMSO-d$_6$) δ 168.31, 155.67, 150.34, 146.11, 138.83, 136.48, 134.61, 133.30, 131.91, 131.15, 128.40, 128.23, 122.44, 122.38, 120.90, 120.58, 115.84, 114.10, 102.96, 62.86, 55.89, 48.91, 46.81, 43.25, 39.11, 22.96, 18.83, 16.02. $C_{29}H_{36}N_6O_2$, MS calculated for m/z $[M+H]^+$: 501.2978 (calculated), 501.2985 (found).

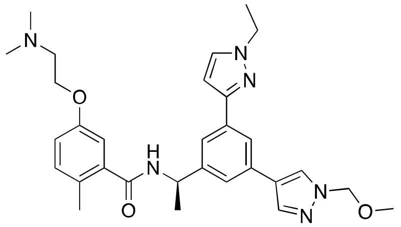

***(R)-5-(2-(dimethylamino)ethoxy)-N-(1-(3-(1-ethyl-1H-pyrazol-3-yl)-5-(1-(methoxymethyl)-1H-pyrazol-3-yl)phenyl)ethyl)-2-methylbenzamide*** (**Jun12924**). White solid, 79% yield for two steps. $^1$H NMR (400 MHz, DMSO-d$_6$) δ 8.78 (d, J = 8.2 Hz, 1H), 8.42 (s, 1H), 8.04 (s, 1H), 7.88 (d, J = 2.0 Hz, 1H), 7.80 (d, J = 2.3 Hz, 1H), 7.73 (s, 1H), 7.58 (s, 1H), 7.19 (d, J = 9.0 Hz, 1H), 6.98 (d, J = 7.0 Hz, 2H), 6.76 (d, J = 2.4 Hz, 1H), 5.42 (s, 2H), 5.22-5.15 (m, 1H), 4.31 (t, J = 5.0 Hz, 2H), 4.18 (q, J = 7.3 Hz, 2H), 3.51 (q, J = 4.6 Hz, 2H), 3.28 (s, 3H), 2.86 (d, J = 3.4 Hz, 6H), 2.24 (s, 3H), 1.51 (d, J = 7.0 Hz, 3H), 1.42 (t, J = 7.2 Hz, 3H). $^{13}$C NMR (101 MHz, DMSO-d$_6$) δ 168.32, 155.67, 150.29, 146.17, 138.83, 137.85, 134.68, 132.87, 131.91, 131.15, 128.52, 128.24, 123.29, 122.63, 121.22, 120.81, 115.85, 114.10, 103.01, 81.79, 62.87, 56.59, 55.91, 48.92, 46.81, 43.26, 22.96, 18.81, 16.00. $C_{30}H_{38}N_6O_3$, MS calculated for m/z $[M+H]^+$: 531.3084 (calculated), 531.3091 (found).

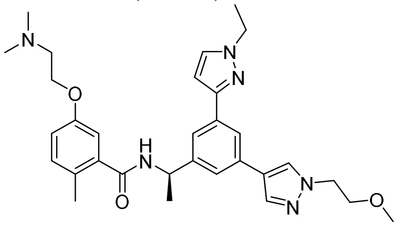

***(R)-5-(2-(dimethylamino)ethoxy)-N-(1-(3-(1-ethyl-1H-pyrazol-3-yl)-5-(1-(2-methoxyethyl)-1H-pyrazol-4-yl)phenyl)ethyl)-2-methylbenzamide*** (**Jun12763**). White solid, 74% yield for two steps. $^1$H NMR (400 MHz, DMSO-d$_6$) δ 8.72 (d, J = 8.3 Hz, 1H), 8.14 (s, 1H), 7.87-7.84 (m, 1H), 7.76 (t, J = 1.6 Hz, 1H), 7.73 (d, J = 2.3 Hz, 1H), 7.62 (d, J = 1.6 Hz, 1H), 7.46 (d, J = 1.7 Hz, 1H), 7.12 (d, J = 8.2 Hz, 1H), 6.90 (d, J = 7.4 Hz, 2H), 6.68 (d, J = 2.3 Hz, 1H), 5.10 (m, 1H), 4.23 (q, J = 5.5 Hz, 4H), 4.11 (q, J = 7.2 Hz, 2H), 3.66 (t, J = 5.3 Hz, 2H), 3.42 (d, J = 6.3 Hz, 2H), 3.18 (s, 3H), 2.84-2.75 (m, 6H), 2.17 (s, 3H), 1.42 (d, J = 7.0 Hz, 3H), 1.35 (t, J = 7.3 Hz, 3H). $^{13}$C NMR (101 MHz, DMSO-d$_6$) δ 168.30, 155.67, 150.34, 146.11, 138.84, 136.63, 134.60, 133.28, 131.91, 131.14, 128.21, 128.18, 122.39, 122.21, 120.92, 120.55, 115.83, 114.08, 102.98, 70.95, 62.85, 58.44, 55.88, 51.78, 48.92, 46.81, 43.24, 22.98, 18.82, 16.03. $C_{31}H_{40}N_6O_3$, MS calculated for m/z $[M+H]^+$: 545.3240 (calculated), 545.3245 (found).

### HNMR and CNMR spectra of A-2

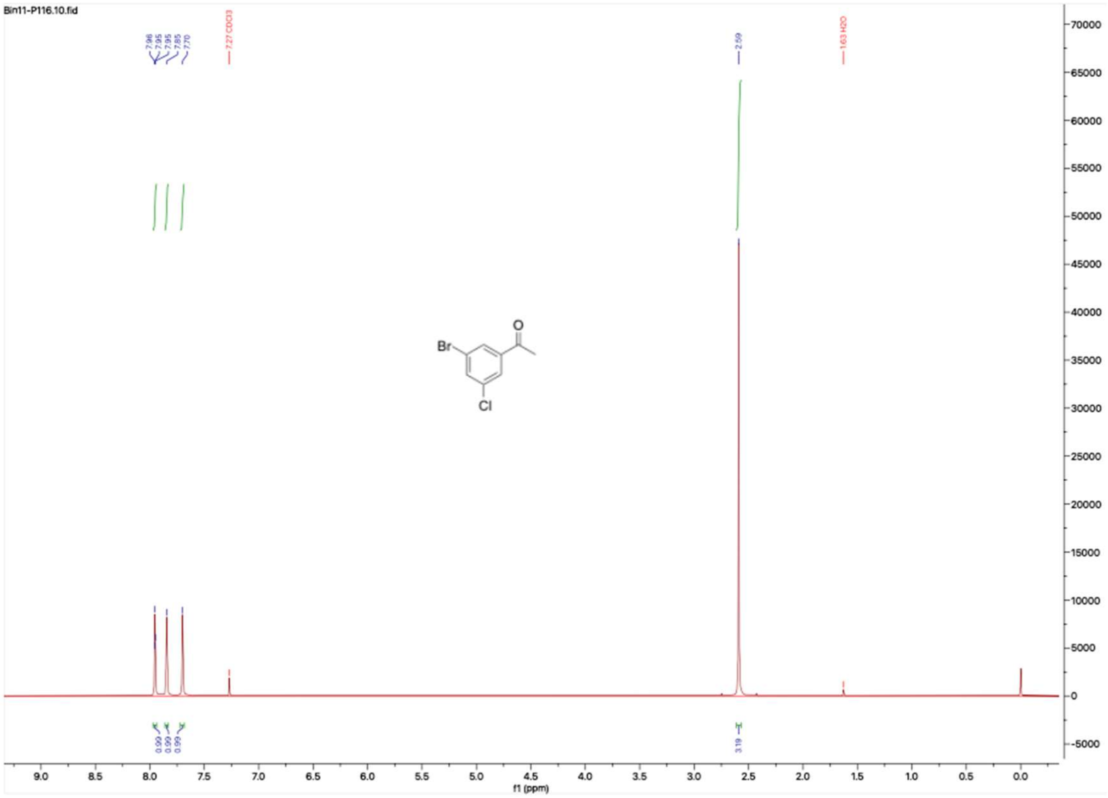


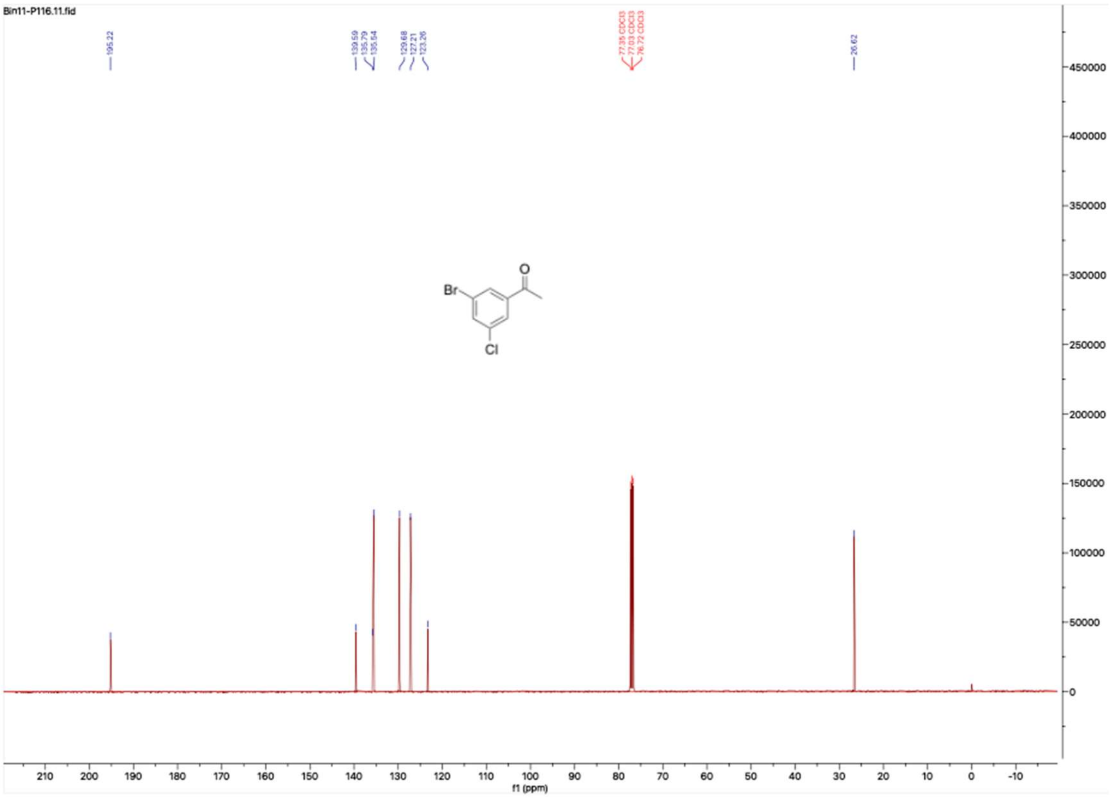

**HNMR and CNMR spectra of A-3**

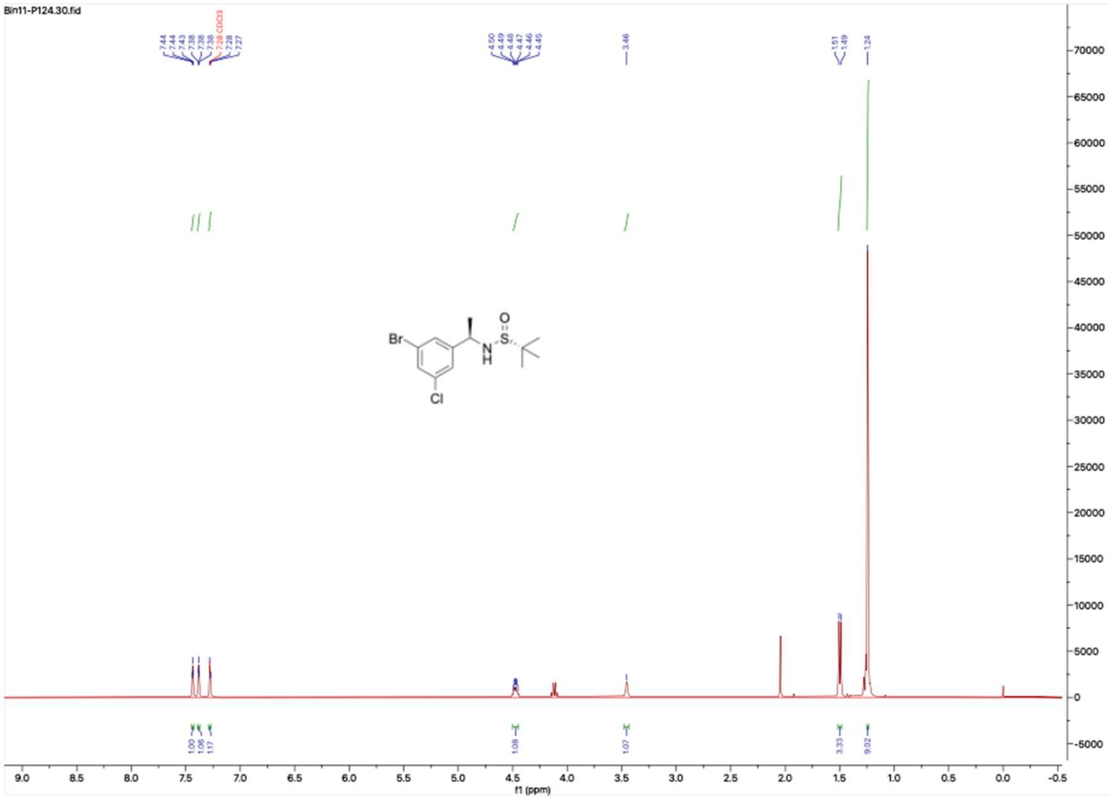

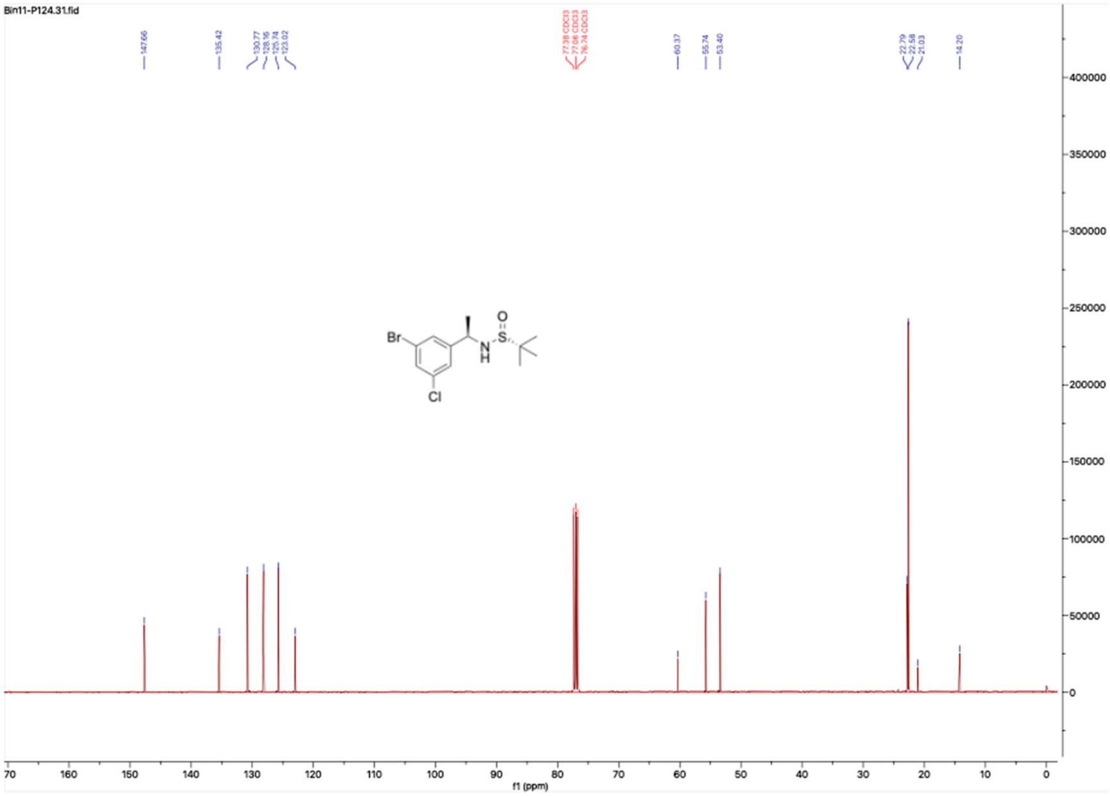

**HNMR and CNMR spectra of A-4**

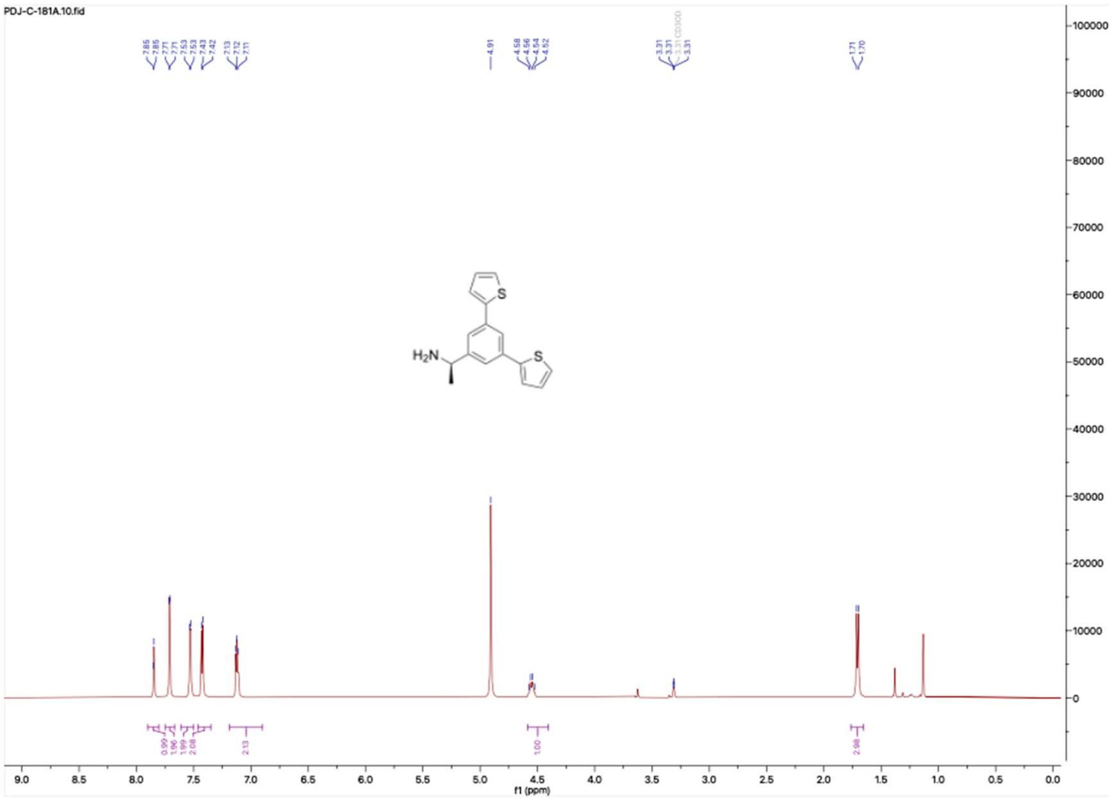


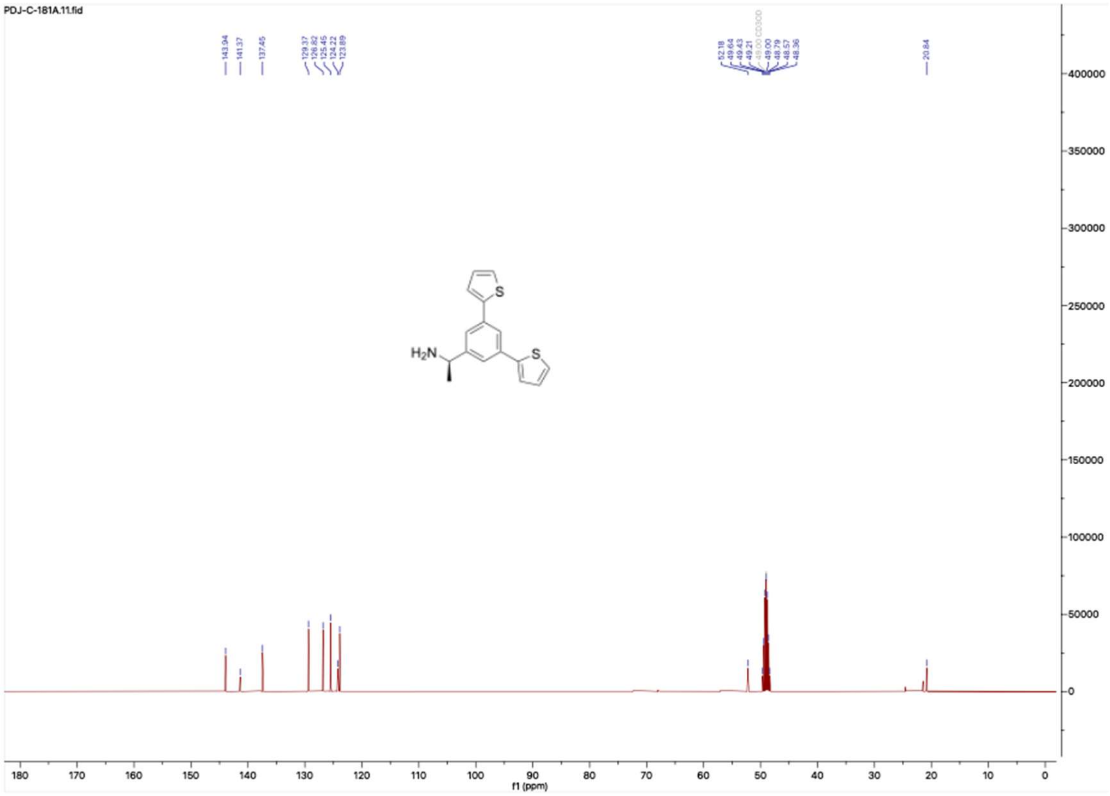

#### HNMR and CNMR spectra of A-5

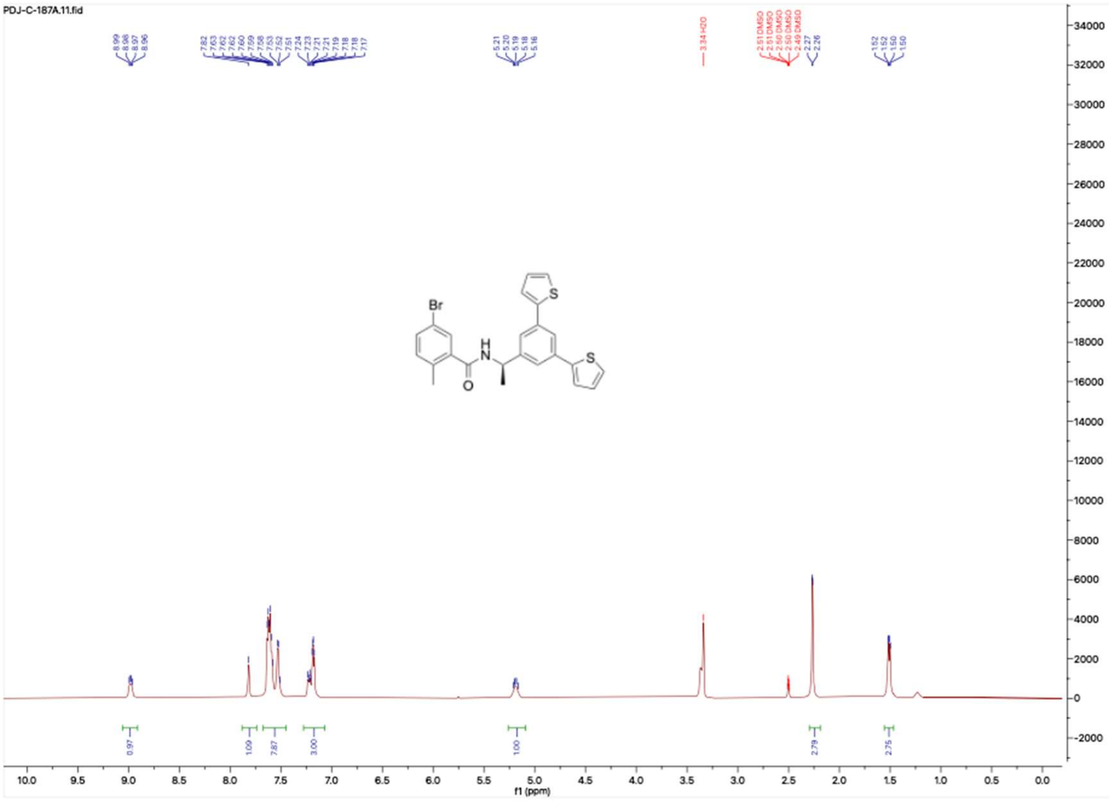


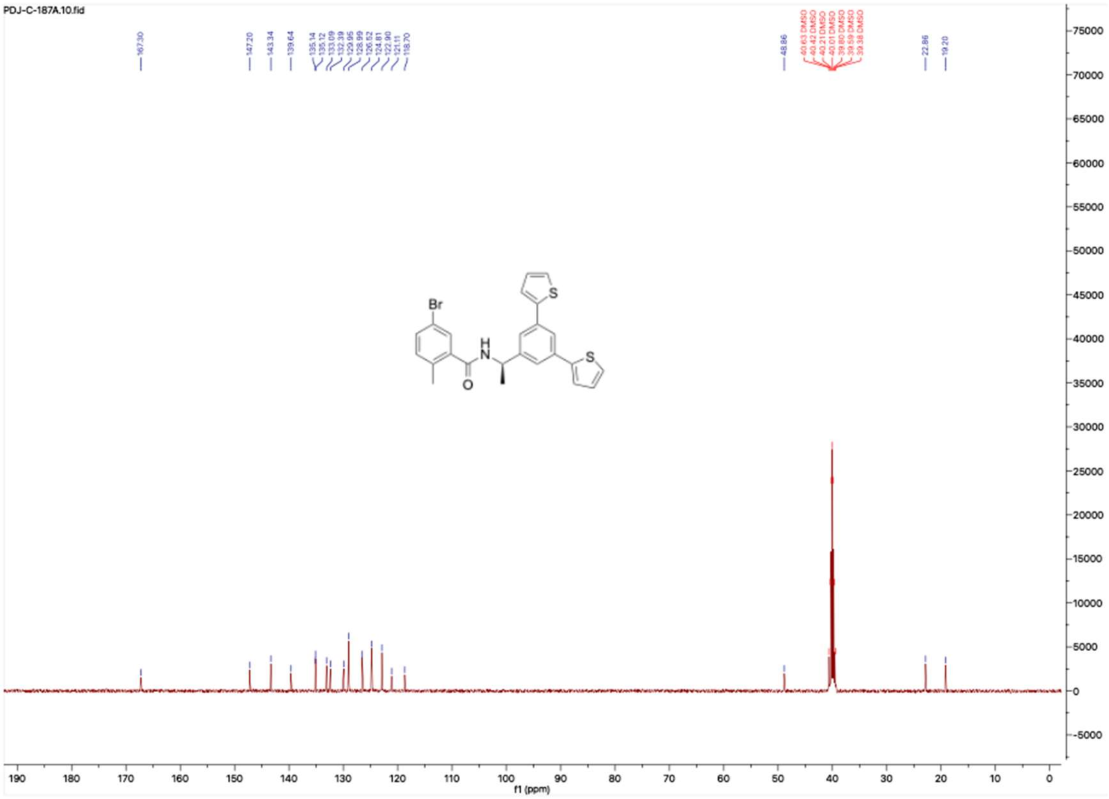

**HNMR and CNMR spectra of Jun1238**

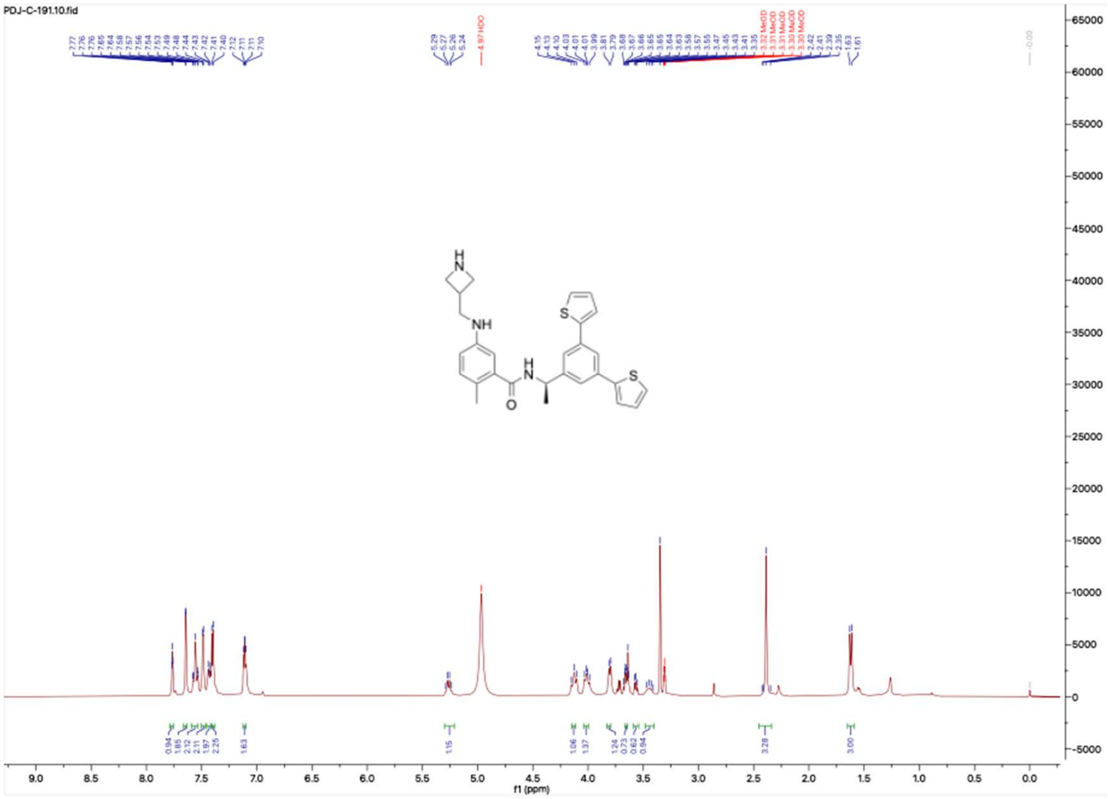

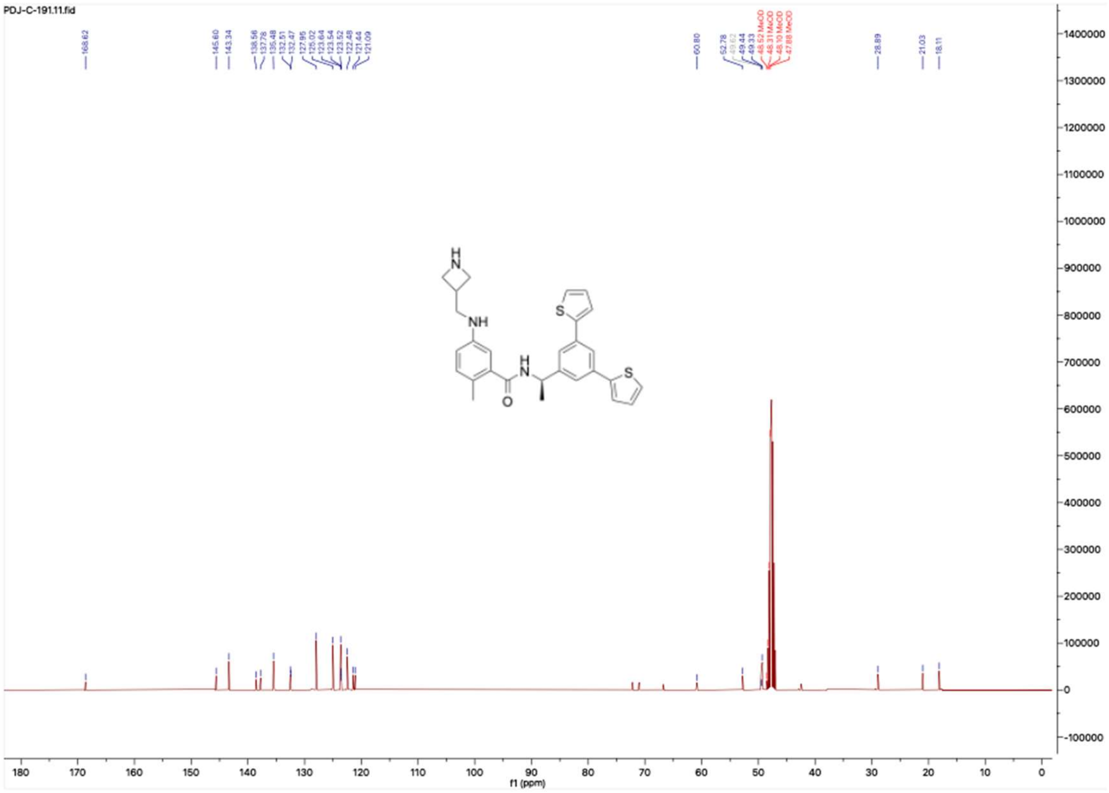

**HNMR and CNMR spectra of Jun1271**

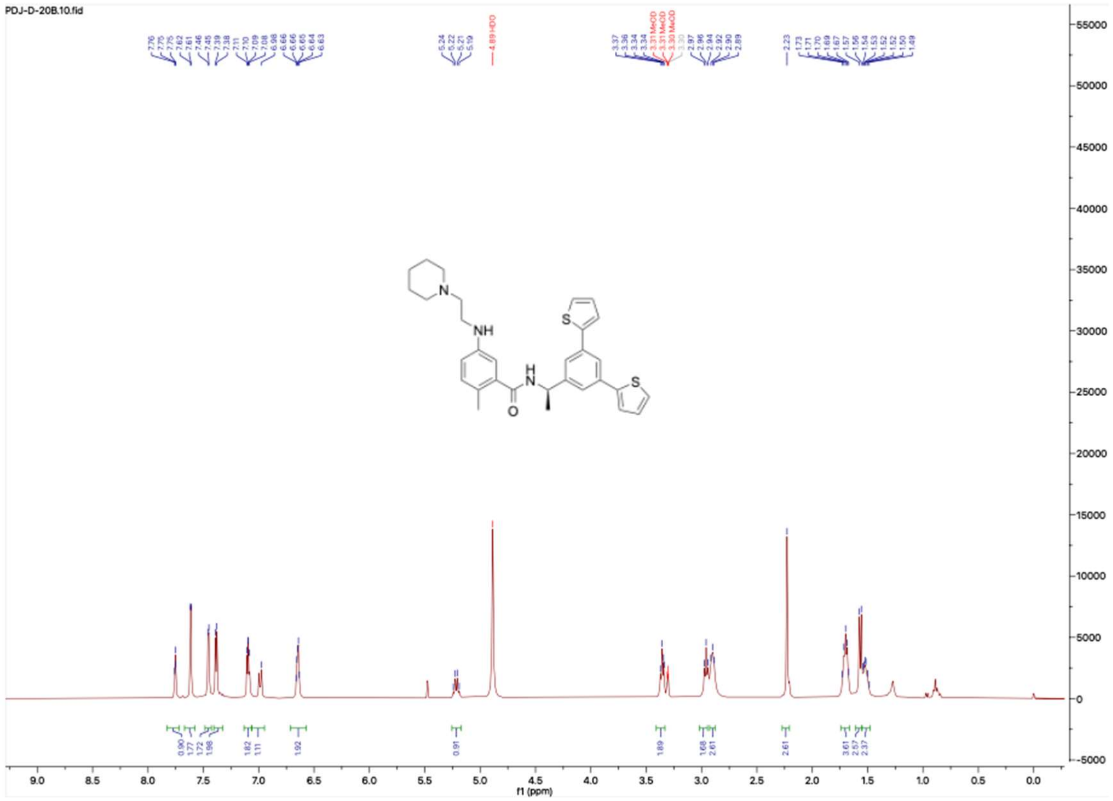

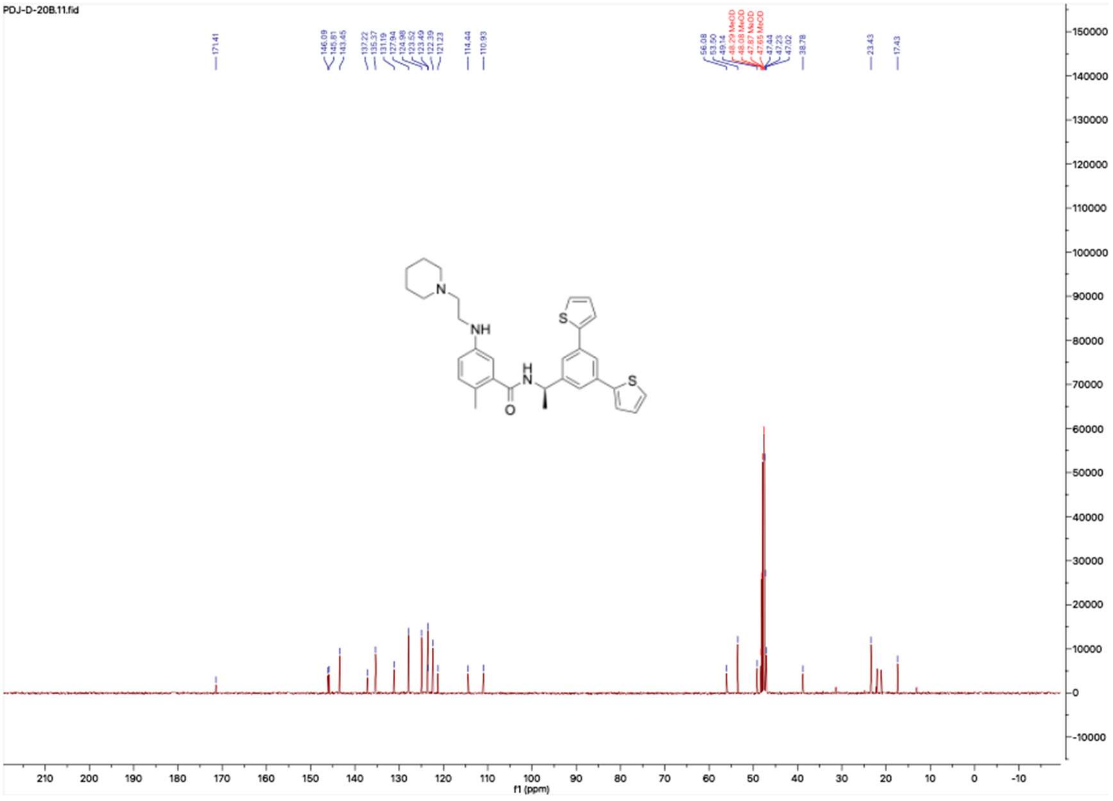

**HNMR and CNMR spectra of Jun11924**

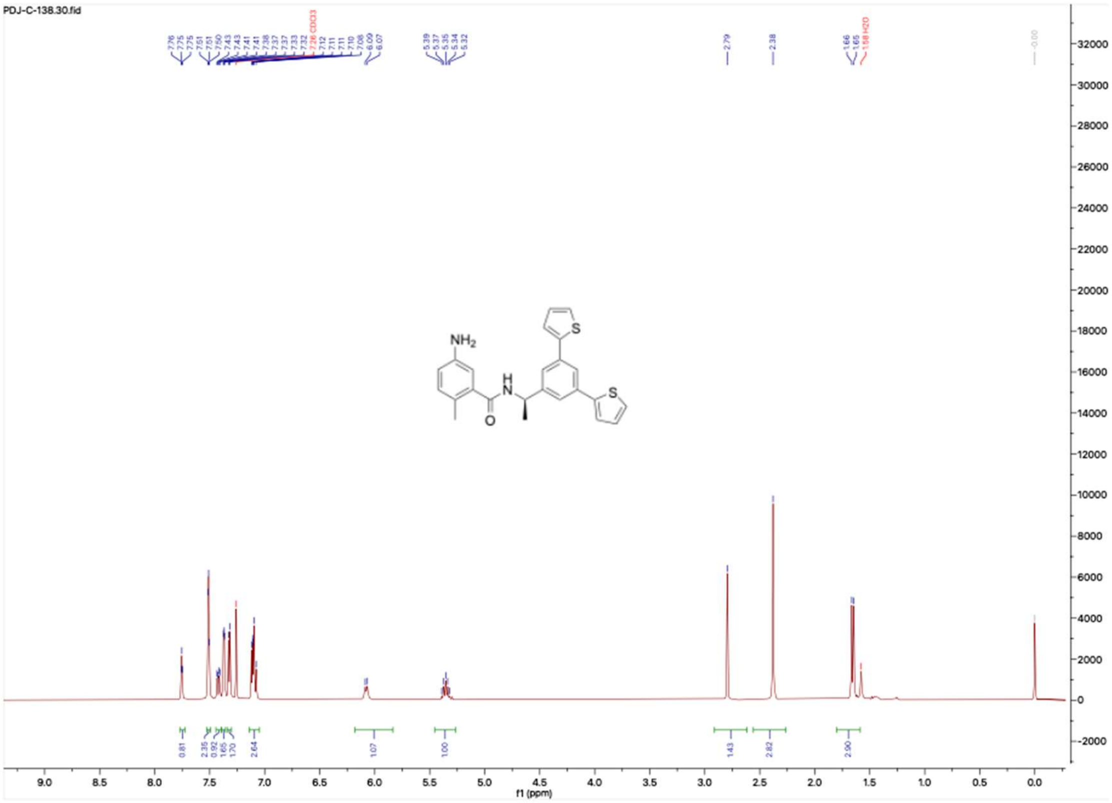

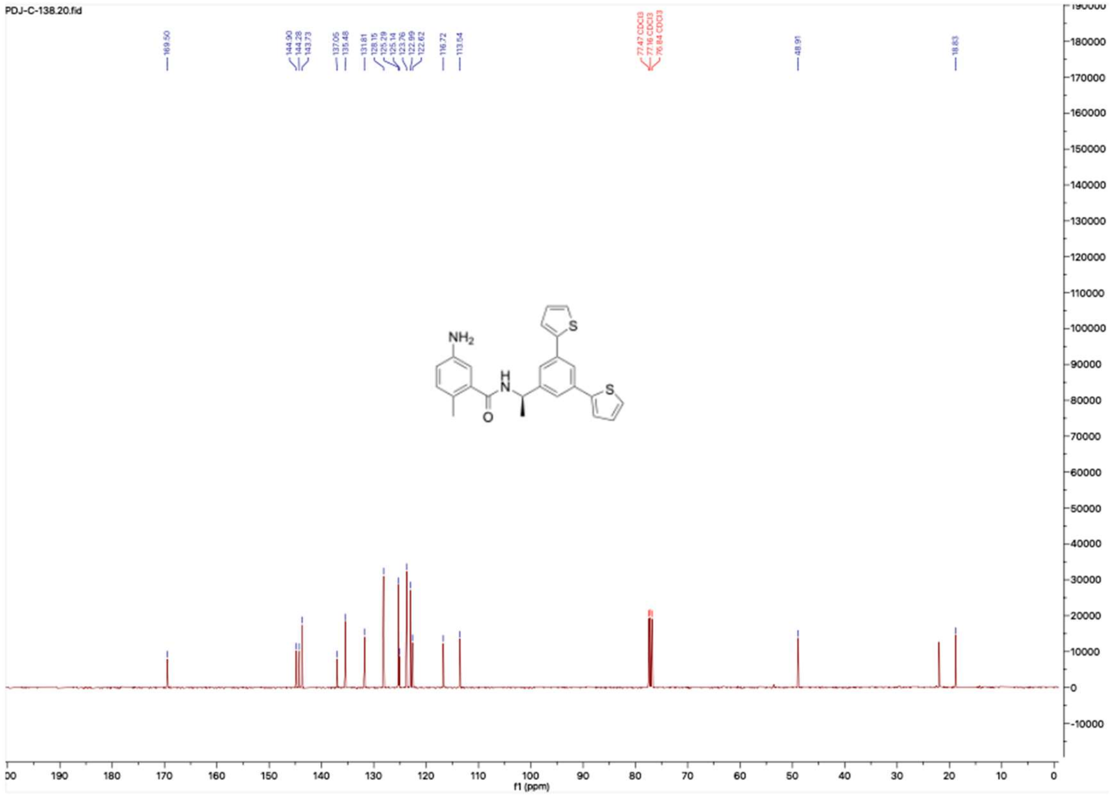

**HNMR and CNMR spectra of Jun11922**

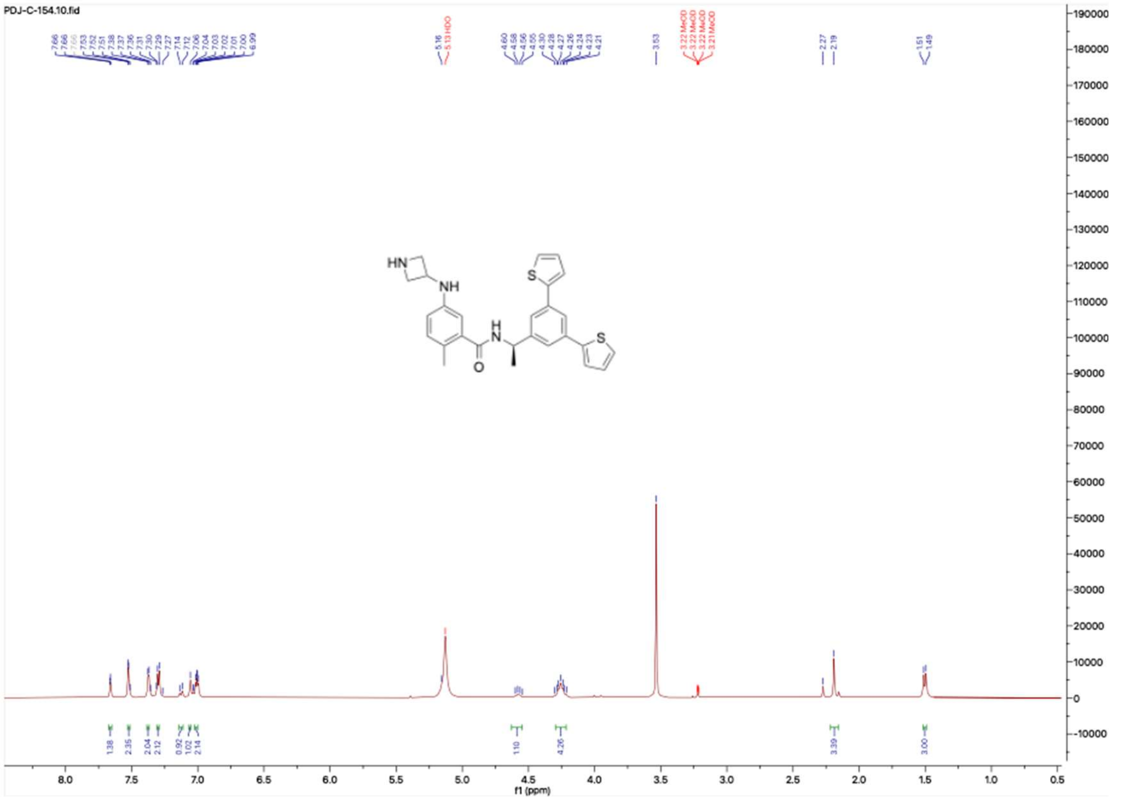

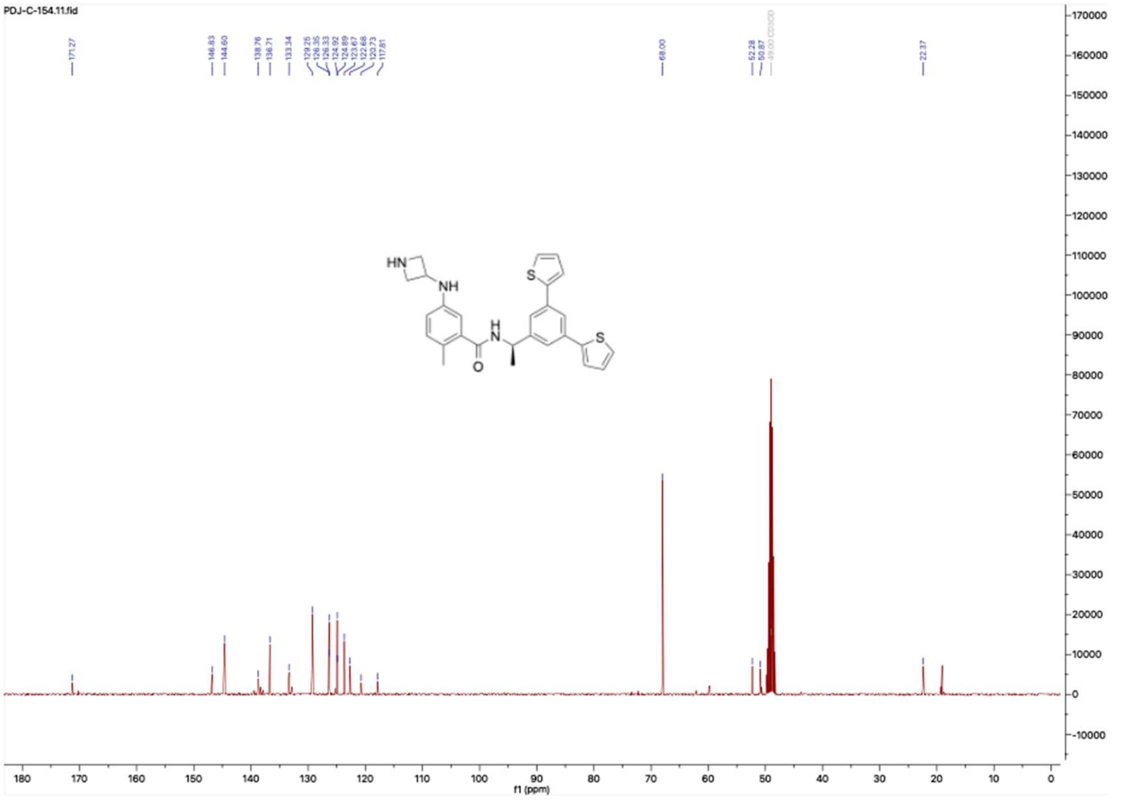

**HNMR and CNMR spectra of Jun12112**

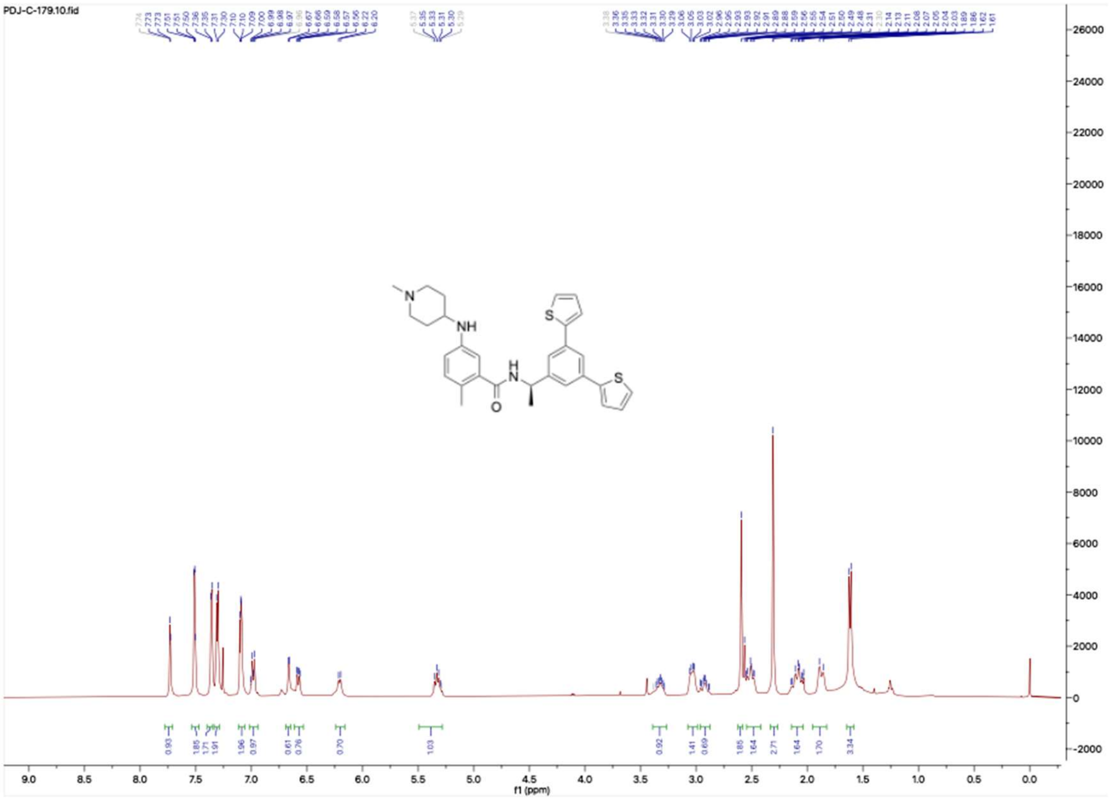

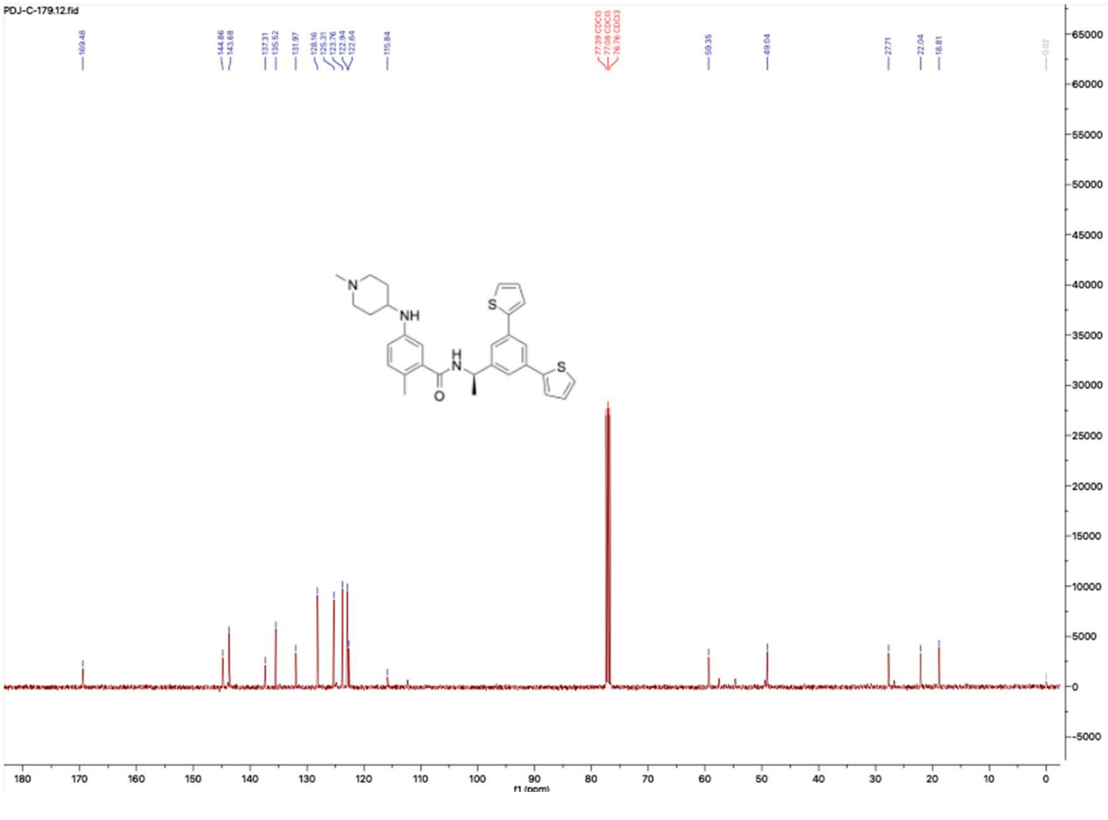

**HNMR and CNMR spectra of Jun11923**

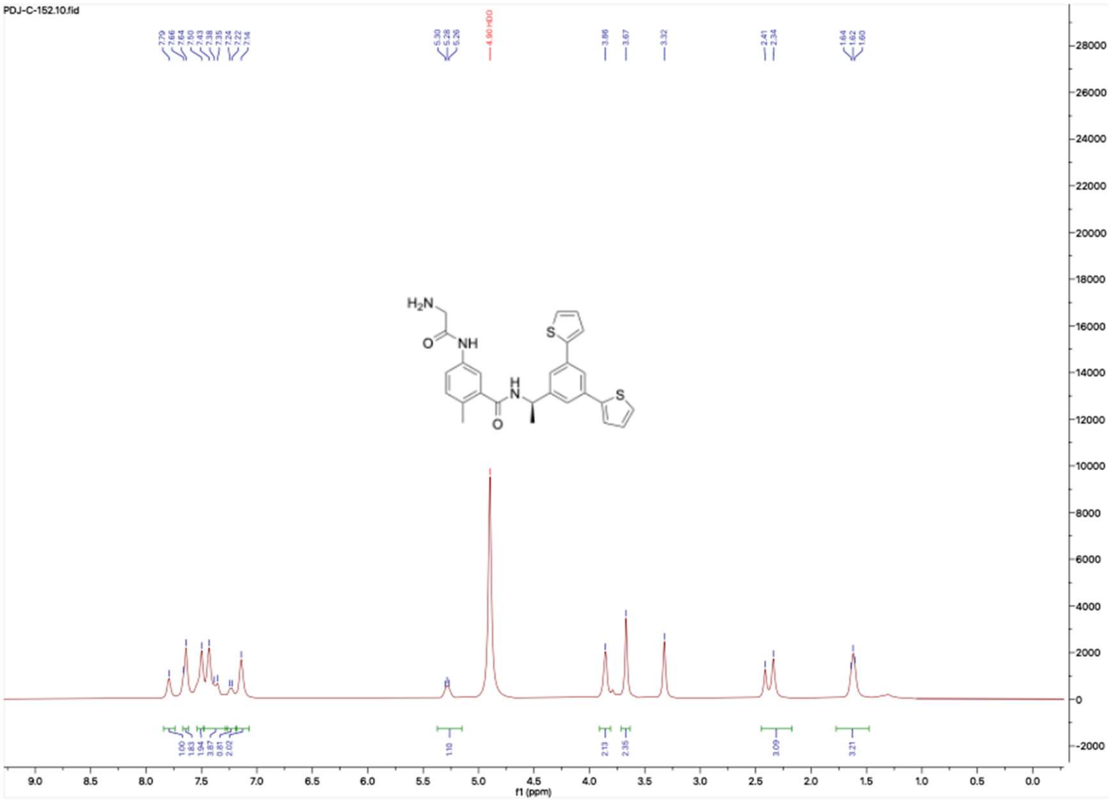

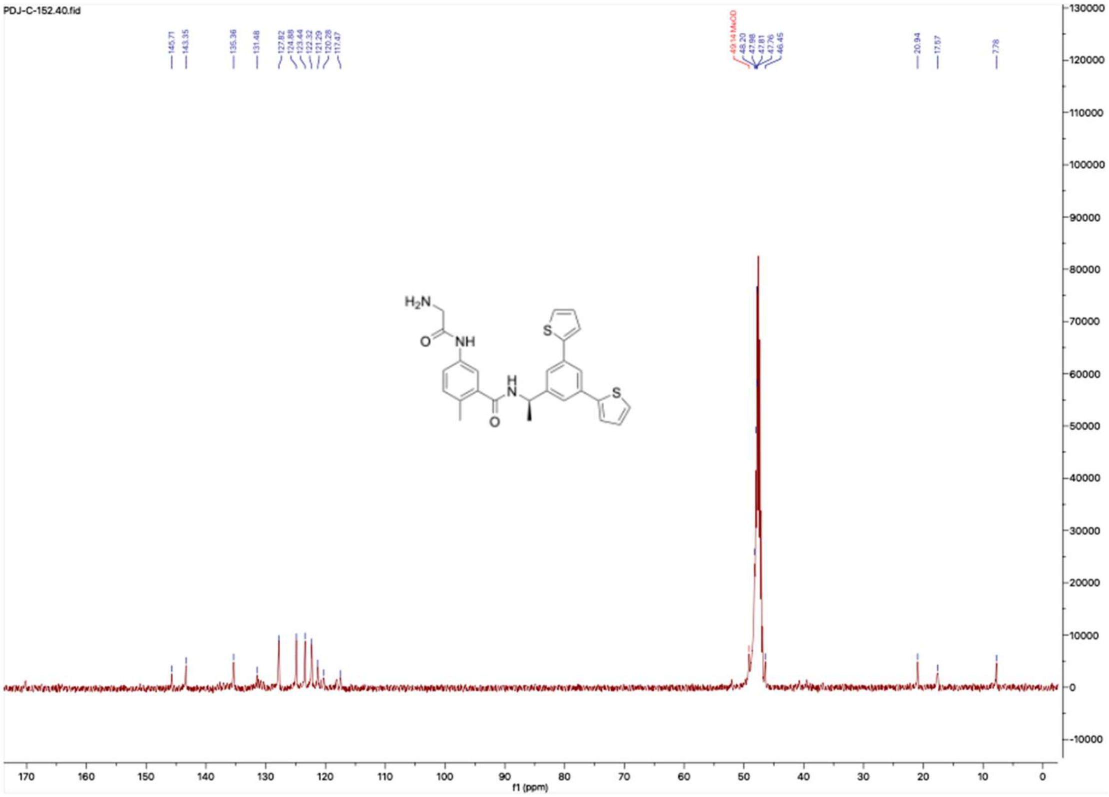

**HNMR and CNMR spectra of Jun12111**

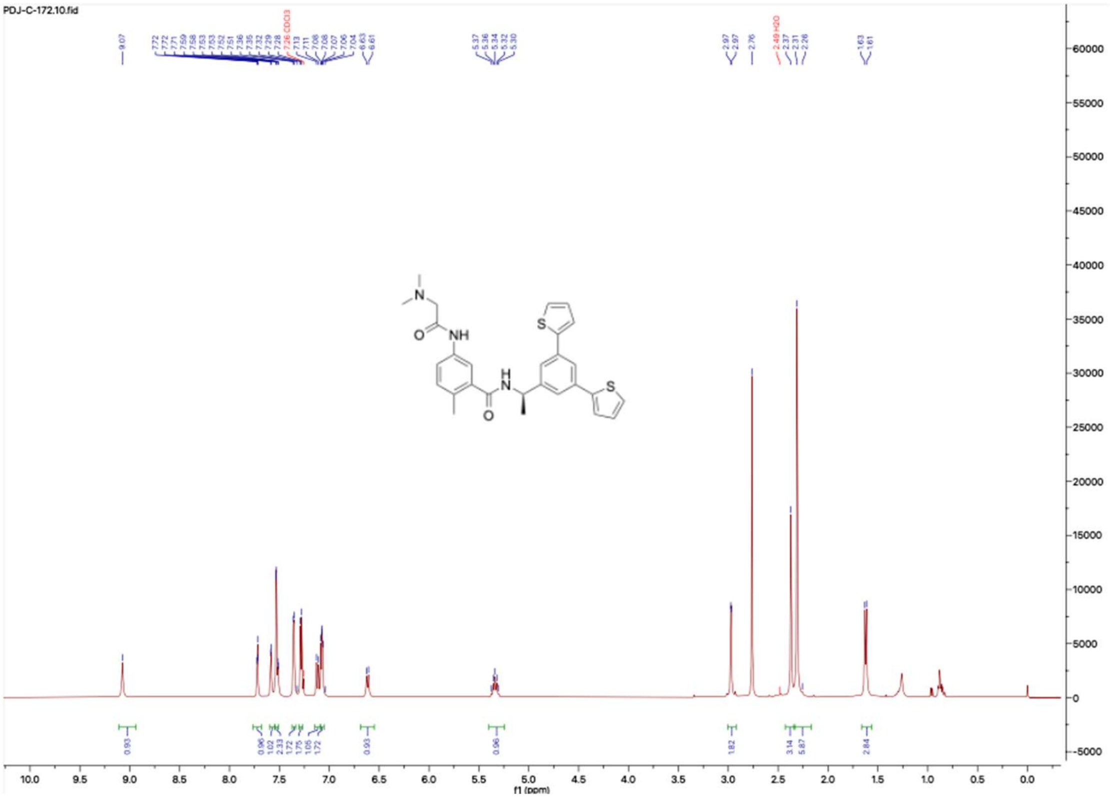

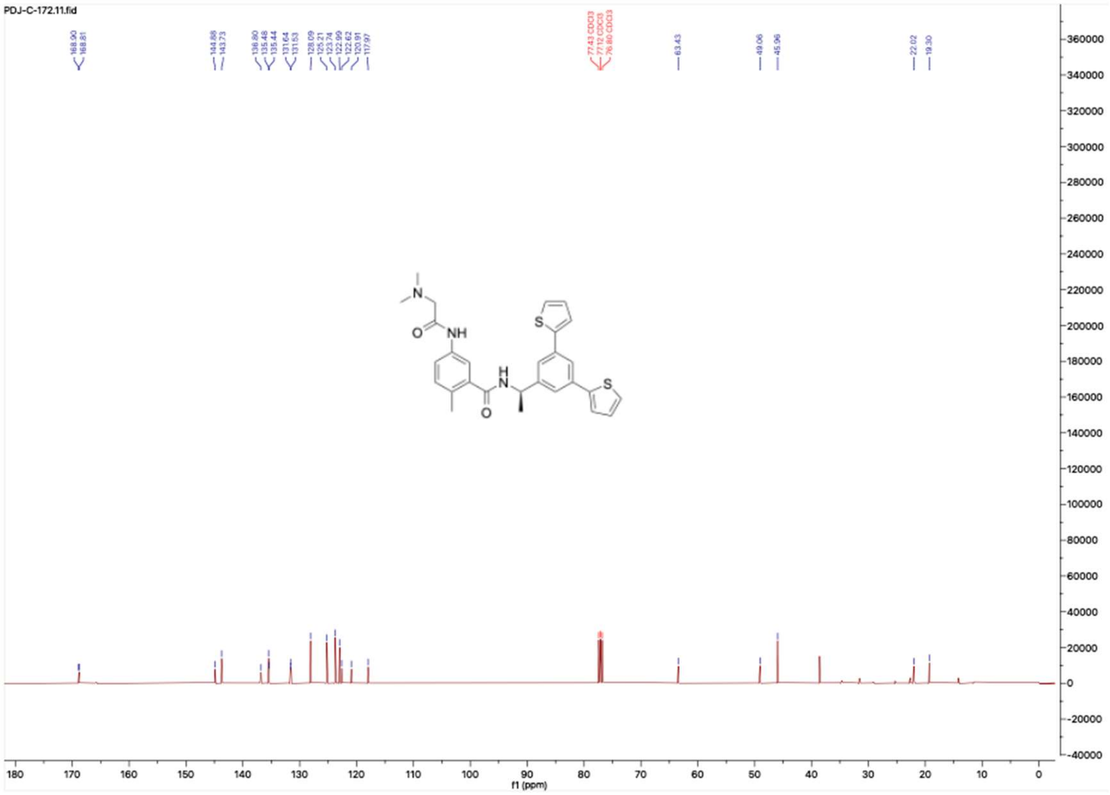

#### HNMR and CNMR spectra of Jun11945

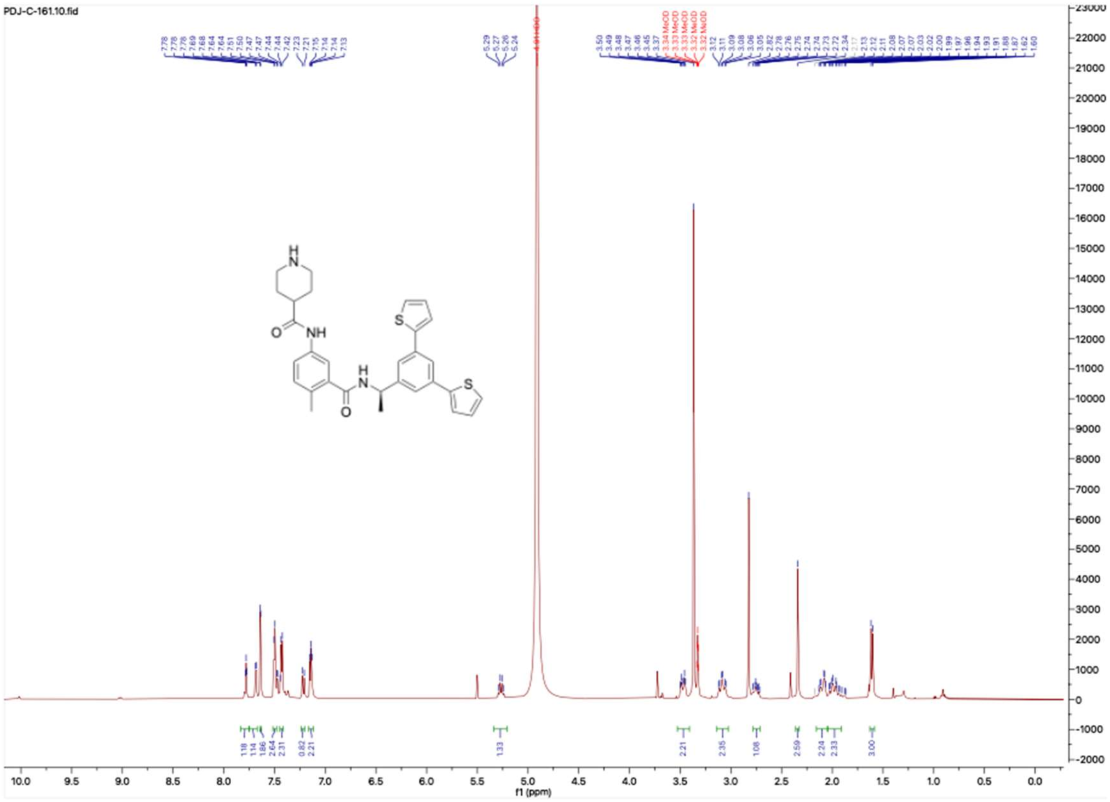


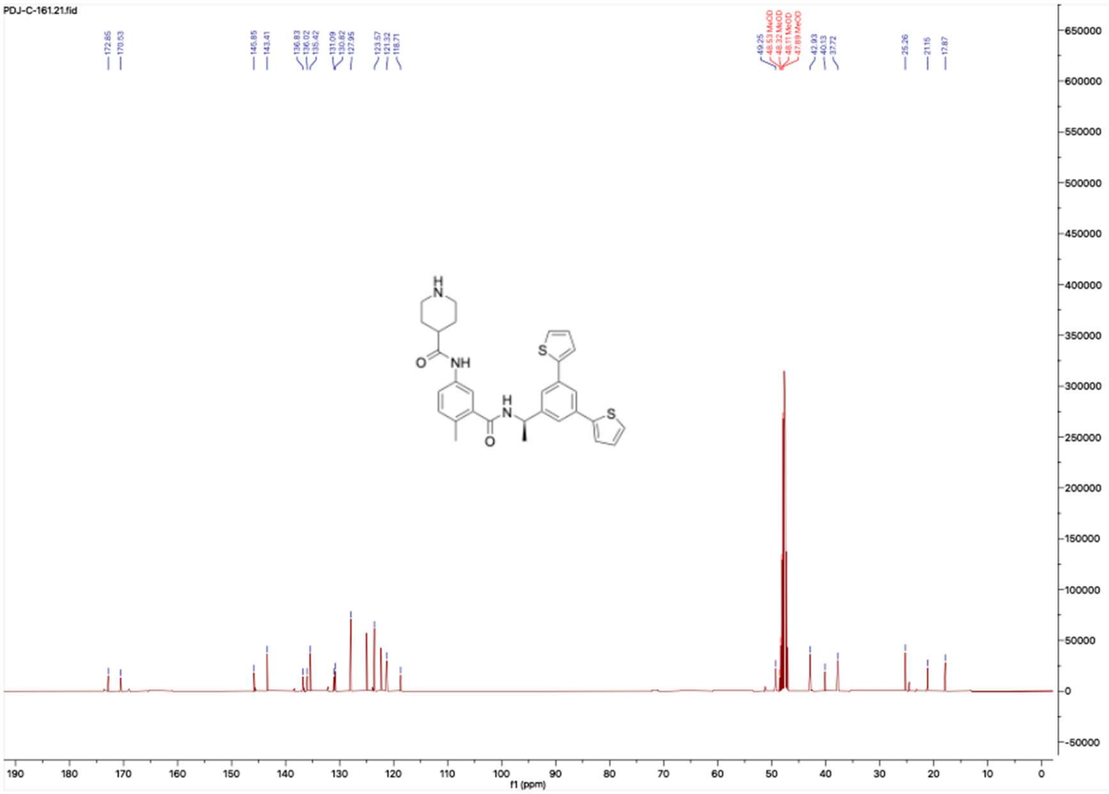

**HNMR and CNMR spectra of A-6**

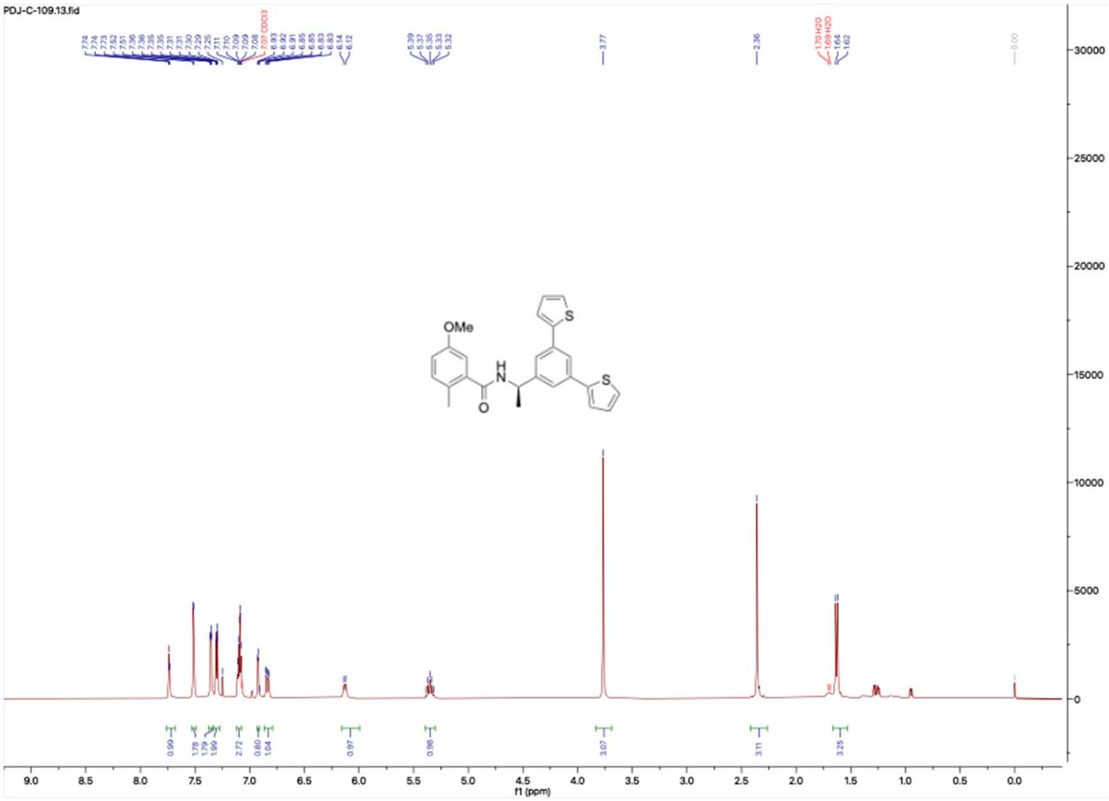

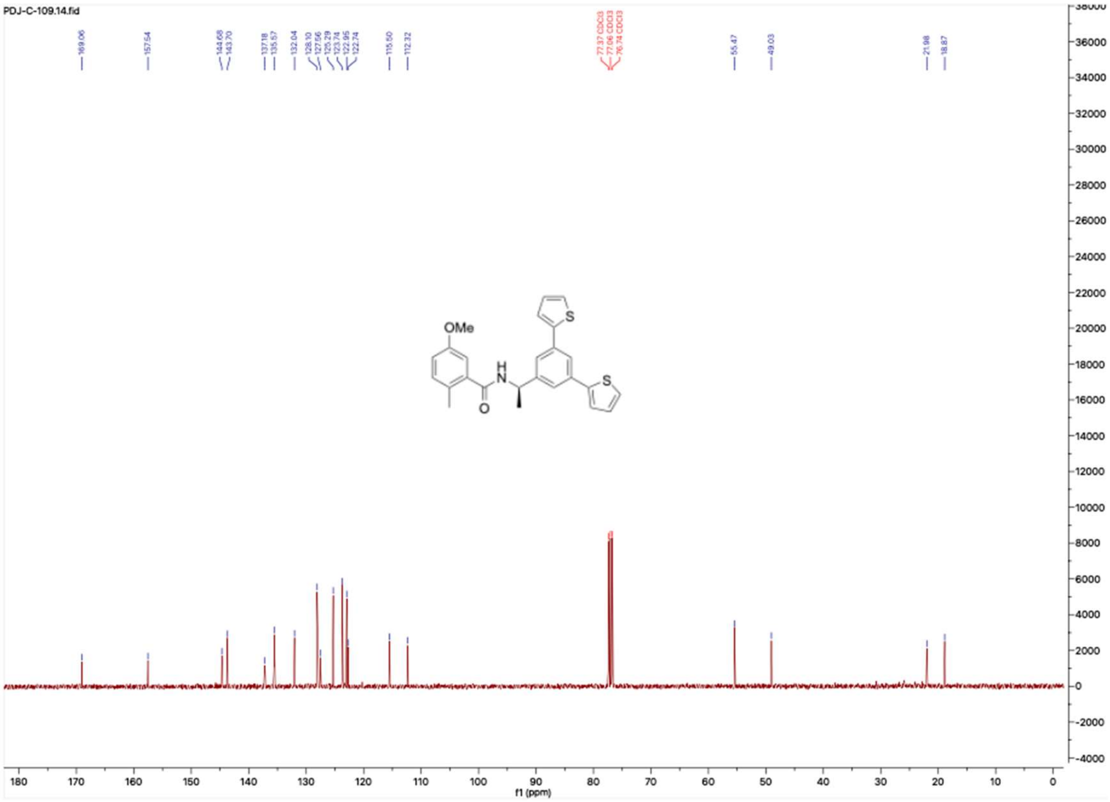

**HNMR and CNMR spectra of A-7**

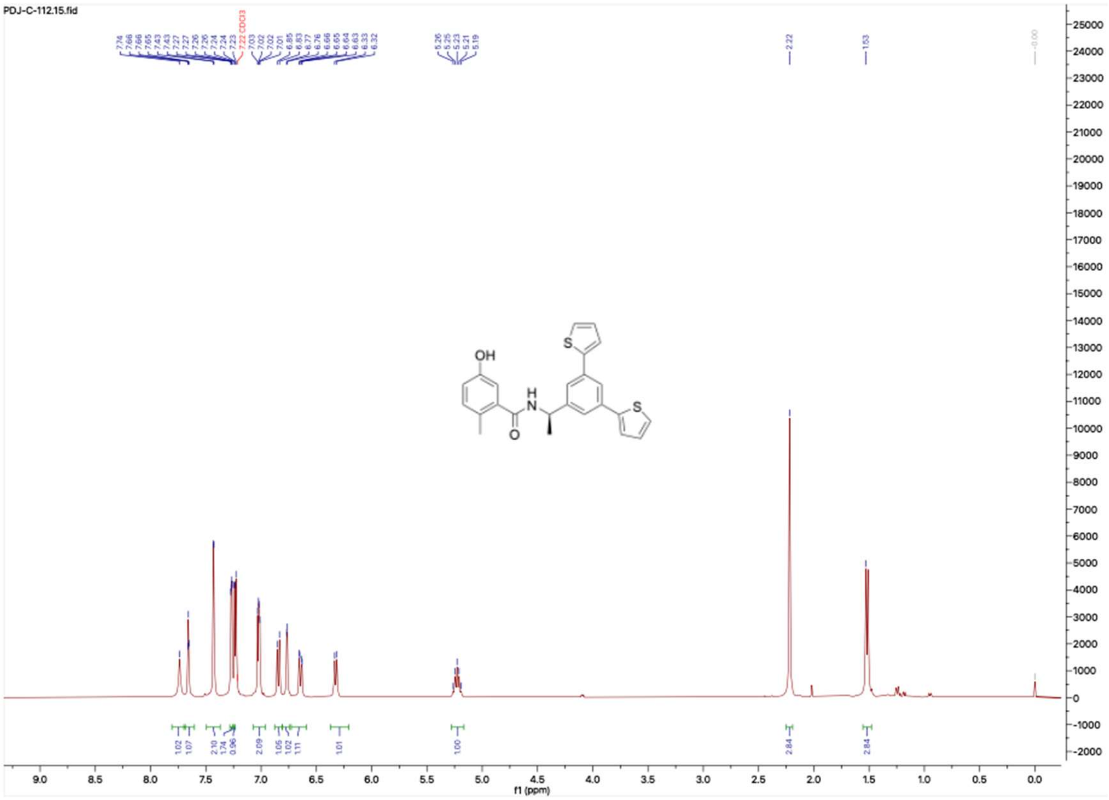


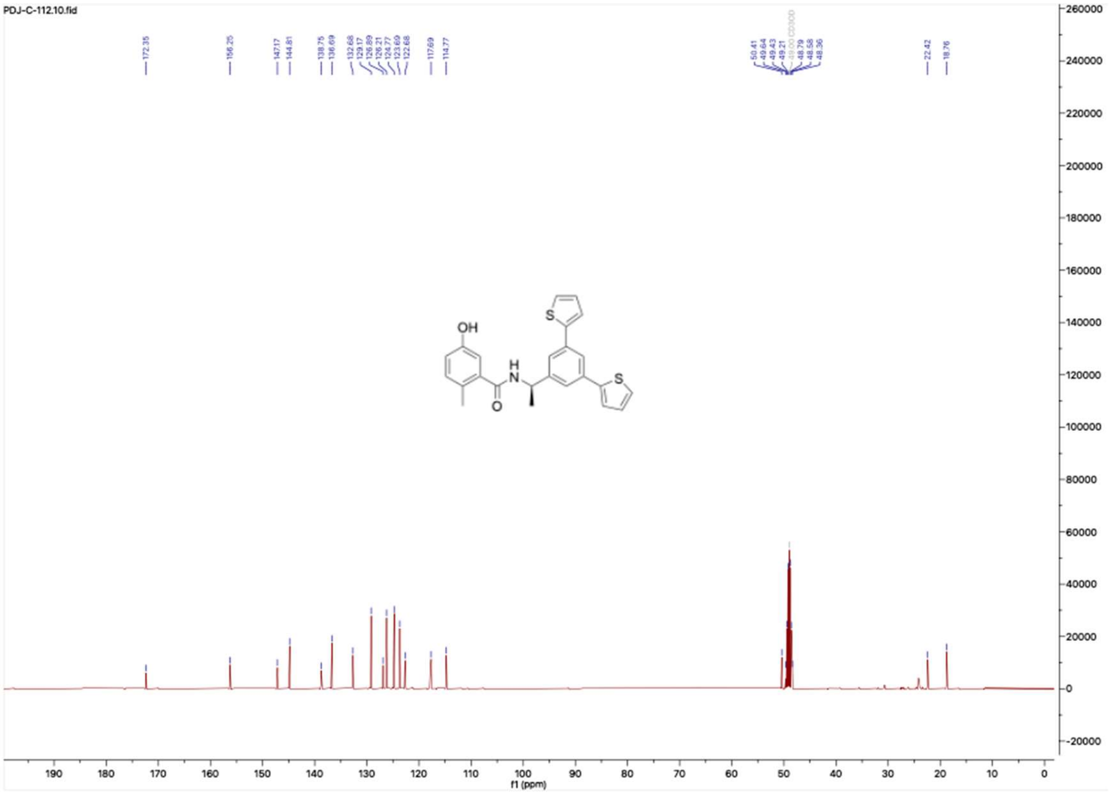

**HNMR and CNMR spectra of Jun11874**

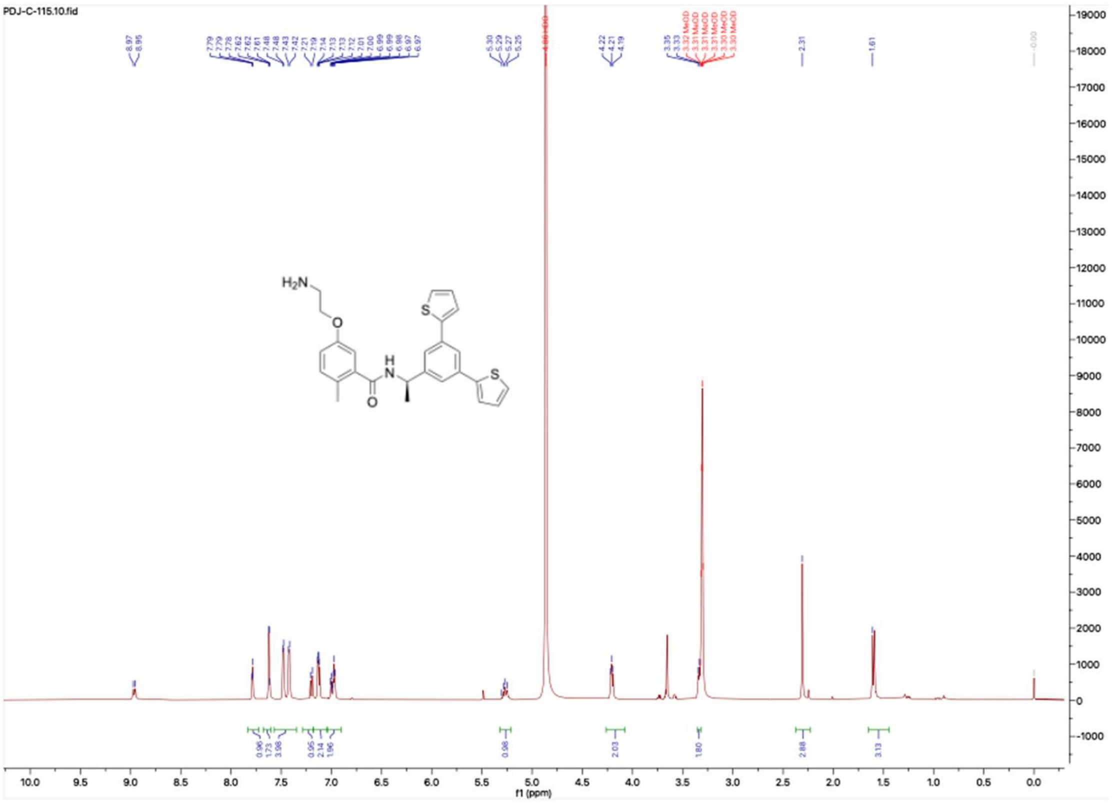

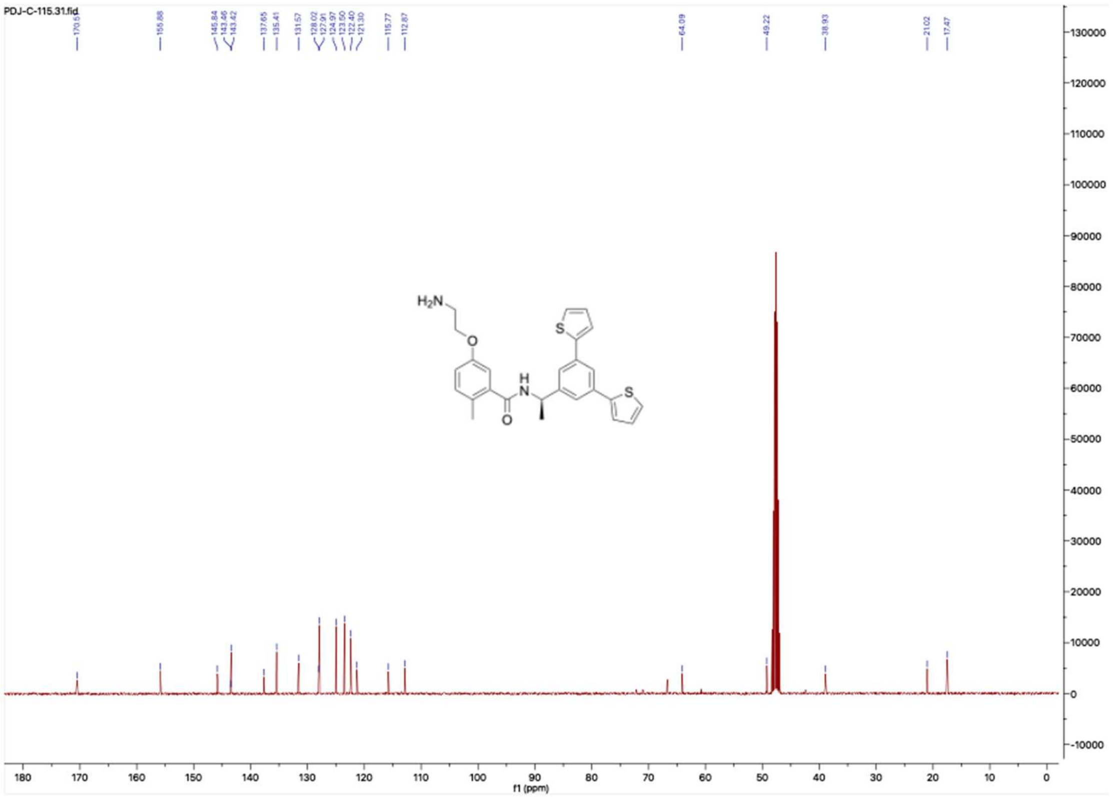

**HNMR and CNMR spectra of Jun11875**

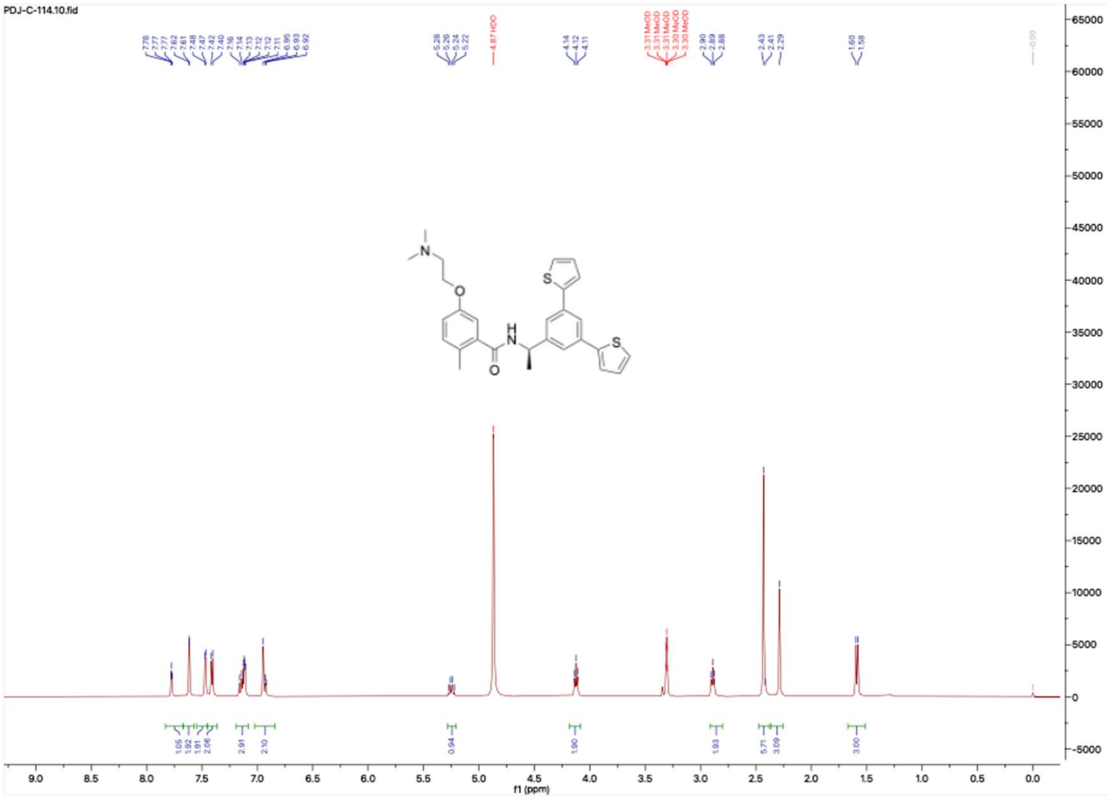


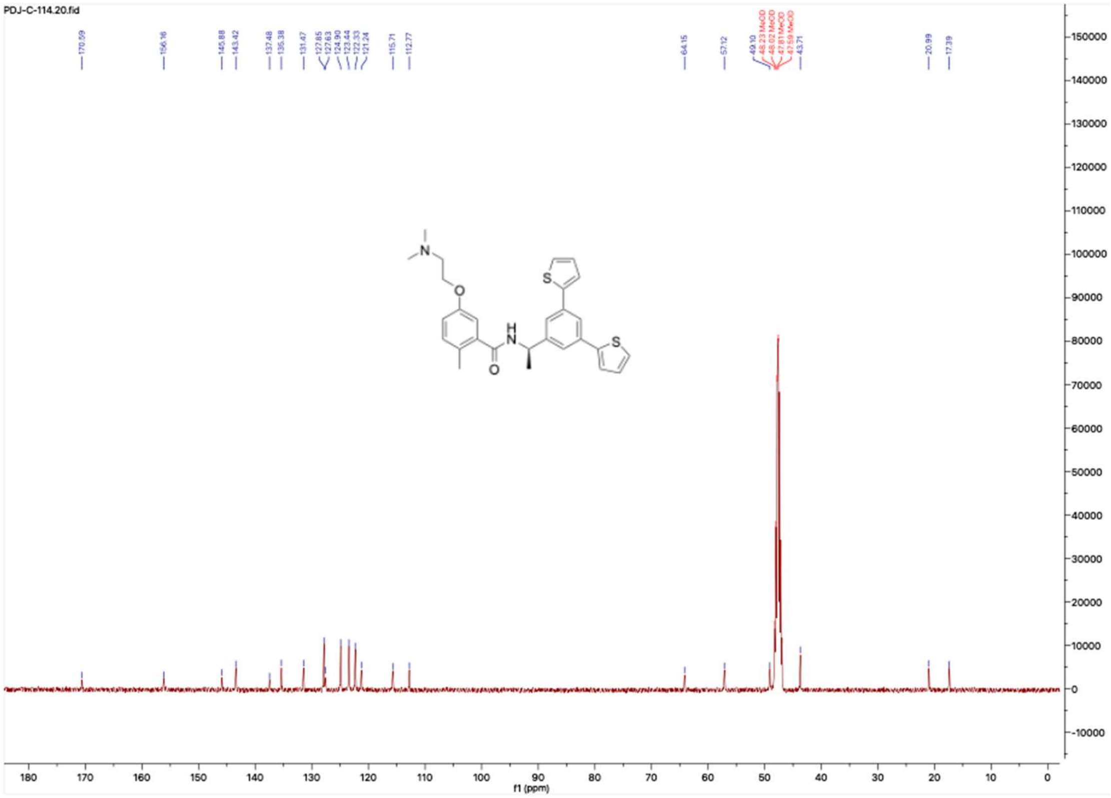

**HNMR and CNMR spectra of Jun11898**

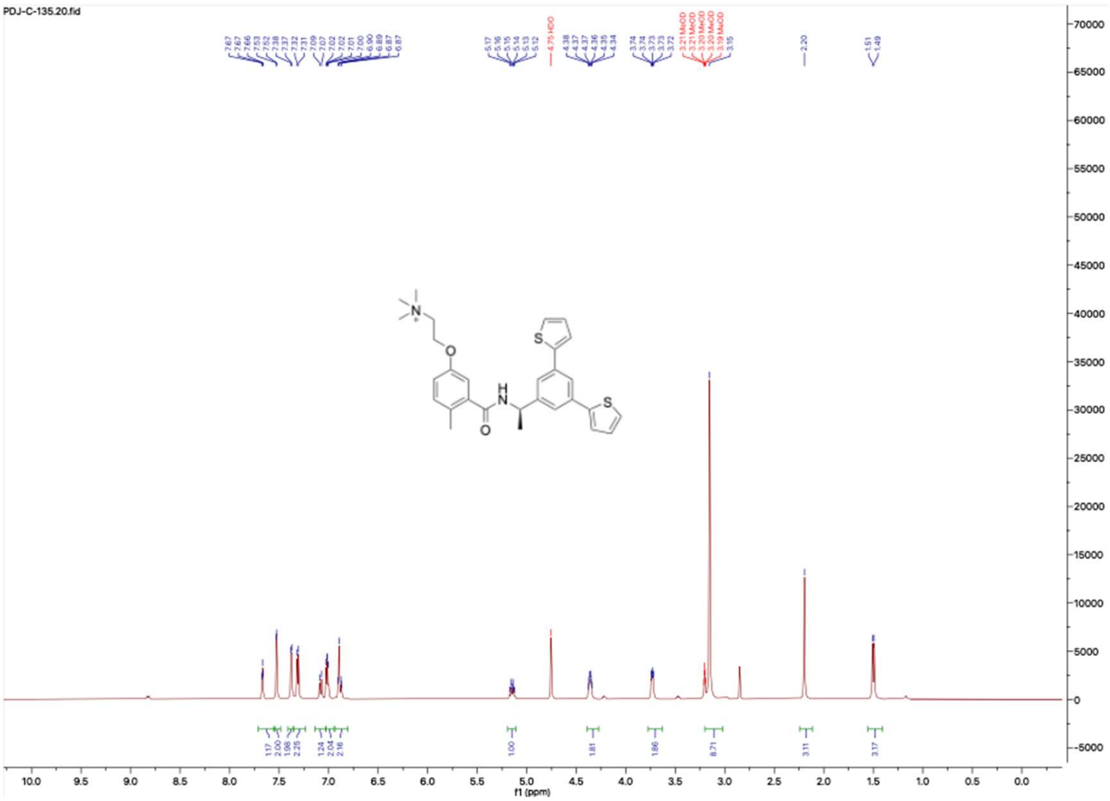

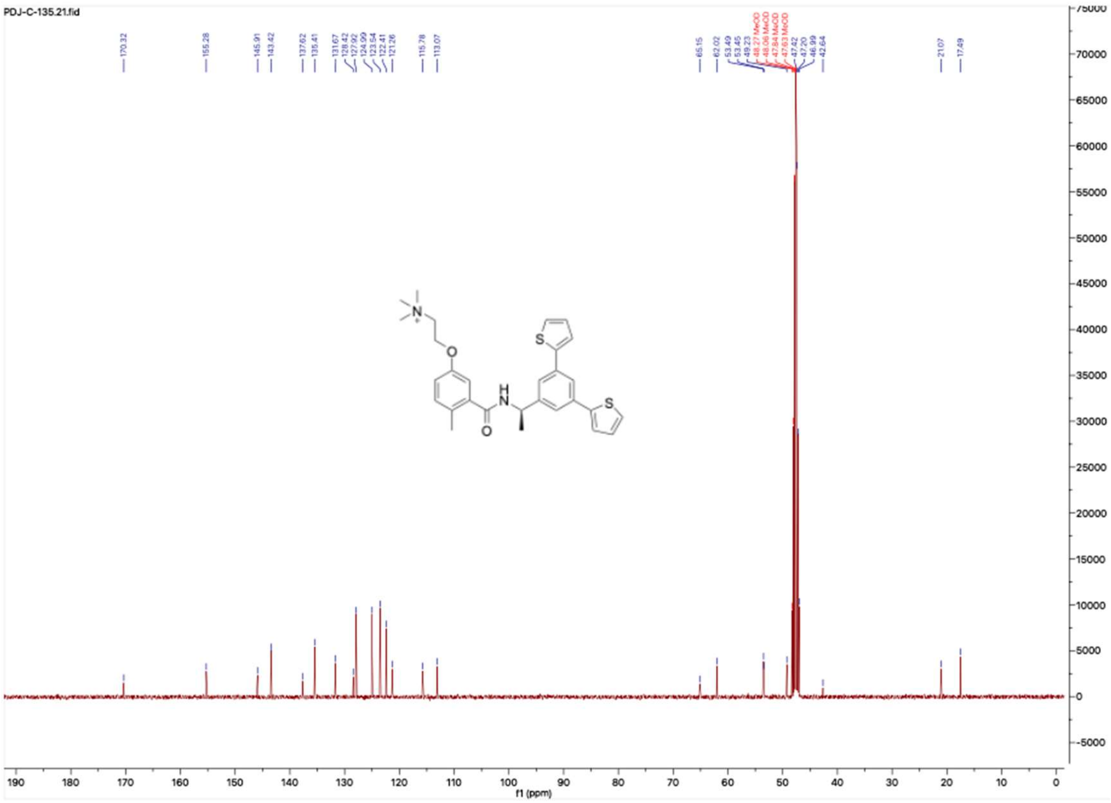

**HNMR and CNMR spectra of Jun11899**

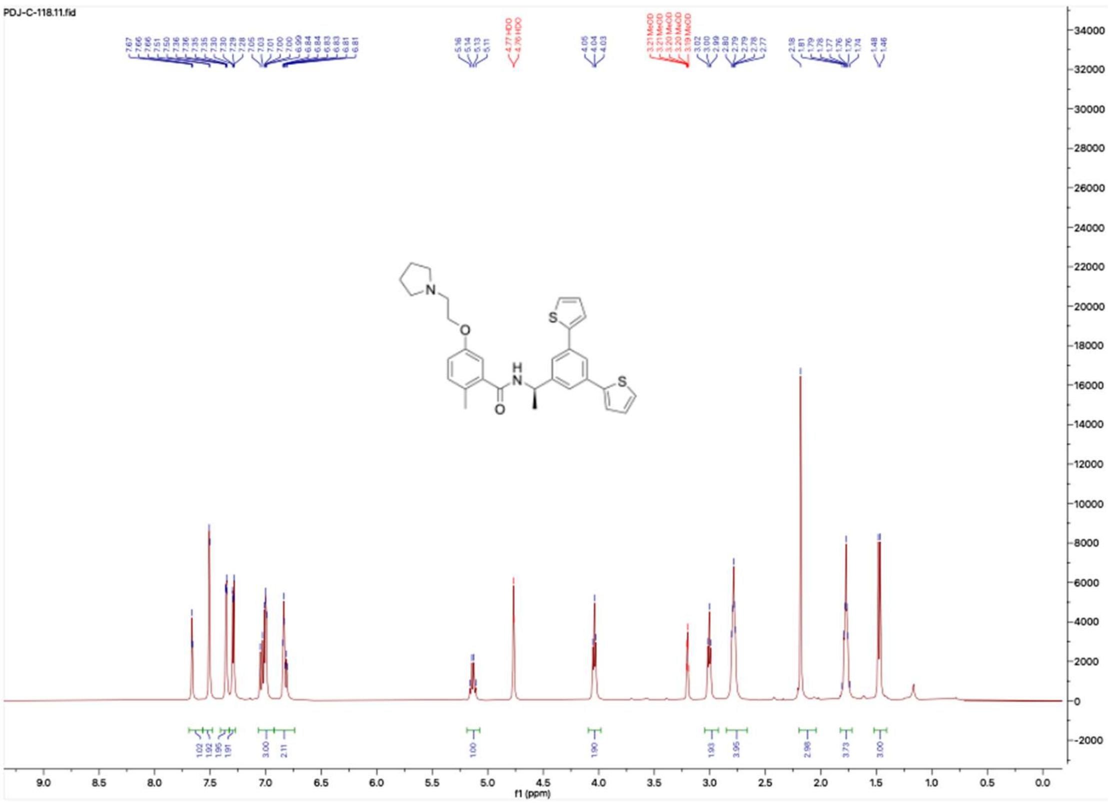


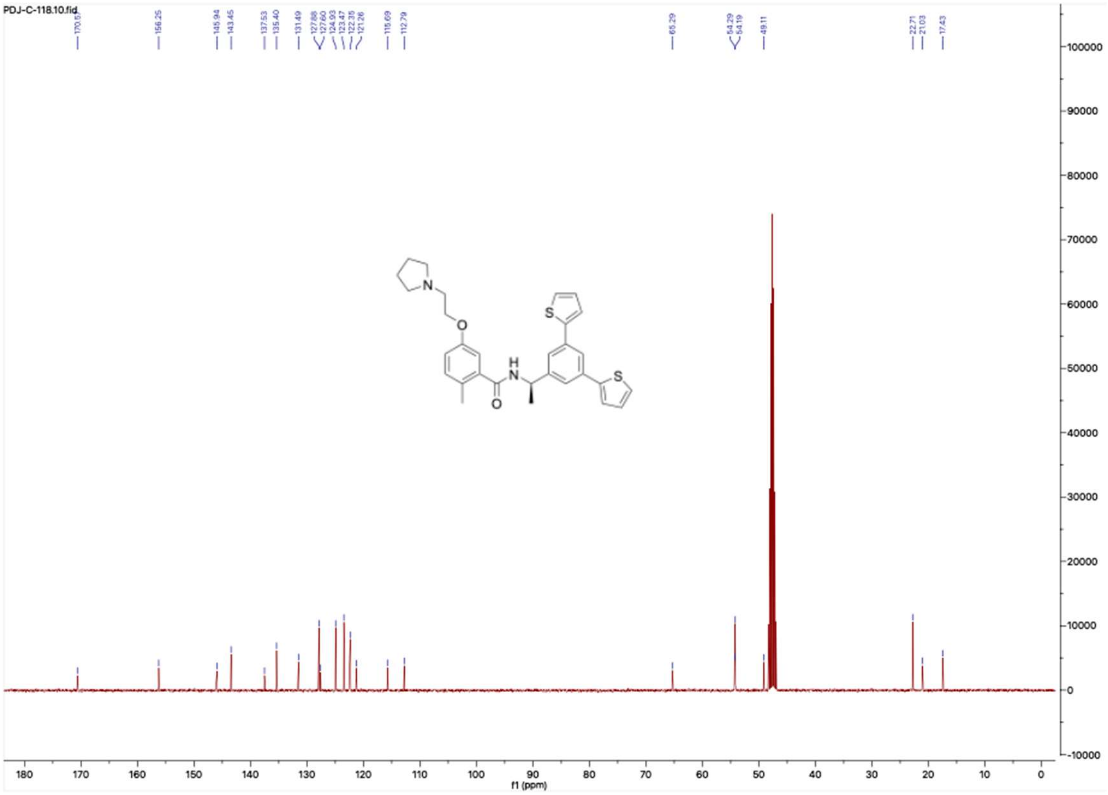

**HNMR and CNMR spectra of Jun118910**

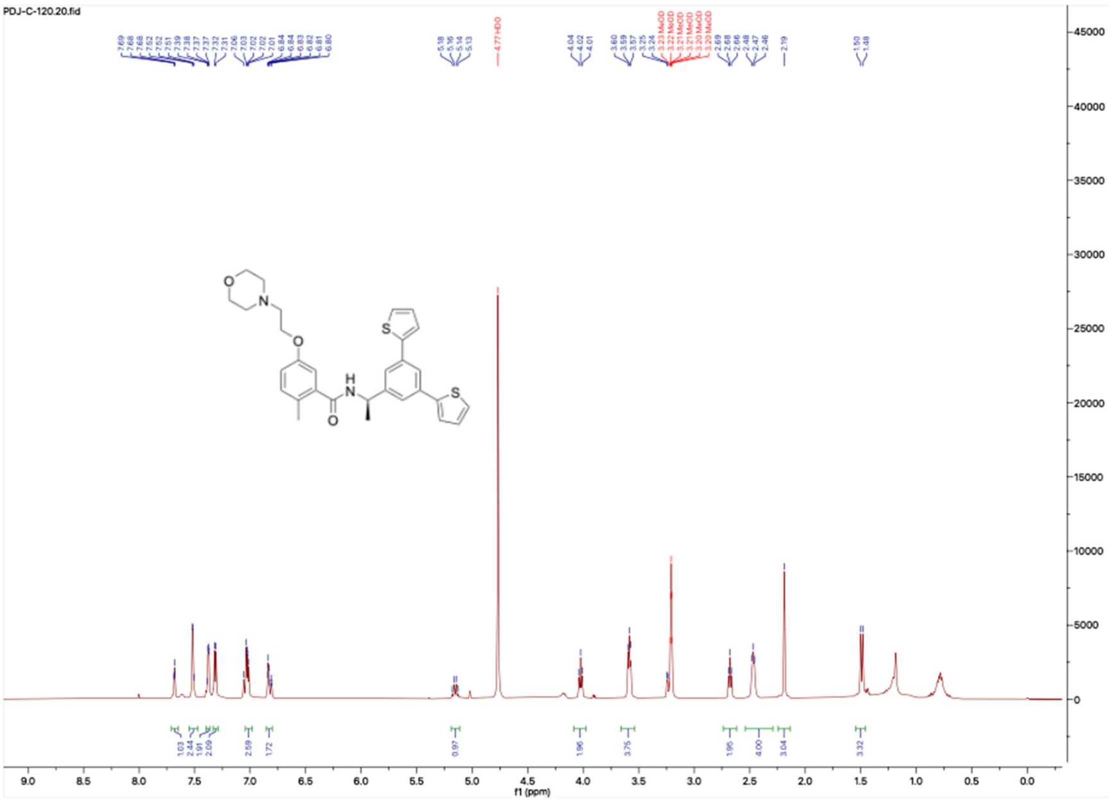

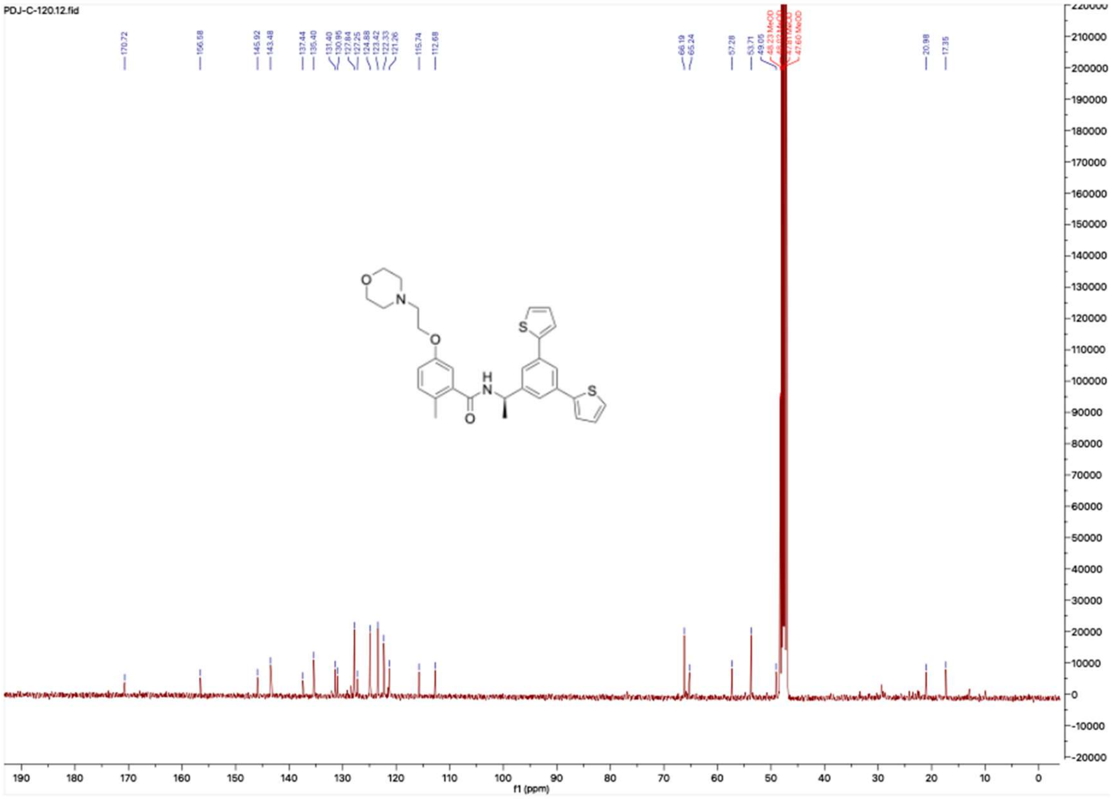

**HNMR and CNMR spectra of B-2**

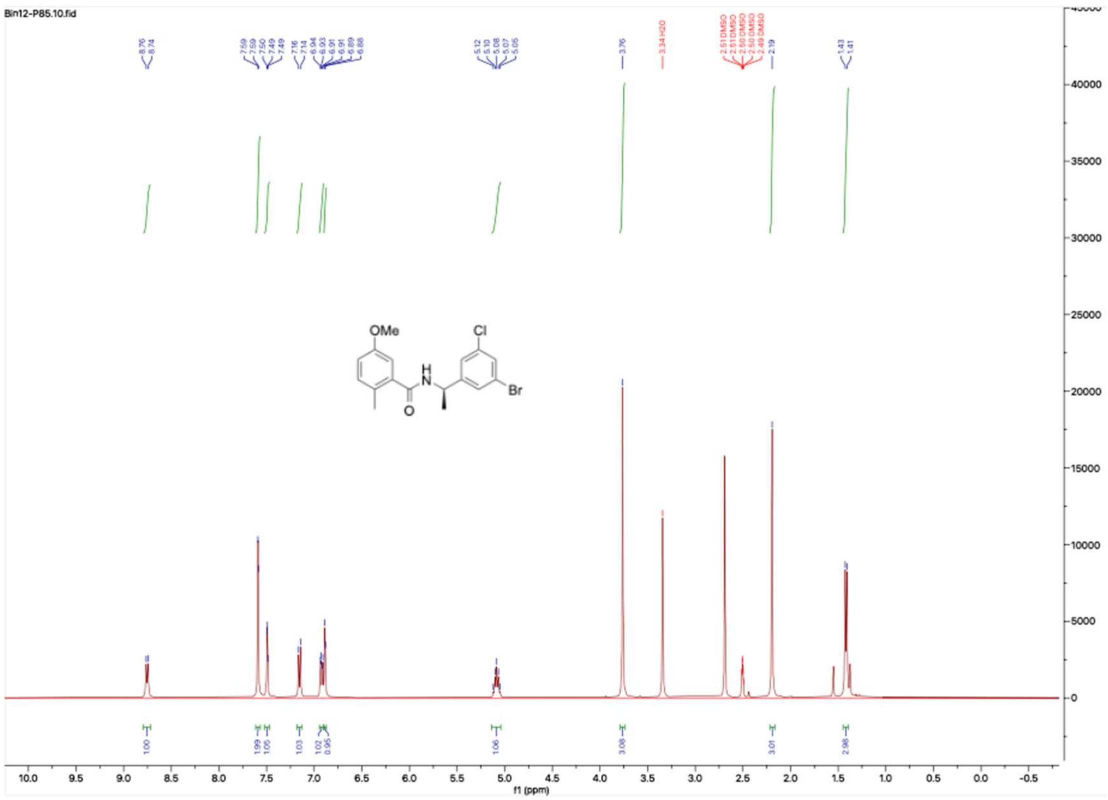

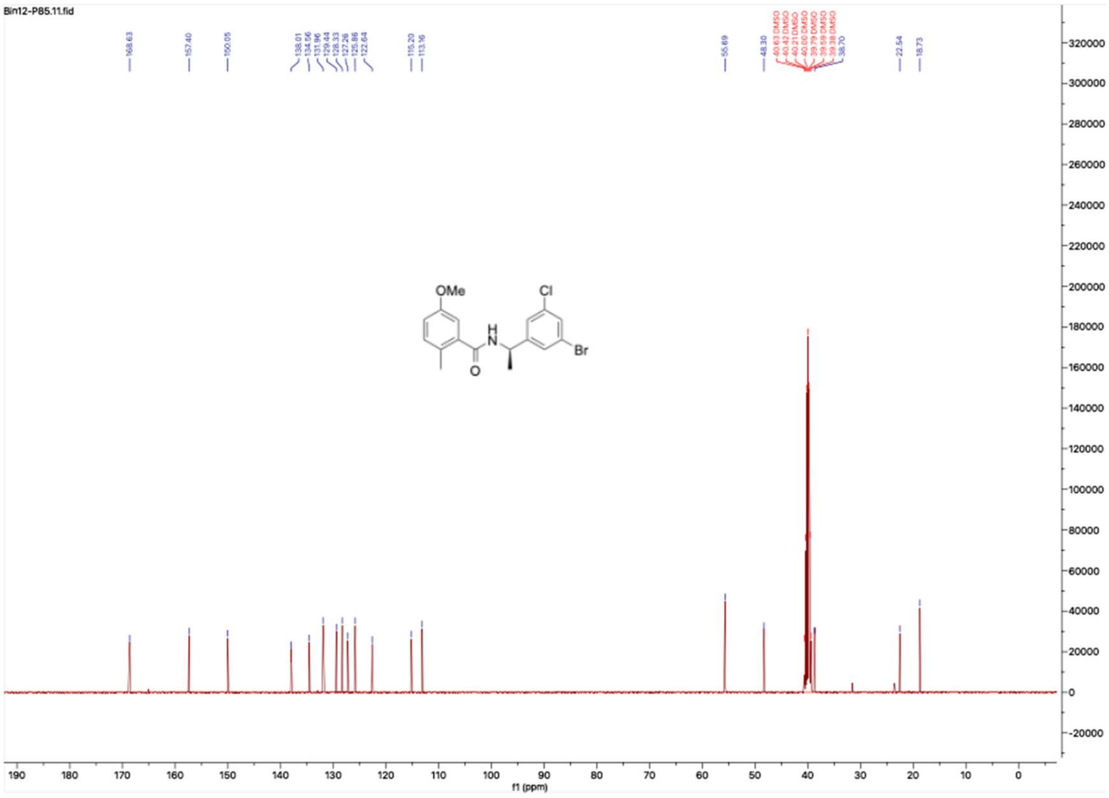

**HNMR and CNMR spectra of B-3**

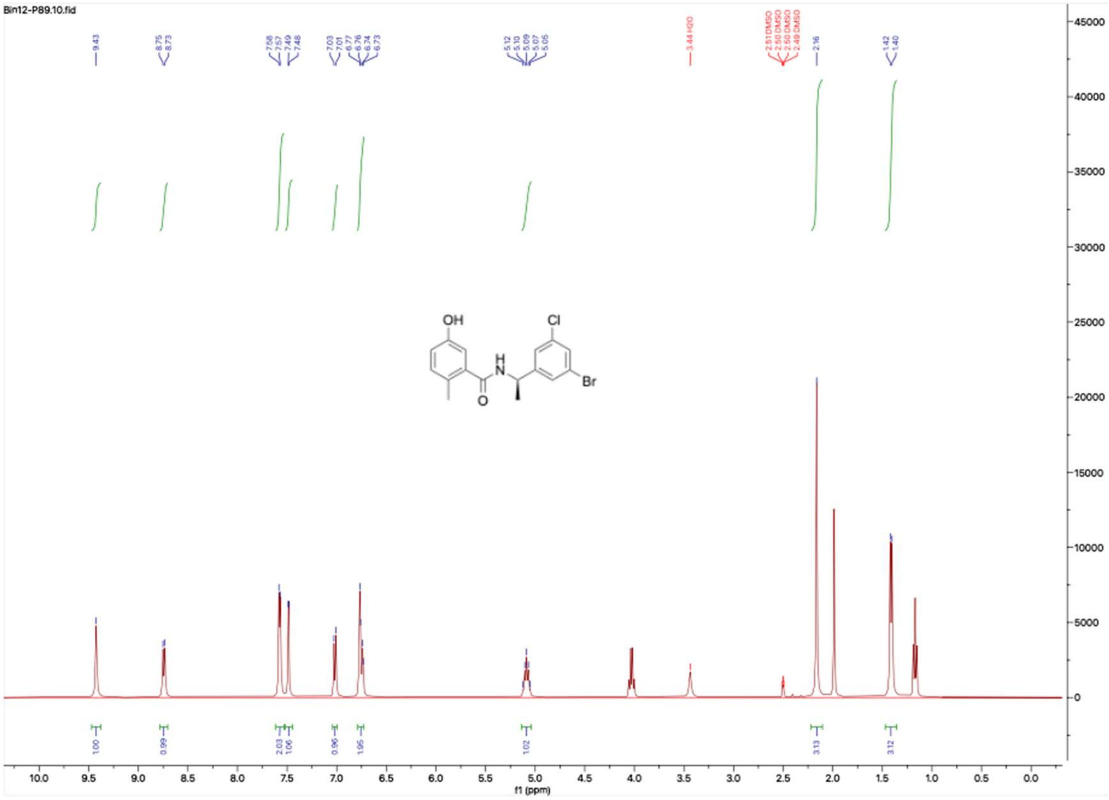


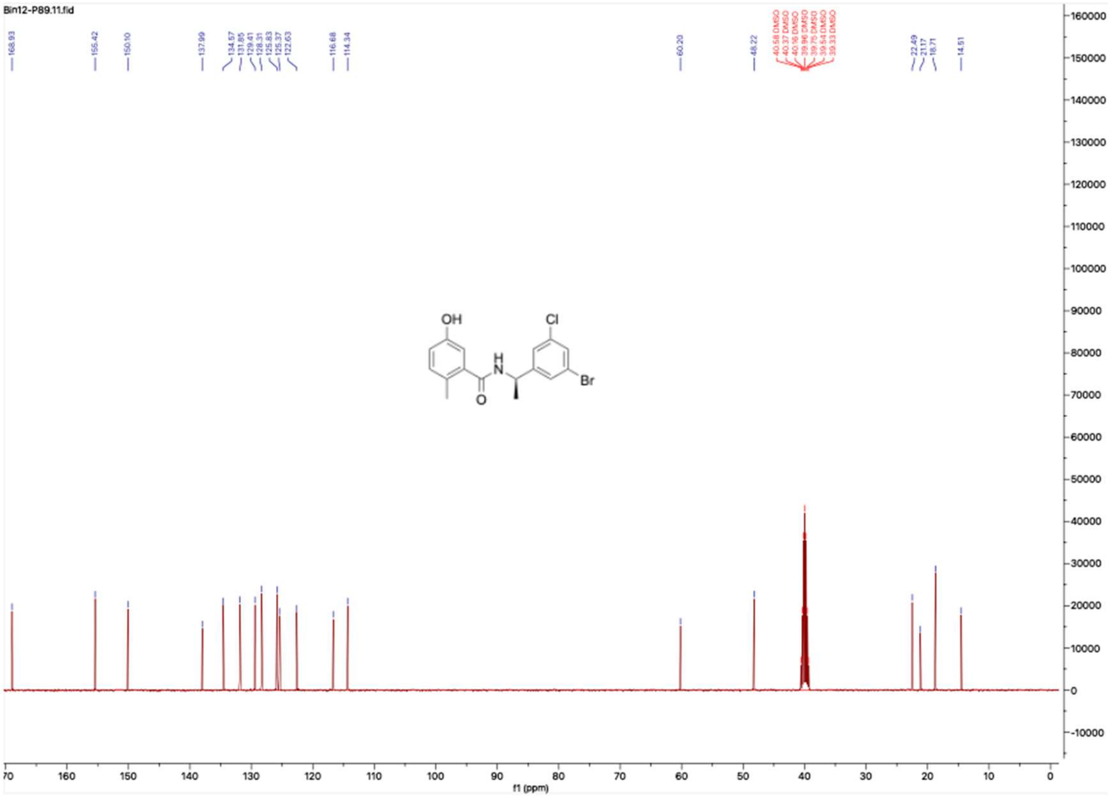

#### HNMR and CNMR spectra of B-4

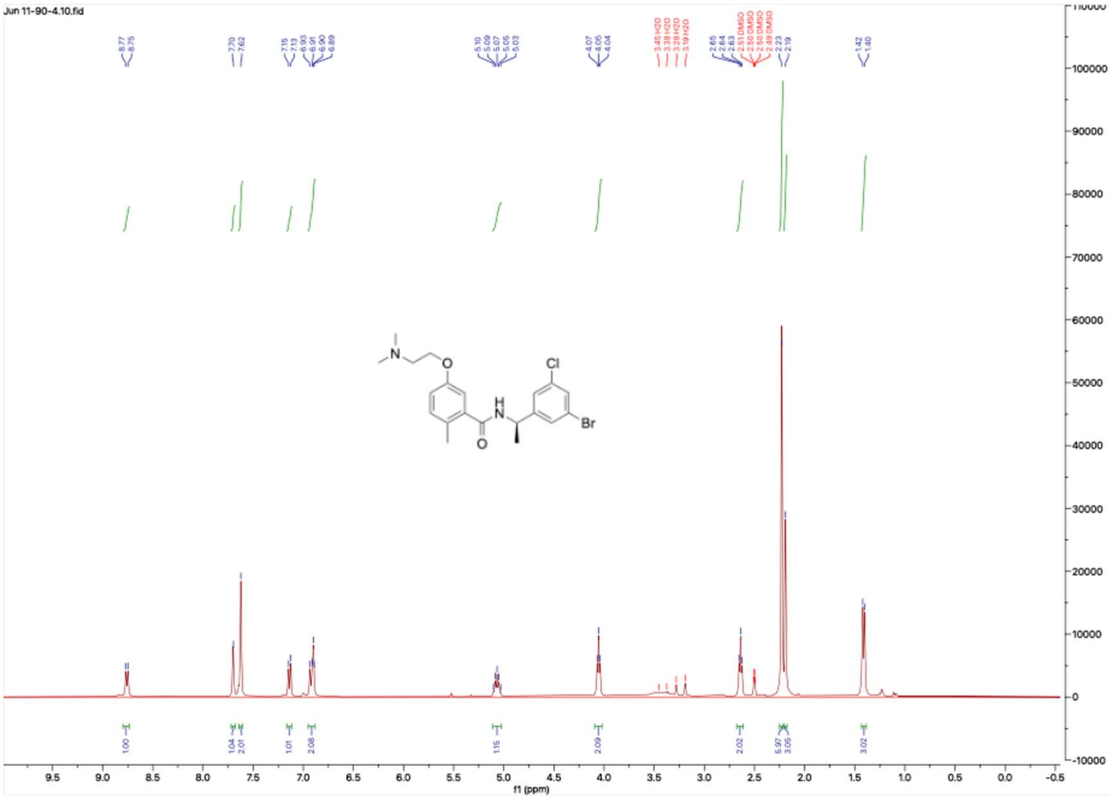

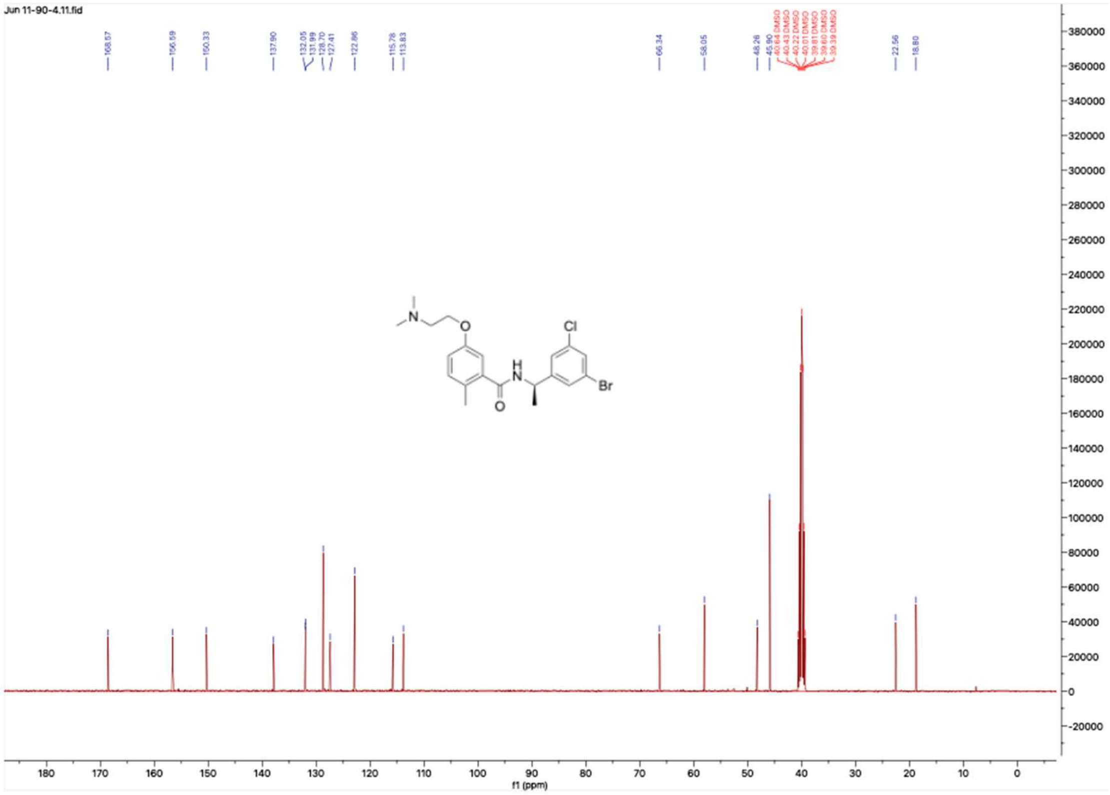

**HNMR and CNMR spectra of Jun11931**

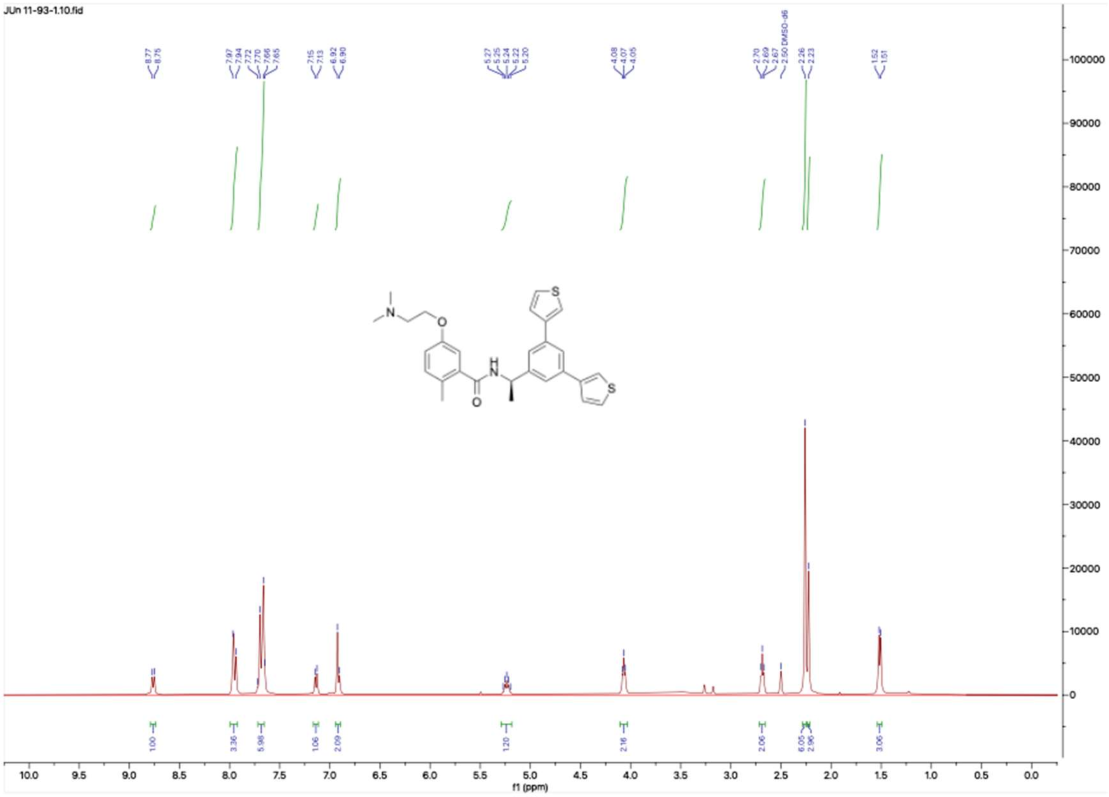


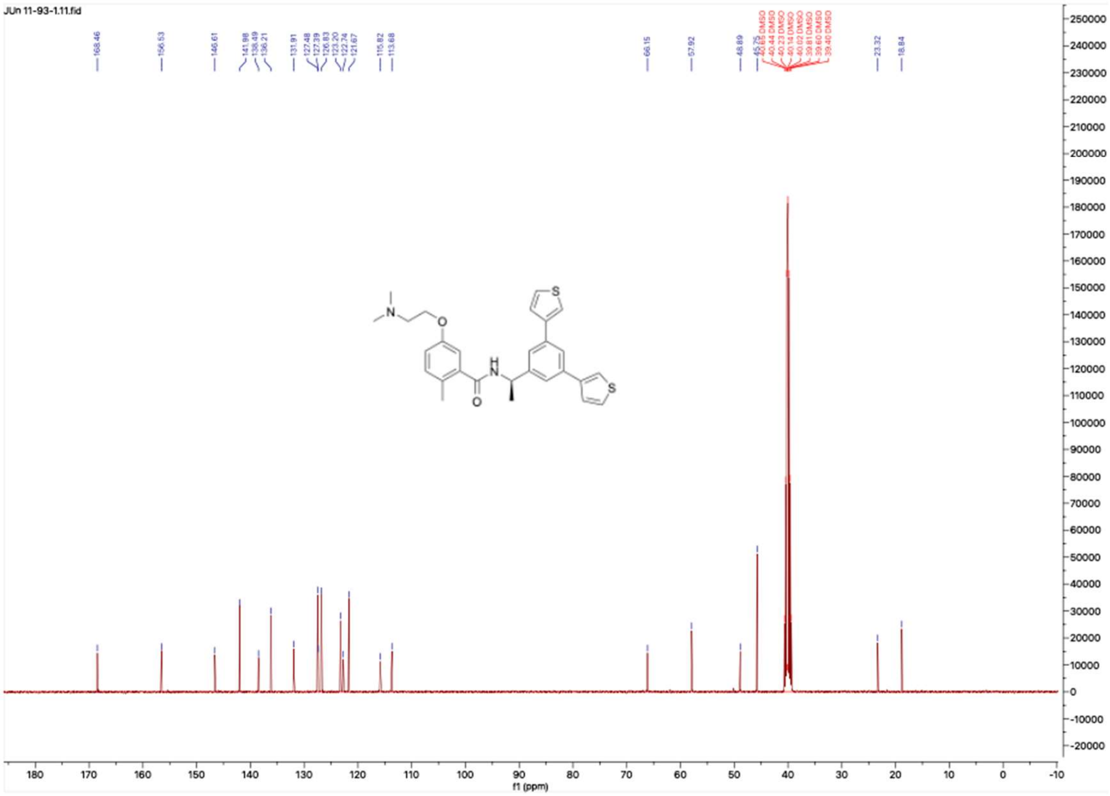

**HNMR and CNMR spectra of Jun119415**

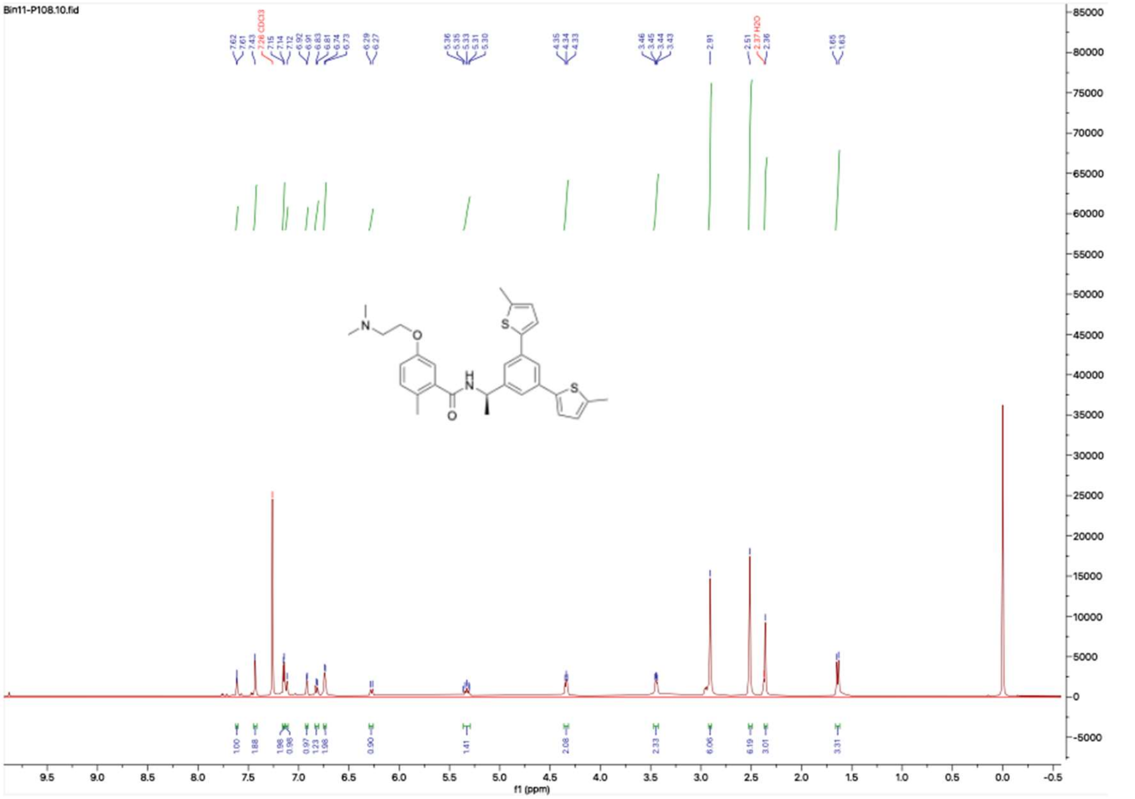


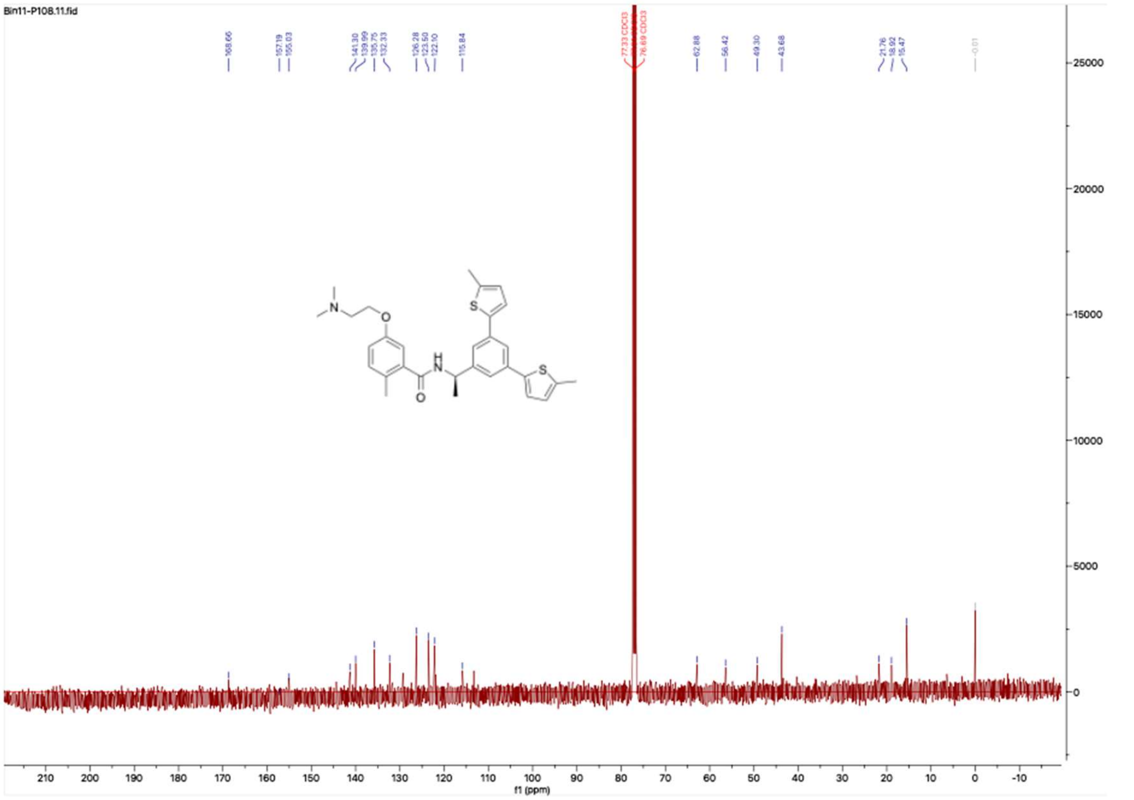

**HNMR and CNMR spectra of Jun11903**

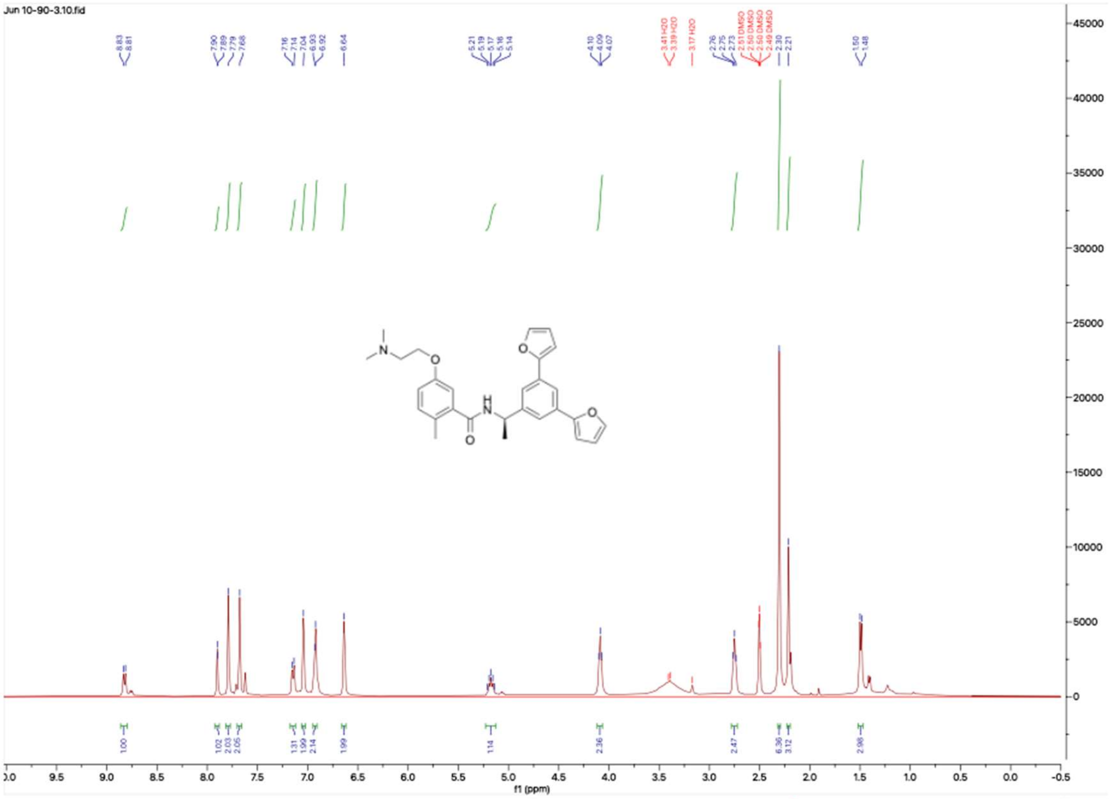

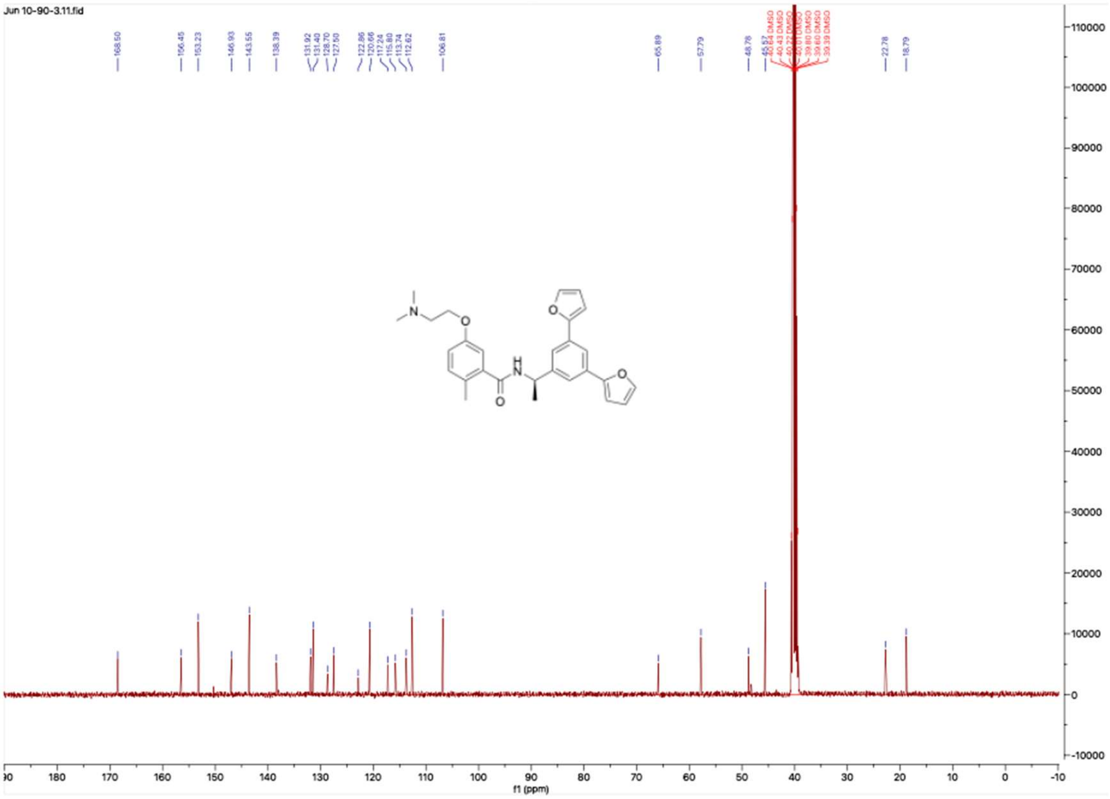

**HNMR and CNMR spectra of Jun11932**

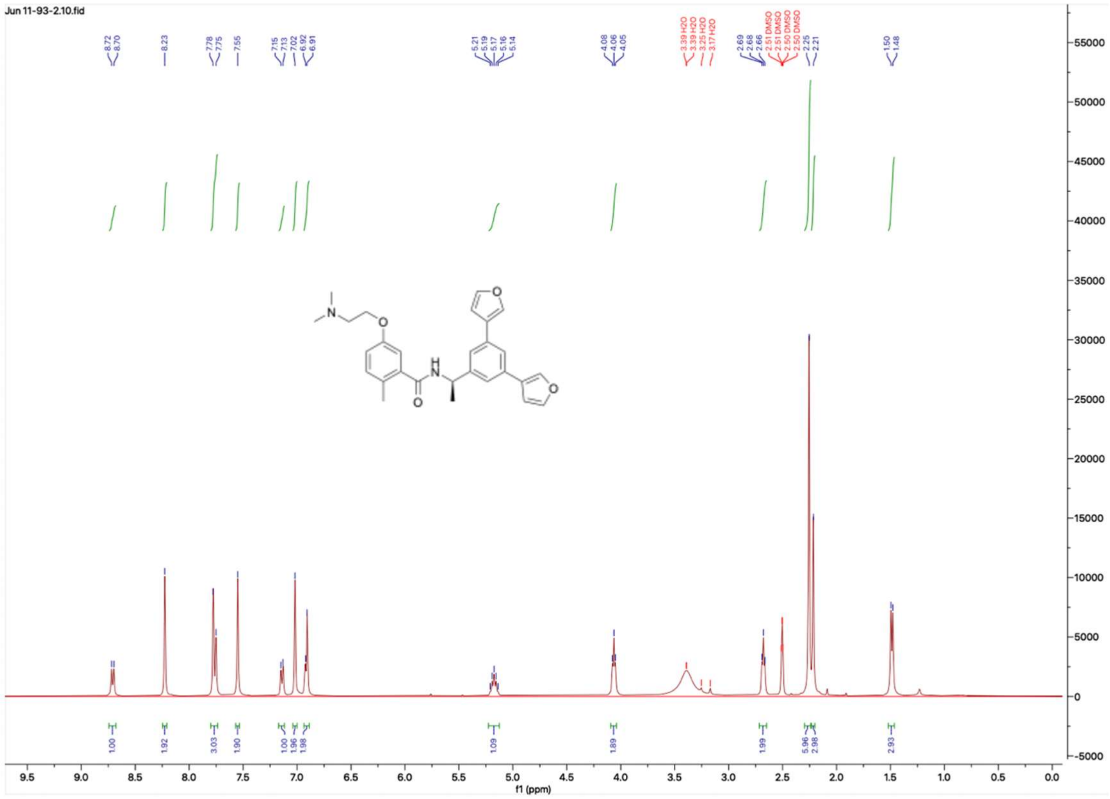


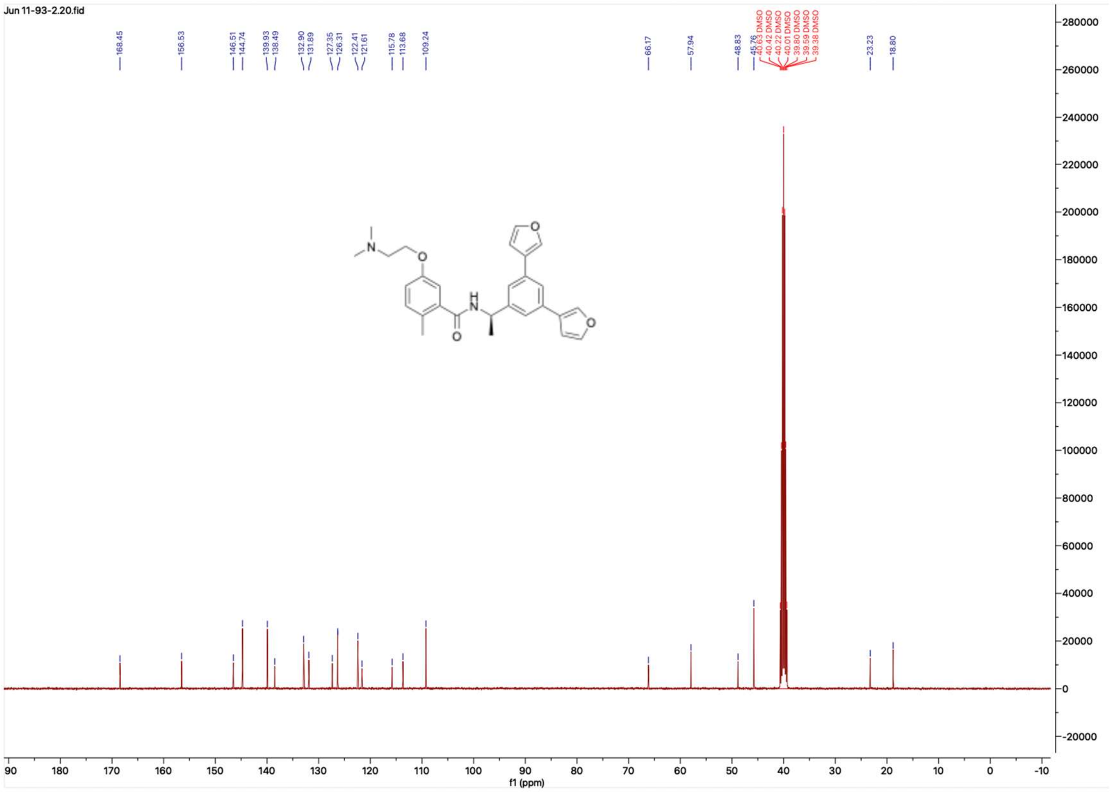

**HNMR and CNMR spectra of Jun11934**

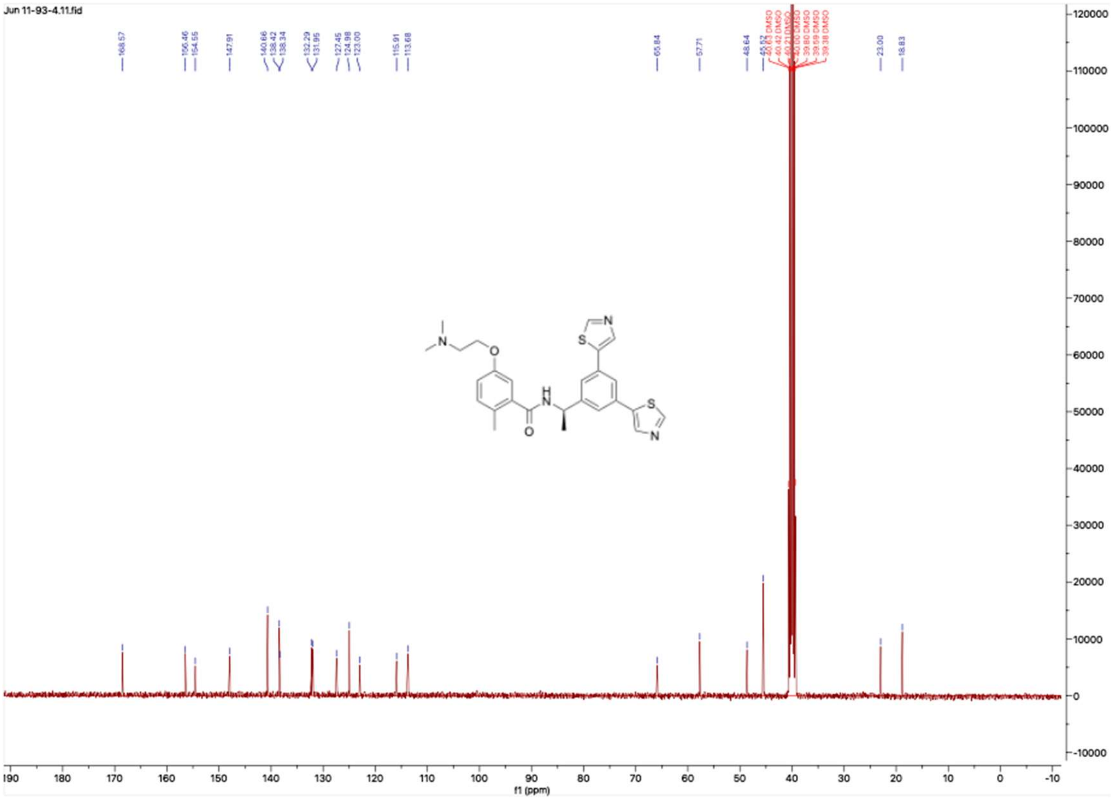

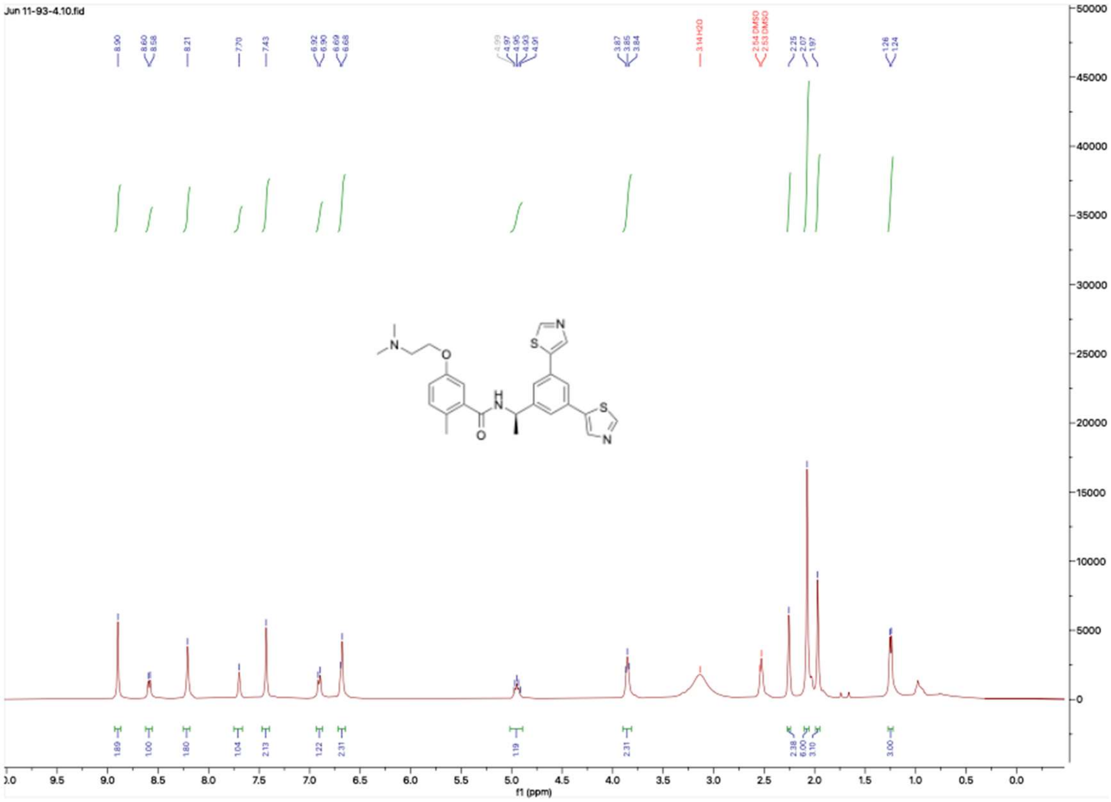

**HNMR and CNMR spectra of Jun11933**

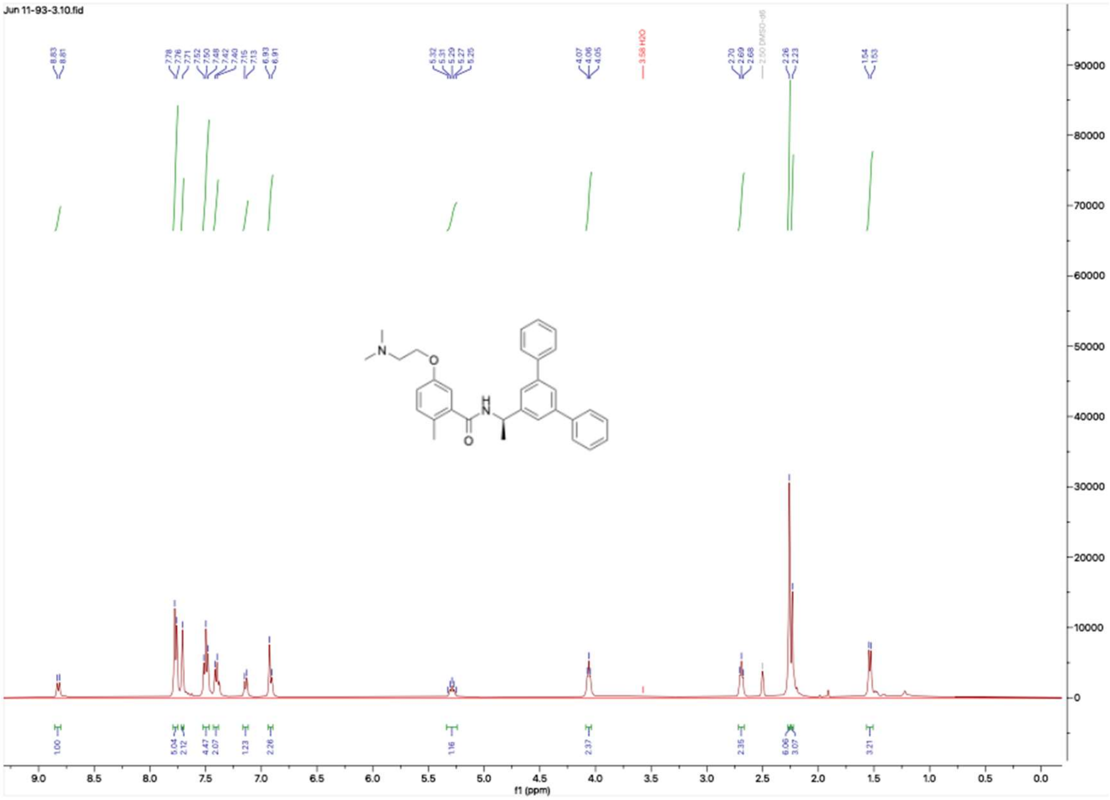

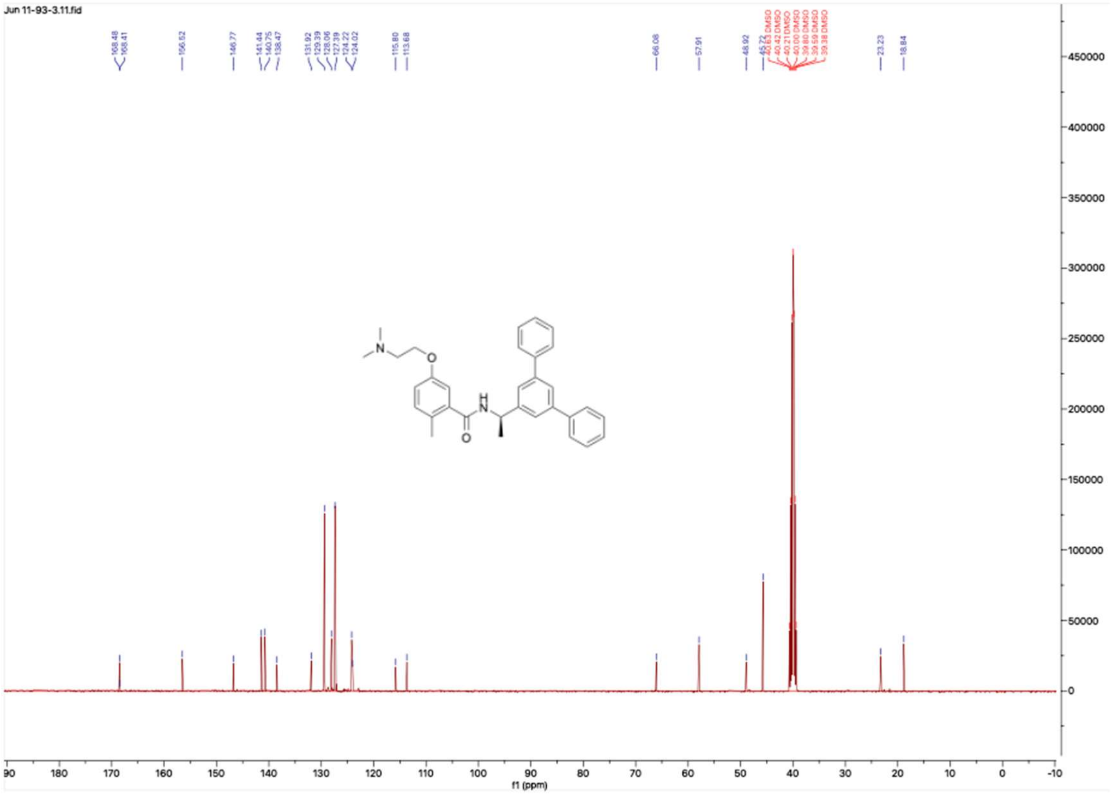

**HNMR and CNMR spectra of Jun11943**

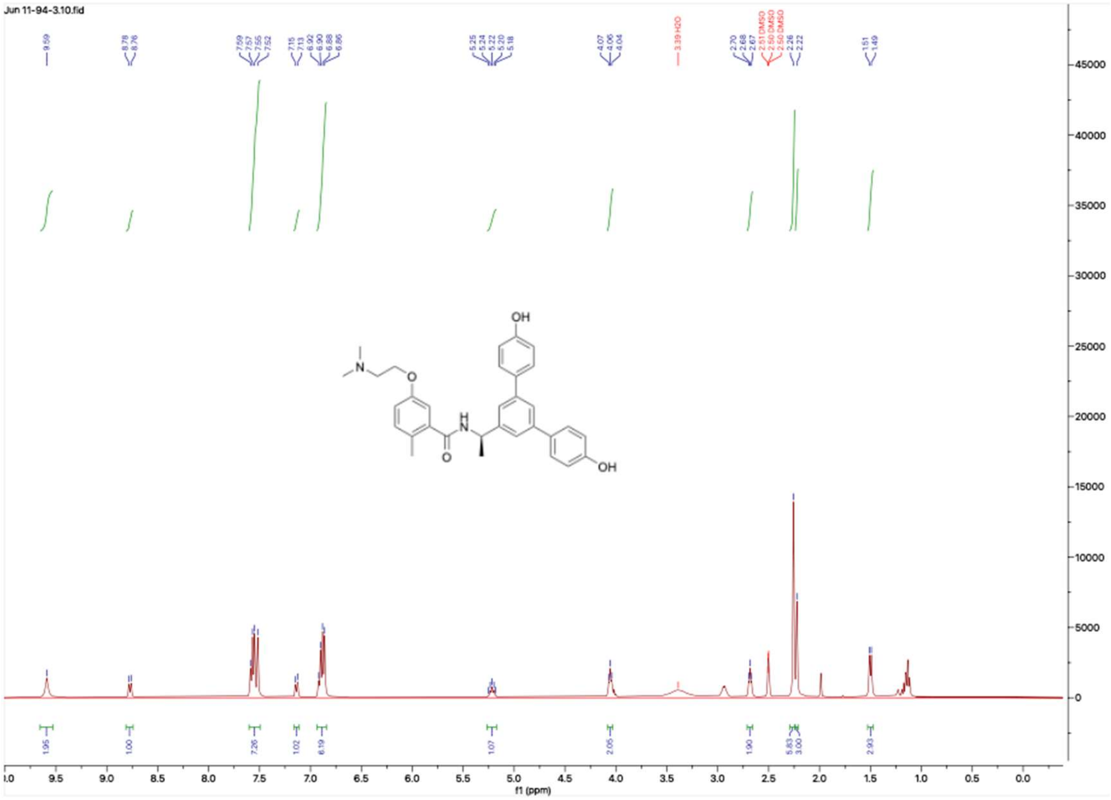


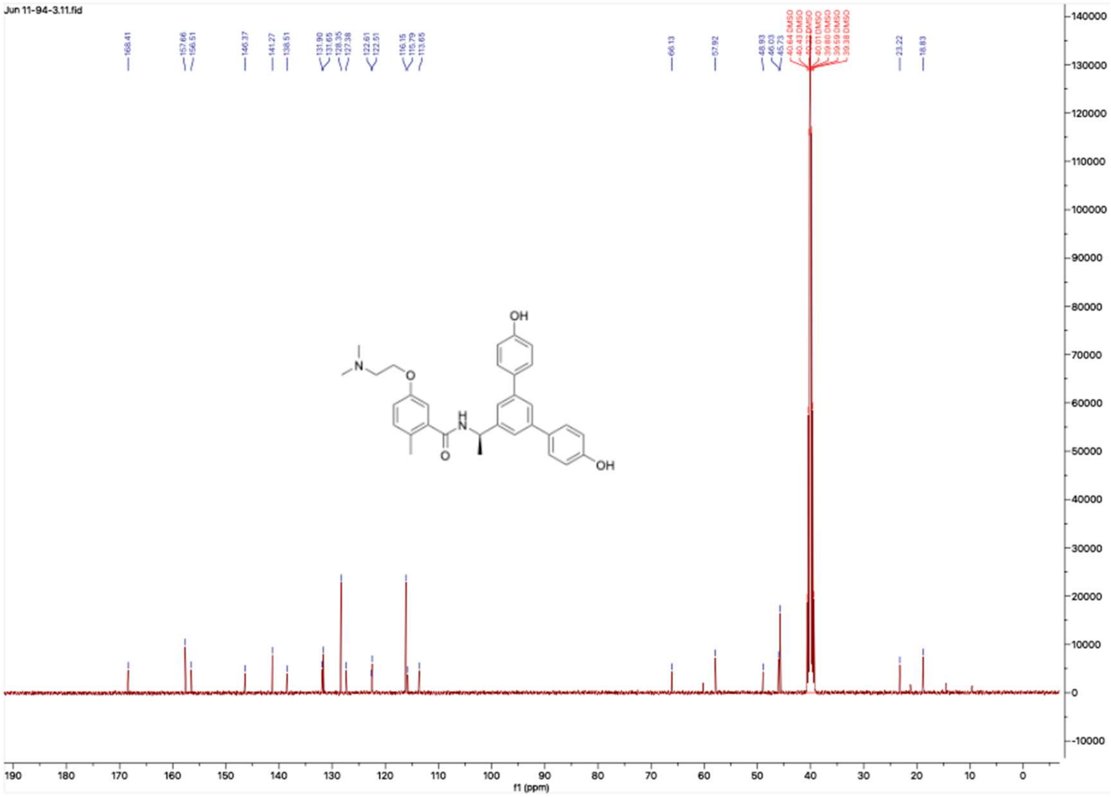

**HNMR and CNMR spectra of Jun1218**

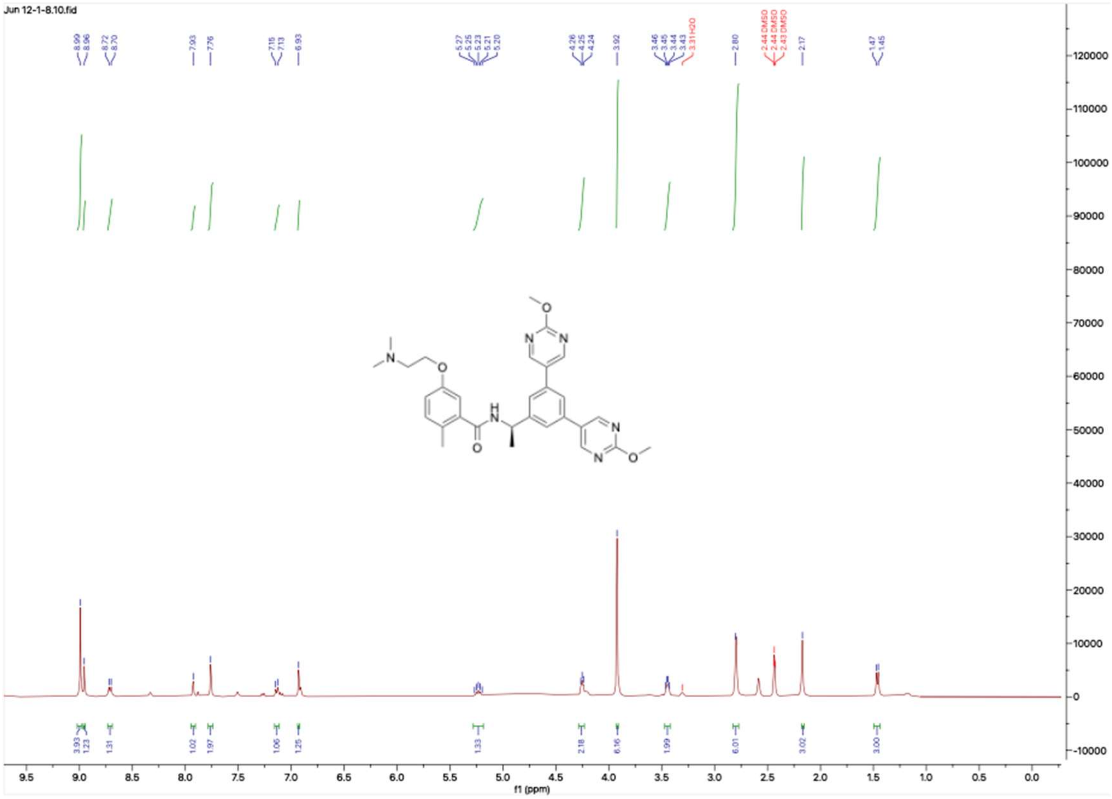

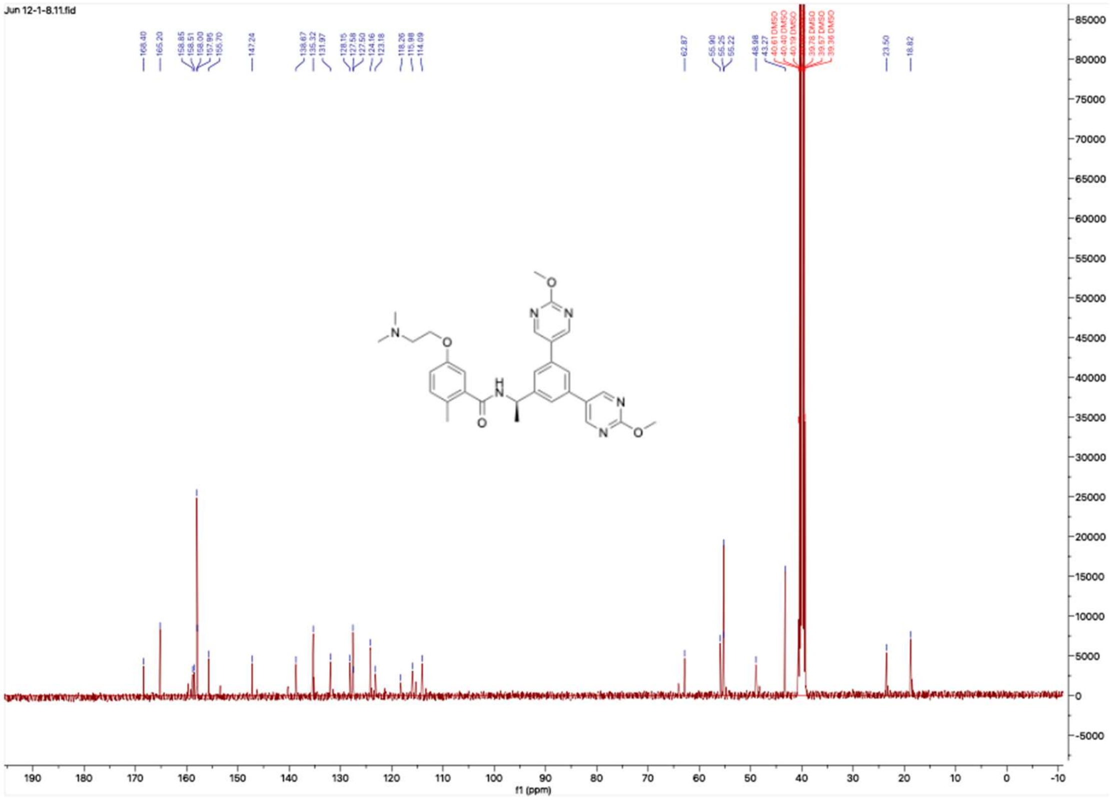

**HNMR and CNMR spectra of Jun11941**

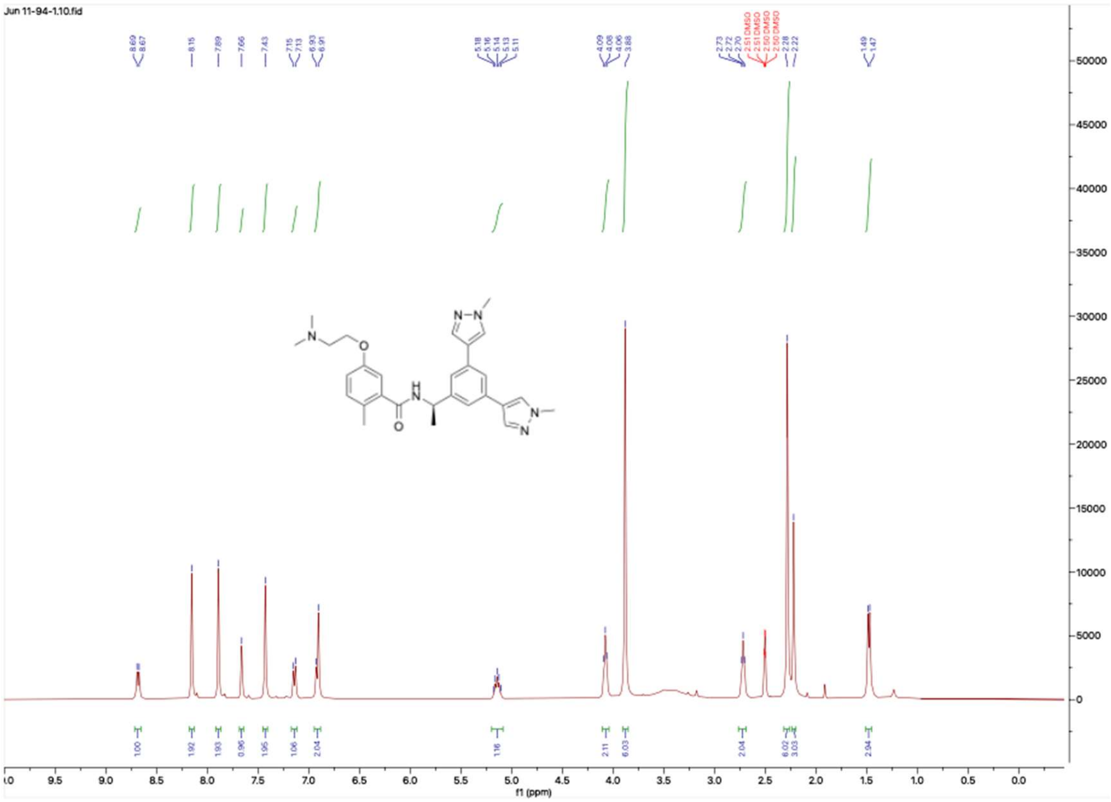


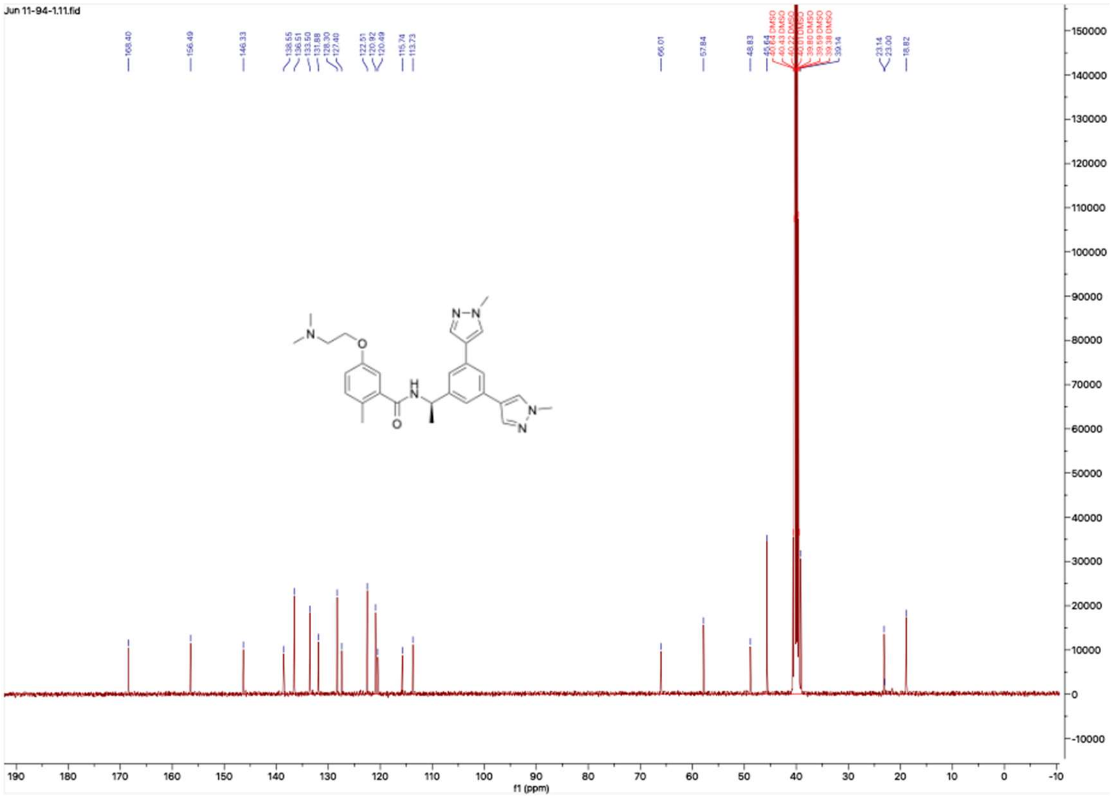

**HNMR and CNMR spectra of Jun11942**

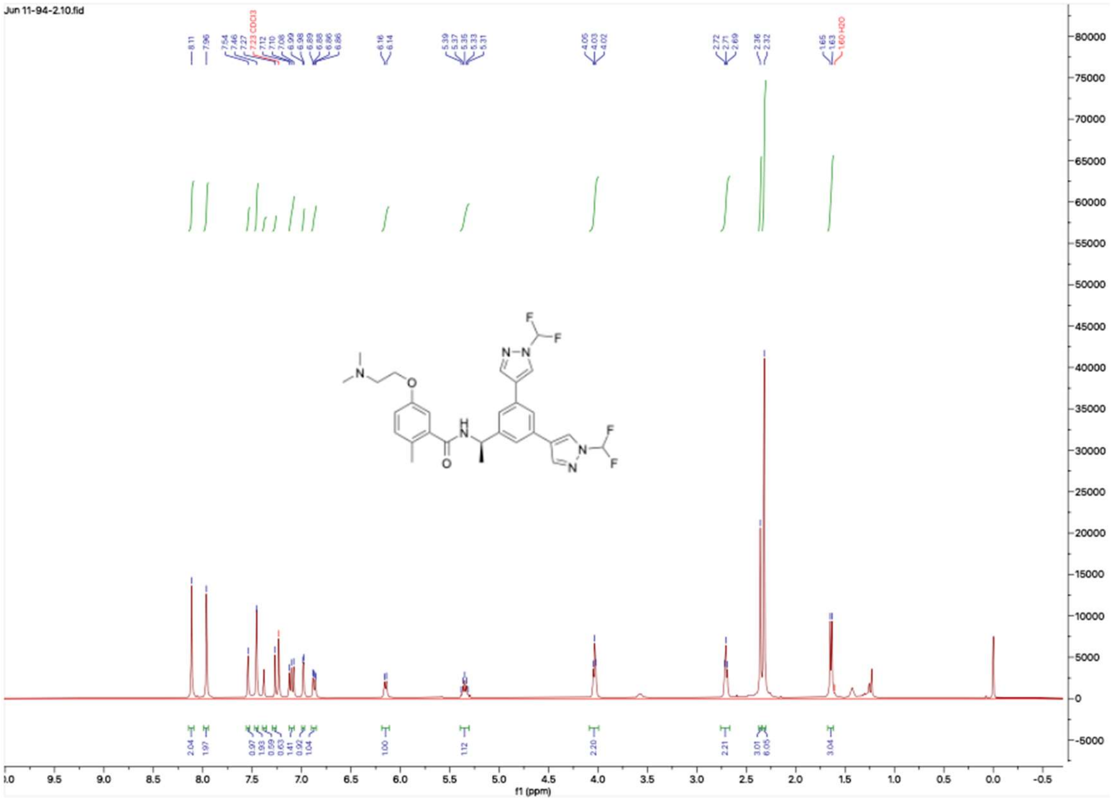


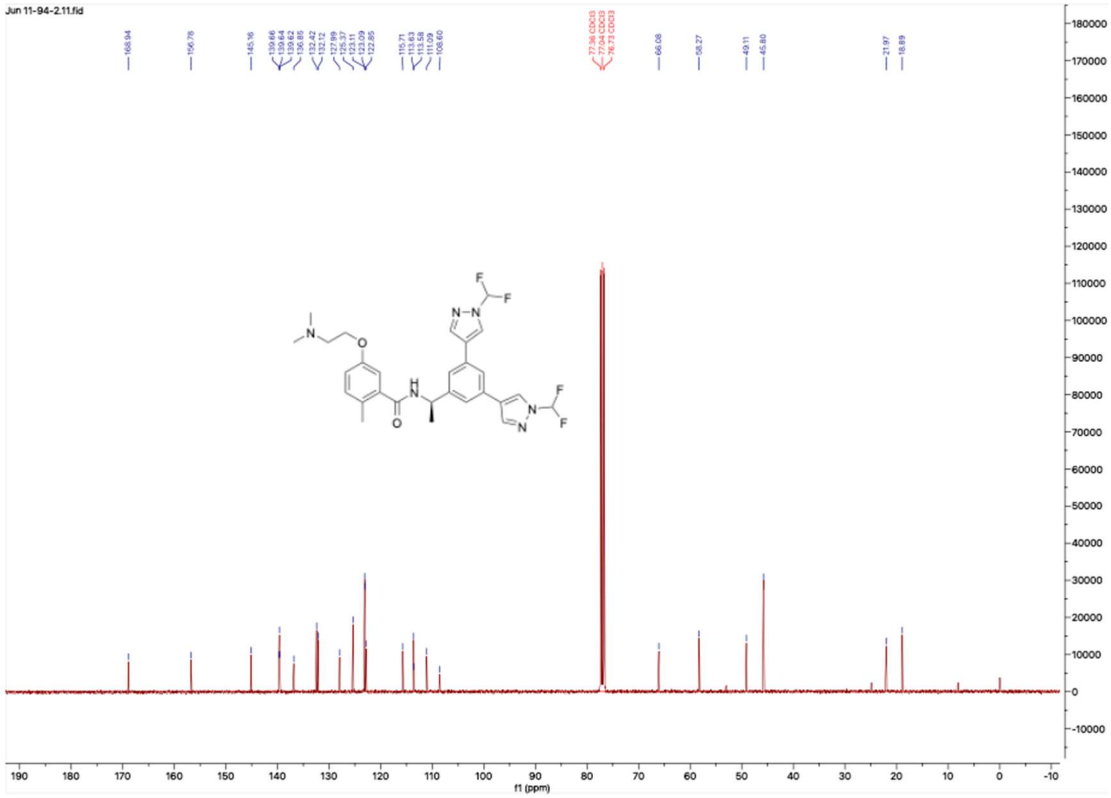

**HNMR and CNMR spectra of Jun1212**

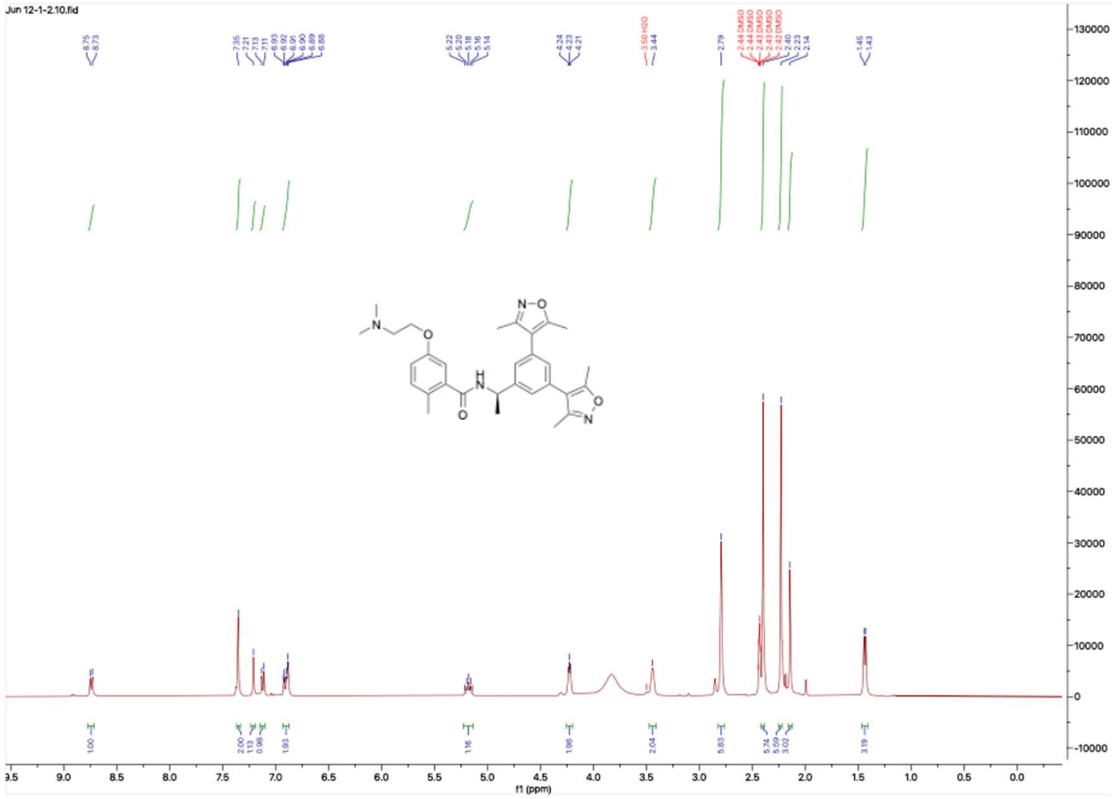

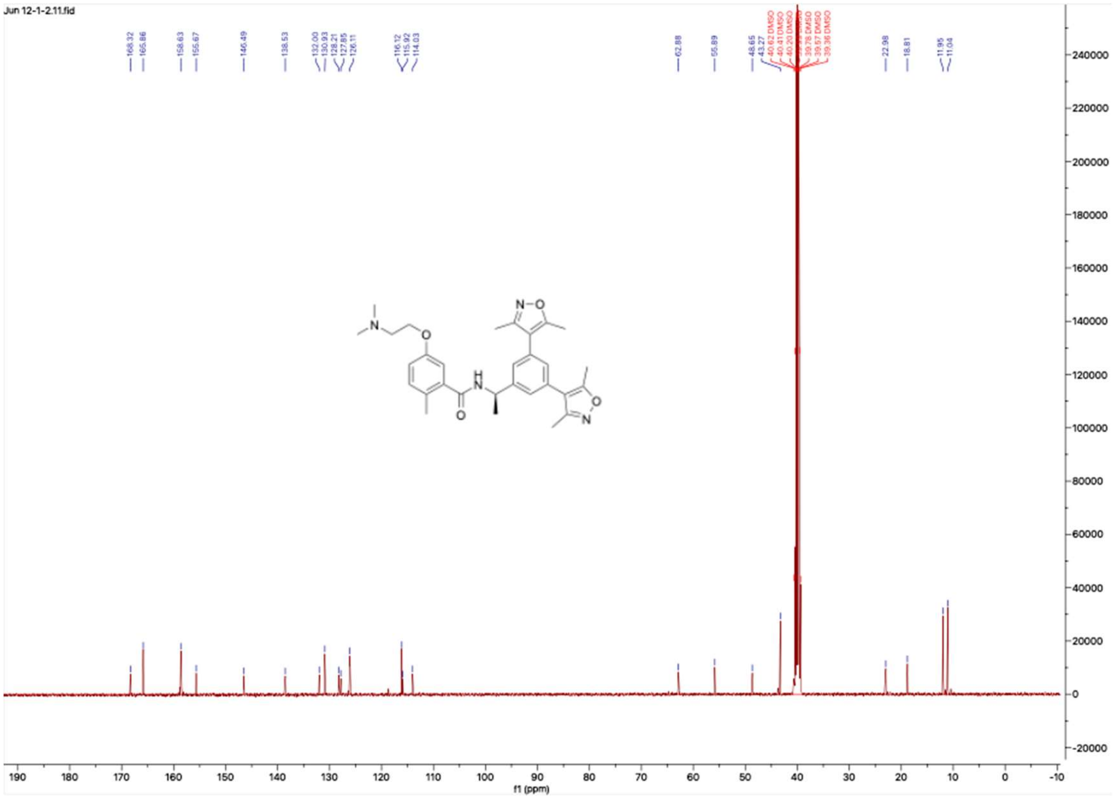

**HNMR and CNMR spectra of Jun1215**

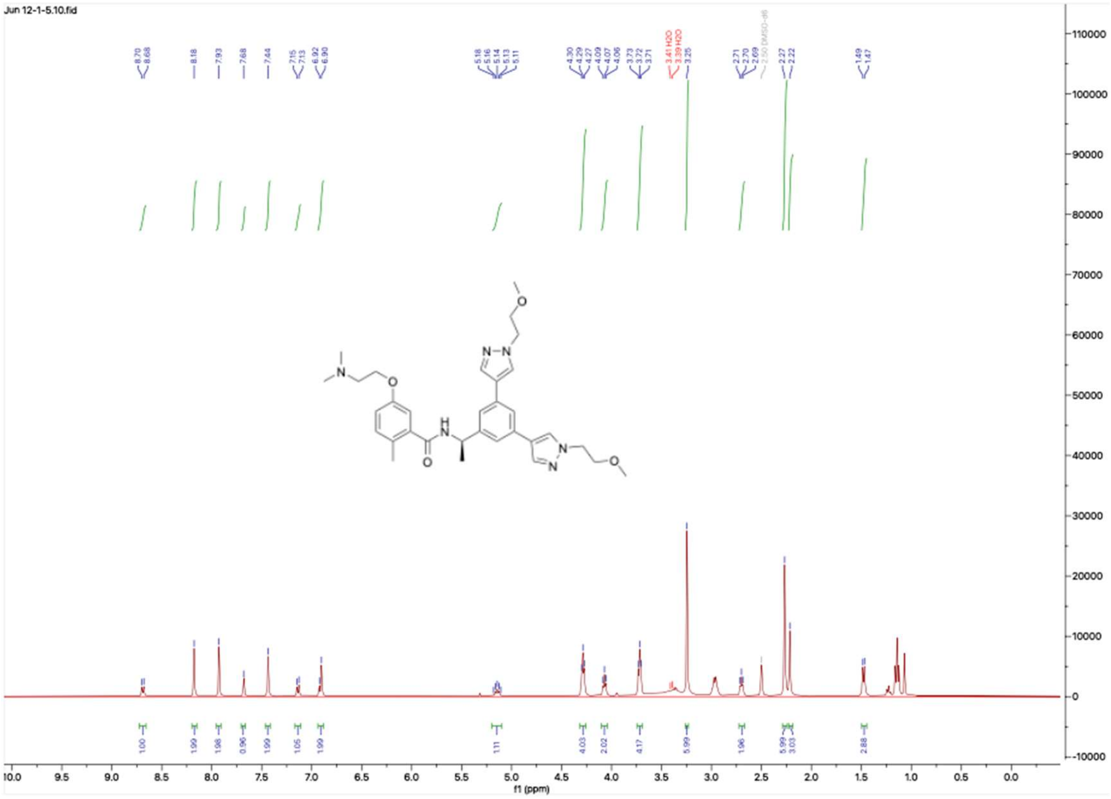

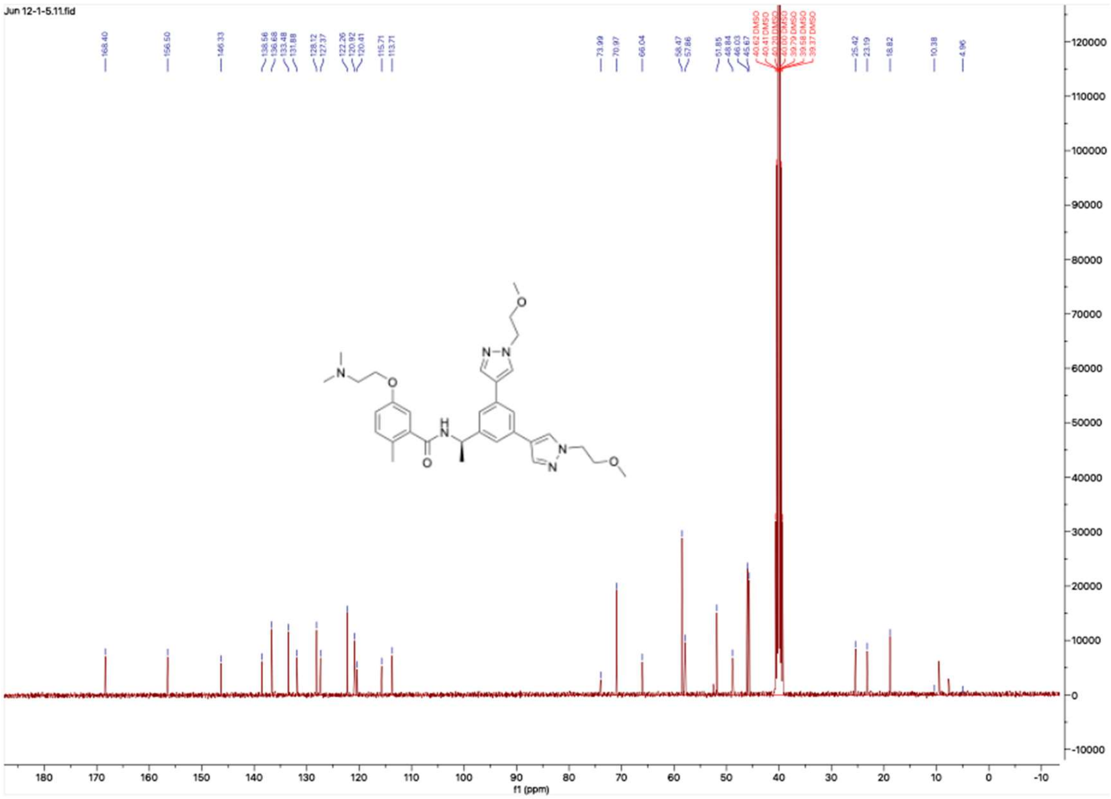

### HNMR and CNMR spectra of Jun1247

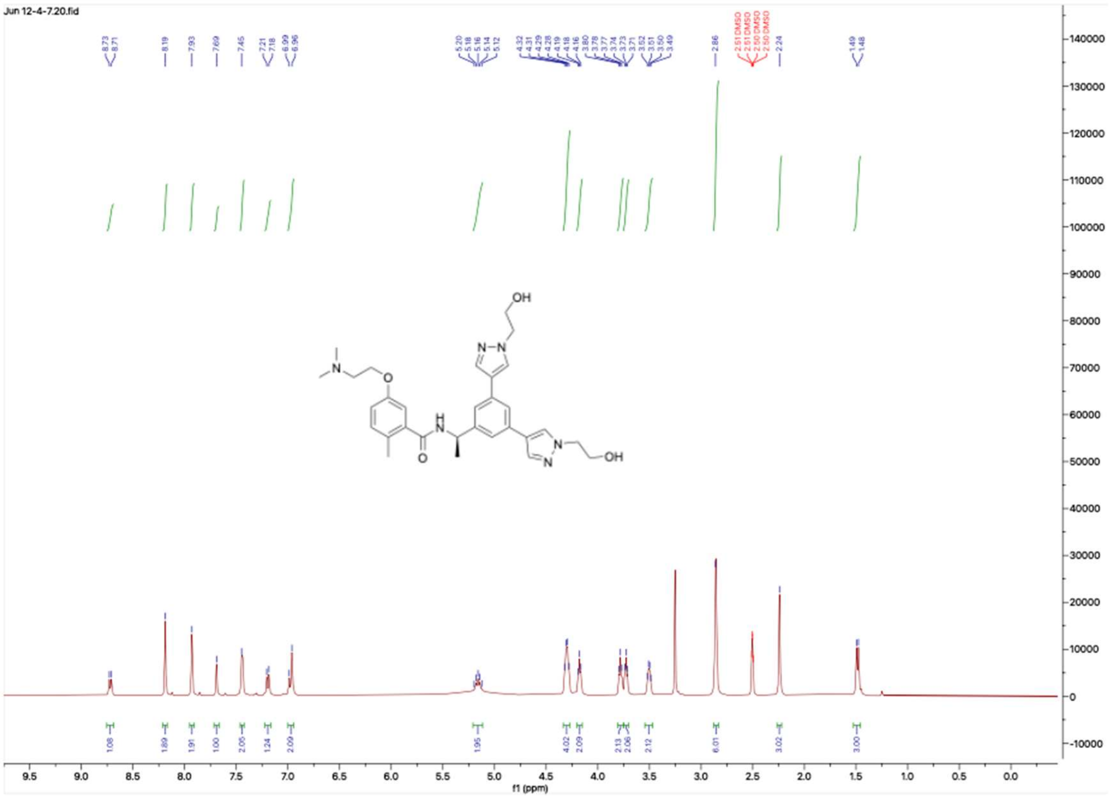

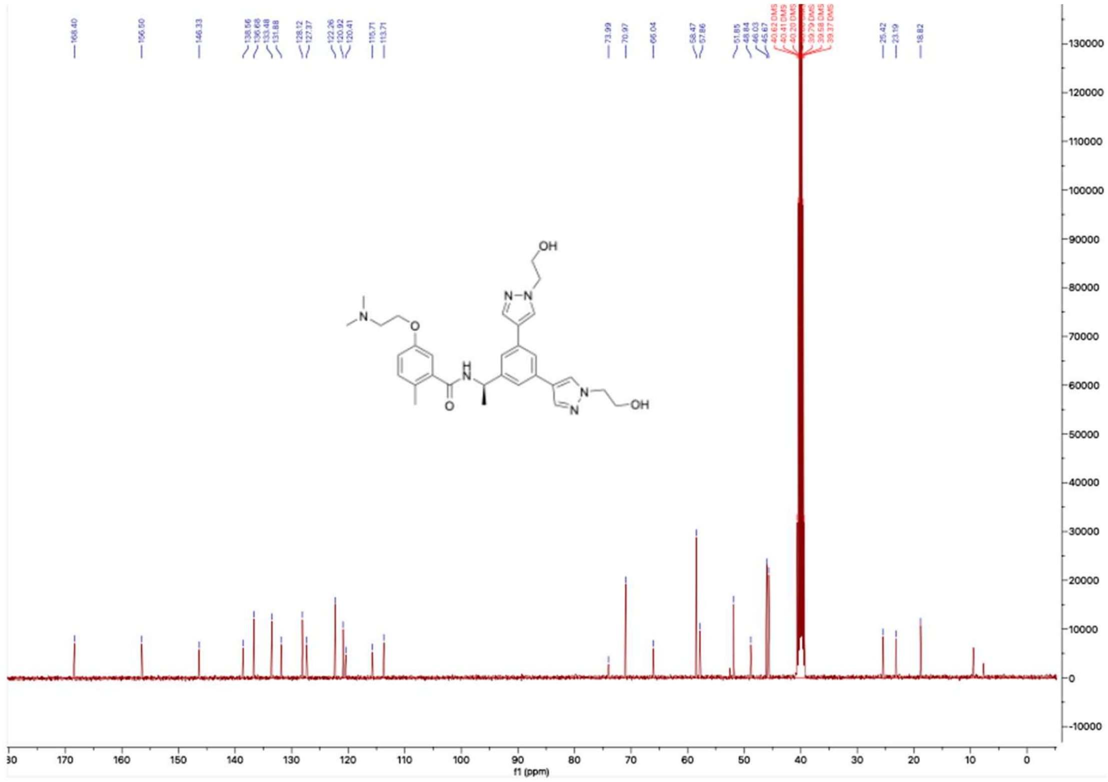

**HNMR and CNMR spectra of Jun12382**

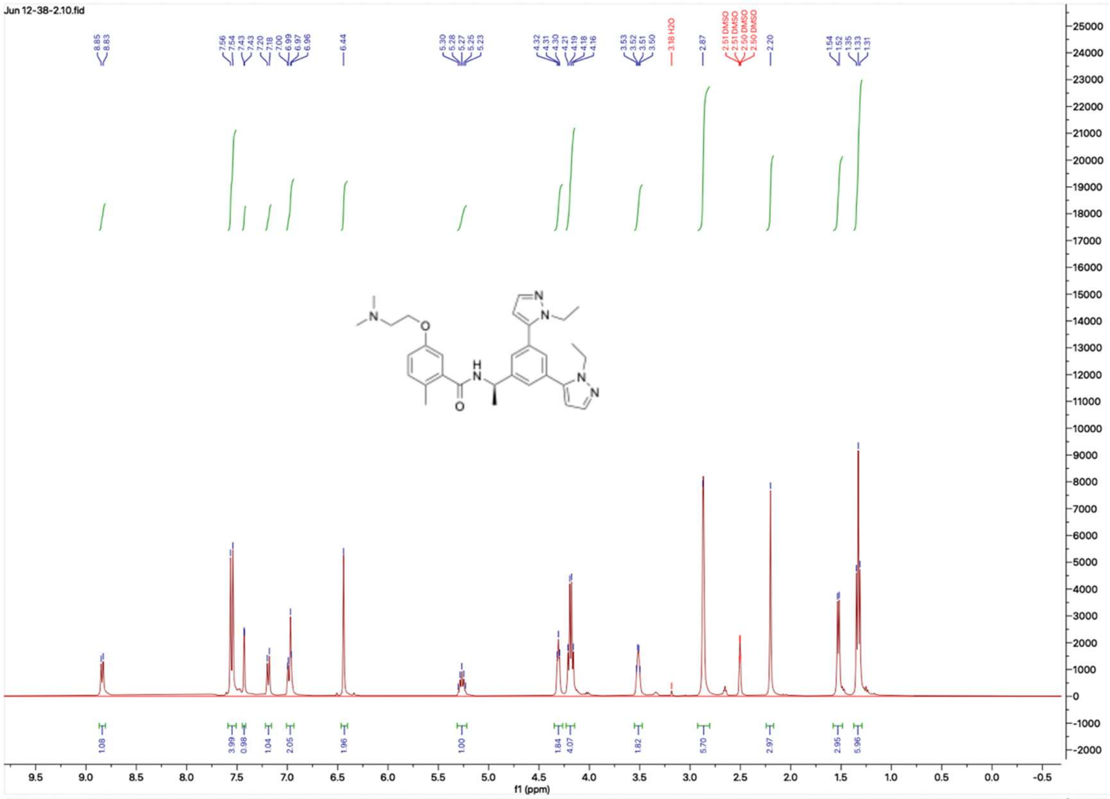

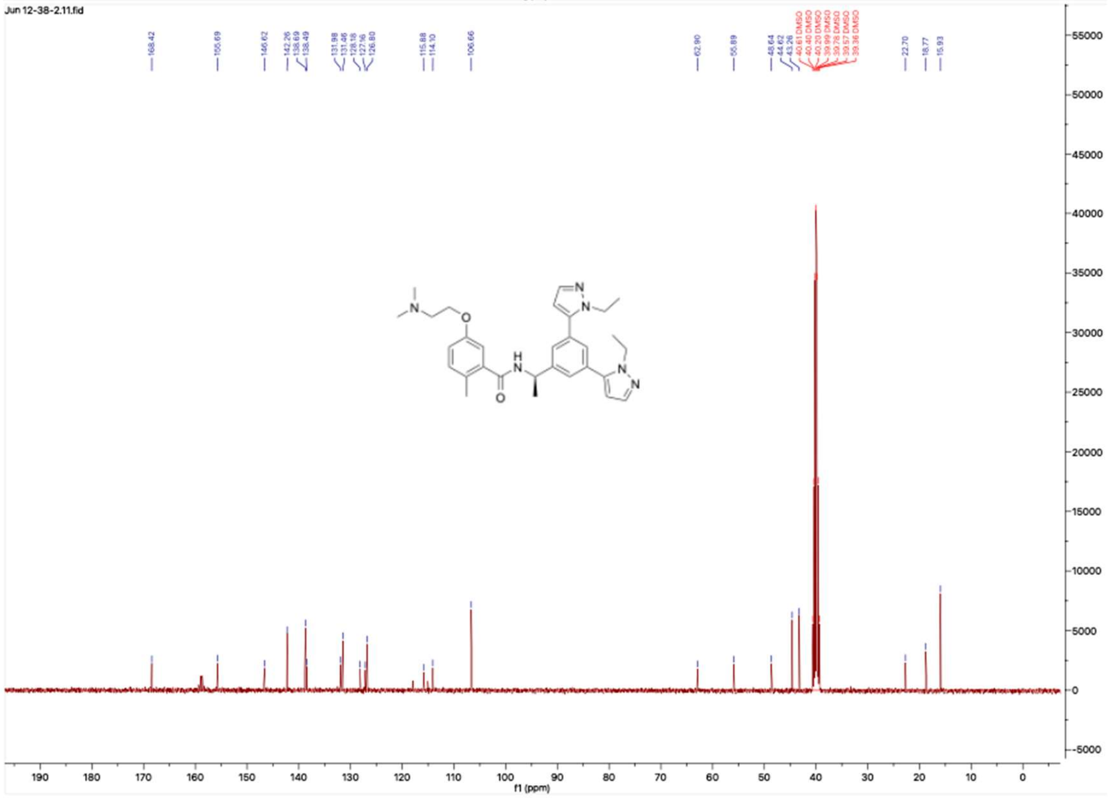

#### HNMR and CNMR spectra of Jun1246

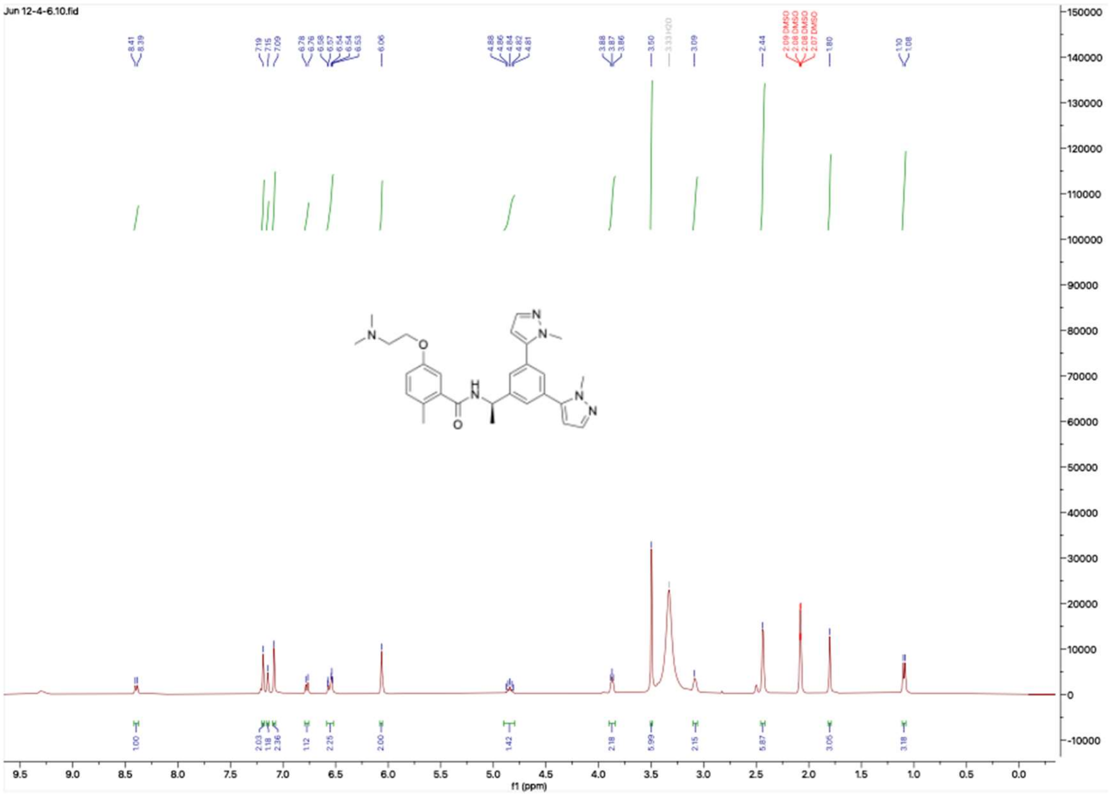

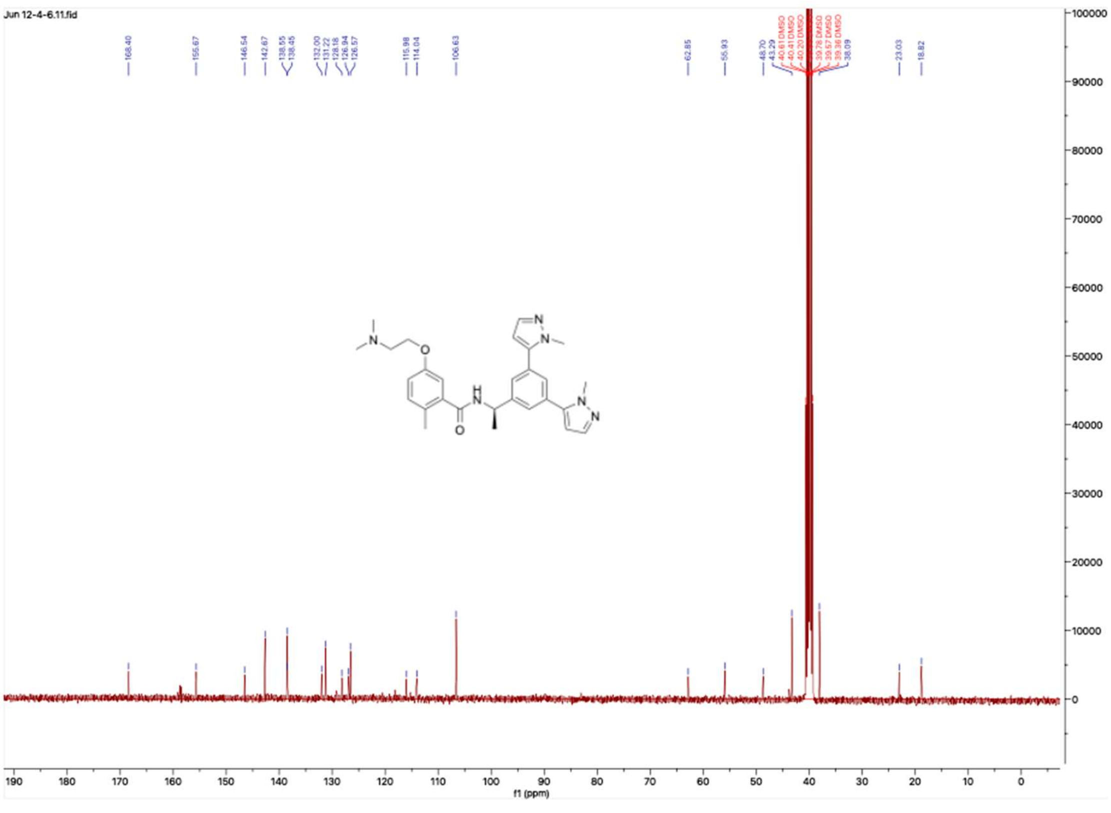

**HNMR and CNMR spectra of Jun1211**

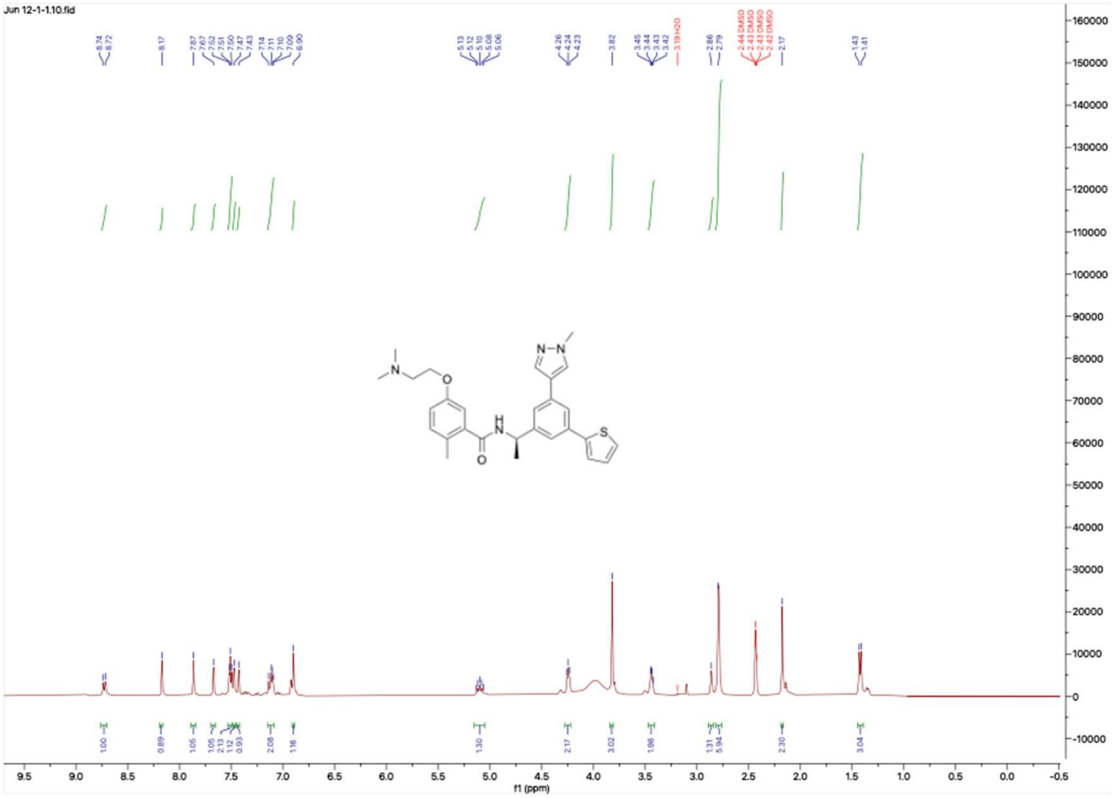

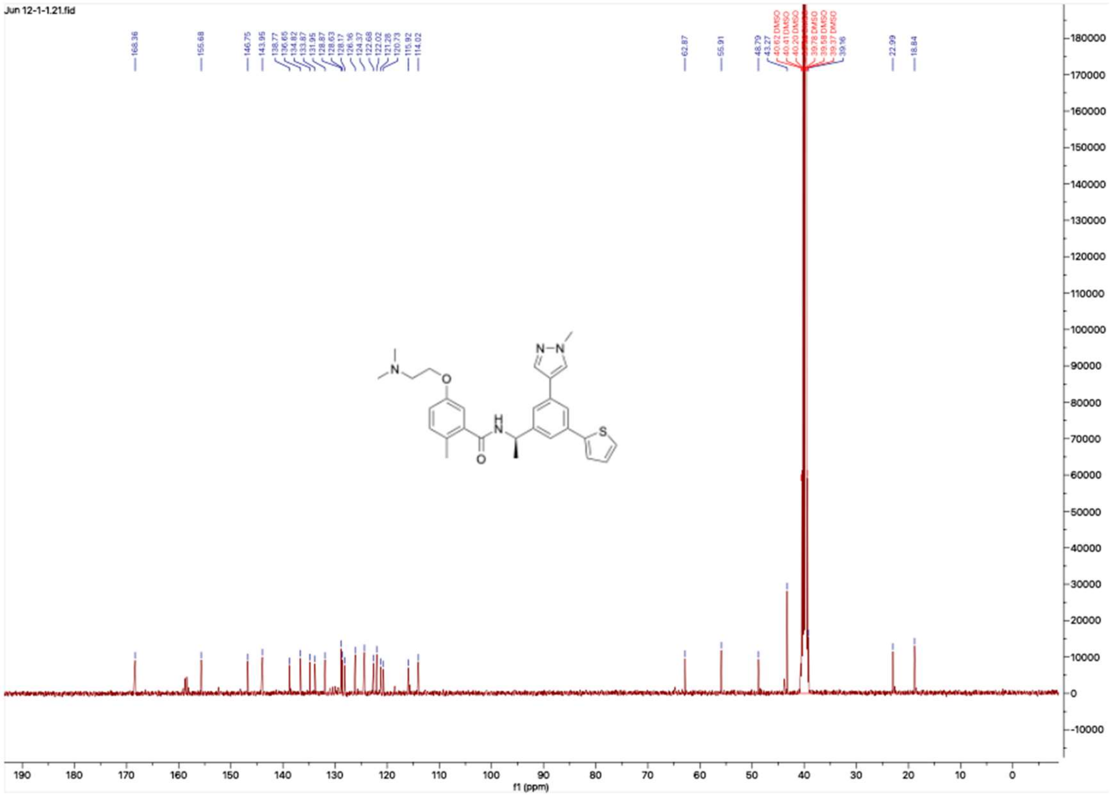

**HNMR and CNMR spectra of Jun1244**

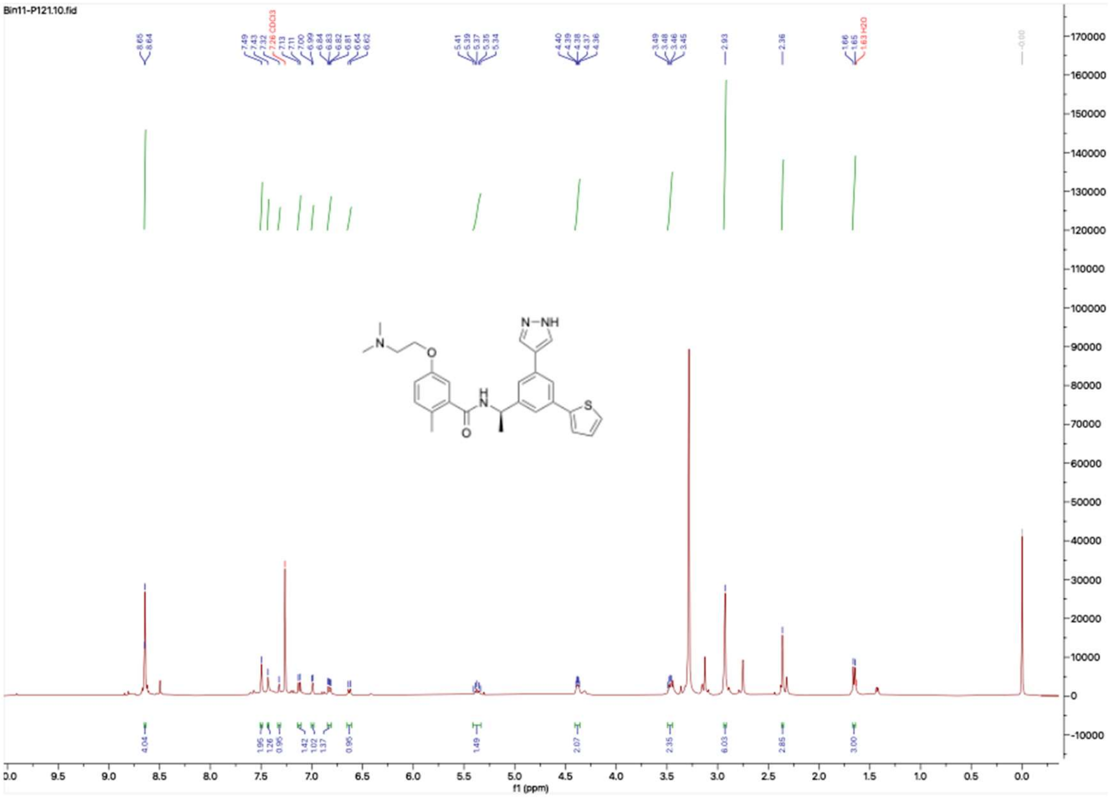


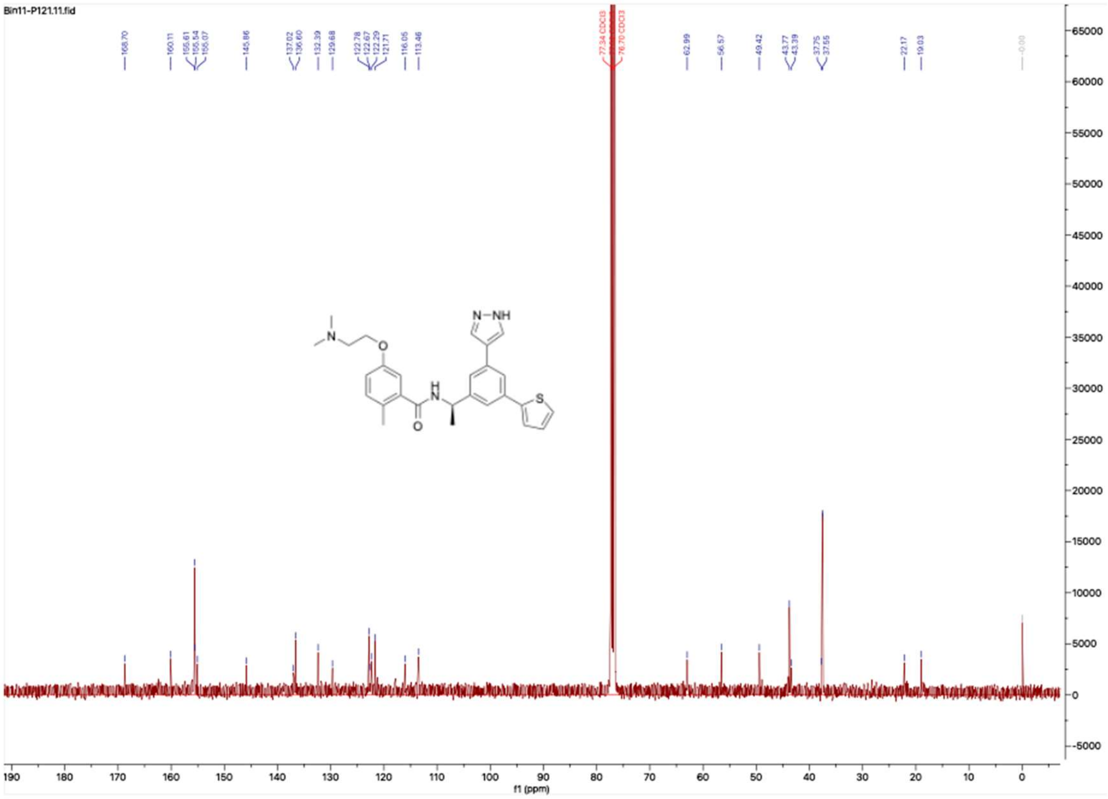

**HNMR and CNMR spectra of Jun1251**

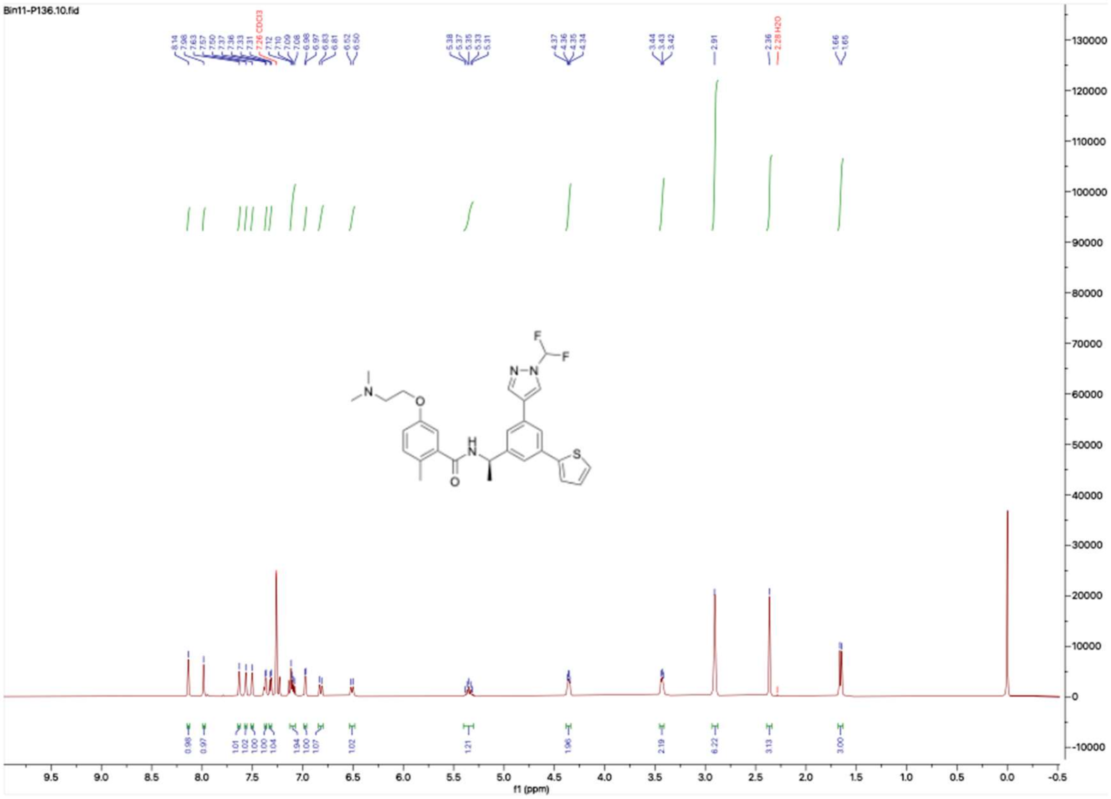

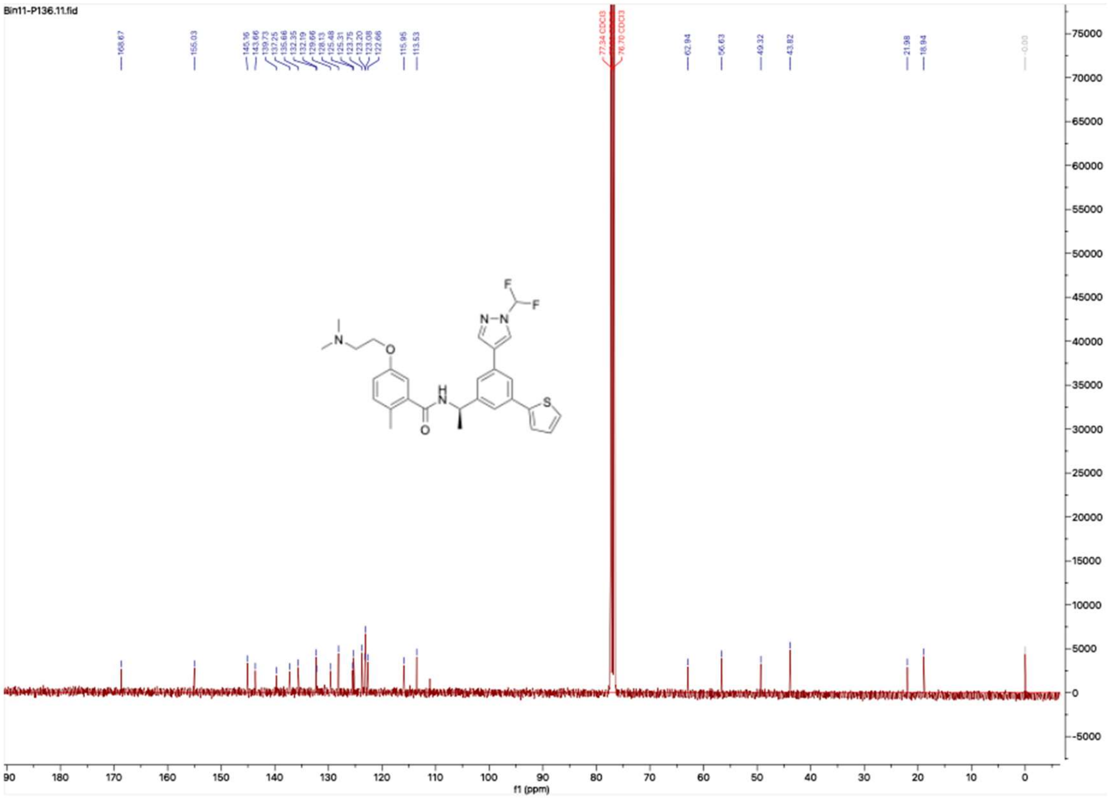

**HNMR and CNMR spectra of Jun1252**

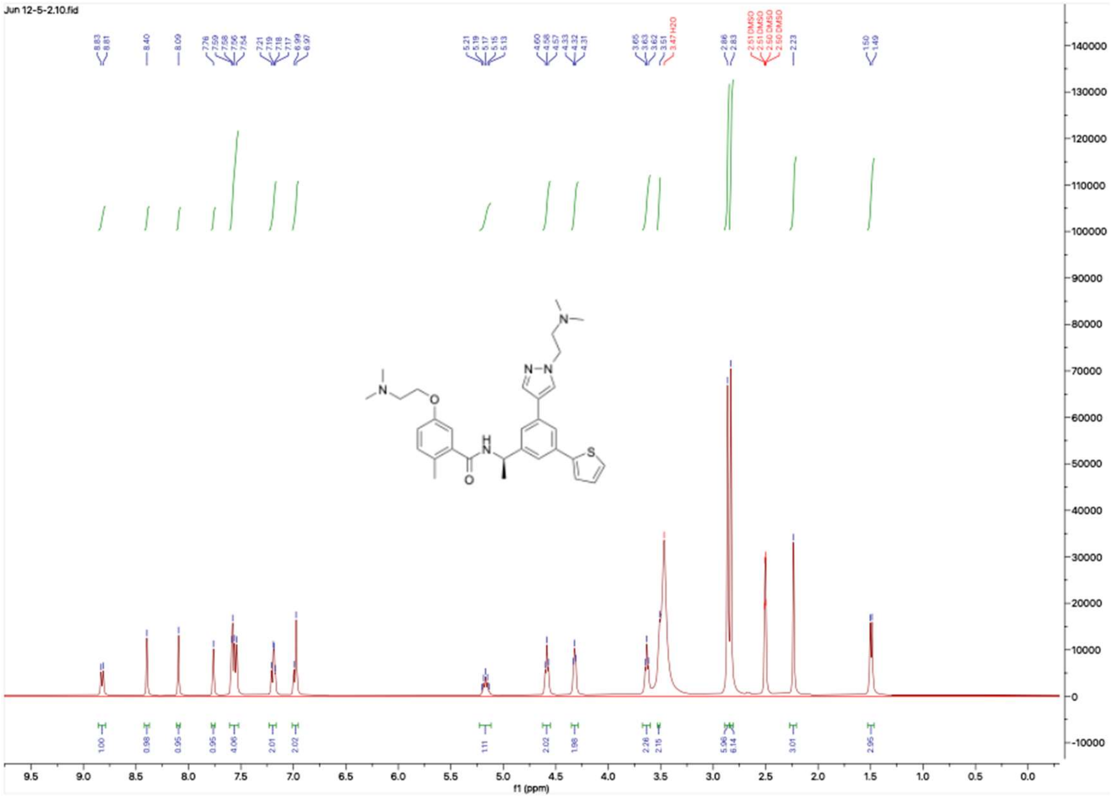


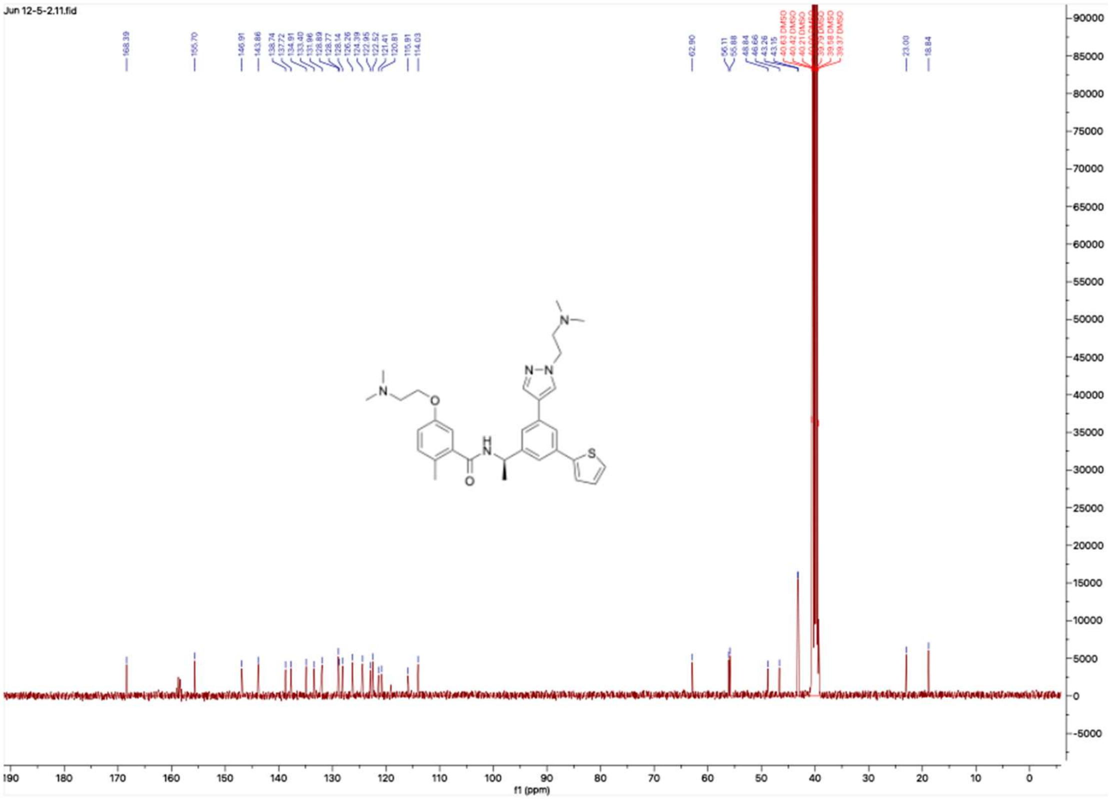

**HNMR and CNMR spectra of Jun1254**

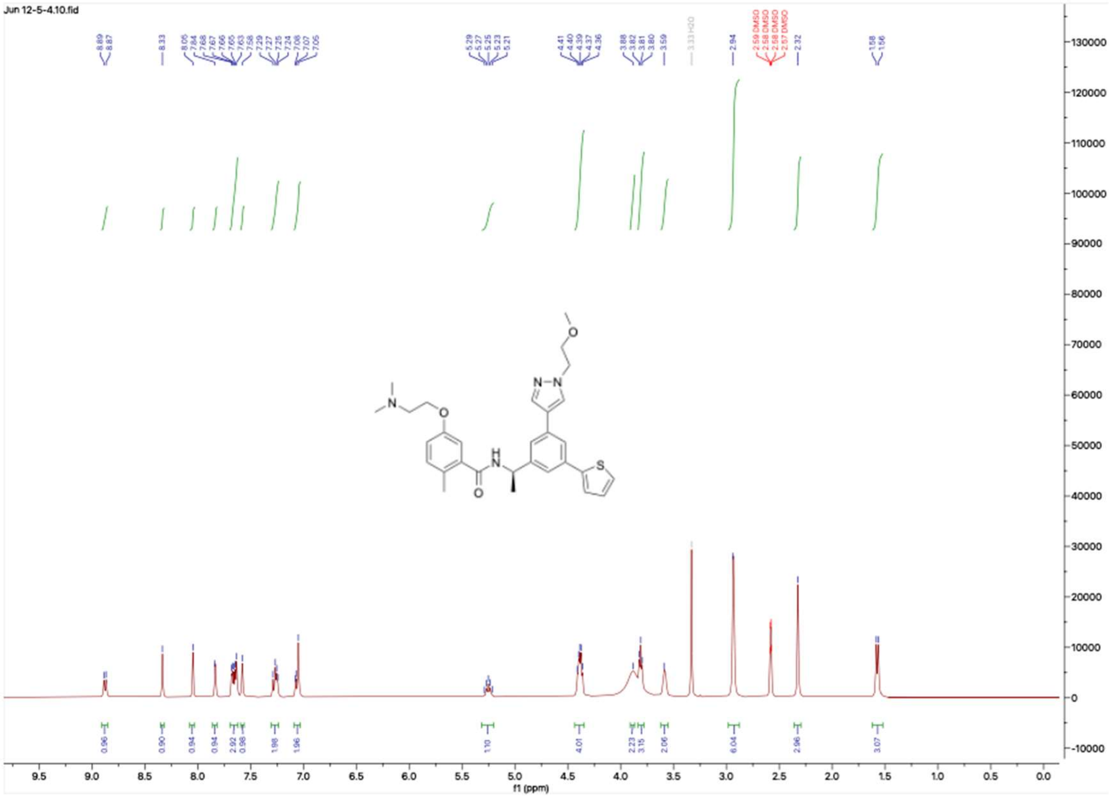


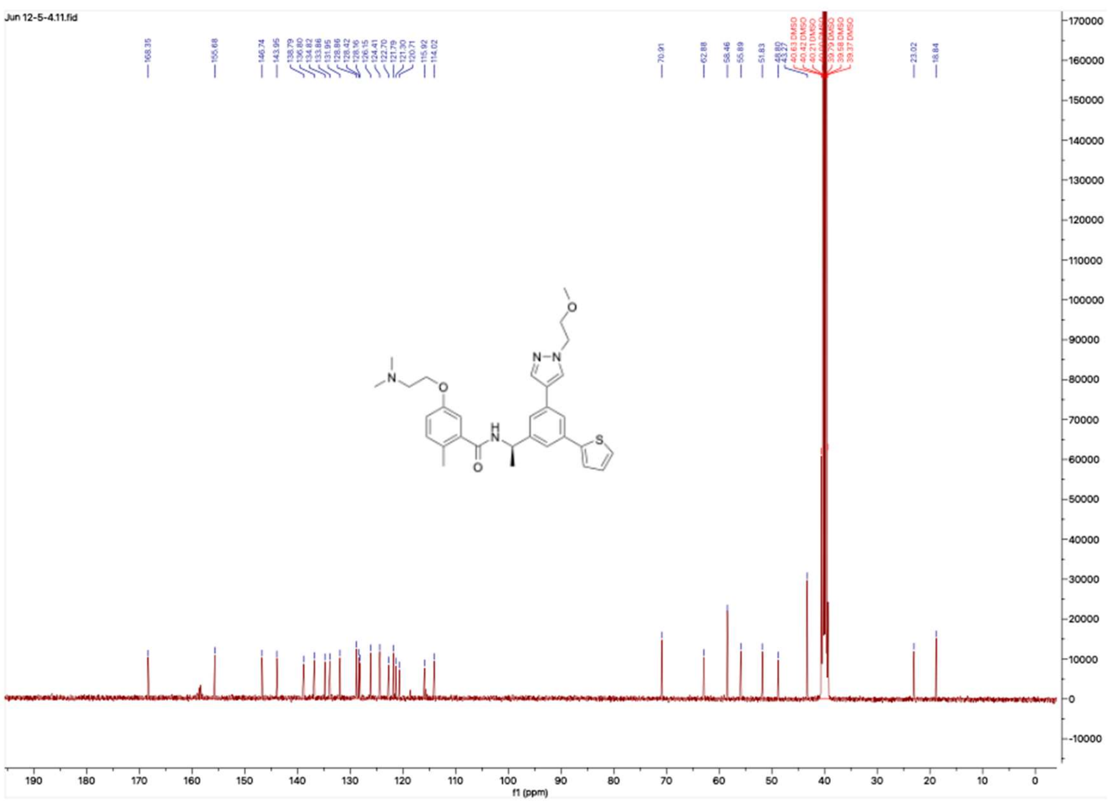

**HNMR and CNMR spectra of Jun1257**

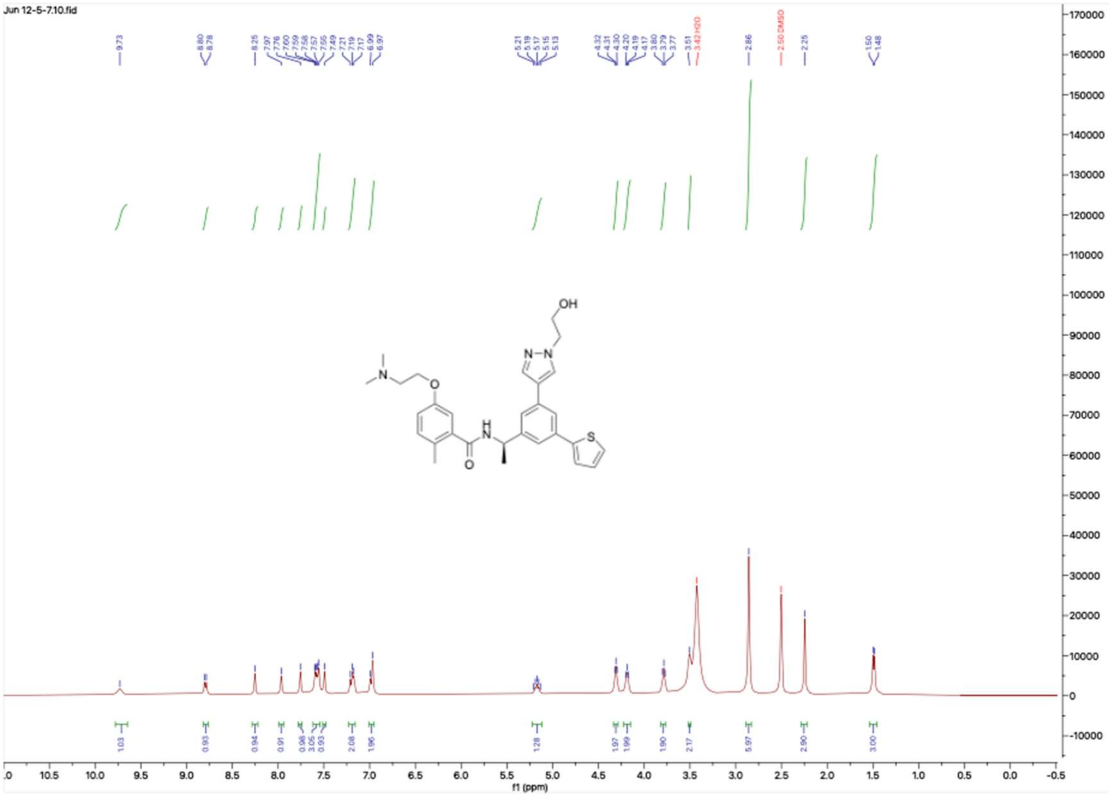


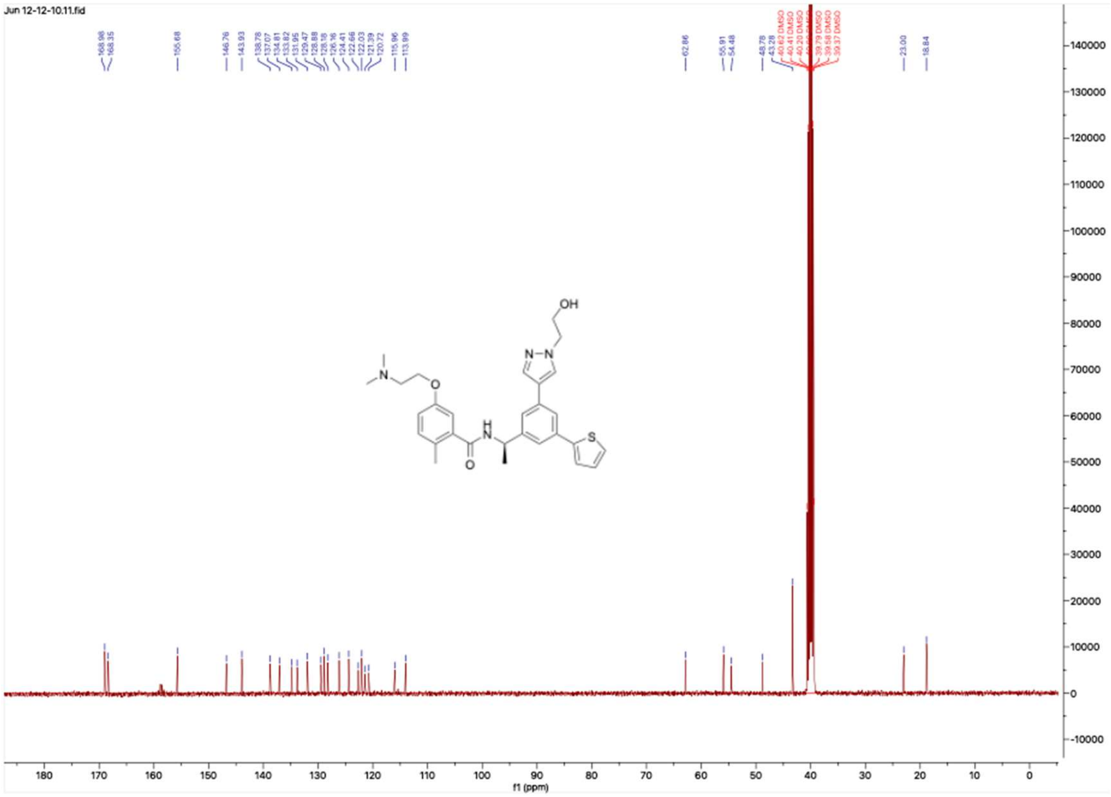

**HNMR and CNMR spectra of Jun121210**

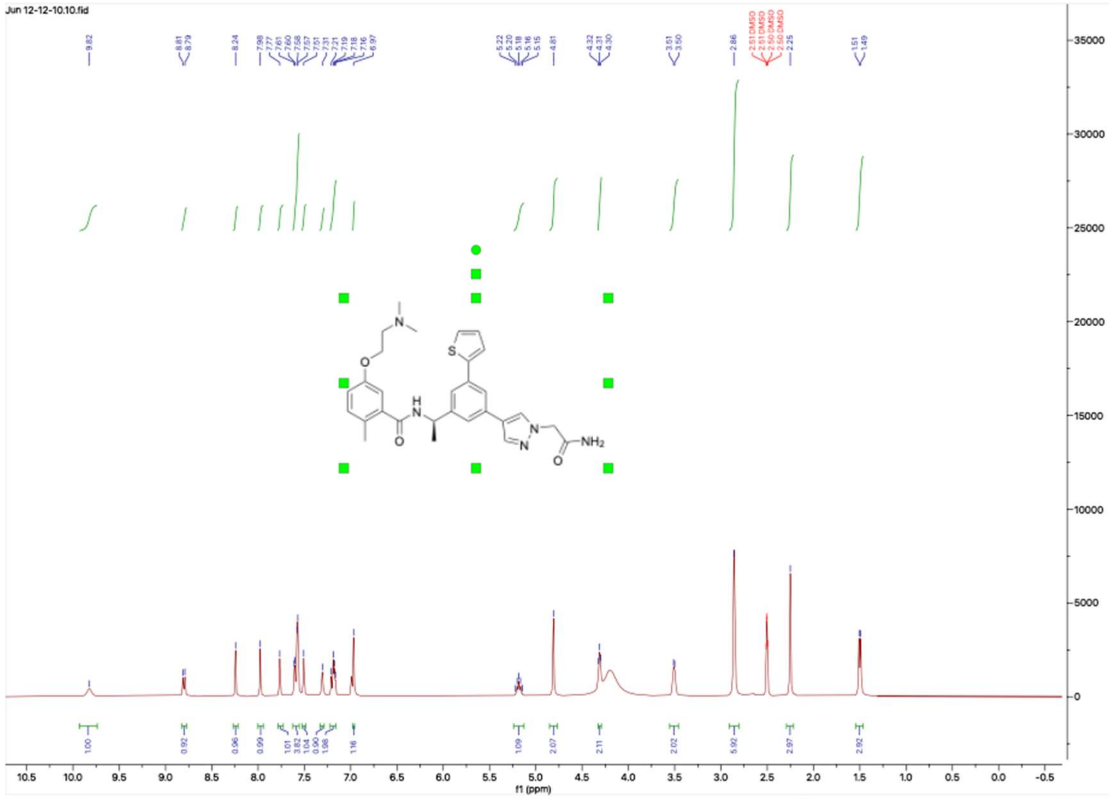

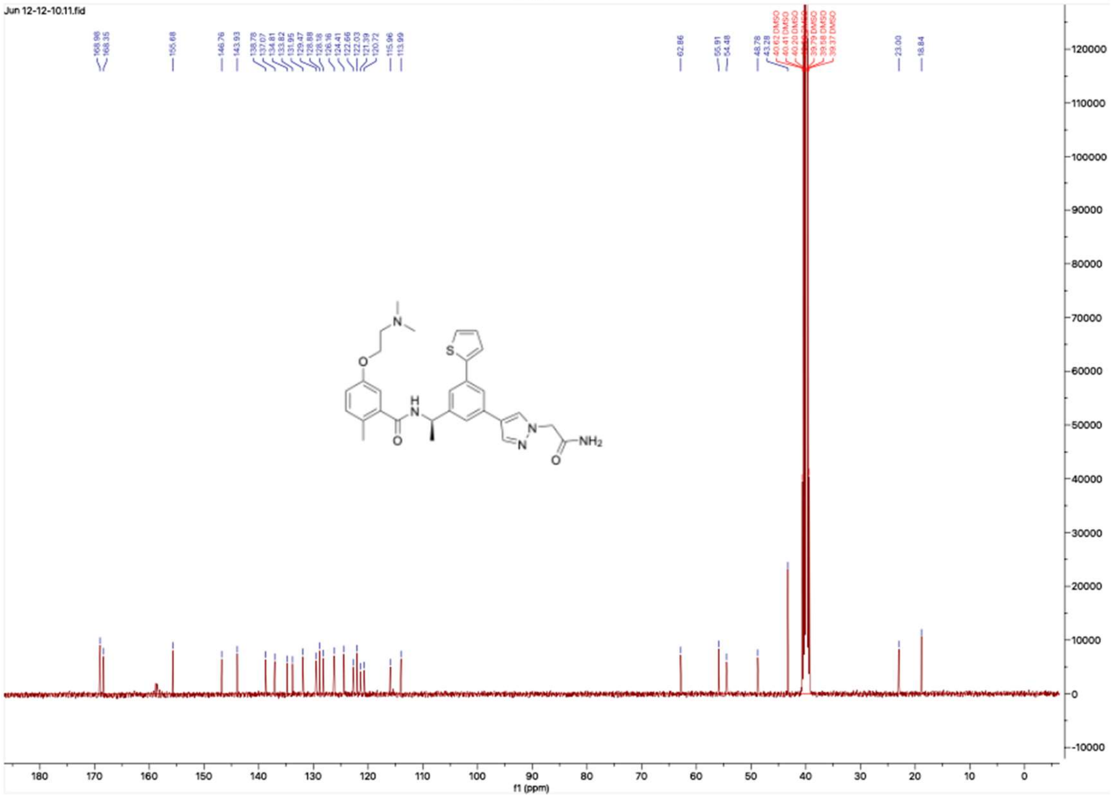

**HNMR and CNMR spectra of Jun12149**

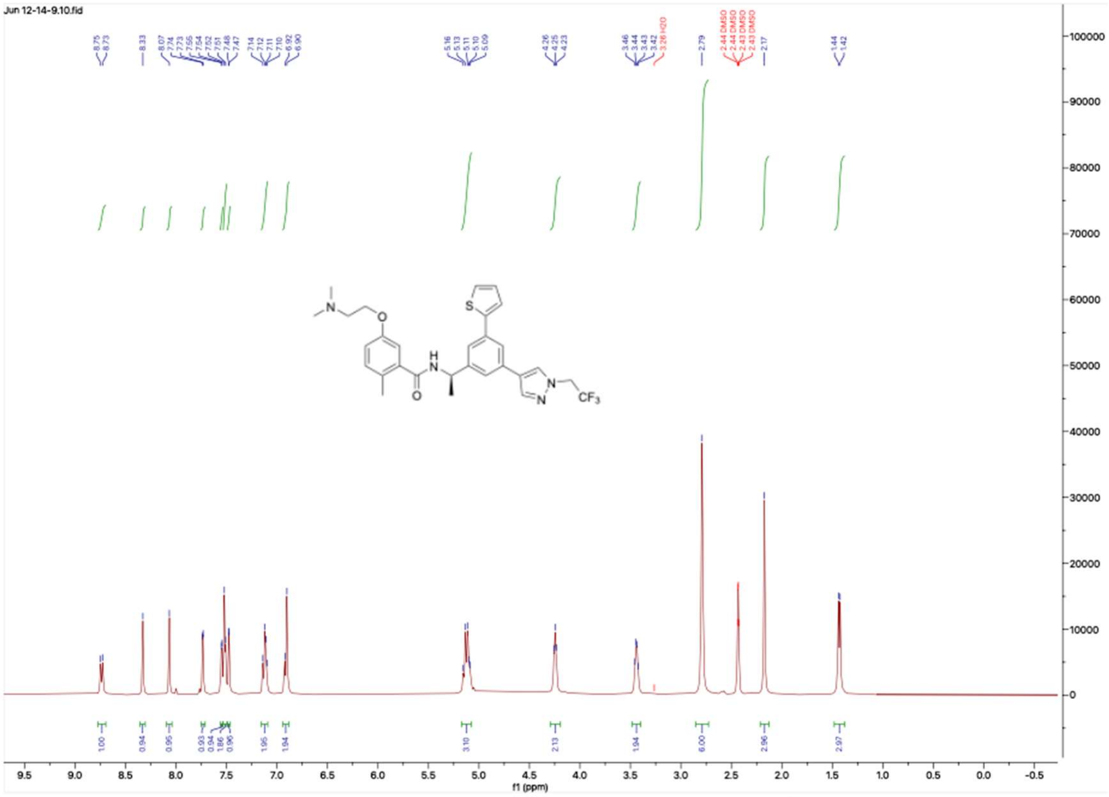

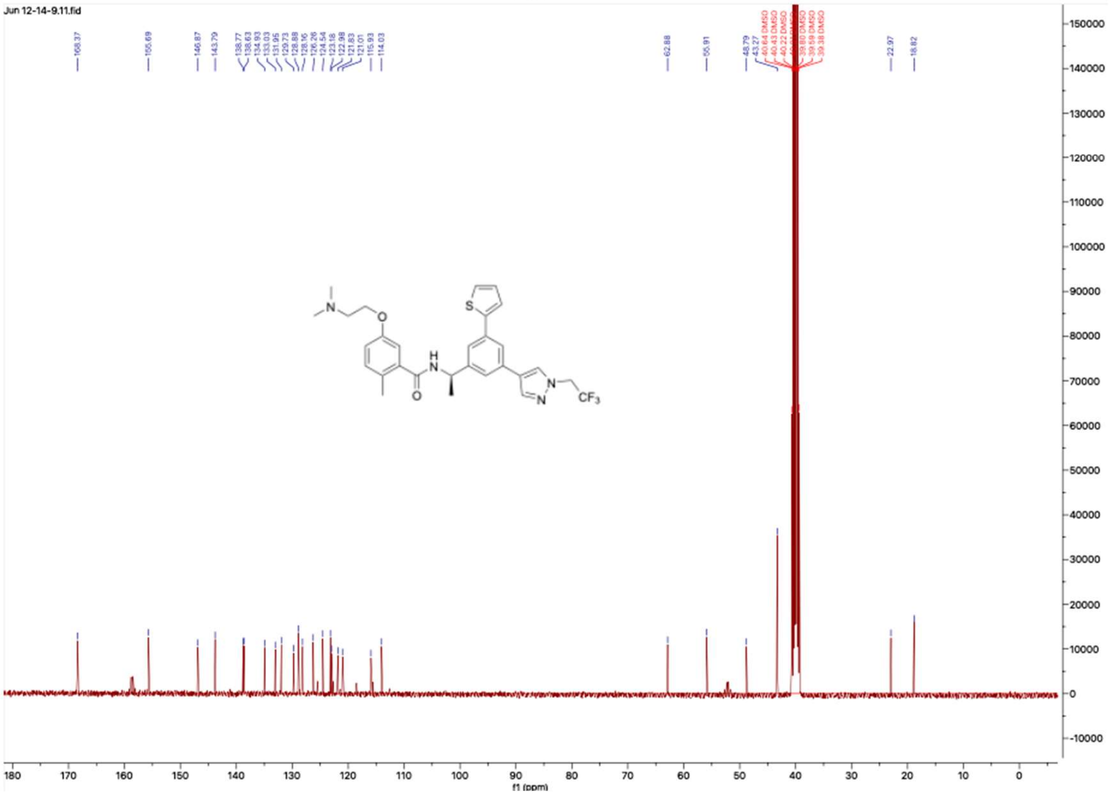

**HNMR and CNMR spectra of Jun12129**

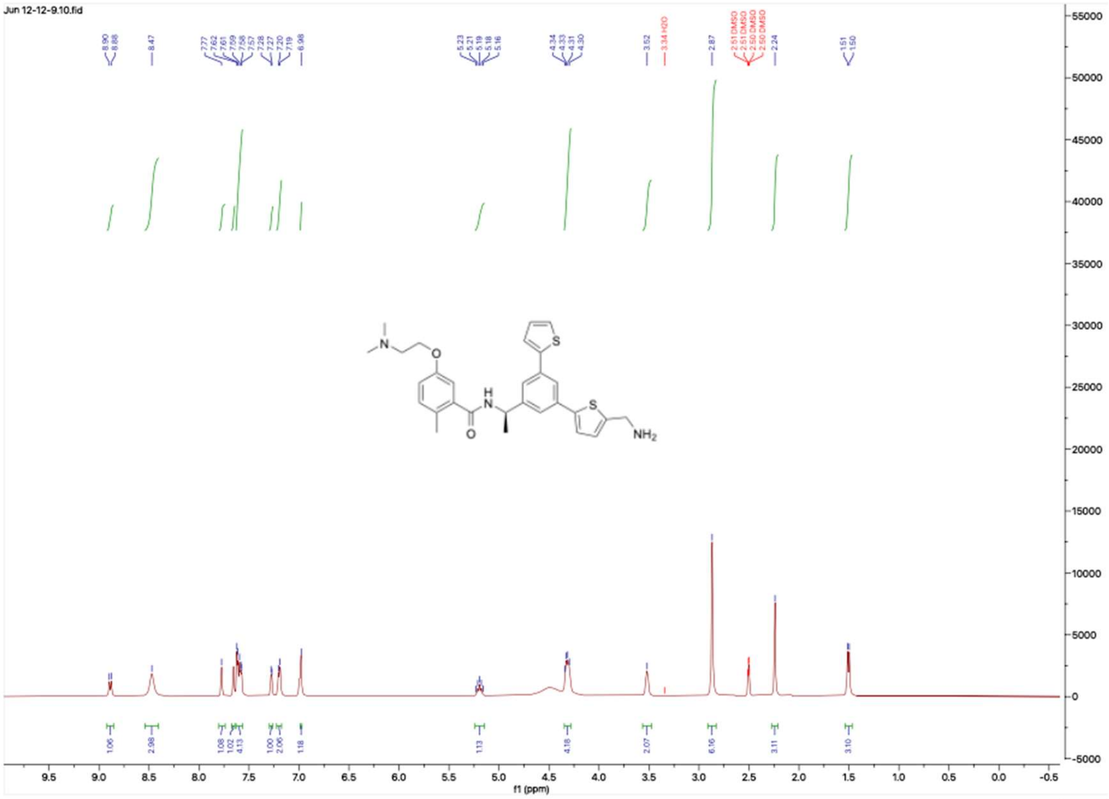


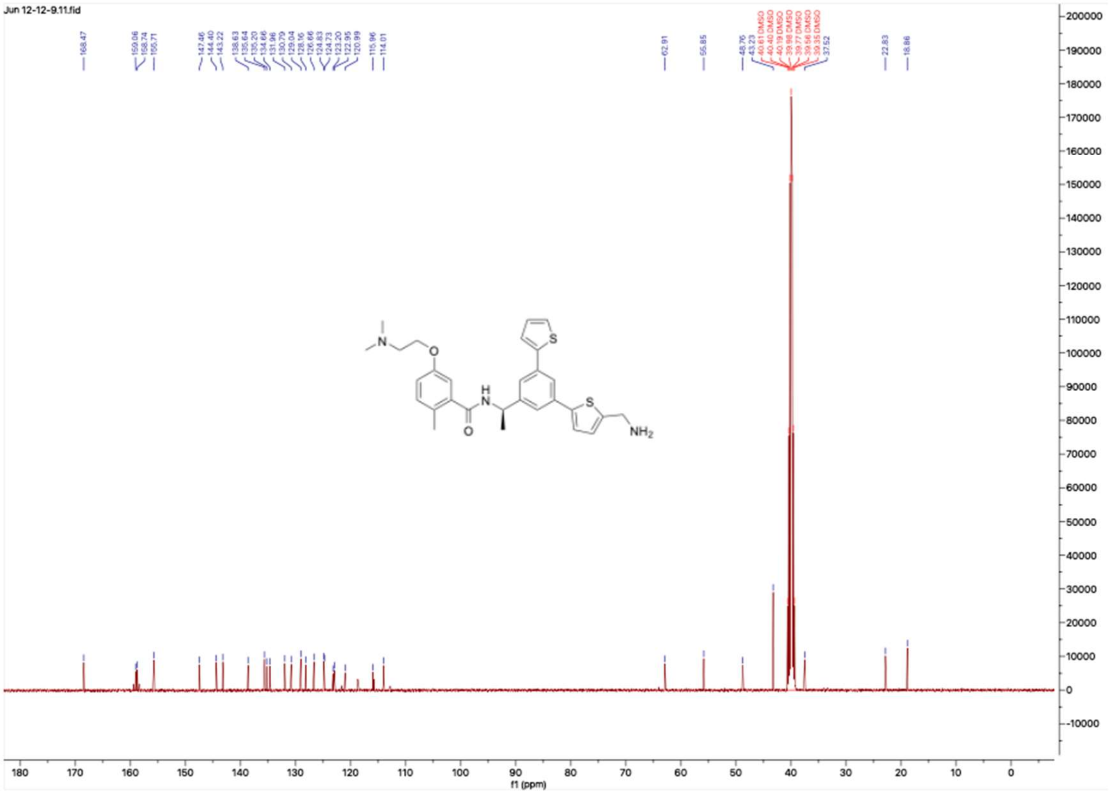

### HNMR and CNMR spectra of Jun121910

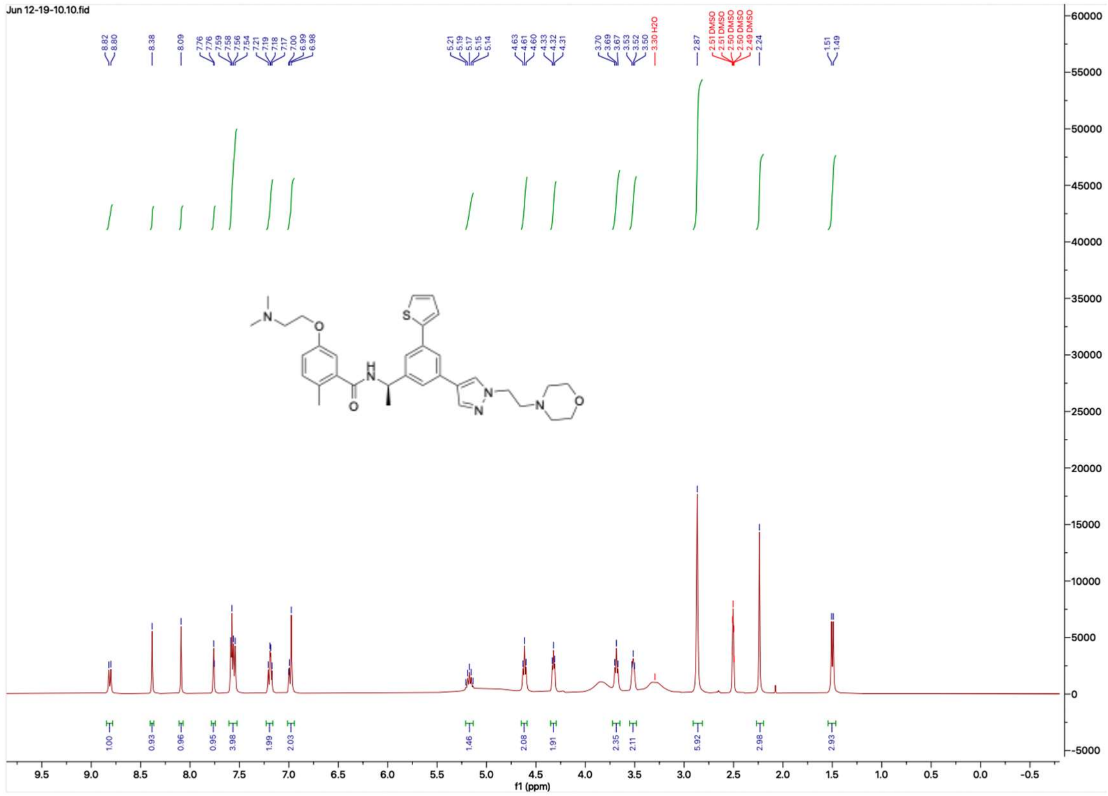


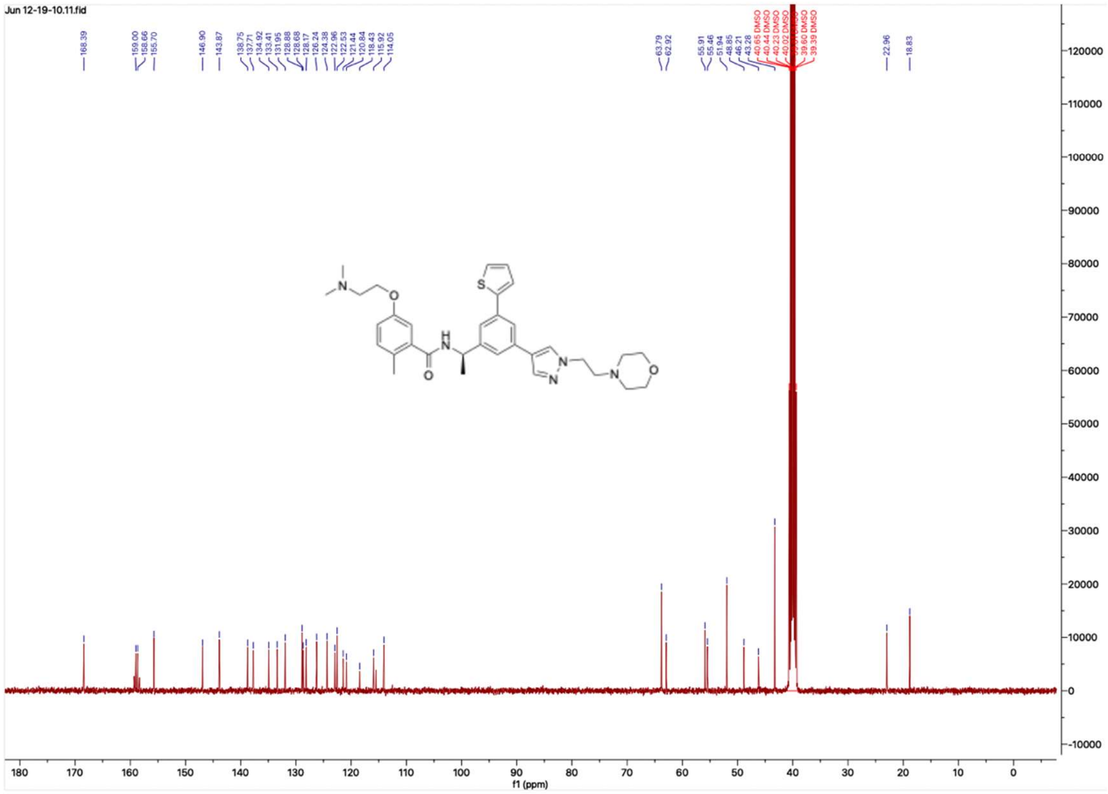

**HNMR and CNMR spectra of Jun12511**

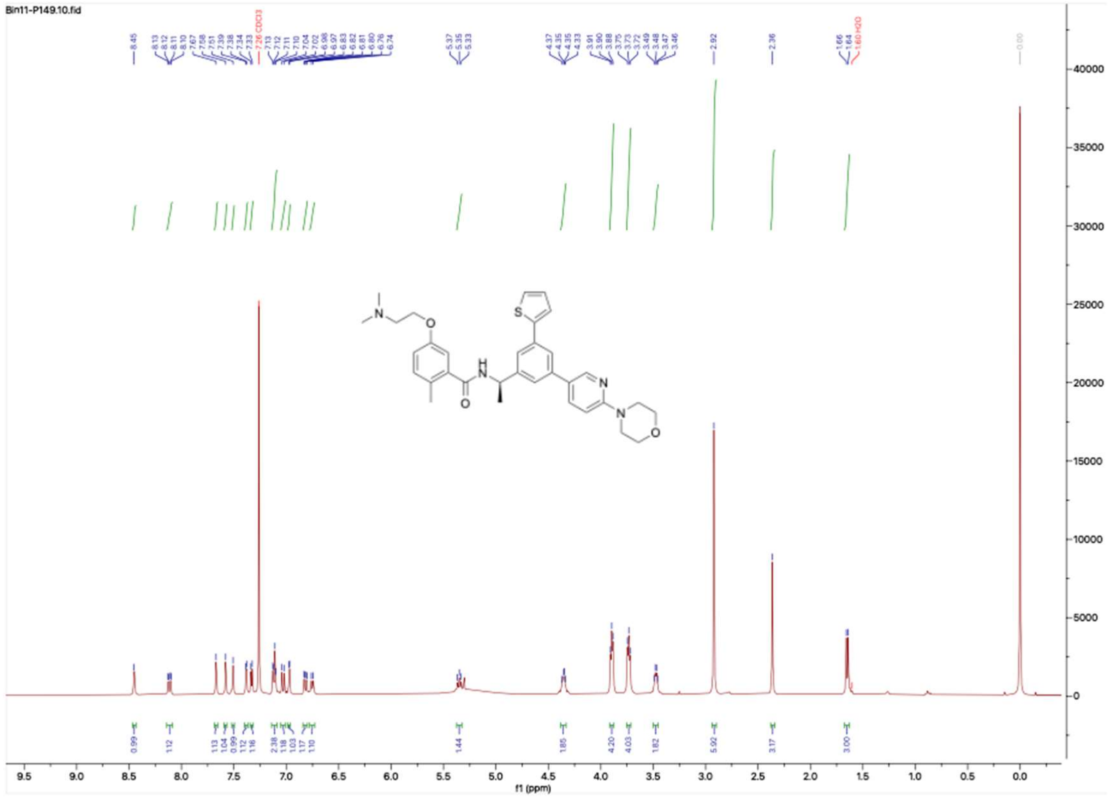

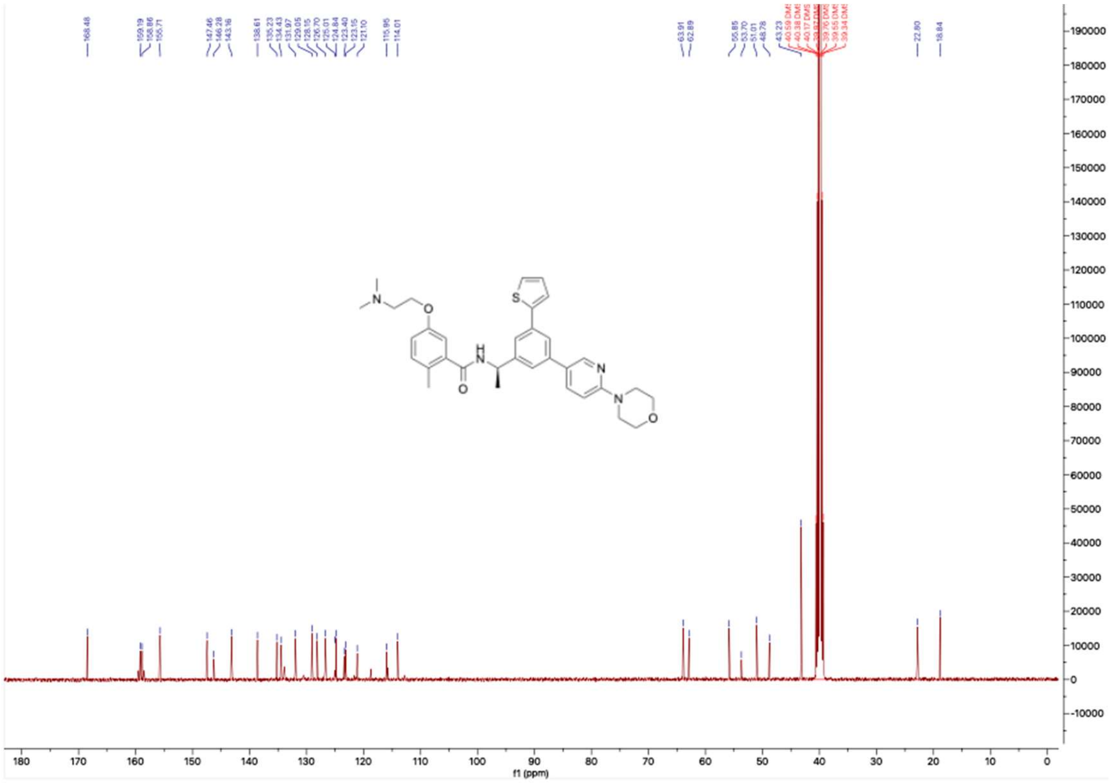

#### HNMR and CNMR spectra of Jun12338

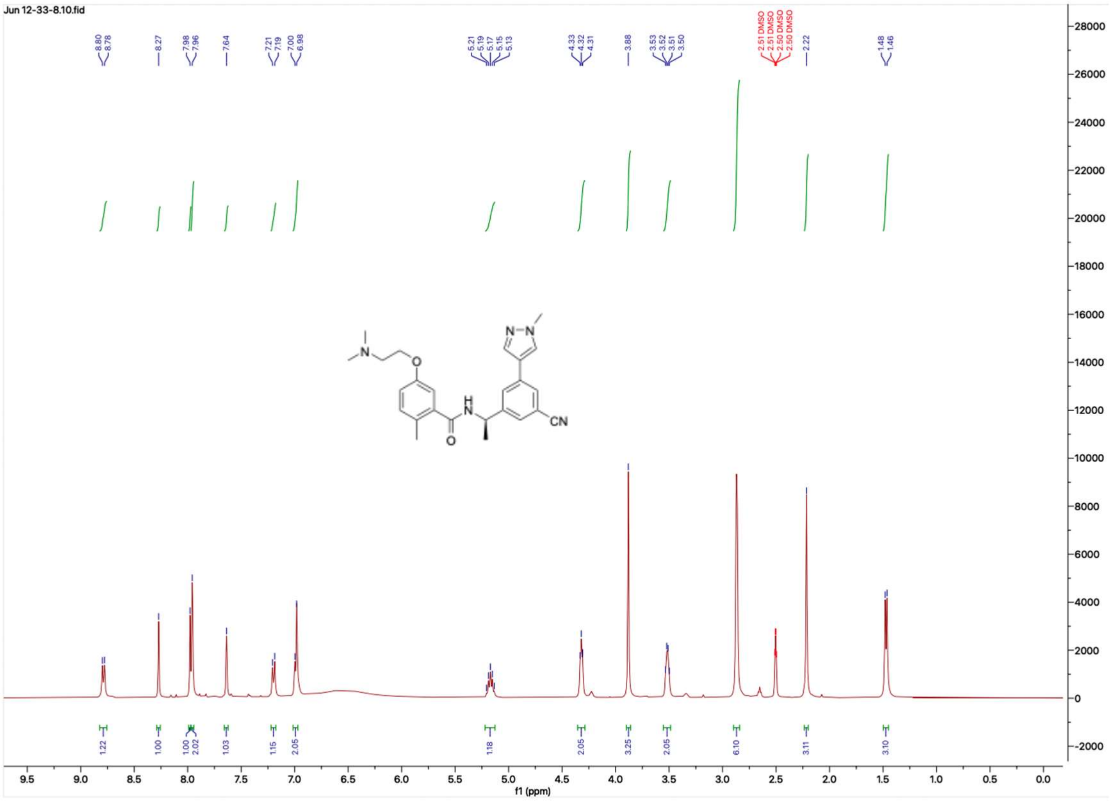

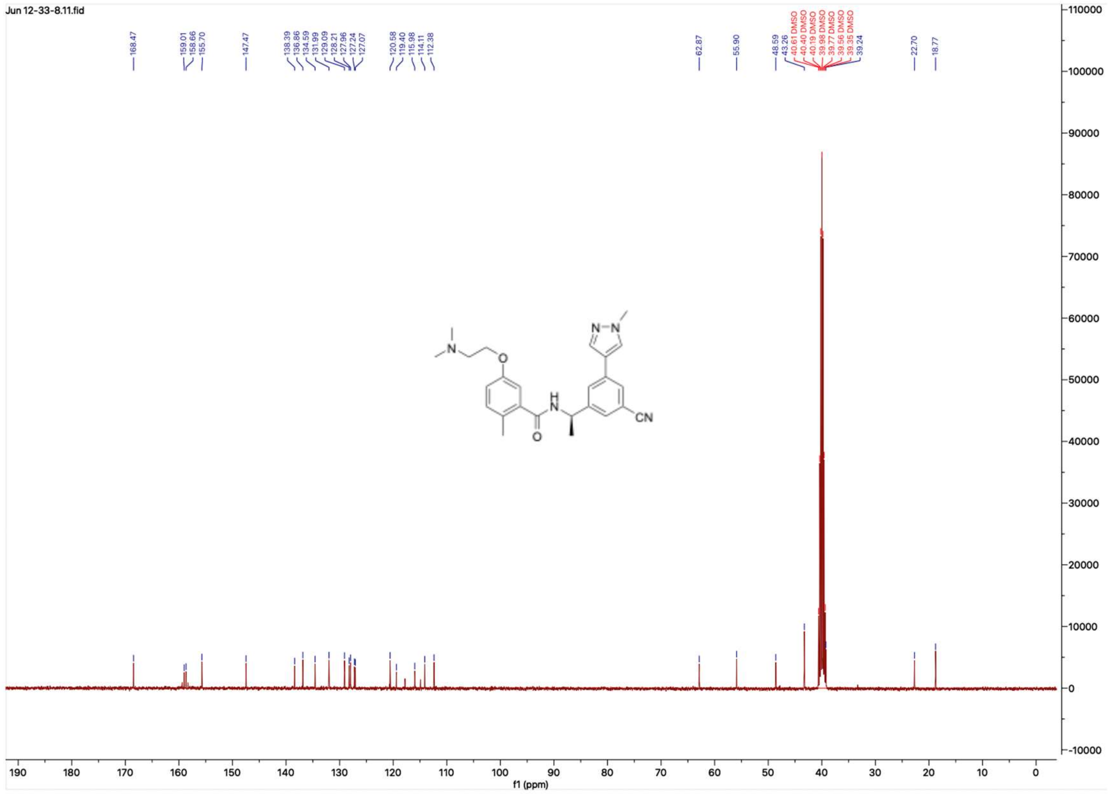

**HNMR and CNMR spectra of Jun12145**

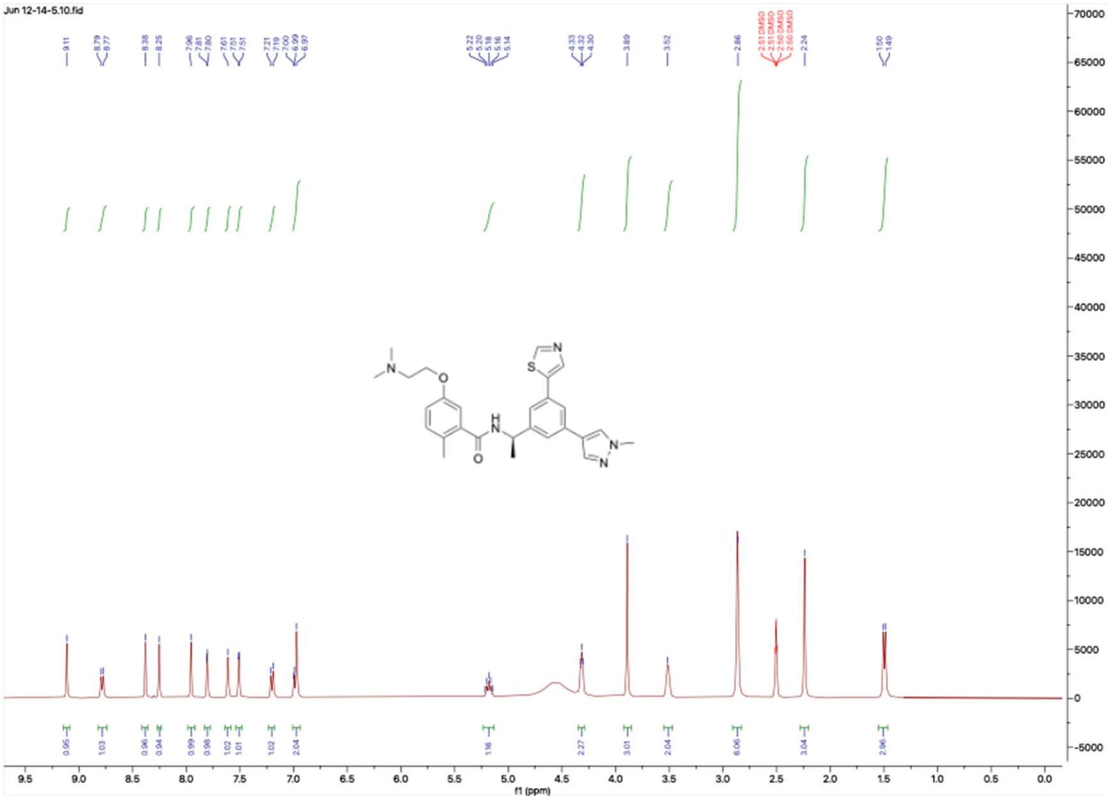

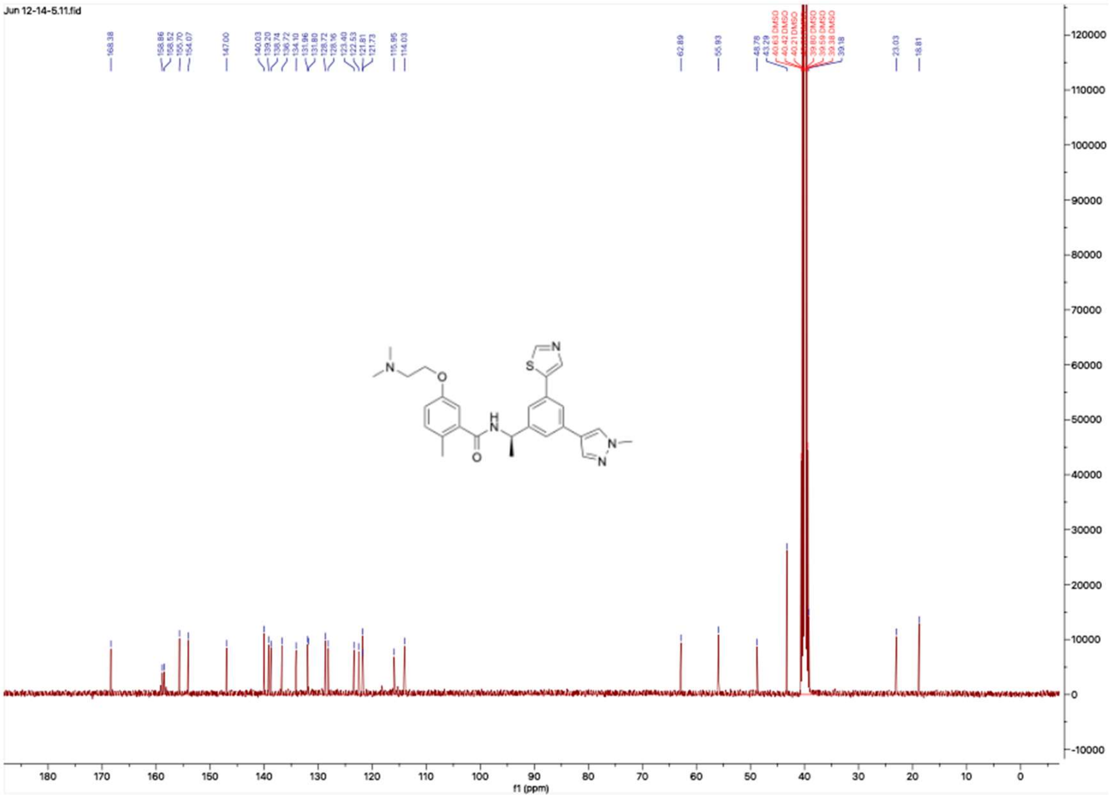

**HNMR and CNMR spectra of Jun12681**

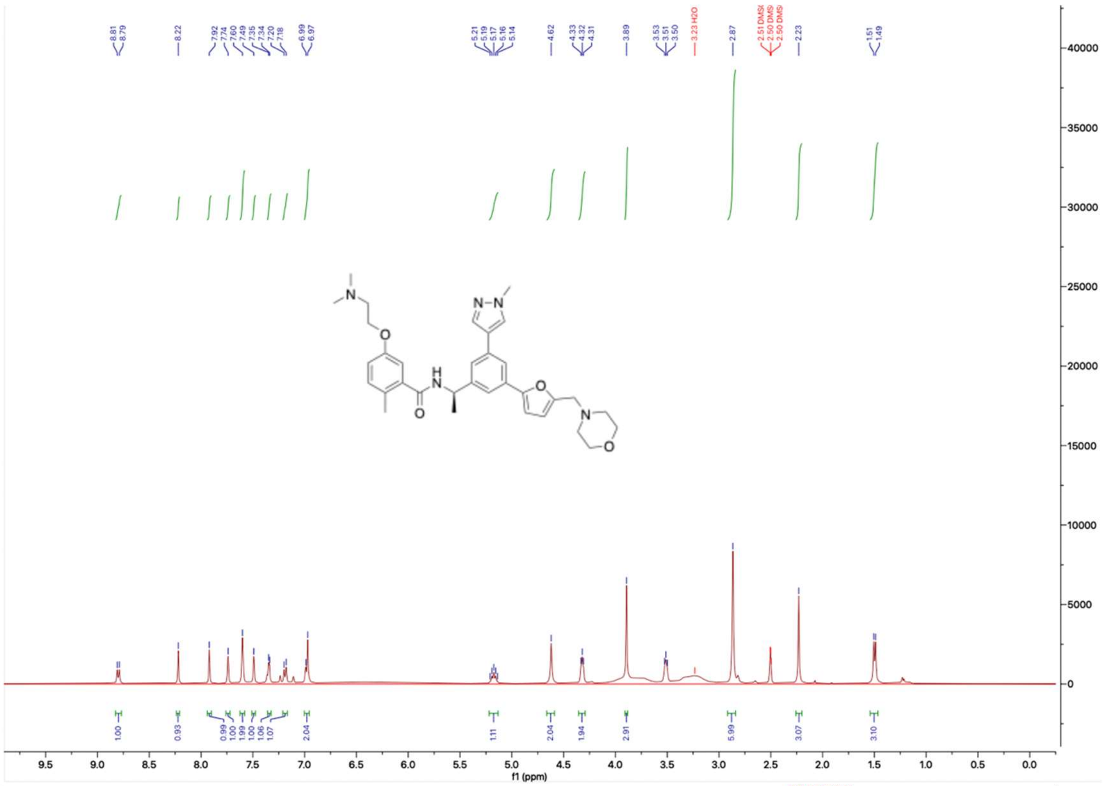

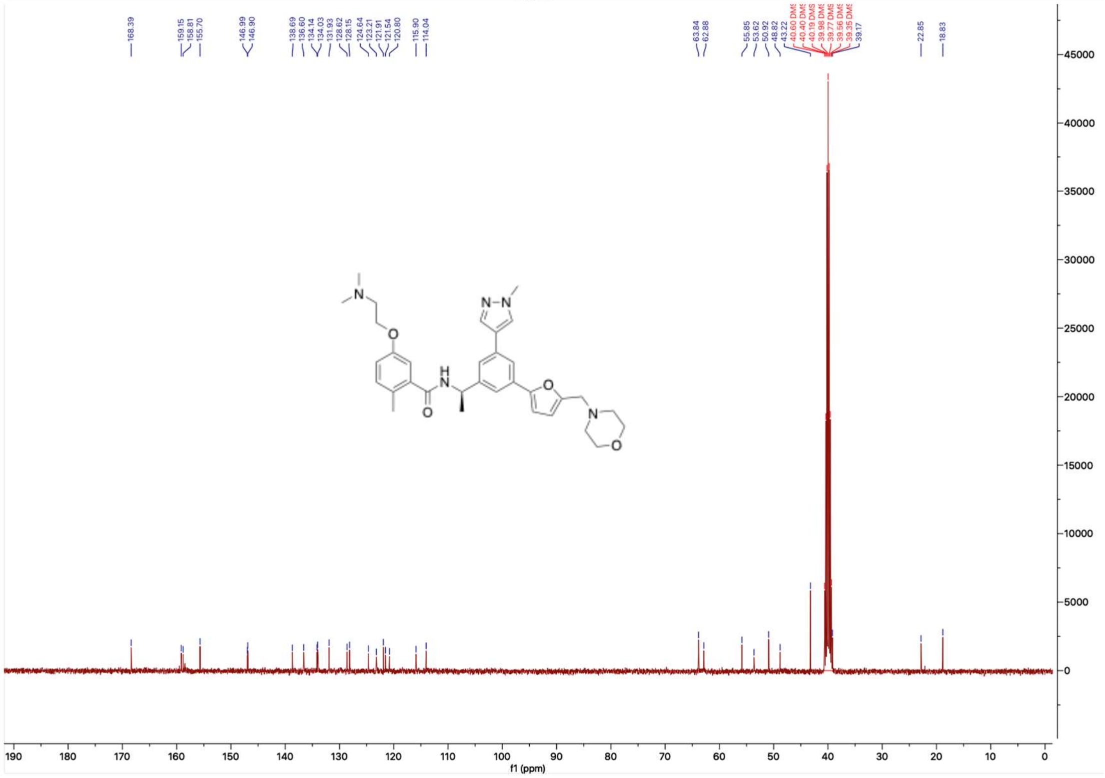

#### HNMR and CNMR spectra of Jun121911

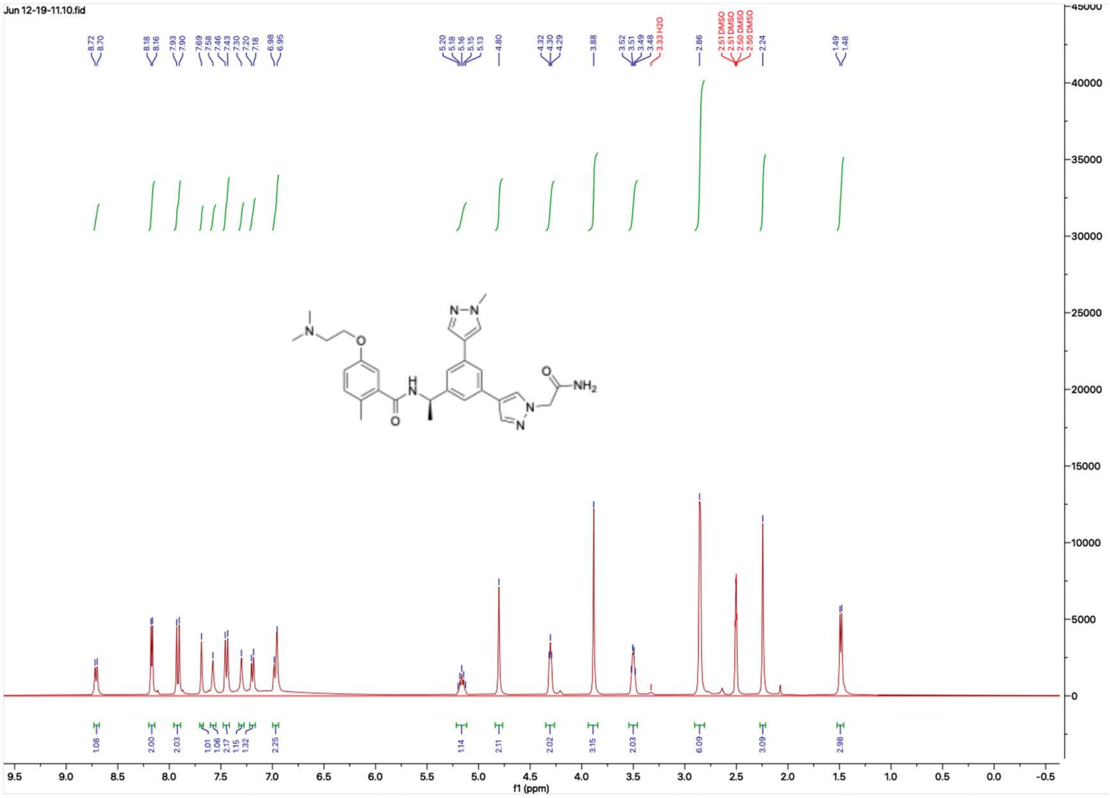

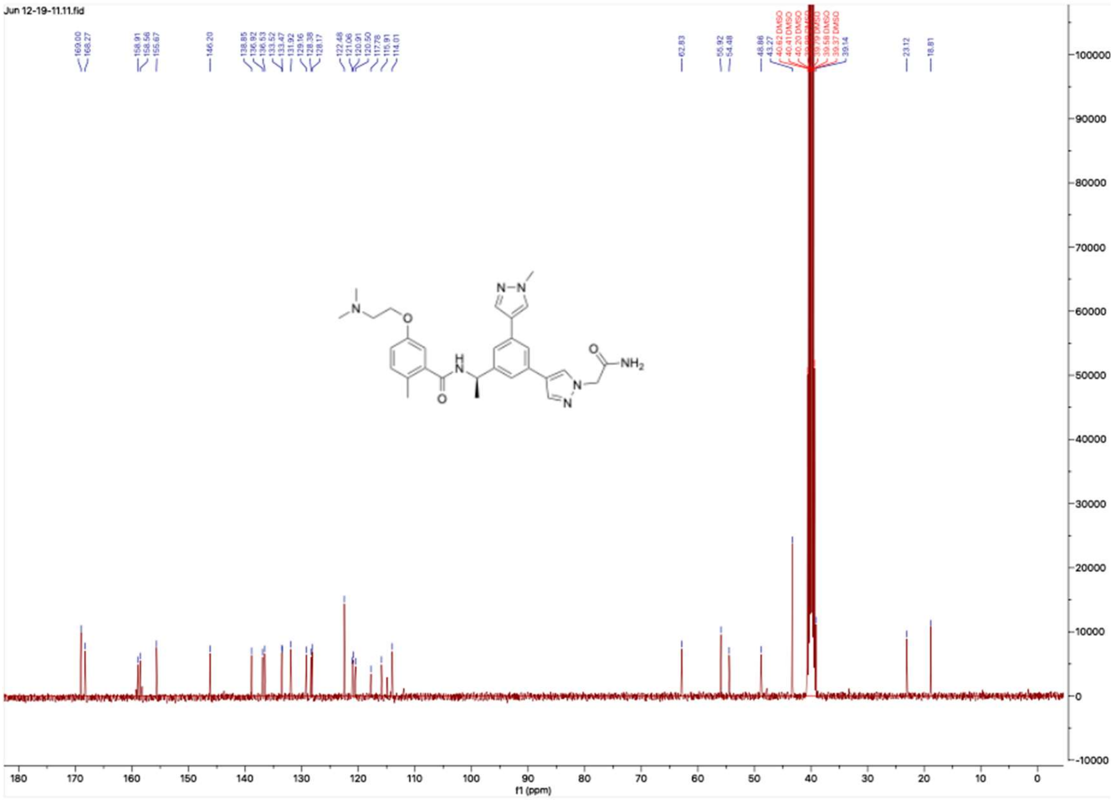

### HNMR and CNMR spectra of Jun12336

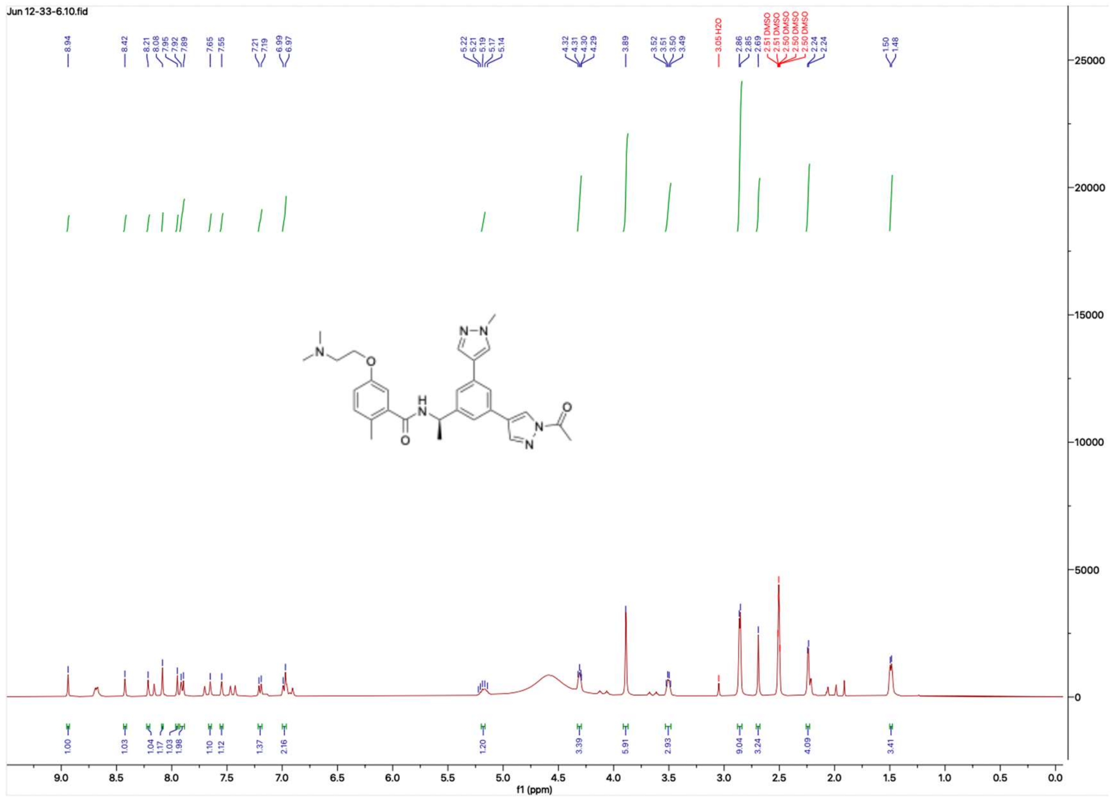

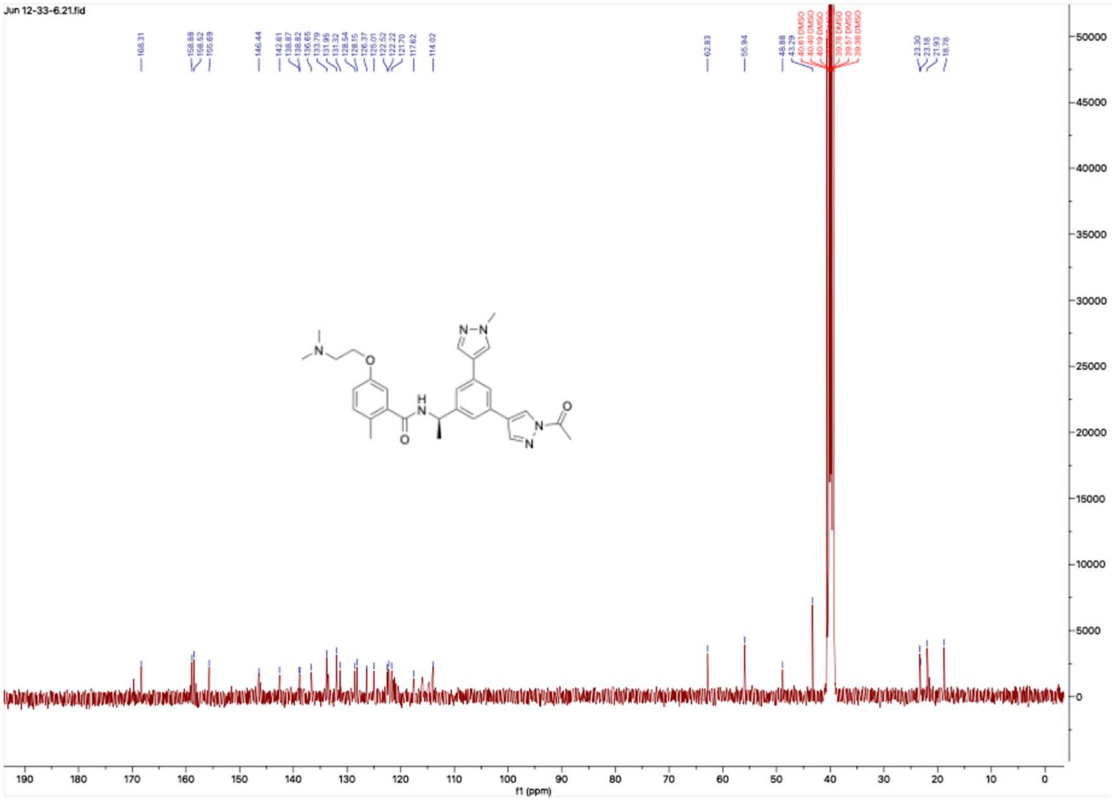

#### HNMR and CNMR spectra of Jun12467

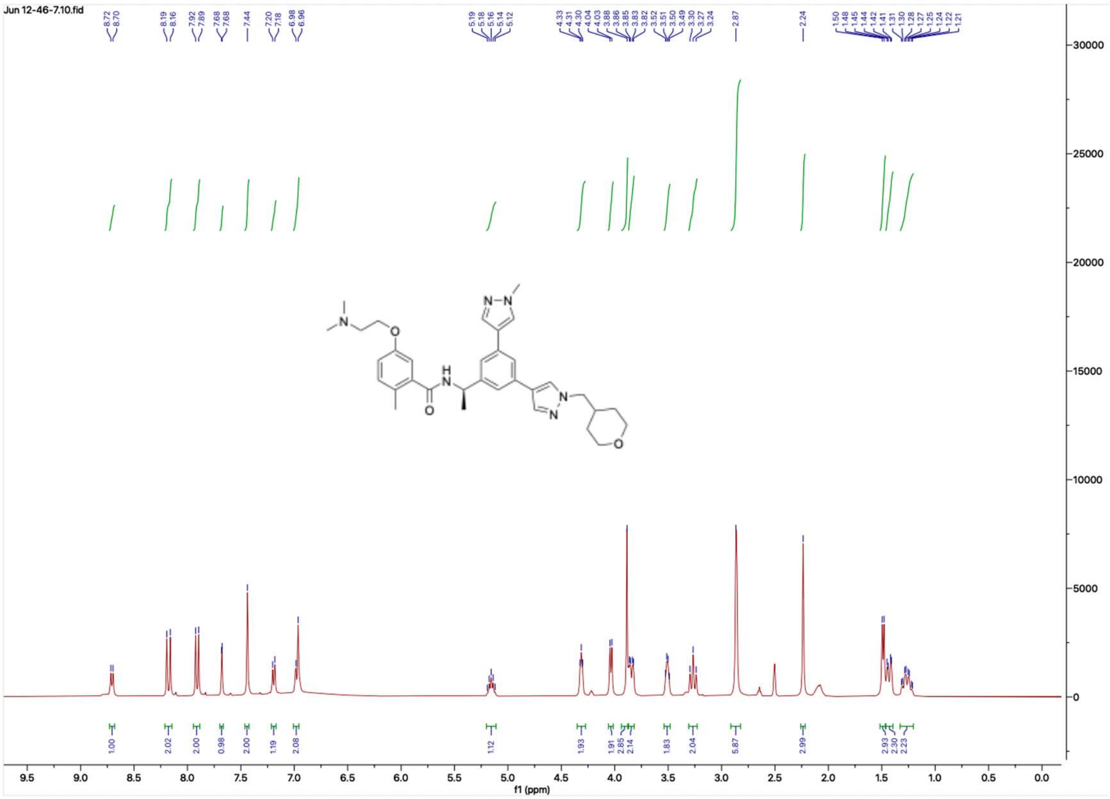


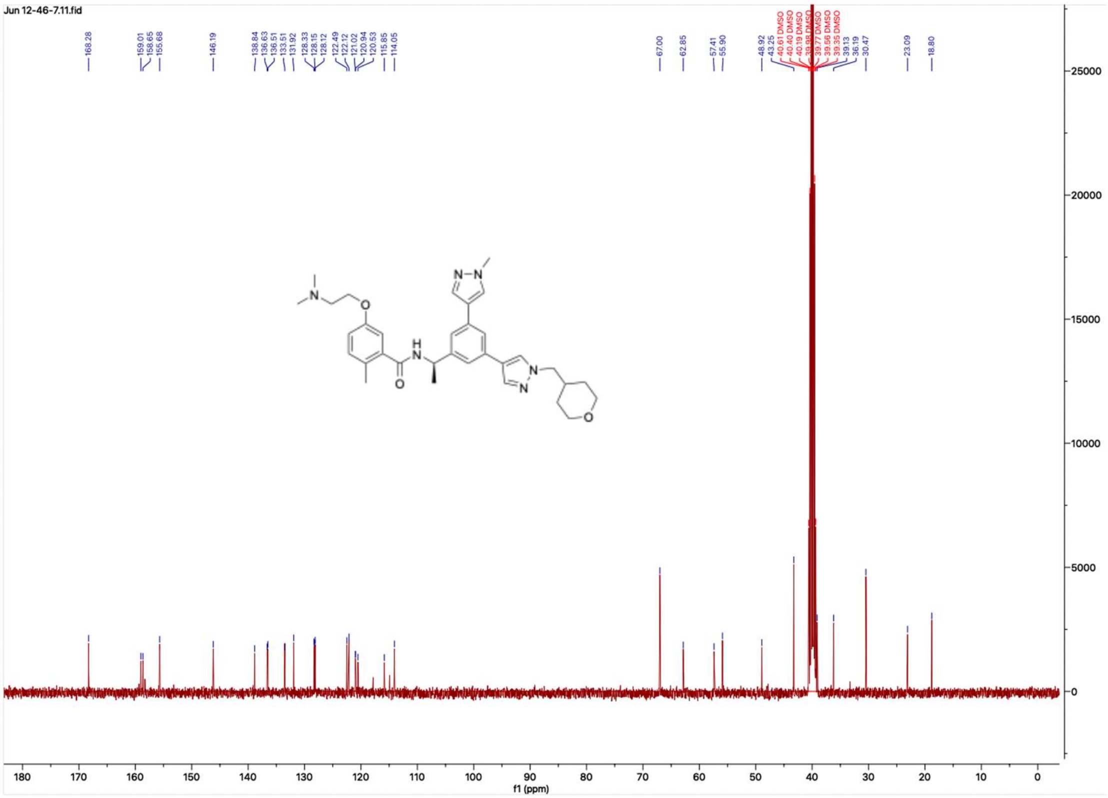

#### HNMR and CNMR spectra of Jun12351

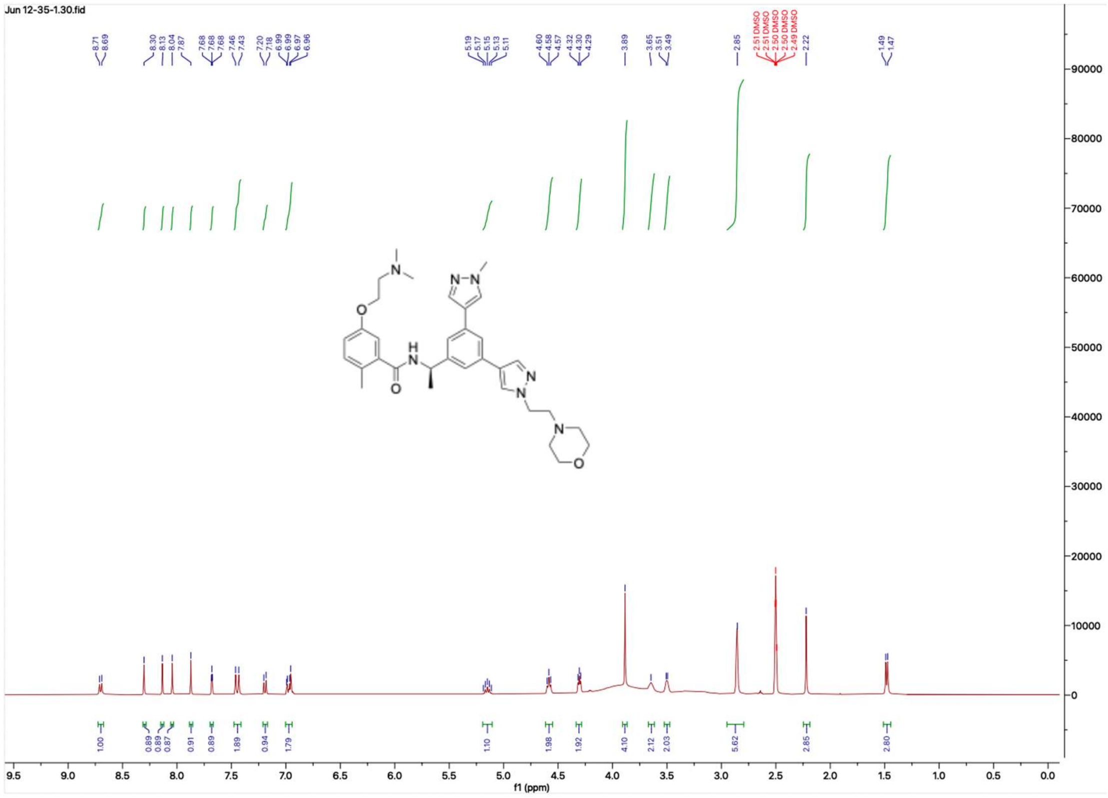


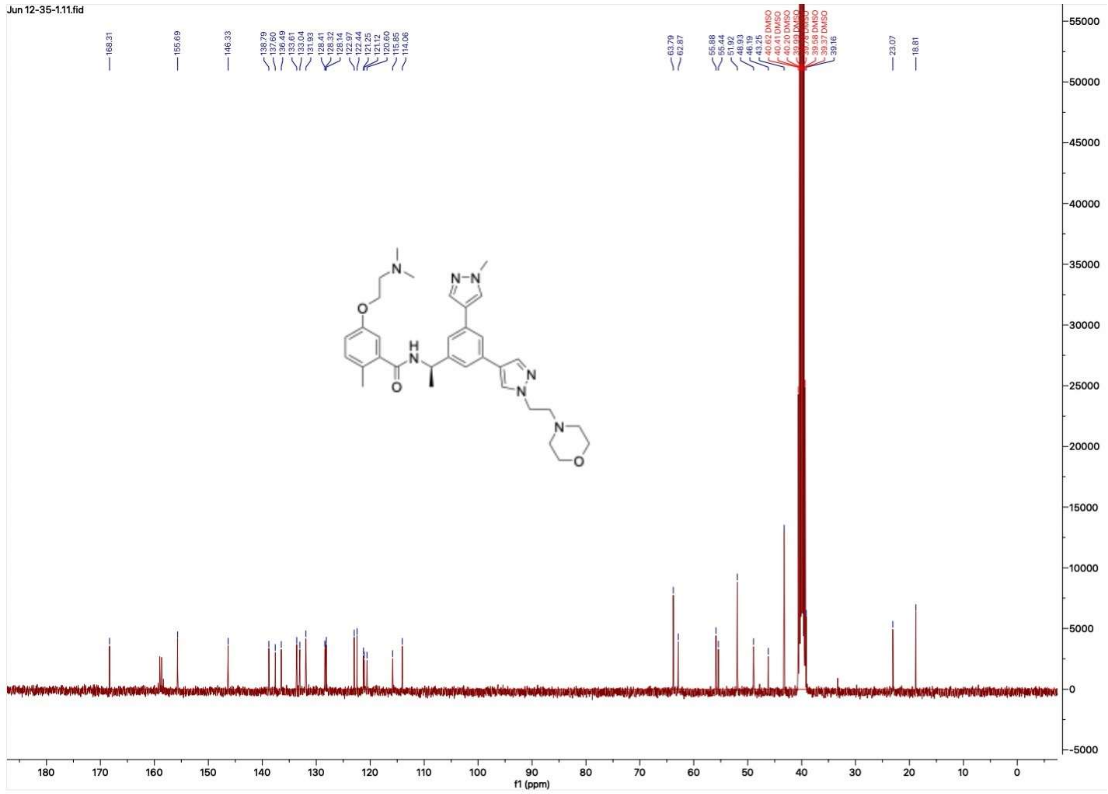

### HNMR and CNMR spectra of Jun12199

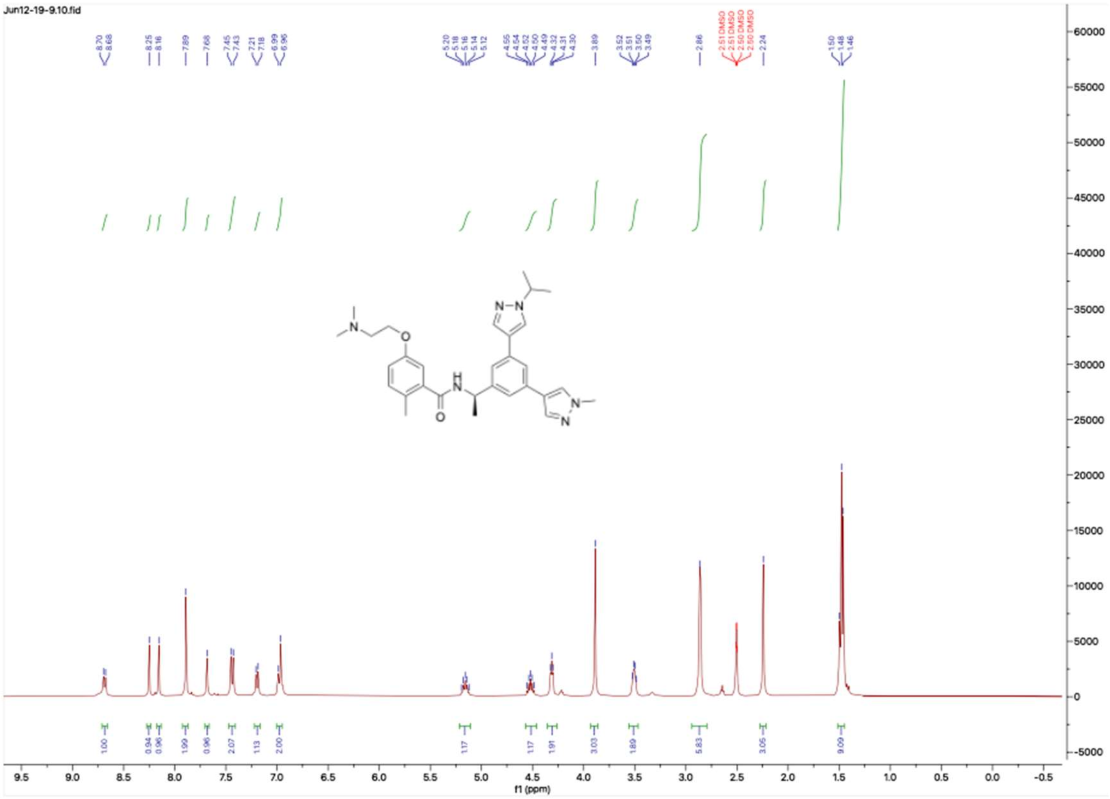


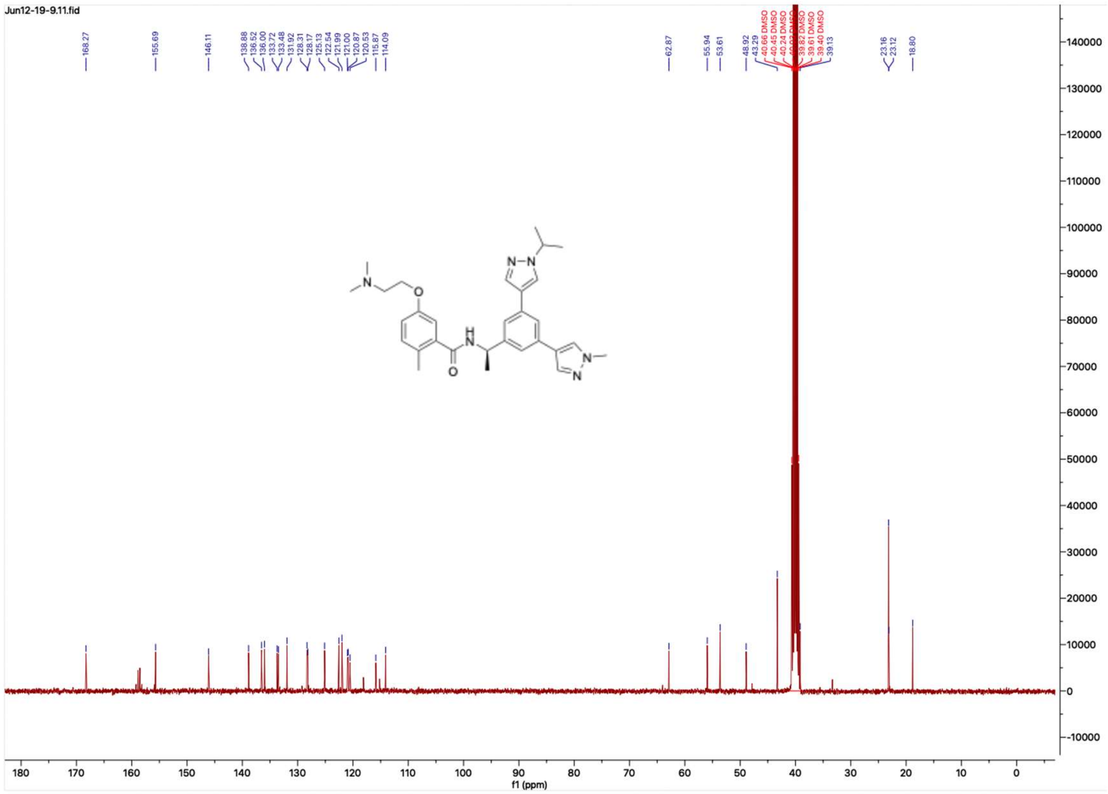

**HNMR and CNMR spectra of Jun12197**

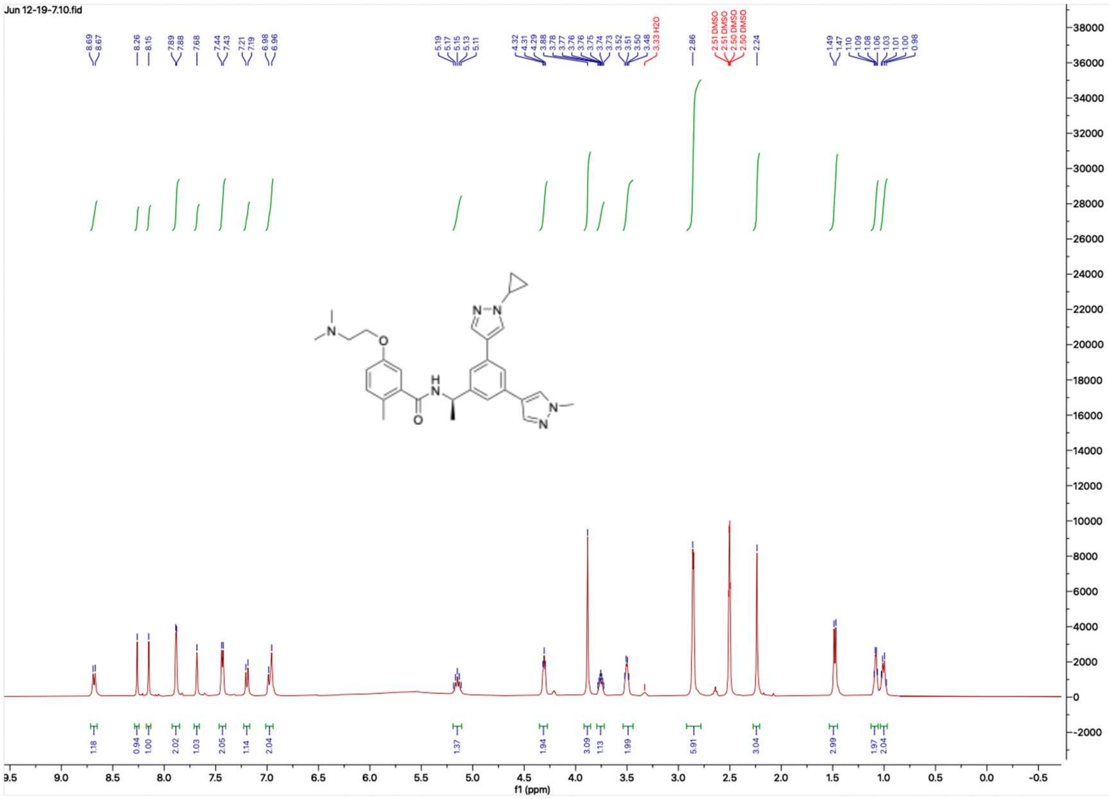

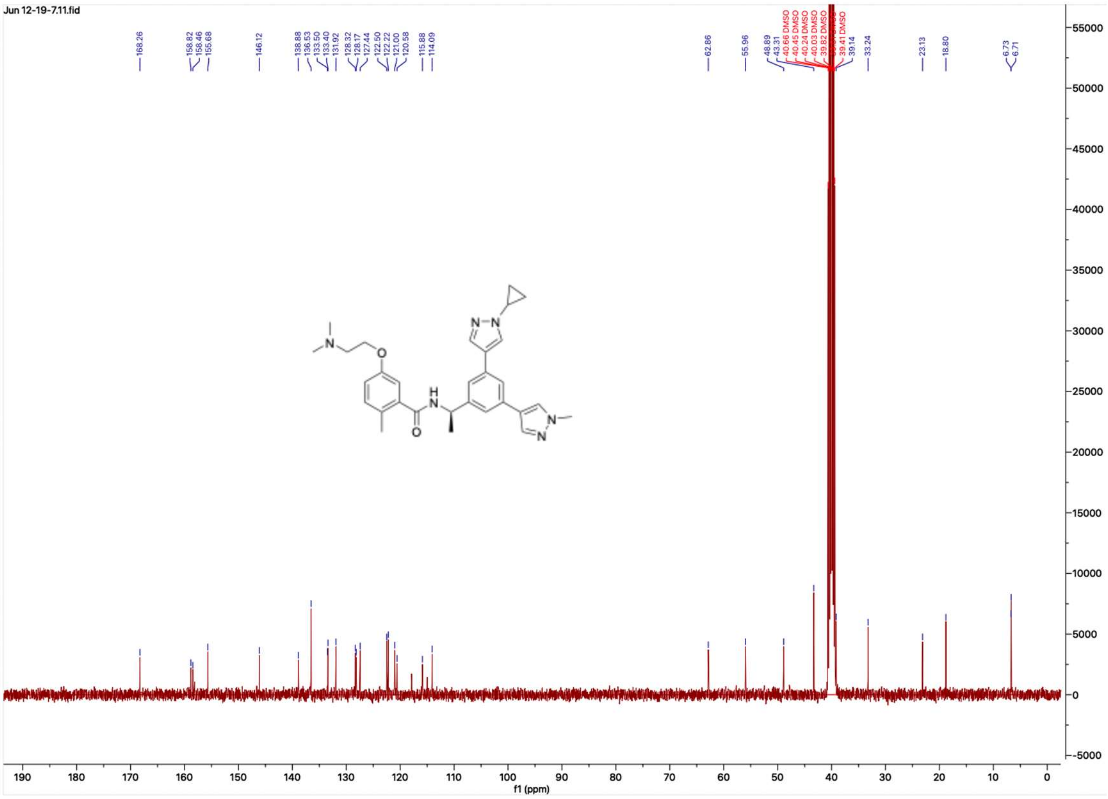

**HNMR and CNMR spectra of Jun12603**

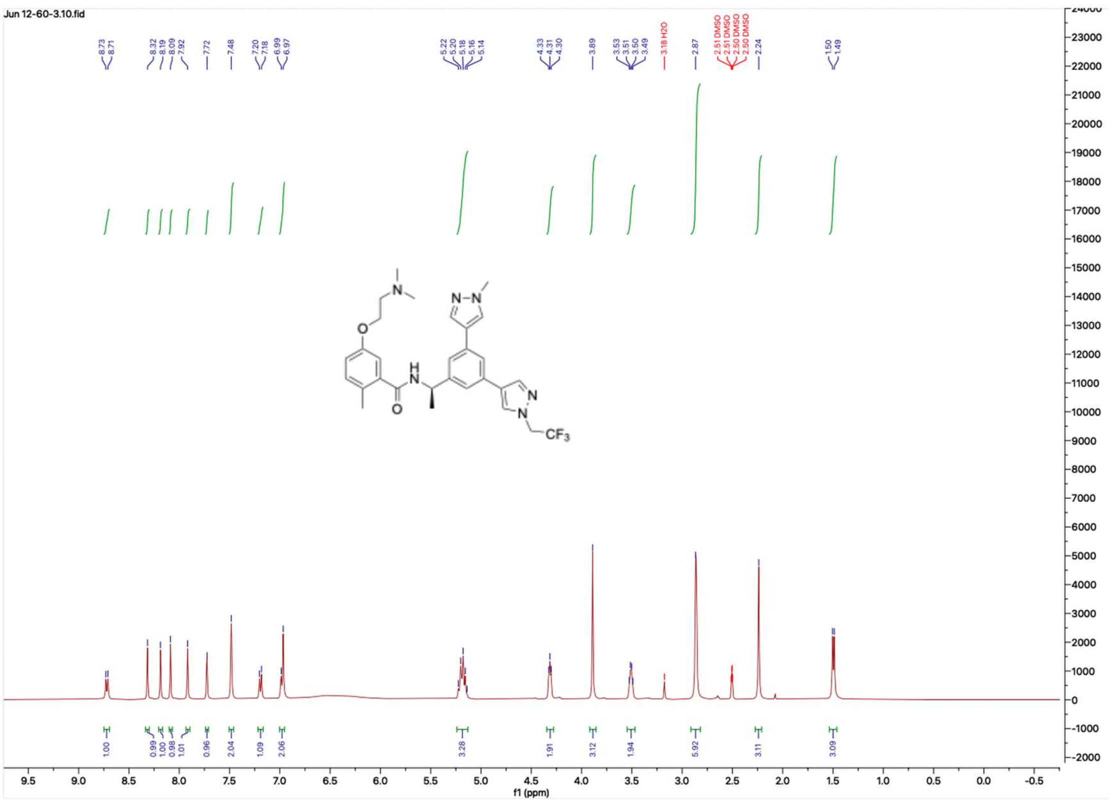

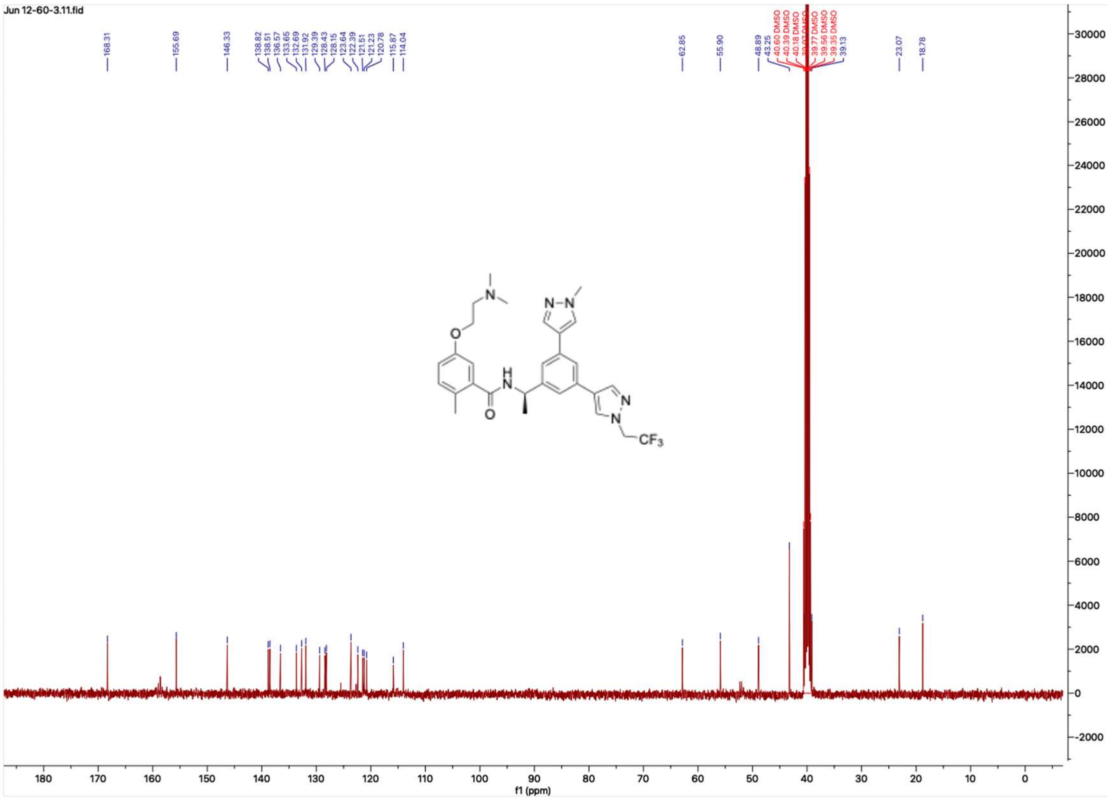

#### HNMR and CNMR spectra of Jun12446

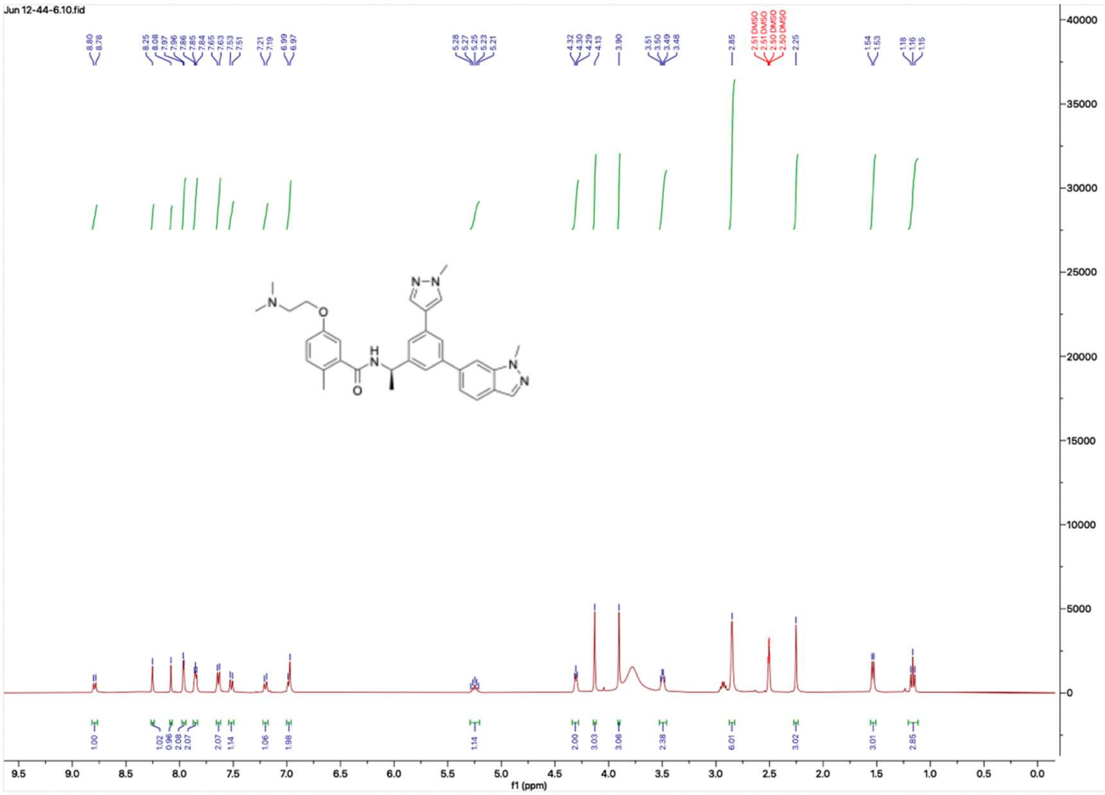


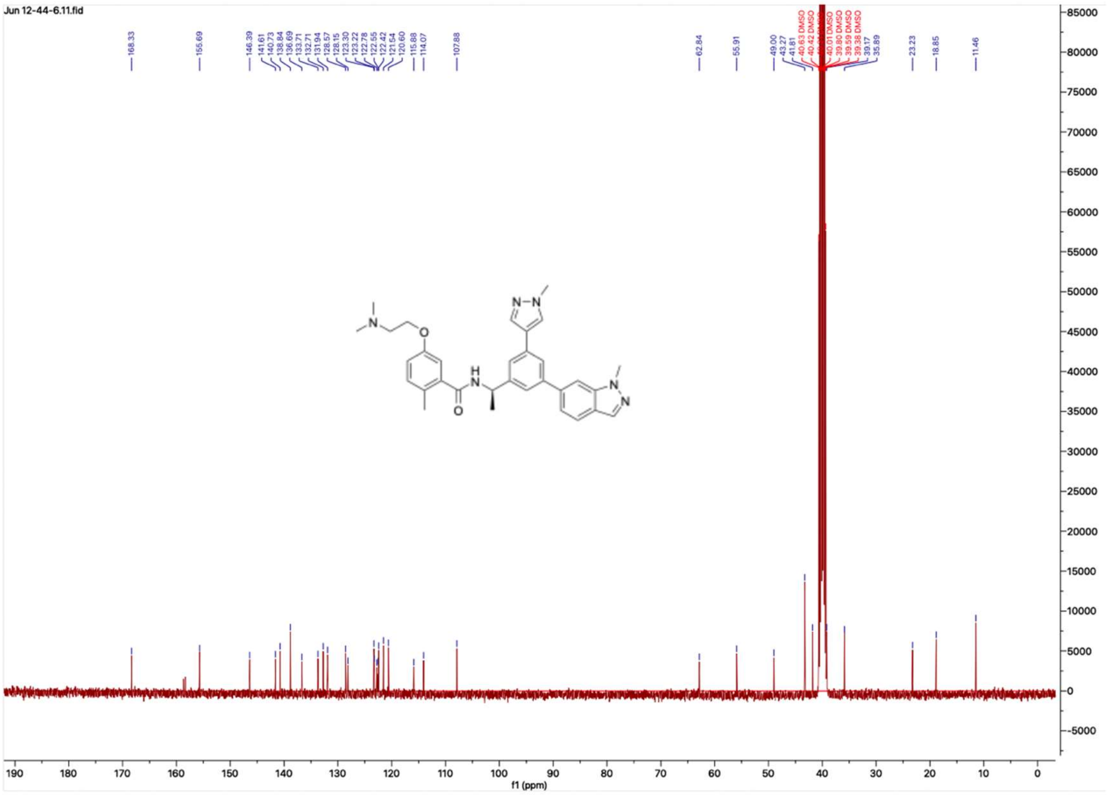

**HNMR and CNMR spectra of Jun12278**

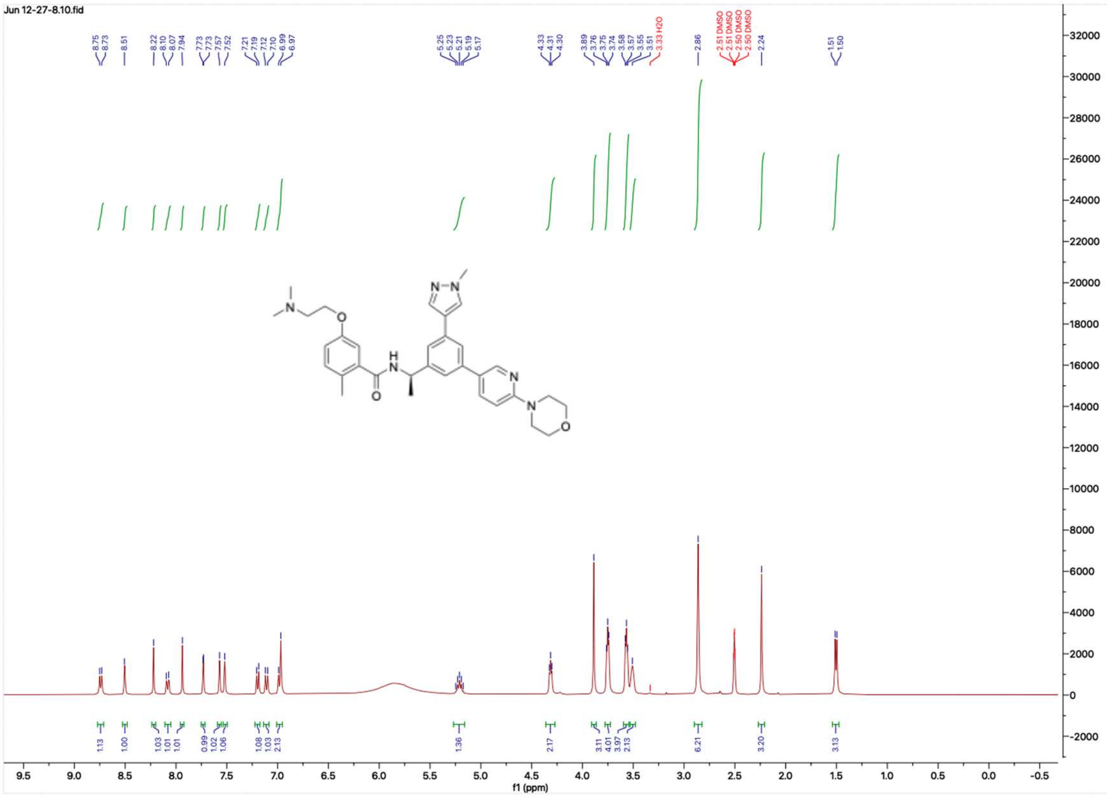


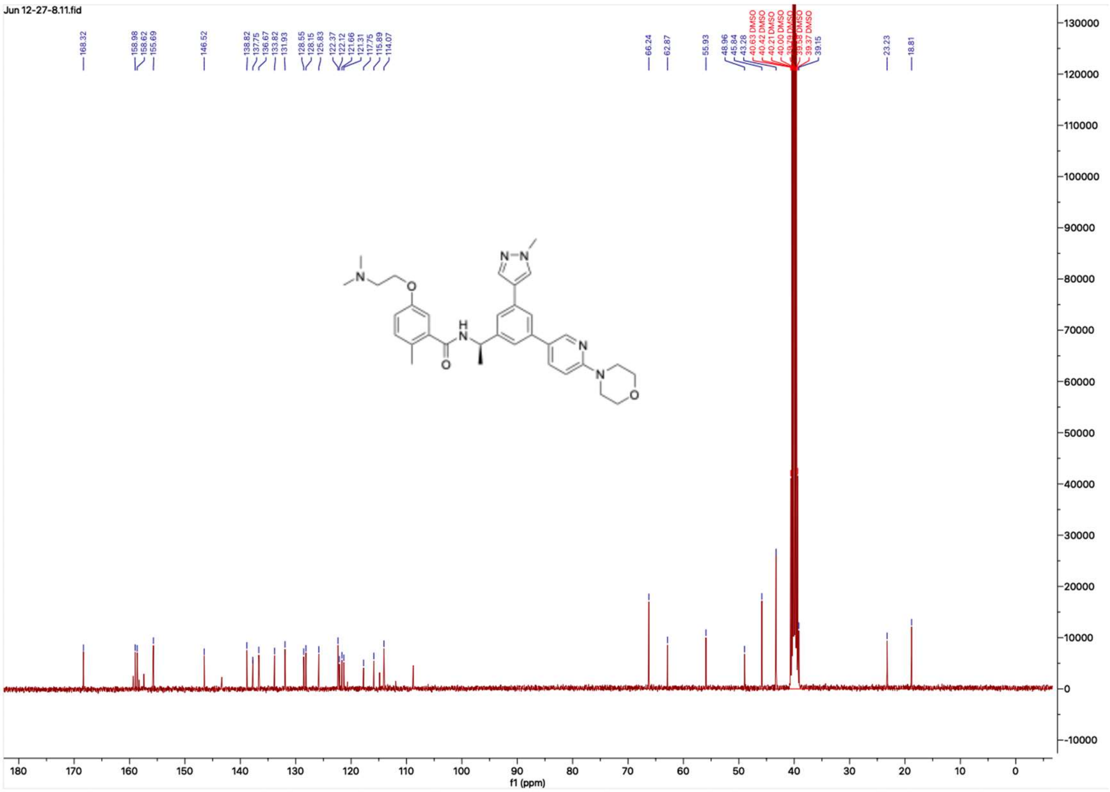

**HNMR and CNMR spectra of Jun12162**

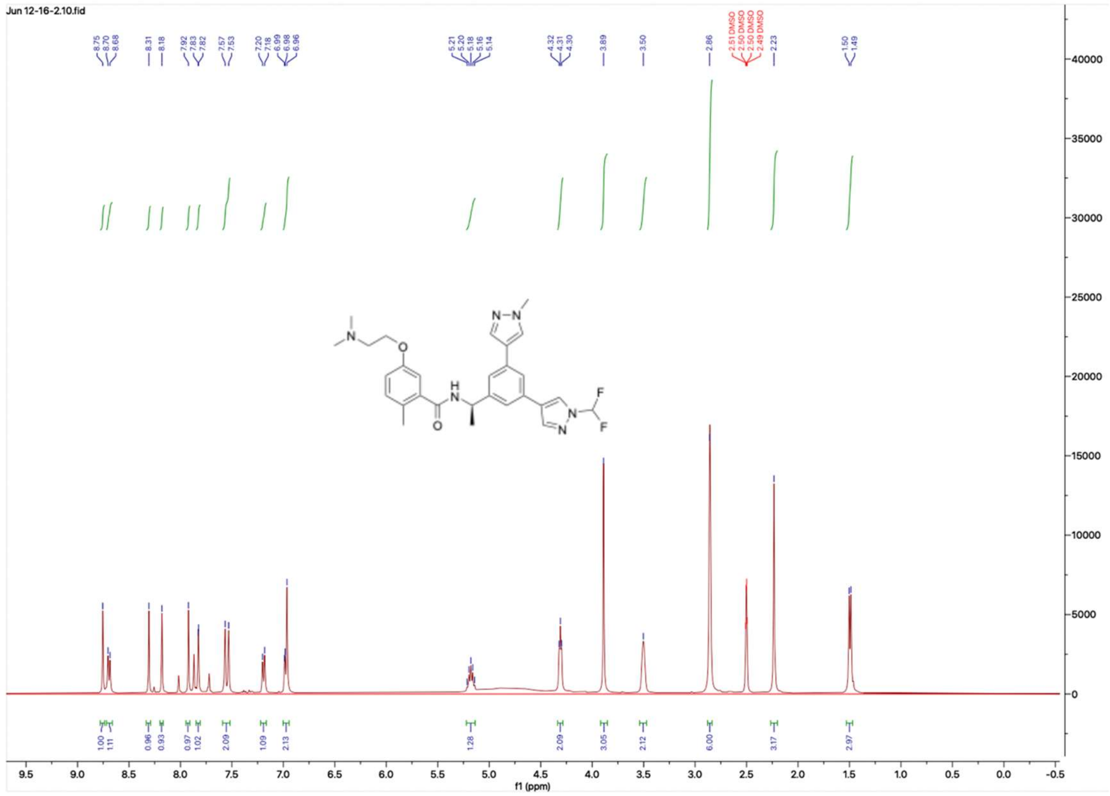

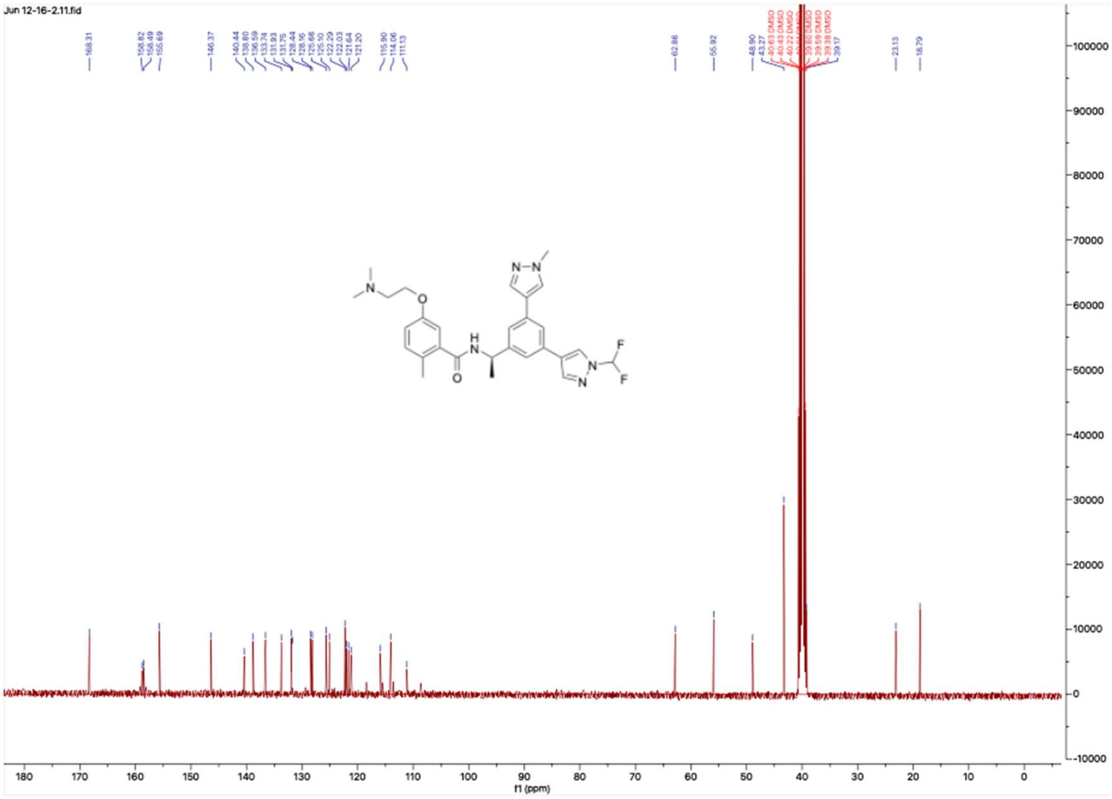

**HNMR and CNMR spectra of Jun12208**

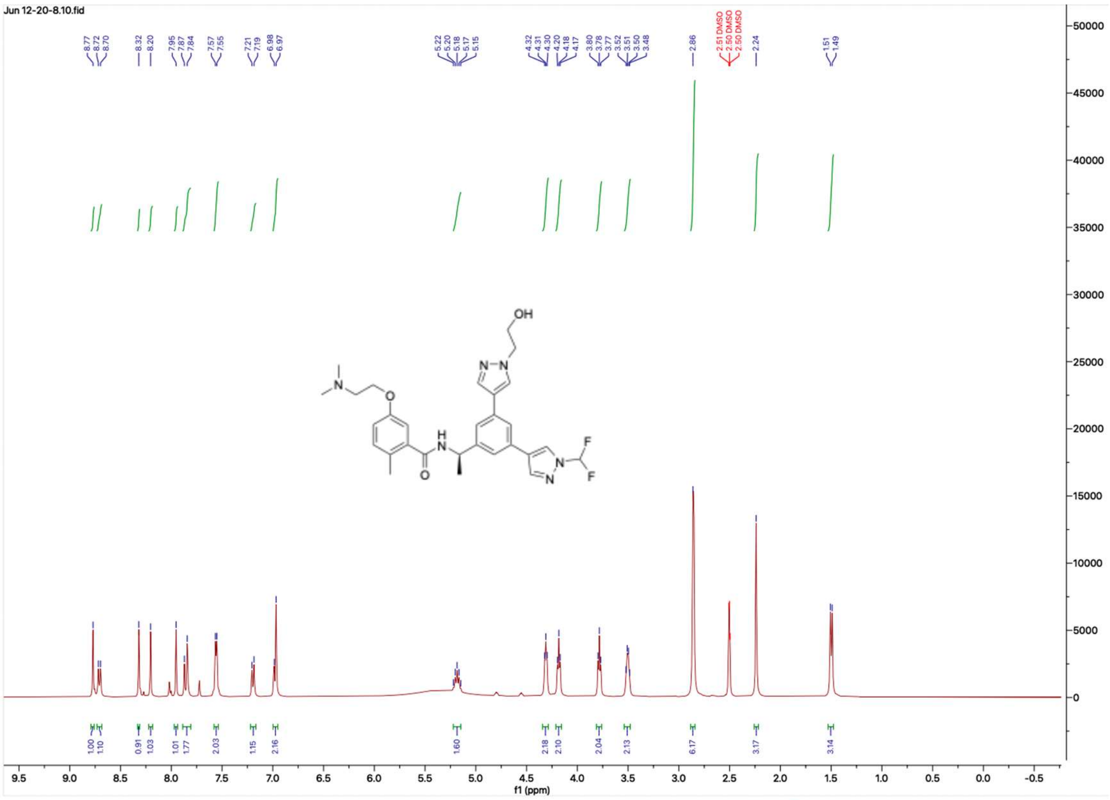

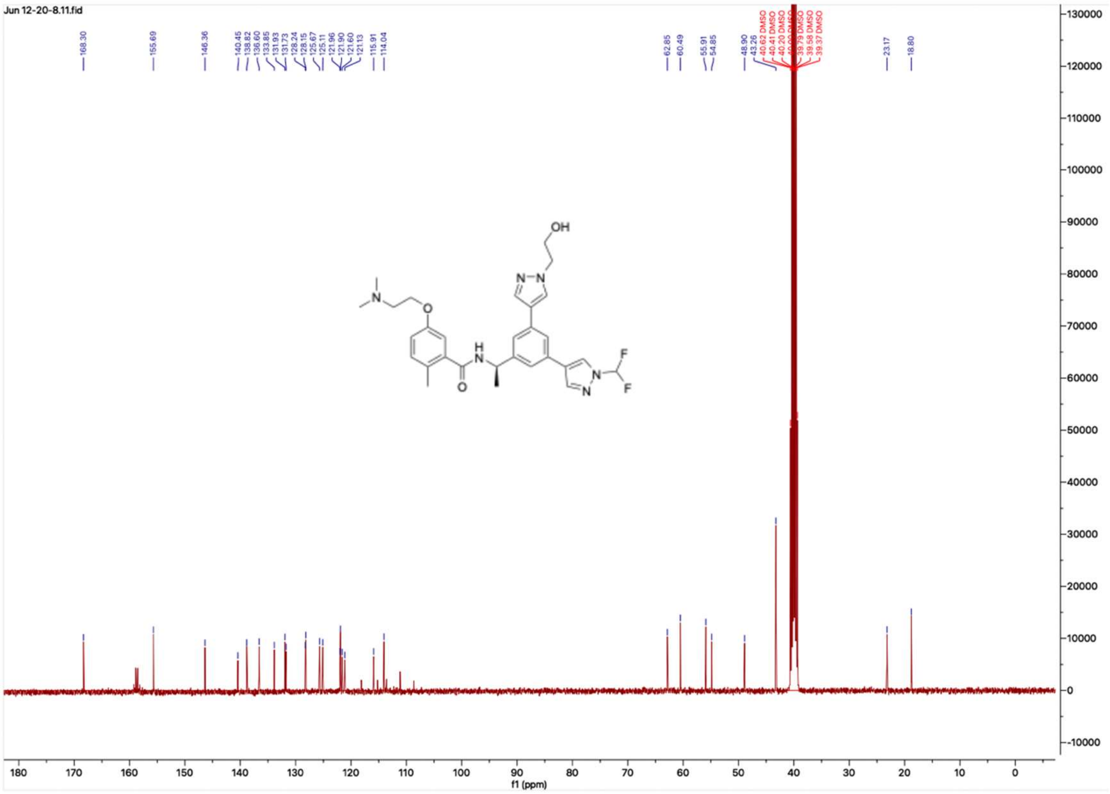

#### HNMR and CNMR spectra of Jun12235

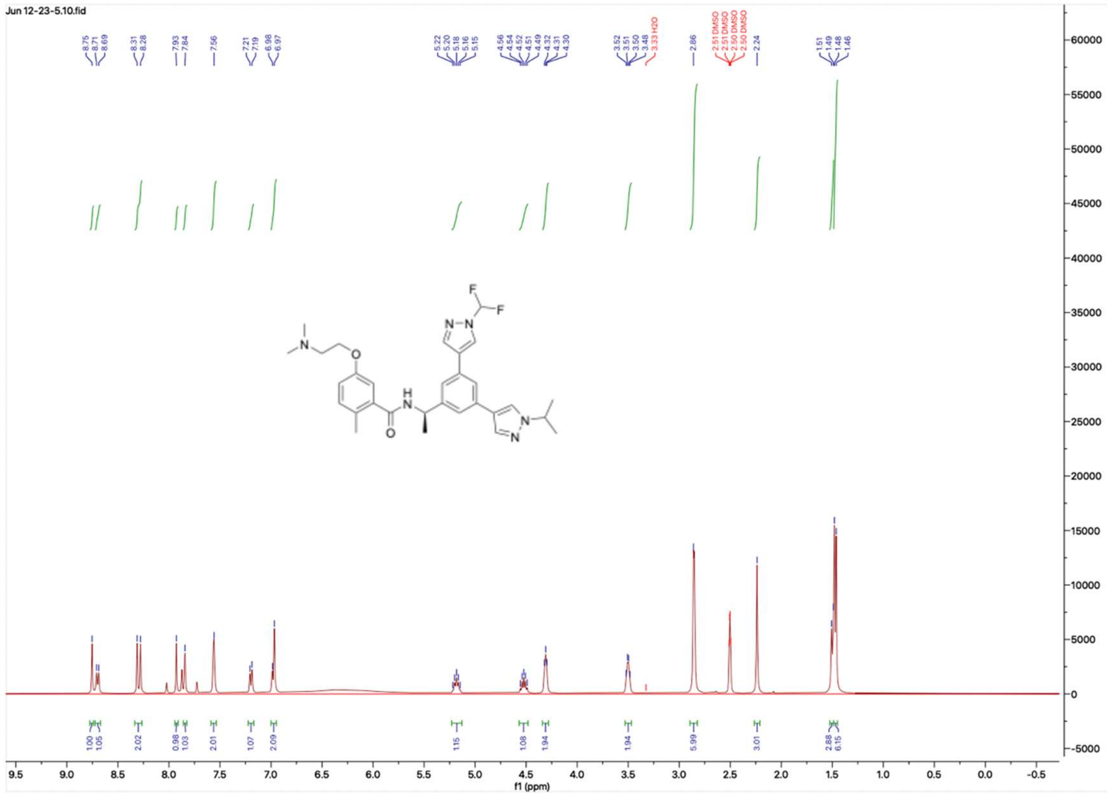


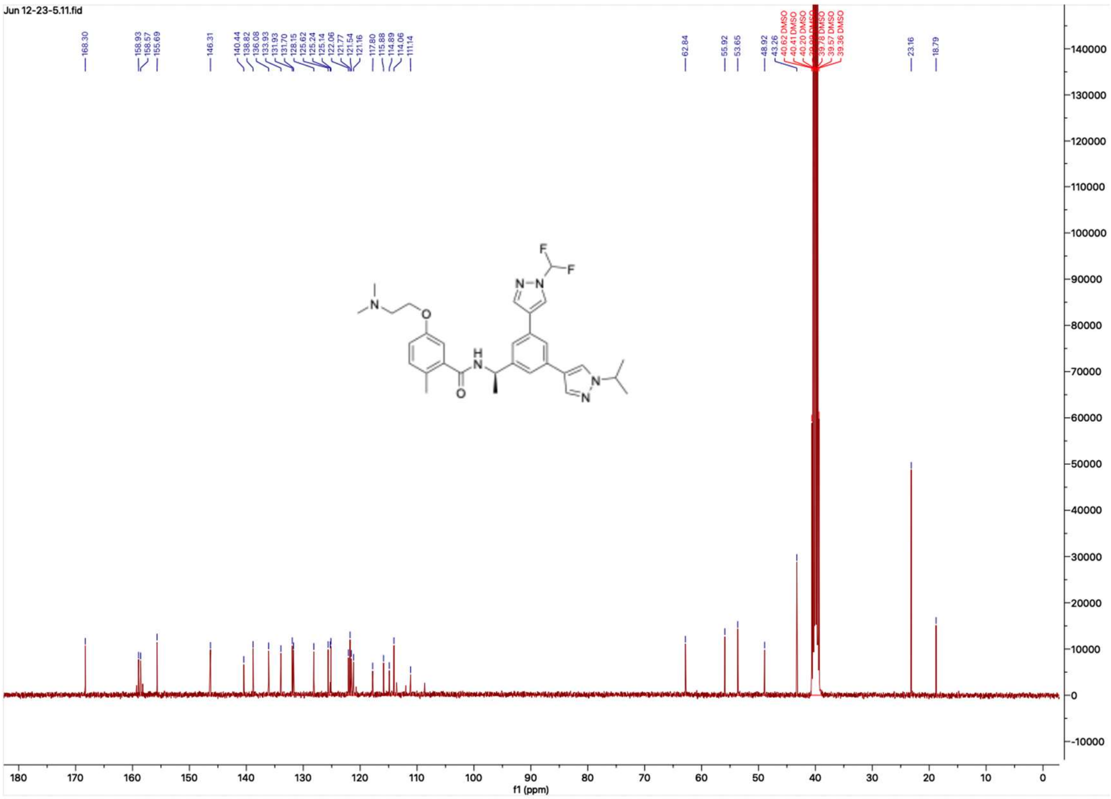

**HNMR and CNMR spectra of Jun12281**

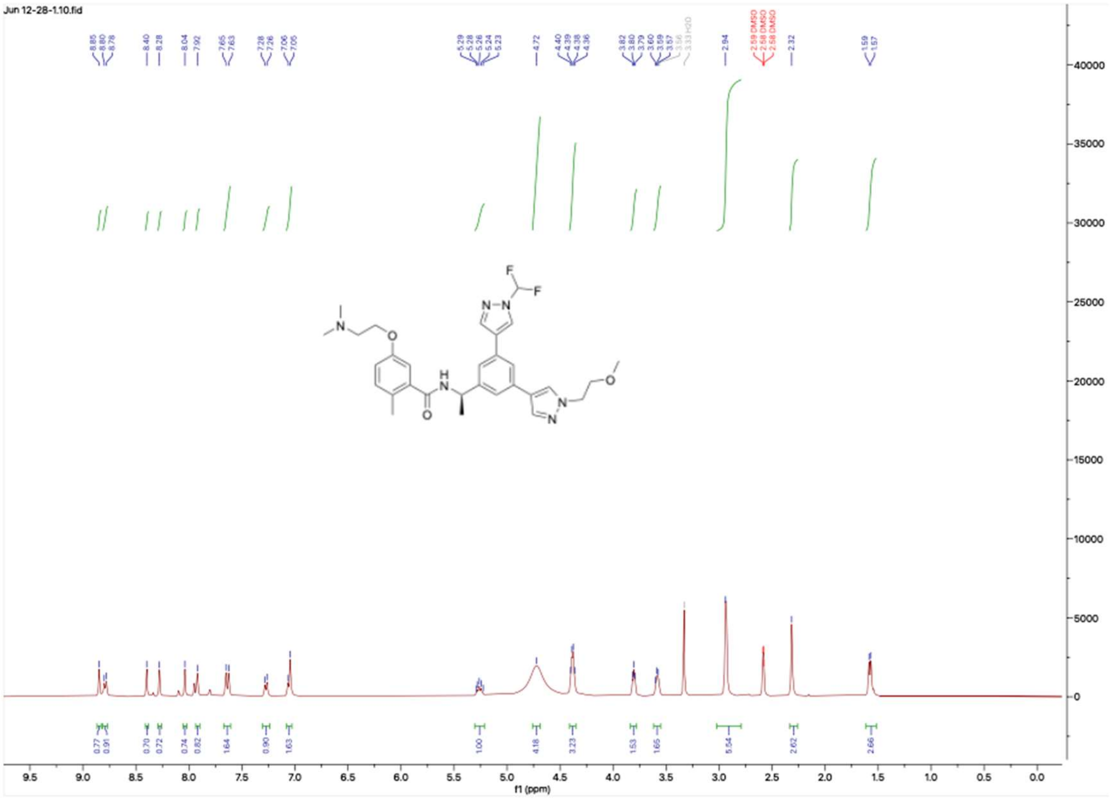


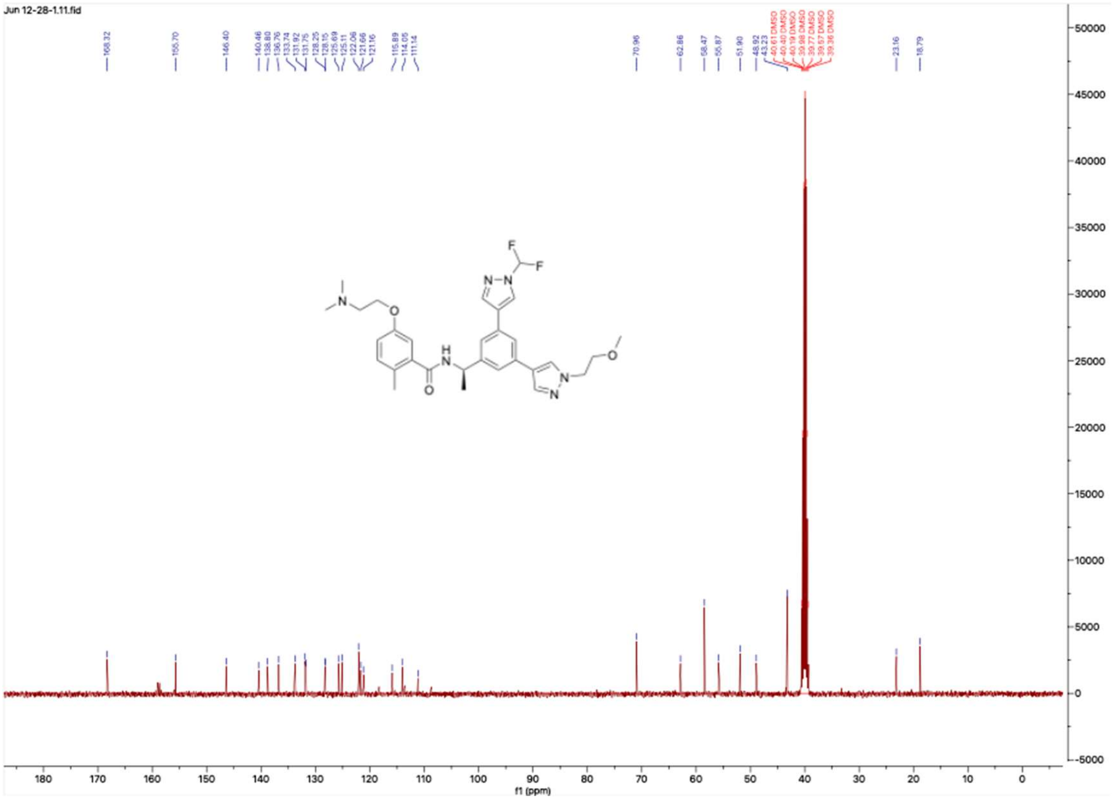

#### HNMR and CNMR spectra of Jun12303

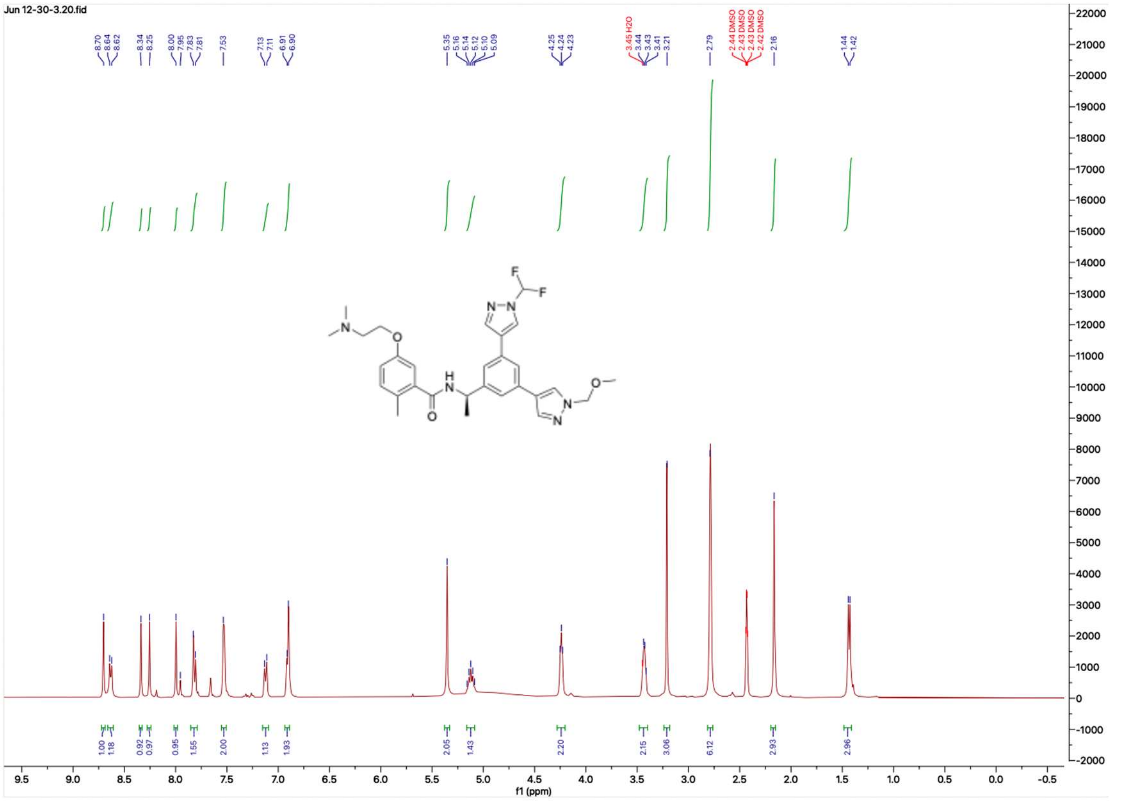

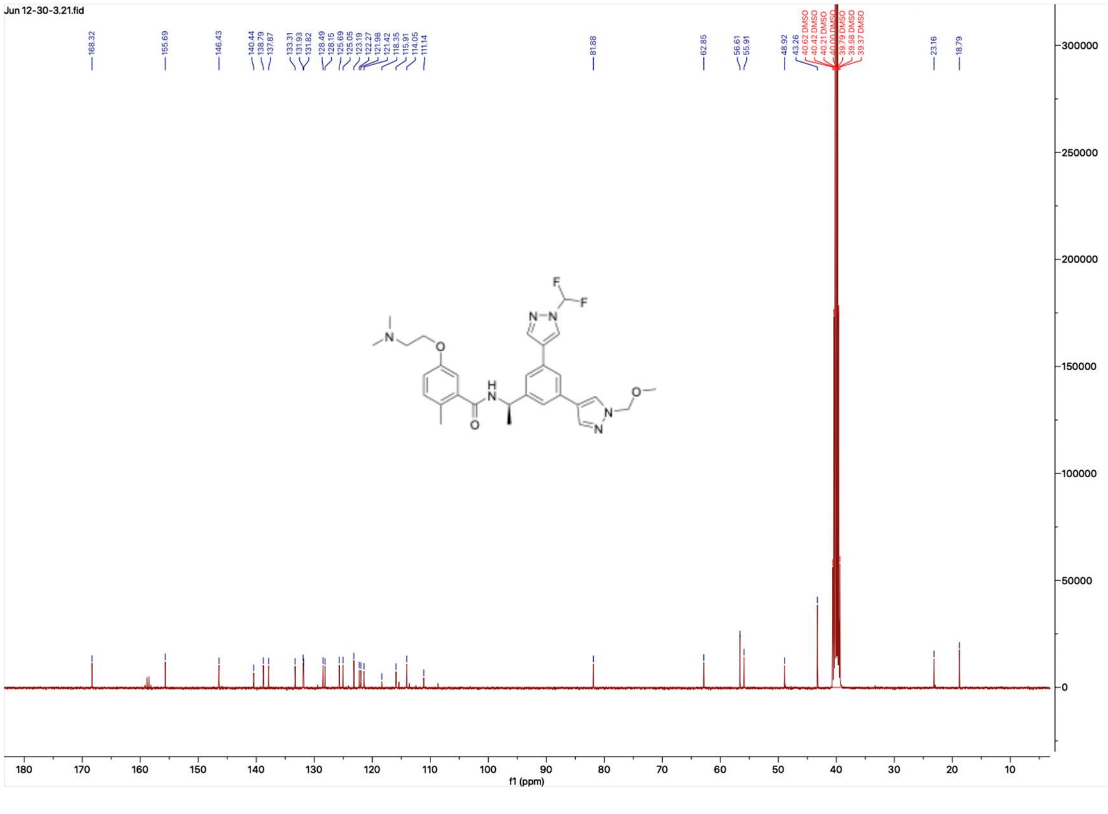

**HNMR and CNMR spectra of Jun12284**

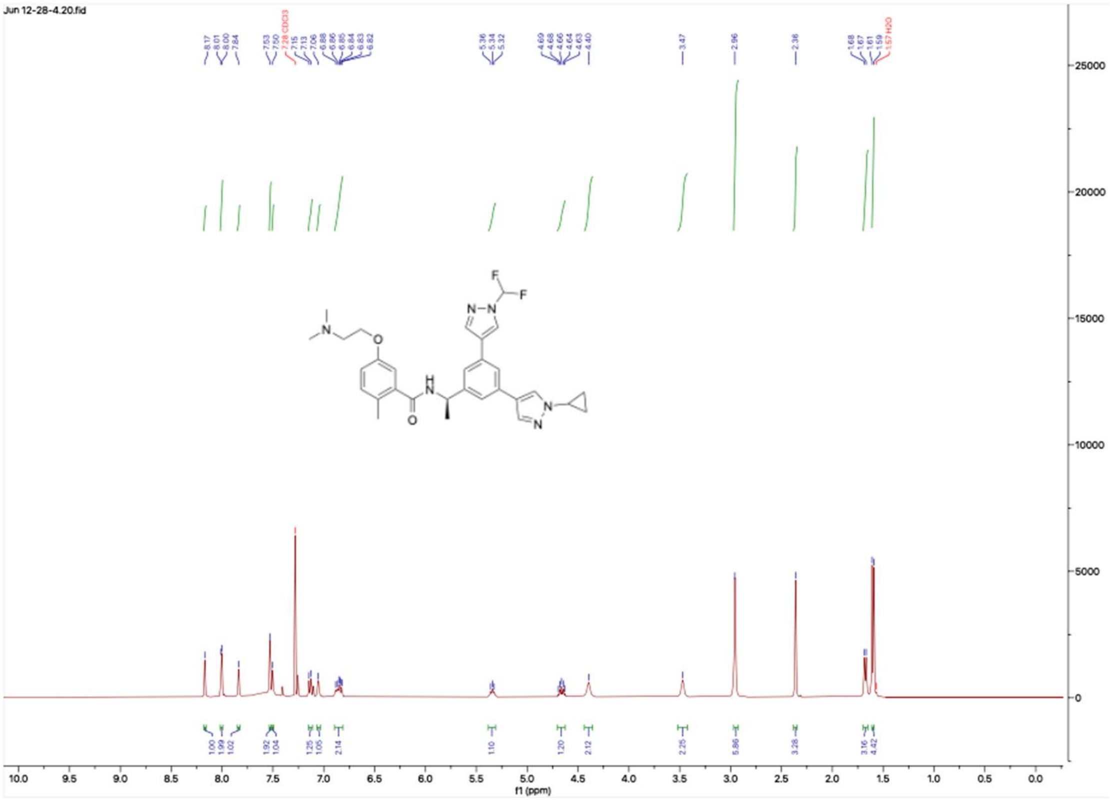

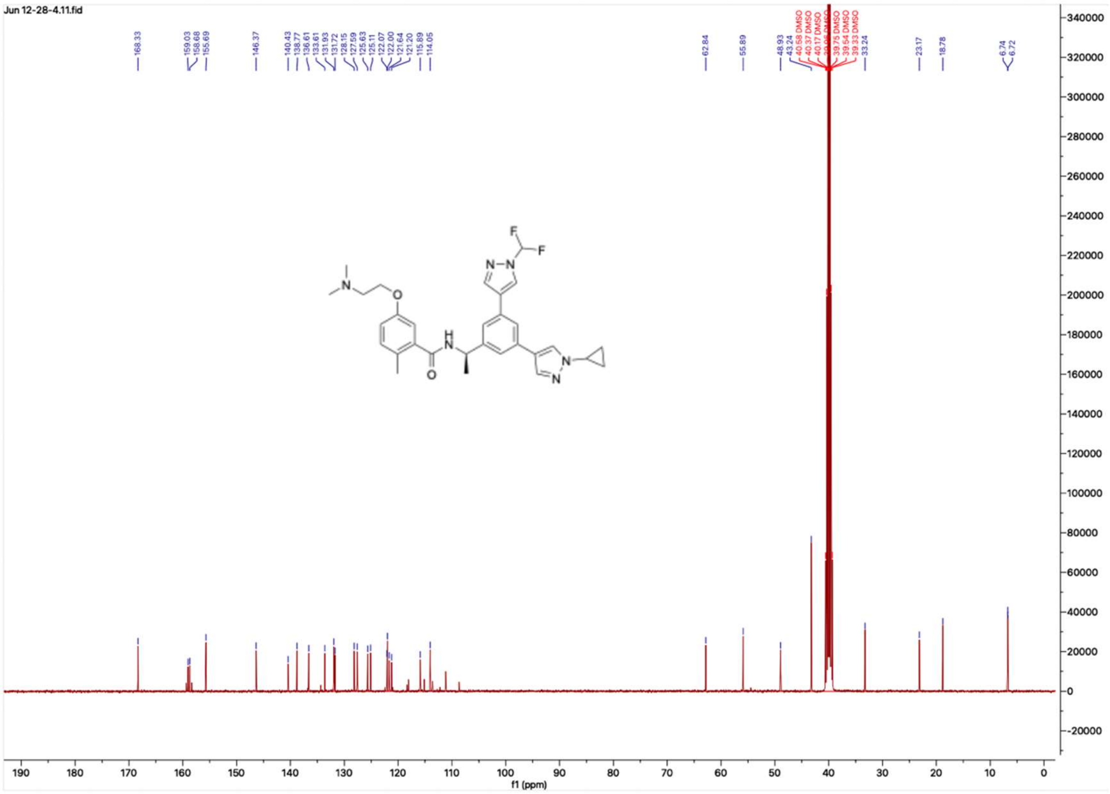

### HNMR and CNMR spectra of Jun12165

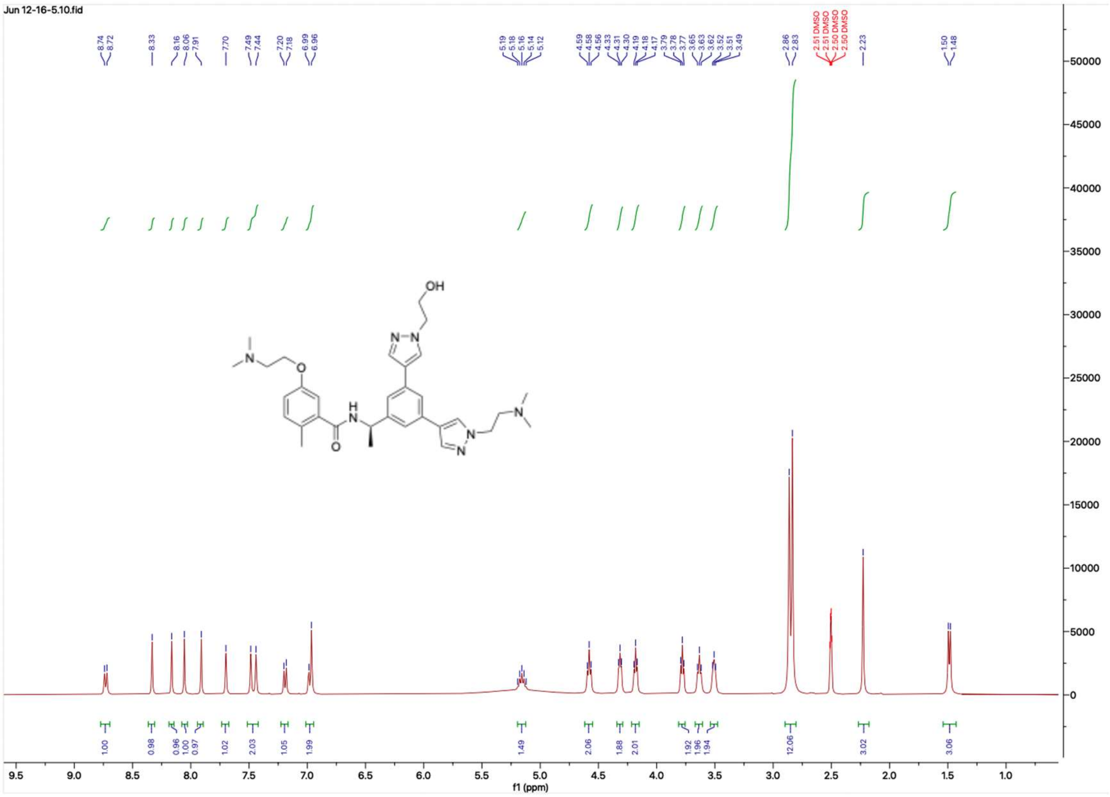


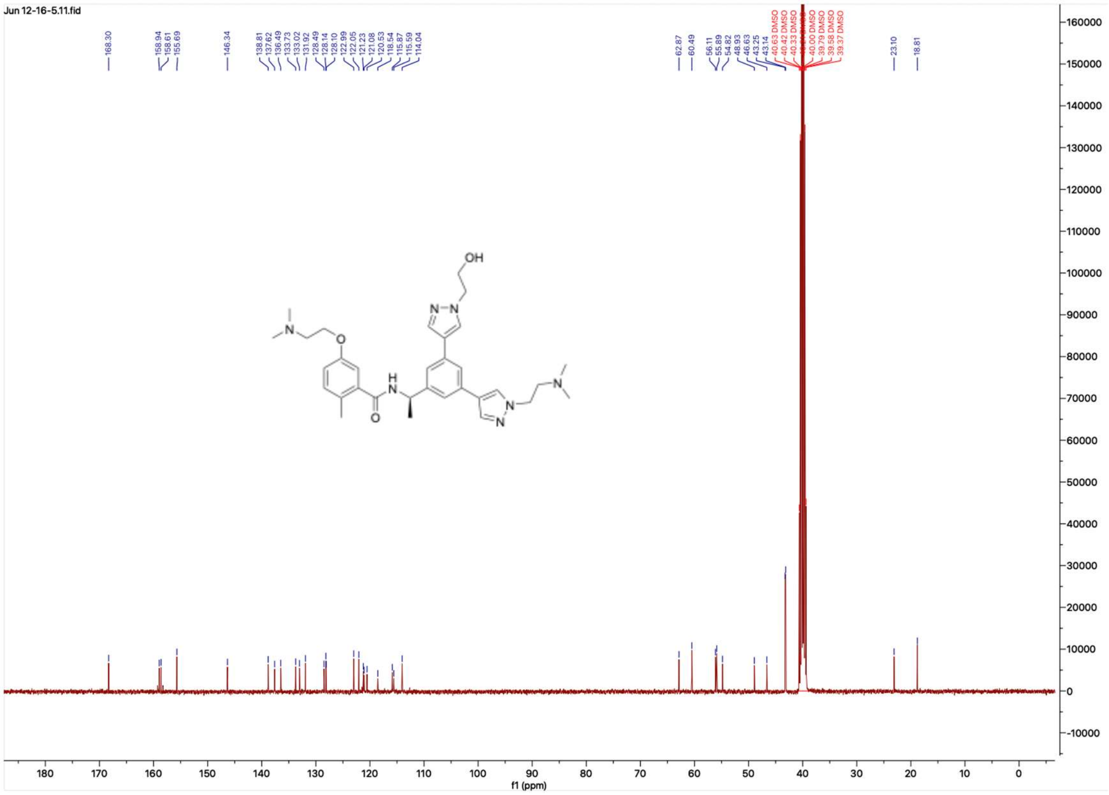

**HNMR and CNMR spectra of Jun12168**

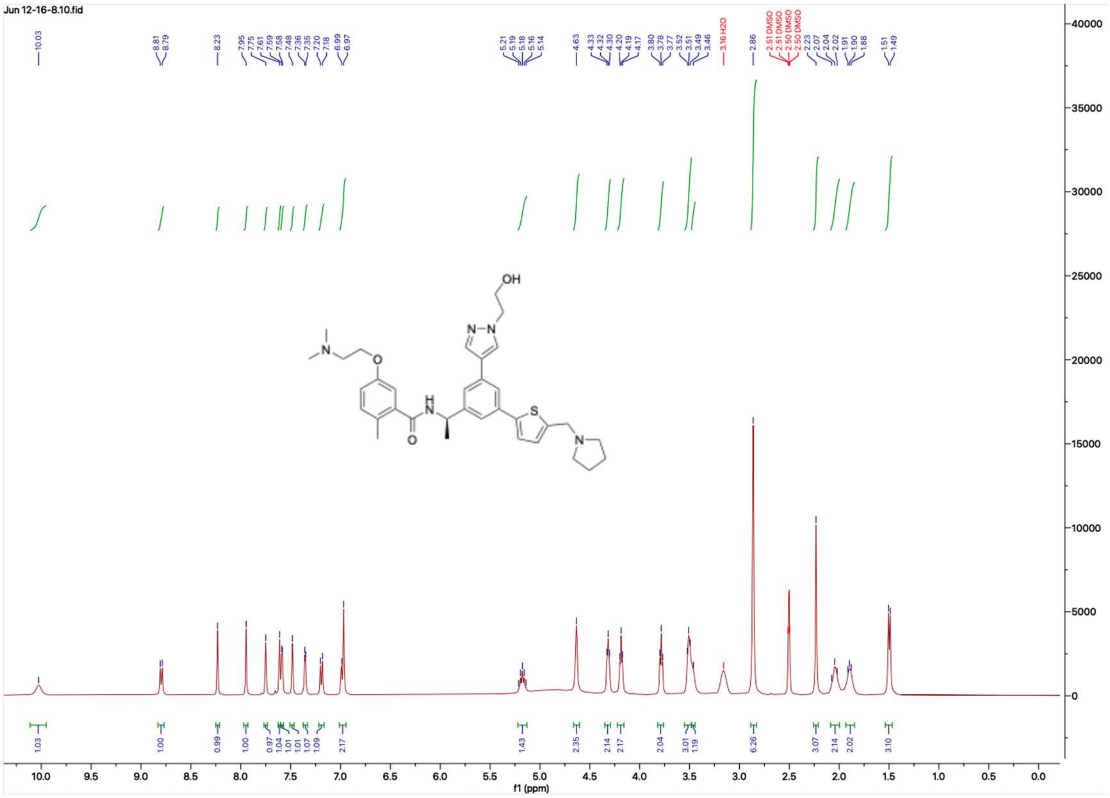

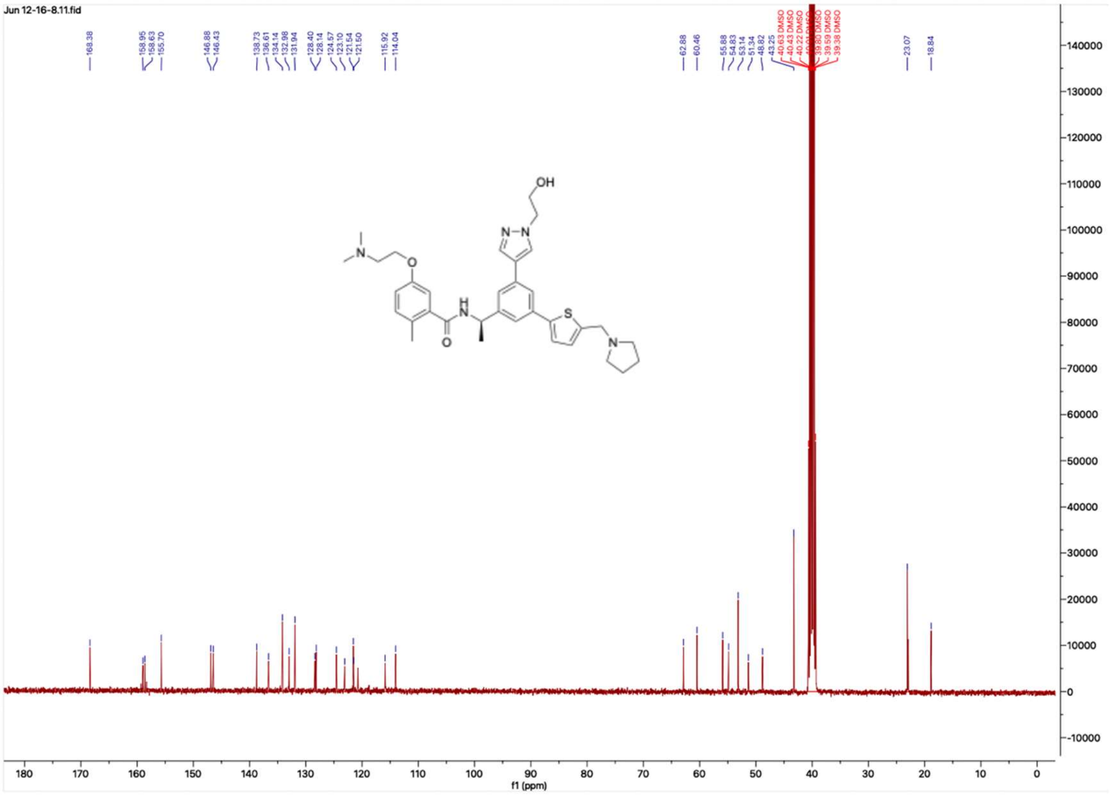

**HNMR and CNMR spectra of Jun12163**

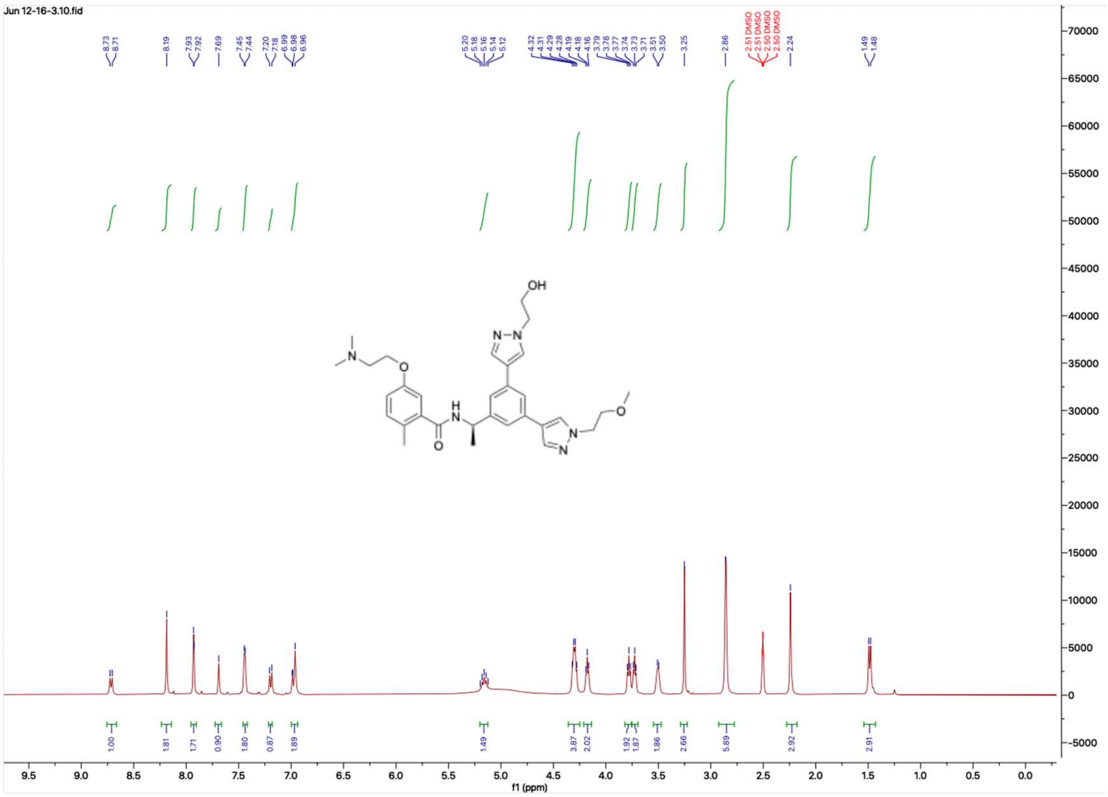

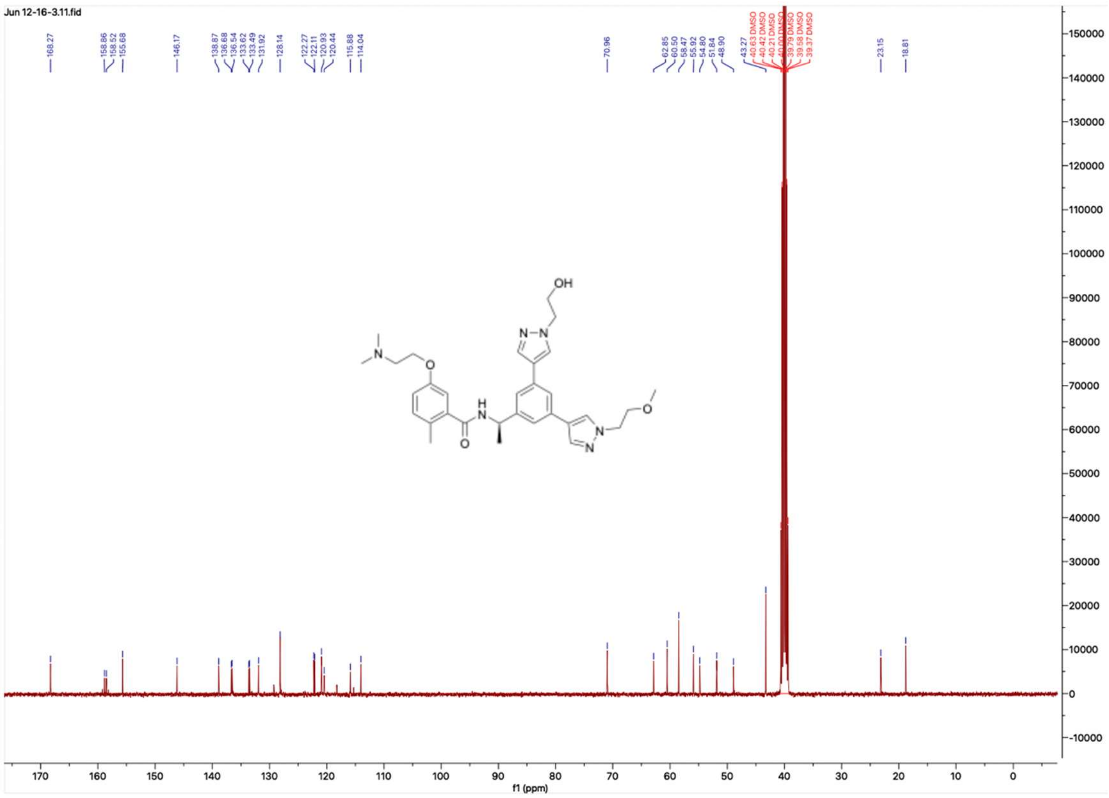

**HNMR and CNMR spectra of Jun12164**

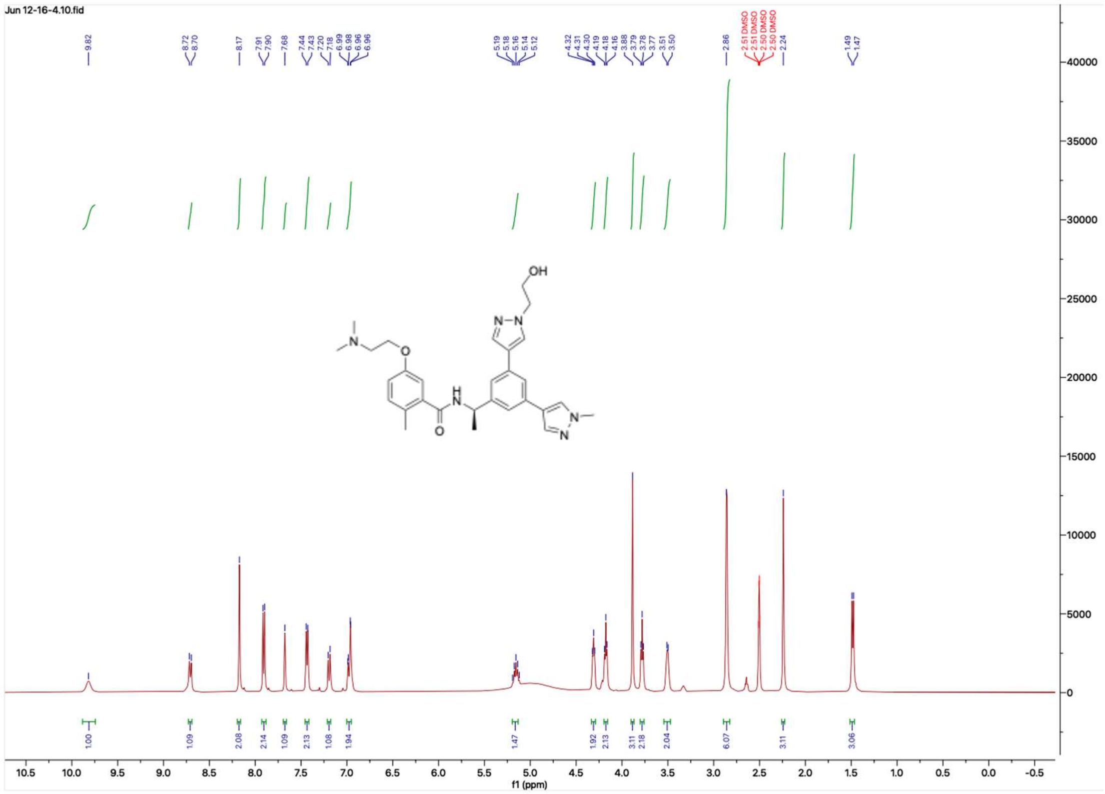


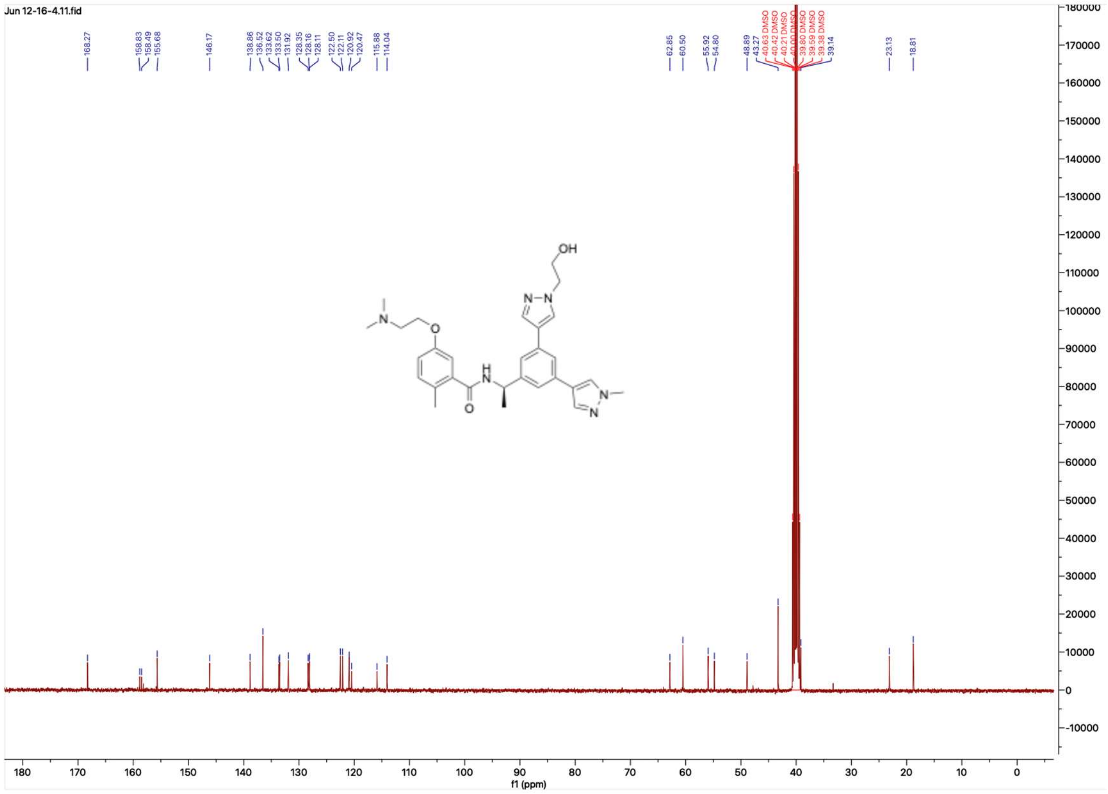

**HNMR and CNMR spectra of Jun12198**

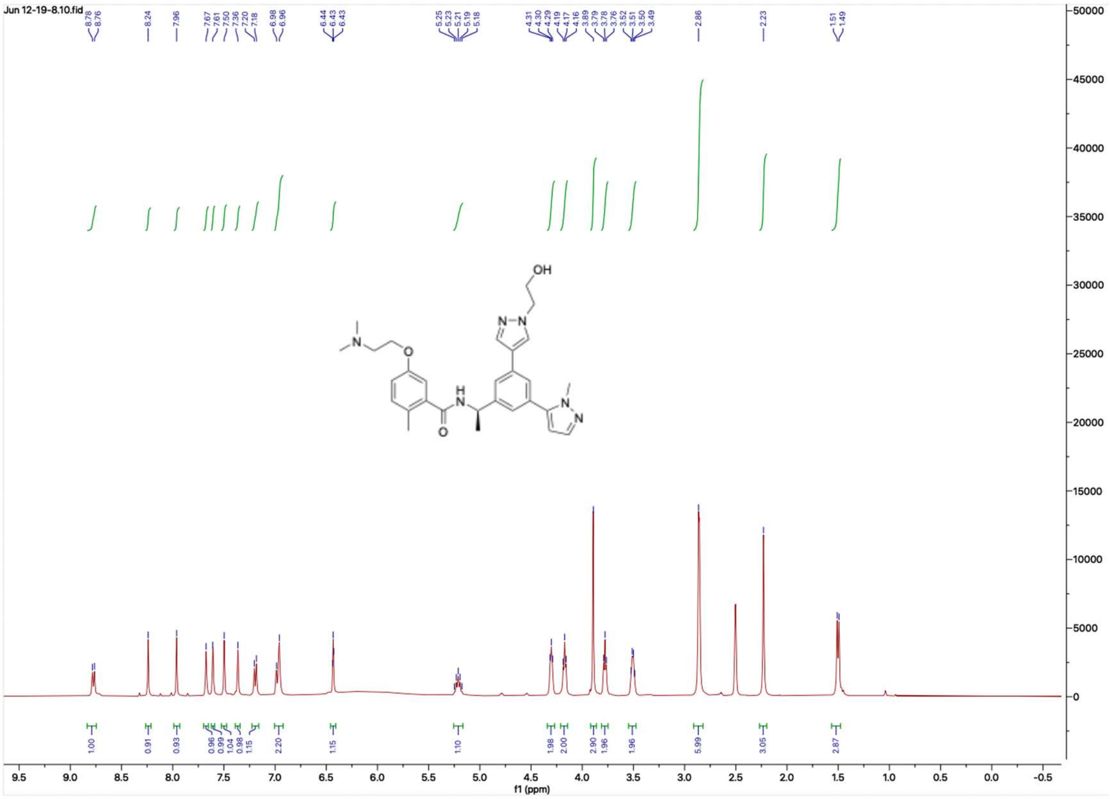


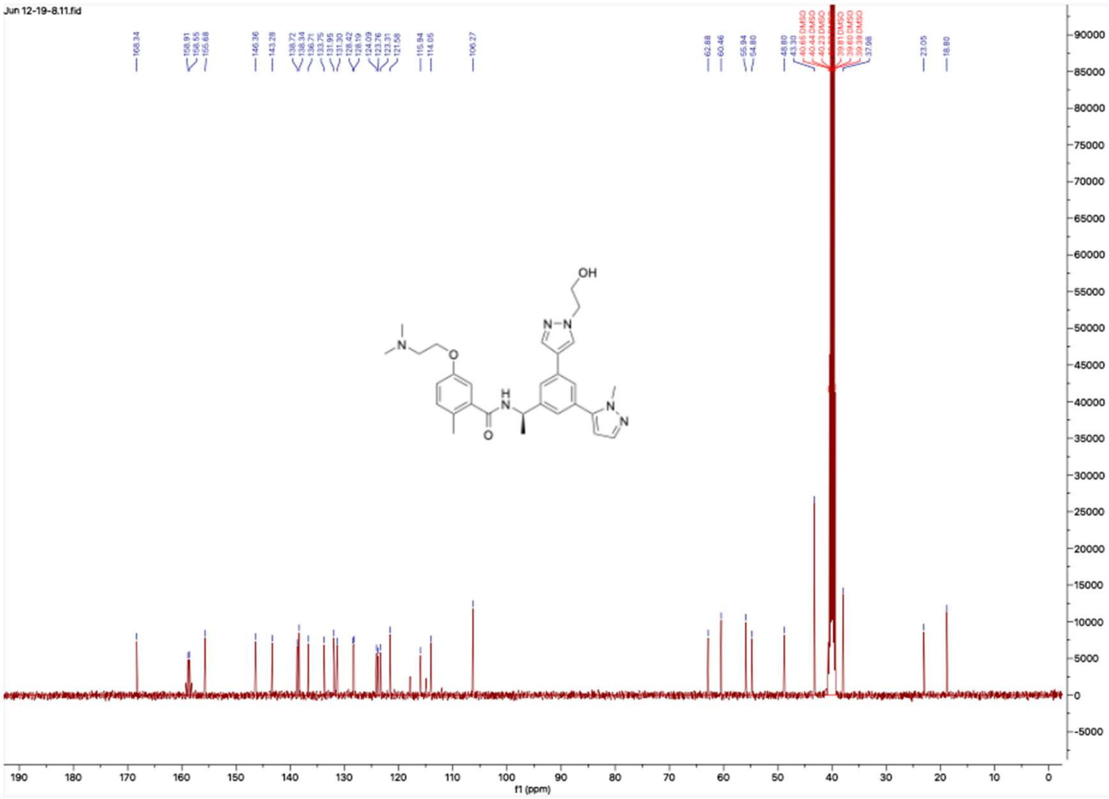

#### HNMR and CNMR spectra of Jun12142

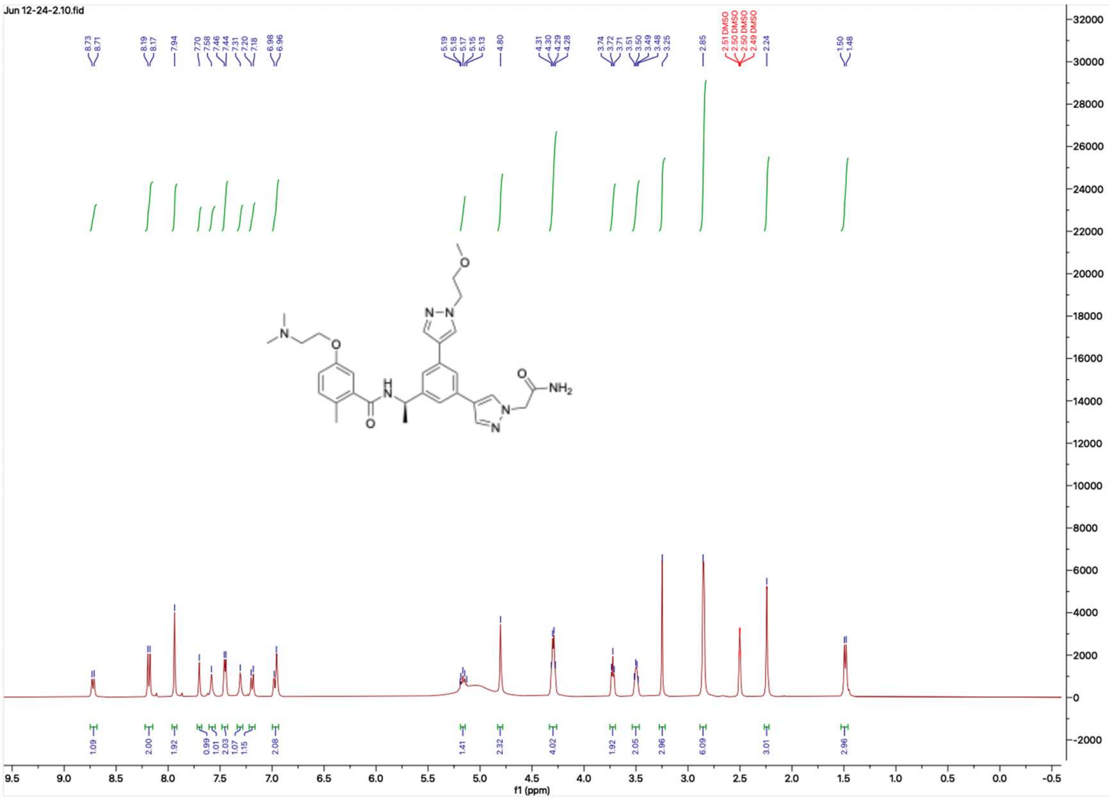


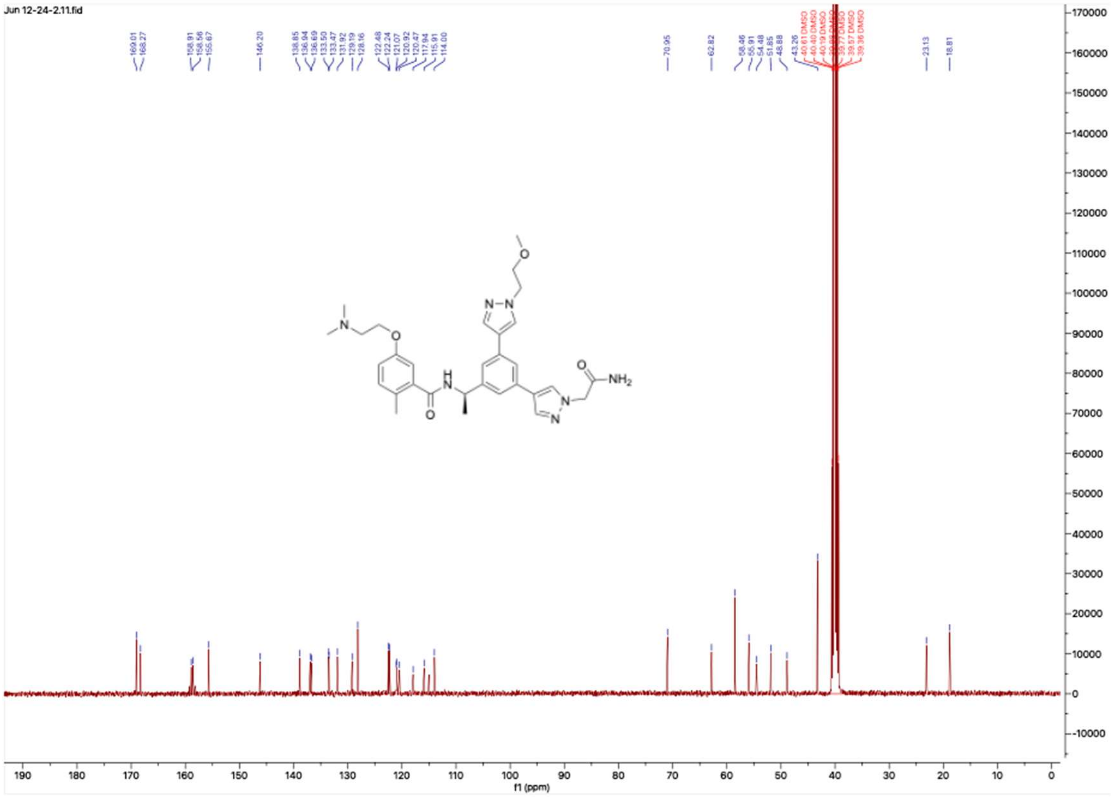

#### HNMR and CNMR spectra of Jun12147

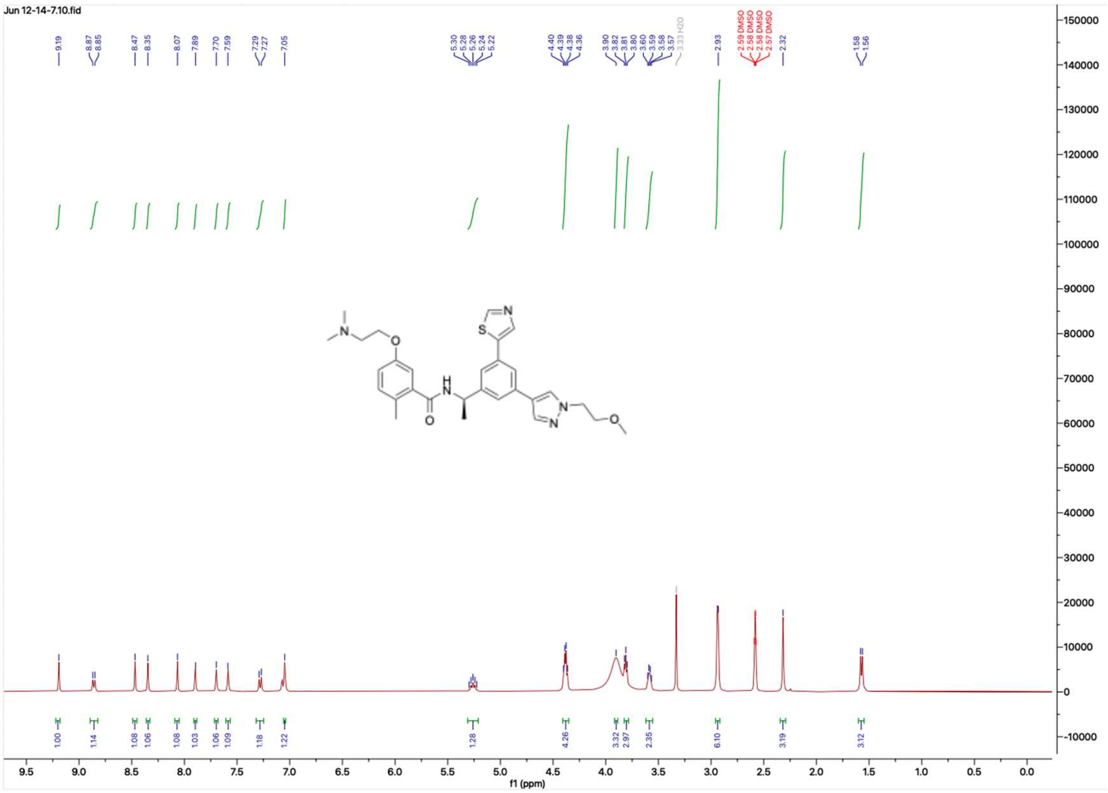

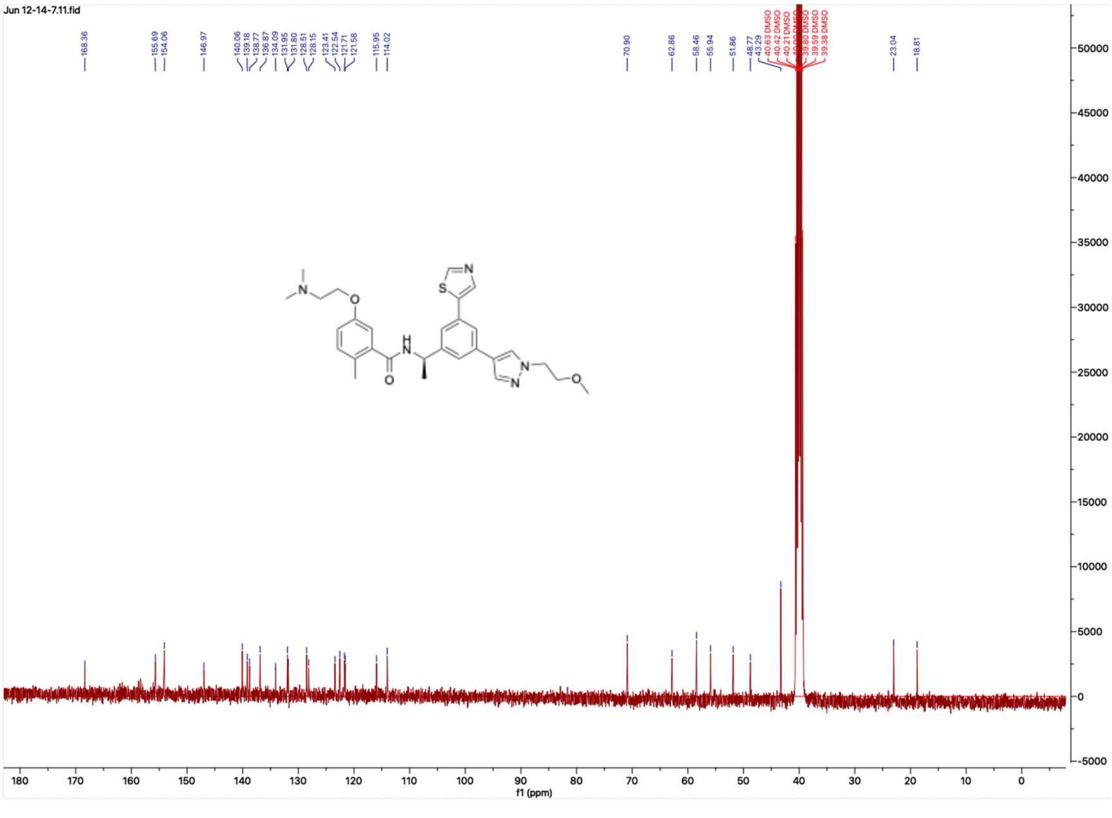

**HNMR and CNMR spectra of Jun12759**

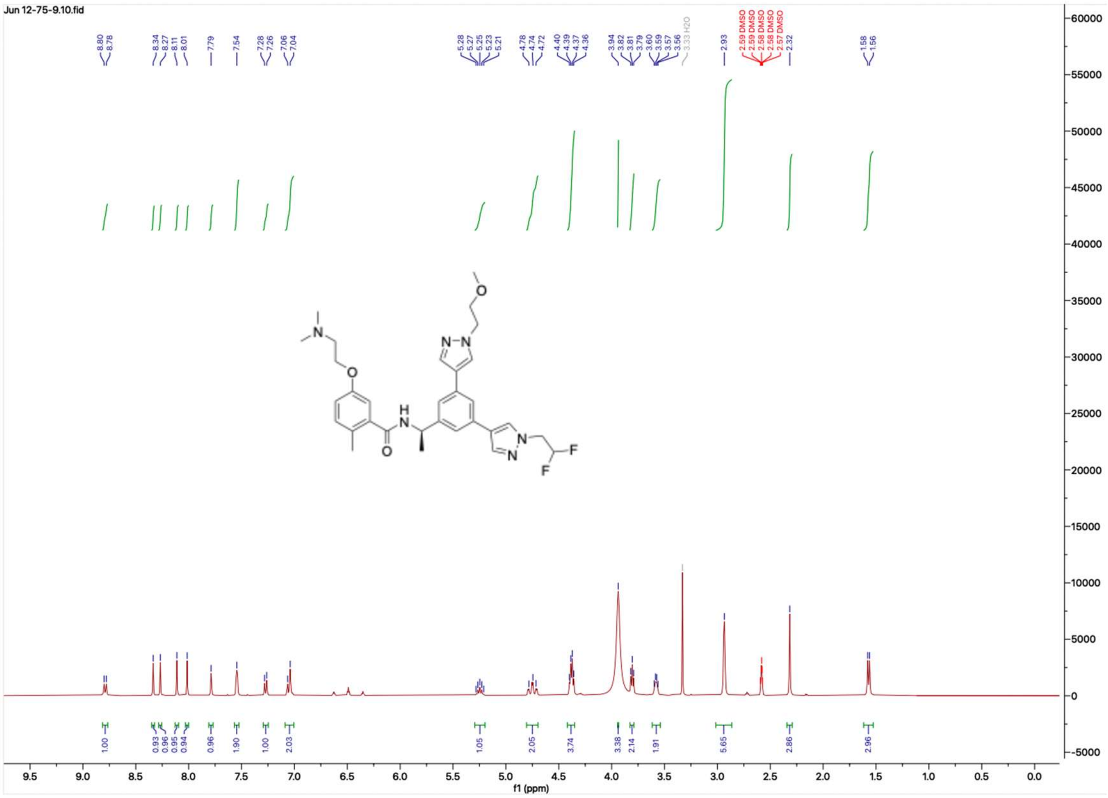

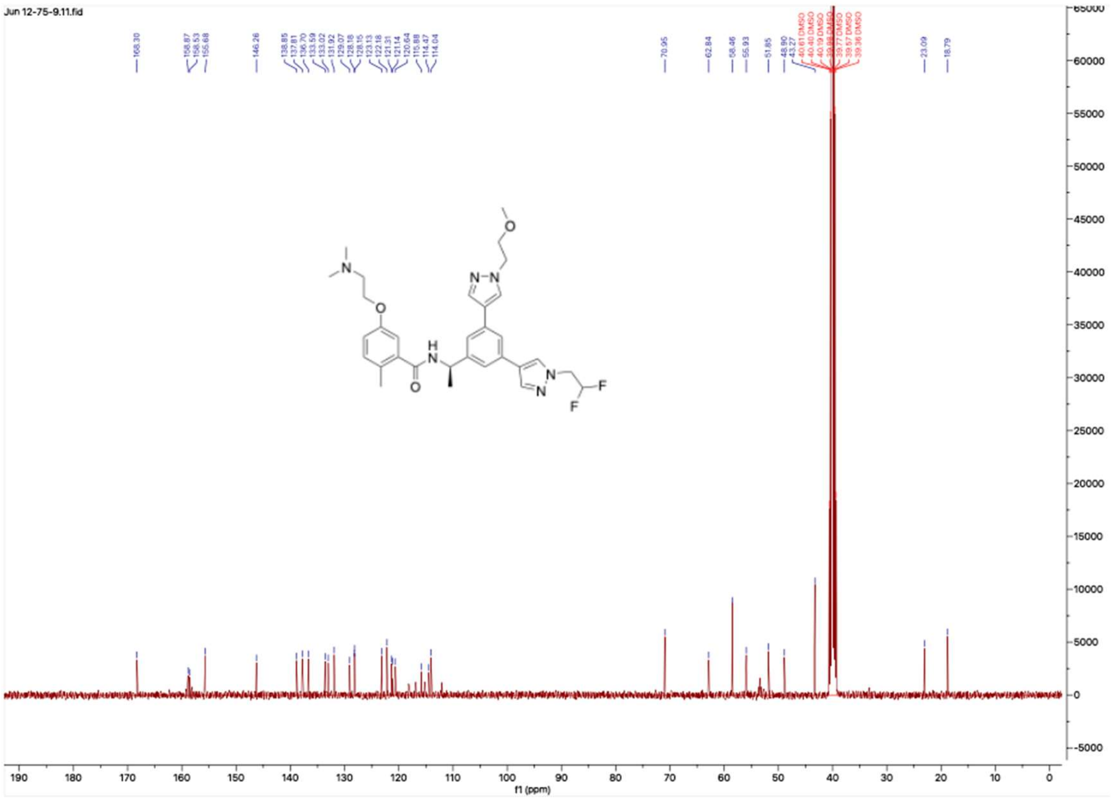

**HNMR and CNMR spectra of Jun12379**

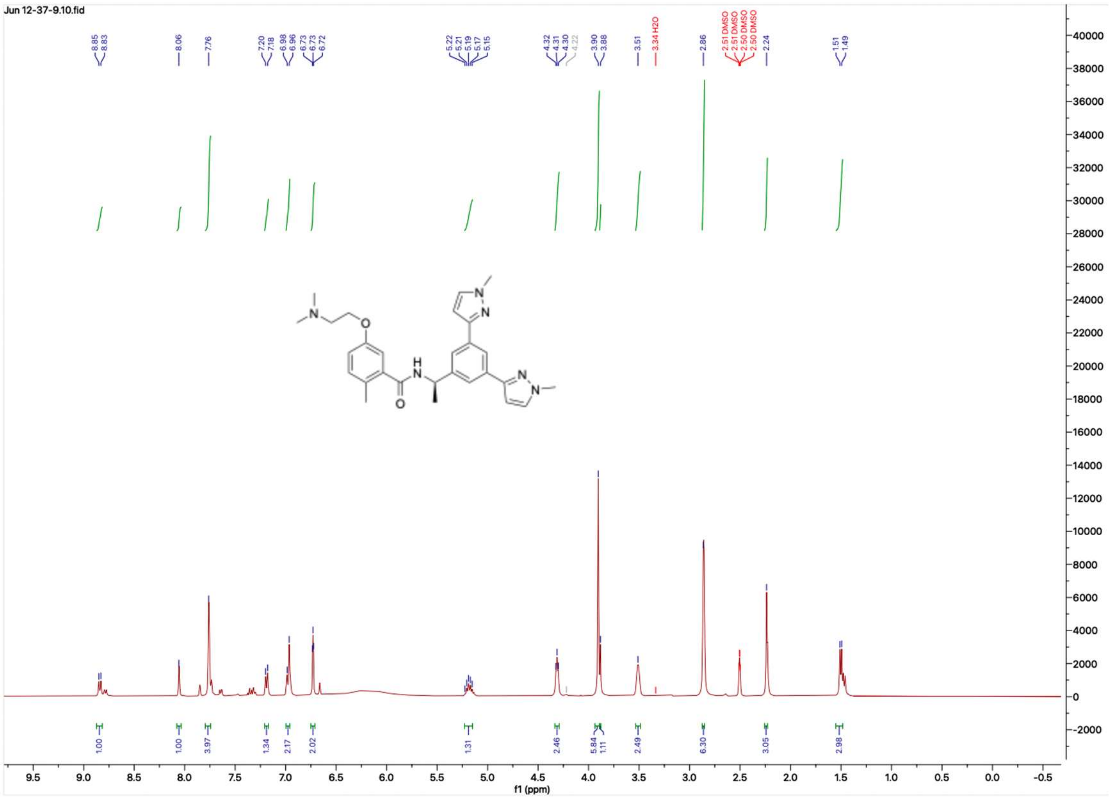


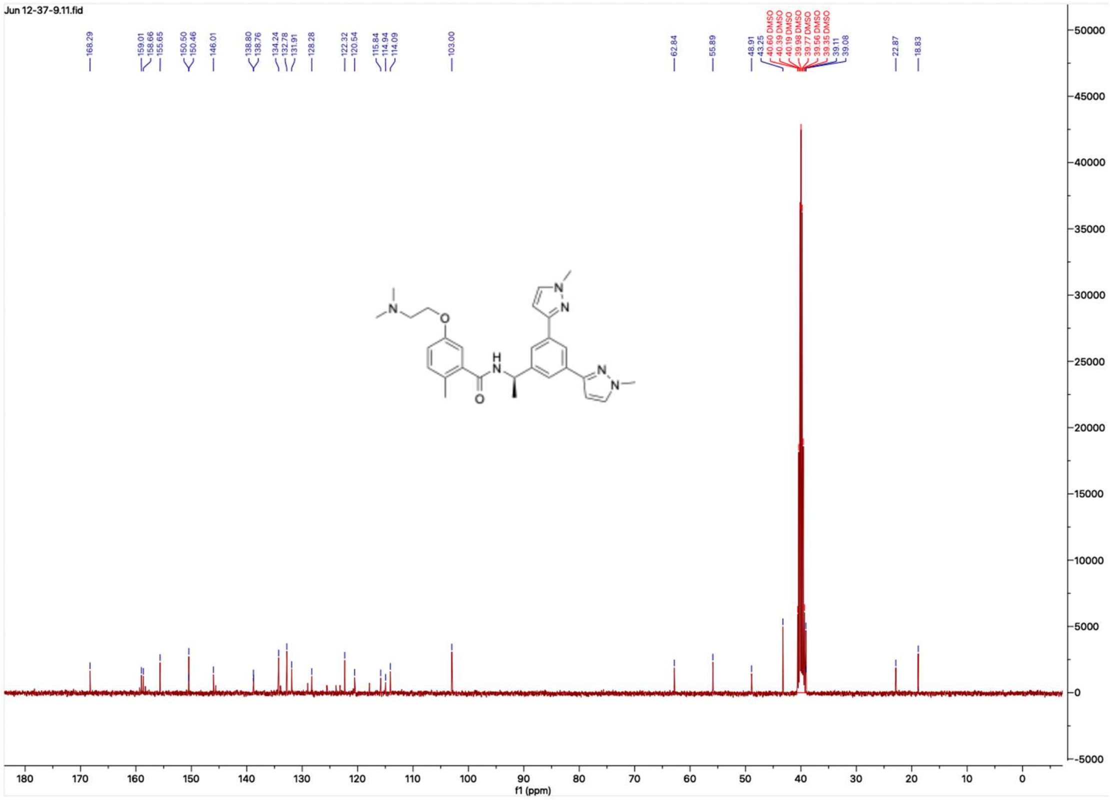

**HNMR and CNMR spectra of Jun12528**

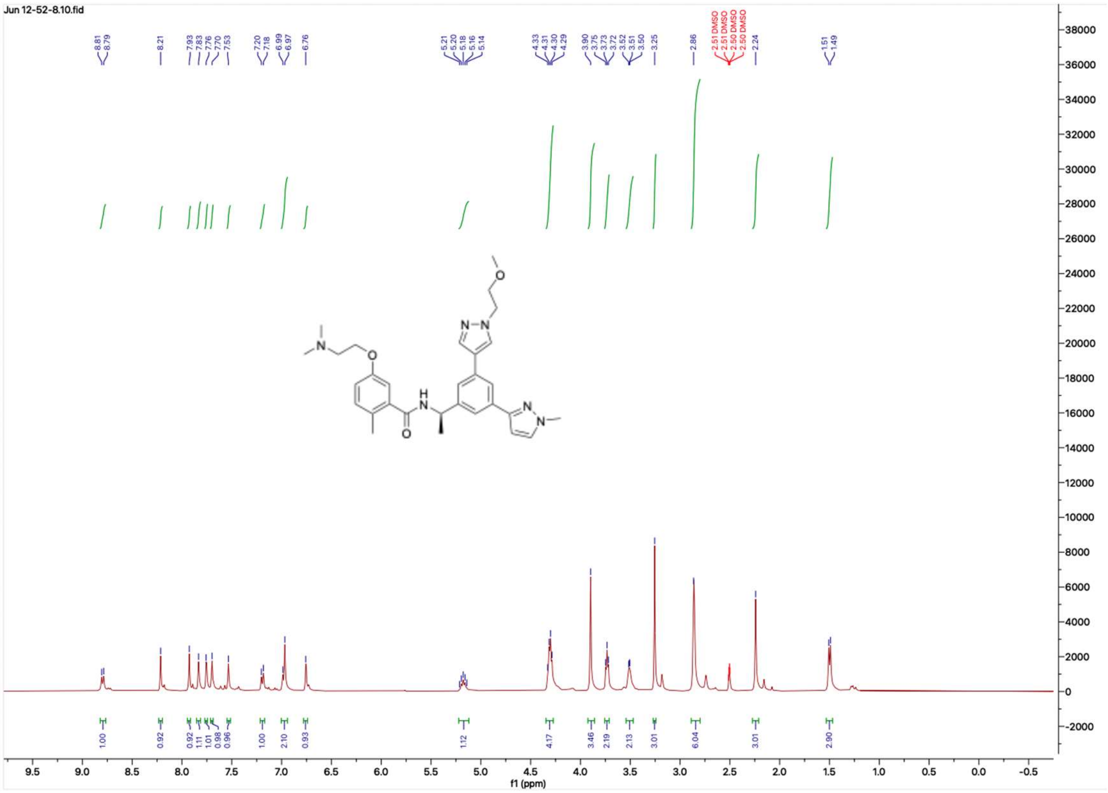


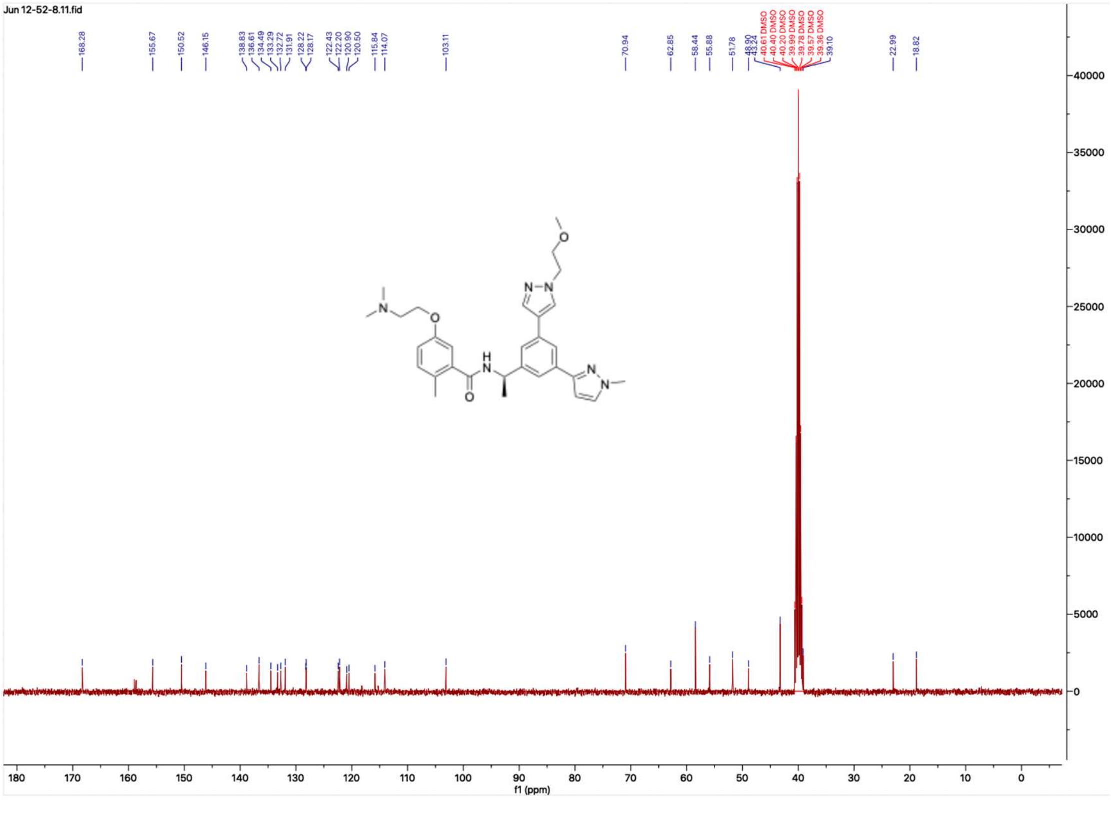

### HNMR and CNMR spectra of Jun12529

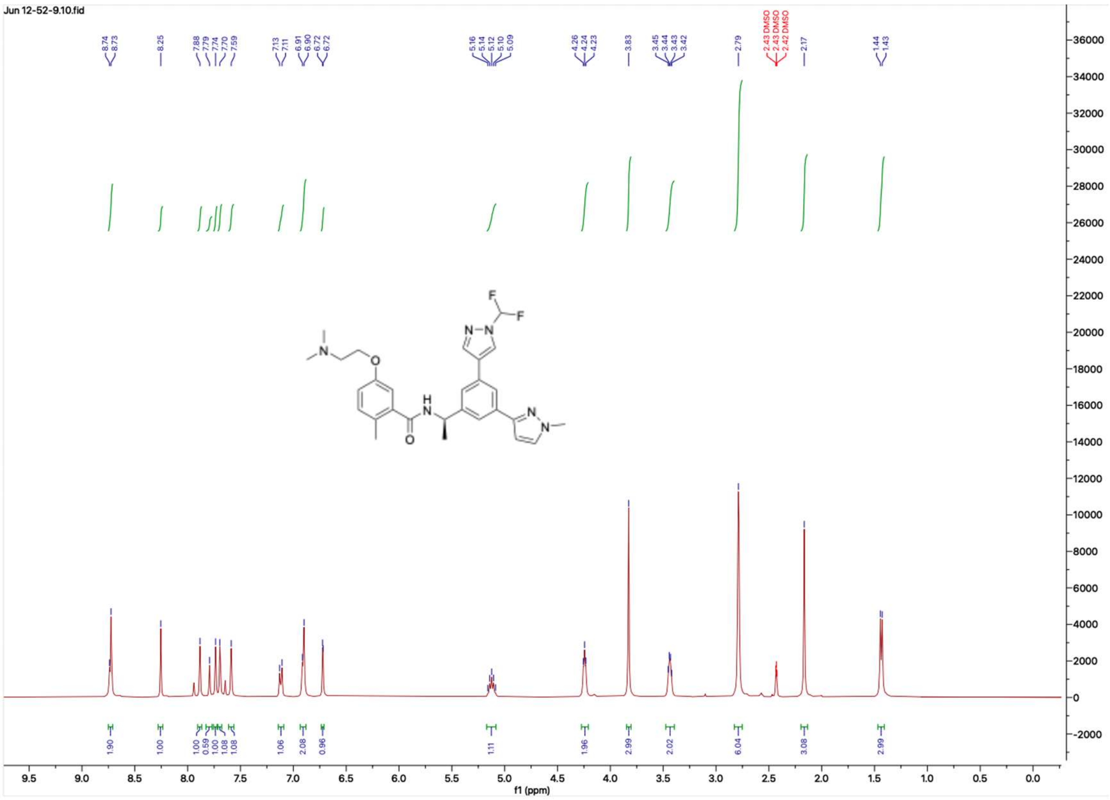

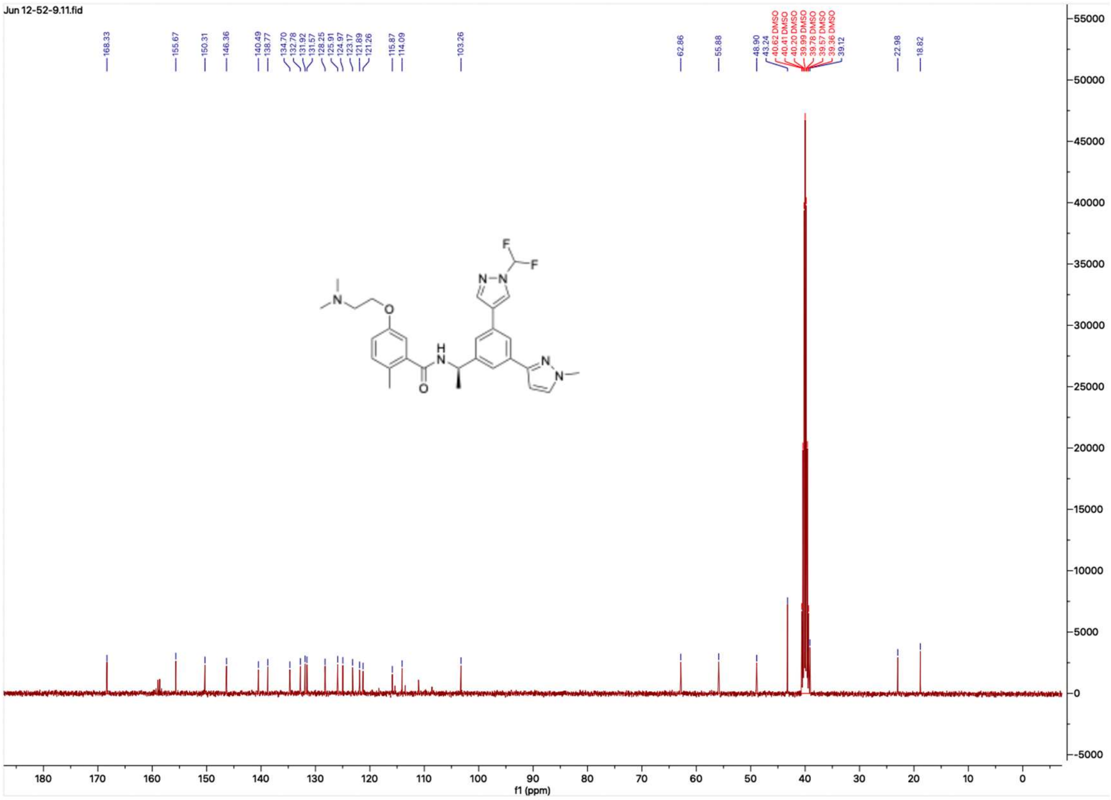

**HNMR and CNMR spectra of Jun12531**

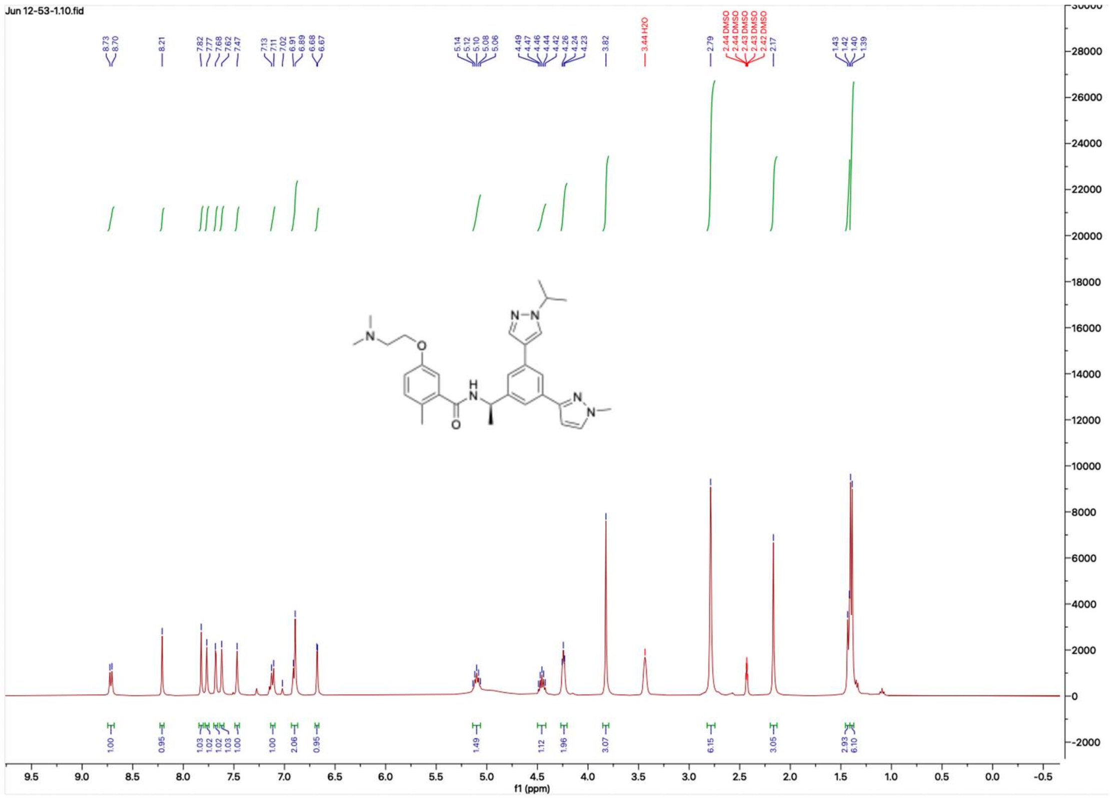

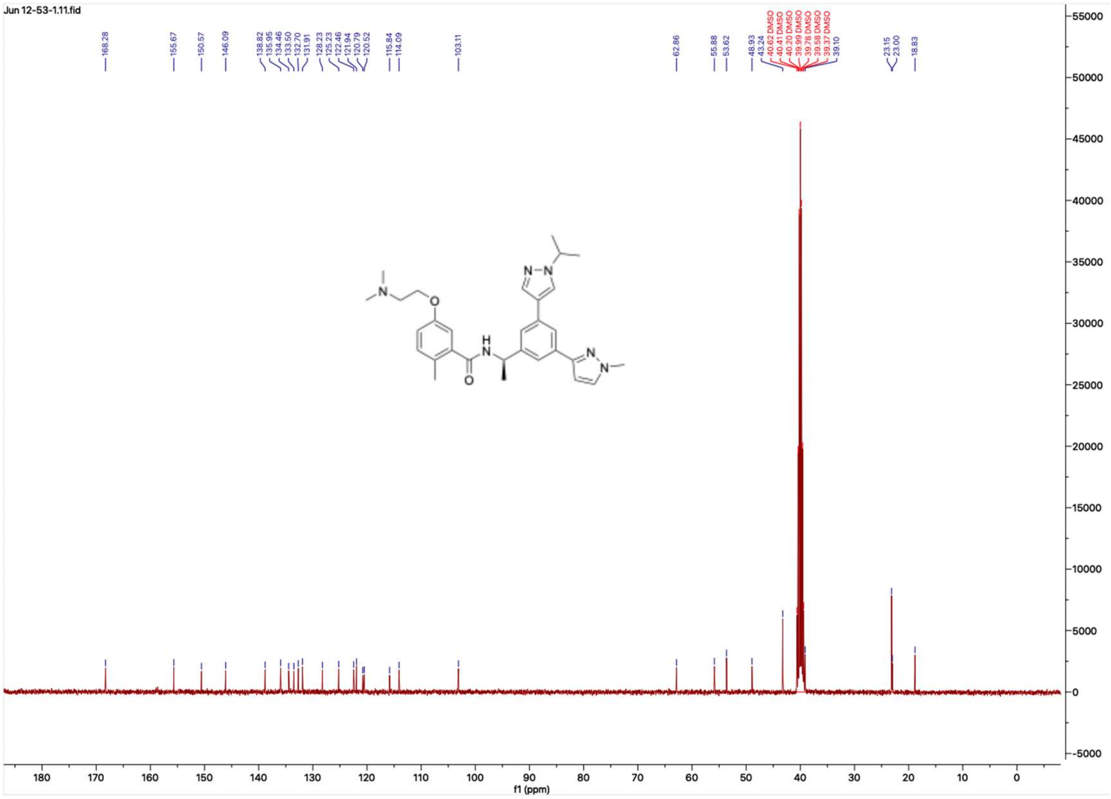

#### HNMR and CNMR spectra of Jun12584

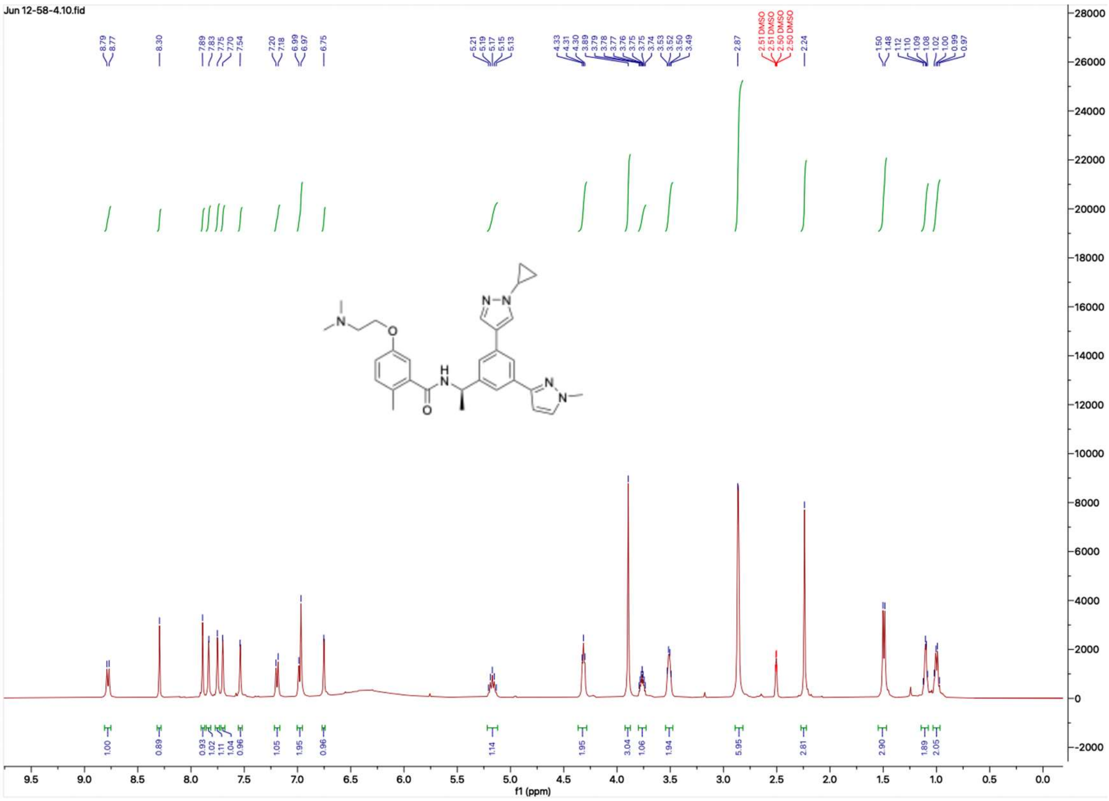


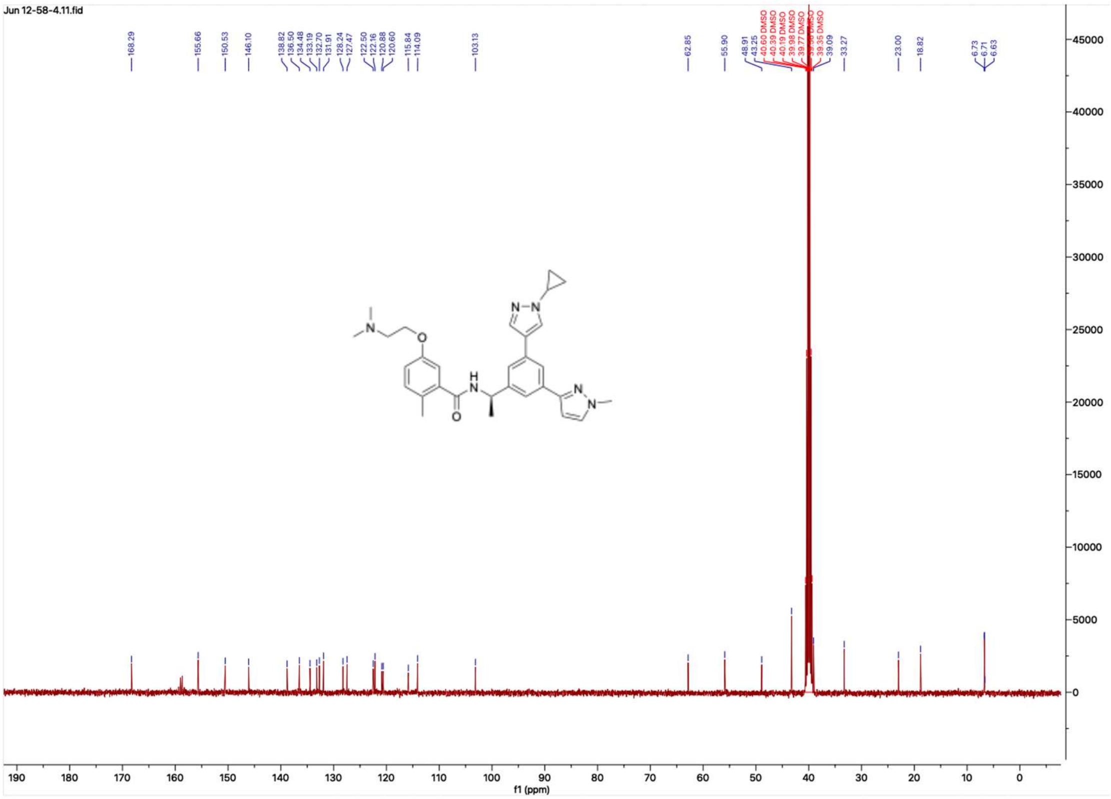

#### HNMR and CNMR spectra of Jun12602

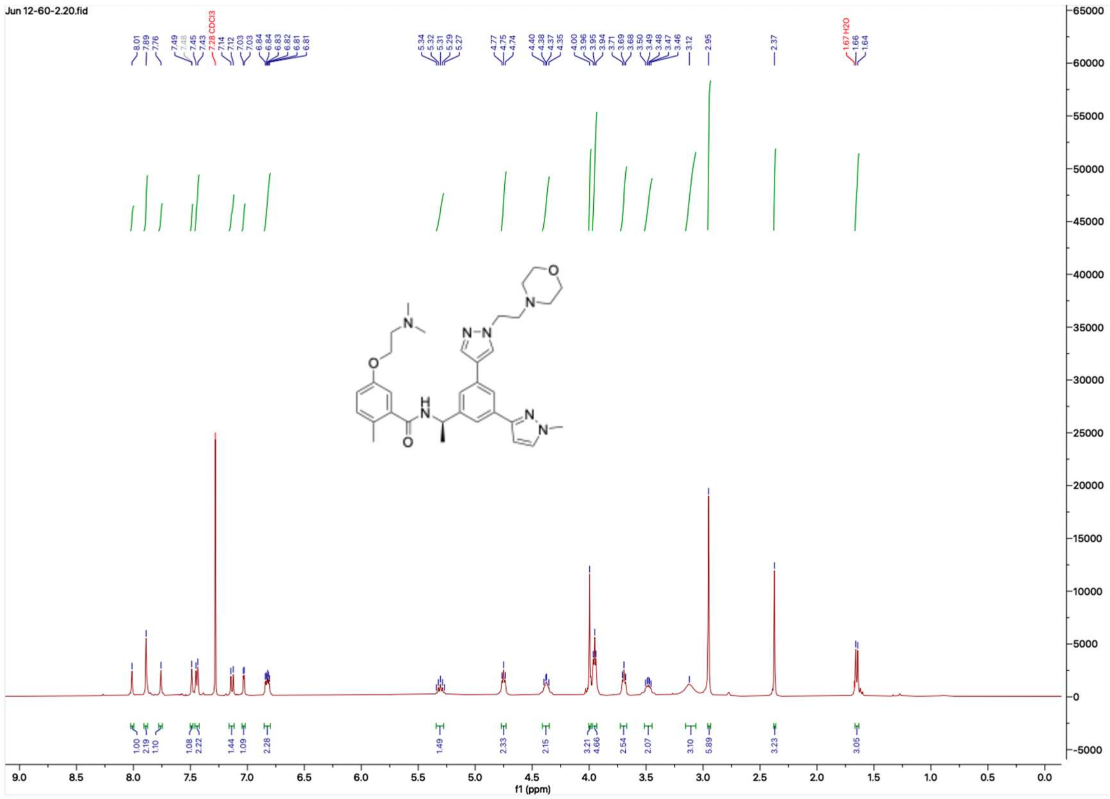

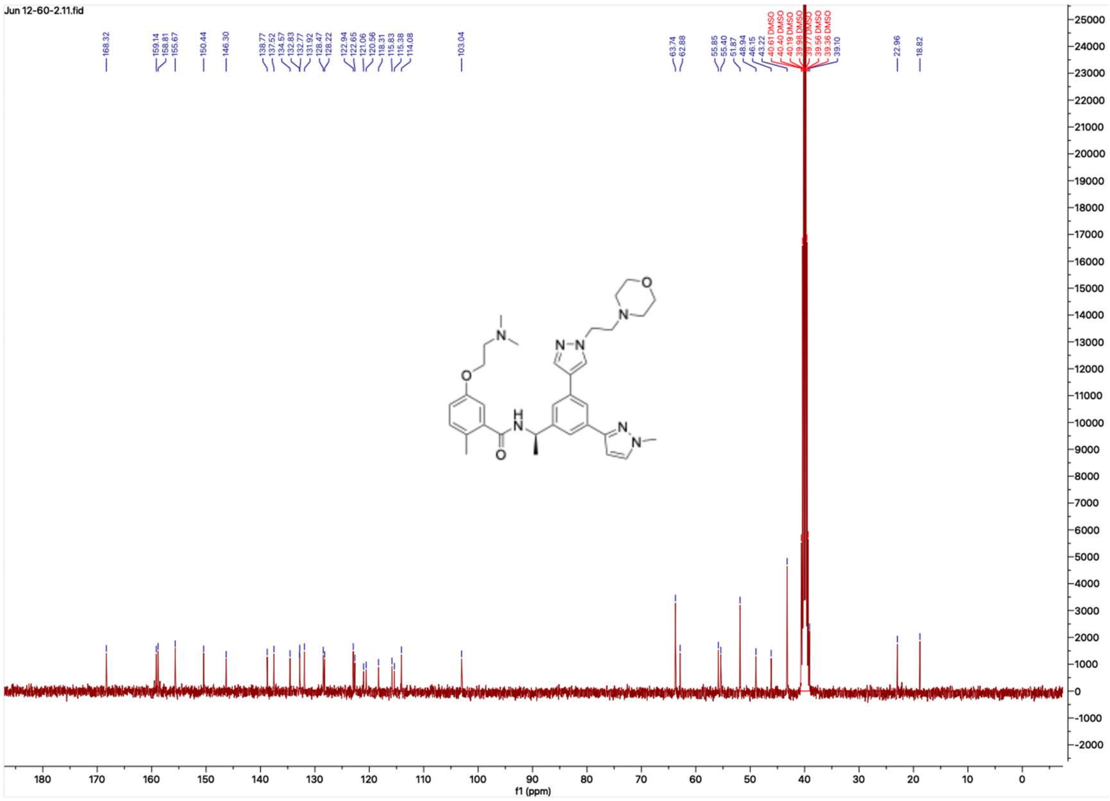

**HNMR and CNMR spectra of Jun12703**

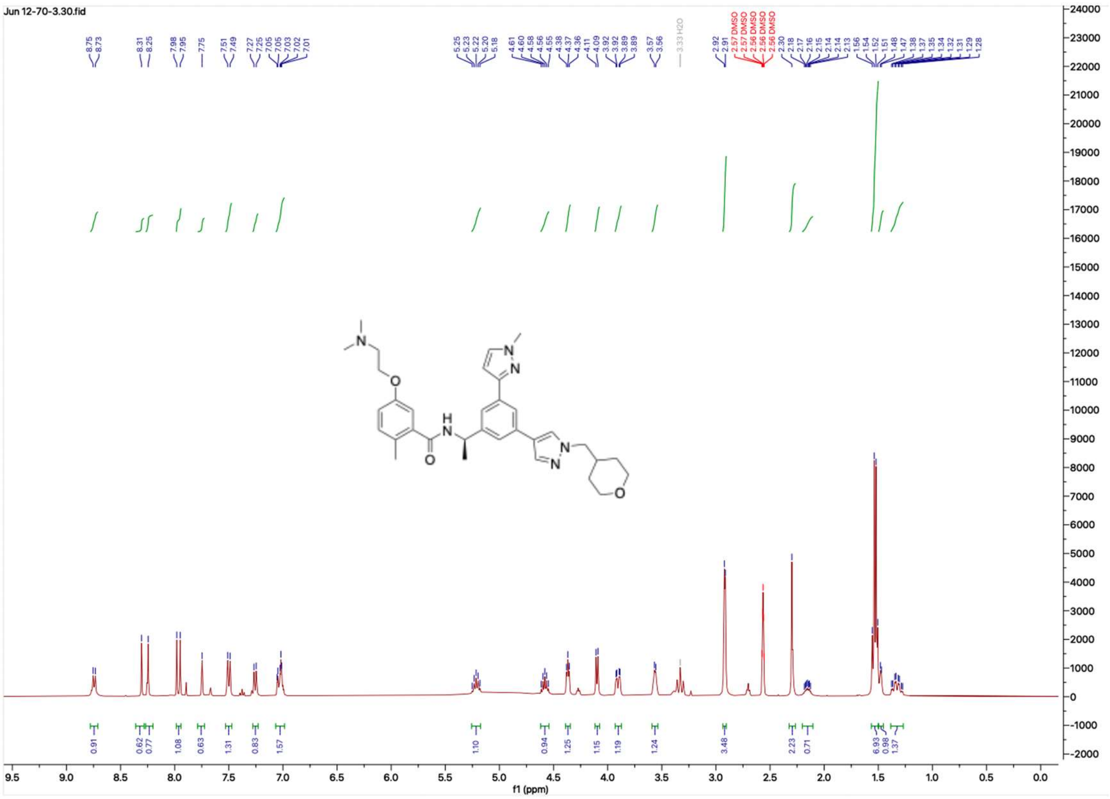

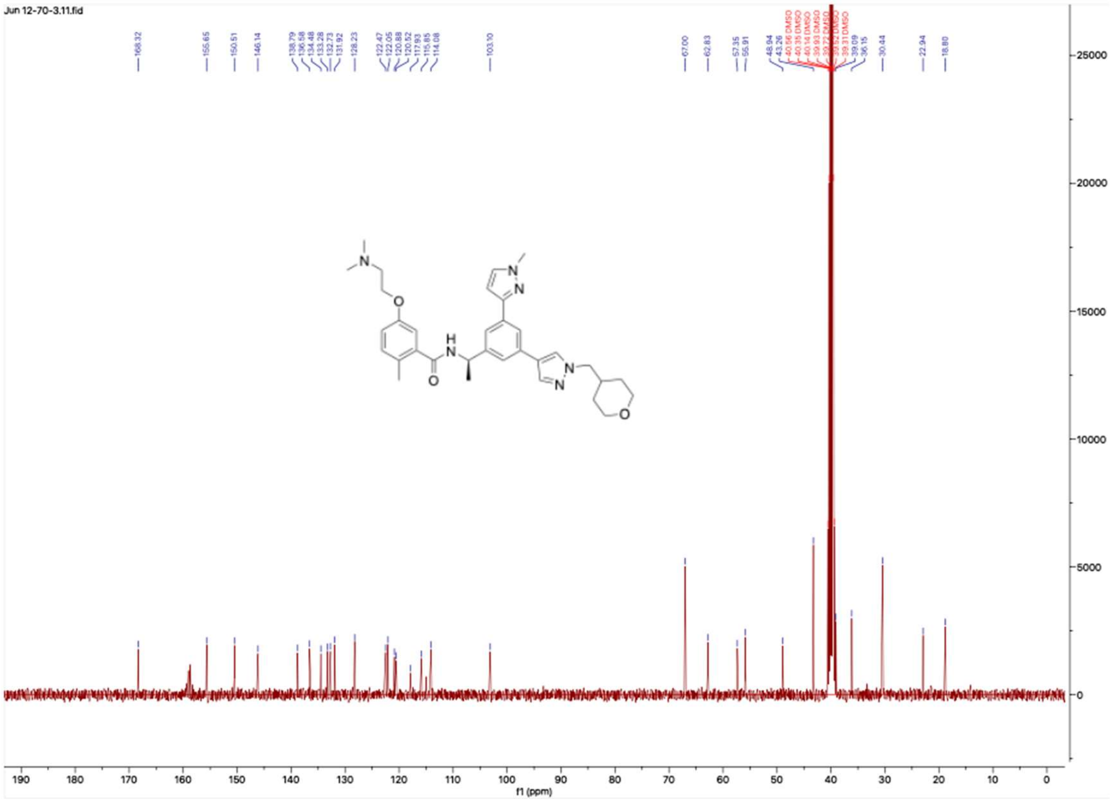

#### HNMR and CNMR spectra of Jun12713

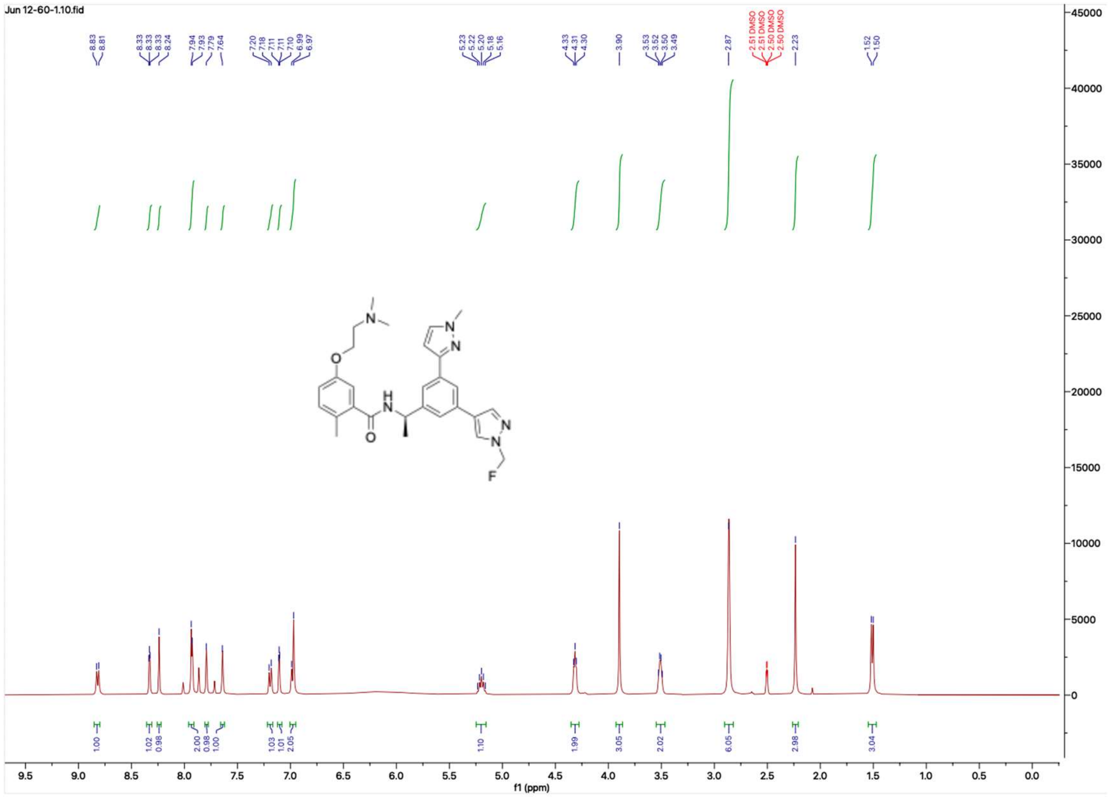

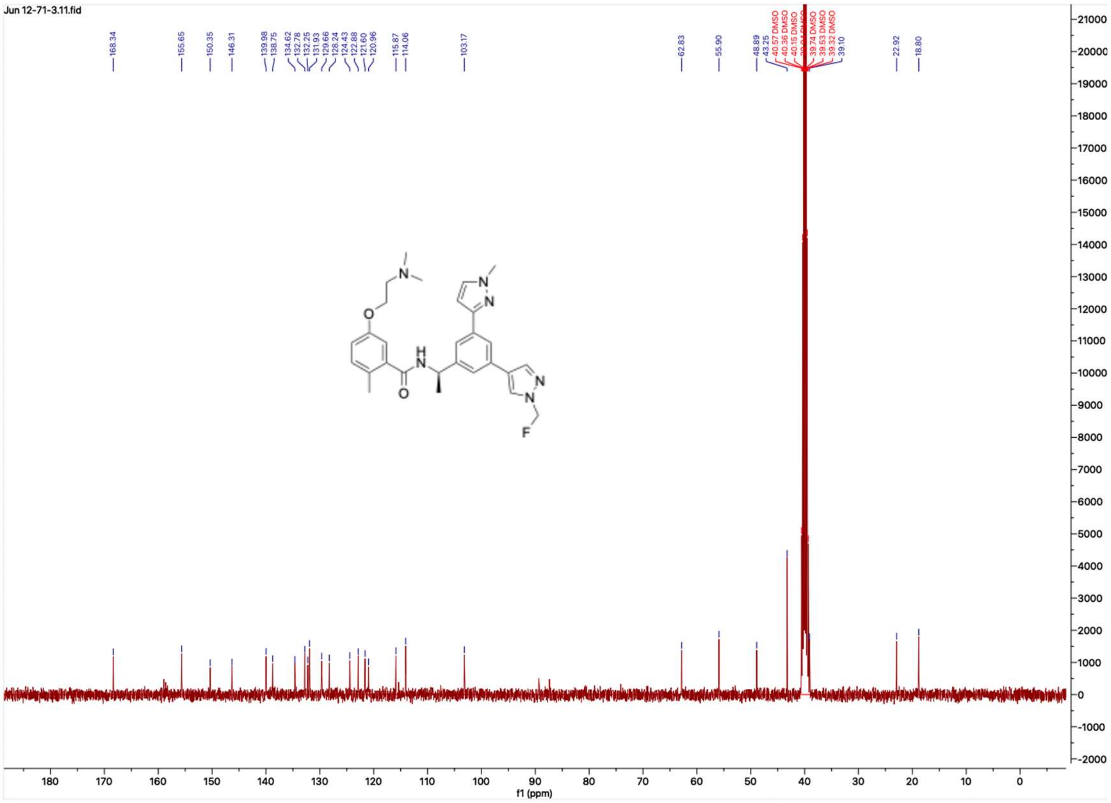

**HNMR and CNMR spectra of Jun12308**

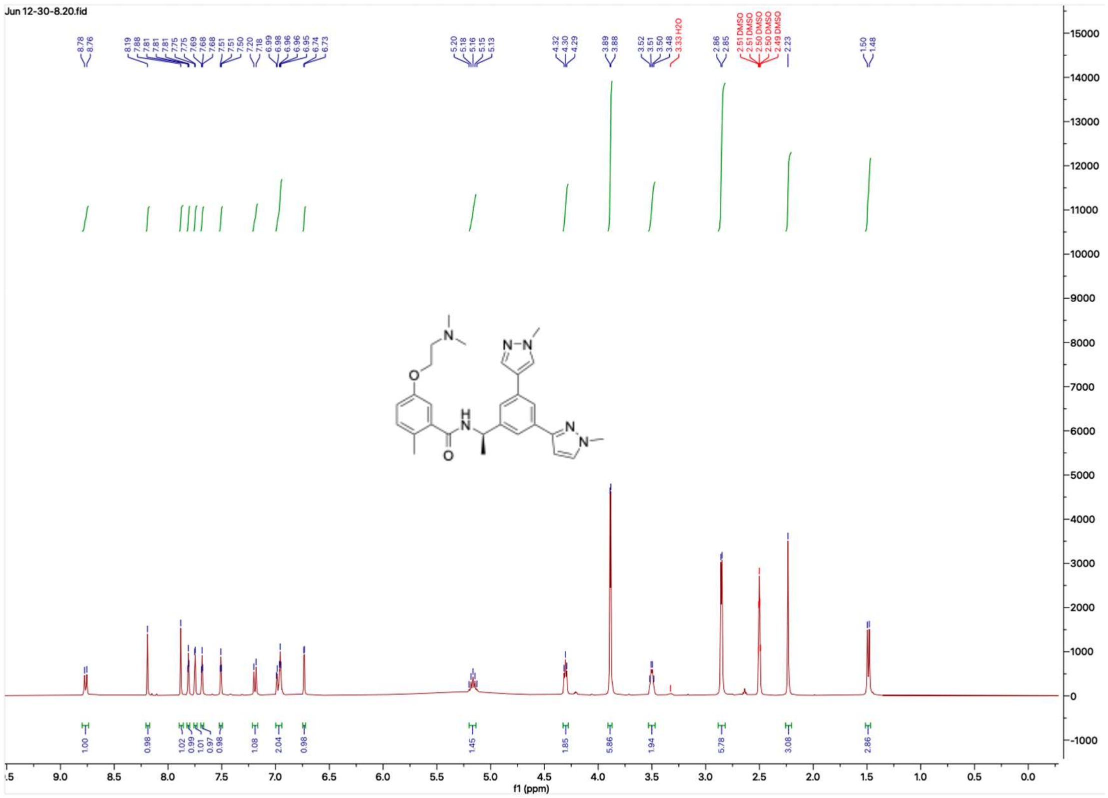

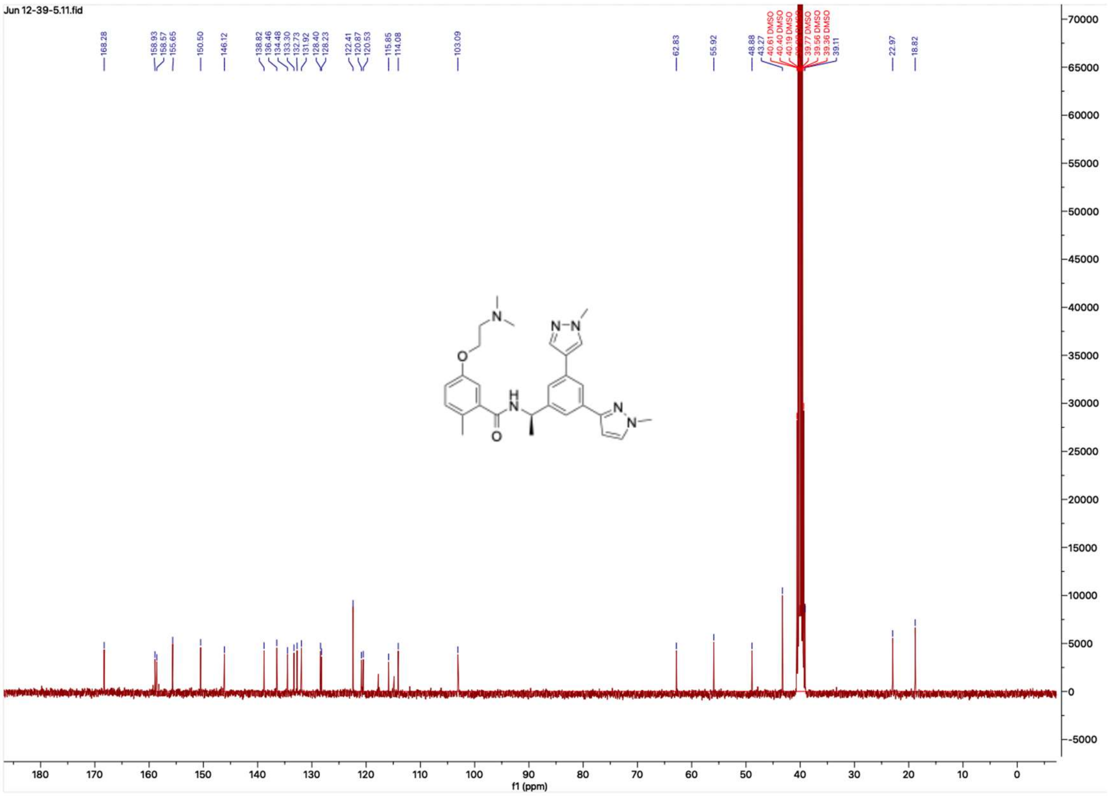

**HNMR and CNMR spectra of Jun12827**

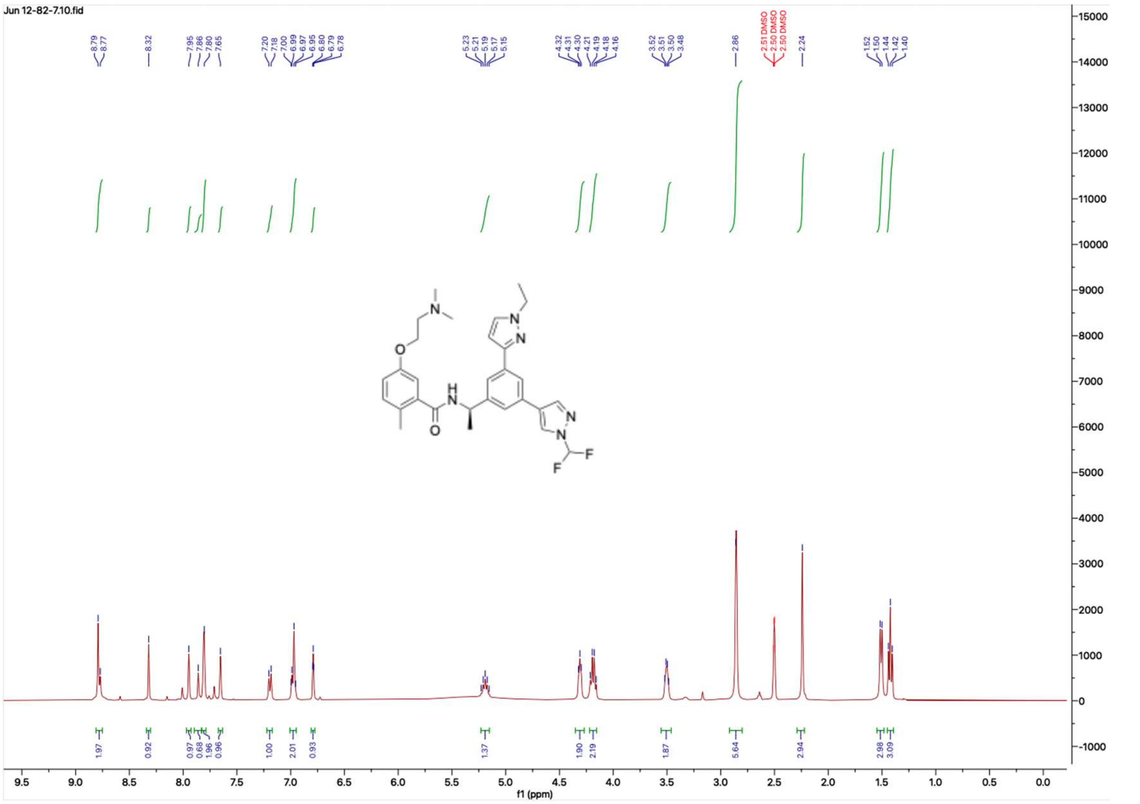

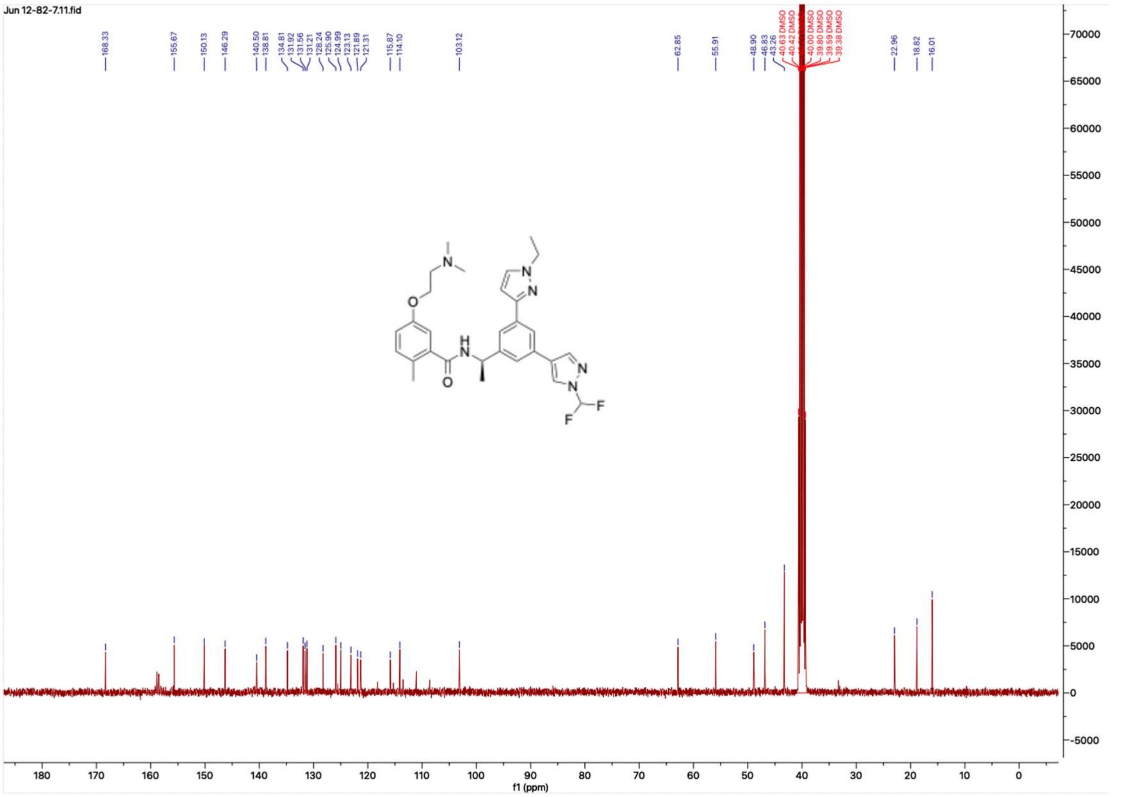

### HNMR and CNMR spectra of Jun12831

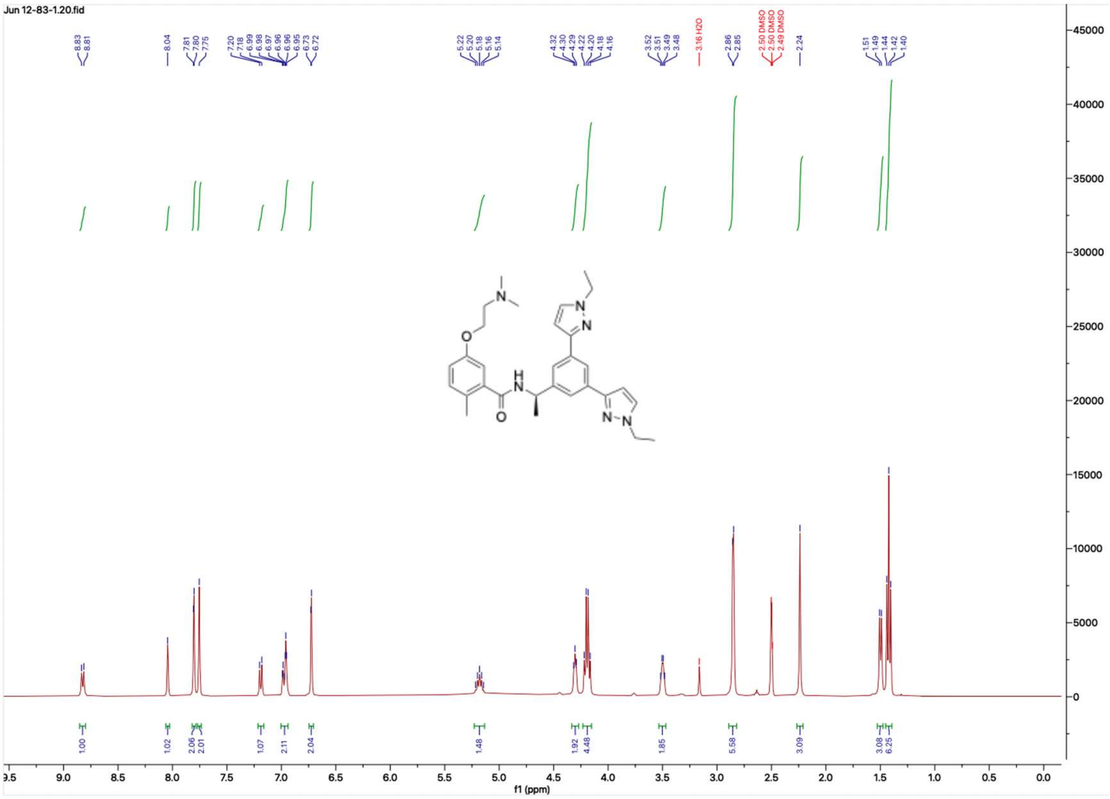


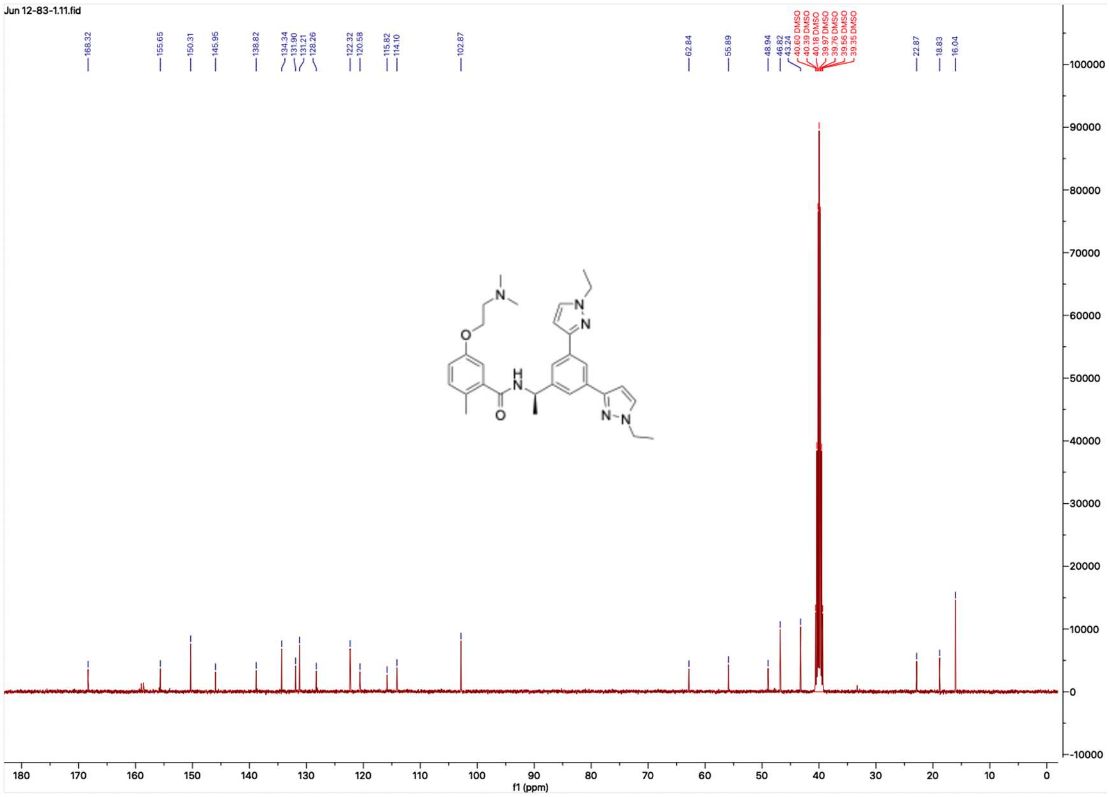

**HNMR and CNMR spectra of Jun12833**

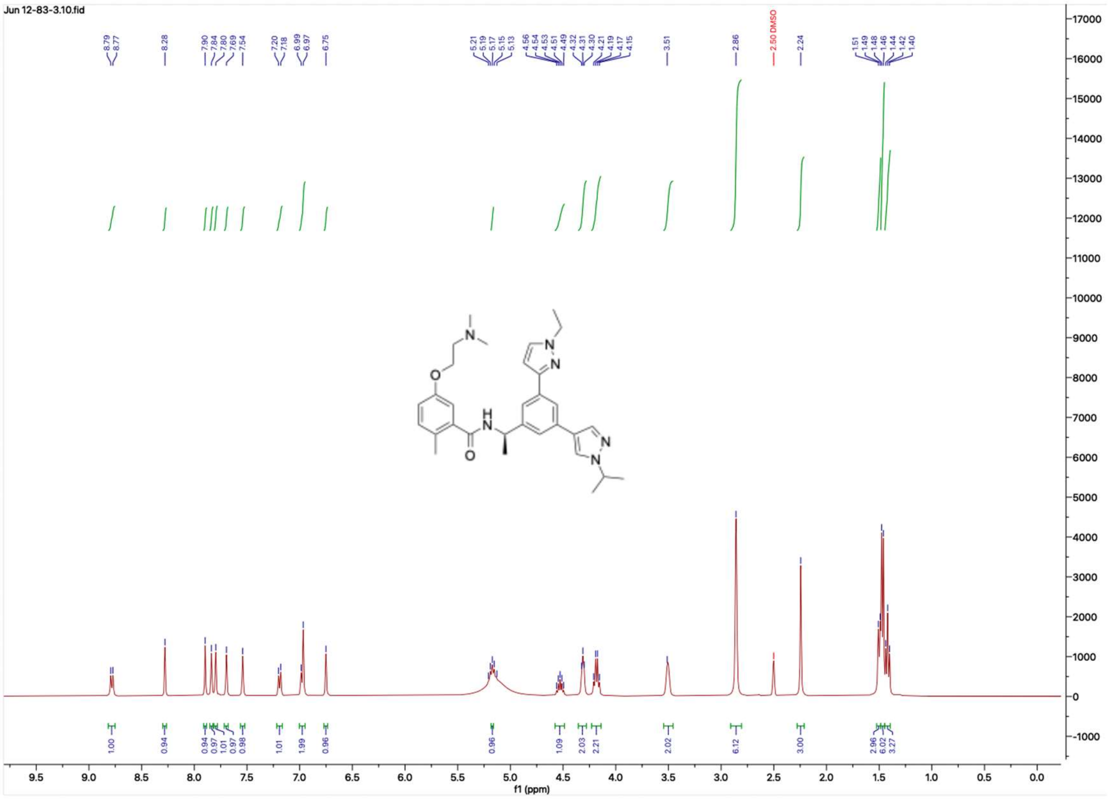

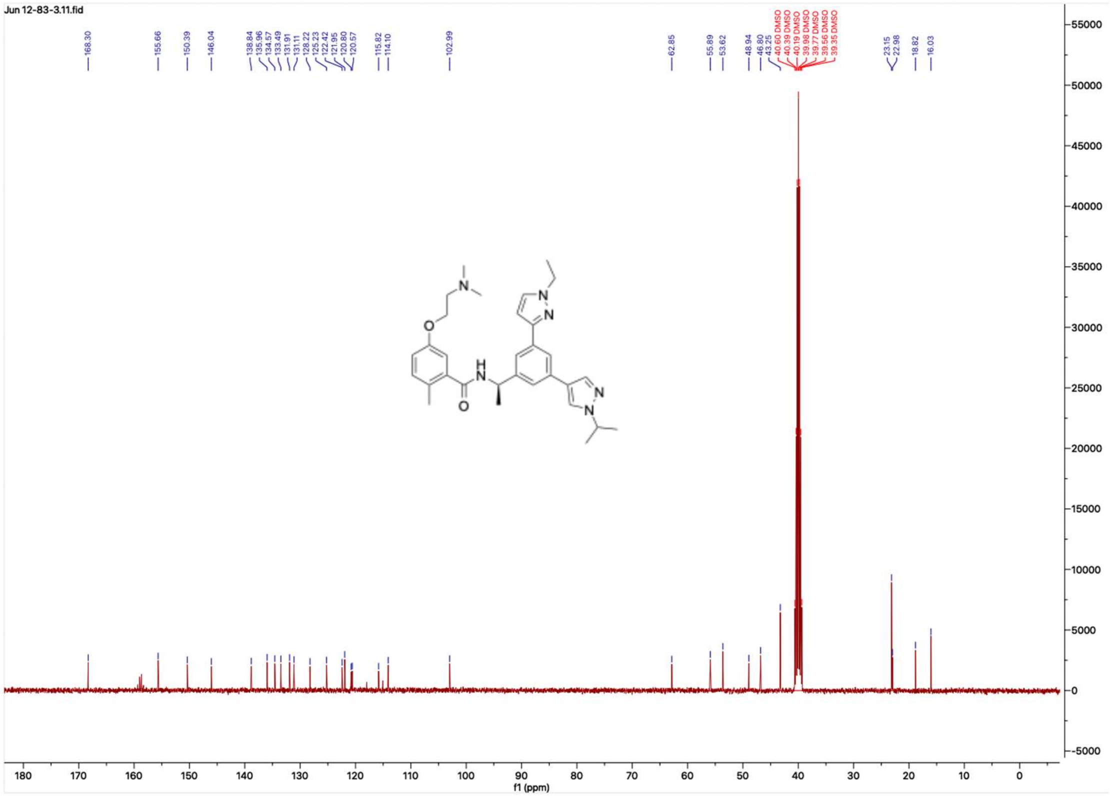

**HNMR and CNMR spectra of Jun12834**

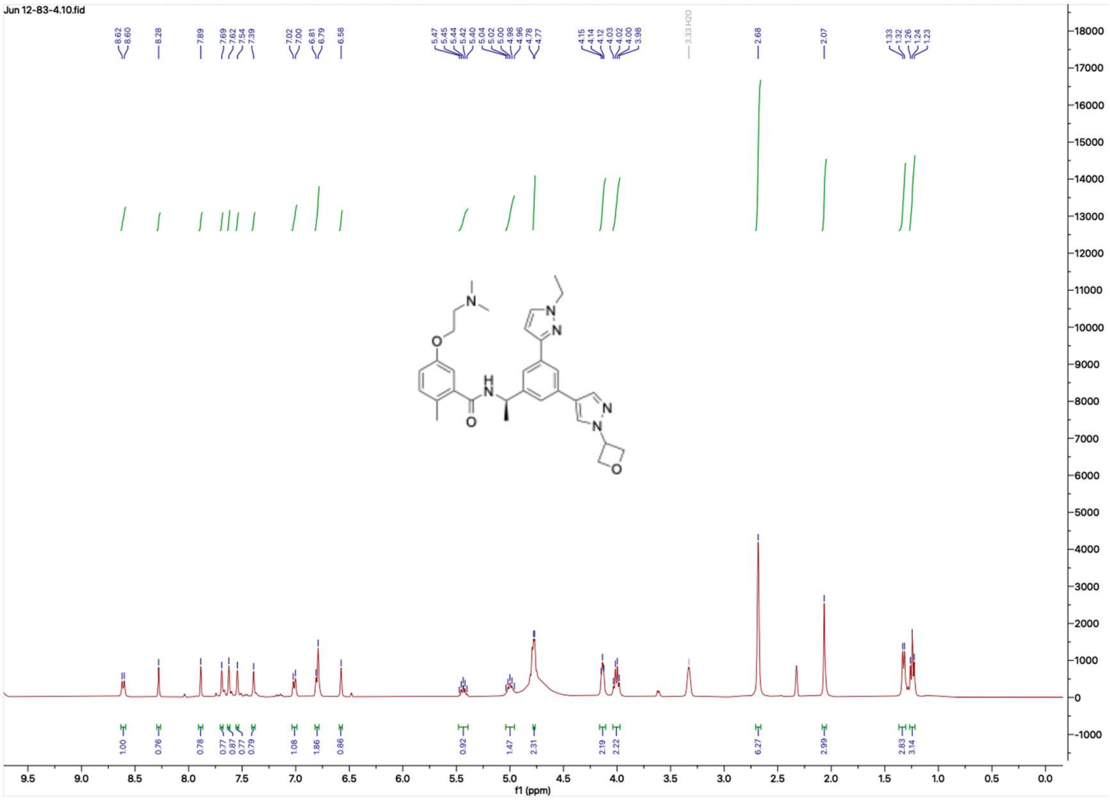


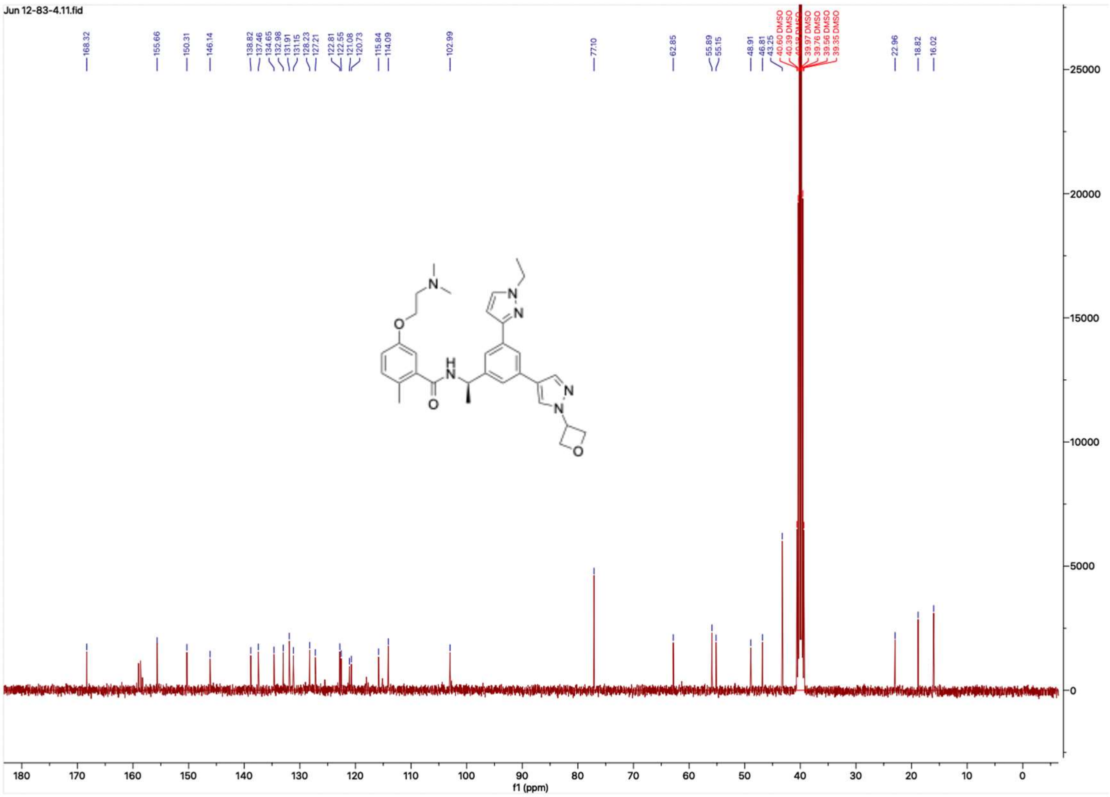

#### HNMR and CNMR spectra of Jun12835

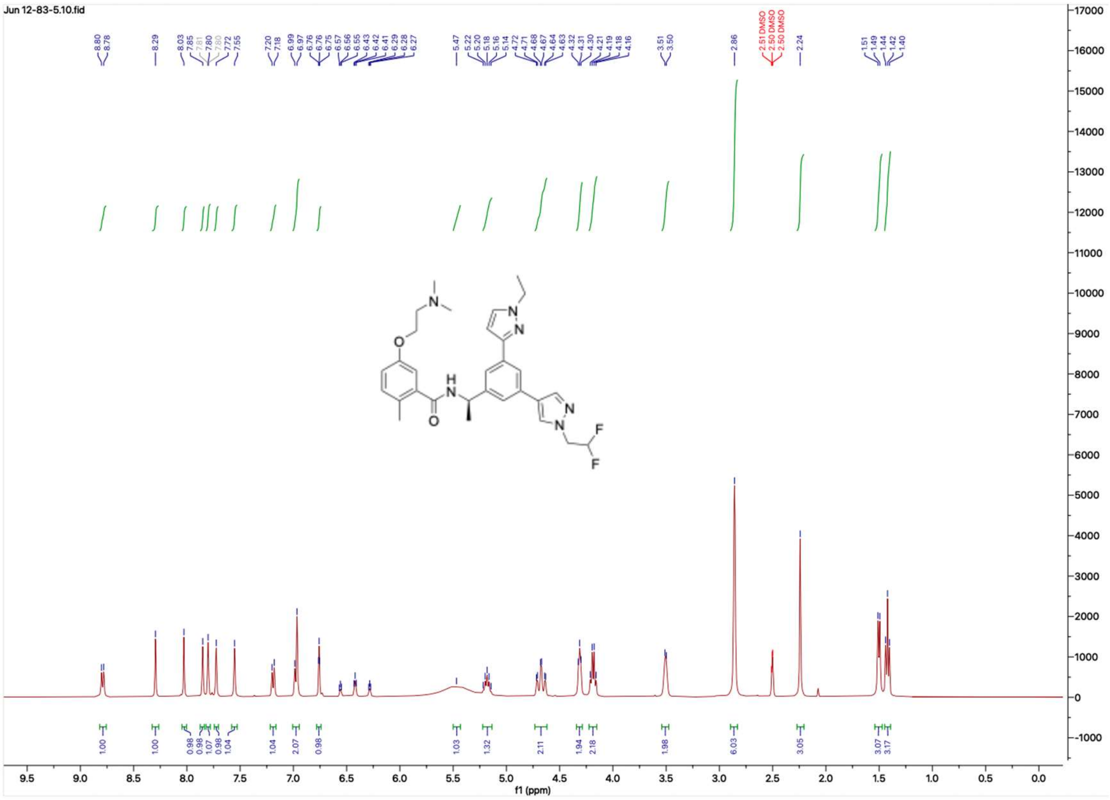


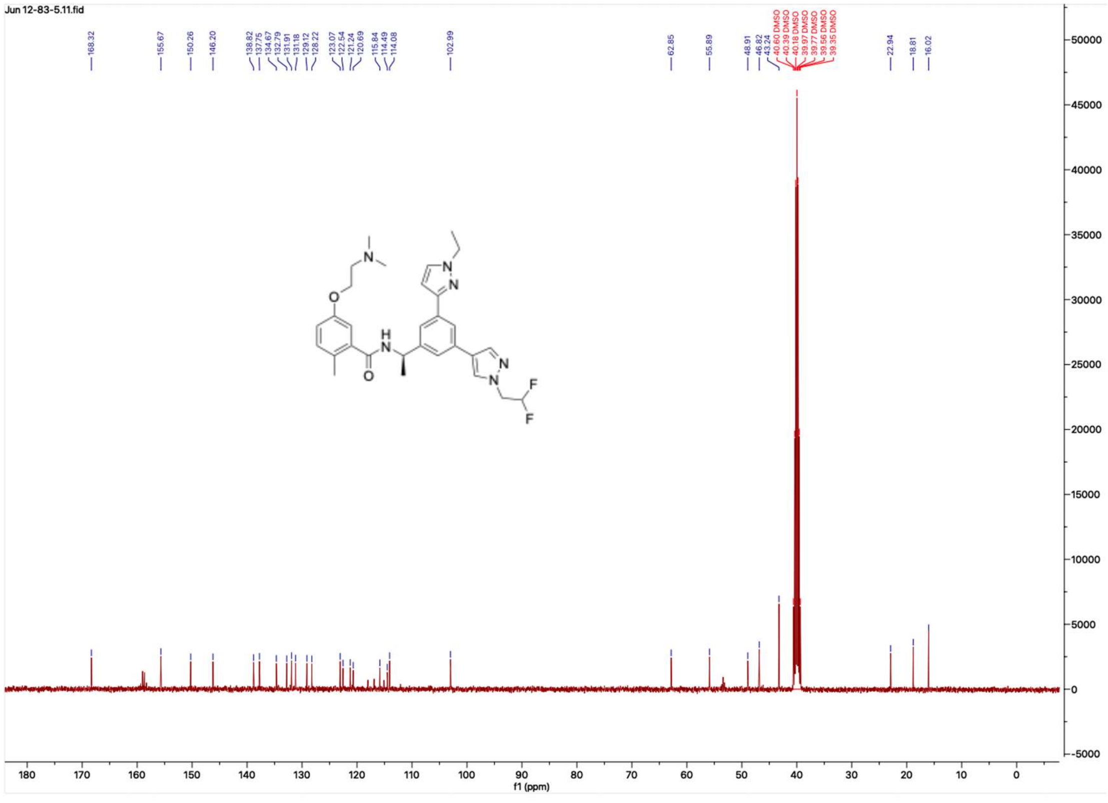

#### HNMR and CNMR spectra of Jun12836

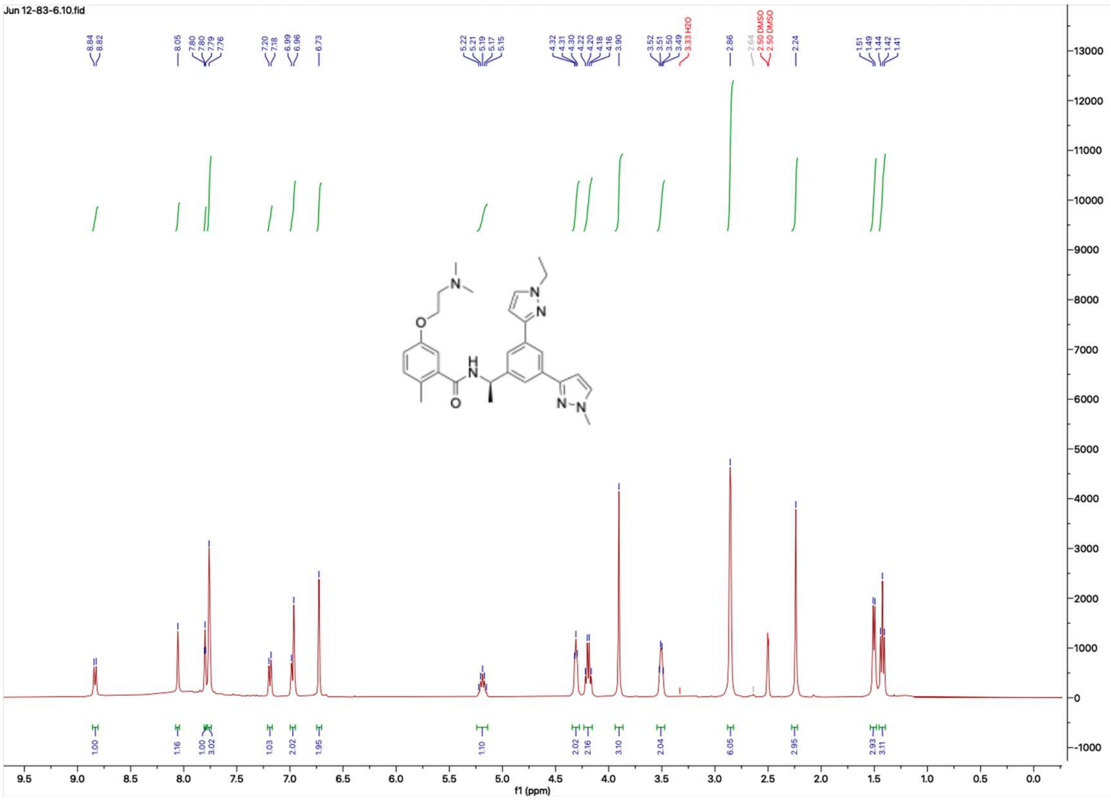


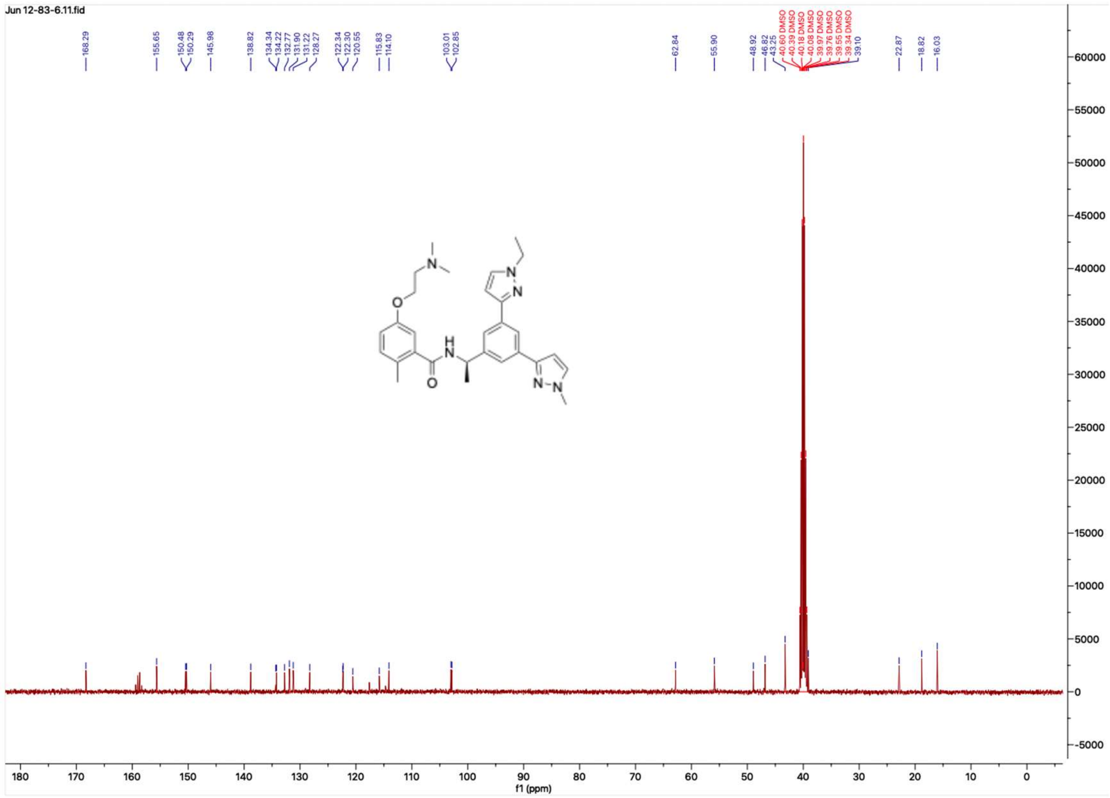

### HNMR and CNMR spectra of Jun12837

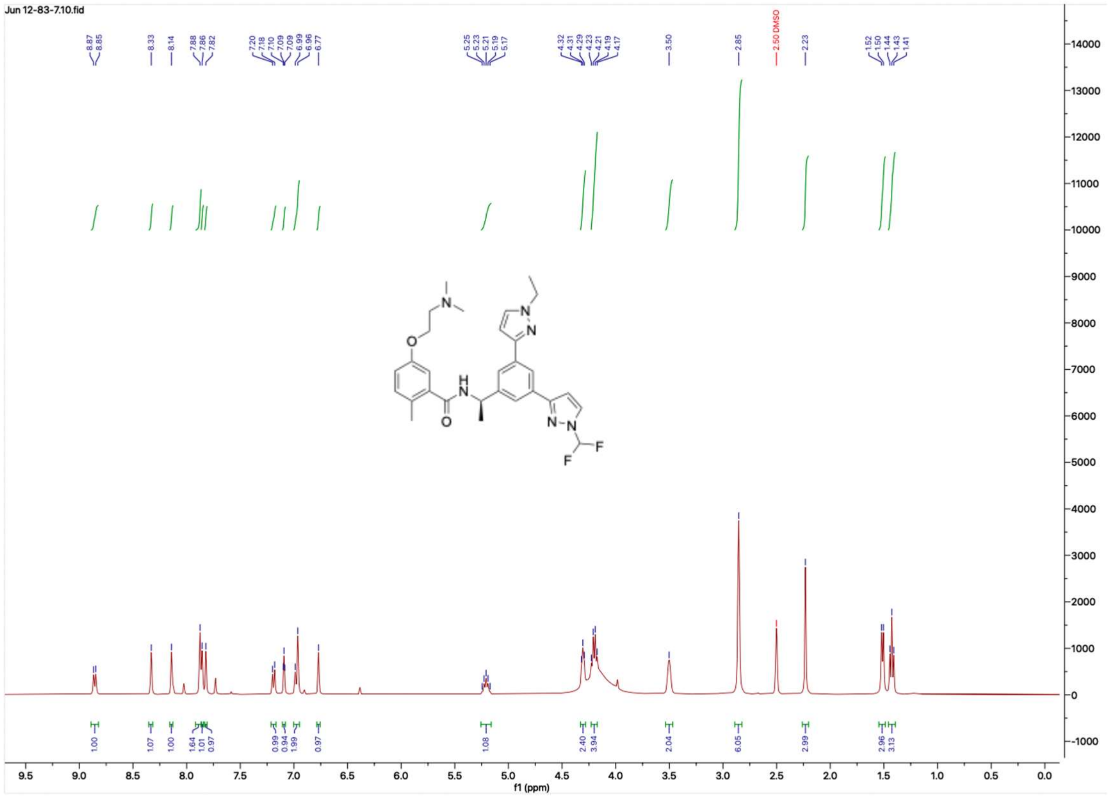


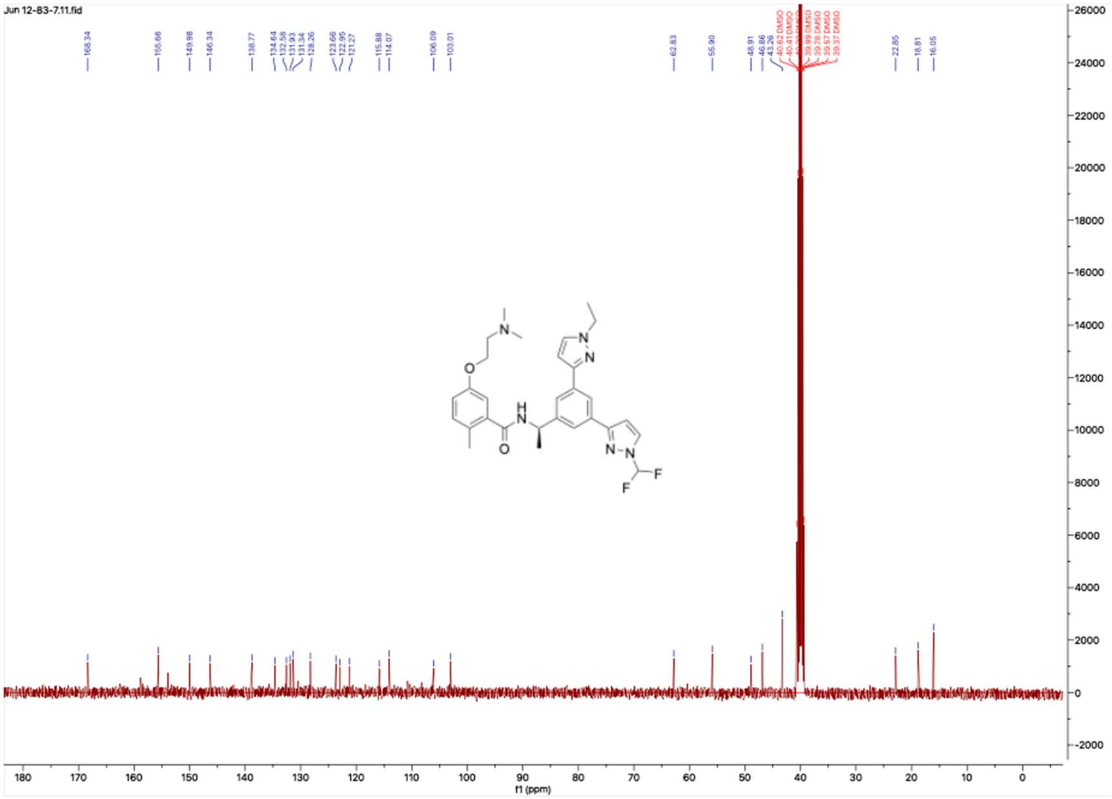

#### HNMR and CNMR spectra of Jun1314

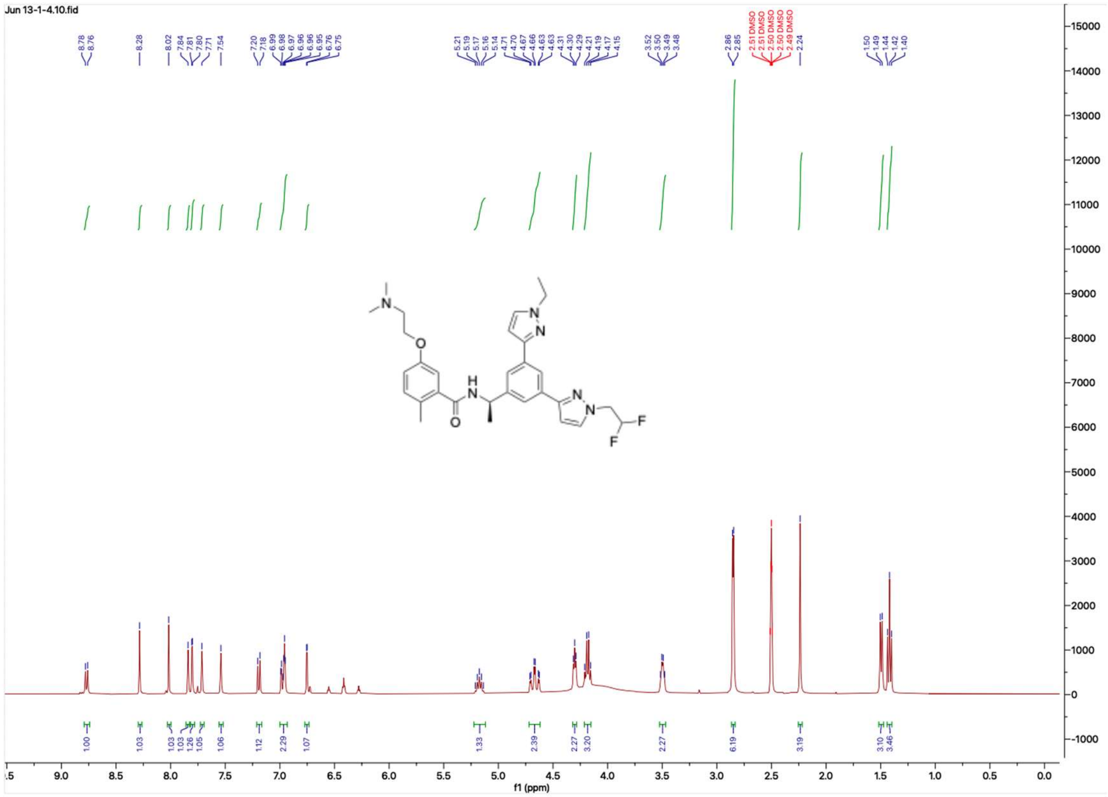


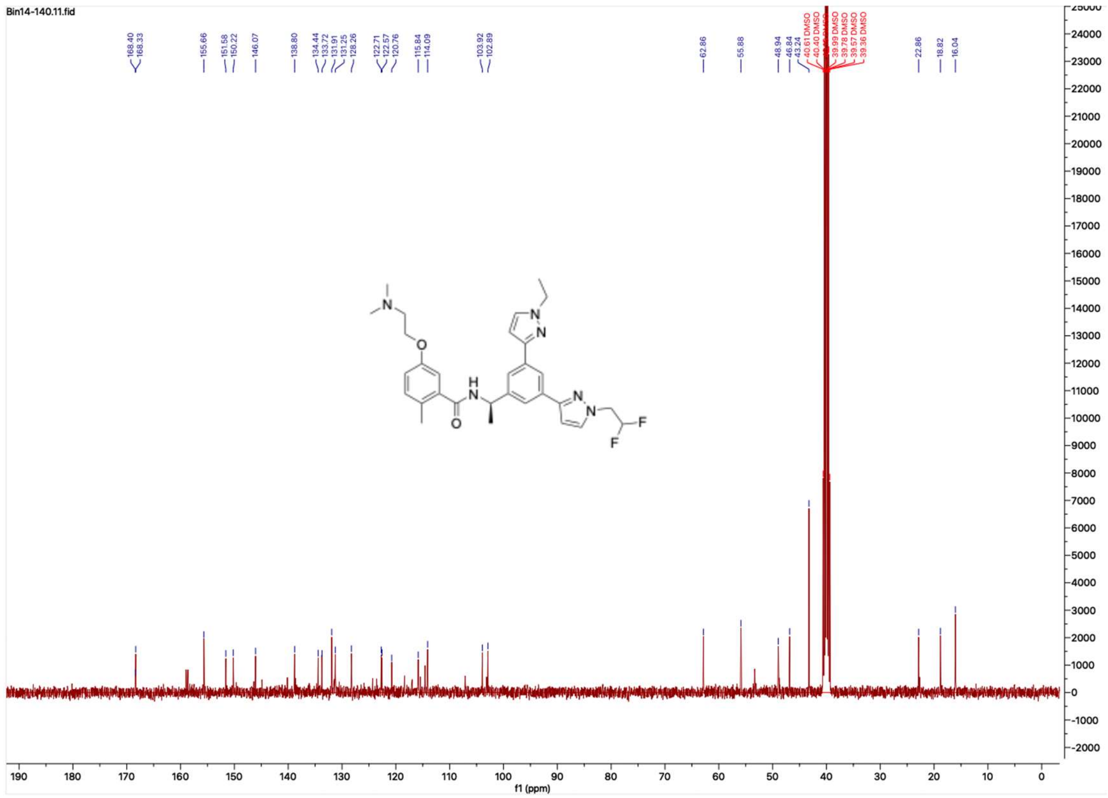

**HNMR and CNMR spectra of Jun12839**

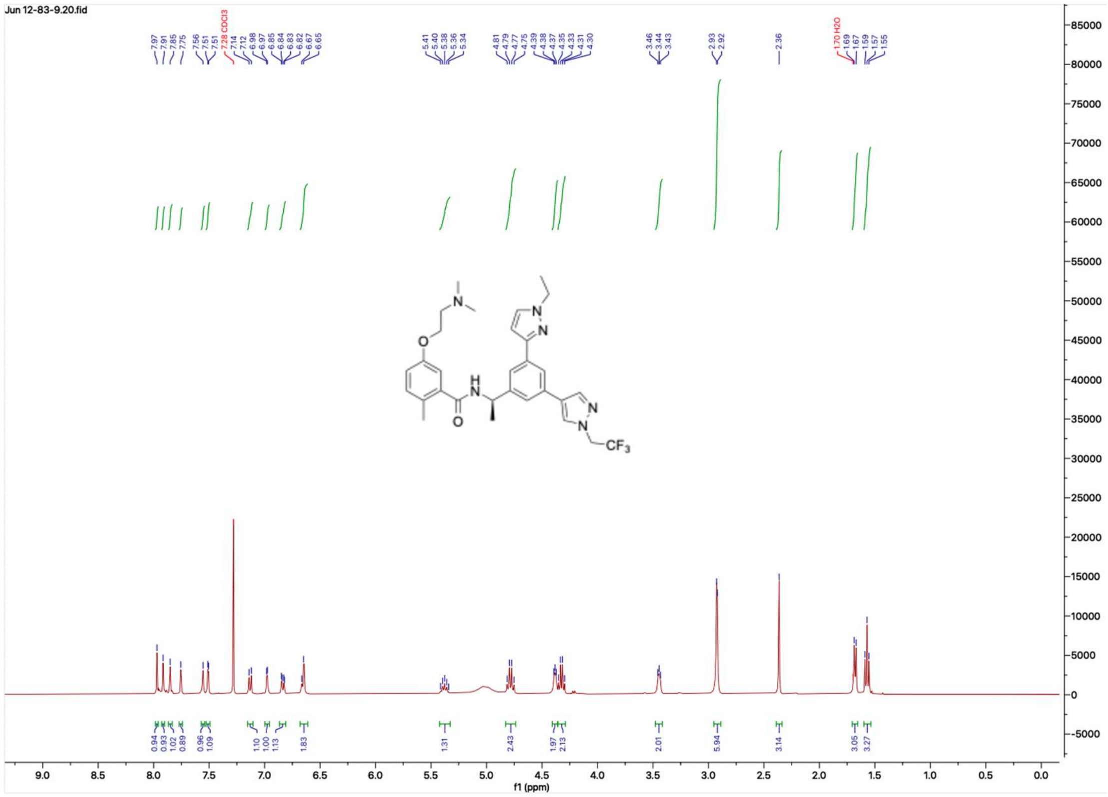

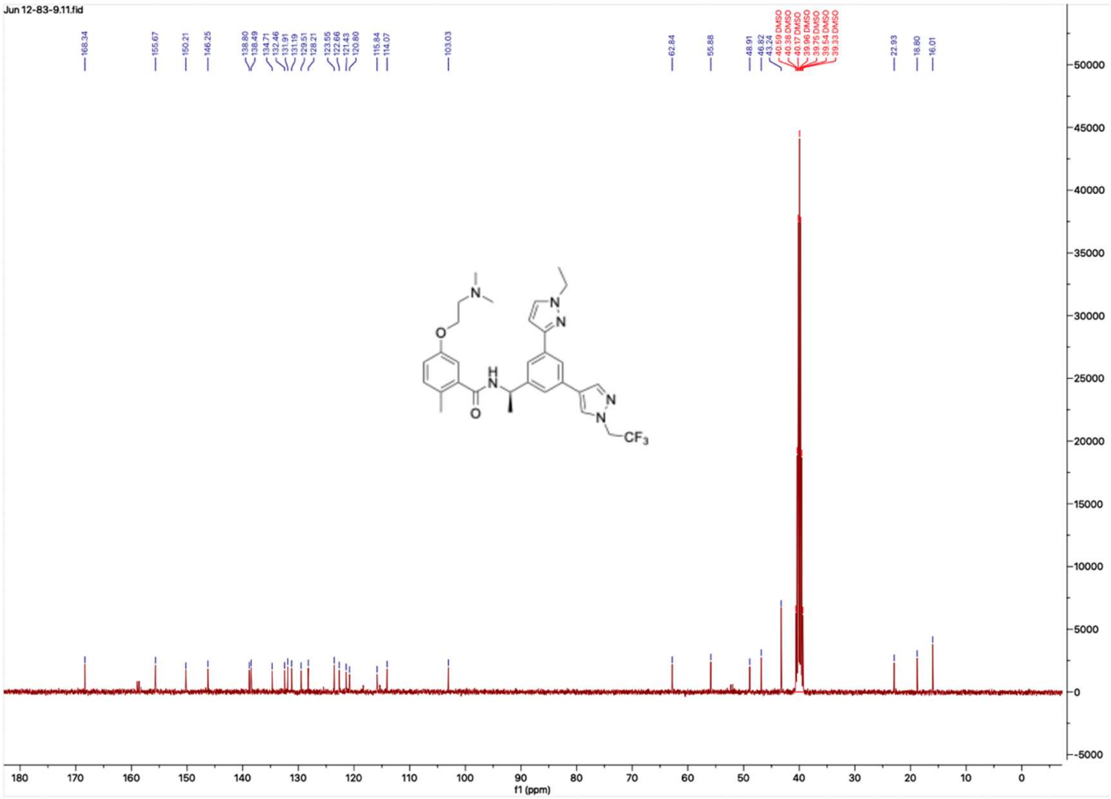

**HNMR and CNMR spectra of Jun12682**

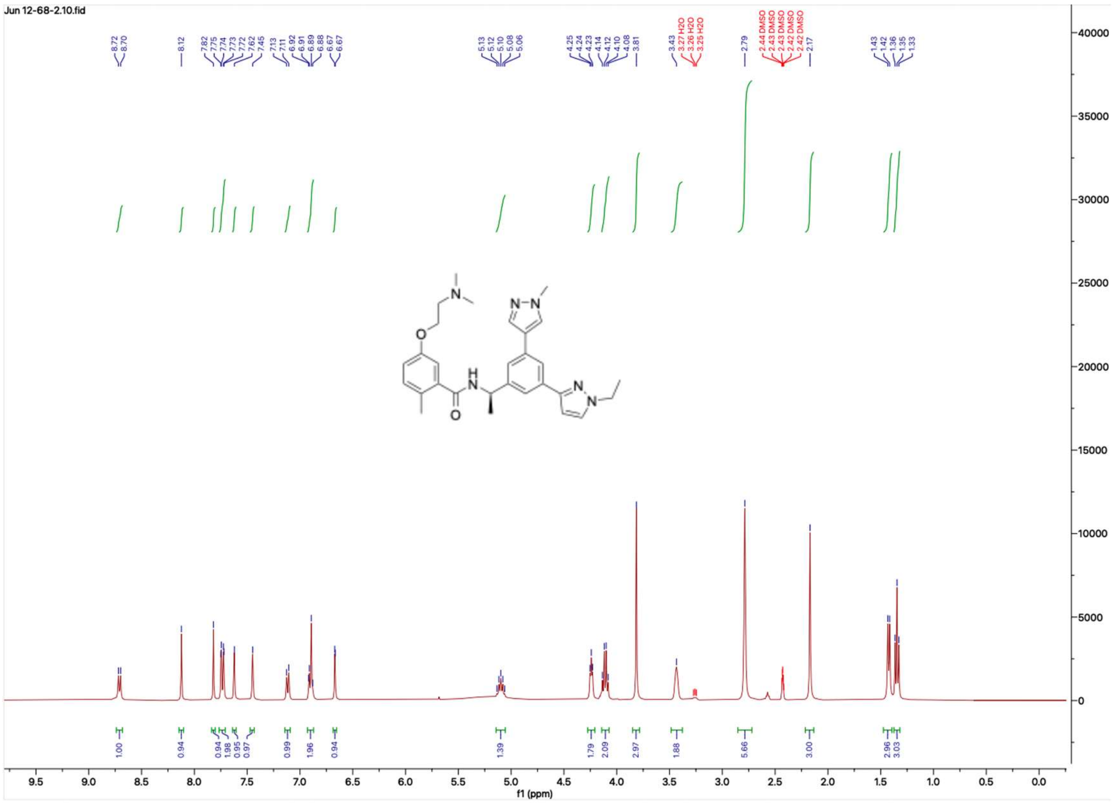

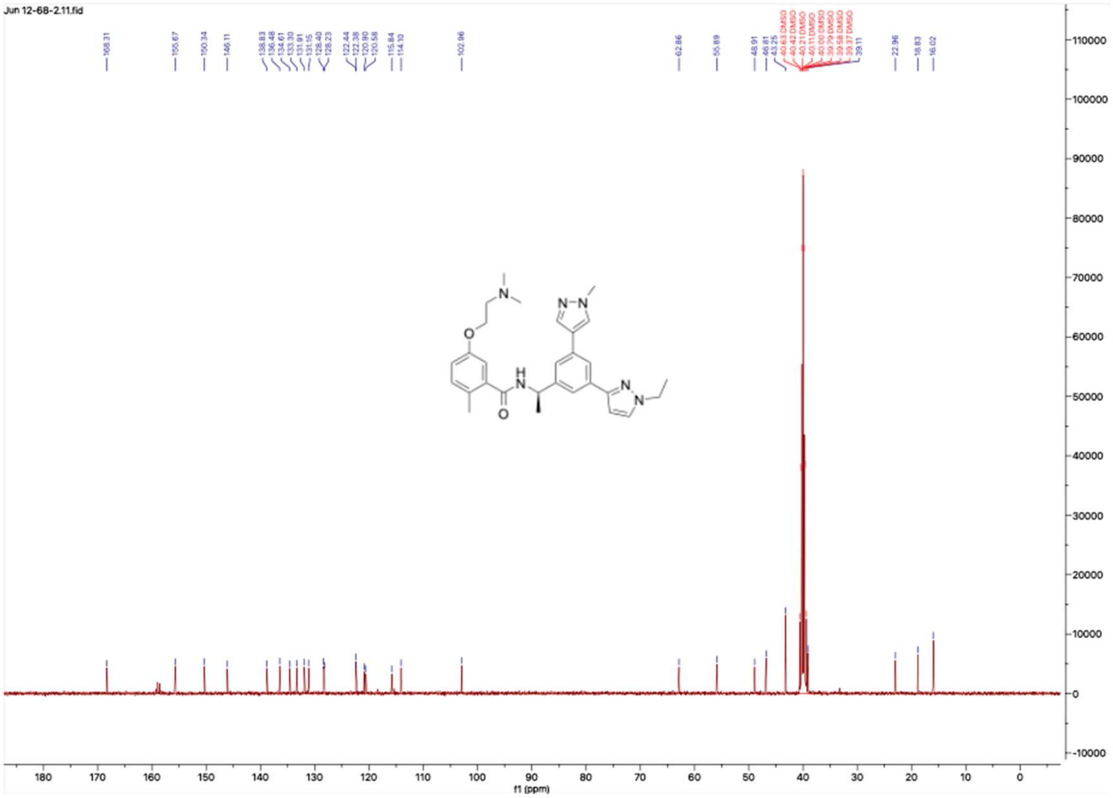

#### HNMR and CNMR spectra of Jun12924

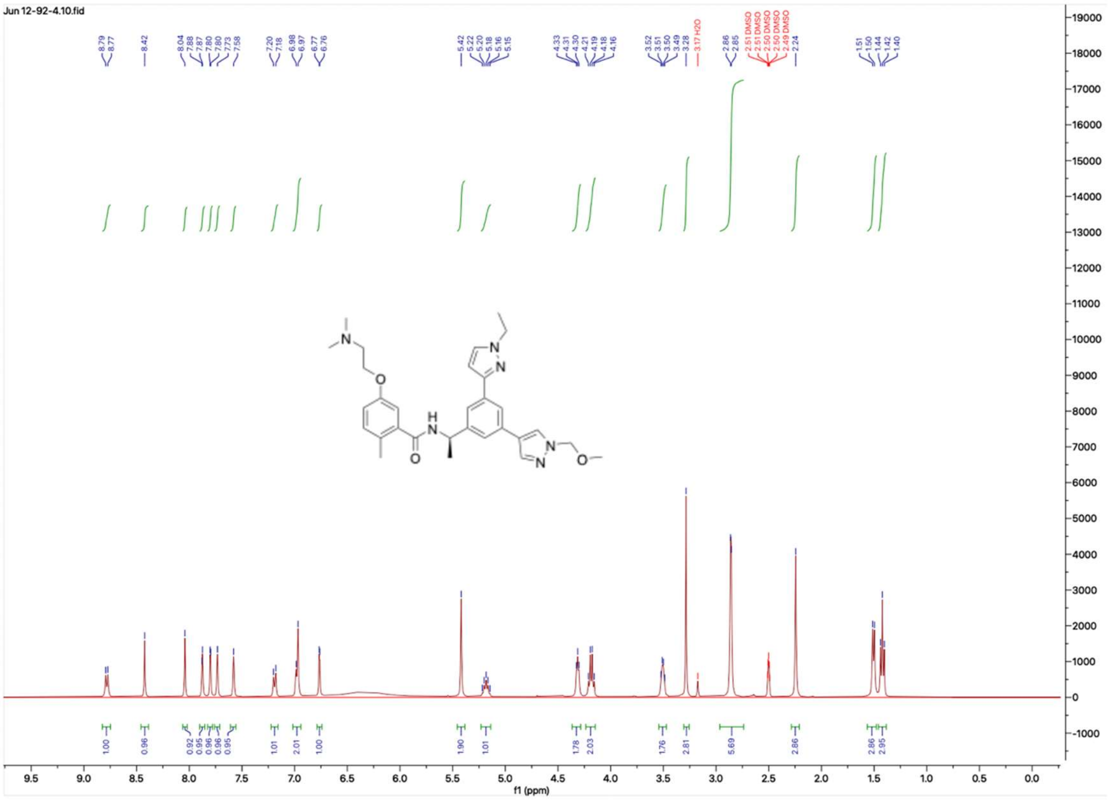


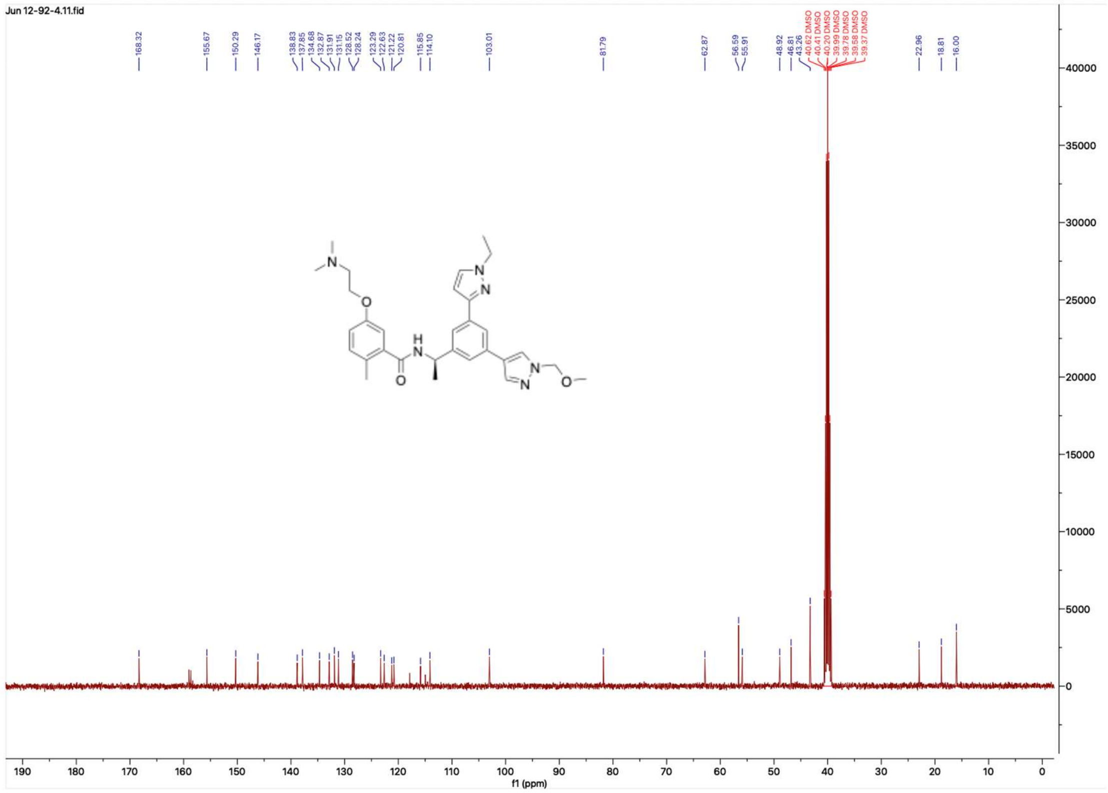

### HNMR and CNMR spectra of Jun12763

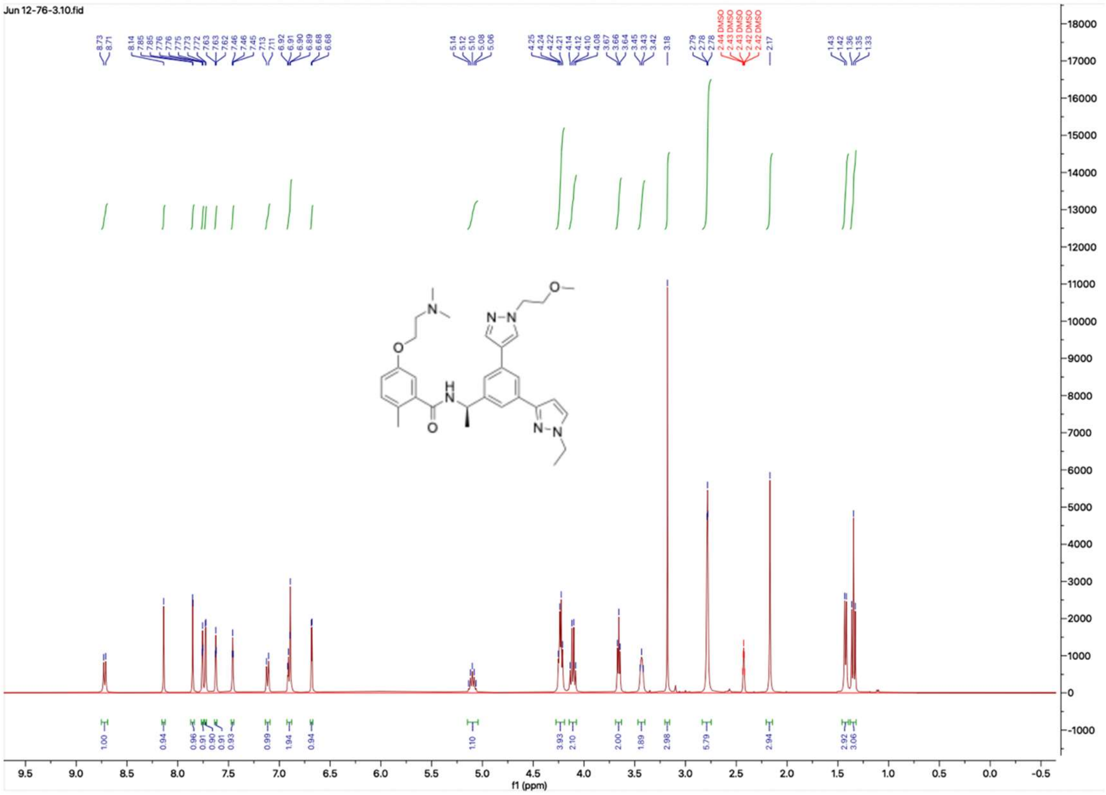


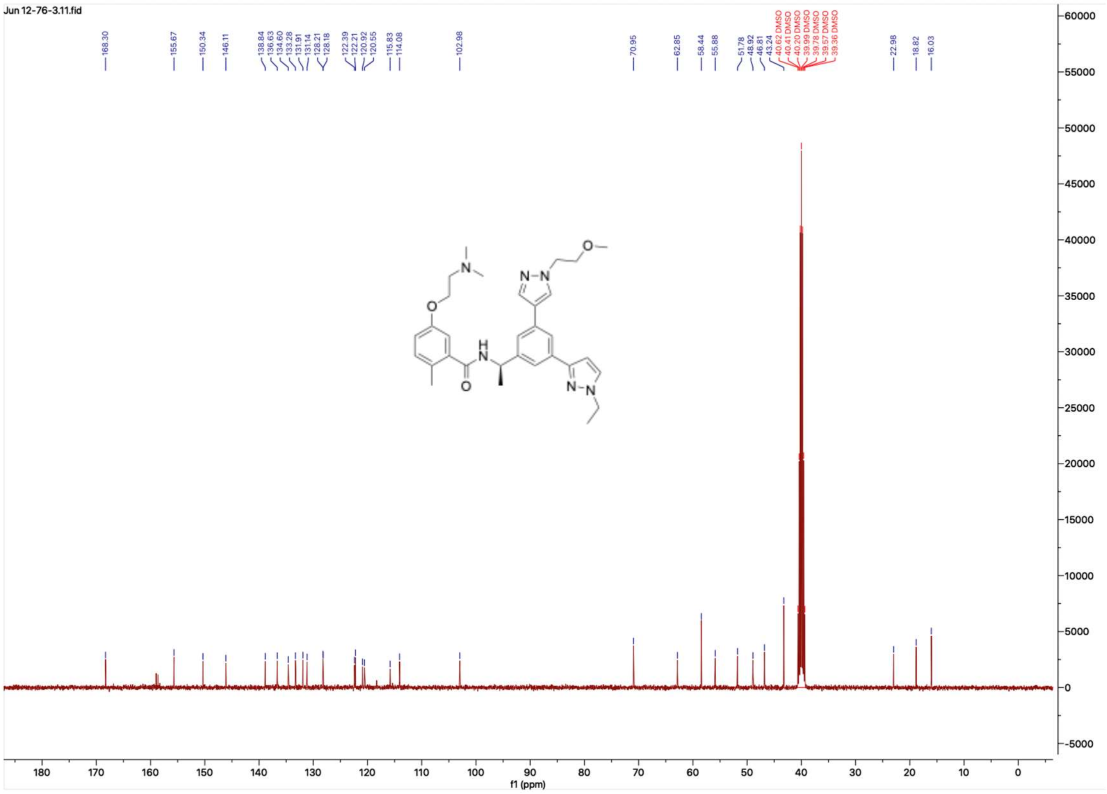
